# Alveolar macrophage chromatin is modified to orchestrate host response to *Mycobacterium bovis* infection

**DOI:** 10.1101/520098

**Authors:** Thomas Jonathan Hall, Douglas Vernimmen, John Andrew Browne, Michael P. Mullen, Stephen Vincent Gordon, David Evan MacHugh, Alan Mark O’Doherty

**Affiliations:** Animal Genomics Laboratory, UCD School of Agriculture and Food Science, College Dublin, Belfield, Dublin, D04 V1W8, Ireland; The Roslin Institute, University of Edinburgh, Easter Bush Campus, Midlothian, EH25 9RG, UK; Bioscience Research Institute, Athlone Institute of Technology, Dublin Road, Athlone, Co. Westmeath, N37 HD68, Ireland; UCD School of Veterinary Medicine, University College Dublin, Belfield, Dublin, D04 W6F6, Ireland; UCD Conway Institute of Biomolecular and Biomedical Research, University College Dublin, Belfield, Dublin, D04 V1W8, Ireland

**Author notes:** **Corresponding authors:** Douglas Vernimmen, David E. MacHugh.

**Keywords:** ChIP-seq, chromatin, macrophage, mycobacteria, *Mycobacterium bovis*, tuberculosis

## Abstract

**Background:** Bovine tuberculosis is caused by infection with *Mycobacterium bovis*, which can also cause disease in a range of other mammals, including humans. Alveolar macrophages are the key immune effector cells that first encounter *M. bovis* and how the macrophage epigenome responds to mycobacterial pathogens is currently not well understood.

**Results:** Here, we have used chromatin immunoprecipitation sequencing (ChIP-seq), RNA-seq and miRNA-seq to examine the effect of *M. bovis* infection on the bovine alveolar macrophage (bAM) epigenome. We show that H3K4me3 is more prevalent, at a genome-wide level, in chromatin from *M. bovis*-infected bAM compared to control non-infected bAM; this was particularly evident at the transcriptional start sites of genes that determine programmed macrophage responses to mycobacterial infection (e.g. M1/M2 macrophage polarisation). This pattern was also supported by the distribution of RNA Polymerase II (PolII) ChIP-seq results, which highlighted significantly increased transcriptional activity at genes demarcated by permissive chromatin. Identification of these genes enabled integration of high-density GWAS data, which revealed genomic regions associated with resilience to infection with *M. bovis* in cattle.

**Conclusions:** Through integration of these data, we show that bAM transcriptional reprogramming occurs through differential distribution of H3K4me3 and PolII at key immune genes. Furthermore, this subset of genes can be used to prioritise genomic variants from a relevant GWAS data set.

## Background

Bovine tuberculosis (bTB) is a chronic infectious disease of livestock, particularly domestic cattle (*Bos taurus* and *Bos indicus*), which causes more than $3 billion in losses to global agriculture annually [1, 2]. The disease can also significantly impact wildlife including, for example, several deer species, American bison (*Bison bison*), African buffalo (*Syncerus caffer*), the brushtail possum (*Trichosurus vulpecula*) and the European badger (*Meles meles*) [3-6]. The etiological agent of bTB is *Mycobacterium bovis*, a pathogen with a genome sequence that is 99.95% identical to *M. tuberculosis*, the primary cause of human tuberculosis (TB) [7]. In certain agroecological milieus *M. bovis* can also cause zoonotic TB with serious implications for human health [8-10].

Previous studies have shown that the pathogenesis of bTB disease in animals is similar to TB disease in humans and many of the features of *M. tuberculosis* infection are also characteristic of *M. bovis* infection in cattle [11-13]. Transmission is via inhalation of contaminated aerosol droplets and the primary site of infection is the lungs where the bacilli are phagocytosed by alveolar macrophages, which normally can contain or destroy intracellular bacilli [14, 15]. Disease-causing mycobacteria, however, can persist and replicate within alveolar macrophages via a bewildering range of evolved mechanisms that subvert and interfere with host immune responses [16-19]. These mechanisms include: recruitment of cell surface receptors on the host macrophage; blocking of macrophage phagosome-lysosome fusion; detoxification of reactive oxygen and nitrogen intermediates (ROI and RNI); harnessing of intracellular nutrient supply and metabolism; inhibition of apoptosis and autophagy; suppression of antigen presentation; modulation of macrophage signalling pathways; cytosolic escape from the phagosome; and induction of necrosis, which leads to immunopathology and shedding of the pathogen from the host [20-25].

Considering the dramatic perturbation of the macrophage by intracellular mycobacteria, we and others have demonstrated that bovine and human alveolar macrophage transcriptomes are extensively reprogrammed in response to infection with *M. bovis* and *M. tuberculosis* [26-31]. These studies have also revealed that differentially expressed gene sets and dysregulated cellular networks and pathways are functionally associated with many of the macrophage processes described above that can control or eliminate intracellular microbes.

For many intracellular pathogens it is now also evident that the infection process involves alteration of epigenetic marks and chromatin remodelling that may profoundly alter host cell gene expression [32-35]. For example, distinct DNA methylation changes are detectable in macrophages infected with the intracellular protozoan *Leishmania donovani*, which causes visceral leishmaniasis [36]. Recent studies using cells with a macrophage phenotype generated from the THP-1 human monocyte cell line have provided evidence that infection with *M. tuberculosis* induces alterations to DNA methylation patterns at specific inflammatory genes [37] and across the genome in a non-canonical fashion [38].

With regards to host cell histones, the intracellular pathogen *Chlamydia trachomatis* secretes NEU, a SET domain-containing effector protein that functions as a histone methyltransferase and induces chromatin modifications favourable to the pathogen [39]. It has also been shown that *Legionella pneumophila*—the causative agent of Legionnaires’ disease—secretes a SET domain-containing histone methyltransferase, RomA, which targets histone H3 to downregulate host genes and promote intracellular replication [40]. In the context of mycobacterial infections, Yaseen et al. have shown that the Rv1988 protein, secreted by virulent mycobacteria, localises to the chromatin upon infection and mediates repression of host cell genes through methylation of histone H3 at a non-canonical arginine residue [41]. In addition, chromatin immunoprecipitation sequencing (ChIP-seq) analysis of H3K4 monomethylation (a marker of poised or active enhancers), showed that regulatory sequence motifs embedded in subtypes of Alu SINE transposable elements are key components of the epigenetic machinery modulating human macrophage gene expression during *M. tuberculosis* infection [42].

In light of the profound macrophage reprogramming induced by mycobacterial infection, and previous work demonstrating a role for host cell chromatin modifications, we have used ChIP-seq and RNA sequencing (RNA-seq) to examine gene expression changes that reflect host-pathogen interaction in bovine alveolar macrophages (bAM) infected with *M. bovis*. The results obtained support an important role for dynamic chromatin remodelling in the macrophage response to mycobacterial infection, particularly with respect to M1/M2 polarisation. Genes identified from ChIP-seq and RNA-seq results were also integrated with GWAS data to prioritise genomic regions and SNPs associated with bTB resilience. Finally, the suitability of bAM for ChIP-seq assays and the results obtained demonstrate that these cells represent an excellent model system for unravelling the epigenetic and transcriptional circuitry perturbed during mycobacterial infection of vertebrate macrophages.

## Results

### *M. bovis* infection induces trimethylation of H3K4 at key immune function related loci in bovine alveolar macrophages

Previous studies have shown that bAM undergo extensive gene expression reprogramming following infection of *M. bovis* [29], with approximately one third of the genome significantly differentially expressed within bovine macrophages 24 hours after infection [28]. Changes of this magnitude are comparable to those observed in previous experiments that have examined the chromatin remodelling that accompanies mycobacterial infection of macrophages, where trimethylation of lysine 4 of Histone H3 (H3K4me3) was shown to correlate with active transcription [42, 43].

We used chromatin immunoprecipitation sequencing (ChIP-seq) to examine histone modification changes that occur after *M. bovis* infection of bAM from sex-and aged-matched Holstein-Friesian cattle. The aim was to determine genome-wide changes in the distribution of H3K4me3 and H3K27me3, and PolII occupancy at the response genes [44]. After data quality control and filtering, ∼760 million paired end reads were aligned to the UMD 3.1 bovine genome build at an average alignment rate of 96.23%. Correlation plots of genome-wide H3K4me3, H3K27me3 and PolII sequencing reads from infected and non-infected bAMs showed high correlation between samples (Pearson’s correlation coefficient: 0.93– 0.97) for all three ChIP-seq targets (supplementary figure 1); indicating that the observed differences in histone modifications between samples are limited and localised to specific regions of the genome.

**Figure 1.**
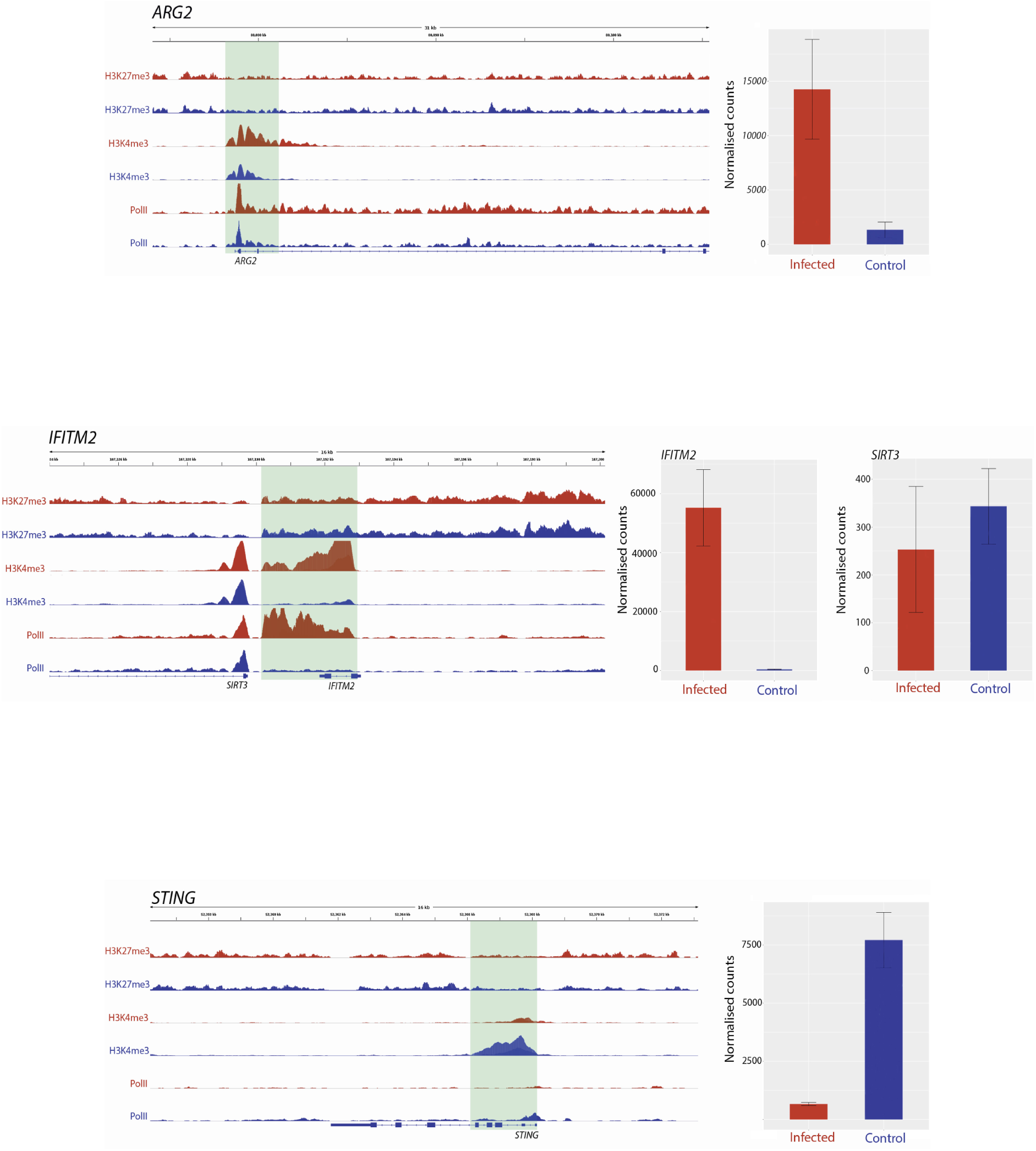
Track visualization of *M. bovis* induced H3K4me3 and PolII occupancy with relative change in expression at three immune response associated genes. Examples of signal tracks illustrating peaks of H3K27me3 (top two tracks), H3K4me3 (middle two tracks) and PolII (bottom two tracks) in infected (red) and non-infected (blue) bovine alveolar macrophages, with the bovine reference genome on the bottom of each panel reading left to right. Accompanying each track image is the expression of the corresponding gene, with normalised counts of infected cells in red and control in blue. The *ARG2* gene exhibited an increase in H3K4me3 at 24 hpi as evidenced by the larger red H3K4me3 and red PolII peaks. The *IFITM2* gene also exhibited larger H3K4me3 and PolII peaks in infected samples; however, in contrast to this, *SIRT3*, which is located ∼20kb upstream from *IFITM2* gene, had no significant change in either peak. *TMEM173* (aka *STING*) exhibits an opposite pattern to most genes identified as having differential H3K4me3, where a larger peak is observed in control samples rather than infected.

Differential peaks between conditions were called, compared and visualised with IGV to determine where differences in H3K4me3, H3K27me3 and PolII occupancy occur between control and infected bAM (Figure 1). ChIP seq peaks are defined as areas of the genome enriched by read counts after alignment to the reference genome.

Peak differences for H3K4me3 occurred at multiple locations across the genome and were estimated by the fold enrichment of a peak normalised against input control DNA that had not undergone antibody enrichment. Differential peaks in each condition were defined by several criteria: 1) the fold enrichment of each peak had to be larger than 10 in at least one condition [45]; 2) the identified peaks had a *P*-value cut off of 0.05; 3) the peaks being compared in each condition were no more than 500 bp up-and downstream of each other; 4) the peaks were classified as different using log-likelihood ratios and affinity scores, with using MACS2 and diffBind, respectively; and 5) visual inspection of the tracks of the peaks confirmed the computationally determined differences in each condition.

Peaks that occurred in a particular sample indicate that H3K4me3 and PolII are highly correlated with condition (Figure 2A); this demonstrates that the differences in H3 modifications are a result of infection rather than genomic differences between animals. Analysis of genome-wide H3K4me3 revealed significant peak differences between control and infected samples at multiple sites in the genome under these criteria, with some of these differences occurring at the transcriptional start site of 233 genes. (Figure 2 A-D and supplementary figure 4). Supplementary figure 1 demonstrates that the differences in H3K4me3 and Pol II peaks are minor, with cells from both conditions sharing most peaks and differing by only 1.8–2.95% in peaks across the genome. Principal component analysis (PCA) of the H3K4me3 mark and PolII data indicated that these H3K4me3 and PolII peak differences are strongly associated with *M. bovis* infection of bAM (supplementary figure 3).

**Figure 2.**
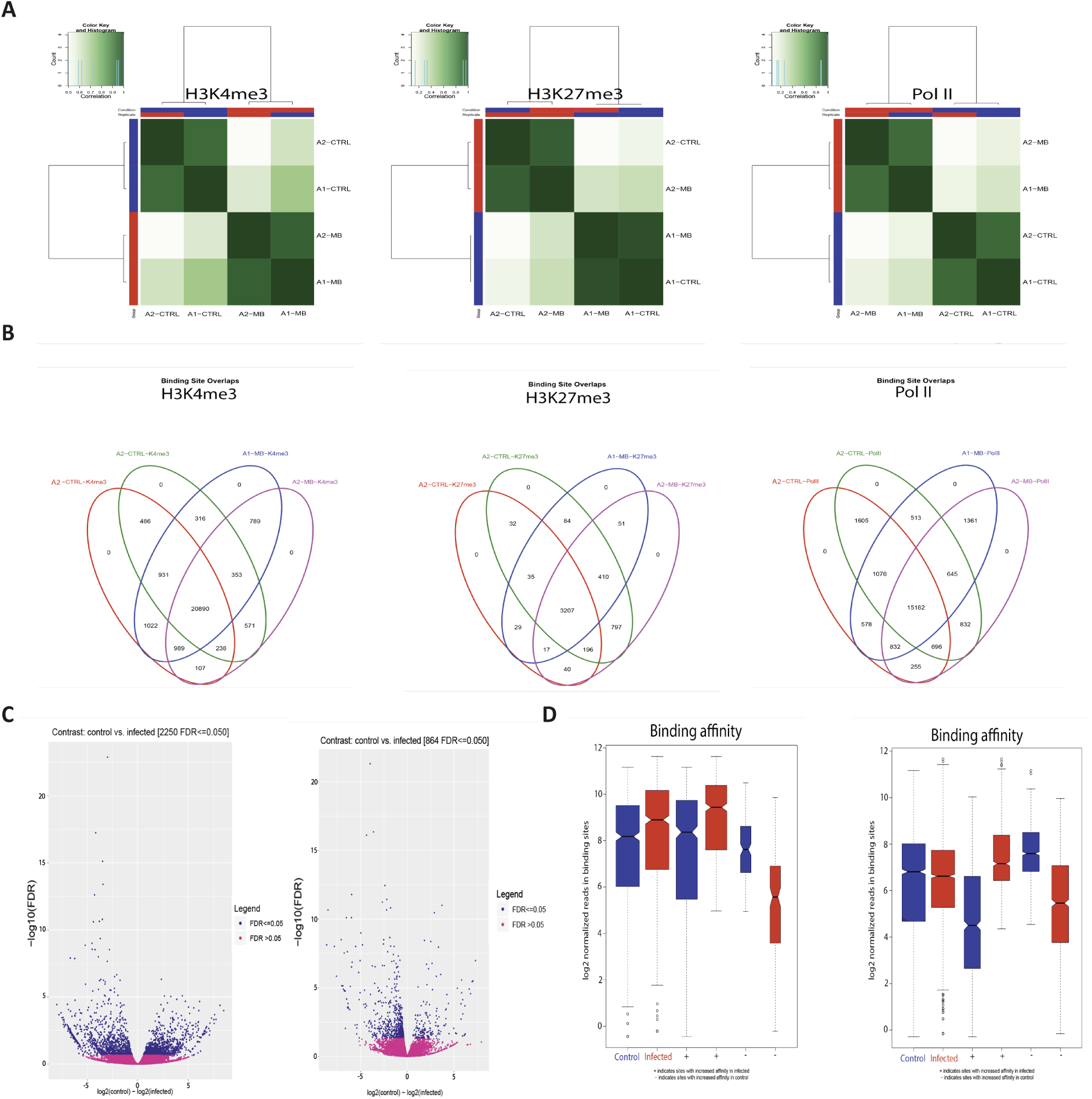
*M.bovis* induced histone modifications occur genome wide at key immune loci. (**A**) Correlation heatmaps of differential peaks for H3K4me3, H3K27me3 and PolII. Every peak location that is not consistent between each animal in each condition (i.e. a peak only occurs in the control group) is compared to determine if these inconsistent peaks are correlated with the animal or the condition. The differential peaks in H3K4me3 and PolII correlate highly with condition, whereas there was no significant global differences in the distribution of H3K27me3. (**B**) Venn diagrams of differential peaks for H3K4me3, H3K27me3 and PolII. Each condition shares the majority of peaks. Where differences occur at TSS of genes, these genes are frequently associated with immune function. (**C**) Volcano plots of differential peaks for H3K4me3 and PolII. The y-axis shows significance as FDR and the x-axis indicates increase in affinity for control (left) and infected (right). Significant sites are denoted in blue. (**E**) Boxplots of differential peaks for H3K4me3 and PolII. Infected bAM are shown in red and control bAM are shown in blue. The left two boxes of each plot show distribution of reads over all differentially bound sites in the infected and control groups. The middle two boxes of each plot show the distribution of reads in differentially bound sites that increase in affinity in the control group. The far right boxes in each plot show the distribution of reads in differentially bound sites that increase in affinity in the infected group.

**Figure 3.**
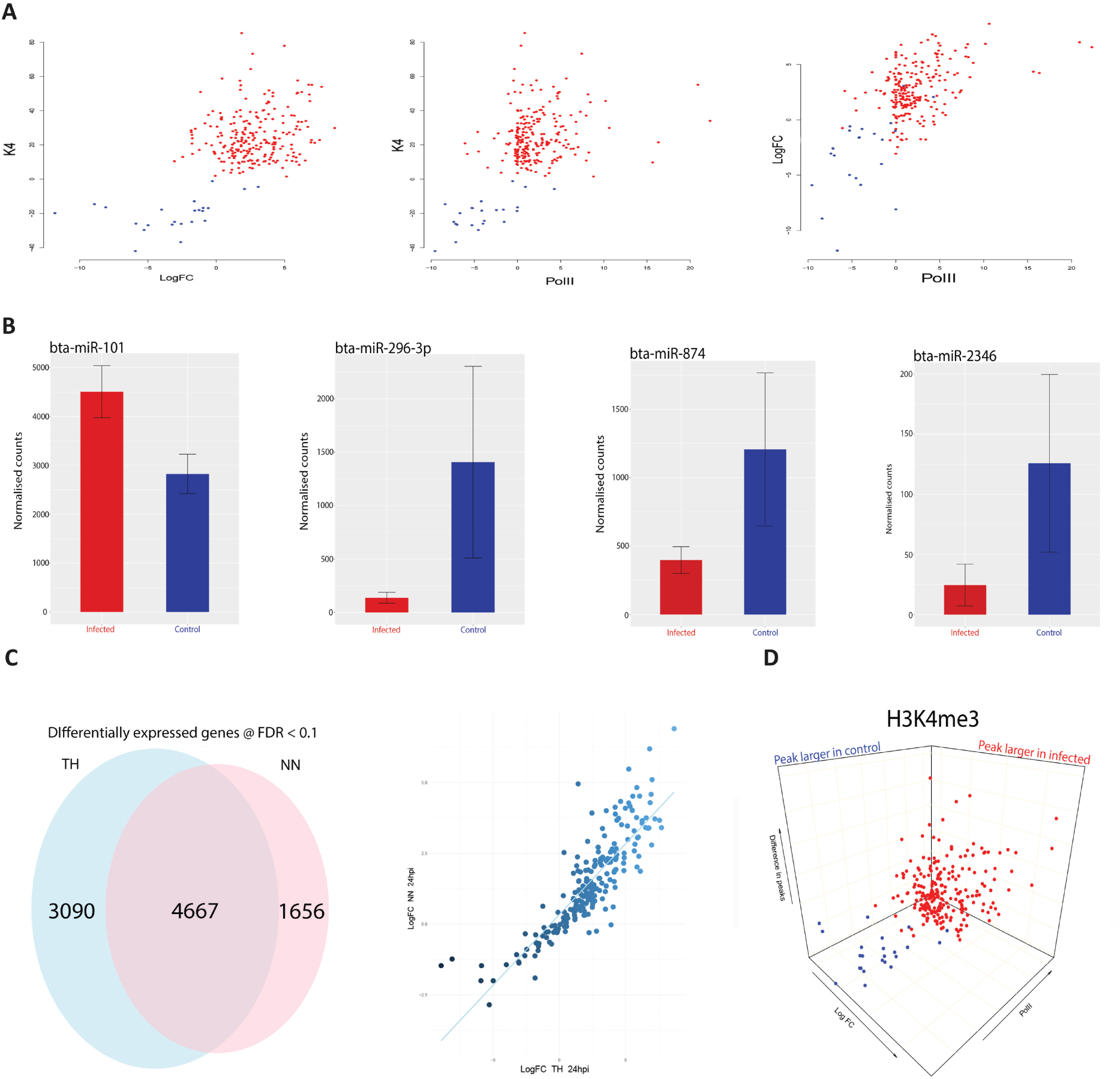
H3K4me3 is accompanied by functional changes in PolII occupancy, gene expression and gene regulation. (**A**) Scatter plots of H3K4me3 against PolII occupancy and gene expression. The first plot is the difference of peaks for H3K4me3 between conditions, ranging from negative to positive values, with negative being a larger peak in the control samples (blue dots) and the positive values being a larger peak in the infected samples (red dots), on the y-axis. The x-axis represents the log2FC for each of the 232 genes, with each gene as a single data point. The second plot also has H3K4me3 on the y-axis but with peak differences in PolII on the x-axis, with negative and positive values corresponding to greater occupancy in the control and infected samples, respectively. The final plot shows log2FC relative to PolII occupancy. (**B**) Plots of normalised miRNA-seq counts. Each plot represents the normalised counts of a miRNA that was detected as exhibiting differential expression. Bta-miR-101 interacts with *ARG2*, bta-miR-296-3p with *TMEM173* (aka *STING*), bta-miR-874 with *BCL2A1* and bta-miR-2346 with *STAT1*. Red bars indicate infected and blue represent control samples. (**C**) Correlation and Venn diagram for both RNA-seq studies. The x-axis of the scatter plot represents the log2FC for each of the 232 genes from this study and the y-axis represents the log2FC for each of the 232 genes from the previous study [29]. The Venn diagram shows the global overlap of differentially expressed genes from both studies with an FDR cut off of <0.1. (**D**) 3-D plots for all three data sets. A combination of all three scatter plots from Figure 2A. Data points are genes. Blue genes are those that exhibited greater H3K4me3 in control bAM, red exhibited greater H3K4me3 in infected bAM.

**Figure 4.**
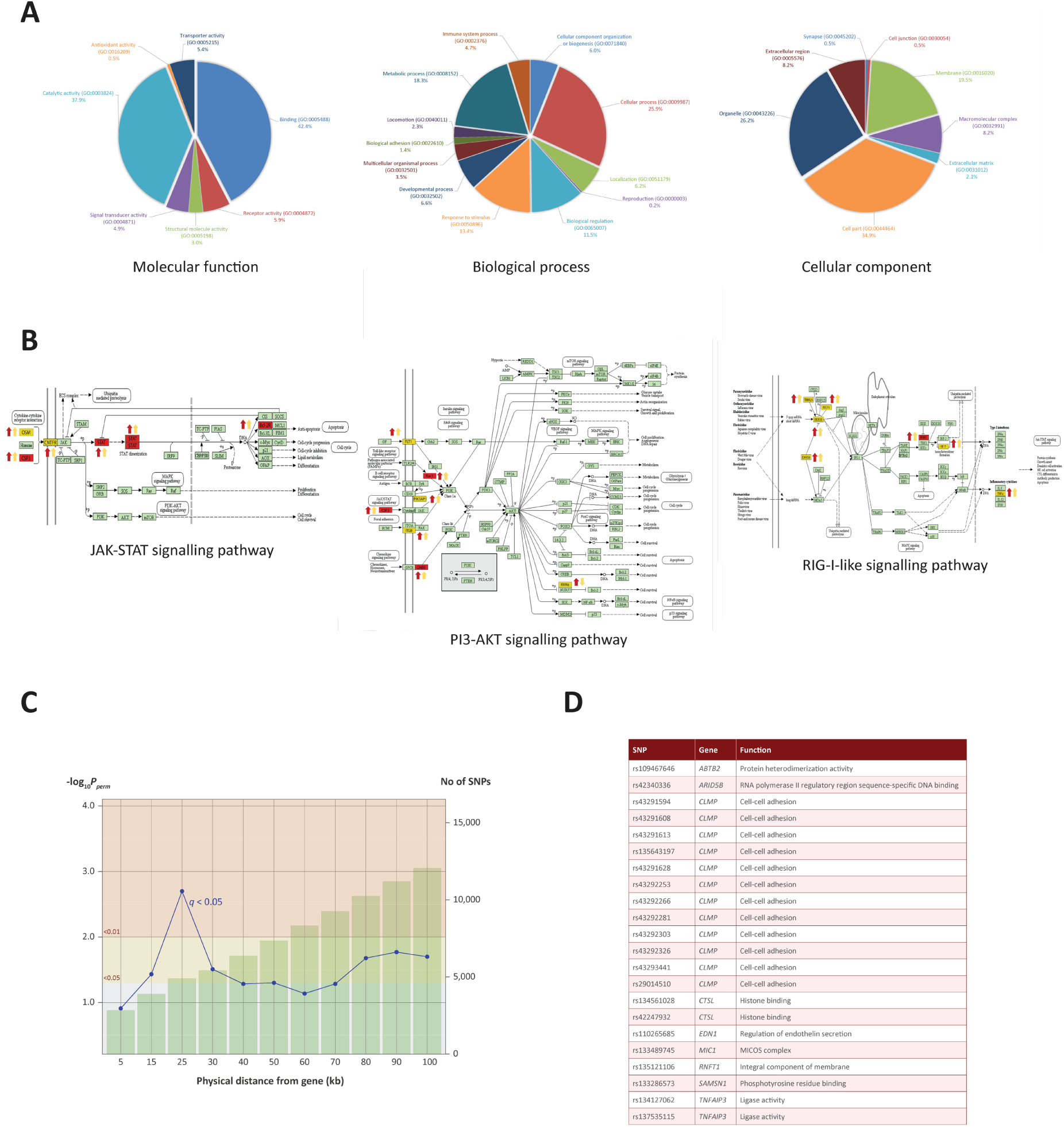
Gene ontology enrichment and pathway analysis. (**A**) Gene ontology pie charts generated through PANTHER pathway analysis. 232 genes cluster by gene ontology under three main categories: *Biological process, Cellular component* and *Molecular function*. (**B**) KEGG pathway images containing genes identified from the ChIP-seq and RNA-seq analysis. Gene symbols coloured in yellow were identified in the ChIP-seq and RNA-seq analysis. Gene symbols coloured in red were also targeted by one or more differentially expressed miRNAs. Up or down red arrows indicate greater H3K4me3 in infected or control, respectively. Up or down yellow arrows indicate log2FC increase or decrease of the associated gene, respectively. (**C**) Line graph showing different genomic ranges from genes that are enriched for significant SNPs from GWAS data for bTB resilience. The bars represent the number of SNPs that occupy each range from each ChIP-seq enriched gene, with more SNPs correlating with a greater distance. The blue plotted line represents the negative log10 probability that the significant SNPs found at each distance at 0.05 FDR *q* value are significant by chance, with SNPs at 25 kb exhibiting the lowest probability. The null SNP *P* value distribution for each data point was generated from 1000 permutations of random SNPs corresponding to the number of SNPs observed in a particular genomic range. (D) Genes enriched for SNPs significantly associated with resilience to *M. bovis* infection. SNP IDs and functional information obtained from the GeneCards^®^ database [107] are also shown.

### Changes in H3K4me3 are accompanied with immune related transcriptional reprogramming

Previous studies have shown that increased H3K4me3 is frequently accompanied by an increase in PolII occupancy and elevated expression of proximal genes [46, 47]. In the present study, we observed that H3K4me3 is accompanied by an increase in PolII occupancy (Figure 1 and supplementary figure 4). For a small number of genes (24 out of 233) where the H3K4me3 peak was larger in the control than the infected samples, PolII occupancy was greater in control bAM for 20 genes (83.3%) and greater in infected bAM for three genes (12.5%). Conversely, where the H3K4me3 peak was larger in the infected bAM, PolII occupancy was greater in the infected samples for 127 genes (60.4%) and greater in the control bAM for 14 genes (6.6%). The remaining 60 genes (25%) did not exhibit H3K4me-associated PolII occupancy in either control or infected samples. Figure 3A illustrates this trend, showing that PolII occupancy normally accompanies H3K4me3.

To establish if H3K4me3 mark patterns were correlated with changes in gene expression, control non-infected bAM and bAM infected *M. bovis* AF2122/97 from four animals 24 hpi (including the two animals used for ChIP-seq) were used to generate eight RNA-seq libraries. After quality control and filtering, ∼250 million reads were mapped to the bovine genome, with 72% total read mapping, overall. RNA-seq analysis revealed 7,757 differentially expressed genes (log2FC > 0: 3,723 genes; log2FC < 0: 4,034 genes; FDR < 0.1). Of the 233 genes identified in the ChIP-seq analysis, 232 (99.6%) were differentially expressed with these criteria (see supplementary file 2). Of the genes that exhibited H3K4me3 peaks that were larger in the infected bAMs, 21 (10%) were downregulated and 189 (90%) were upregulated. Of the genes that exhibited larger H3K4me3 peaks in the control group, 22 (91.6%) were downregulated and 2 (8.4%) were upregulated (Figure 3A). This pattern of directional gene expression correlating with H3K4me3 for the control and infected samples is consistent with the literature [46, 47].

Existing published RNA-seq data generated by our group using *M. bovis*-infected (*n* = 10) and control non-infected bAM (*n* = 10) at 24 hpi [29], was also examined in light of the results from the present study. For the 232 genes identified here, a Pearson correlation coefficient of 0.85 was observed for two data sets (Figure 3D), thus demonstrating that gene expression differences between *M. bovis*-infected and control non-infected bAM are consistent across experiments, even where samples sizes are markedly different.

### Transcriptional reprogramming is coupled with differential microRNA expression

We have previously demonstrated that differential expression of immunoregulatory microRNAs (miRNAs) is evident in bAM infected with *M. bovis* compared to non-infected control bAM [31, 48]. To investigate the expression of miRNA in bAM used for the ChIP-seq analyses, miRNA was extracted and sequenced from the samples used for the RNA-seq analysis. After quality control and filtering, ∼100 million reads were mapped to the bovine genome, with 79% total reads mapping, overall. Twenty-three differentially expressed miRNAs were detected at 24 hpi (log2FC > 0: 13; log2FC < 0: 10; FDR < 0.10). Of the 232 genes identified in the ChIP-seq/RNA-seq analysis, 93 are potential targets for the 23 differentially expressed miRNAs (supplementary data 3). Further examination revealed that multiple immune genes, such as *BCL2A1* (bta-mir-874), *ARG2* (bta-mir-101), *TMEM173* (aka *STING*) (bta-mir-296-3p) and *STAT1* (bta-mir-2346), are potential regulatory targets for these miRNAs (Figure 3B). This observation therefore supports the hypothesis that miRNAs function in parallel with chromatin modifications to modulate gene expression in response to infection by *M. bovis*.

The H3K4me3, PolII, K27me3 and RNA-seq data were subsequently integrated to evaluate the relationship between histone modifications and gene expression changes. Three dimensional plots were generated to visualize the global differences between H3K4me3, PolII and gene expression in infected and non-infected bAM (Figure 3D). These plots show that reduction of H3K4me3 in infected cells is associated with a decrease in gene expression and an absence of PolII occupancy. Genome-wide H3K27me3 was also investigated to determine whether methylation of this residue was altered in response to *M. bovis* infection and if it was related to gene expression. No significant differences for H3K27me3 between control and infected bAM were detected, indicating that repression of gene expression through H3K27me3 does not play a role in the bAM response to *M. bovis* at 24hpi. However, supplementary figure 2 indicates that the presence of a H3K27me3 peak in both control and infected cells at the TSS of a H3k4me3 enriched gene correlated well with a lower or complete lack of Pol II occupancy.

### Pathway analysis reveals H3K4me3 marks are enriched for key immunological genes

To identify biological pathways associated with genes identified through the ChIP-seq analyses, we integrated the ChIP-seq, RNA-seq and miRNA-seq data and created a list of 232 genes that were present in each data set. Pathway analyses were carried out using three pathway tools: Ingenuity Pathway Analysis (IPA), Panther and DAVID [49-51]. IPA revealed an association with *respiratory illness* and the *innate immune response* (supplementary file 2). Panther was used to examine the gene ontology categories of the 232 genes (Figure 4A); this revealed enrichment for metabolic processes, response to stimuli and cellular processes, indicating that increased H3K4me3 in response to *M. bovis* infection occurs at TSS of genes associated with the immune response.

The final part of the pathway analysis was performed using DAVID [51]. DAVID uses a list of background genes and query genes (in this case the 232 common genes across data sets) and identifies enriched groups of genes with shared biological functions. The DAVID analysis demonstrated that the 232 genes are involved in several signalling pathways, including the PI3K/AKT/mTOR, JAK-STAT and RIG-I-like signalling pathways (Figure 4B and the top 10 pathways are detailed in supplementary file 3).

### GWAS integration prioritises bovine SNPs associated with resilience to *M. bovis* infection

Previous work used high-density SNP (597,144 SNPs) data from 841 Holstein-Friesian bulls for a genome-wide association study (GWAS) to detect SNPs associated with susceptibility/resistance to *M. bovis* infection [52]. Using a permutation-based approach to generate null SNP distributions, we leveraged these data to show that genomic regions within 100 kb up-and downstream of each of the 232 genes exhibiting differential H3K4me3 ChIP-seq peaks are significantly enriched for additional SNPs associated with resilience to *M. bovis* infection.

In total, 12,056 SNPs within the GWAS data set were located within 100 kb of the 232 H3K4me3 genes. Of these SNPs, up to 26 were found to be significantly associated with bTB susceptibility, depending on the distance interval of each gene. Interestingly, 22 SNPs found within 25 kb of 11 genes were found to be most significant at *P* and *q* values < 0.05, with declining significance of association as the region extended beyond 25 kb (Figure 4C and supplementary file 3). Significant SNPs were detected in proximity to the following genes: *SAMSN1, CTSL, TNFAIP3, CLMP, ABTB2, RNFT1, MIC1, MIC2, EDN1, ARID5B*, all of which had significant differential enrichment of H3K4me3.

## Discussion

### H3K4me3 mark occurs at key immune genes

Our study has generated new information regarding host-pathogen interaction during the initial stages of *M. bovis* infection. We demonstrate that chromatin is remodelled through differential H3K4me3 and that PolII occupancy is altered at key immune genes in *M. bovis*-infected bAM. This chromatin remodelling correlates with changes in the expression of genes that are pivotal for the innate immune response to mycobacteria [28, 29, 53]. Our work supports the hypothesis that chromatin modifications of the host macrophage genome play an essential role during intracellular infections by mycobacterial pathogens [54, 55].

The top pathways identified were the JAK-STAT signalling pathway, the PI3K/AKT/mTOR signalling pathway and the RIG-I-like receptor signalling pathway. In mammals, the JAK-STAT pathway is the principal signalling pathway that modulates expression of a wide array of cytokines and growth factors, involved in cell proliferation and apoptosis [56]. The JAK-STAT signalling pathway and its regulators are also associated with coordinating an effective host response to mycobacterial infection [57, 58]. Two JAK-STAT associated stimulating factors and a ligand receptor that exhibited increased H3K4me3 marks in infected samples were encoded by the *OSM, CSF3* and *CNTFR* genes, respectively [59, 60]. *OSM* has previously been shown to be upregulated in cells infected with either *M. bovis* or *M. tuberculosis* [29, 61, 62]. Our work shows that this increased expression in response to mycobacteria is facilitated by H3K4me3-mediated chromatin accessibility. The protein encoded by *CSF3* has also been implicated as an immunostimulator in the response to mycobacterial infection due to its role in granulocyte and myeloid haematopoiesis [63]. *CNTFR* encodes a ligand receptor that stimulates the JAK-STAT pathway and shows increased expression in other studies of mycobacterial infection [28, 29]. Following stimulation of *JAK* through ligand receptor binding, *STAT1* expression is increased. STAT1, a signal transducer and transcription activator that mediates cellular responses to interferons (IFNs), cytokines and growth factors, is a pivotal JAK-STAT component and a core component in the response to mycobacterial infection [64]. Here, the TSS of *STAT1* was associated with an increased deposition of H3K4me3. Interestingly, upregulation of *STAT1* was associated with a downregulation of bta-miR-2346, predicted to be a negative regulator of STAT1 (see supplementary file 3). Overall, these results show that major components of the JAK-STAT pathway undergo chromatin remodelling mediated via H3K4me3, thereby facilitating activation and propagation of the JAK-STAT pathway through chromatin accessibility.

Key genes encoding components of the PI3K/AKT/mTOR pathway, such as *IRF7, RAC1* and *PIK3AP1*, were also identified as having increased H3K4me3 in *M. bovis* infected macrophages. PI3K/AKT/mTOR signalling contributes to a variety of processes that are critical in mediating aspects cell growth and survival [65]. Phosphatidylinositol-3 kinases (PI3Ks) and the mammalian target of rapamycin (mTOR) are integral to coordinating innate immune defences [66]. The PI3K/AKT/mTOR pathway is an important regulator of type I interferon production via activation of the interferon-regulatory factor 7, IRF7. RAC1 is a key activator of the PI3K/AKT/mTOR pathway and, in its active state, binds to a range of effector proteins to regulate cellular responses such as secretory processes, phagocytosis of apoptotic cells, and epithelial cell polarization [67]. In addition, *in silico* analysis of our differentially expressed miRNAs predicted that several miRNAs, such as bta-miR-1343-3p, bta-miR-2411-3p and bta-miR-1296, regulate *RAC1. PIK3AP1* expression was also increased, in line with previous mycobacterial infection studies [28, 29]. Hence as observed with the JAK-STAT pathway, H3K4me3 at these key PI3K/AKT/mTOR pathway genes acts to regulate the innate response to mycobacterial infection.

Initiation of the RIG-I-like receptor signalling pathway generally occurs following viral infection through sensing of viral RNAs by cytoplasmic RIG-I-like receptors (RLRs) and activation of transcription factors that drive production of type I IFNs [68]. A number of bacterial species induce type I IFN independently of TLRs, potentially through the RIG-I-like pathway [69, 70]. While type I IFNs have a well characterised role in the inhibition of viral replication, their role during bacterial infection is less well defined [71-73]. In humans and mice, *M. tuberculosis*-induced type I IFN is associated with TB disease progression and impairment of host resistance [74-76]. In our study, genes encoding multiple components of the RIG-I-like receptor signalling pathway, such as *TRIM25, ISG15, IRF7* and *IKBKE*, were enriched for K4me3 and PolII occupancy in *M. bovis-*infected bAM. These results demonstrate that the RIG-I-like pathway activation in *M. bovis*-infected bAM is driven, to a large extent, by reconfiguration of the host chromatin.

H3K4me3 enriched loci are also flanked by genomic polymorphisms associated with resilience to *M. bovis* infection. Integration of our data with GWAS data from 841 bulls that have robust phenotypes for bTB susceptibility/resistance revealed 22 statistically significant SNPs within 25 kb of 11 H3K4me3 enriched genes. Most of these genes are involved in host immunity, with *CTSL, TNFAIP3*, and *RNFT1* directly implicated in the human response to *M. tuberculosis* infection [77-79]. The reprioritisation of genomic regions and array-based SNPs using integrative genomics approaches will be relevant for genomic prediction and genome-enabled breeding and may facilitate fine mapping efforts and the identification of targets for genome editing of cattle resilient to bTB.

### H3K4me3 deposition at host macrophage genes and immunological evasion by *M. bovis*

The present study has revealed elevated H3K4me3 deposition and PolII occupancy at key immune genes that are involved in the innate response to mycobacterial infection. In addition, we also identified several immune genes that had differential H3K4me3 and expression, where the expression change may be detrimental to the ability the host macrophage to clear infection. An example of this is *ARG2*, which exhibited increased H3K4me3 deposition, PolII occupancy and expression (Log2FC = 3.415, *P*adj = 7.52 ×10^-16^) in infected cells. However, it is also interesting to note that the integrated expression output of *ARG2* may also be determined by the bta-miR-101 miRNA, a potential silencer of *ARG2* expression, which was observed to be upregulated in infected cells. Elevated levels of arginase 2, the protein product of the *ARG2* gene, have previously been shown to shift macrophages to an M2 phenotype [80, 81], which is anti-inflammatory and exhibits decreased responsiveness to IFN-gamma and decreased bactericidal activity [82]. Hence, it may be hypothesised that *M. bovis* infection triggers H3K4me3 deposition at the TSS of *ARG2* to drive an M2 phenotype and generate a more favourable niche for the establishment of infection. Similarly to *ARG2*, increased expression of *BCL2A1* in *M. bovis*-infected bAM may also facilitate development of a replicative milieu for intracellular mycobacteria. Increased expression of *BCL2A1* is associated with decreased macrophage apoptosis [83], which would otherwise restrict replication of intracellular pathogens.

In comparison to control non-infected bAM, the *TMEM173* (aka *STING*) gene exhibited substantially decreased expression in *M. bovis*-infected bAM (Log2FC = −3.225, *P*_adj_ = 8.64 ×10^-11^). *TMEM173* encodes transmembrane protein 173, which drives IFN production and as such is a major regulator of the innate immune response to viral and bacterial infections, including *M. bovis* and *M. tuberculosis* [28, 84, 85]. Downregulation of *TMEM173* indicates that *M. bovis* can actively reduce or block methylation of H3K4 at this gene in infected macrophages, thereby enhancing the pathogen’s intracellular survival. In this regard, we have recently shown that infection of bAM with *M. tuberculosis*, which is attenuated in cattle, causes increased *TMEM173* expression compared to infection with *M. bovis* [28].

The molecular mechanisms that pathogens employ to manipulate the host genome to subvert or evade the immune response are yet to be fully elucidated. Hijacking the host’s own mechanisms for chromatin modulation is one potential explanation that has garnered attention in recent years [33, 35]. These modulations of the host chromatin in bAMs may be mediated through *M. bovis*-derived signals transmitted through bacterial metabolites, RNA-signalling or secreted peptides [41, 86-88].

## Conclusions

Elucidation of the mechanisms used by pathogens to establish infection, and ultimately cause disease, requires an intimate knowledge of host-pathogen interactions. Using transcriptomics and epigenomics, we have identified altered expression of major host immune genes following infection of primary bovine macrophages with *M. bovis.* We have shown that reprogramming of the alveolar macrophage transcriptome occurs mainly through increased deposition of H3K4me3 at key immune function genes, with additional gene expression modulation via miRNA differential expression. This modulation of gene expression drives a shift of the macrophage phenotype towards the more replicate-permissive M2 macrophage phenotype. We have also identified that alveolar macrophages infected with *M. bovis* exhibit differentially expressed genes (in regions with modified chromatin) that are enriched for significant SNPs from GWAS data for bTB resilience; this shows the power of integrative approaches to identify key loci to be targeted for selective breeding programmes. Our data confirm the emerging concept that pathogens can hijack host chromatin, through manipulation of H3K4me3, to subvert host immunity and to establish infection.

## Methods

### Preparation and infection of bAMs

Alveolar macrophages and *M. bovis* 2122 were prepared as described previously [89] with minor adjustments, 2 × 10^6^ macrophages were seeded in 60 mm tissue culture plates and challenged with *M. bovis* at an MOI of 10:1 (2 × 10^7^ bacteria per plate) for 24 h; parallel non-infected controls were prepared simultaneously.

### Preparation of nucleic acids for sequencing

Sheared fixed chromatin was prepared exactly as described in the truChIP(tm) Chromatin Shearing Kit (Covaris) using 2 × 10^6^ macrophage cells per AFA tube. Briefly, cells were washed in cold PBS and 2.0 ml of fixing Buffer A was added to each plate, to which 200 µl of freshly prepared 11.1% formaldehyde solution was added. After 10 min on a gentle rocker the crosslinking was halted by the addition of 120 µl of quenching solution E, cells were washed with cold PBS, released from the plate using a cell scraper and re-suspended in 300 µl Lysis Buffer B for 10 min with gentle agitation at 4^°^C to release the nuclei. The nuclei were pelleted and washed once in Wash Buffer C and three times in Shearing Buffer D3 (X3) prior to been resuspended in a final volume of 130 µl of Shearing Buffer D3. The nuclei were transferred to a micro AFA tube and sonicated for 8 min each using the Covaris E220e as per the manufacturer’s instructions. Chromatin immunoprecipitation of sonicated DNA samples was carried out using the Chromatin Immunoprecipitation (ChIP) Assay Kit (Merck KGaD) and anti-H3K4me3 (05-745R) (Merck KGaD), Pol II (H-224) (Santa Cruz Biotechnology, Inc.) or anti-H3K27me3 (07-449) (Merck KGaD) as previously described [90]. RNA was extracted from infected (*n* = 4) and control (*n* = 4) bAM samples using the RNeasy Plus Mini Kit (Qiagen) as previously described [91]. All 8 samples exhibited excellent RNA quality metrics (RIN >9).

### Sequencing

Illumina TruSeq Stranded mRNA and TruSeq Small RNA kits were used for mRNA-seq and small RNA-seq library preparations and the NEB Next Ultra ChIPseq Library Prep kit (New England Biolabs) was used for ChIP-seq library preparations. Pooled libraries were sequenced by Edinburgh Genomics (http://genomics.ed.ac.uk) as follows: paired-end reads (2 × 75 bp) were obtained for mRNA and ChIP DNA libraries using the HiSeq 4000 sequencing platform and single-end read (50 bp) were obtained for small RNA libraries using the HiSeq 2500 high output version 4 platform.

### ChIP-seq bioinformatics analysis

Computational analyses for all bioinformatic processes were performed on a 72-CPU compute server with Linux Ubuntu (version 16.04.4 LTS). An average of 54 M paired end 75bp reads were obtained for each histone mark. At each step of data processing, read quality was assessed via FastQC (version 0.11.5) [92]. Any samples that indicated adapter contamination were trimmed via Cutadapt (version 1.15) [93]. Raw read correlation plots were generated via EaSeq (version 1.05) [94]. The raw reads were aligned to UMD 3.1.1 bovine genome assembly using Bowtie2 (version 2.3.0) [95]. The mean alignment rate for the histone marks was 96.23%. The resulting SAM files were converted and indexed into BAM files via Samtools (version 1.3.1) [96]. After alignment, samples were combined and sorted into 14 files, based on the animal (A1 or A2), the histone mark (K4/K27/PolII) and treatment (control or infected) i.e. A1-CTRL-K4. Peaks were called by using alignment files to determine where the reads have aligned to specific regions of the genome, and then comparing that alignment to the input samples as a normalization step.

The peak calling was carried out via MACS (version 2.1.1.20160309) [97]. The K4me3 mark was called in sharp peak mode and K27me3 and Pol II were called in broad peak mode, as per the user guide. Peak tracks were generated in macs and visualized with the Integrative Genome Viewer (version 2.3) [98]. Union peaks were generated by combining and merging overlapping peaks in all samples for each histone mark. Differential peak calling was called via macs using the bdgdiff function. Peaks images were generated by visually assessing all three marks in tandem across the entire bovine genome with IGV. The significance of peaks was determined by sorting peaks for each mark in each treatment by *P* value and then fold enrichment with a cut-off of 2 and a *P* value threshold of 0.05 [99]. Peaks from each animal in each condition for each mark were cross referenced with the IGV images and differential peak caller to determine a difference in fold enrichment for each observed peak difference between conditions. This required comparing peak start and end sites, chromosomes, *P* and *q* values for each summit, summit locations and normalised fold enrichment of a peak against the input sample (see supplementary file 1 for peak sets). Any peaks that exhibited a difference of 4 or greater fold enrichment, a *P* value of less than 0.05, an FDR (*q* value) less than 0.05 and that were also identified by the differential peak caller were selected for further analysis (see supplementary file 1 for peaks at TSS that met some but not all of the above criteria). Peaks that were then classified to be different between conditions in all three data sets were examined to determine their proximity to TSS. Differential peaks were also called using the R package DiffBind (version 2.80) [100]. DiffBind includes functions to support the processing of peak sets, including overlapping and merging peak sets, counting sequencing reads overlapping intervals in peak sets, and identifying statistically significantly differentially bound sites based on evidence of binding affinity (measured by differences in read densities, see supplementary info 1). For H3K27me3 DiffBind differential peak calling, the initial MACS2 peak list, consisting of 64,264 total peaks (see supplementary info 1), was merged and reduced to a smaller group of larger, broader peaks to reduce noise and false positive discovery (Figure 2B).

### RNA-seq bioinformatics analysis

An average of 44 M paired end 75 bp reads were obtained for each of the eight samples (four control, four infected). Adapter sequence contamination and paired-end reads of poor quality were removed from the raw data. At each step, read quality was assessed with FastQC (version 0.11.5). Any samples that indicated adapter contamination were trimmed via Cutadapt (version 1.15). The raw reads were aligned to the UMD 3.1.1 bovine transcriptome using Salmon (version 0.8.1) [101]. Aligned reads were also counted in Salmon and the resulting quantification files were annotated at gene level via tximport (version 3.7) [102]. The annotated gene counts were then normalised and differential expression analysis performed with DESeq2 (version 1.20.0) [103], correcting for multiple testing using the Benjamini-Hochberg method [104]. Genes identified from ChIP-seq as exhibiting differential histone modifications were cross referenced with the RNA-seq data set to determine significant log2FC between *M. bovis*-infected and control non-infected. Additionally, this RNA-seq data was cross referenced with RNA-seq data from a previous study that investigated bAM infected with *M. bovis* [29].

### MicroRNA-seq bioinformatics analysis

A mean of 26 M paired-end 50 bp reads were obtained for each of the eight samples (four control, four infected). At each step of data processing, read quality was assessed via fastqc (version 0.11.5). Any samples that exhibited adapter contamination were trimmed via Cutadapt (version 1.15) and all reads smaller than 17 bp were removed from the analysis. Raw reads were mapped to UMD3.1 using Bowtie (version 1.2.2). miRNA detection, identification and quantification was carried out with mirdeep2 (version 0.0.91). Isoform analysis was also performed using mirdeep2. Differential expression analysis was performed using DESeq2, correcting for multiple testing with the Benjamini-Hochberg method. Any miRNAs that were significantly differentially expressed (FDR < 0.10) were selected for further analysis. To determine if significantly differentially expressed miRNAs target genes selected in the ChIP seq analysis, miRmap [105] was used to predict the likelihood a specific miRNA targets one or more of the genes based on three criteria: delta G binding, probability exact and phylogenetic conservation of seed site, which is then combined into a single scoring metric (miRmap score). Any predicted gene targets with miRmap score ≥ 0.70 were included in the analysis (see supplementary info 3).

### Pathway analysis

Pathway analysis was carried out on any gene that had a differential peak between control and infected samples. Pathway analysis and gene ontology was carried out by DAVID (version 6.8), Ingenuity pathway analysis (01-13) and PANTHER (version 13.1) [49, 106]. KEGG pathways were selected by choosing pathways that had the highest number of genes identified in the ChIP seq data and had a FDR < 0.05.

### Integration of GWAS data

GWAS data for genetic susceptibility to *M. bovis* infection previously generated by Richardson and colleagues [52] were analysed to determine if subsets of SNPs selected according to their distance to H3K4me3 and PolII active loci were enriched for significant GWAS hits. The nominal *P* values used in this study were generated using single SNP regression analysis in a mixed animal model as described previously [52]. In summary, high-density genotypes (*n* = 597,144) of dairy bulls (*n* = 841) used for artificial insemination were associated with deregressed estimated breeding values for bTB susceptibility that had been calculated from epidemiological information on 105,914 daughters and provided by the Irish Cattle Breeding Federation (ICBF). In this study, the significance of the distribution of SNP nominal *P* values (from Richardson et al 2016) within and up to 100 kb up and downstream to genes identified as having differential H3K4me3 and PolII activity on bTB susceptibility were estimated in R using *q* value (FDRTOOL) and permutation analysis (custom scripts). A total of 1000 samplings (with replacement) from the HD GWAS *P* value data set (*n* = 597,144) representing the size of each of selected SNP subsets were generated. The *q* values for each SNP *P* value subset and all its permuted equivalents were calculated using the FDRTOOL library in R. The subsequent significance level (*P*_perm_) assigned to each of the SNP subsets was equivalent to the proportion of permutations in which at least the same number of *q* values < 0.05 as the SNP subset were obtained, i.e. by chance.

## Abbreviations

bAM: bovine alveolar macrophage/s;
bTB: bovine tuberculosis;
ChIP: chromatin immunoprecipitation;
FDR: false discovery rate;
GWAS: genome-wide association study;
H3K27me3: histone H3 lysine 27 tri-methylation;
H3K4me3: histone H3 lysine 4 tri-methylation;
hpi: hours post infection;
log_2_FC: log_2_ fold change;
PolII: RNA Polymerase II;
SNP: single nucleotide polymorphism;
TB: tuberculosis.

## Acknowledgments

The authors would also like to thank FAANG–Europe for awarding A.M.O.D. a short-term scientific mission (STSM) grant. We would also like to acknowledge Edinburgh Genomics for generation of sequencing data.

## Author Contributions

Conceptualization, A.M.O.D., D.E.M. and T.J.H.; Software and Formal Analysis, T.J.H. and M.P.M.; Investigation, A.M.O.D., D.V. and J.A.B.; Resources, D.E.M., S.V.G. and D.V.; Data Curation, T.J.H.; Writing – original draft, T.J.H, A.M.O.D. and D.E.M.; Writing – review and editing, D.V., S.V.G and M.P.M.; Supervision and Project Administration, D.E.M. and A.M.O.D.; Funding Acquisition; D.E.M, S.V.G, D.V. and A.M.O.D.

## Ethics Statement and Consent to Participate

All animal procedures were performed according to the provisions of Statutory Instrument No. 543/2012 (under Directive 2010/63/EU on the Protection of Animals used for Scientific Purposes). Ethical approval was obtained from the University College Dublin Animal Ethics Committee (protocol number AREC-13-14-Gordon). No human subjects were used for this study.

## Consent for Publication

Not applicable.

