## Supplementary Figures for "Alveolar macrophage chromatin is modified to orchestrate host response to *Mycobacterium bovis* infection"

**Supplementary Figure 1 (following 3 pages): Genome wide signal intensity correlation histograms of raw ChIP-seq reads.**

2D-histograms of the global levels of (A) H3K4me3, (B) PolII and (C) H3K27me3. Reads from each animal and each condition are correlated based on read position and alignment. Plots and values were obtained using the 'Correlate' tool of easeq bioinformatics tool with the window size set to 1000 bp and max values on axes as 200 counts. r values are genome-wide Pearson's correlation coefficients for the datasets.

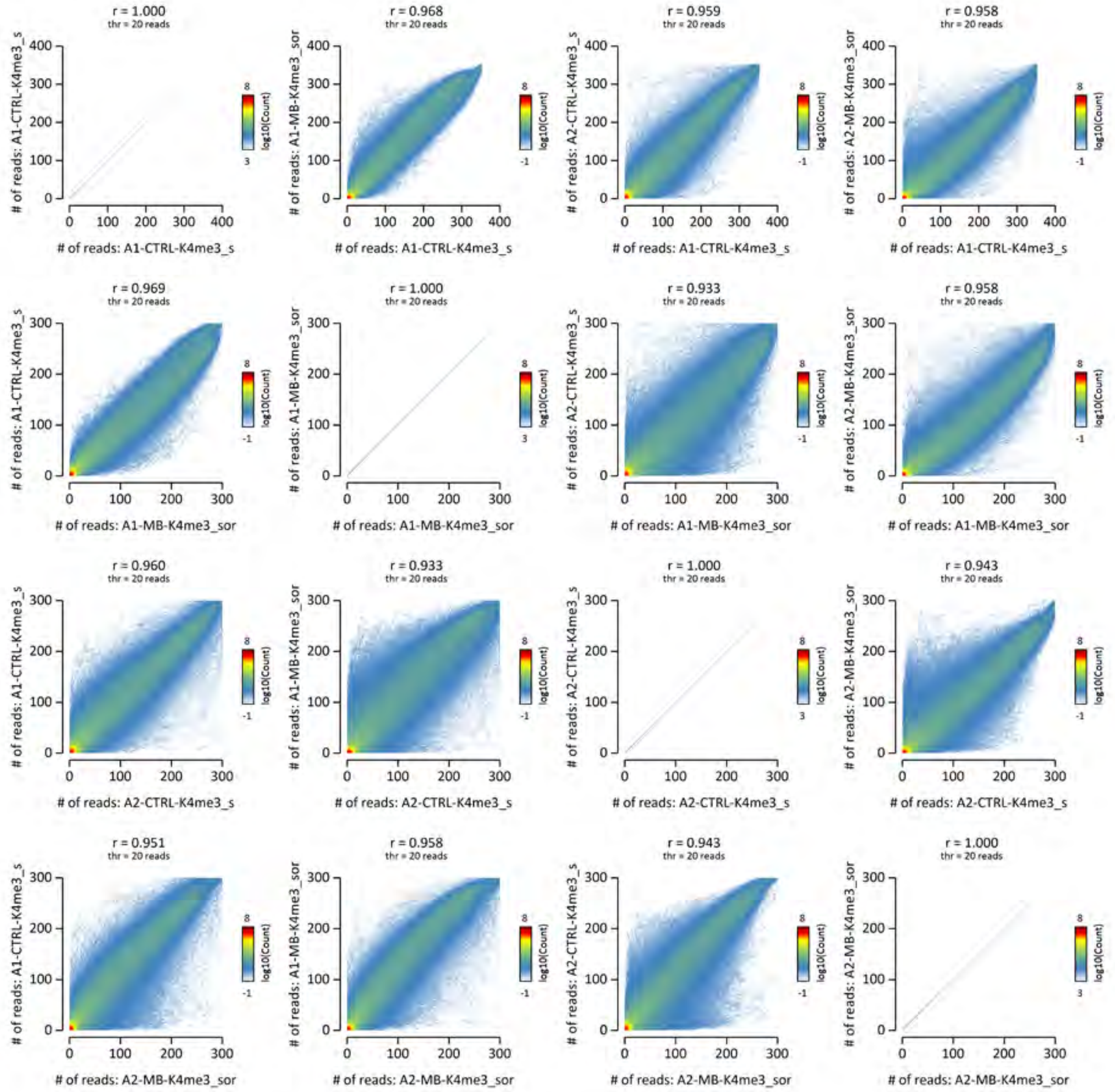
r = 1.000
thr = 20 reads
r = 0.968
thr = 20 reads
r = 0.959
thr = 20 reads
r = 0.958
thr = 20 reads
### of reads: A1-CTRL-K4me3_s
### of reads: A1-MB-K4me3_sor
### of reads: A2-CTRL-K4me3_s
### of reads: A2-MB-K4me3_sor
r = 0.969
thr = 20 reads
r = 1.000
thr = 20 reads
r = 0.933
thr = 20 reads
r = 0.958
thr = 20 reads
r = 0.960
thr = 20 reads
r = 0.933
thr = 20 reads
r = 1.000
thr = 20 reads
r = 0.943
thr = 20 reads
r = 0.951
thr = 20 reads
r = 0.958
thr = 20 reads
r = 0.943
thr = 20 reads
r = 1.000
thr = 20 reads
log10(Count)

A

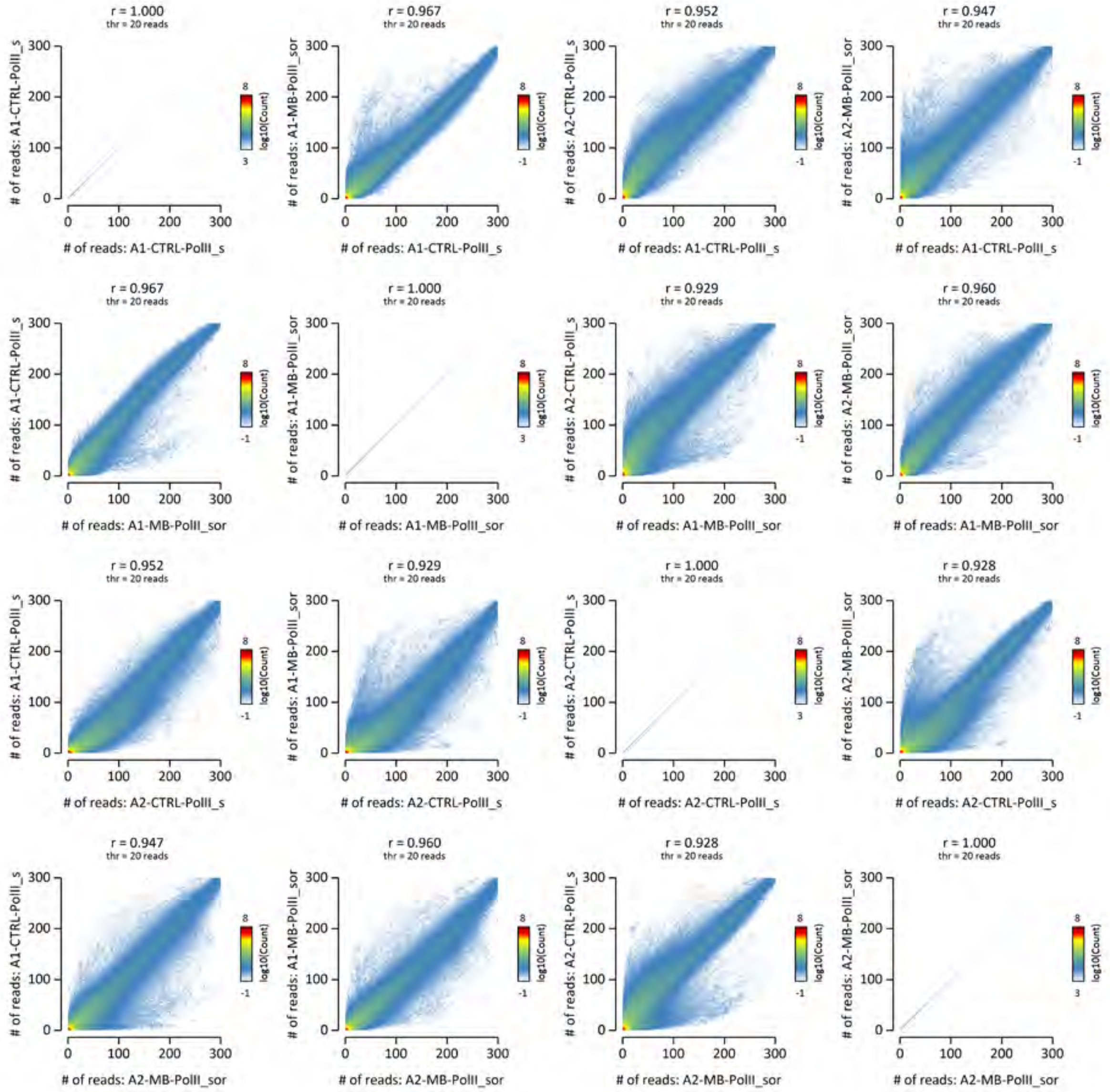
r = 1.000
thr = 20 reads
r = 0.967
r = 0.952
r = 0.947
r = 0.929
r = 0.960
r = 0.928
### of reads: A1-CTRL-PolII_s
### of reads: A1-MB-PolII_sor
### of reads: A2-CTRL-PolII_s
### of reads: A2-MB-PolII_sor
log10(Count)

**B**

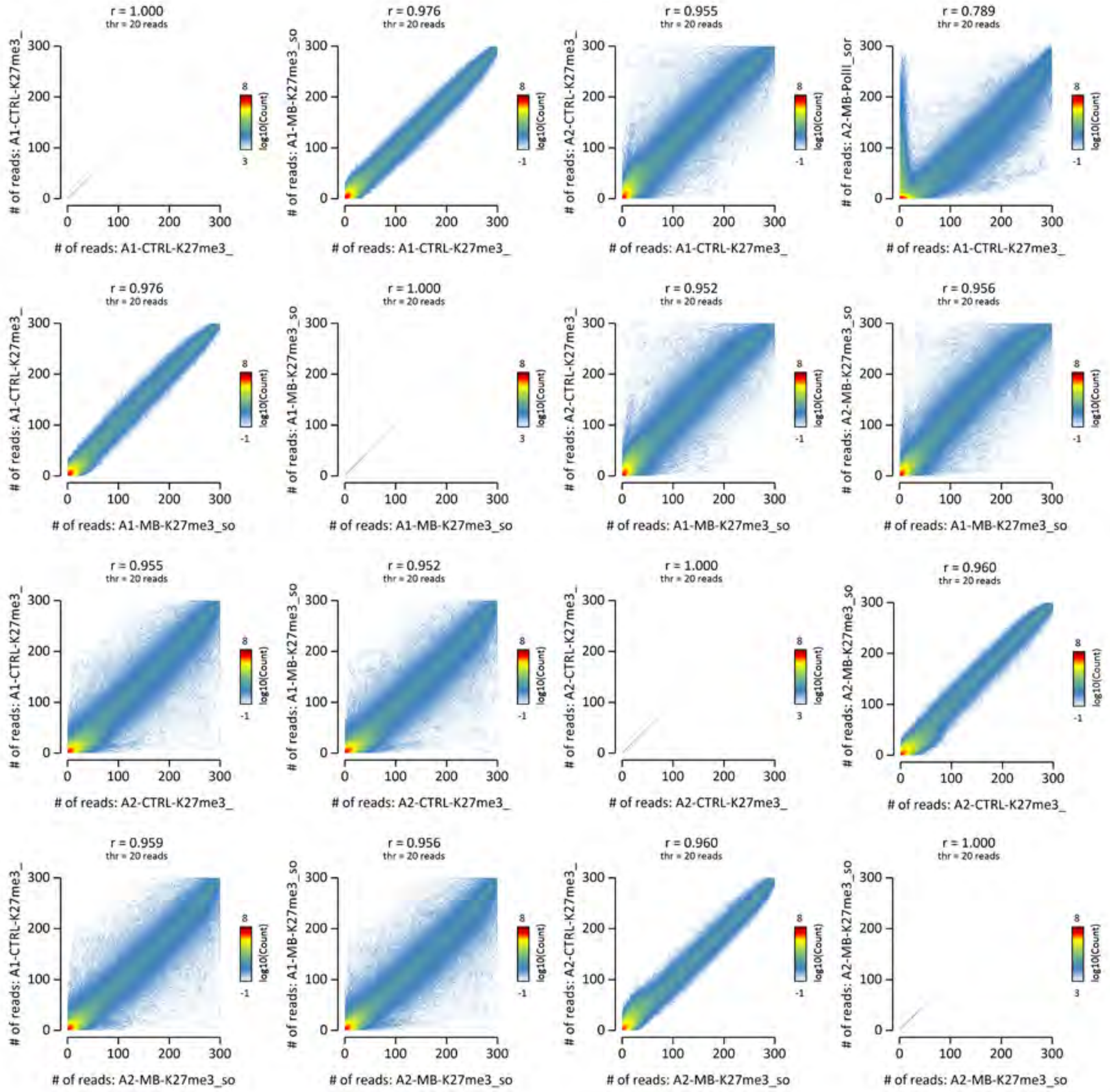
r = 1.000
thr = 20 reads
r = 0.976
r = 0.955
r = 0.789
r = 0.952
r = 0.956
r = 0.960
r = 0.959
### of reads: A1-CTRL-K27me3_
### of reads: A1-MB-K27me3_so
### of reads: A2-CTRL-K27me3_
### of reads: A2-MB-K27me3_so
### of reads: A2-MB-PolII_sor
log10(Count)

C

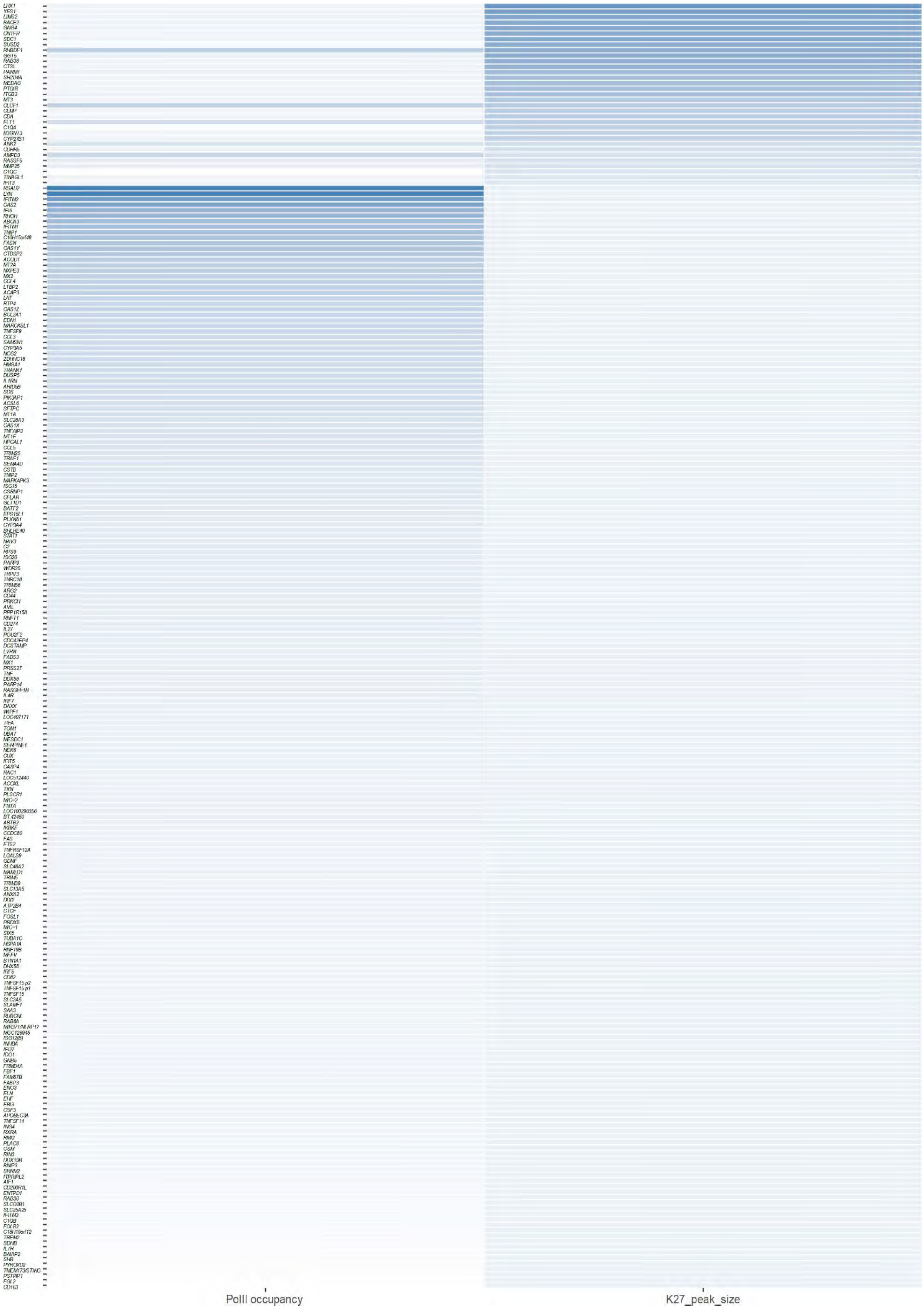
PolII occupancy
K27_peak_size

**Supplementary Figure 2: Correlation heatmap of PolII occupancy and overall H3K27me3 peak size.**

Correlation heatmap demonstrating the relative levels of PolII and H3K27me3 peak density and size with a blue colour scale (white = no peak, dark blue = dense peak) at the TSS of a gene identified as having differential PolII and H3K4me3 peaks (y axis). The plot indicates a negative correlation between the size of PolII peaks and the size of H3K27me3 peaks.

**Supplementary Figure 3 (following 3 pages): Principle component analysis of differential ChIP-seq peaks between conditions.**

PCA plots of (A) H3K4me3, (B) PolII and (C) H3K27me3 ChIP-seq peak differences between control and infected samples generated by R tool DiffBind. H3K4me3 and PolII peaks cluster strongly by condition.

**A**

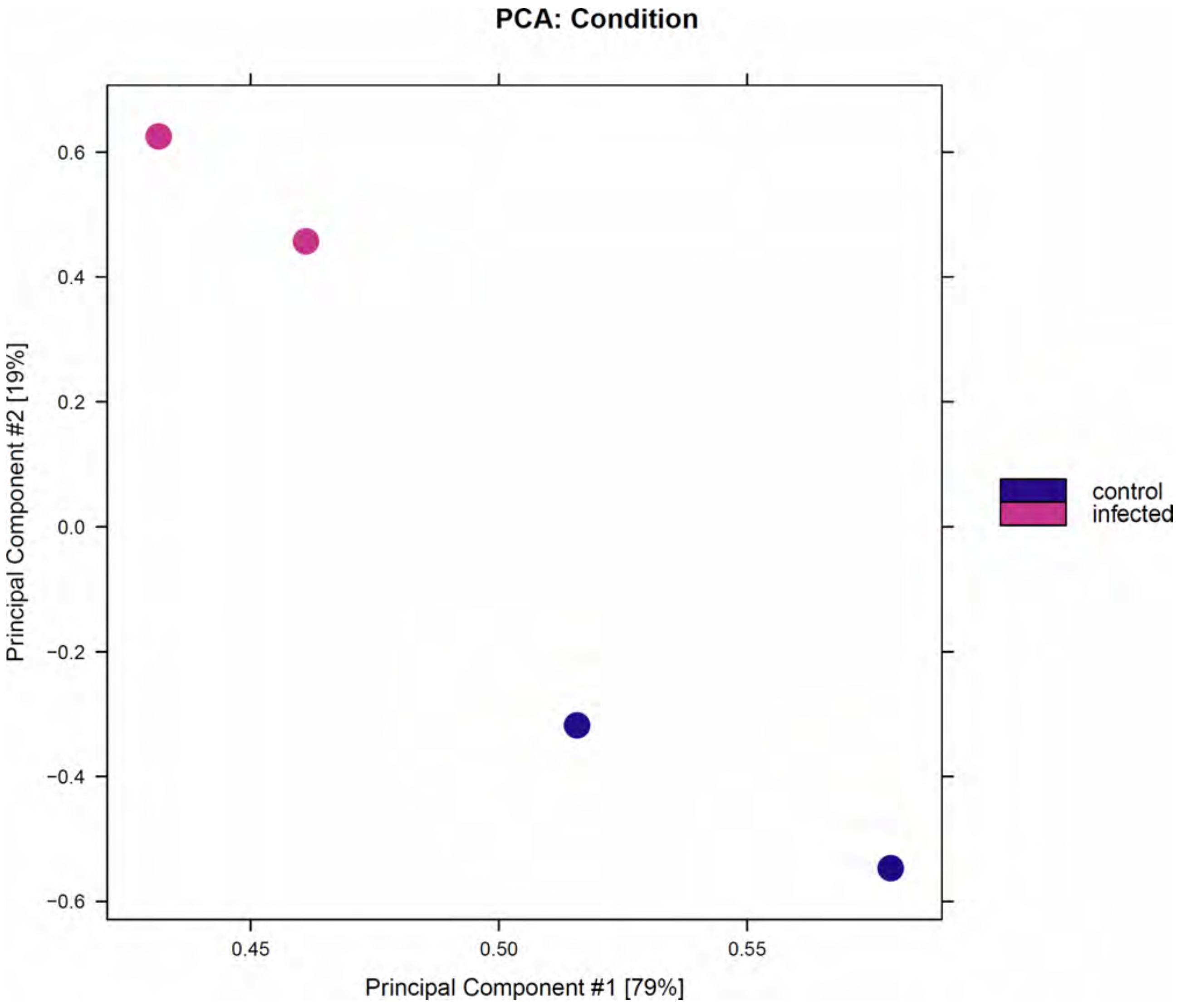
PCA: Condition
Principal Component #2 [19%]
0.6
0.4
0.2
0.0
−0.2
−0.4
−0.6
0.45
0.50
0.55
Principal Component #1 [79%]
control
infected

**B**

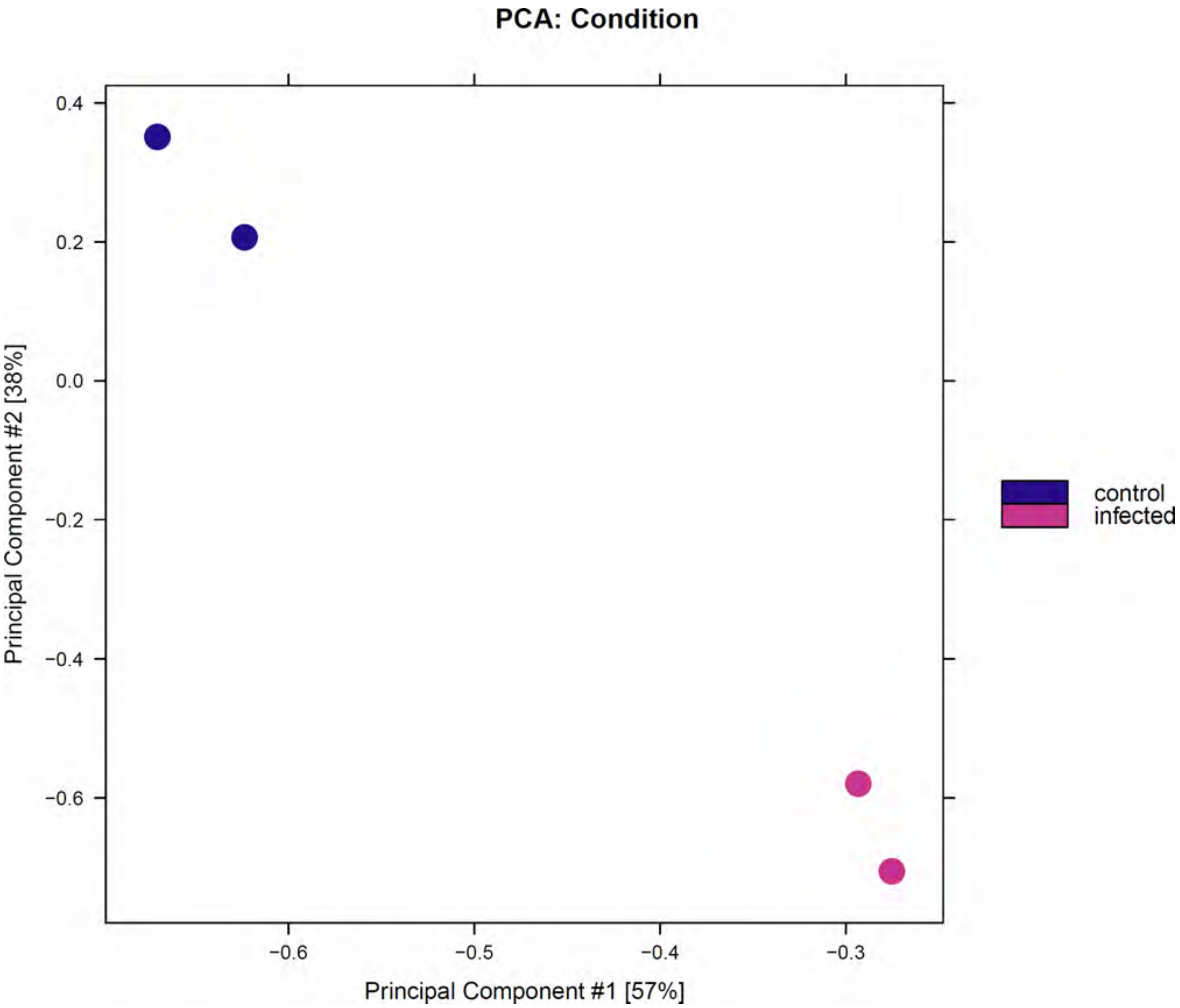
PCA: Condition
0.4
0.2
0.0
−0.2
−0.4
−0.6
Principal Component #2 [38%]
−0.6
−0.5
−0.4
−0.3
Principal Component #1 [57%]
control
infected

**C**

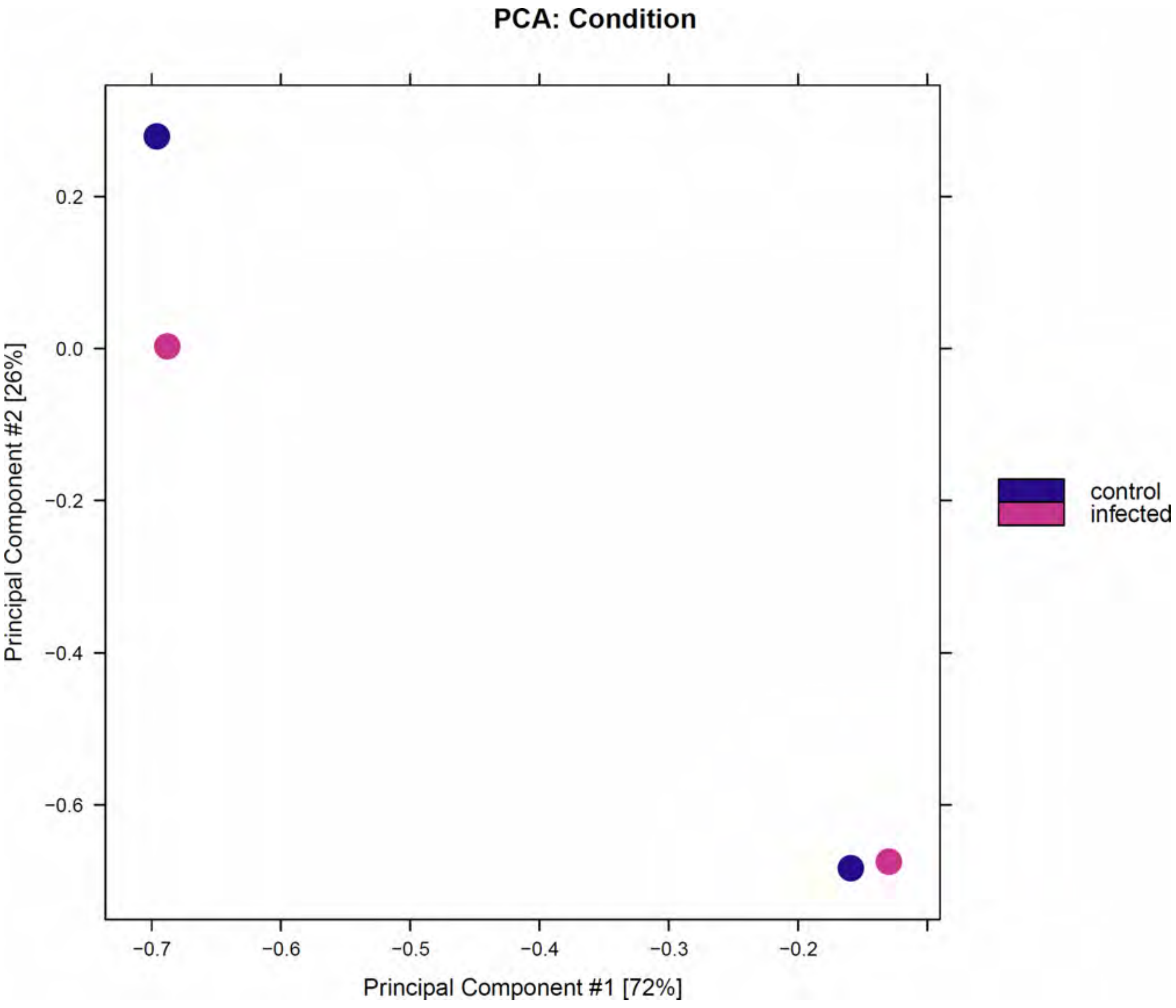
PCA: Condition
0.2
0.0
−0.2
−0.4
−0.6
Principal Component #2 [26%]
−0.7
−0.6
−0.5
−0.4
−0.3
−0.2
Principal Component #1 [72%]
control
infected

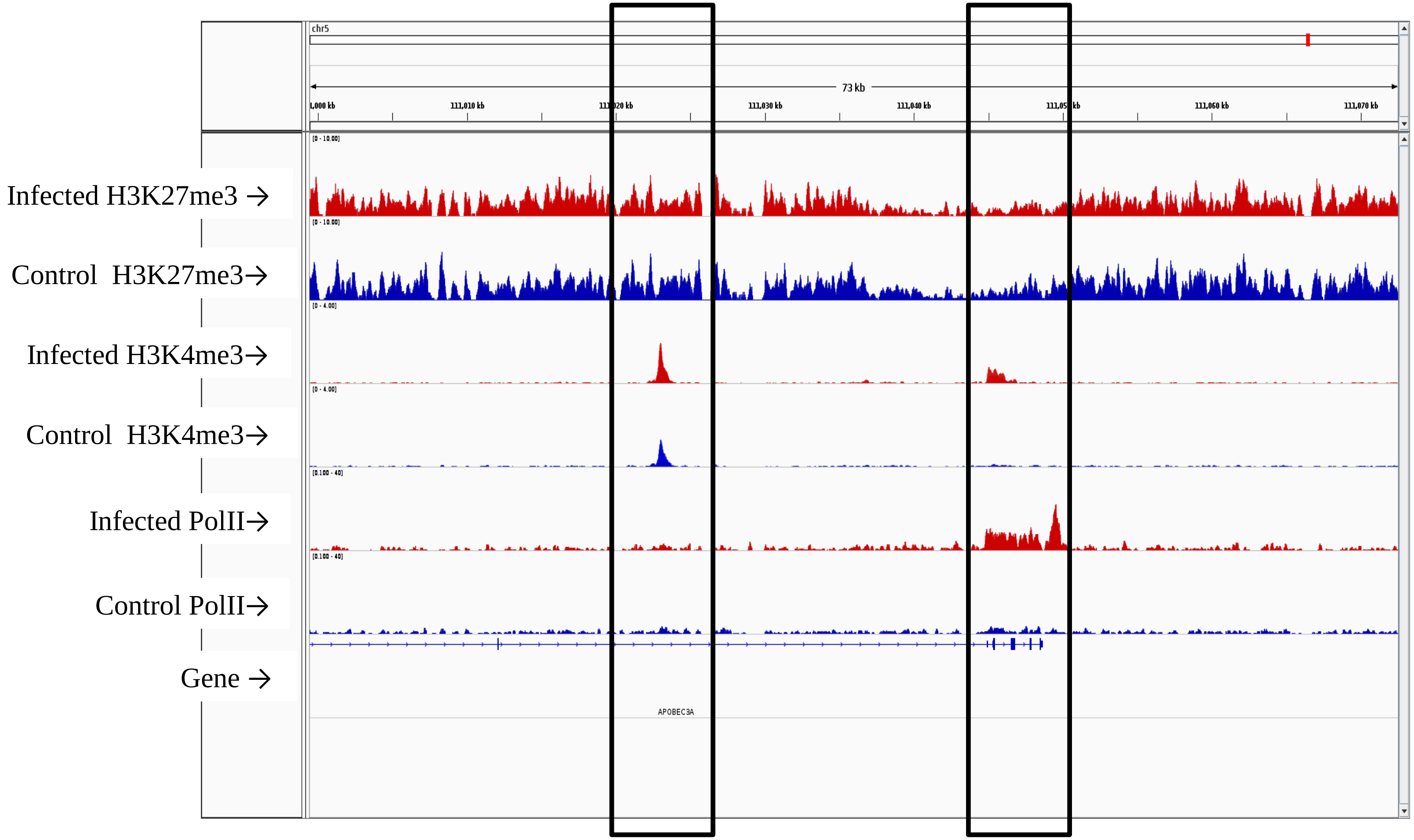


**Supplementary figure 4 (following 235 pages)**. IGV plot showing ChIP-seq tracks. Black open boxes highlight loci that are differentially enriched for methylation at histone 3 lysine 4 (H3K4me3) and/or RNA polymerase II oocupancy.

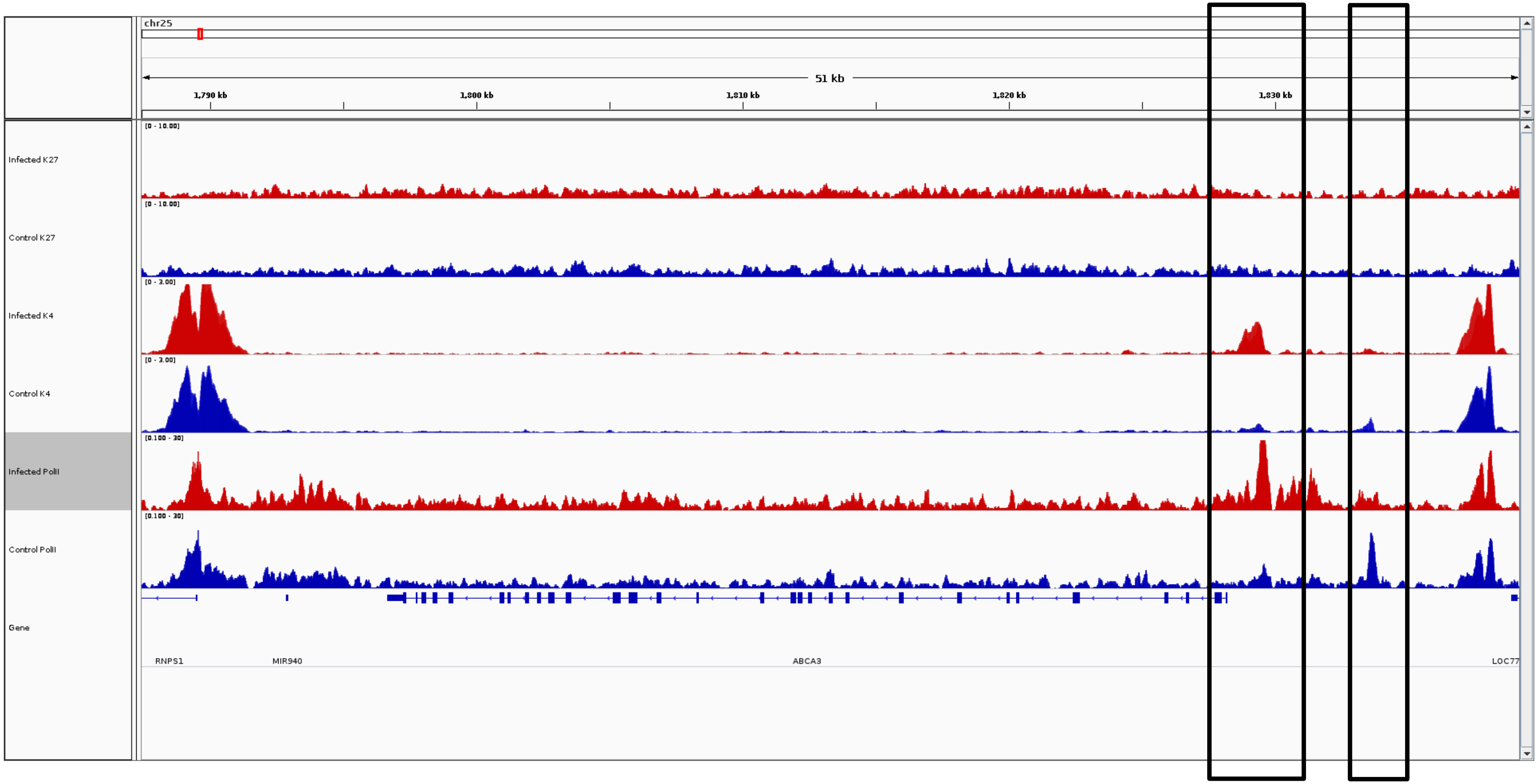
chr25
51 kb
1,790 kb
1,800 kb
1,810 kb
1,820 kb
1,830 kb
Infected K27
Control K27
Infected K4
Control K4
Infected PolII
Control PolII
Gene
RNPS1
MIR940
ABCA3
LOC77

***ABCA3***

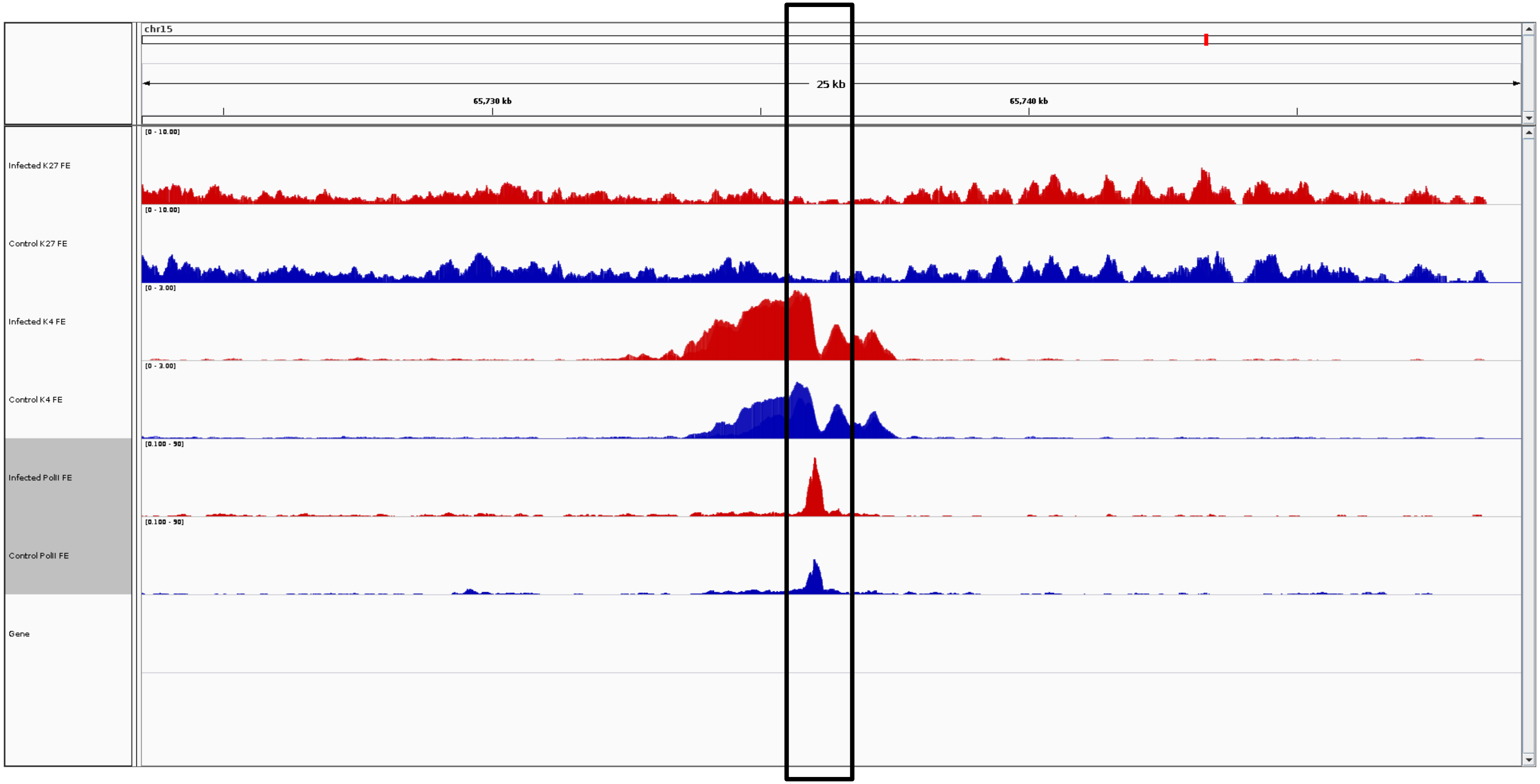

chr15
25 kb
65,730 kb
65,740 kb
[0 - 10.00]
Infected K27 FE
Control K27 FE
[0 - 3.00]
Infected K4 FE
Control K4 FE
[0.100 - 90]
Infected PolII FE
Control PolII FE
Gene


***ABTB2***

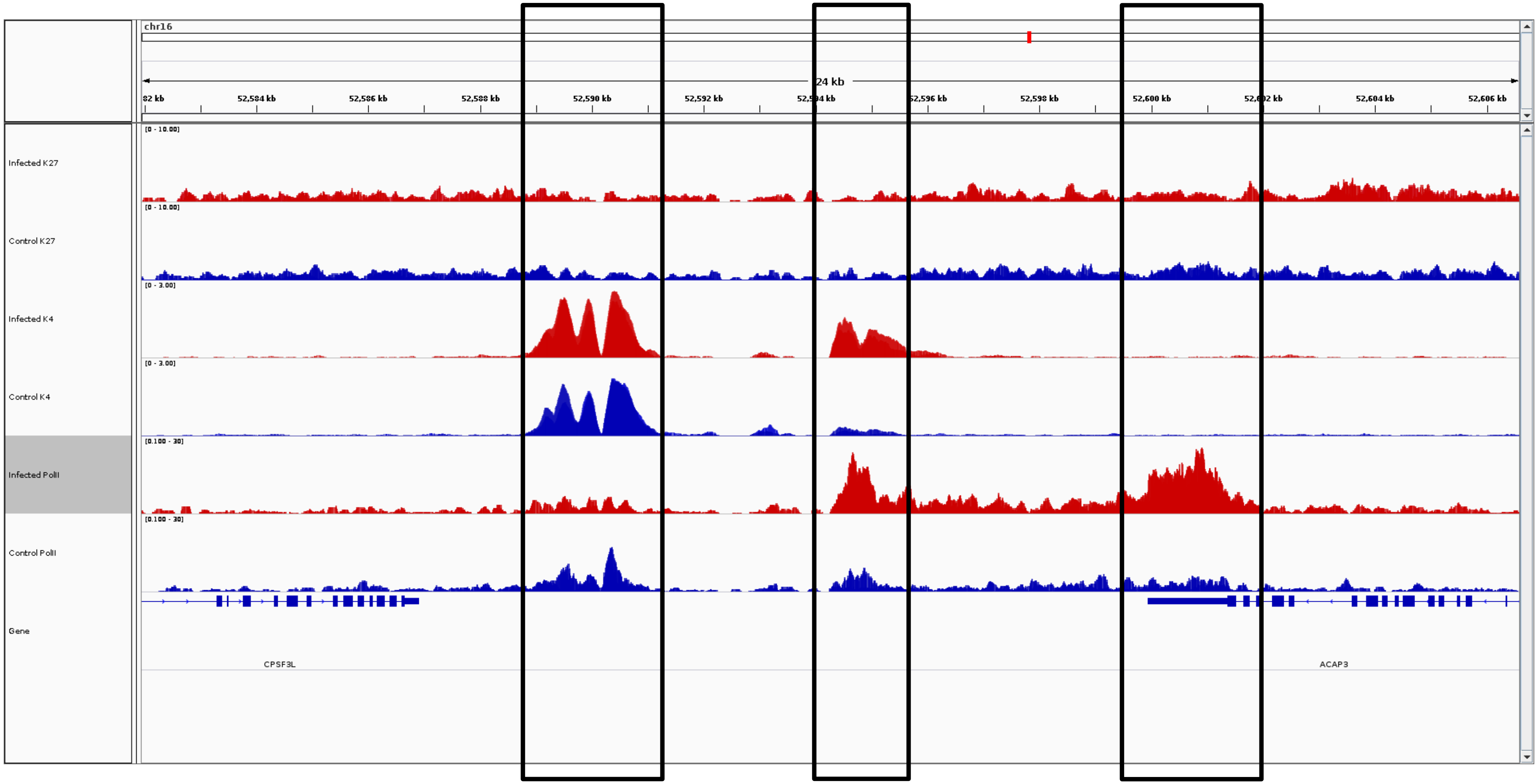
chr16
24 kb
82 kb
52,584 kb
52,586 kb
52,588 kb
52,590 kb
52,592 kb
52,596 kb
52,598 kb
52,600 kb
52,604 kb
52,606 kb
Infected K27
[0 - 10.00]
Control K27
[0 - 10.00]
Infected K4
[0 - 3.00]
Control K4
[0 - 3.00]
Infected PolII
[0.100 - 30]
Control PolII
[0.100 - 30]
Gene
CPSF3L
ACAP3

***ACAP3***

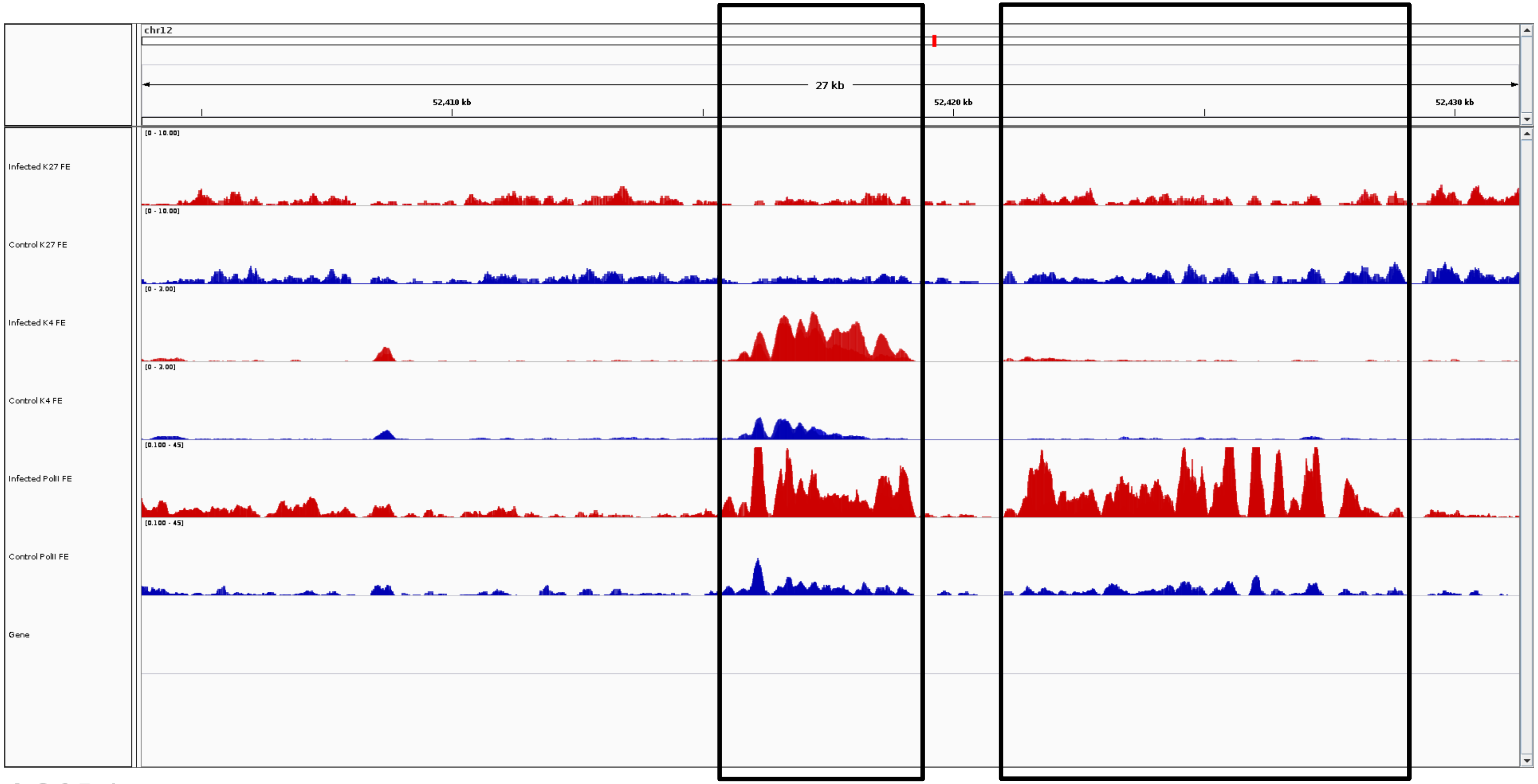
chr12
27 kb
52,410 kb
52,420 kb
52,430 kb
Infected K27 FE
[0 - 10.00]
Control K27 FE
[0 - 10.00]
Infected K4 FE
[0 - 3.00]
Control K4 FE
[0 - 3.00]
Infected PolII FE
[0.100 - 45]
Control PolII FE
[0.100 - 45]
Gene

***ACOD1***

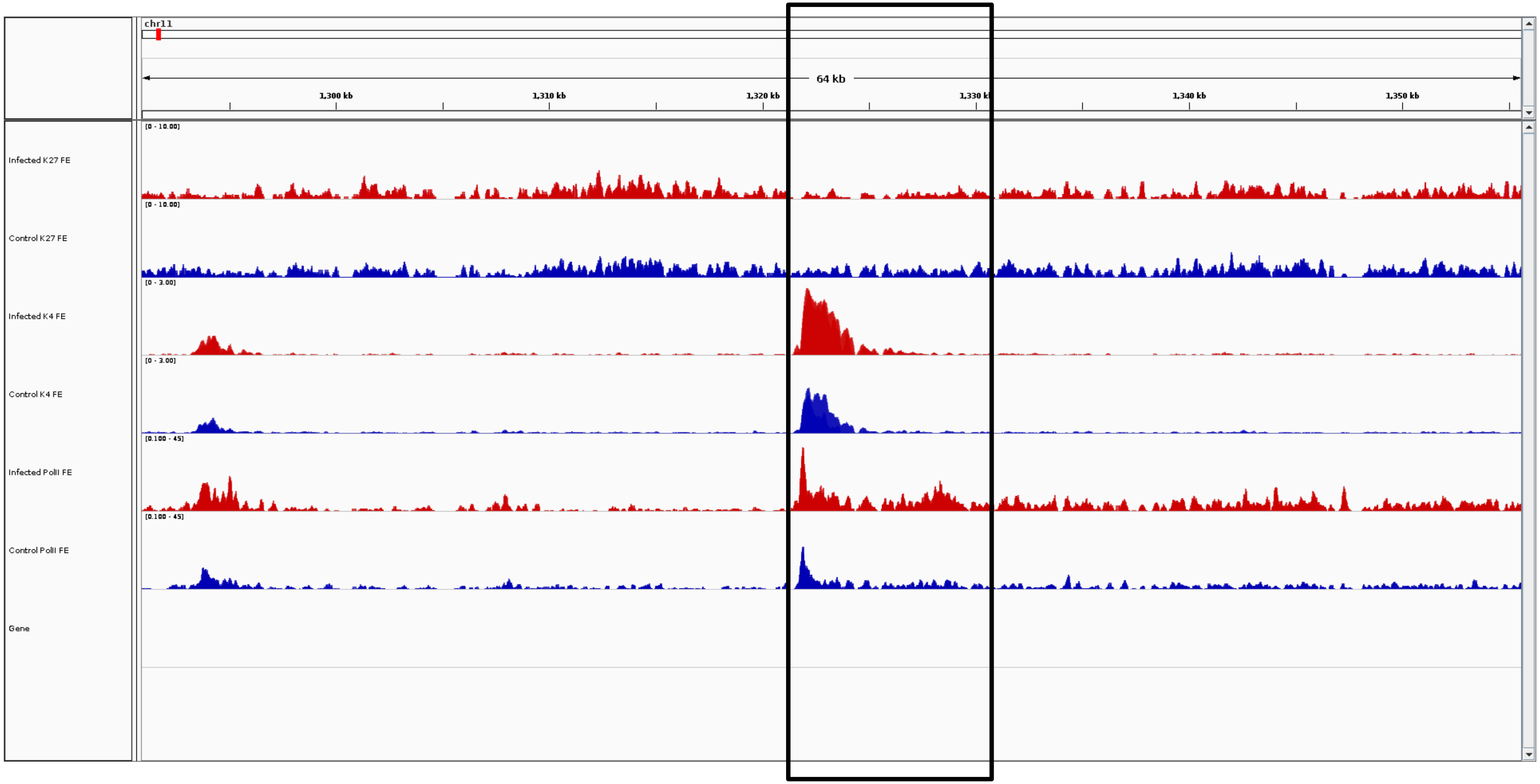
chr11
64 kb
1,300 kb
1,310 kb
1,320 kb
1,340 kb
1,350 kb
Infected K27 FE
[0 - 10.00]
Control K27 FE
[0 - 10.00]
Infected K4 FE
[0 - 3.00]
Control K4 FE
[0 - 3.00]
Infected PolII FE
[0.100 - 45]
Control PolII FE
[0.100 - 45]
Gene

***ACOXL***

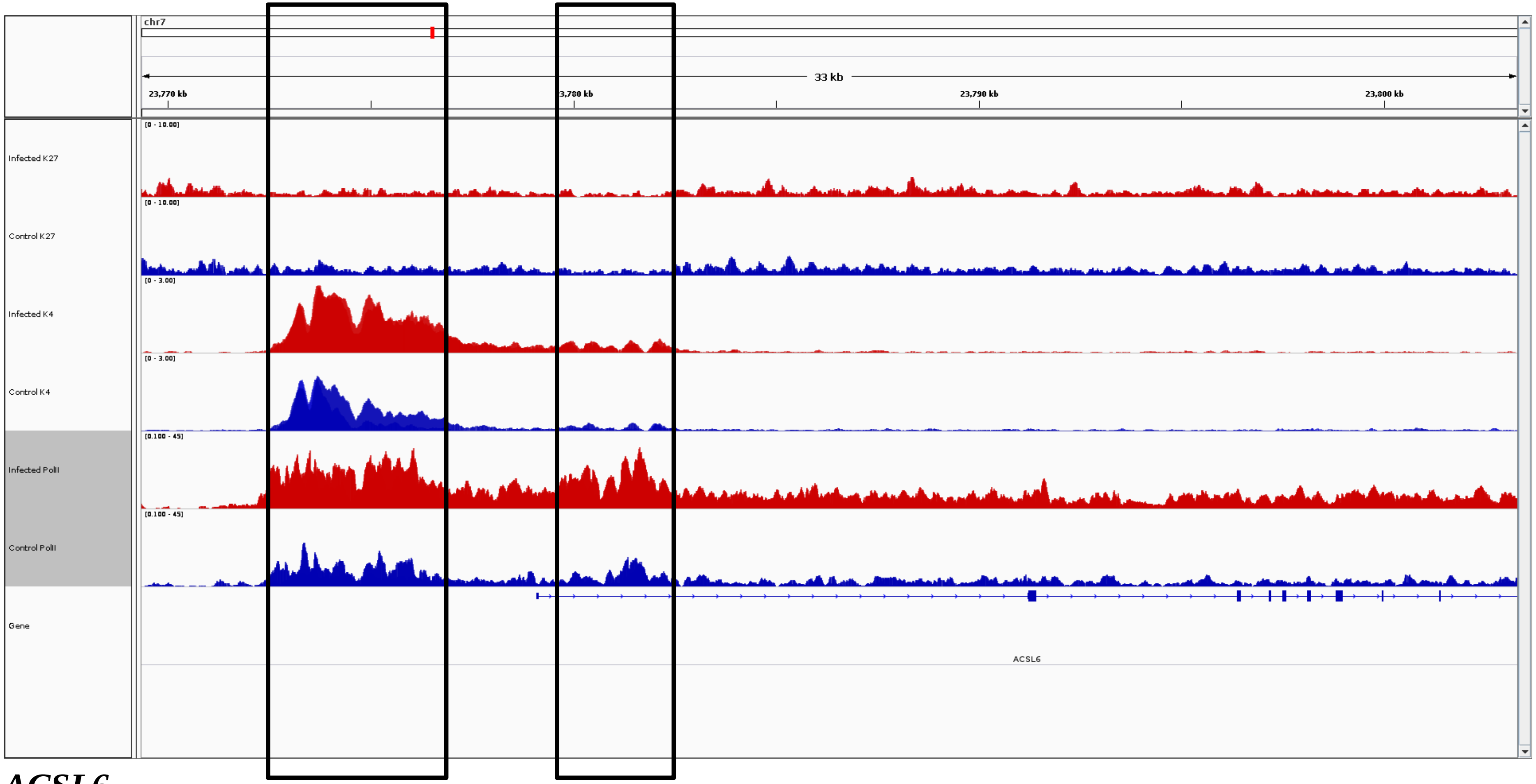
chr7
33 kb
23,770 kb
3,780 kb
23,790 kb
23,800 kb
Infected K27
Control K27
Infected K4
Control K4
Infected PolII
Control PolII
Gene
ACSL6

***ACSL6***

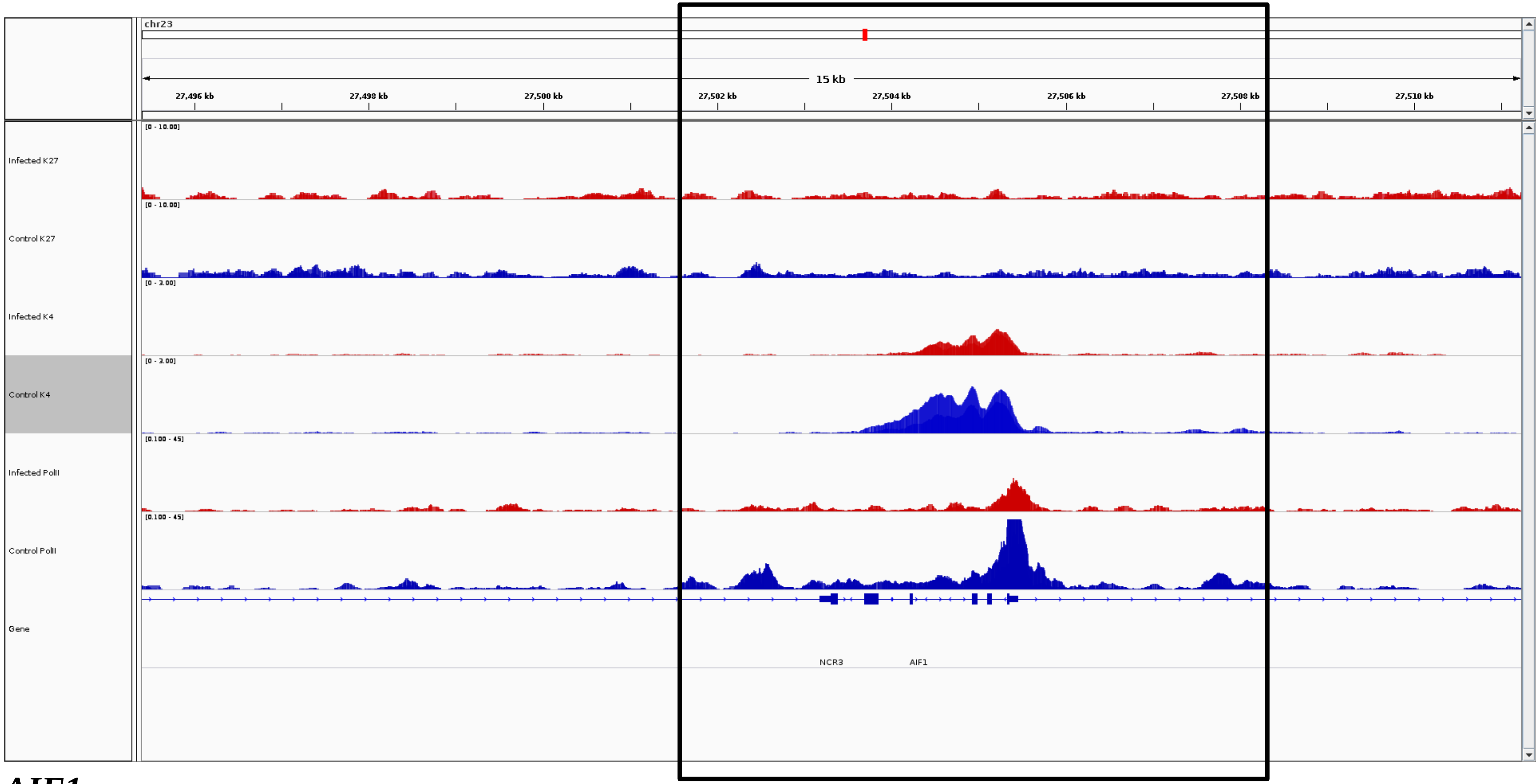
chr23
15 kb
27,496 kb
27,498 kb
27,500 kb
27,502 kb
27,504 kb
27,506 kb
27,508 kb
27,510 kb
Infected K27
[0 - 10.00]
Control K27
[0 - 10.00]
Infected K4
[0 - 3.00]
Control K4
[0 - 3.00]
Infected PolII
[0.100 - 45]
Control PolII
[0.100 - 45]
Gene
NCR3
AIF1

***AIF1***

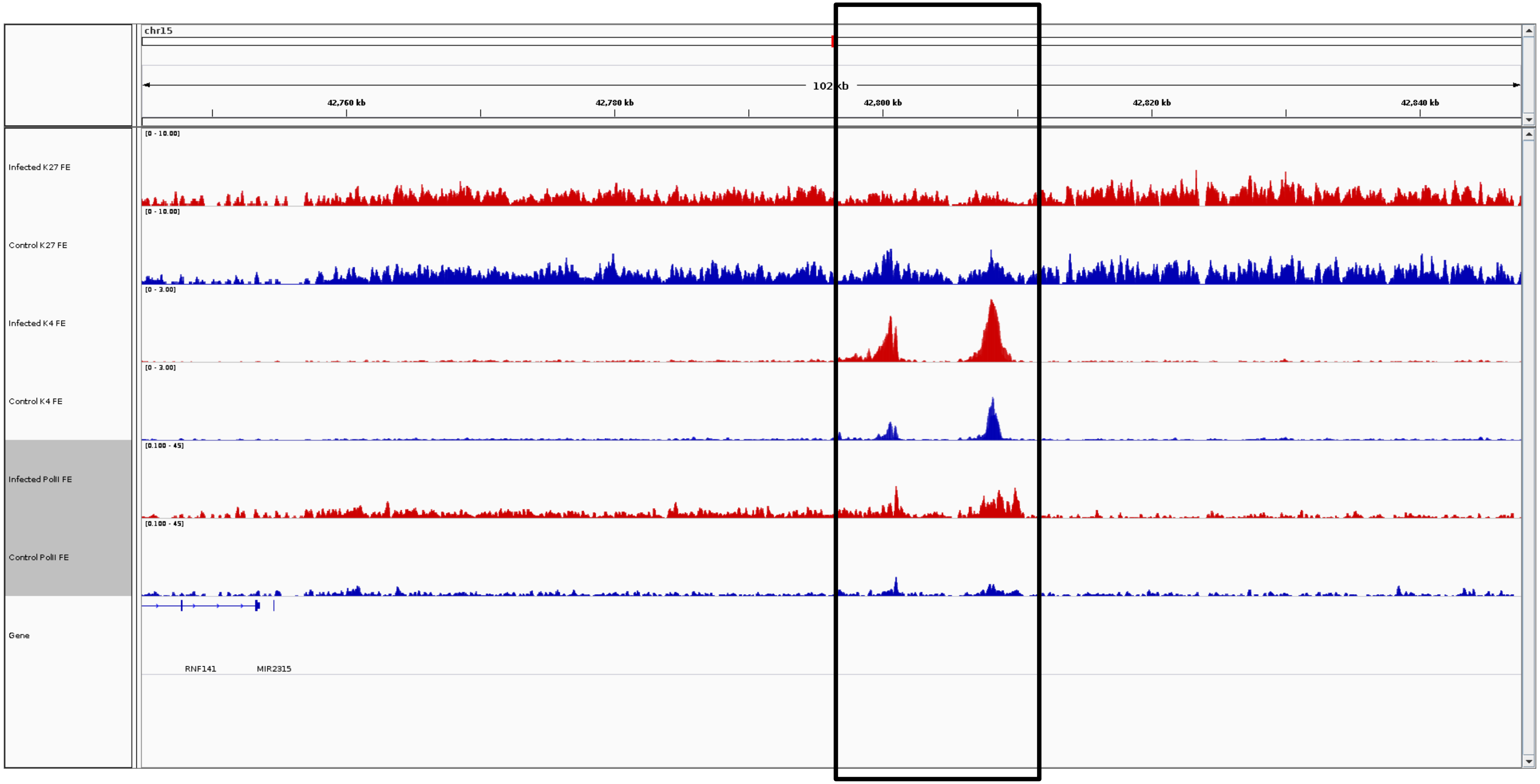
chr15
102 kb
42,760 kb
42,780 kb
42,800 kb
42,820 kb
42,840 kb
Infected K27 FE
Control K27 FE
Infected K4 FE
Control K4 FE
Infected PolII FE
Control PolII FE
Gene
RNF141
MIR2315

***AMPD1***

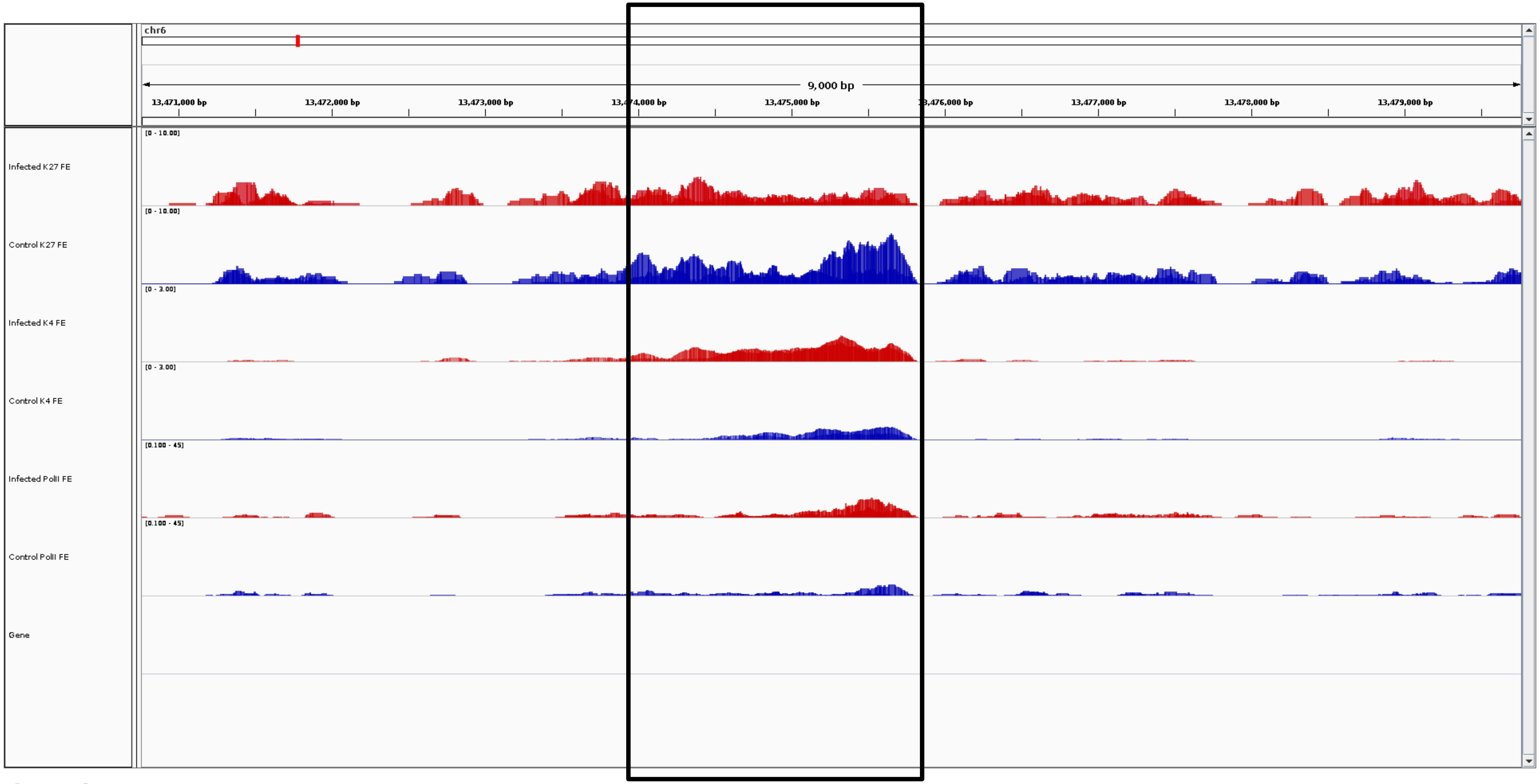
chr6
9,000 bp
13,471,000 bp
13,472,000 bp
13,473,000 bp
13,474,000 bp
13,475,000 bp
13,476,000 bp
13,477,000 bp
13,478,000 bp
13,479,000 bp
Infected K27 FE
[0 - 10.00]
Control K27 FE
[0 - 10.00]
Infected K4 FE
[0 - 3.00]
Control K4 FE
[0 - 3.00]
Infected PolII FE
[0.100 - 45]
Control PolII FE
[0.100 - 45]
Gene

***ANK2***

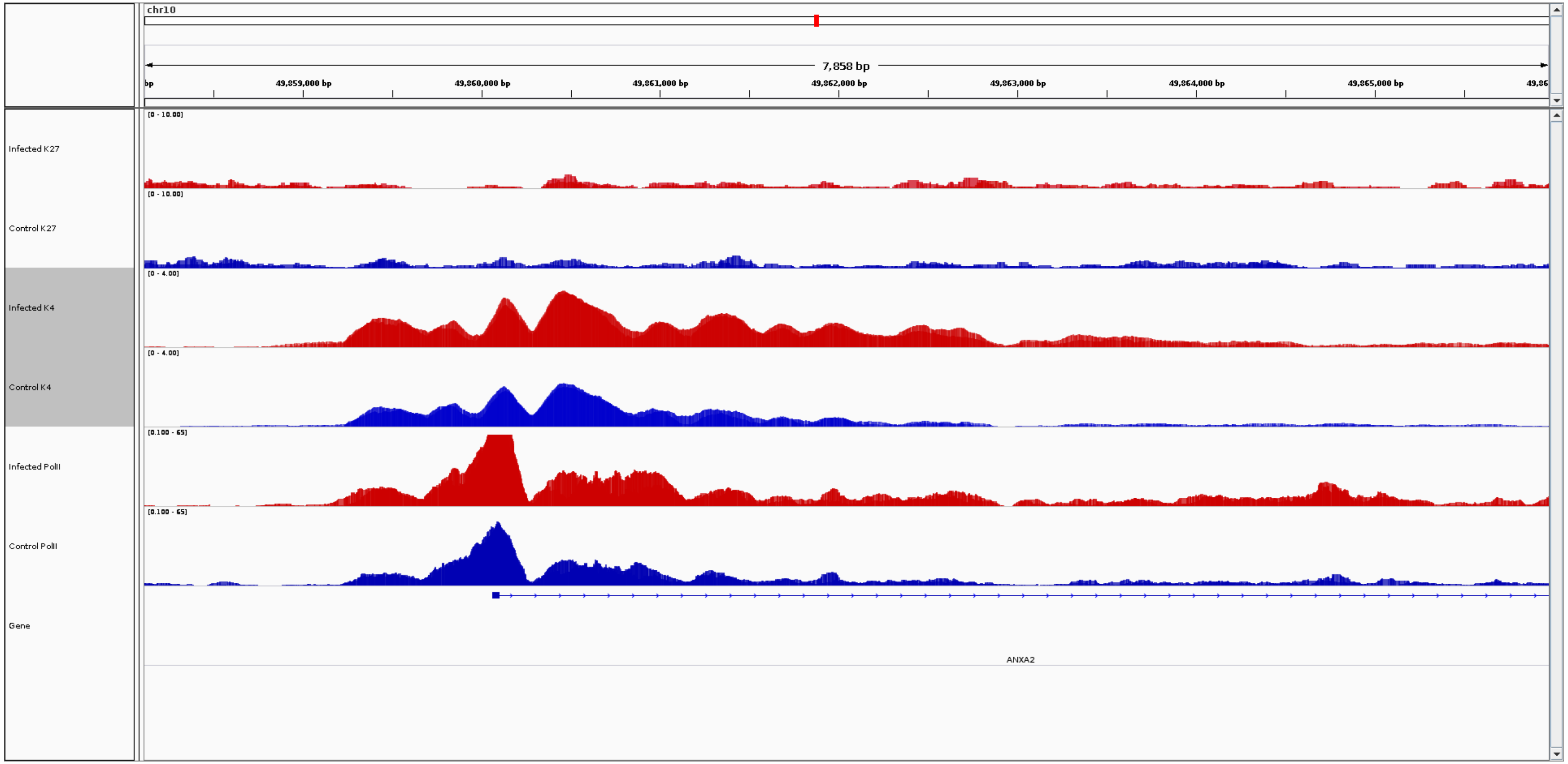
chr10
7,858 bp
49,859,000 bp
49,860,000 bp
49,861,000 bp
49,862,000 bp
49,863,000 bp
49,864,000 bp
49,865,000 bp
Infected K27
[0 - 10.00]
Control K27
[0 - 10.00]
Infected K4
[0 - 4.00]
Control K4
[0 - 4.00]
Infected PolII
[0.100 - 65]
Control PolII
[0.100 - 65]
Gene
ANXA2

***ANXA2***

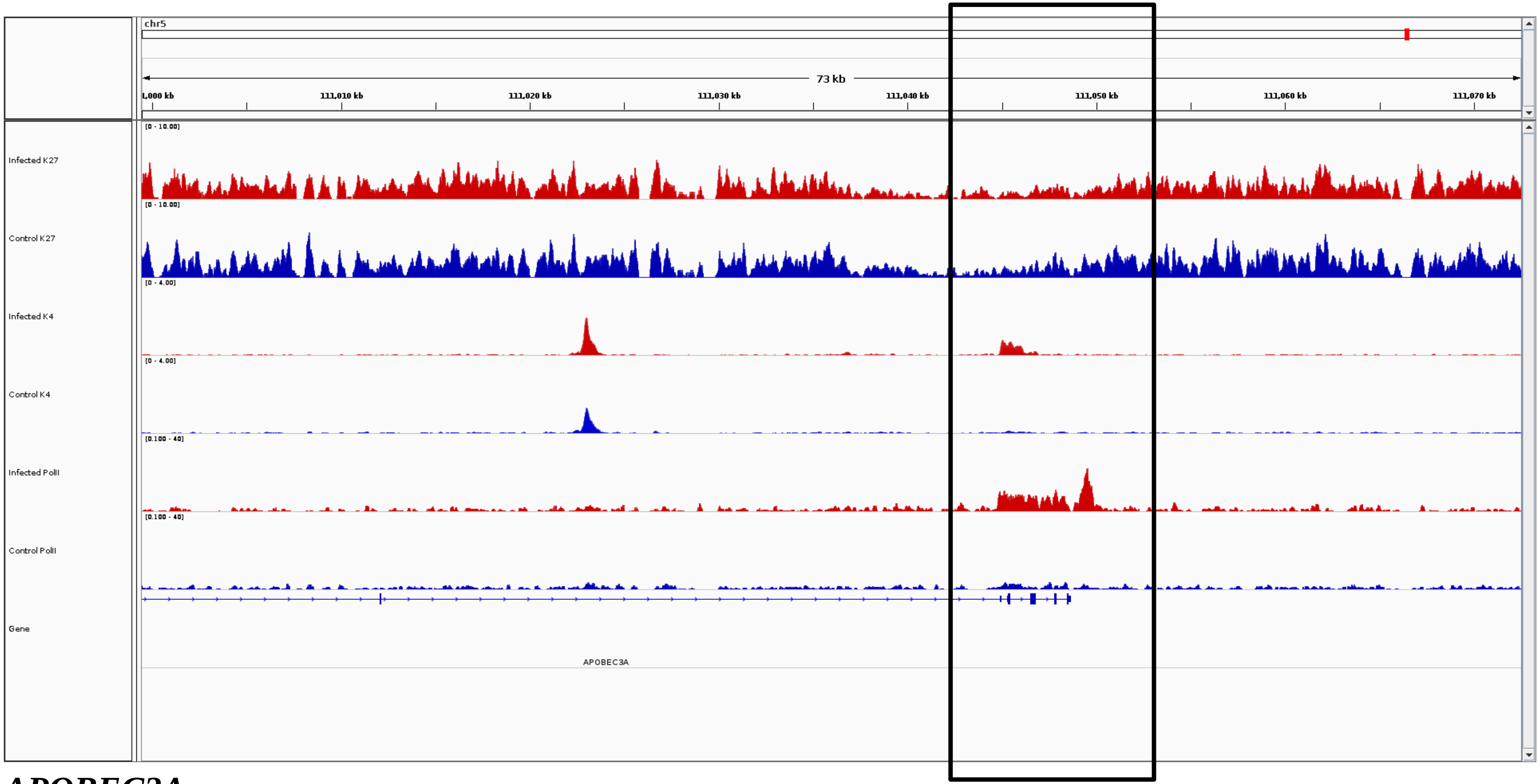
chr5
73 kb
1,000 kb
111,010 kb
111,020 kb
111,030 kb
111,040 kb
111,050 kb
111,060 kb
111,070 kb
Infected K27
[0 - 10.00]
Control K27
[0 - 10.00]
Infected K4
[0 - 4.00]
Control K4
[0 - 4.00]
Infected PolII
[0.100 - 40]
Control PolII
[0.100 - 40]
Gene
APOBEC3A

***APOBEC3A***

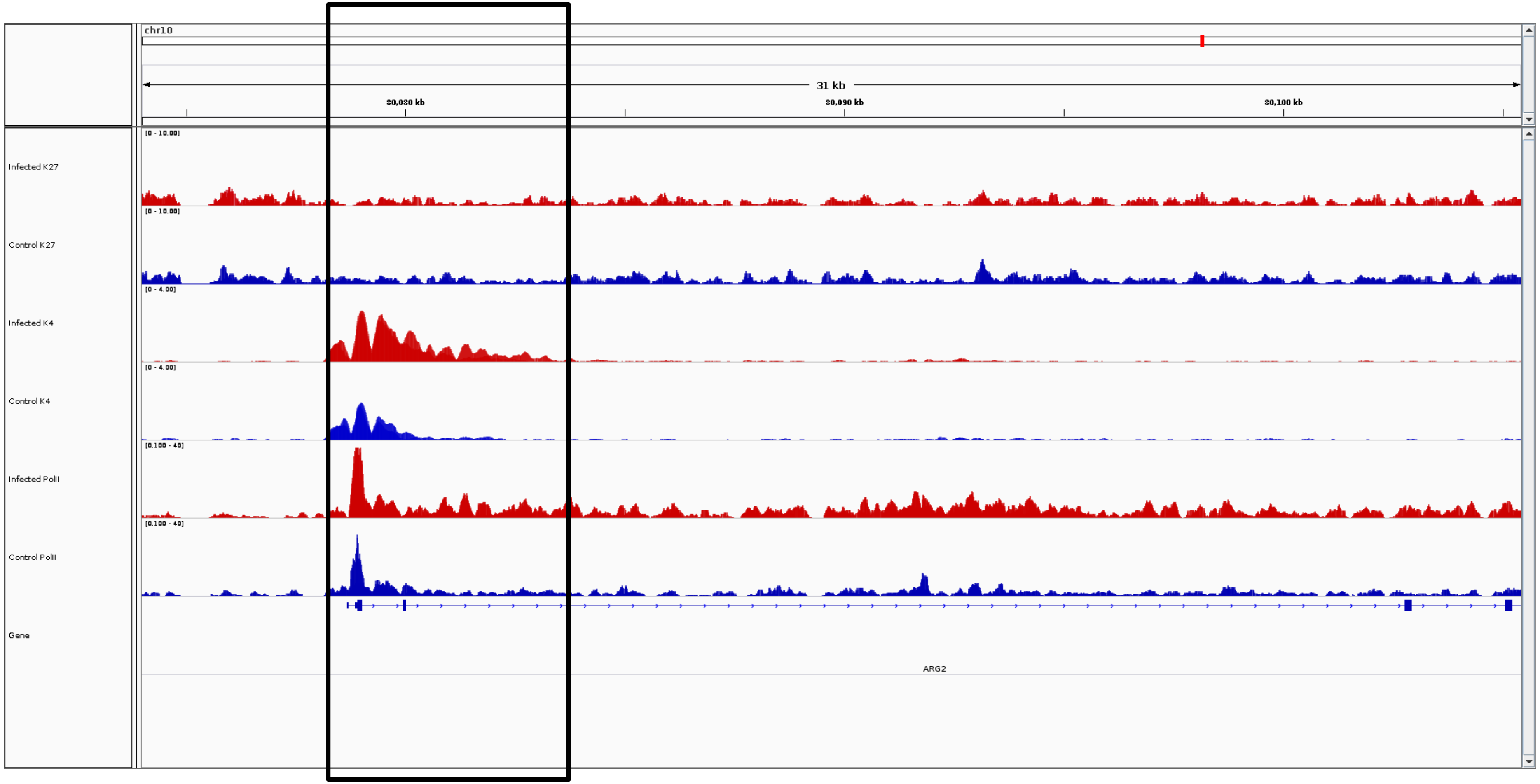
chr10
31 kb
80,080 kb
80,090 kb
80,100 kb
Infected K27
[0 - 10.00]
Control K27
[0 - 10.00]
Infected K4
[0 - 4.00]
Control K4
[0 - 4.00]
Infected PolII
[0.100 - 40]
Control PolII
[0.100 - 40]
Gene
ARG2

***ARG2***

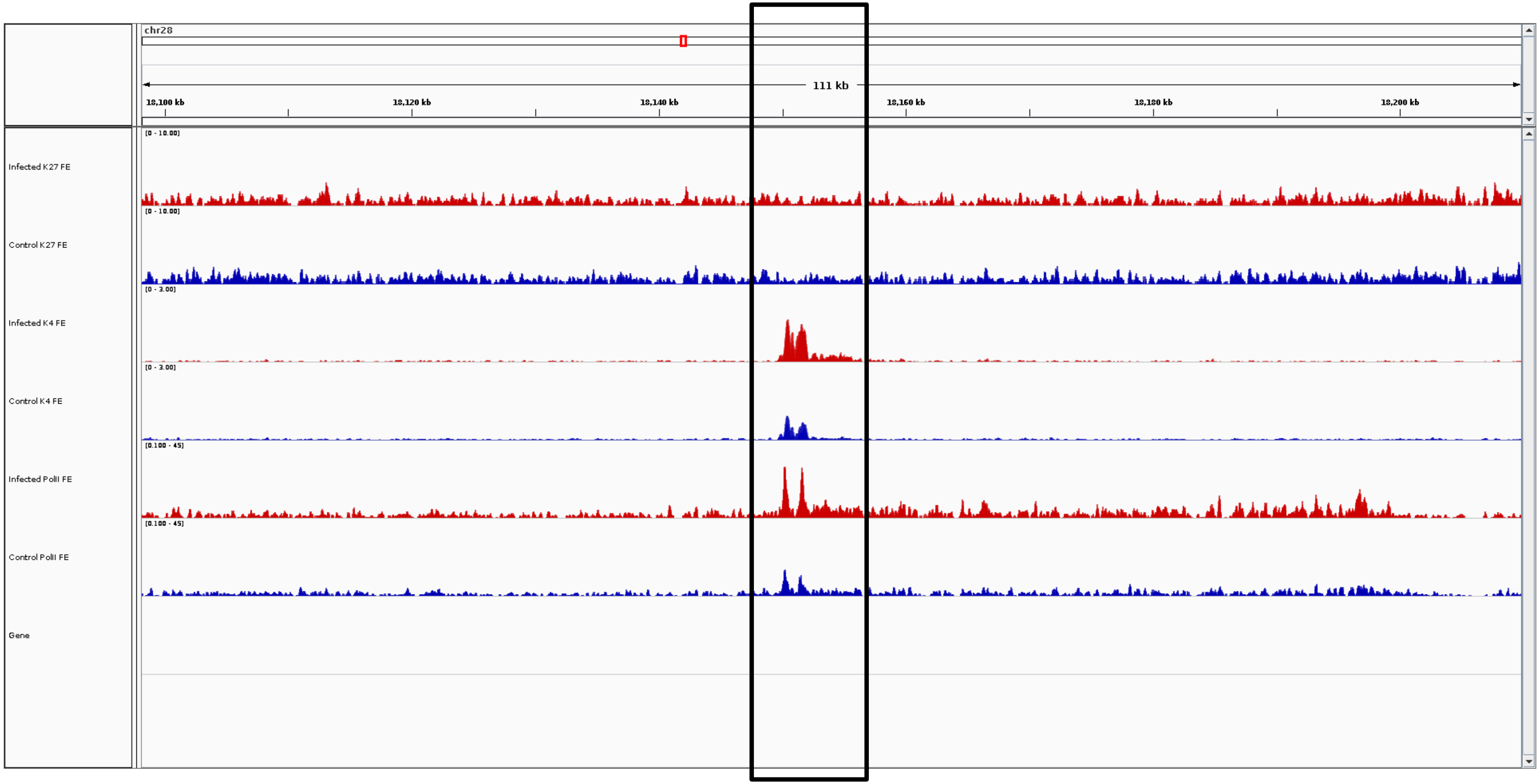
chr28
111 kb
18,100 kb
18,120 kb
18,140 kb
18,160 kb
18,180 kb
18,200 kb
Infected K27 FE
Control K27 FE
Infected K4 FE
Control K4 FE
Infected PolII FE
Control PolII FE
Gene

***ARID5B***

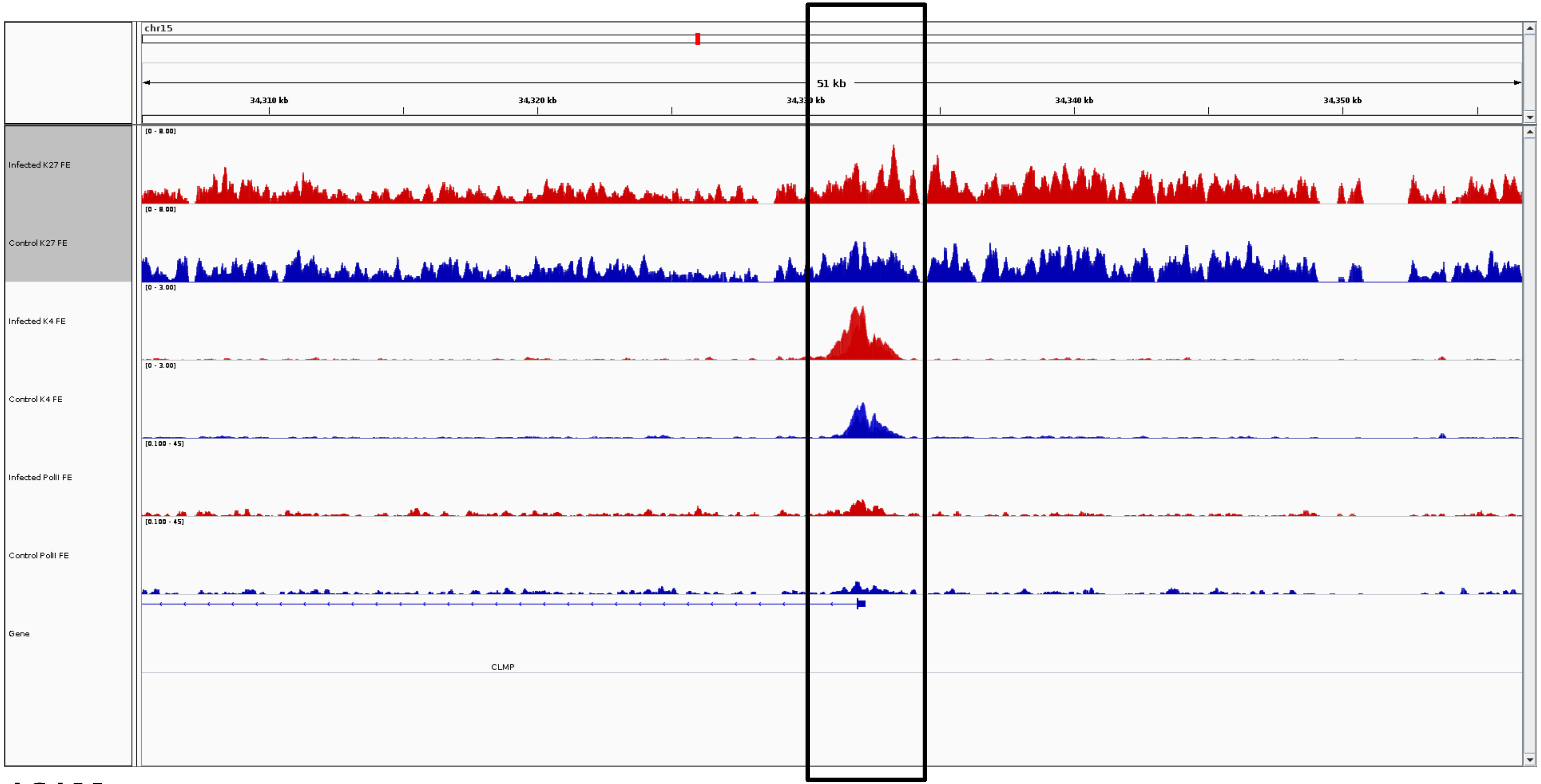
chr15
51 kb
34,310 kb
34,320 kb
34,330 kb
34,340 kb
34,350 kb
[0 - 8.00]
Infected K27 FE
[0 - 8.00]
Control K27 FE
[0 - 3.00]
Infected K4 FE
[0 - 3.00]
Control K4 FE
[0.100 - 45]
Infected PolII FE
[0.100 - 45]
Control PolII FE
Gene
CLMP

***ASAM***

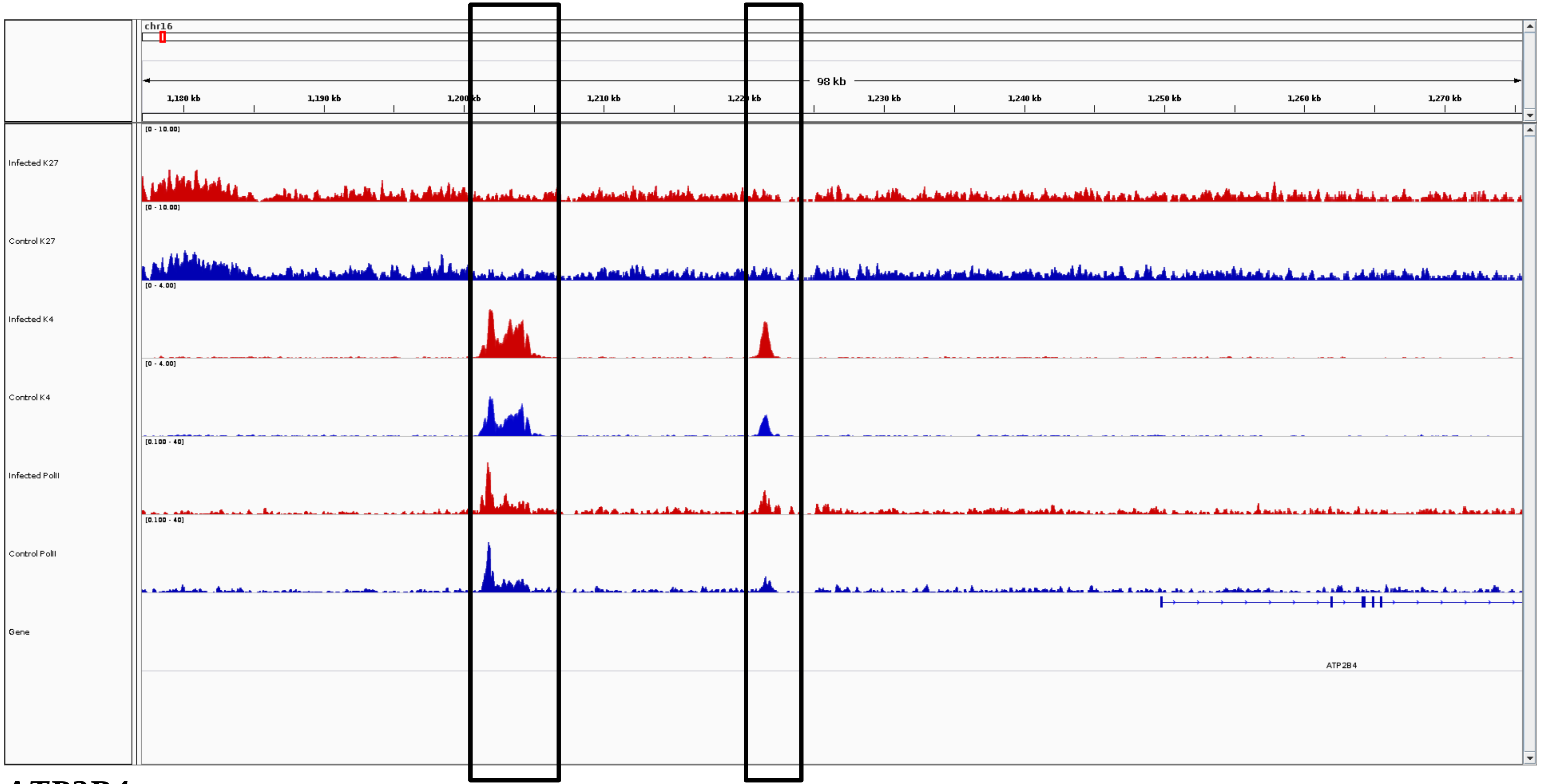
chr16
98 kb
1,180 kb
1,190 kb
1,200 kb
1,210 kb
1,230 kb
1,240 kb
1,250 kb
1,260 kb
1,270 kb
Infected K27
Control K27
Infected K4
Control K4
Infected PolII
Control PolII
Gene
ATP2B4

***ATP2B4***

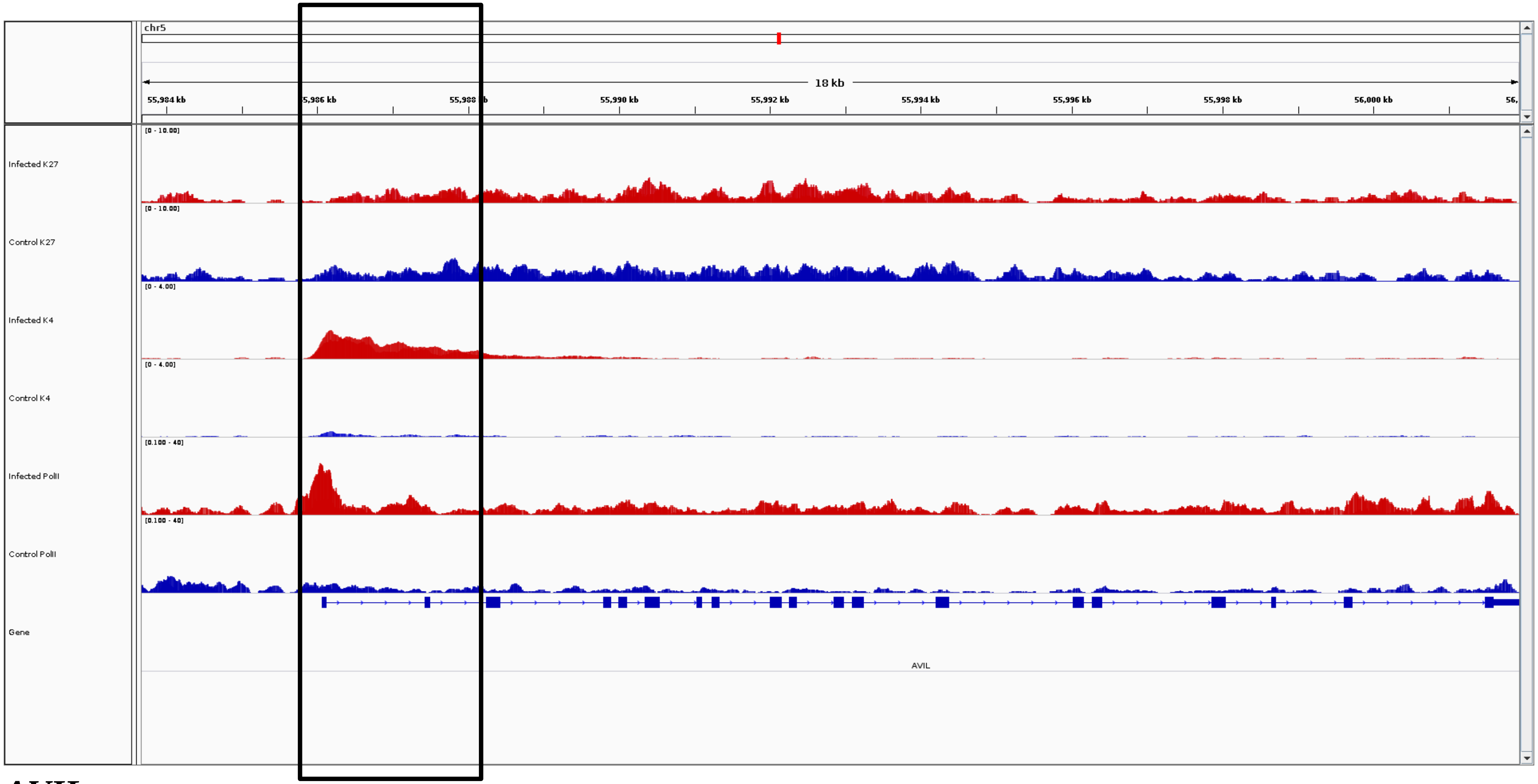
chr5
18 kb
55,984 kb
5,986 kb
55,988
55,990 kb
55,992 kb
55,994 kb
55,996 kb
55,998 kb
56,000 kb
Infected K27
[0 - 10.00]
Control K27
[0 - 10.00]
Infected K4
[0 - 4.00]
Control K4
[0 - 4.00]
Infected PolII
[0.100 - 40]
Control PolII
[0.100 - 40]
Gene
AVIL

***AVIL***

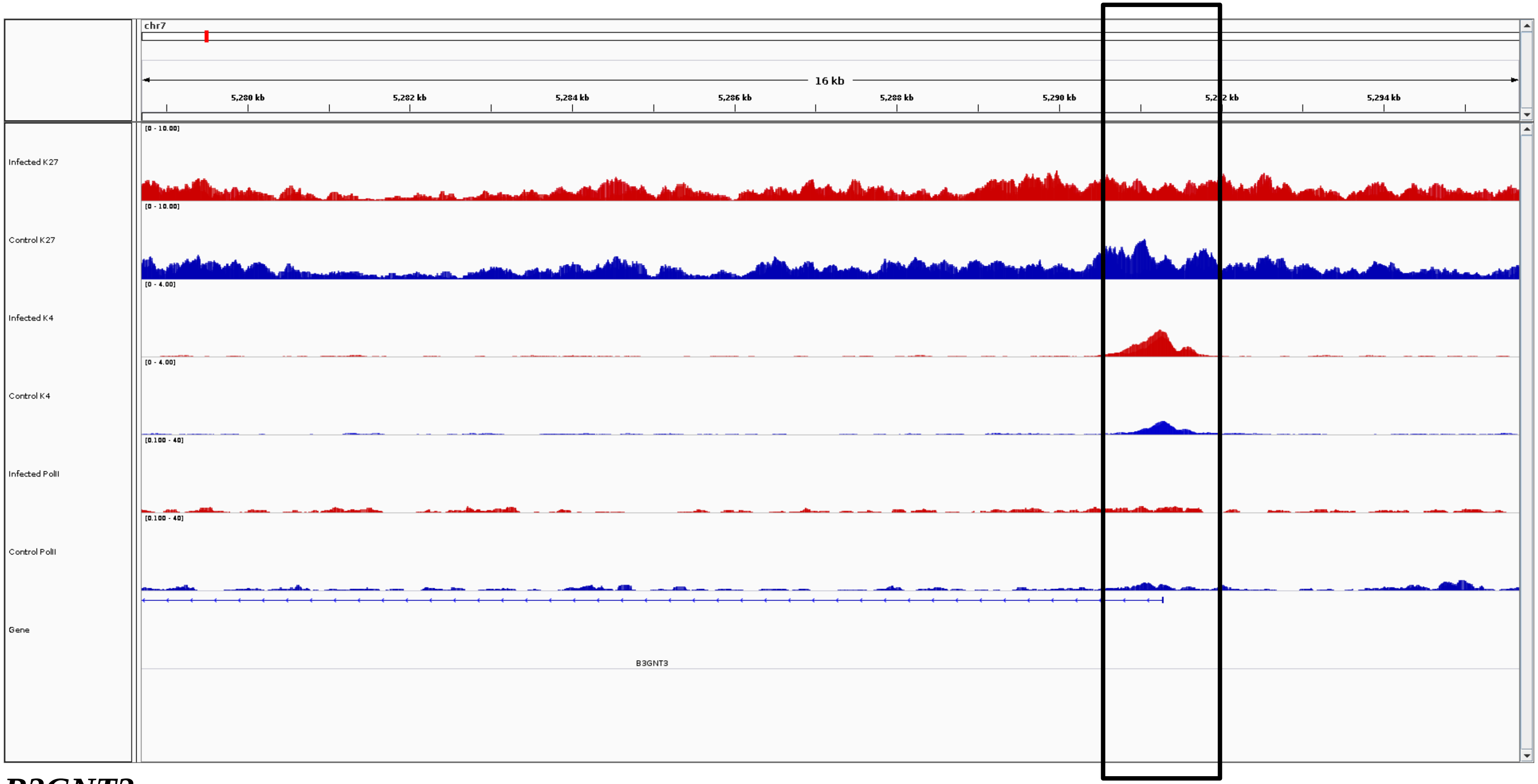
chr7
16 kb
5,280 kb
5,282 kb
5,284 kb
5,286 kb
5,288 kb
5,290 kb
5,294 kb
Infected K27
Control K27
Infected K4
Control K4
Infected PolII
Control PolII
Gene
B3GNT3

***B3GNT3***

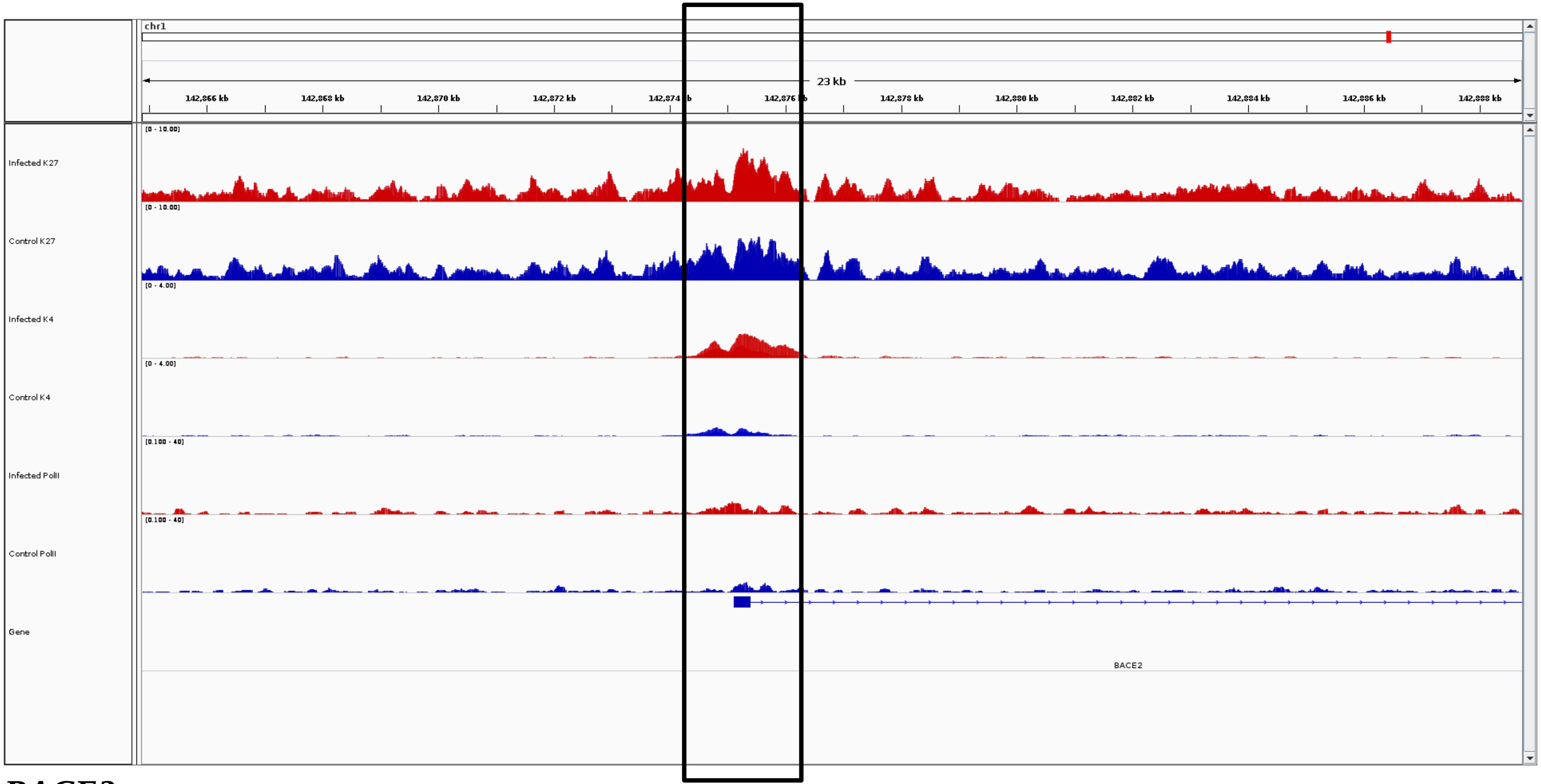
chr1
23 kb
142,866 kb
142,868 kb
142,870 kb
142,872 kb
142,878 kb
142,880 kb
142,882 kb
142,884 kb
142,886 kb
142,888 kb
Infected K27
Control K27
Infected K4
Control K4
Infected PolII
Control PolII
Gene
BACE2

***BACE2***

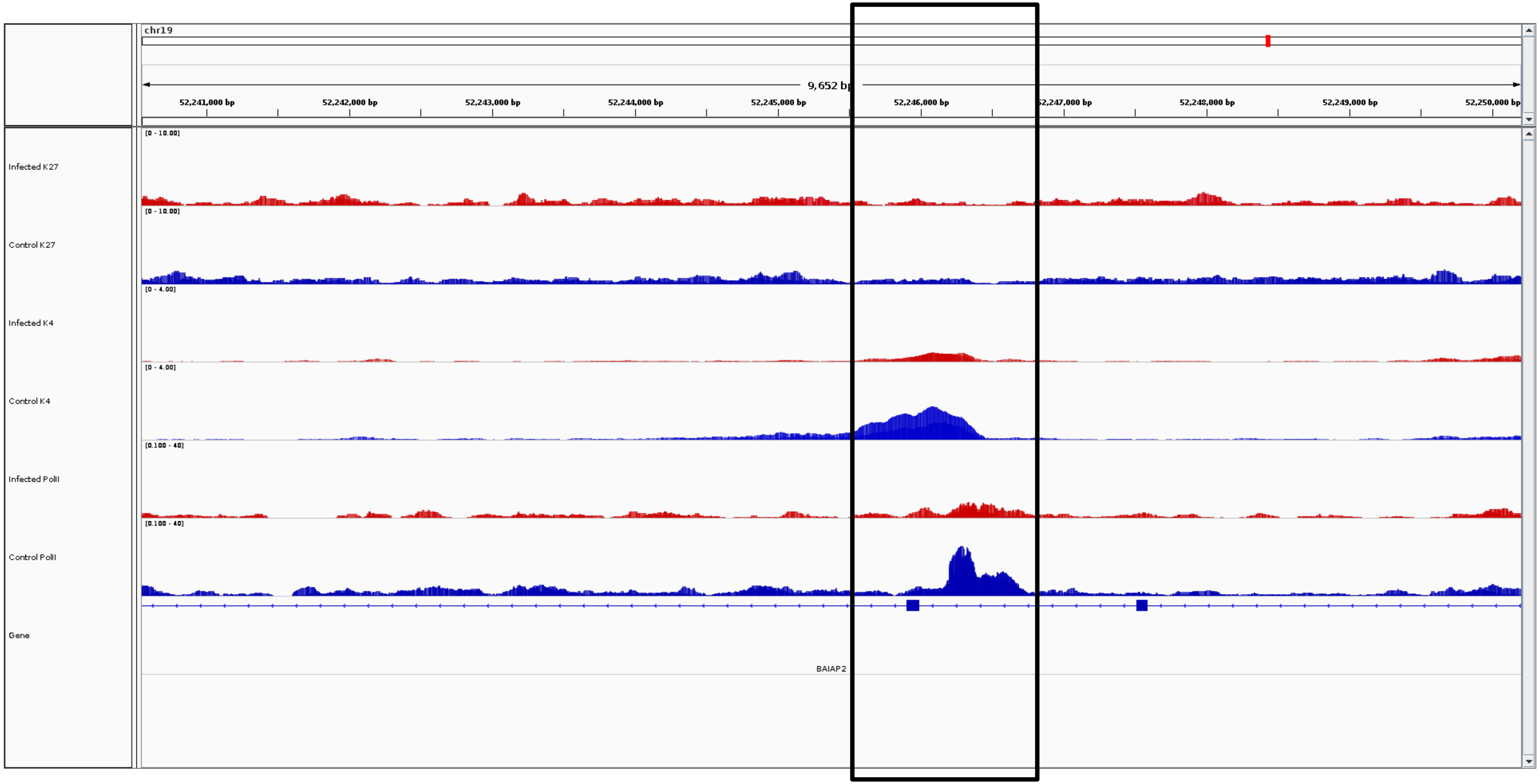
chr19
9,652 bp
52,241,000 bp
52,242,000 bp
52,243,000 bp
52,244,000 bp
52,245,000 bp
52,246,000 bp
52,247,000 bp
52,248,000 bp
52,249,000 bp
52,250,000 bp
Infected K27
[0 - 10.00]
Control K27
[0 - 10.00]
Infected K4
[0 - 4.00]
Control K4
[0 - 4.00]
Infected PolII
[0.100 - 40]
Control PolII
[0.100 - 40]
Gene
BAIAP2

***BAIAP2***

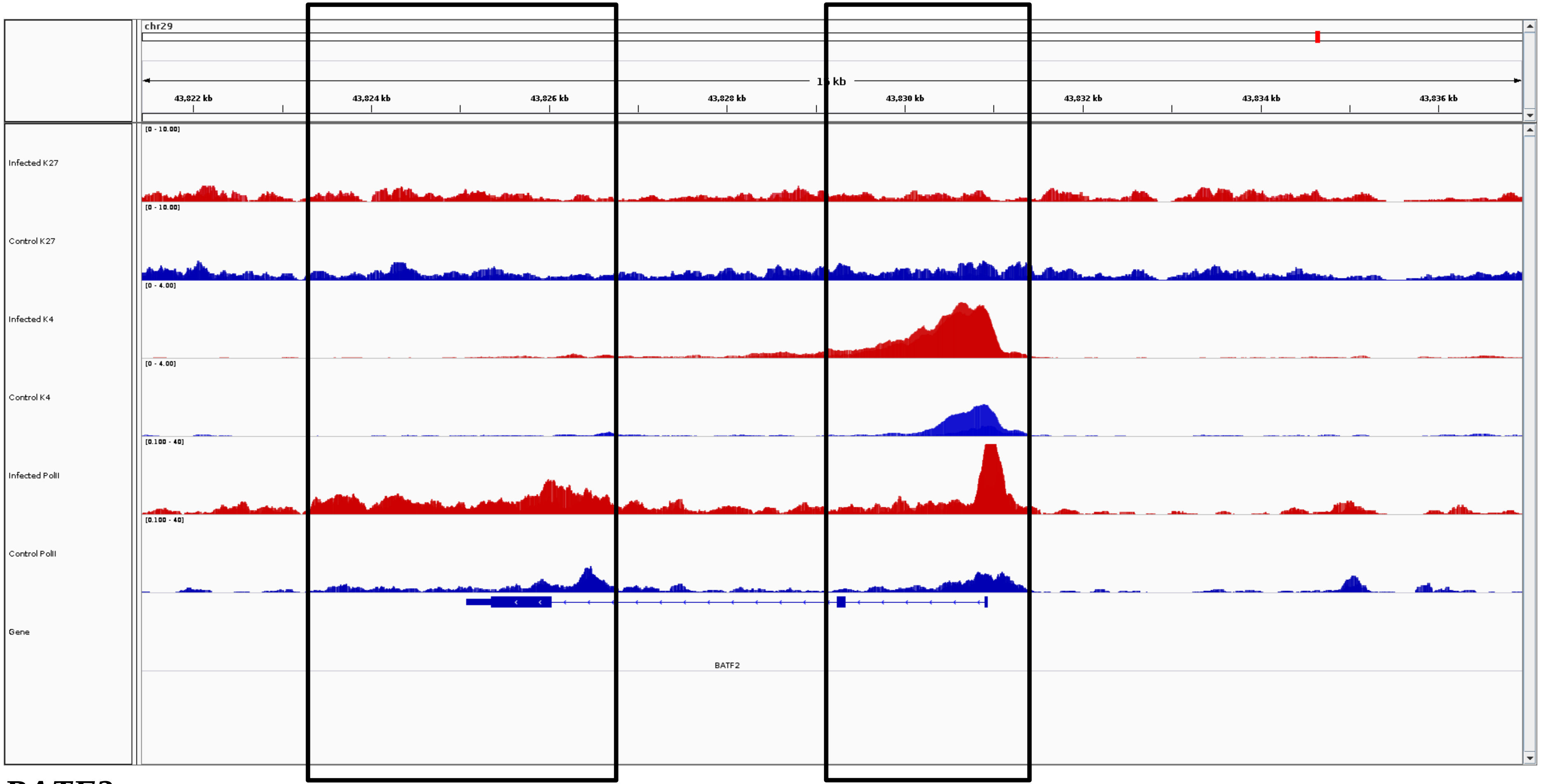
chr29
15 kb
43,822 kb
43,824 kb
43,826 kb
43,828 kb
43,830 kb
43,832 kb
43,834 kb
43,836 kb
Infected K27
Control K27
Infected K4
Control K4
Infected PolII
Control PolII
Gene
BATF2

***BATF2***

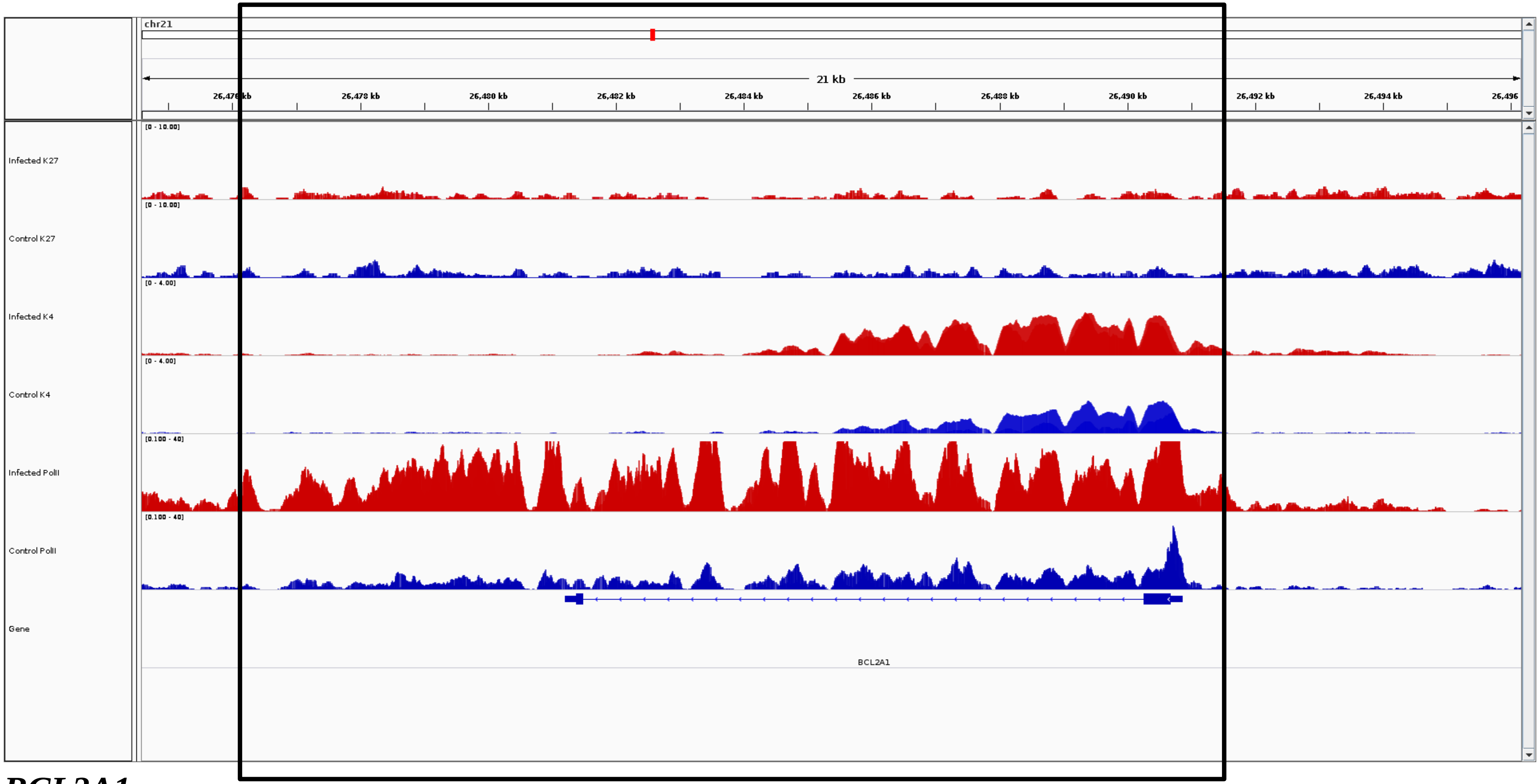
chr21
21 kb
26,476 kb
26,478 kb
26,480 kb
26,482 kb
26,484 kb
26,486 kb
26,488 kb
26,490 kb
26,492 kb
26,494 kb
26,496
Infected K27
Control K27
Infected K4
Control K4
Infected PolII
Control PolII
Gene
BCL2A1

***BCL2A1***

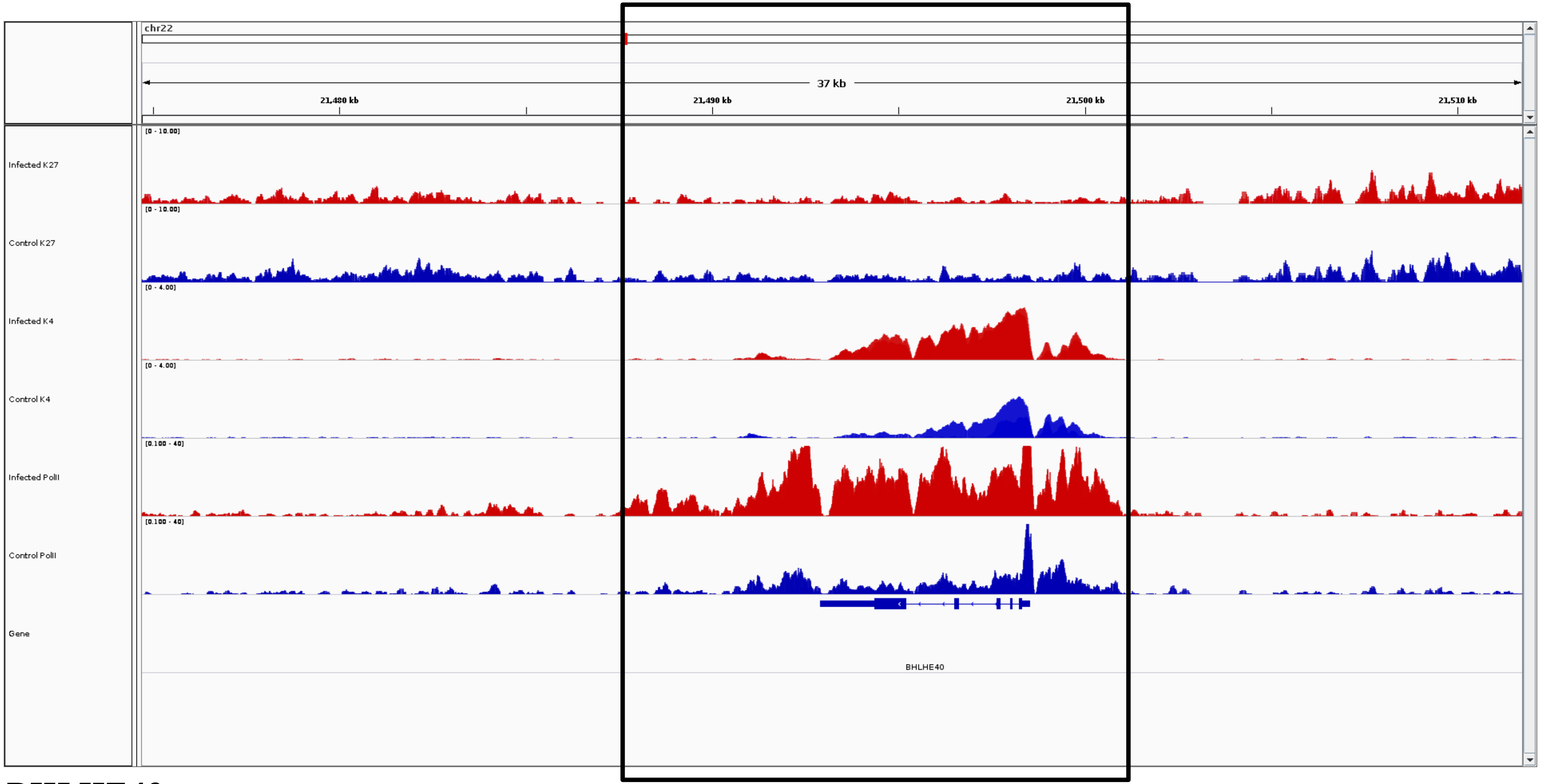
chr22
37 kb
21,480 kb
21,490 kb
21,500 kb
21,510 kb
Infected K27
[0 - 10.00]
Control K27
[0 - 10.00]
Infected K4
[0 - 4.00]
Control K4
[0 - 4.00]
Infected PolII
[0.100 - 40]
Control PolII
[0.100 - 40]
Gene
BHLHE40

***BHLHE40***

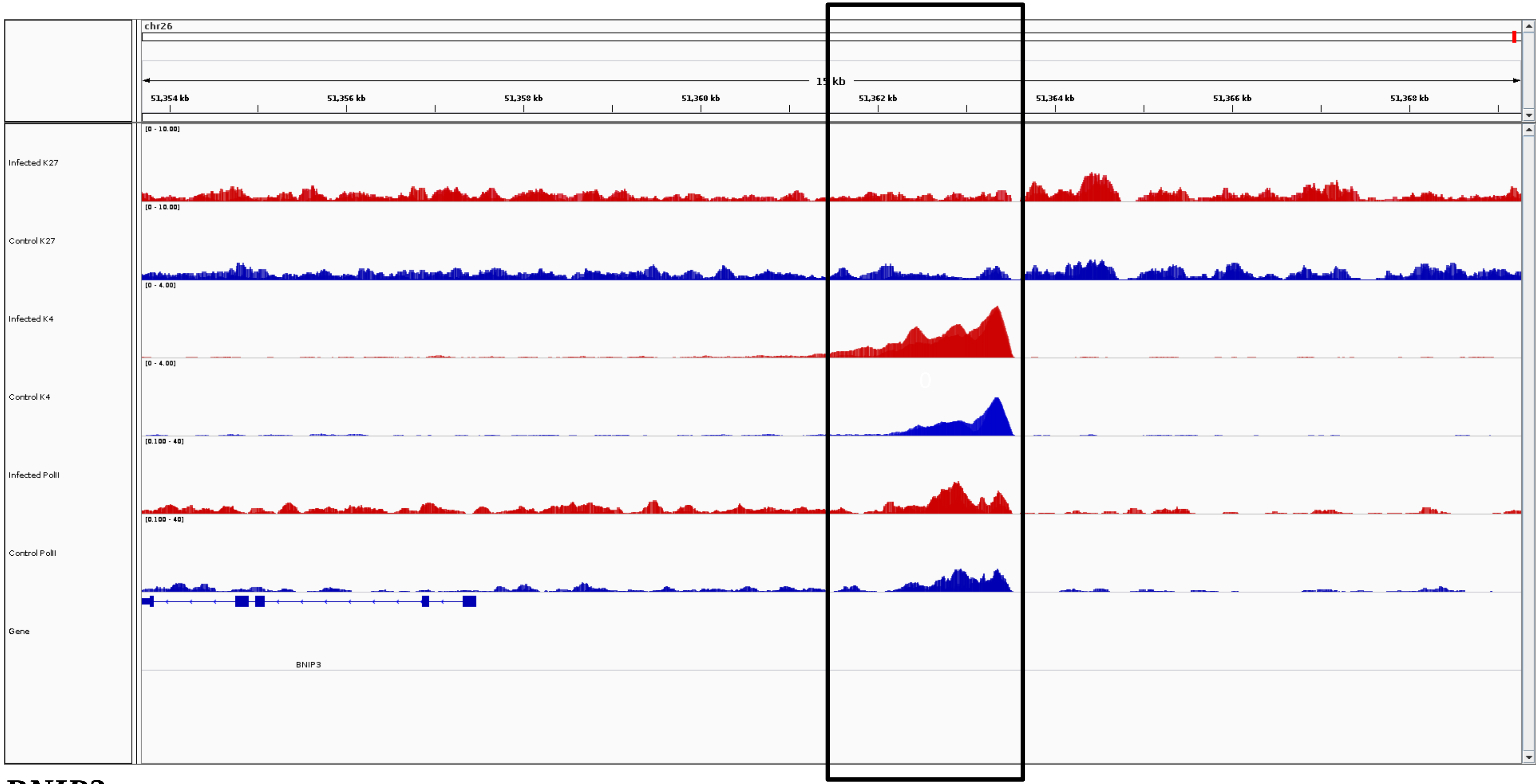
chr26
15 kb
51,354 kb
51,356 kb
51,358 kb
51,360 kb
51,362 kb
51,364 kb
51,366 kb
51,368 kb
Infected K27
[0 - 10.00]
Control K27
[0 - 10.00]
Infected K4
[0 - 4.00]
Control K4
[0 - 4.00]
Infected PolII
[0.100 - 40]
Control PolII
[0.100 - 40]
Gene
BNIP3

***BNIP3***

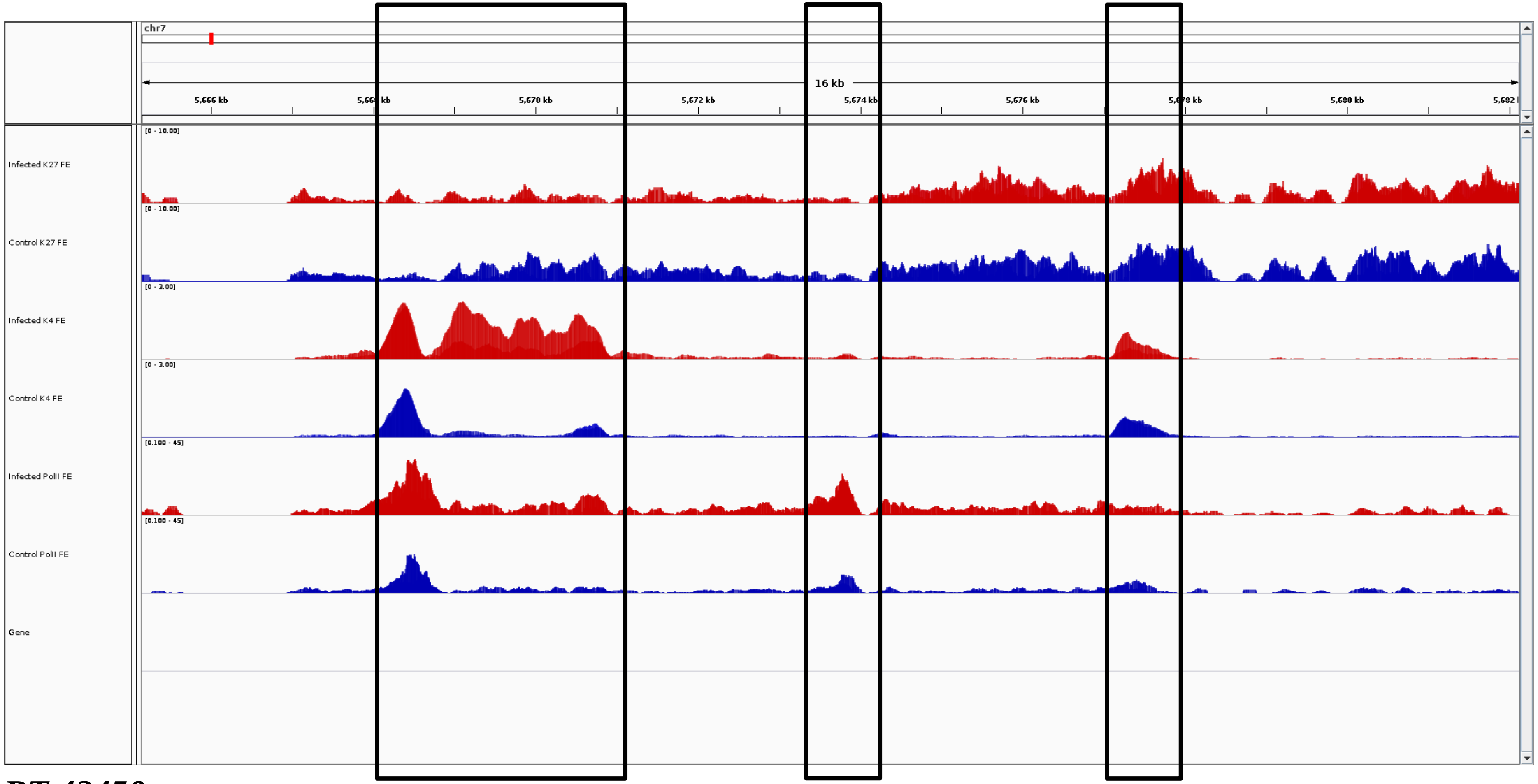
chr7
16 kb
5,666 kb
5,670 kb
5,672 kb
5,674 kb
5,676 kb
5,680 kb
Infected K27 FE
Control K27 FE
Infected K4 FE
Control K4 FE
Infected PolII FE
Control PolII FE
Gene
[0 - 10.00]
[0 - 3.00]
[0.100 - 45]

***BT.42450***

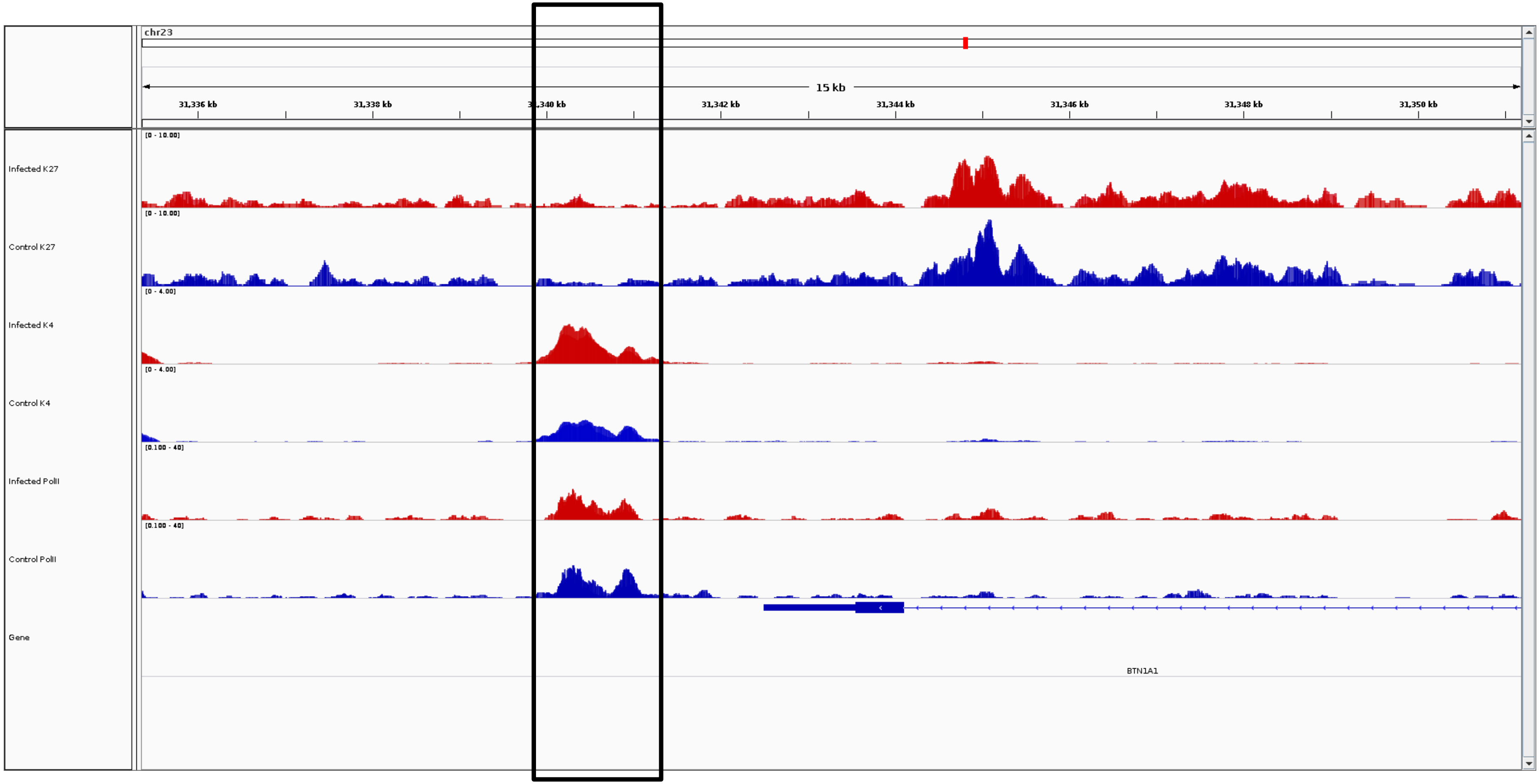
chr23
15 kb
31,336 kb
31,338 kb
31,340 kb
31,342 kb
31,344 kb
31,346 kb
31,348 kb
31,350 kb
Infected K27
Control K27
Infected K4
Control K4
Infected PolII
Control PolII
Gene
BTN1A1

***BTN1A1***

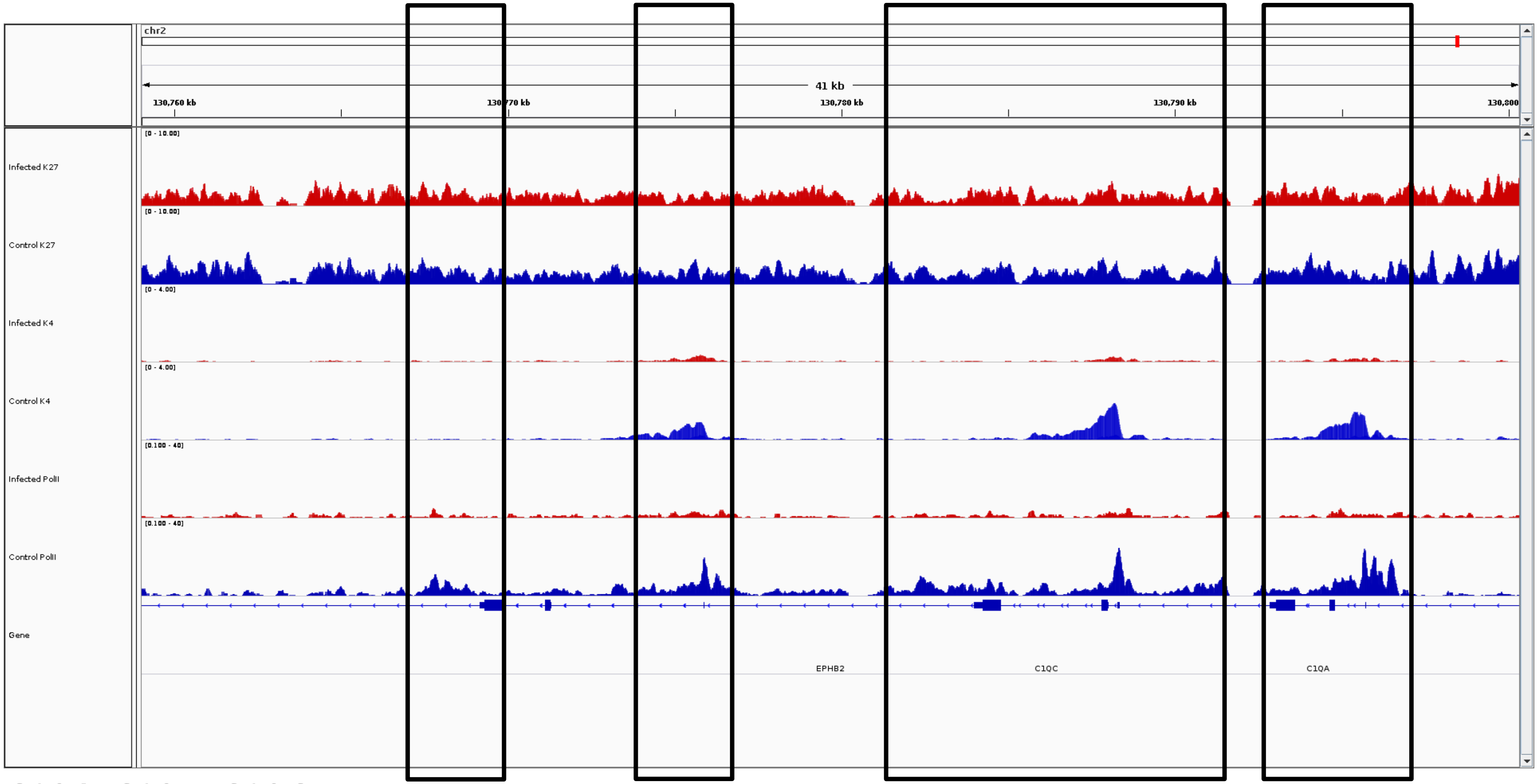


***C1QA, C1QB, C1QC***

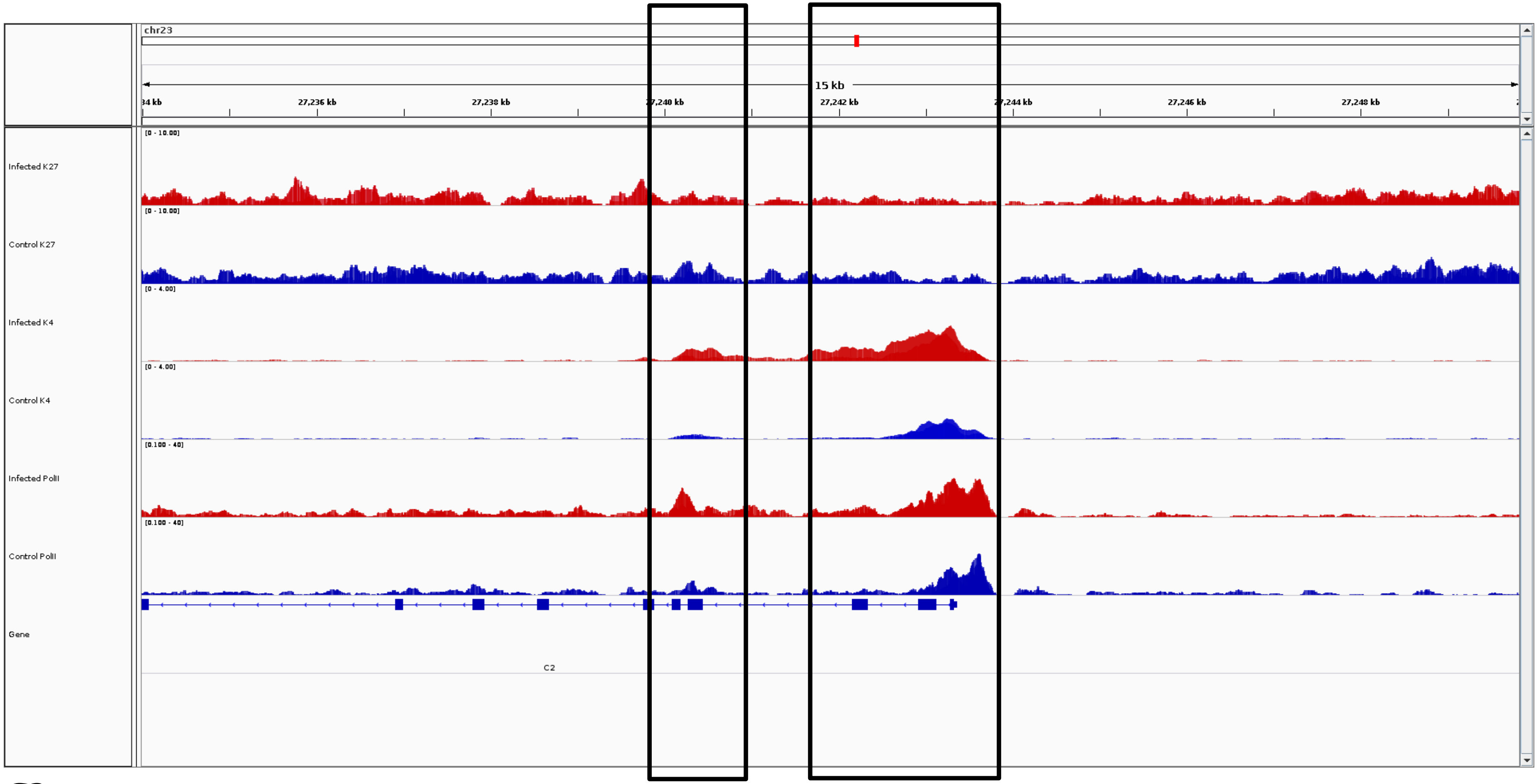
chr23
15 kb
27,236 kb
27,238 kb
27,242 kb
27,246 kb
27,248 kb
Infected K27
Control K27
Infected K4
Control K4
Infected PolII
Control PolII
Gene
C2

*C2*

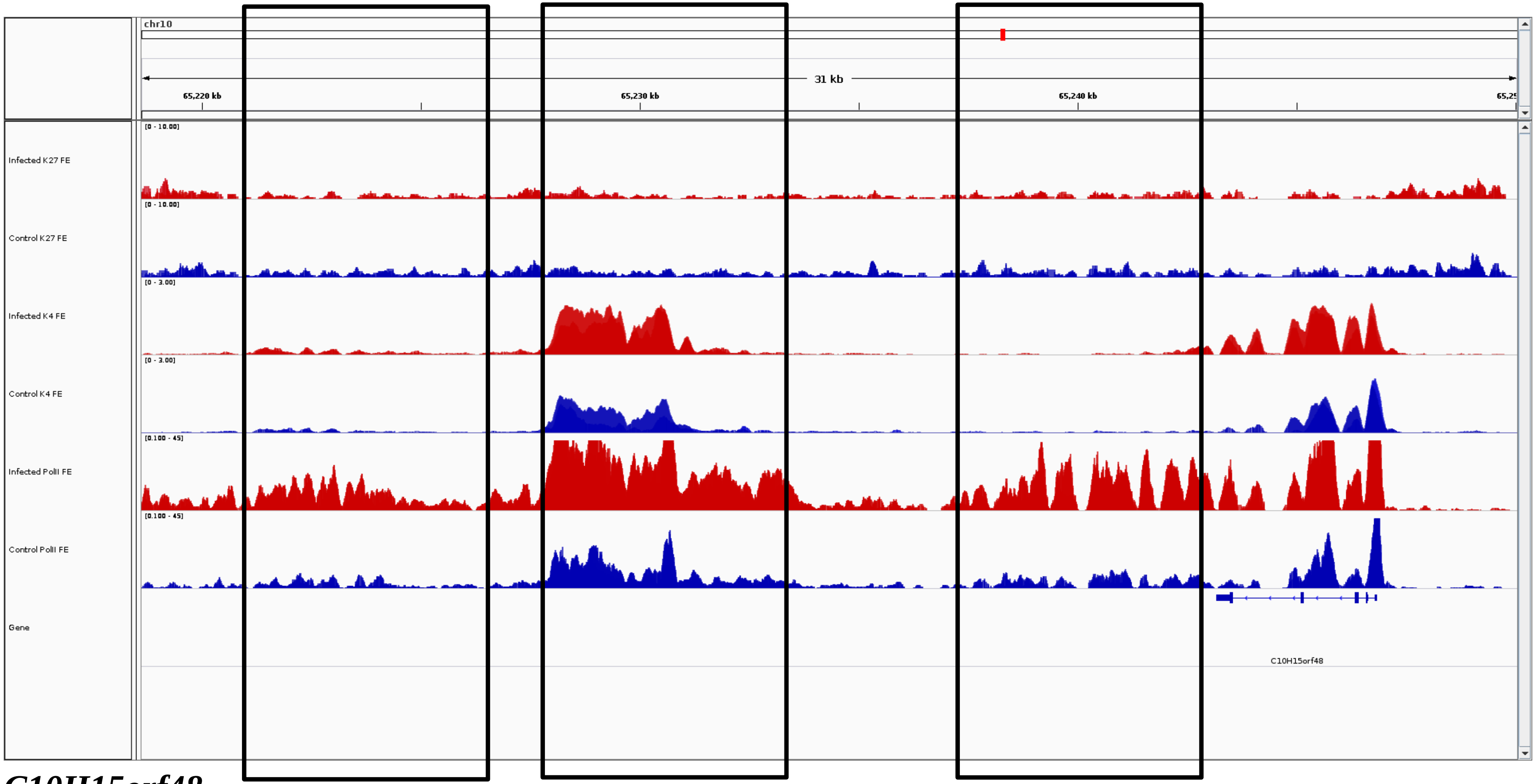


***C10H15orf48***

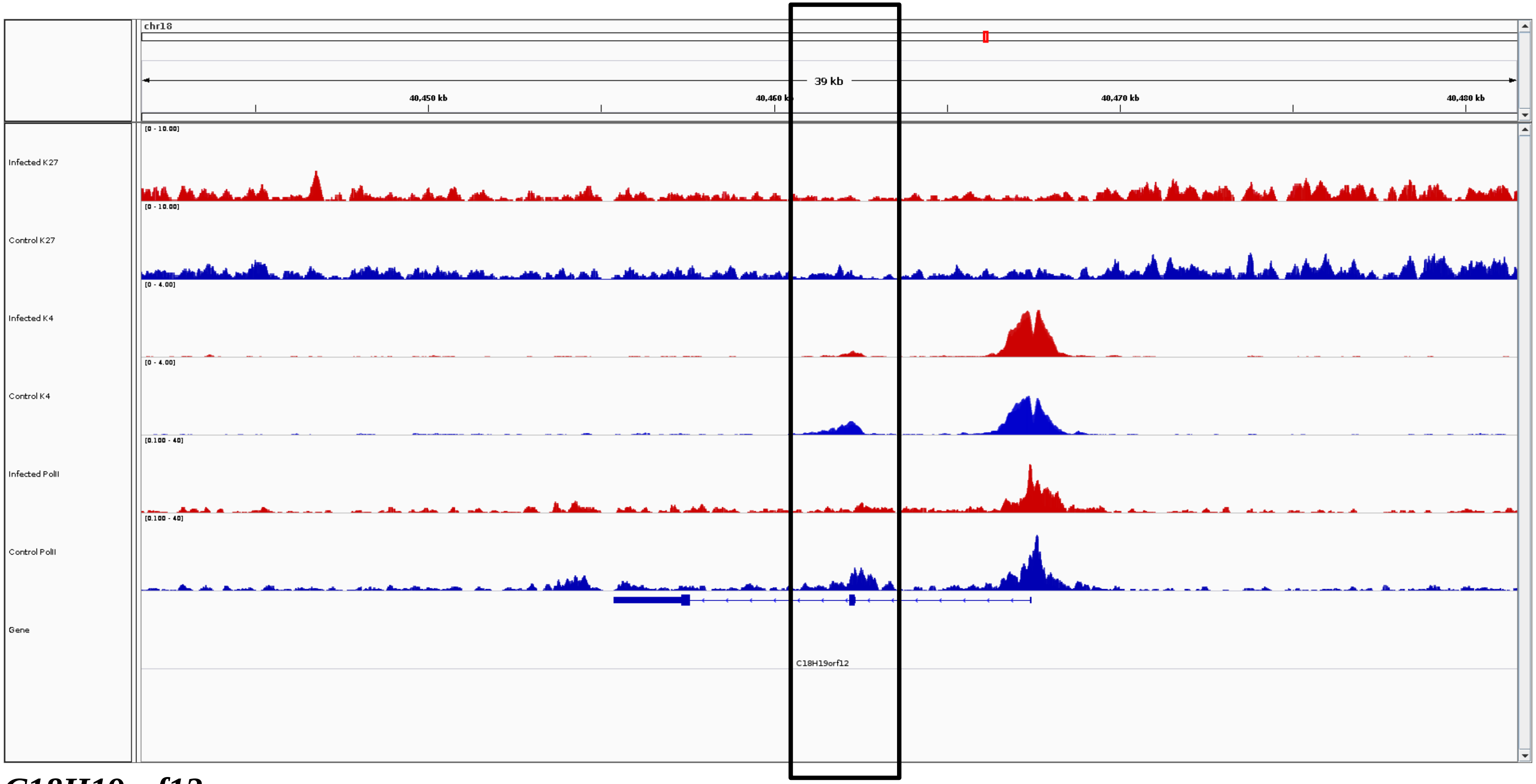


***C18H19orf12***

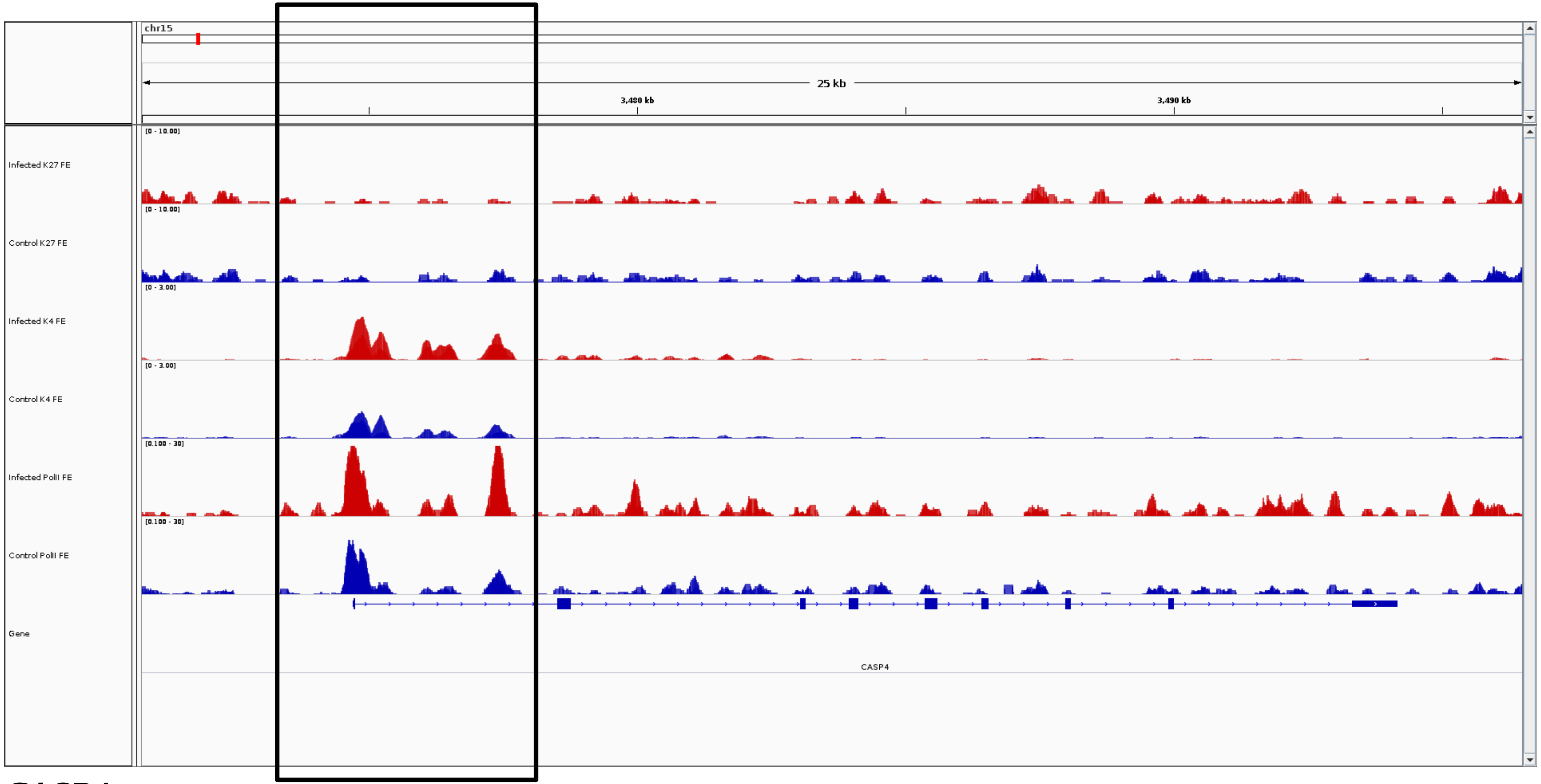
chr15
25 kb
3,480 kb
3,490 kb
Infected K27 FE
[0 - 10.00]
Control K27 FE
[0 - 10.00]
Infected K4 FE
[0 - 3.00]
Control K4 FE
[0 - 3.00]
Infected PolII FE
[0.100 - 30]
Control PolII FE
[0.100 - 30]
Gene
CASP4

***CASP4***

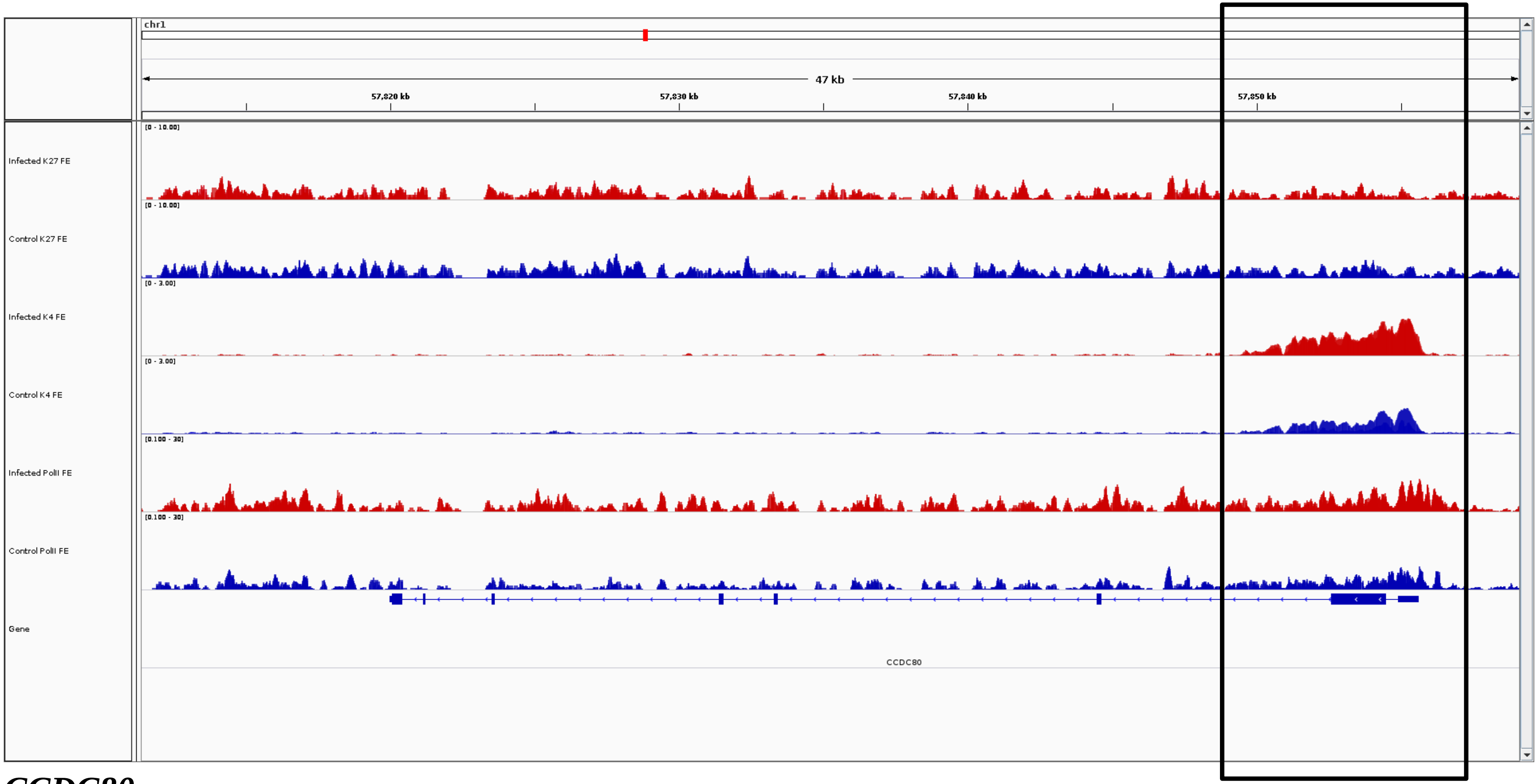
chr1
47 kb
57,820 kb
57,830 kb
57,840 kb
57,850 kb
Infected K27 FE
Control K27 FE
Infected K4 FE
Control K4 FE
Infected PolII FE
Control PolII FE
Gene
CCDC80

***CCDC80***

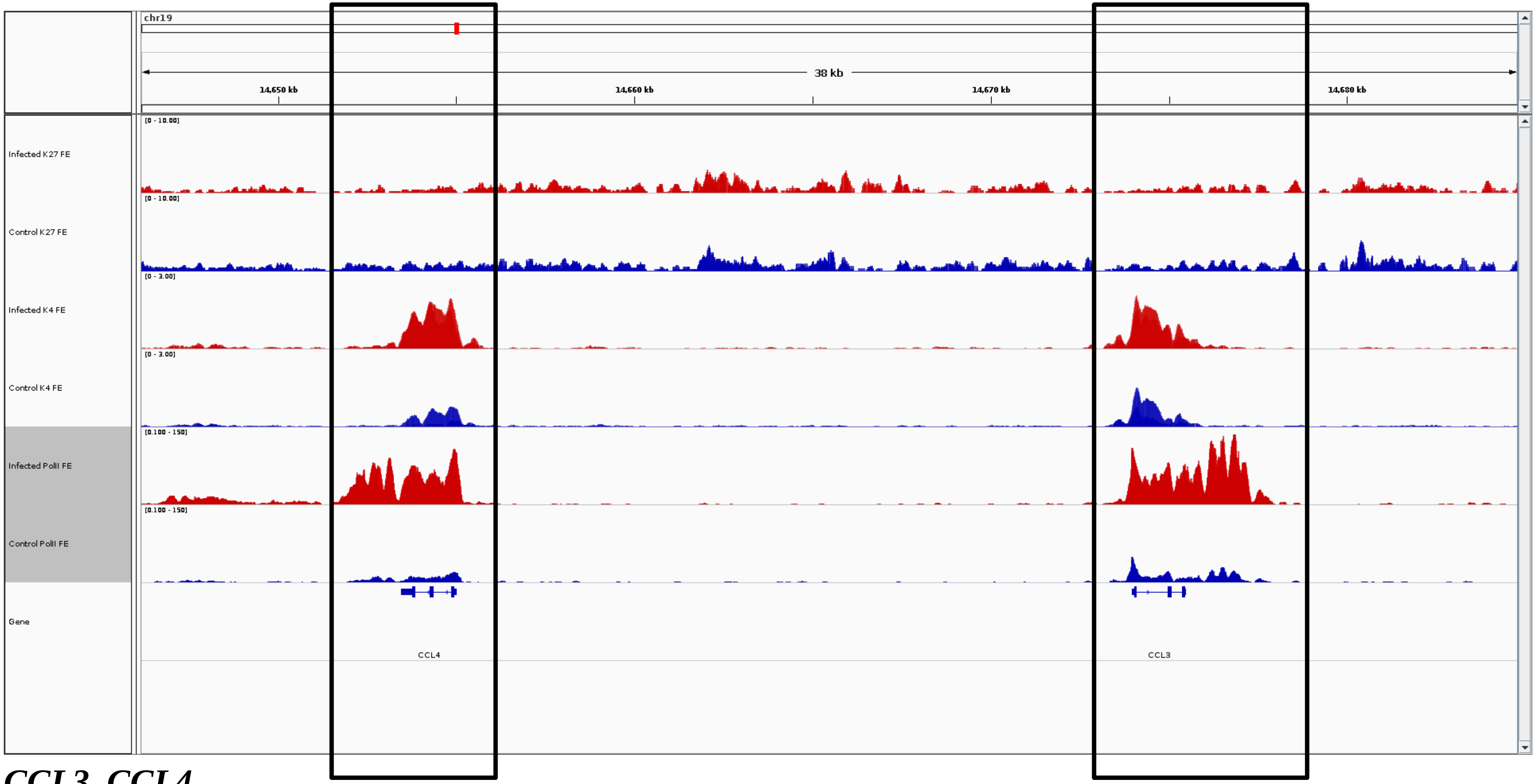
chr19
38 kb
14,650 kb
14,660 kb
14,670 kb
14,680 kb
Infected K27 FE
Control K27 FE
Infected K4 FE
Control K4 FE
Infected PolII FE
Control PolII FE
Gene
CCL4
CCL3

***CCL3, CCL4***

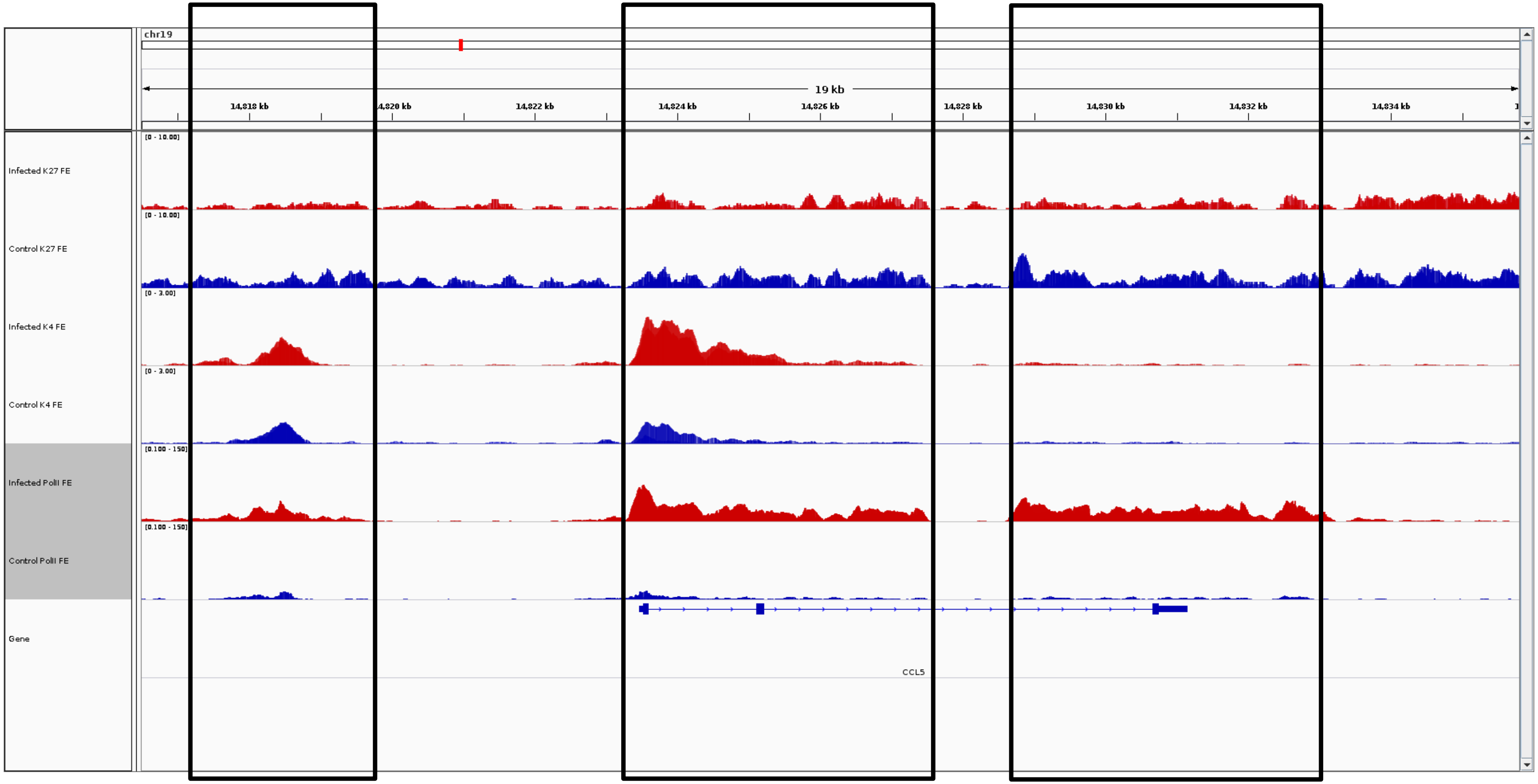
chr19
19 kb
14,818 kb
14,820 kb
14,822 kb
14,824 kb
14,826 kb
14,828 kb
14,830 kb
14,832 kb
14,834 kb
Infected K27 FE
Control K27 FE
Infected K4 FE
Control K4 FE
Infected PolII FE
Control PolII FE
Gene
[0 - 10.00]
[0 - 3.00]
[0.100 - 150]
CCL5

***CCL5***

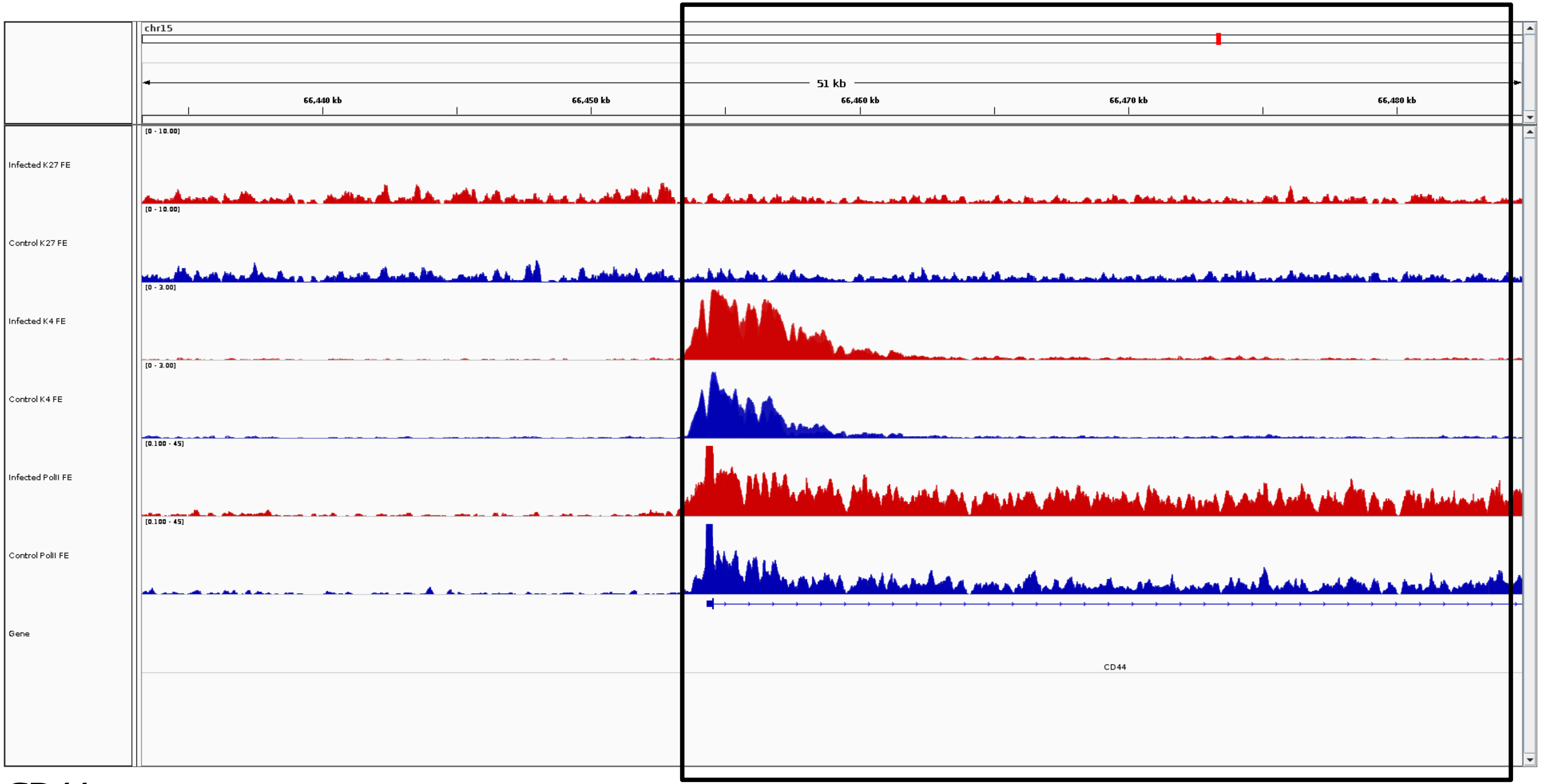
chr15
51 kb
66,440 kb
66,450 kb
66,460 kb
66,470 kb
66,480 kb
Infected K27 FE
Control K27 FE
Infected K4 FE
Control K4 FE
Infected PolII FE
Control PolII FE
Gene
CD44

***CD44***

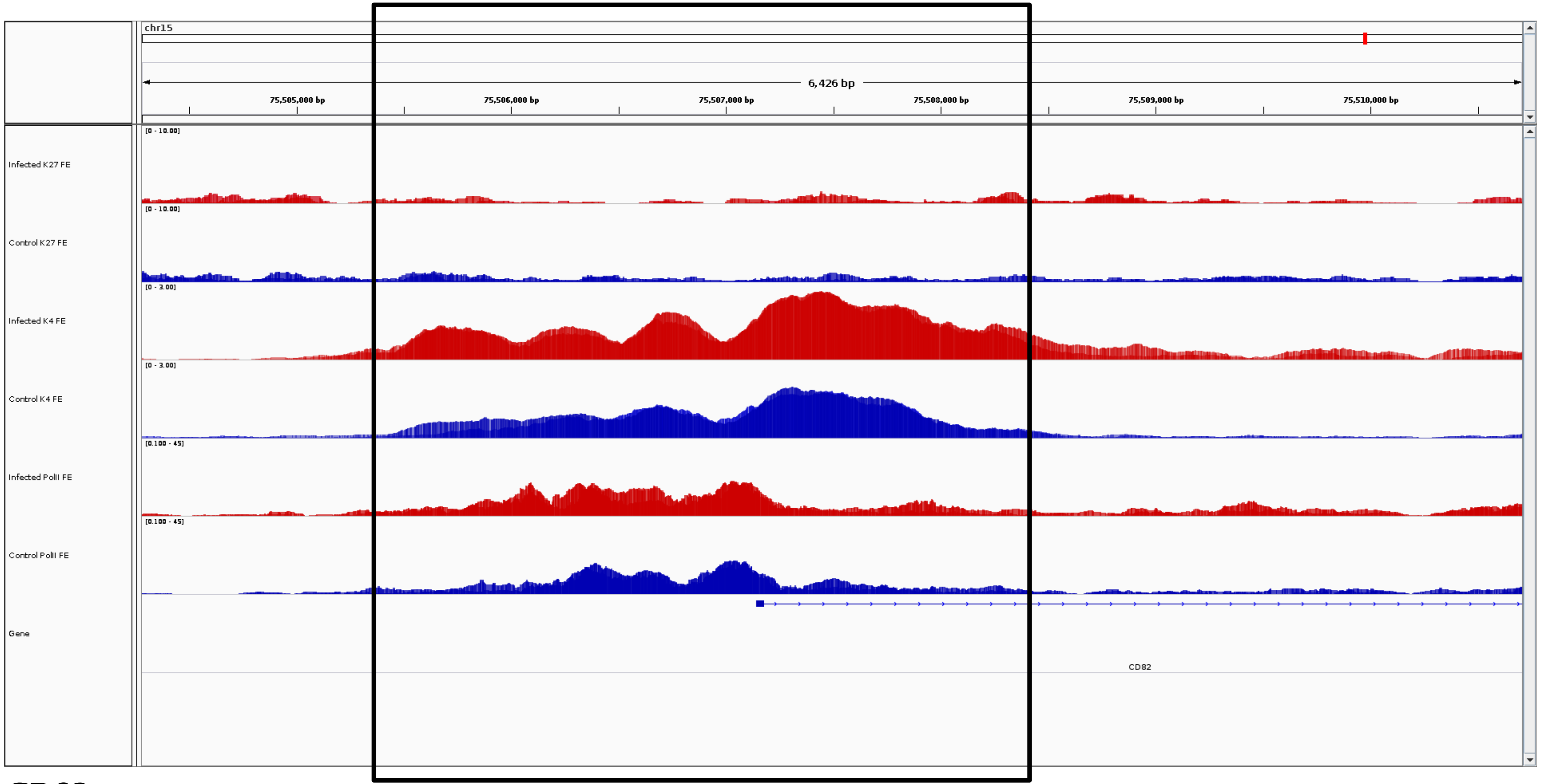
chr15
6,426 bp
75,505,000 bp
75,506,000 bp
75,507,000 bp
75,508,000 bp
75,509,000 bp
75,510,000 bp
Infected K27 FE
[0 - 10.00]
Control K27 FE
[0 - 10.00]
Infected K4 FE
[0 - 3.00]
Control K4 FE
[0 - 3.00]
Infected PolII FE
[0.100 - 45]
Control PolII FE
[0.100 - 45]
Gene
CD82

***CD82***

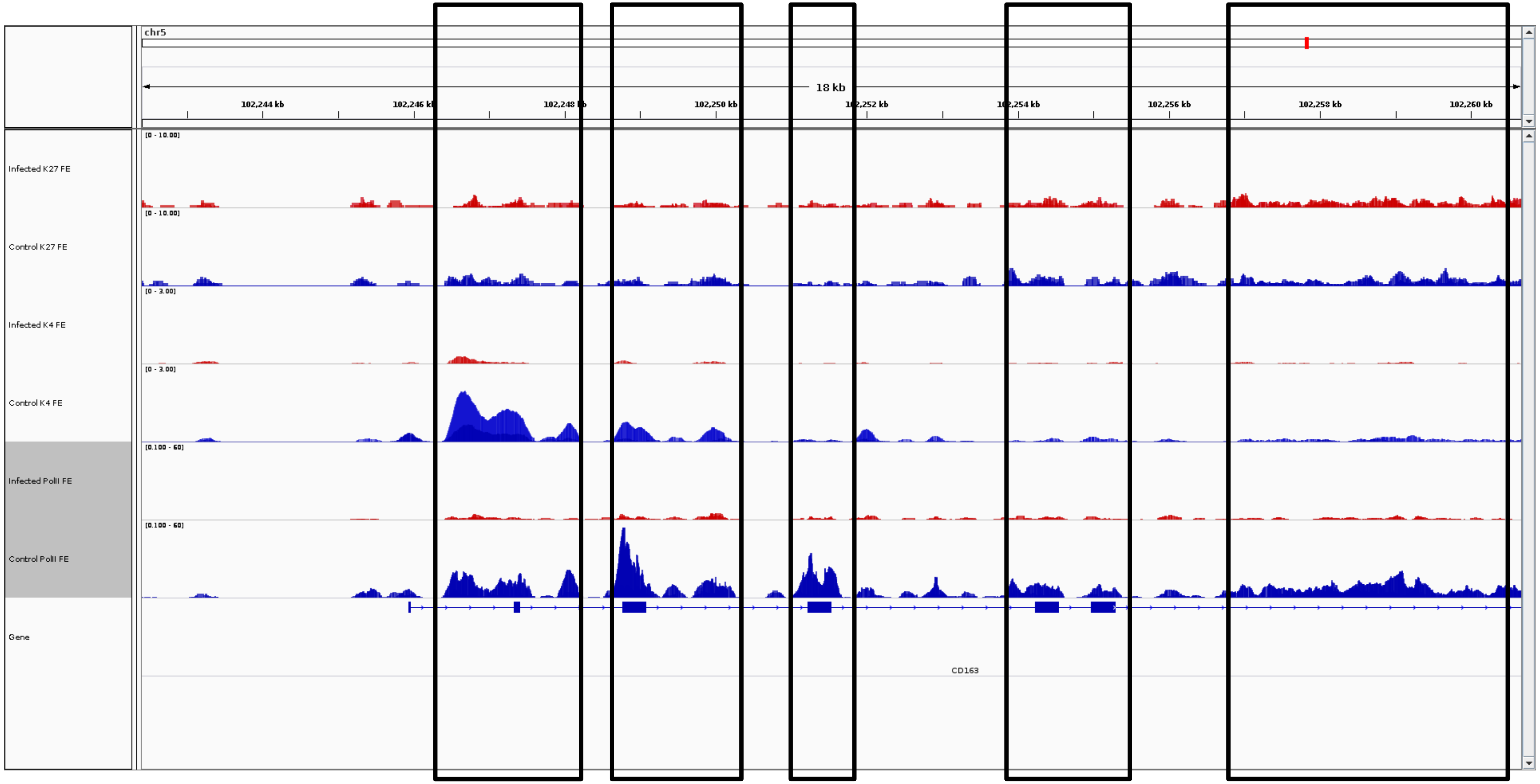
chr5
18 kb
102,244 kb
102,246 kb
102,248 kb
102,250 kb
102,252 kb
102,254 kb
102,256 kb
102,258 kb
102,260 kb
Infected K27 FE
Control K27 FE
Infected K4 FE
Control K4 FE
Infected PolII FE
Control PolII FE
Gene
CD163

***CD163***

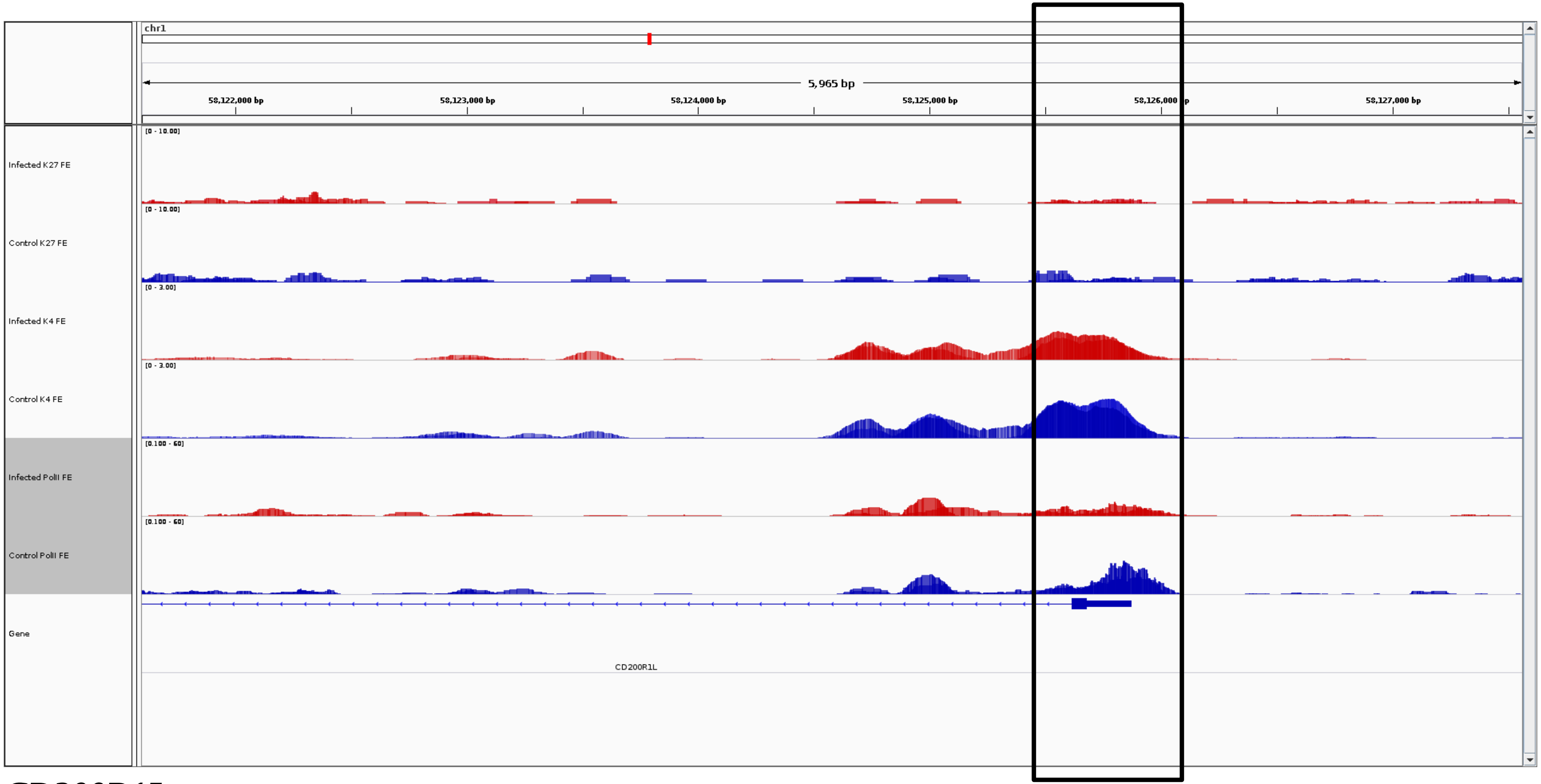
chr1
5,965 bp
58,122,000 bp
58,123,000 bp
58,124,000 bp
58,125,000 bp
58,126,000 bp
58,127,000 bp
Infected K27 FE
[0 - 10.00]
Control K27 FE
[0 - 10.00]
Infected K4 FE
[0 - 3.00]
Control K4 FE
[0 - 3.00]
Infected PolII FE
[0.100 - 60]
Control PolII FE
[0.100 - 60]
Gene
CD200R1L

***CD200R1L***

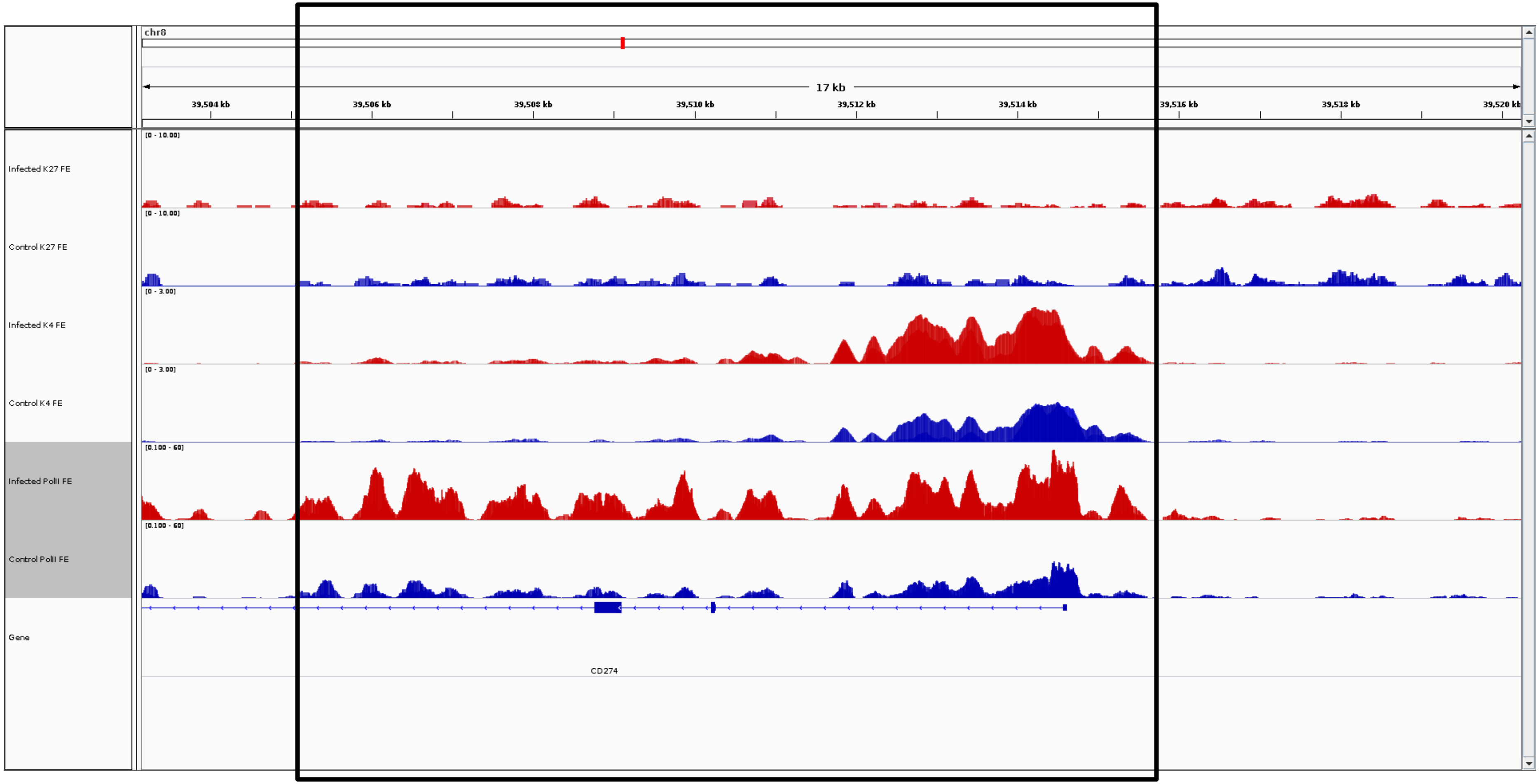
chr8
17 kb
39,504 kb
39,506 kb
39,508 kb
39,510 kb
39,512 kb
39,514 kb
39,516 kb
39,518 kb
39,520 kb
Infected K27 FE
[0 - 10.00]
Control K27 FE
[0 - 10.00]
Infected K4 FE
[0 - 3.00]
Control K4 FE
[0 - 3.00]
Infected PolII FE
[0.100 - 60]
Control PolII FE
[0.100 - 60]
Gene
CD274

***CD274***

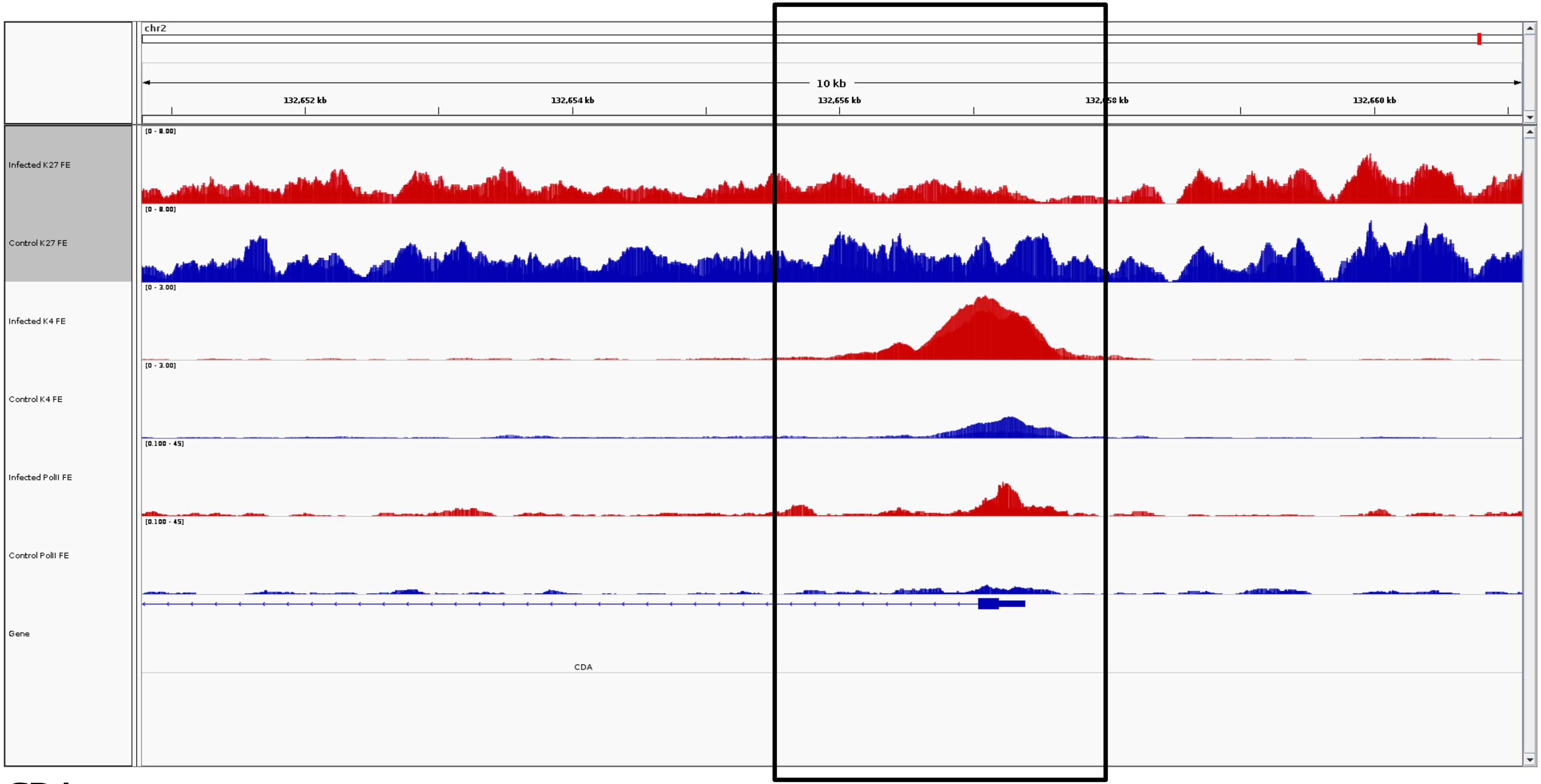
chr2
10 kb
132,652 kb
132,654 kb
132,656 kb
132,660 kb
Infected K27 FE
Control K27 FE
Infected K4 FE
Control K4 FE
Infected PolII FE
Control PolII FE
Gene
CDA

***CDA***

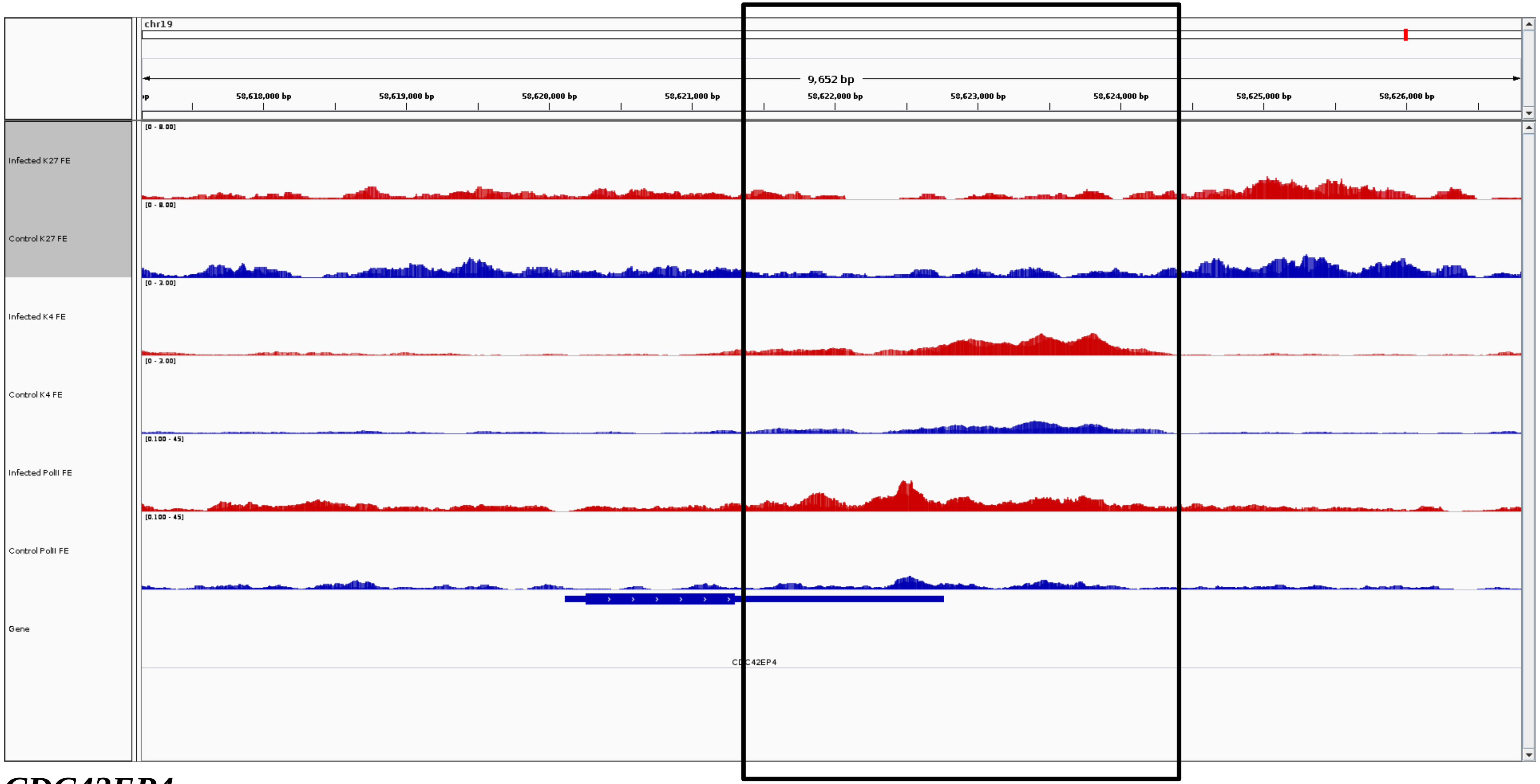
chr19
9,652 bp
58,618,000 bp
58,619,000 bp
58,620,000 bp
58,621,000 bp
58,622,000 bp
58,623,000 bp
58,624,000 bp
58,625,000 bp
58,626,000 bp
Infected K27 FE
[0 - 8.00]
Control K27 FE
[0 - 8.00]
Infected K4 FE
[0 - 3.00]
Control K4 FE
[0 - 3.00]
Infected PolII FE
[0.100 - 45]
Control PolII FE
[0.100 - 45]
Gene
CDC42EP4

***CDC42EP4***

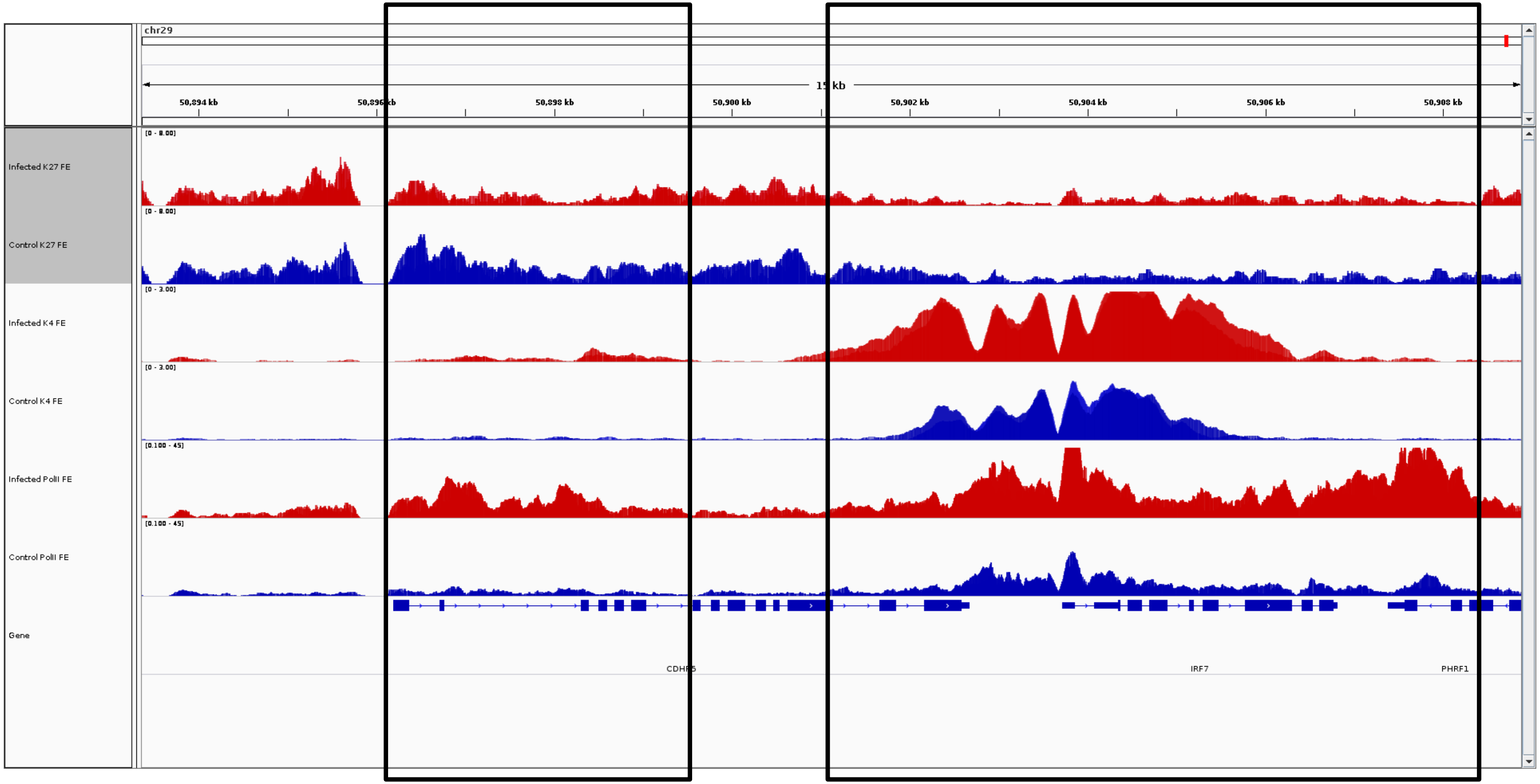
chr29
15 kb
50,894 kb
50,896 kb
50,898 kb
50,900 kb
50,902 kb
50,904 kb
50,906 kb
50,908 kb
Infected K27 FE
Control K27 FE
Infected K4 FE
Control K4 FE
Infected PolII FE
Control PolII FE
Gene
[0 - 8.00]
[0 - 8.00]
[0 - 3.00]
[0 - 3.00]
[0.100 - 45]
[0.100 - 45]
CDHR5
IRF7
PHRF1

***CDHR5***

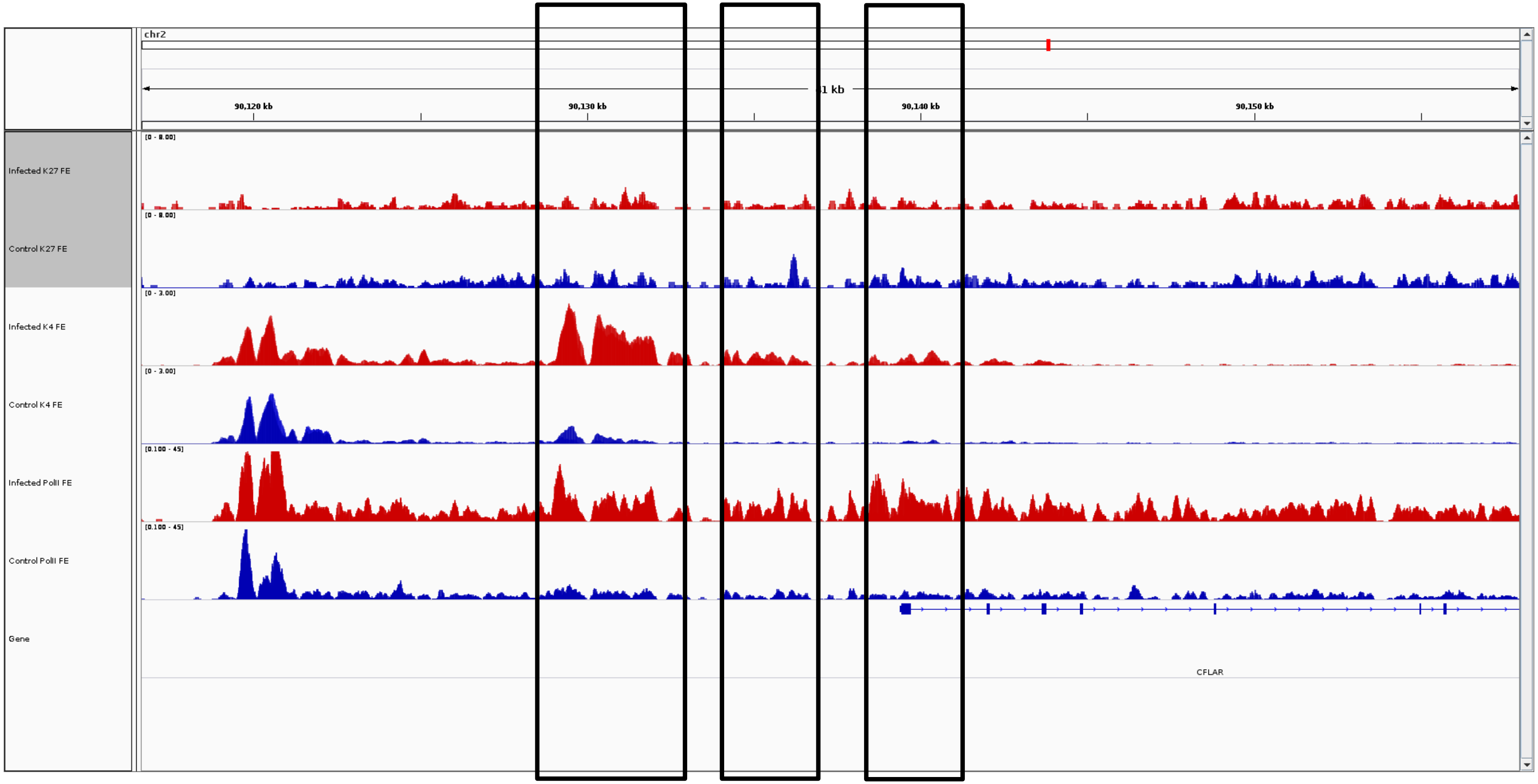
chr2
1 kb
90,120 kb
90,130 kb
90,140 kb
90,150 kb
Infected K27 FE
Control K27 FE
Infected K4 FE
Control K4 FE
Infected PolII FE
Control PolII FE
Gene
CFLAR

***CFLAR***

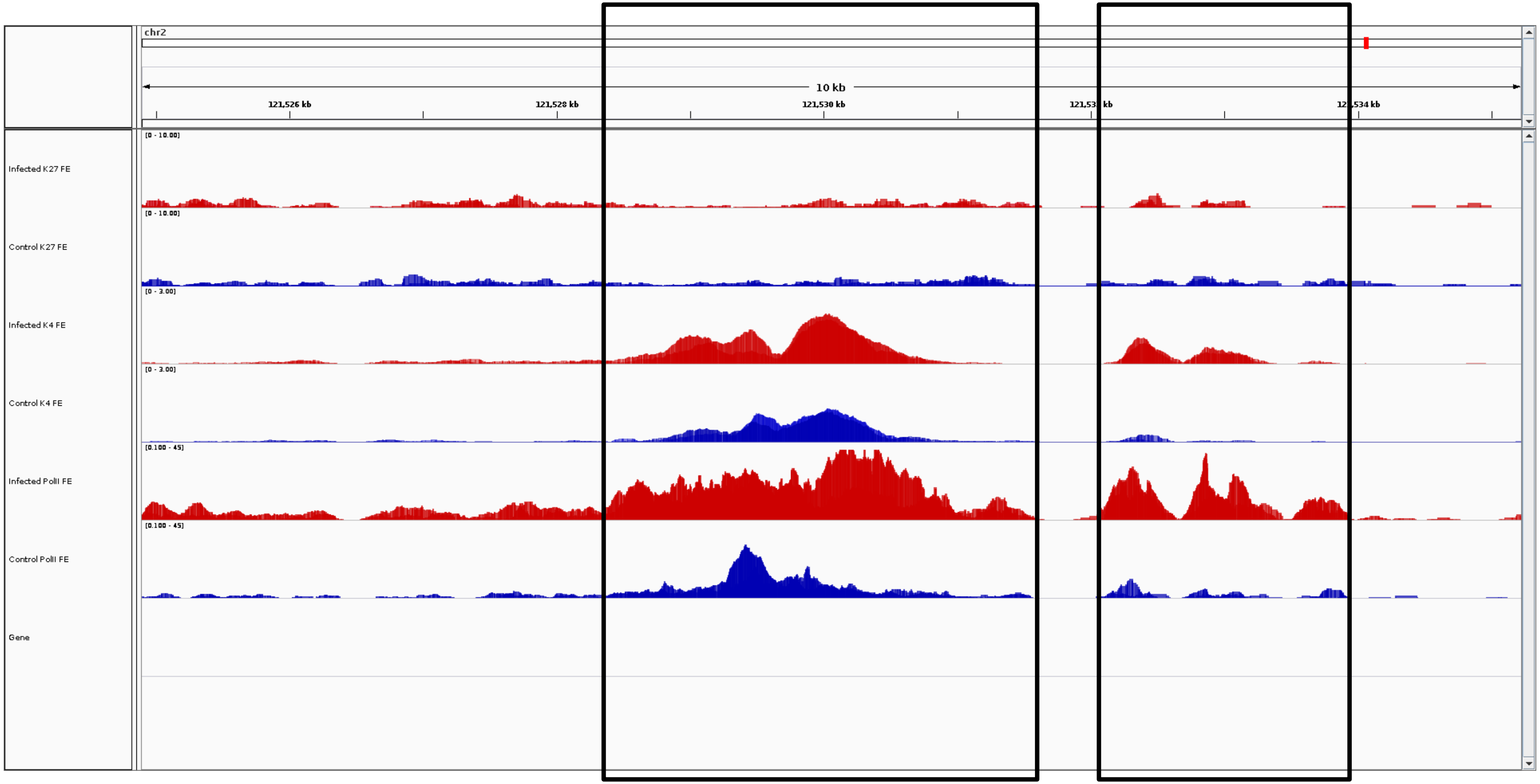


**chr2 121,524,892 – 121,535,228**

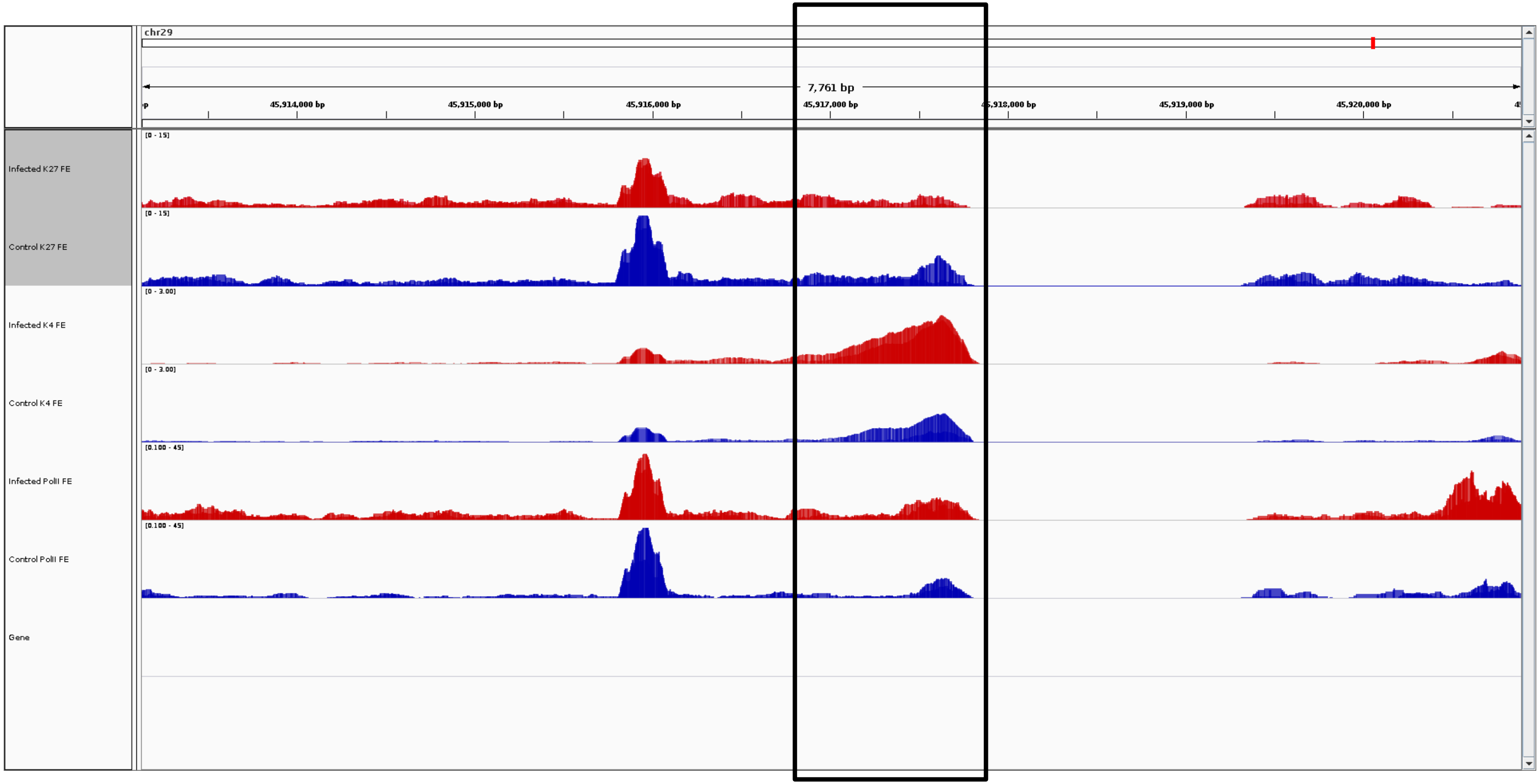
chr29
7,761 bp
45,914,000 bp
45,915,000 bp
45,916,000 bp
45,917,000 bp
45,918,000 bp
45,919,000 bp
45,920,000 bp
Infected K27 FE
Control K27 FE
Infected K4 FE
Control K4 FE
Infected PolII FE
Control PolII FE
Gene
[0 - 15]
[0 - 3.00]
[0.100 - 45]

***CLCF1***

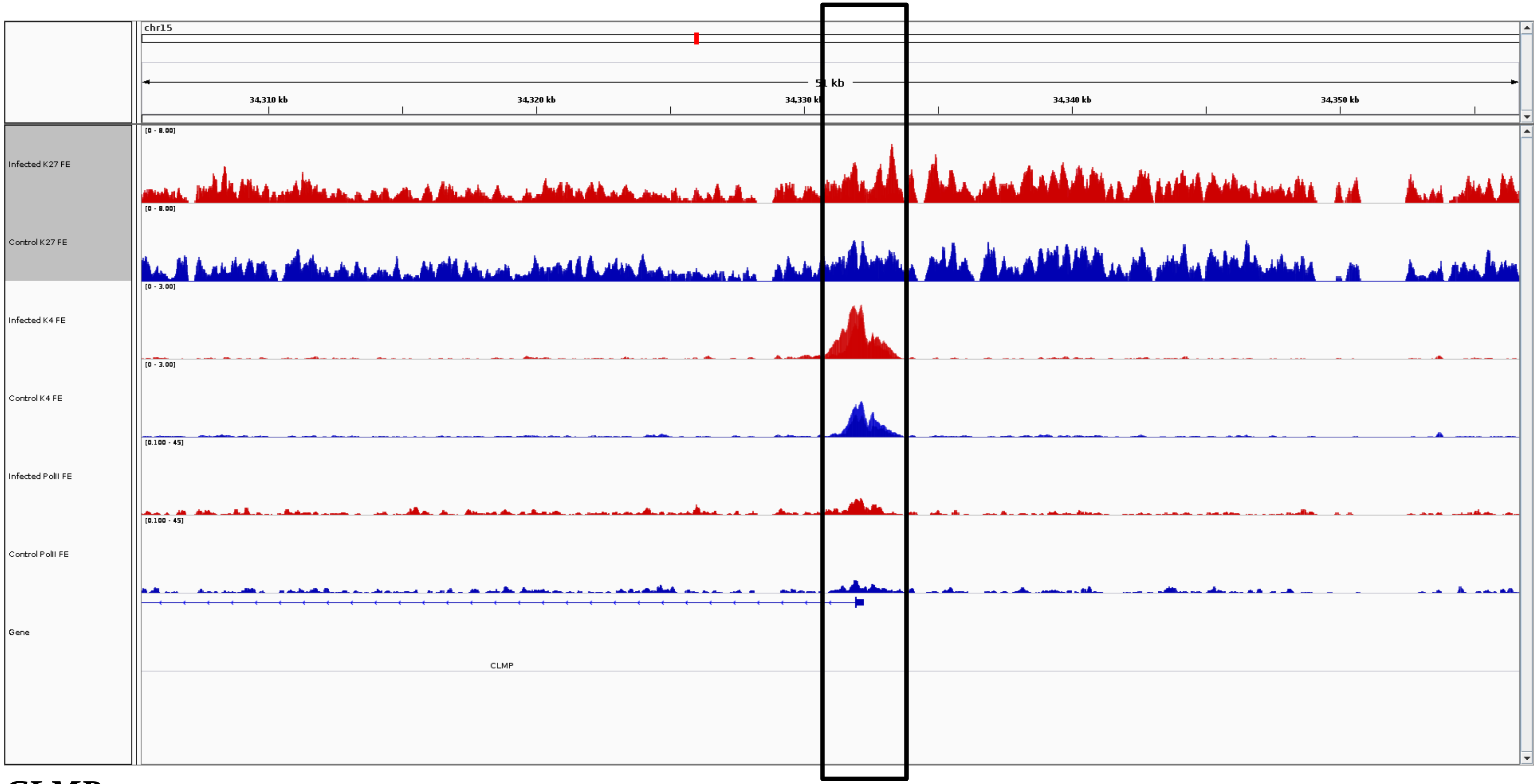
chr15
51 kb
34,310 kb
34,320 kb
34,330 kb
34,340 kb
34,350 kb
Infected K27 FE
Control K27 FE
Infected K4 FE
Control K4 FE
Infected PolII FE
Control PolII FE
Gene
CLMP

***CLMP***

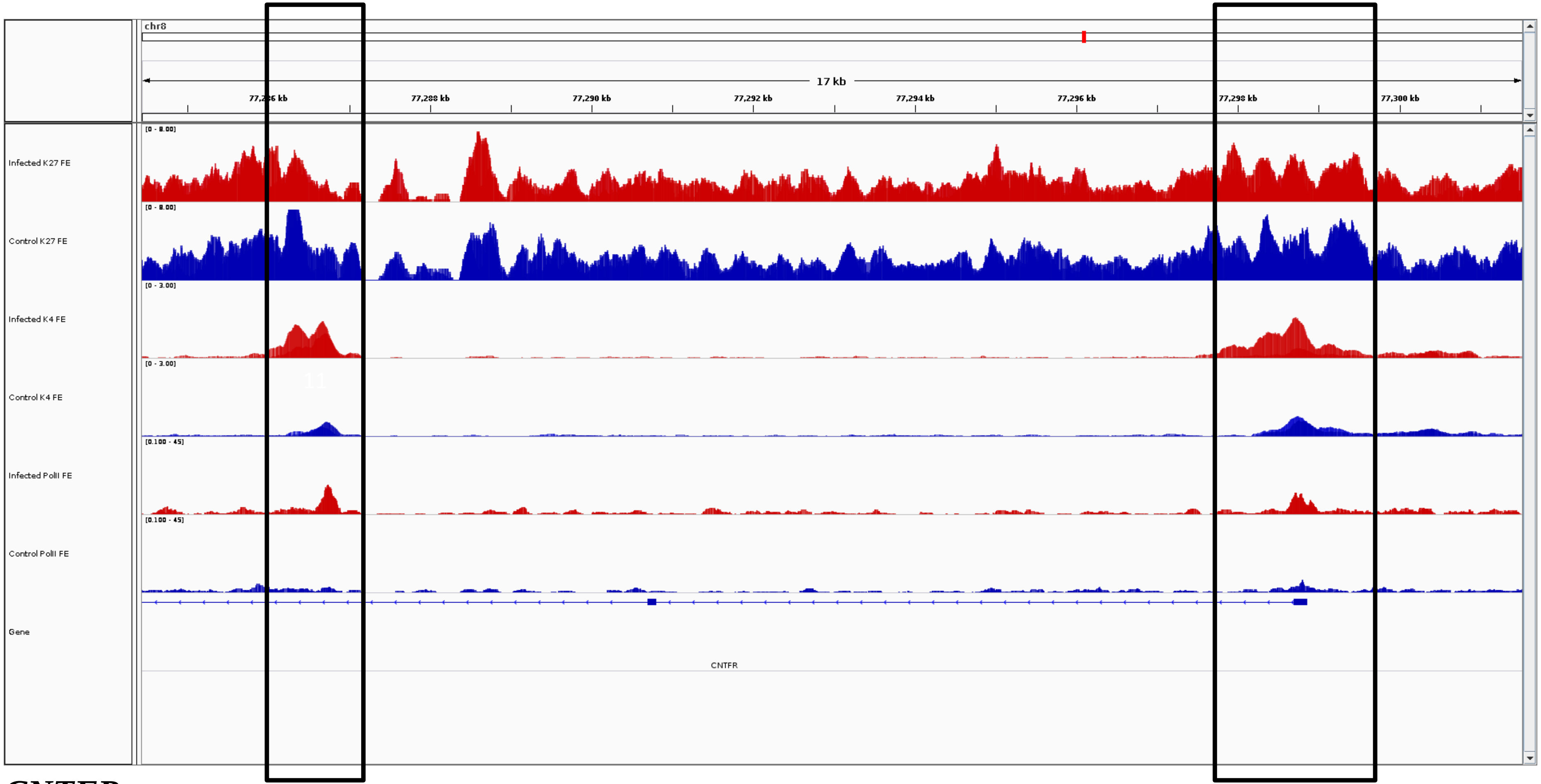
chr8
17 kb
77,286 kb
77,288 kb
77,290 kb
77,292 kb
77,294 kb
77,296 kb
77,298 kb
77,300 kb
[0 - 8.00]
Infected K27 FE
[0 - 8.00]
Control K27 FE
[0 - 3.00]
Infected K4 FE
[0 - 3.00]
Control K4 FE
[0.100 - 45]
Infected PolII FE
[0.100 - 45]
Control PolII FE
Gene
CNTFR

***CNTFR***

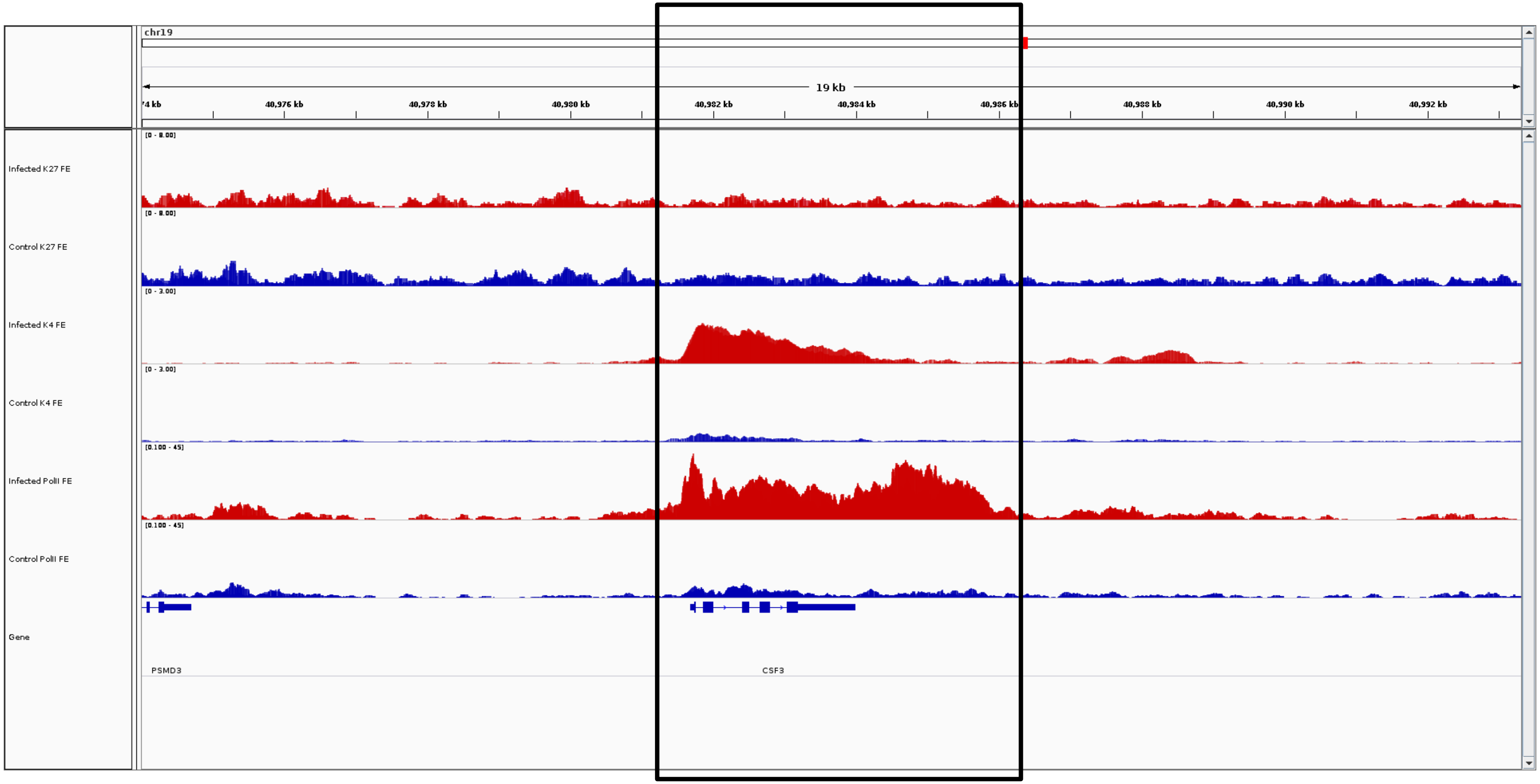
chr19
19 kb
40,976 kb
40,978 kb
40,980 kb
40,982 kb
40,984 kb
40,986 kb
40,988 kb
40,990 kb
40,992 kb
Infected K27 FE
Control K27 FE
Infected K4 FE
Control K4 FE
Infected PolII FE
Control PolII FE
Gene
PSMD3
CSF3

***CSF3***

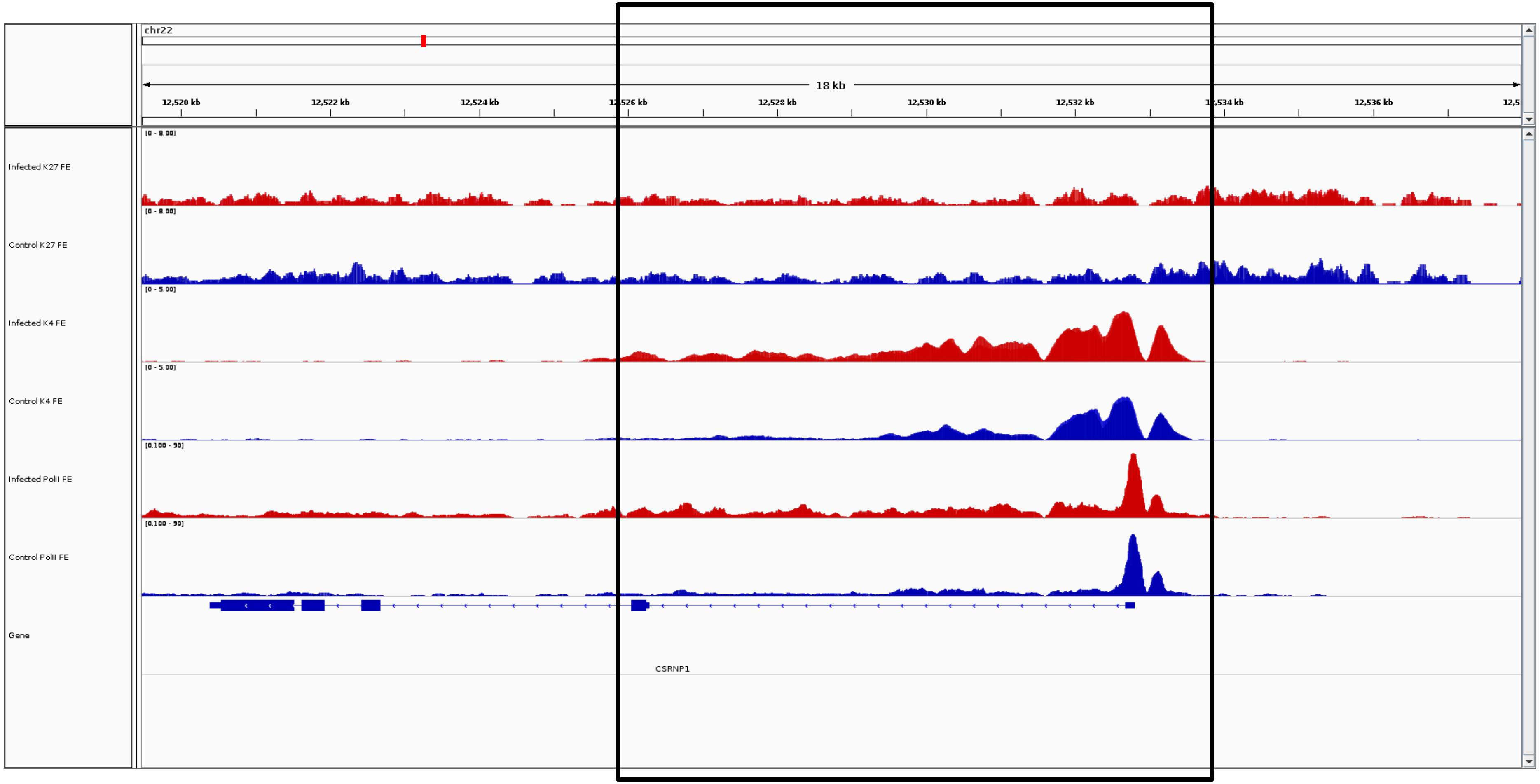
chr22
18 kb
12,520 kb
12,522 kb
12,524 kb
12,526 kb
12,528 kb
12,530 kb
12,532 kb
12,534 kb
12,536 kb
Infected K27 FE
Control K27 FE
Infected K4 FE
Control K4 FE
Infected PolII FE
Control PolII FE
Gene
CSRNP1

***CSRNP1***

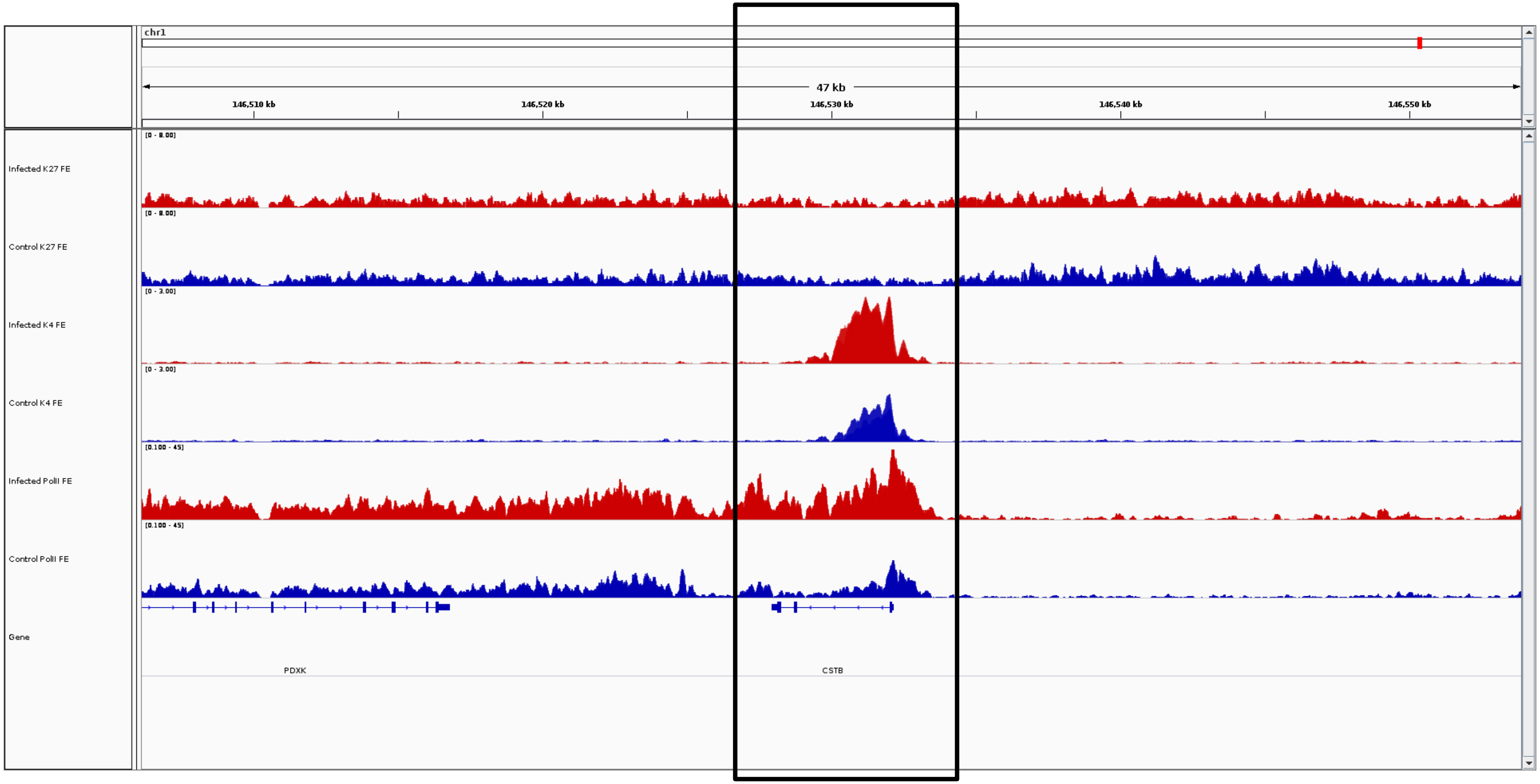
chr1
47 kb
146,510 kb
146,520 kb
146,530 kb
146,540 kb
146,550 kb
Infected K27 FE
[0 - 8.00]
Control K27 FE
[0 - 8.00]
Infected K4 FE
[0 - 3.00]
Control K4 FE
[0 - 3.00]
Infected PolII FE
[0.100 - 45]
Control PolII FE
[0.100 - 45]
Gene
PDXK
CSTB

***CSTB***

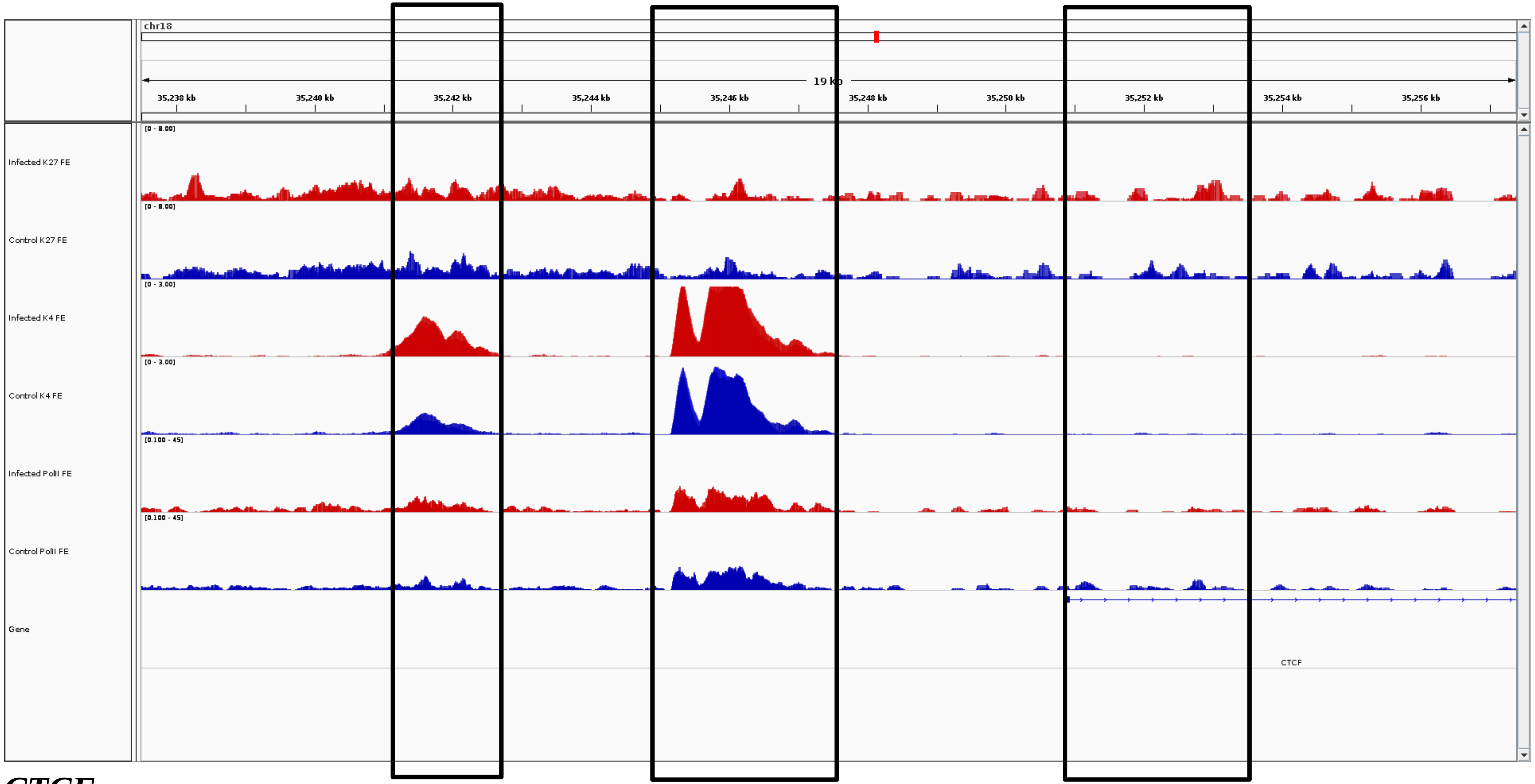
chr18
19 kb
35,238 kb
35,240 kb
35,242 kb
35,244 kb
35,246 kb
35,248 kb
35,250 kb
35,252 kb
35,254 kb
35,256 kb
Infected K27 FE
Control K27 FE
Infected K4 FE
Control K4 FE
Infected PolII FE
Control PolII FE
Gene
CTCF

***CTCF***

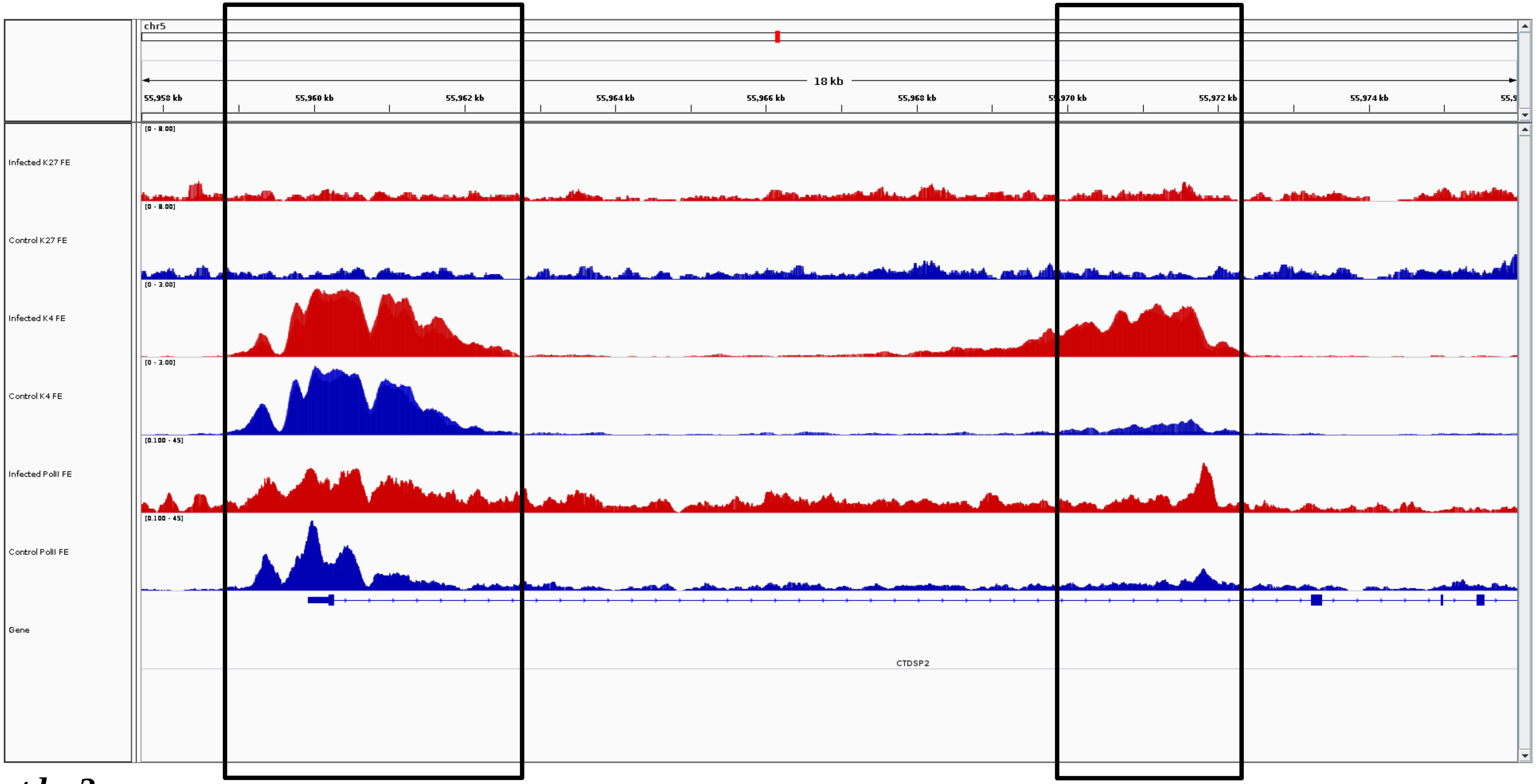
chr5
18 kb
55,958 kb
55,960 kb
55,962 kb
55,964 kb
55,966 kb
55,968 kb
55,972 kb
55,974 kb
[0 - 8.00]
Infected K27 FE
[0 - 8.00]
Control K27 FE
[0 - 3.00]
Infected K4 FE
[0 - 3.00]
Control K4 FE
[0.100 - 45]
Infected PolII FE
[0.100 - 45]
Control PolII FE
Gene
CTDSP2

*ctdsp2*

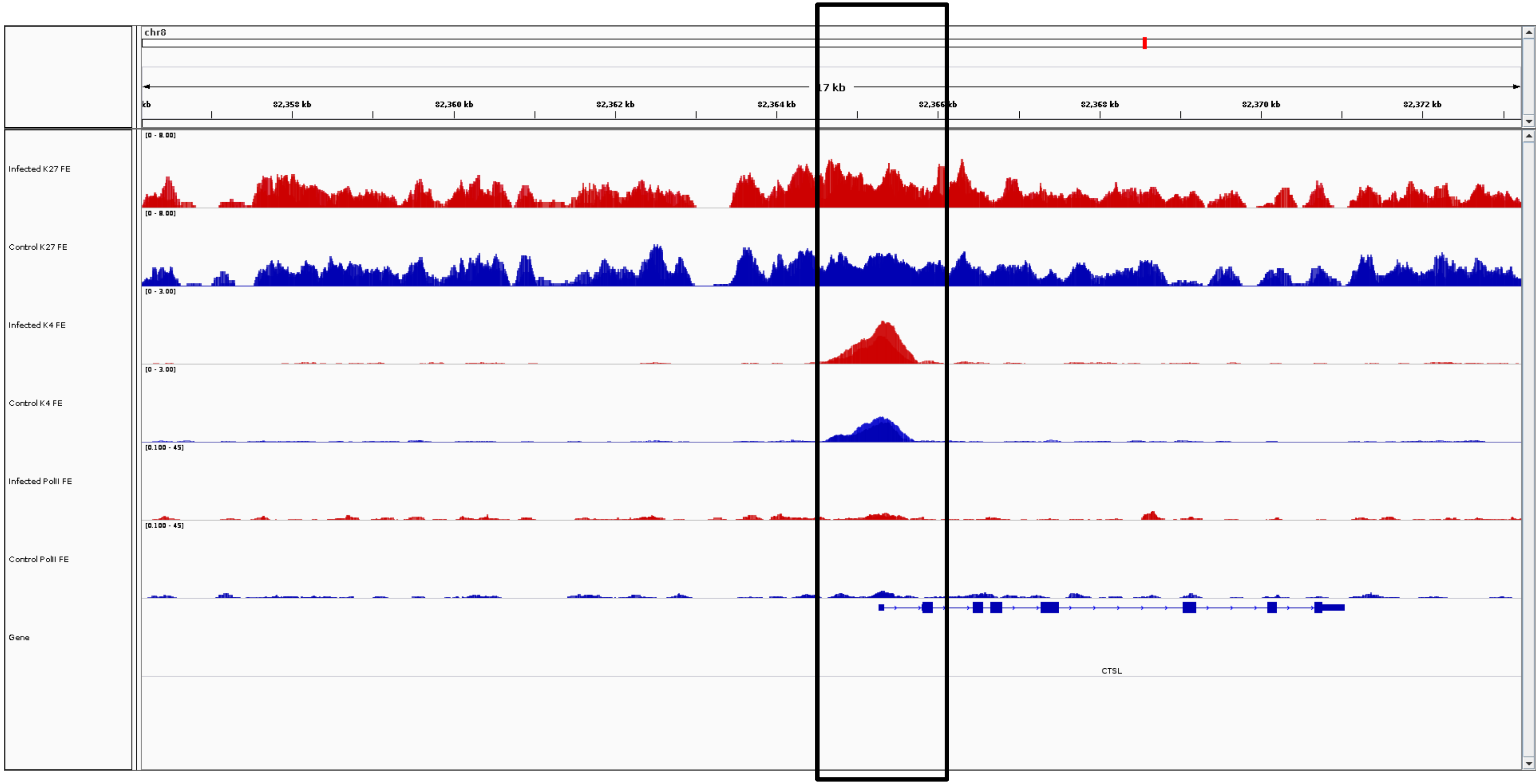
chr8
17 kb
kb
82,358 kb
82,360 kb
82,362 kb
82,364 kb
82,366 kb
82,368 kb
82,370 kb
82,372 kb
Infected K27 FE
[0 - 8.00]
Control K27 FE
[0 - 8.00]
Infected K4 FE
[0 - 3.00]
Control K4 FE
[0 - 3.00]
Infected PolII FE
[0.100 - 45]
Control PolII FE
[0.100 - 45]
Gene
CTSL

***CTSL***

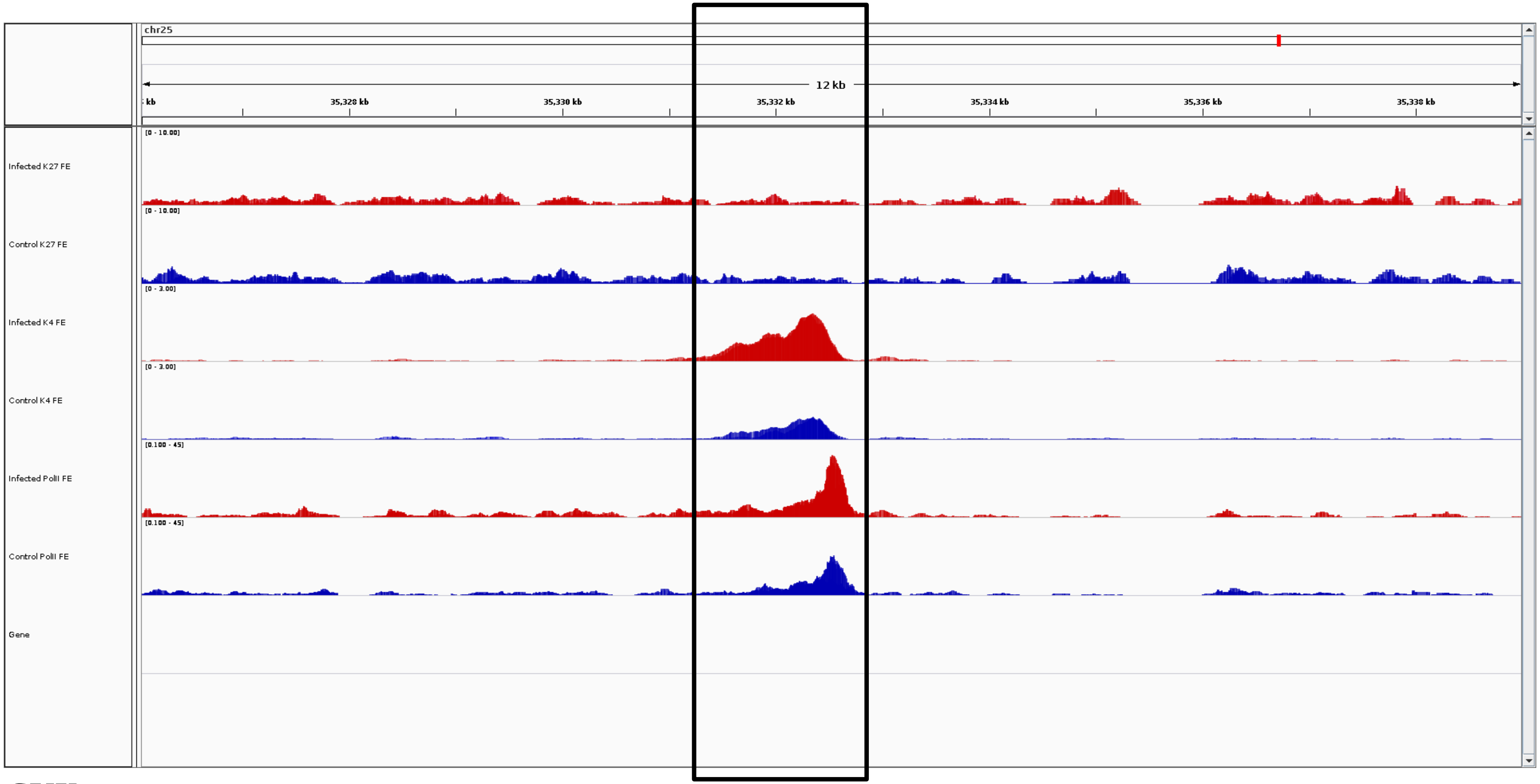

chr25
12 kb
35,328 kb
35,330 kb
35,332 kb
35,334 kb
35,336 kb
35,338 kb
Infected K27 FE
[0 - 10.00]
Control K27 FE
[0 - 10.00]
Infected K4 FE
[0 - 3.00]
Control K4 FE
[0 - 3.00]
Infected PolII FE
[0.100 - 45]
Control PolII FE
[0.100 - 45]
Gene


***CUX***

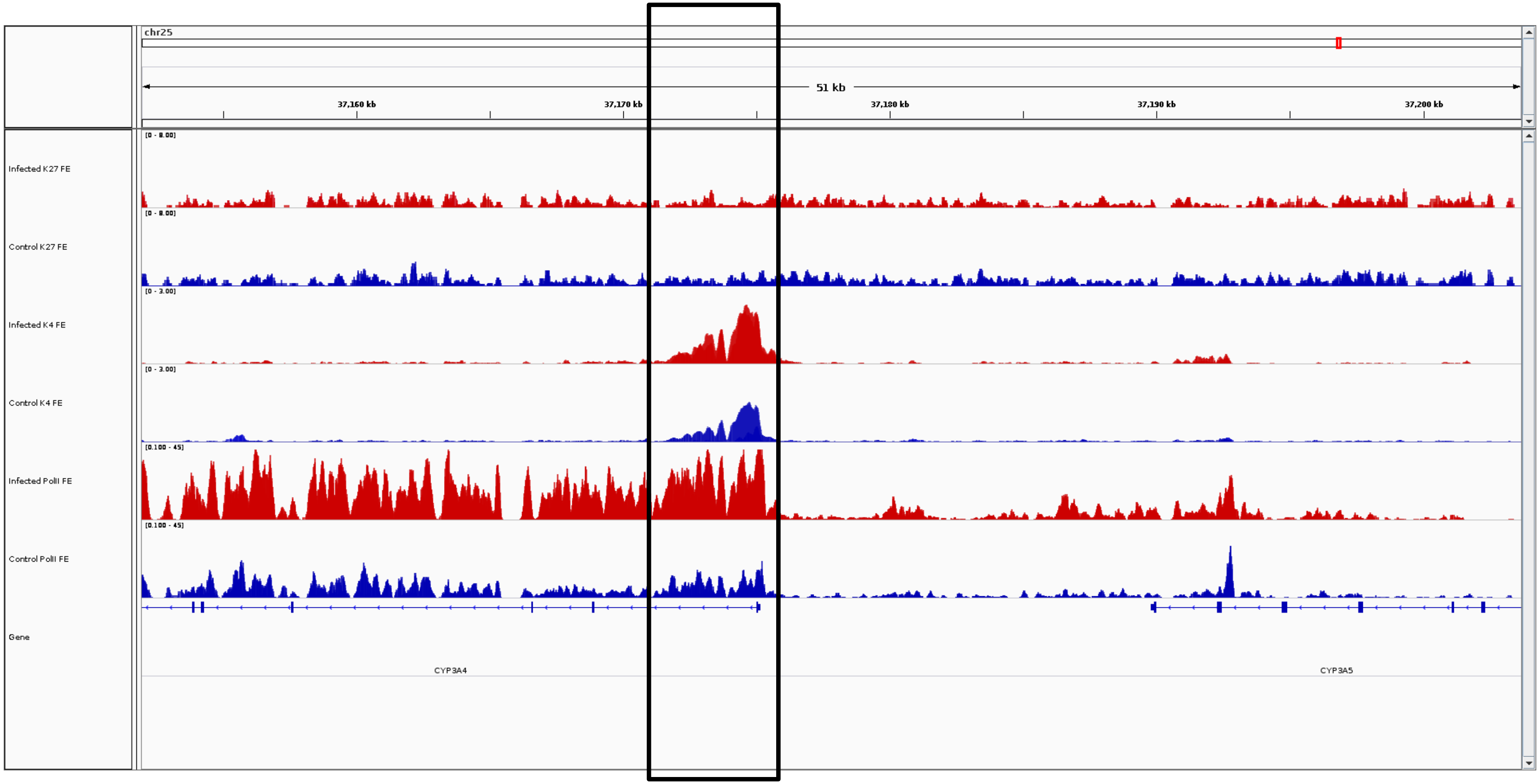
chr25
51 kb
37,160 kb
37,170 kb
37,180 kb
37,190 kb
37,200 kb
Infected K27 FE
Control K27 FE
Infected K4 FE
Control K4 FE
Infected PolII FE
Control PolII FE
Gene
CYP3A4
CYP3A5

***CYP3A4***

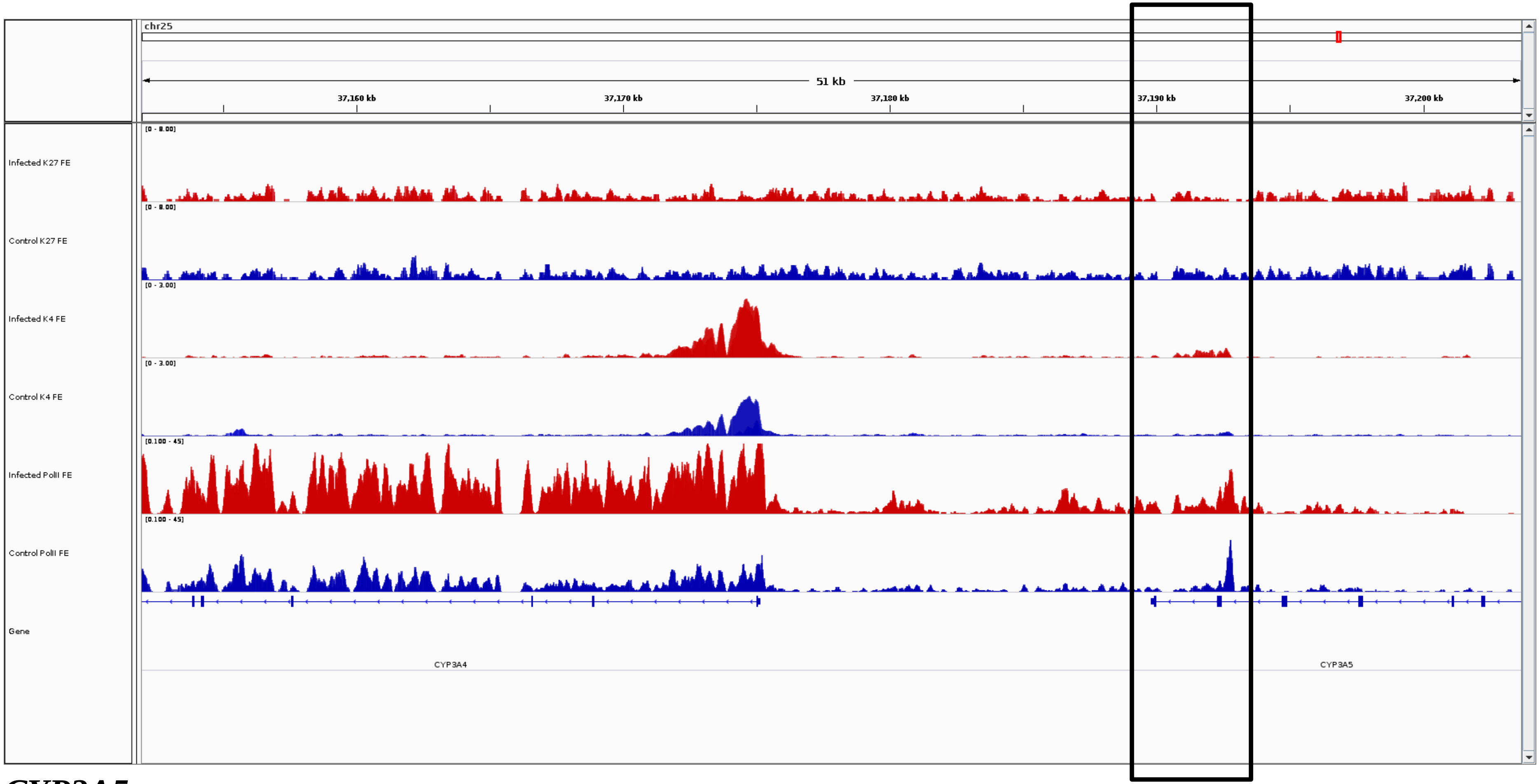
chr25
51 kb
37,160 kb
37,170 kb
37,180 kb
37,190 kb
37,200 kb
Infected K27 FE
[0 - 8.00]
Control K27 FE
[0 - 8.00]
Infected K4 FE
[0 - 3.00]
Control K4 FE
[0 - 3.00]
Infected PolII FE
[0.100 - 45]
Control PolII FE
[0.100 - 45]
Gene
CYP3A4
CYP3A5

***CYP3A5***

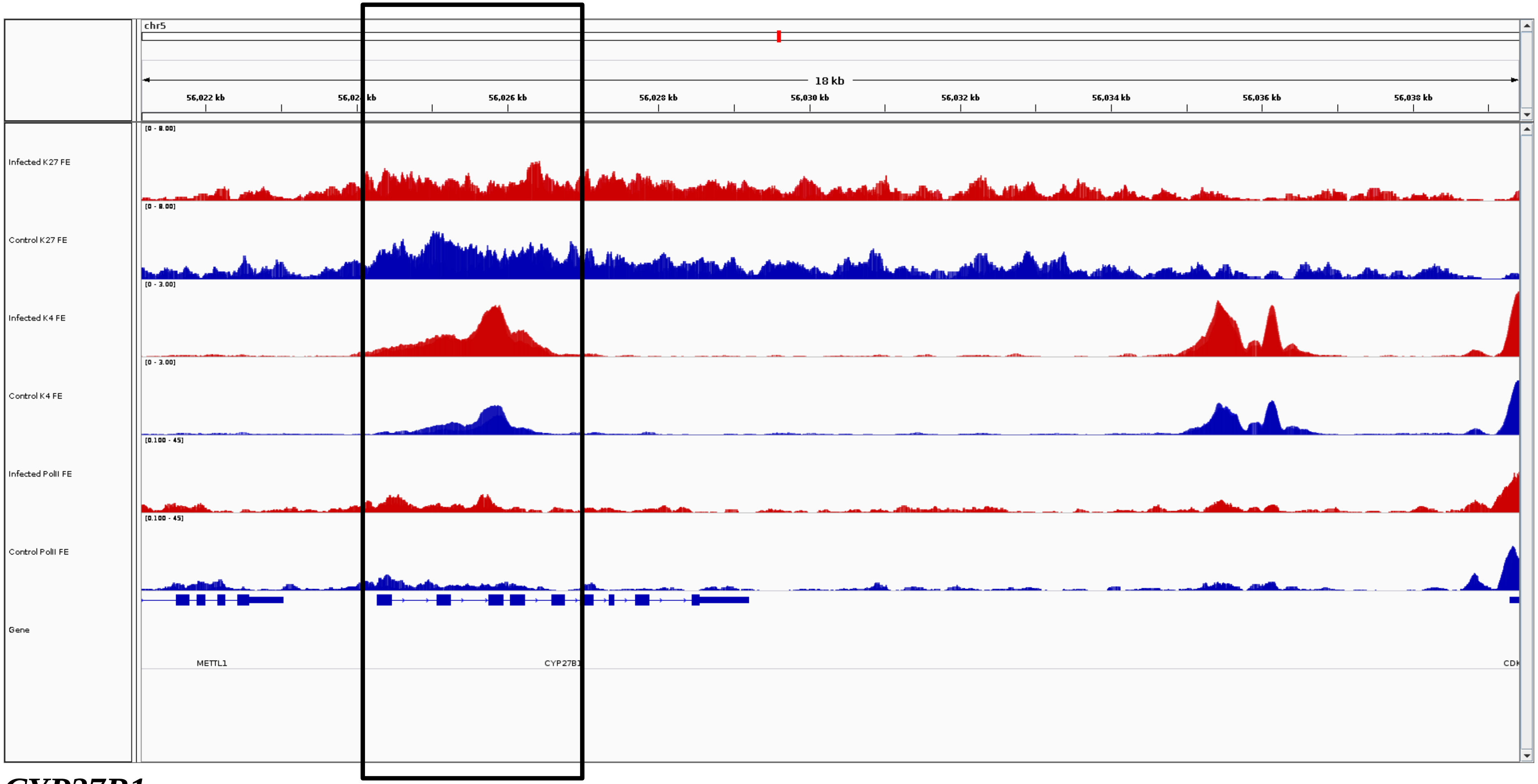
chr5
18 kb
56,022 kb
56,026 kb
56,028 kb
56,030 kb
56,032 kb
56,034 kb
56,036 kb
56,038 kb
Infected K27 FE
Control K27 FE
Infected K4 FE
Control K4 FE
Infected PolII FE
Control PolII FE
Gene
METTL1
CYP27B1

***CYP27B1***

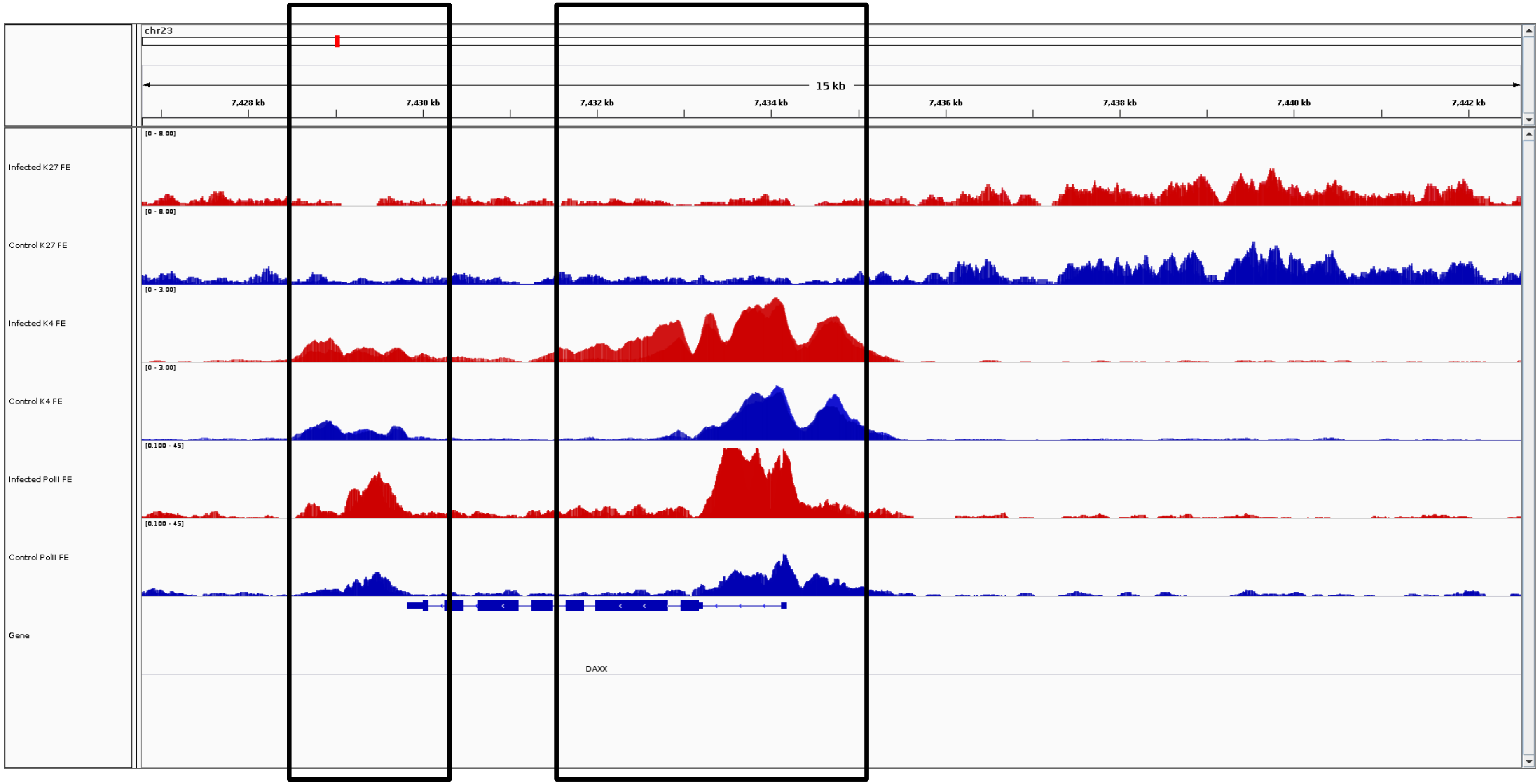
chr23
15 kb
7,428 kb
7,430 kb
7,432 kb
7,434 kb
7,436 kb
7,438 kb
7,440 kb
7,442 kb
Infected K27 FE
Control K27 FE
Infected K4 FE
Control K4 FE
Infected PolII FE
Control PolII FE
Gene
DAXX

***DAXX***

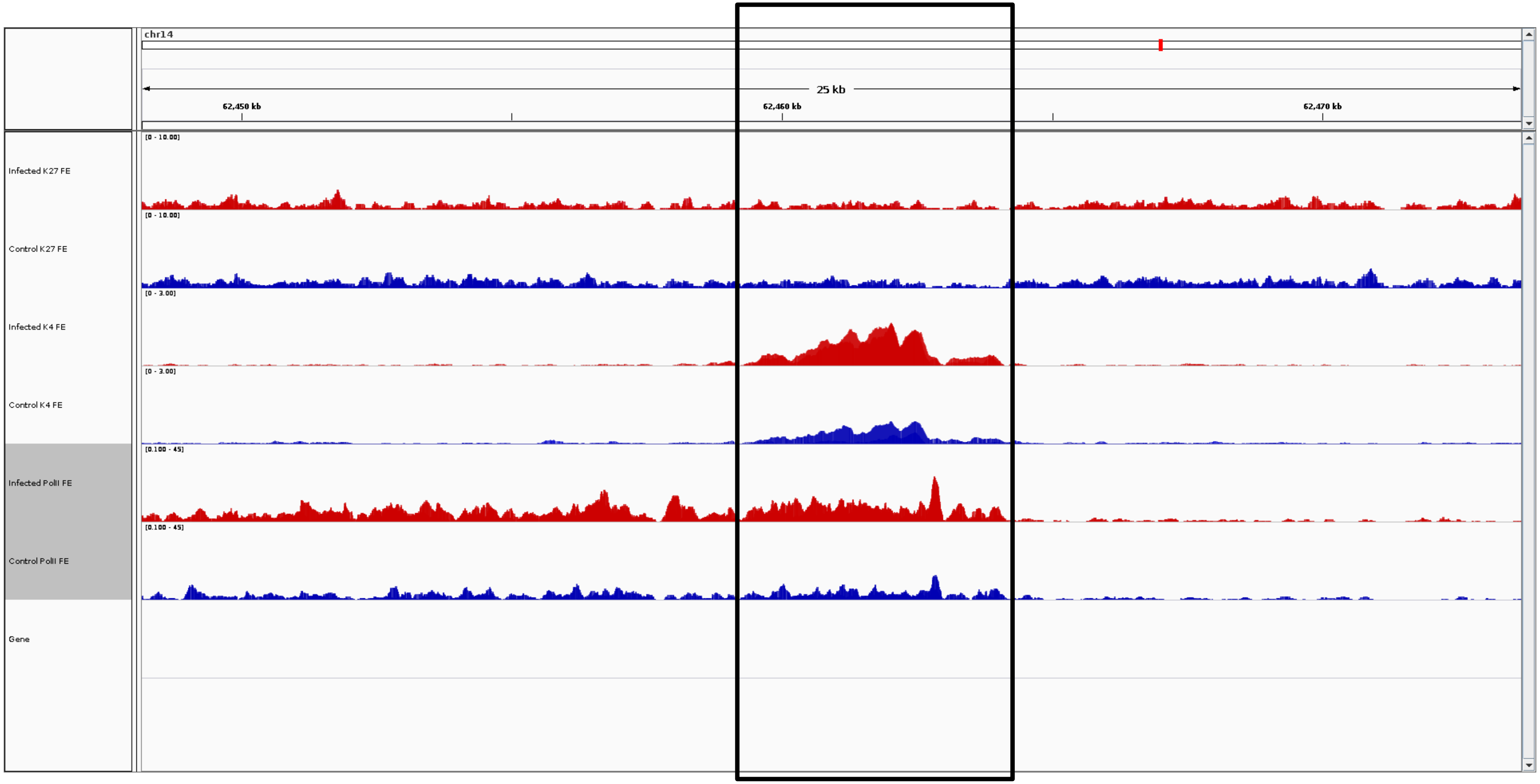
chr14
25 kb
62,450 kb
62,460 kb
62,470 kb
Infected K27 FE
Control K27 FE
Infected K4 FE
Control K4 FE
Infected PolII FE
Control PolII FE
Gene

***DCSTAMP***

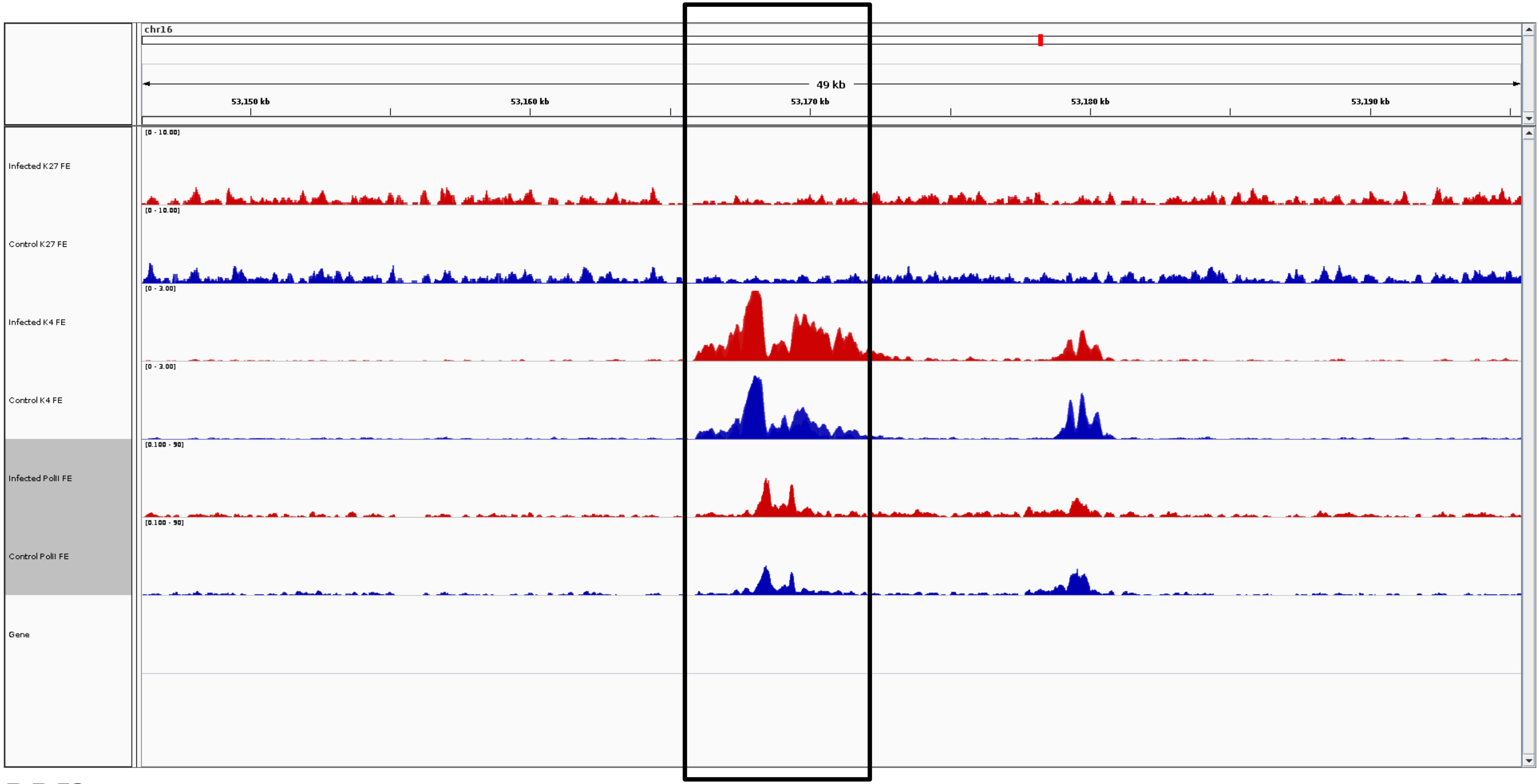
chr16
49 kb
53,150 kb
53,160 kb
53,170 kb
53,180 kb
53,190 kb
Infected K27 FE
[0 - 10.00]
Control K27 FE
[0 - 10.00]
Infected K4 FE
[0 - 3.00]
Control K4 FE
[0 - 3.00]
Infected PolII FE
[0.100 - 50]
Control PolII FE
[0.100 - 50]
Gene

***DDI2***

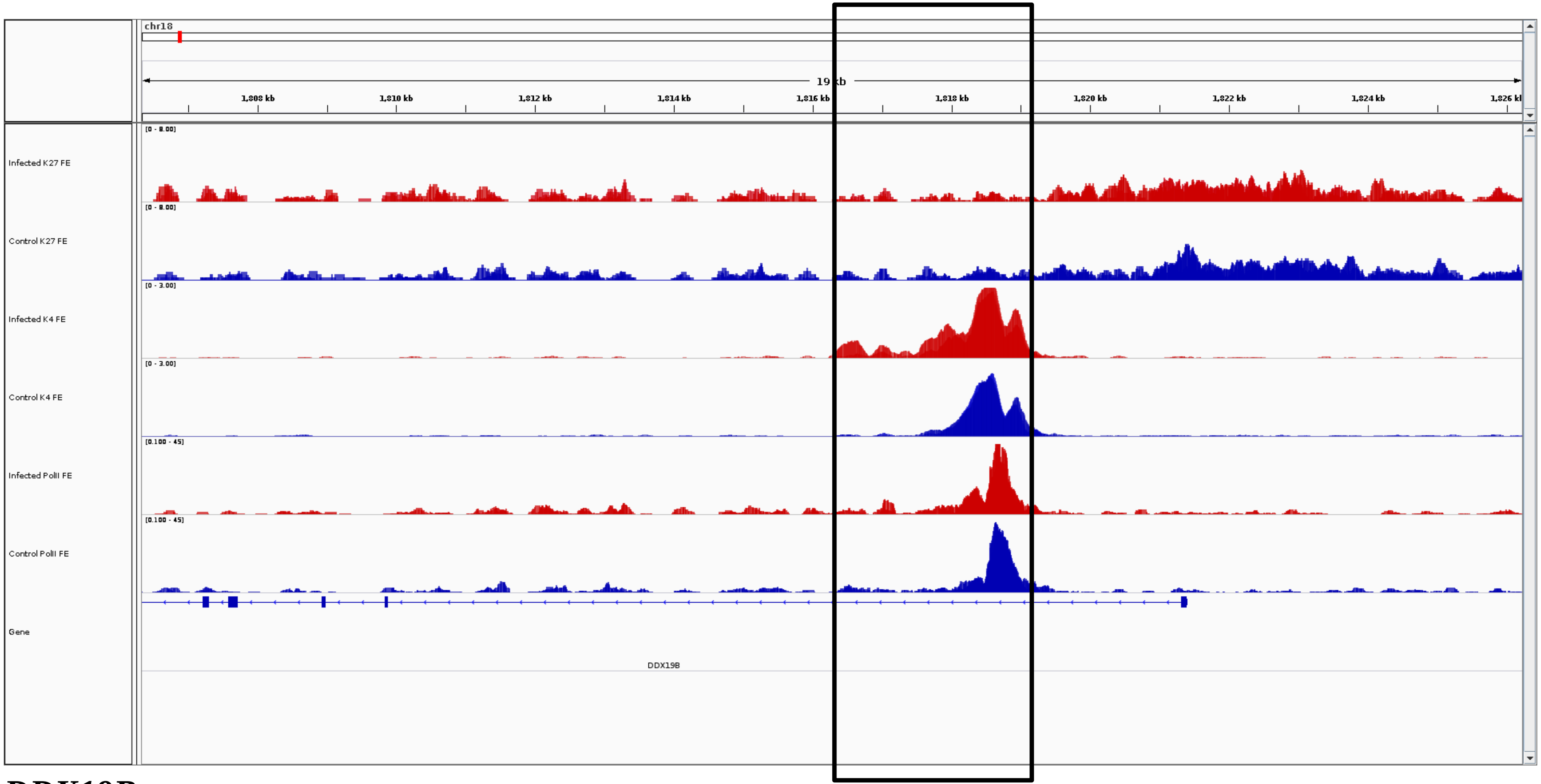
chr18
19 kb
1,808 kb
1,810 kb
1,812 kb
1,814 kb
1,816 kb
1,818 kb
1,820 kb
1,822 kb
1,824 kb
[0 - 8.00]
Infected K27 FE
[0 - 8.00]
Control K27 FE
[0 - 3.00]
Infected K4 FE
[0 - 3.00]
Control K4 FE
[0.100 - 45]
Infected PolII FE
[0.100 - 45]
Control PolII FE
Gene
DDX19B

***DDX19B***

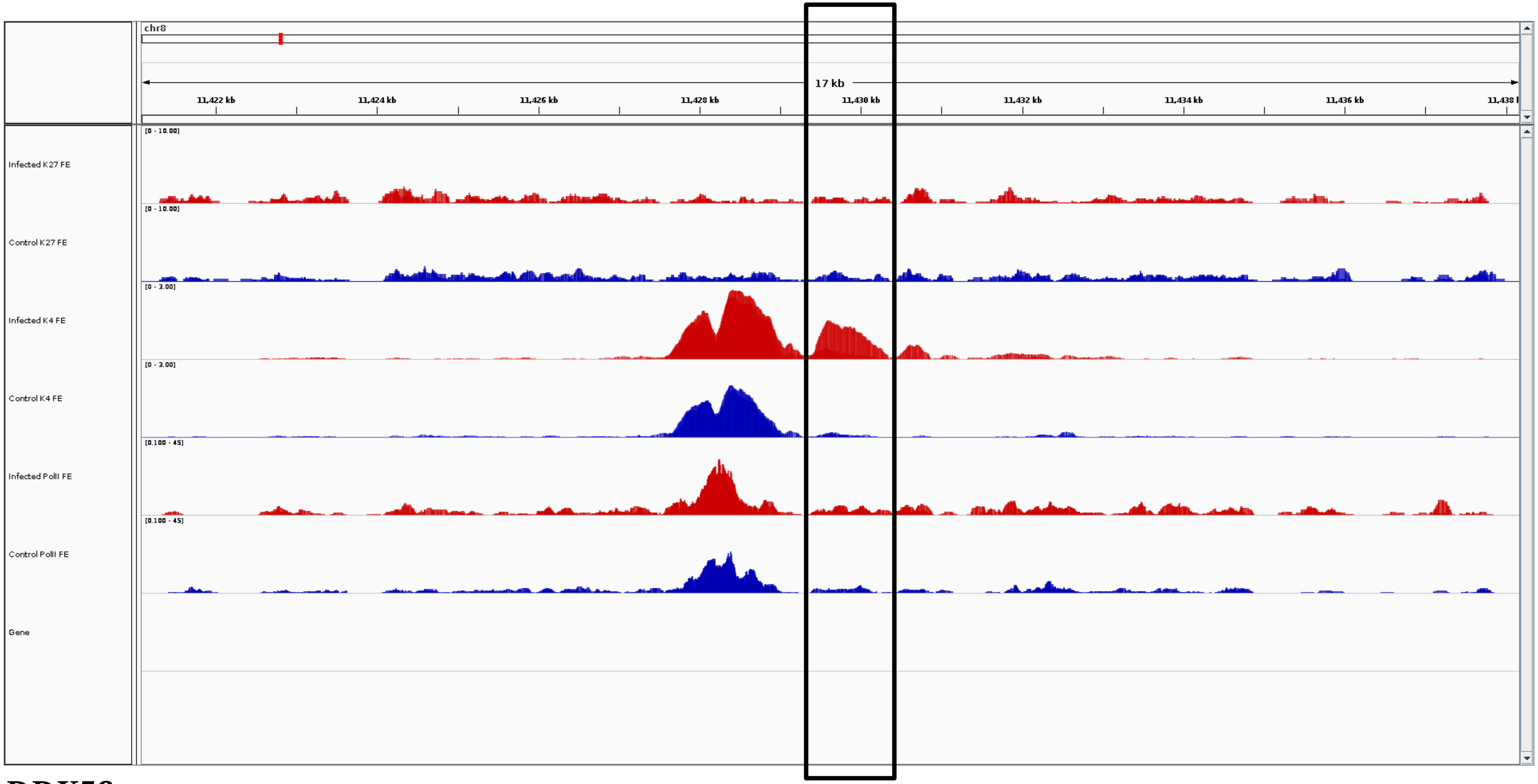
chr8
17 kb
11,422 kb
11,424 kb
11,426 kb
11,428 kb
11,430 kb
11,432 kb
11,434 kb
11,436 kb
Infected K27 FE
Control K27 FE
Infected K4 FE
Control K4 FE
Infected PolII FE
Control PolII FE
Gene

***DDX58***

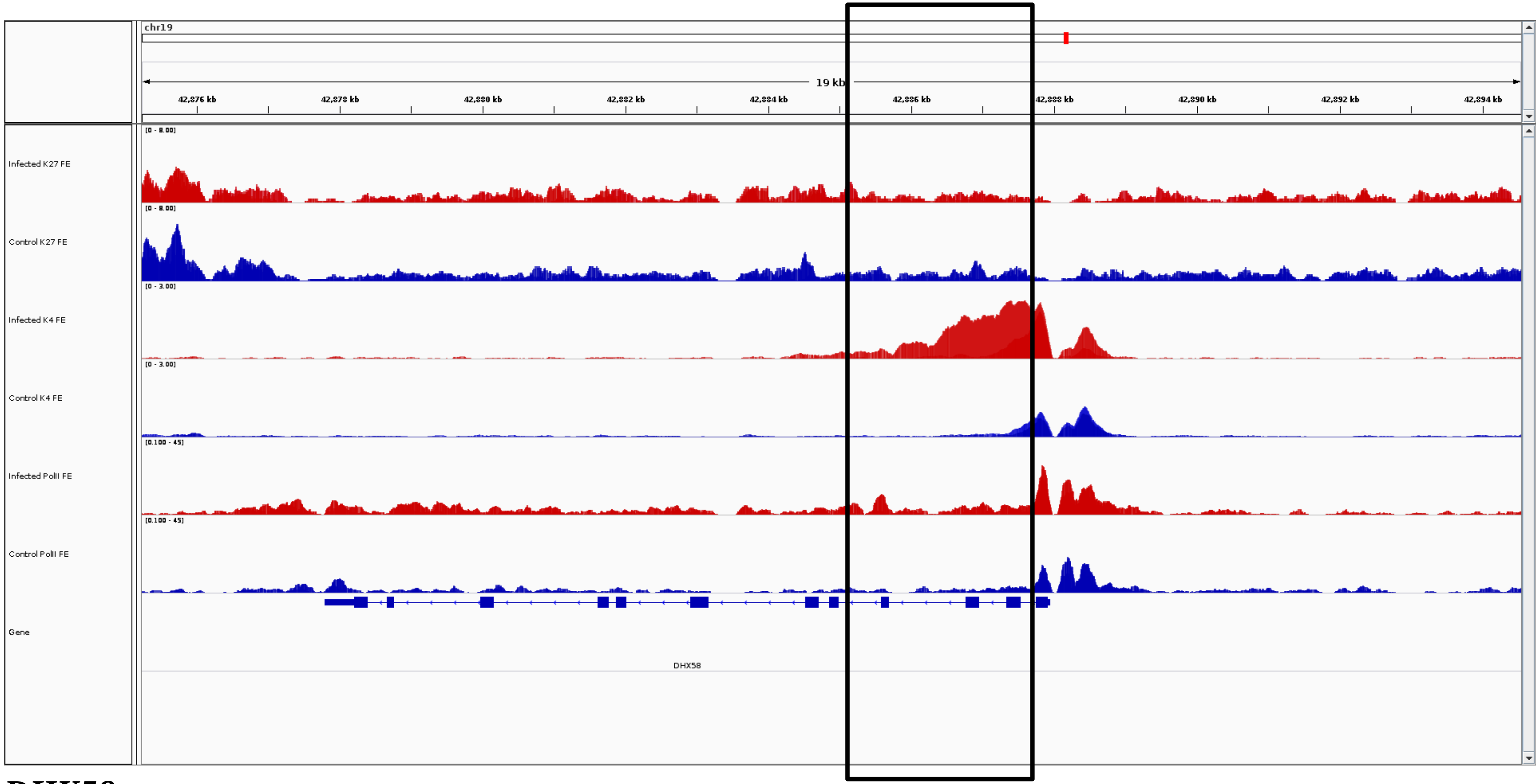
chr19
19 kb
42,876 kb
42,878 kb
42,880 kb
42,882 kb
42,884 kb
42,886 kb
42,888 kb
42,890 kb
42,892 kb
42,894 kb
Infected K27 FE
[0 - 8.00]
Control K27 FE
[0 - 8.00]
Infected K4 FE
[0 - 3.00]
Control K4 FE
[0 - 3.00]
Infected PolII FE
[0.100 - 45]
Control PolII FE
[0.100 - 45]
Gene
DHX58

***DHX58***

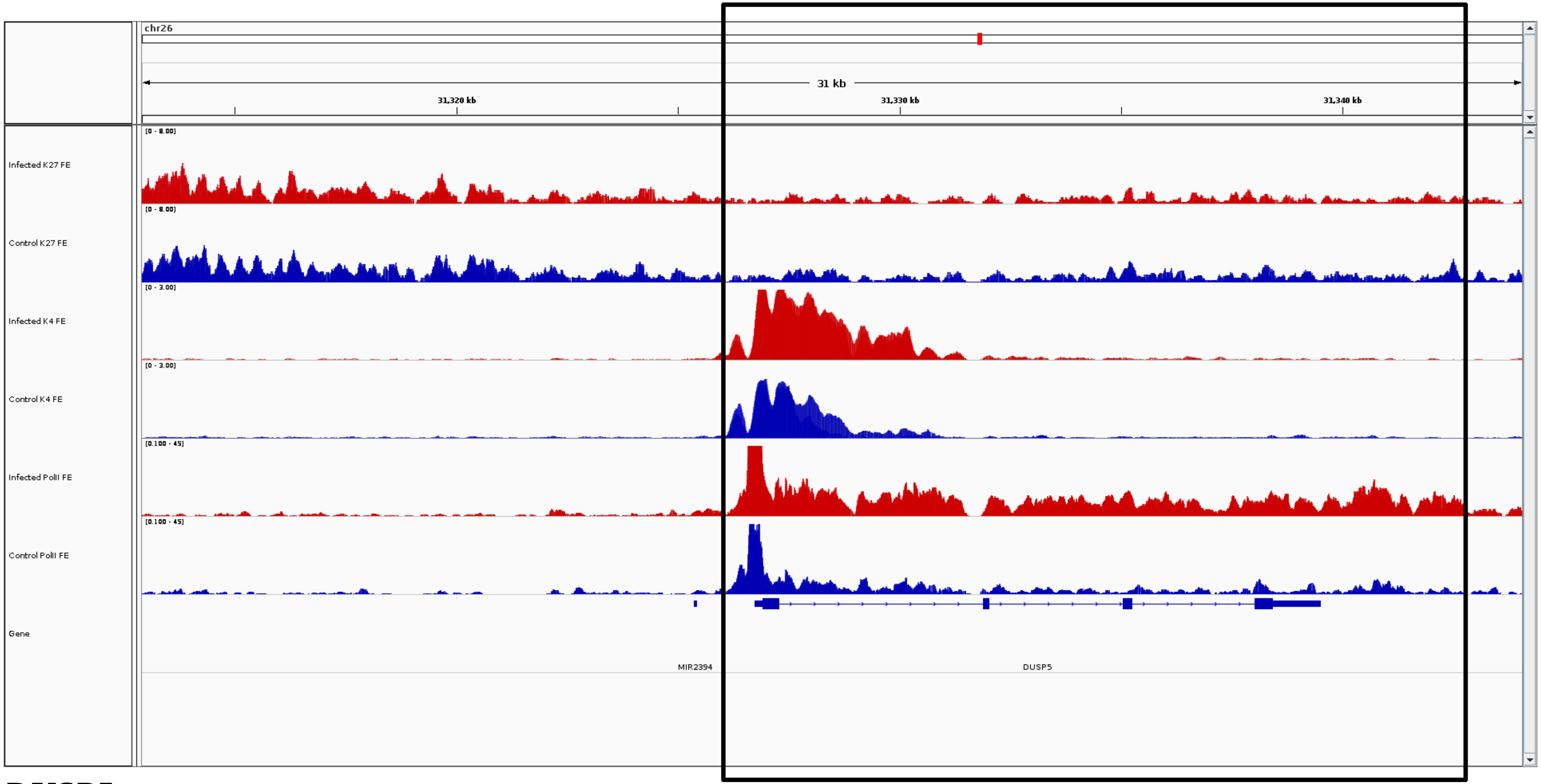
chr26
31 kb
31,320 kb
31,330 kb
31,340 kb
Infected K27 FE
Control K27 FE
Infected K4 FE
Control K4 FE
Infected PolII FE
Control PolII FE
Gene
MIR2394
DUSP5

***DUSP5***

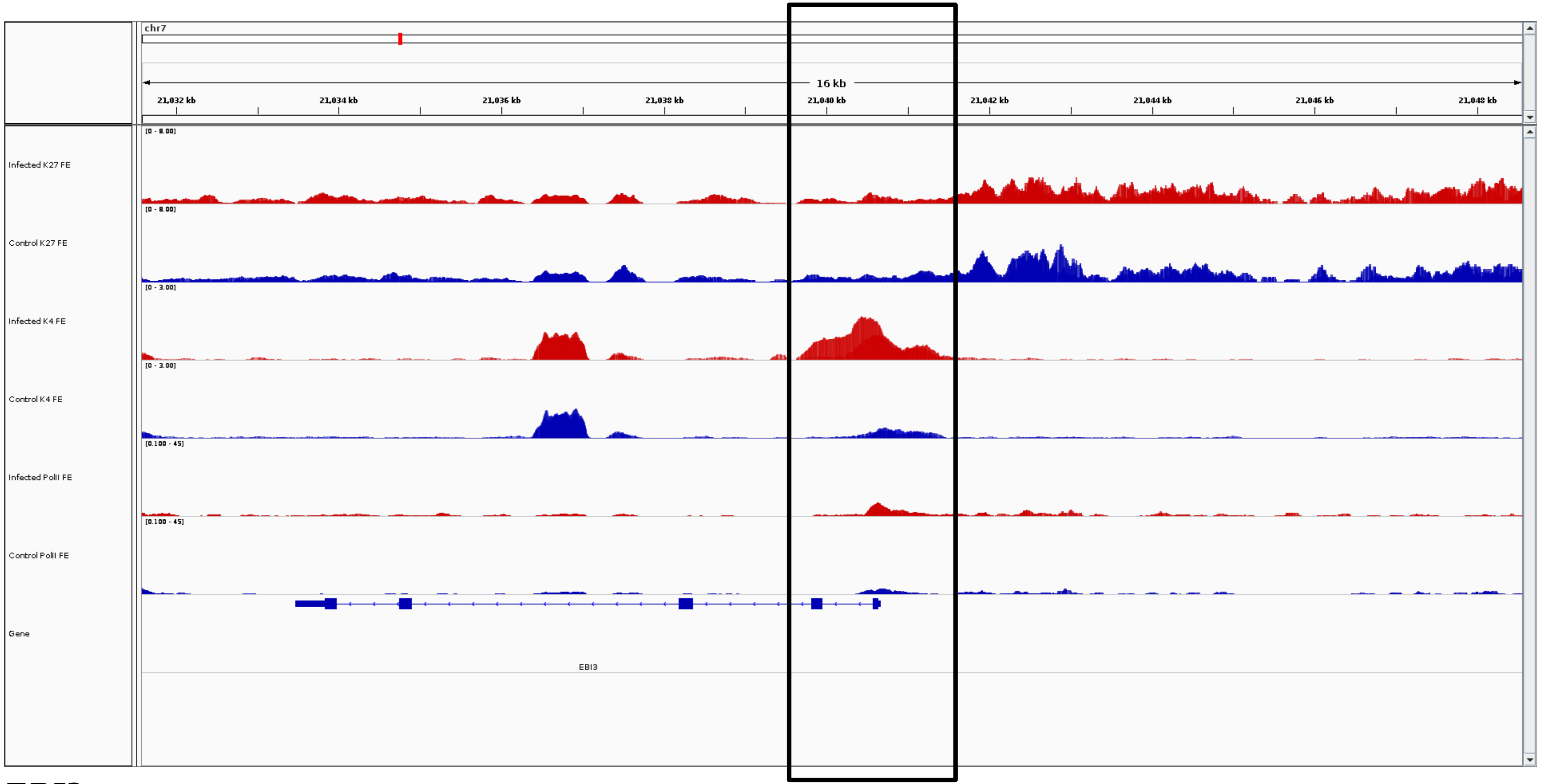
chr7
16 kb
21,032 kb
21,034 kb
21,036 kb
21,038 kb
21,040 kb
21,042 kb
21,044 kb
21,046 kb
21,048 kb
Infected K27 FE
[0 - 8.00]
Control K27 FE
[0 - 8.00]
Infected K4 FE
[0 - 3.00]
Control K4 FE
[0 - 3.00]
Infected PolII FE
[0.100 - 45]
Control PolII FE
[0.100 - 45]
Gene
EBI3

***EBI3***

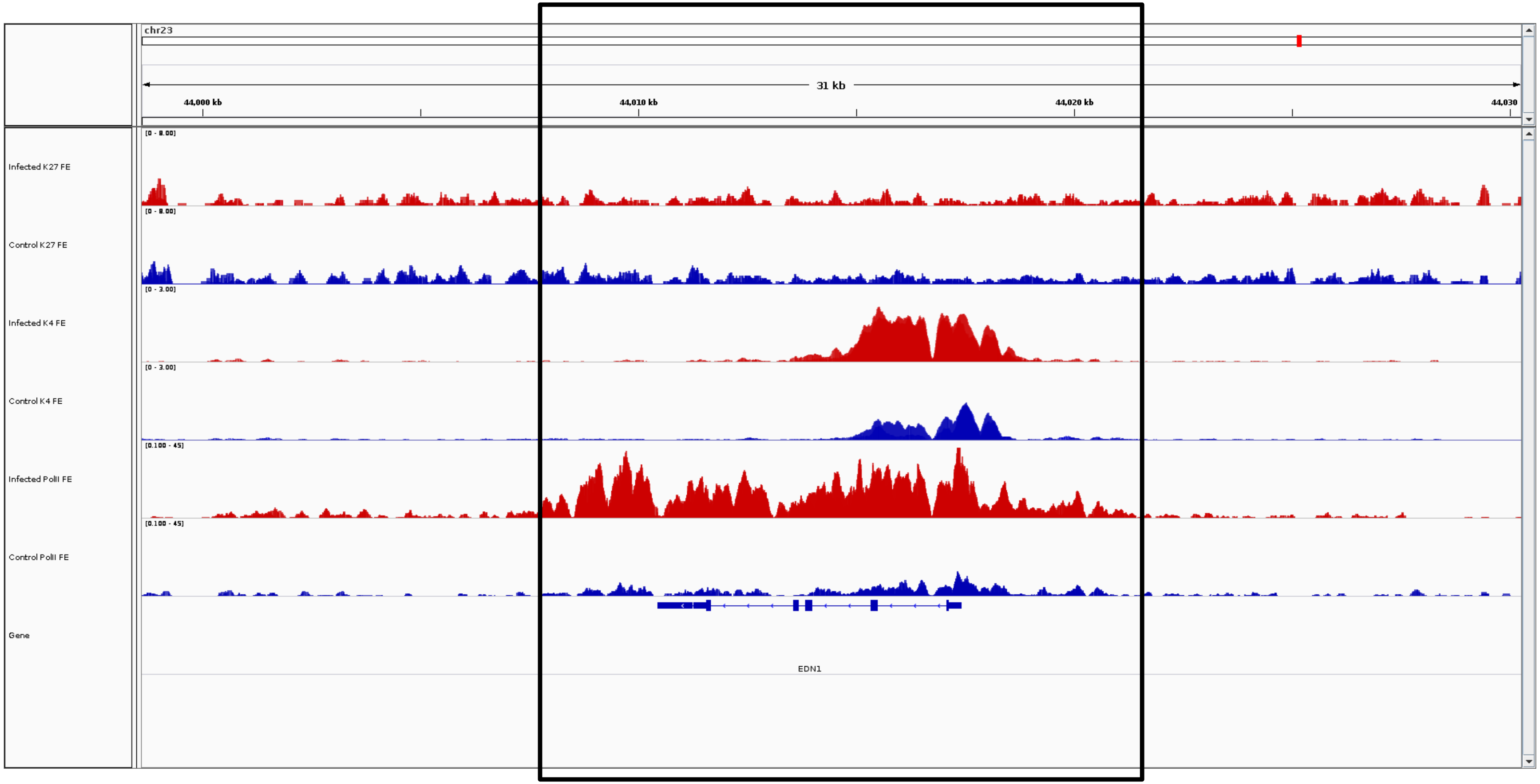
chr23
31 kb
44,000 kb
44,010 kb
44,020 kb
44,030
Infected K27 FE
Control K27 FE
Infected K4 FE
Control K4 FE
Infected PolII FE
Control PolII FE
Gene
EDN1

***EDN1***

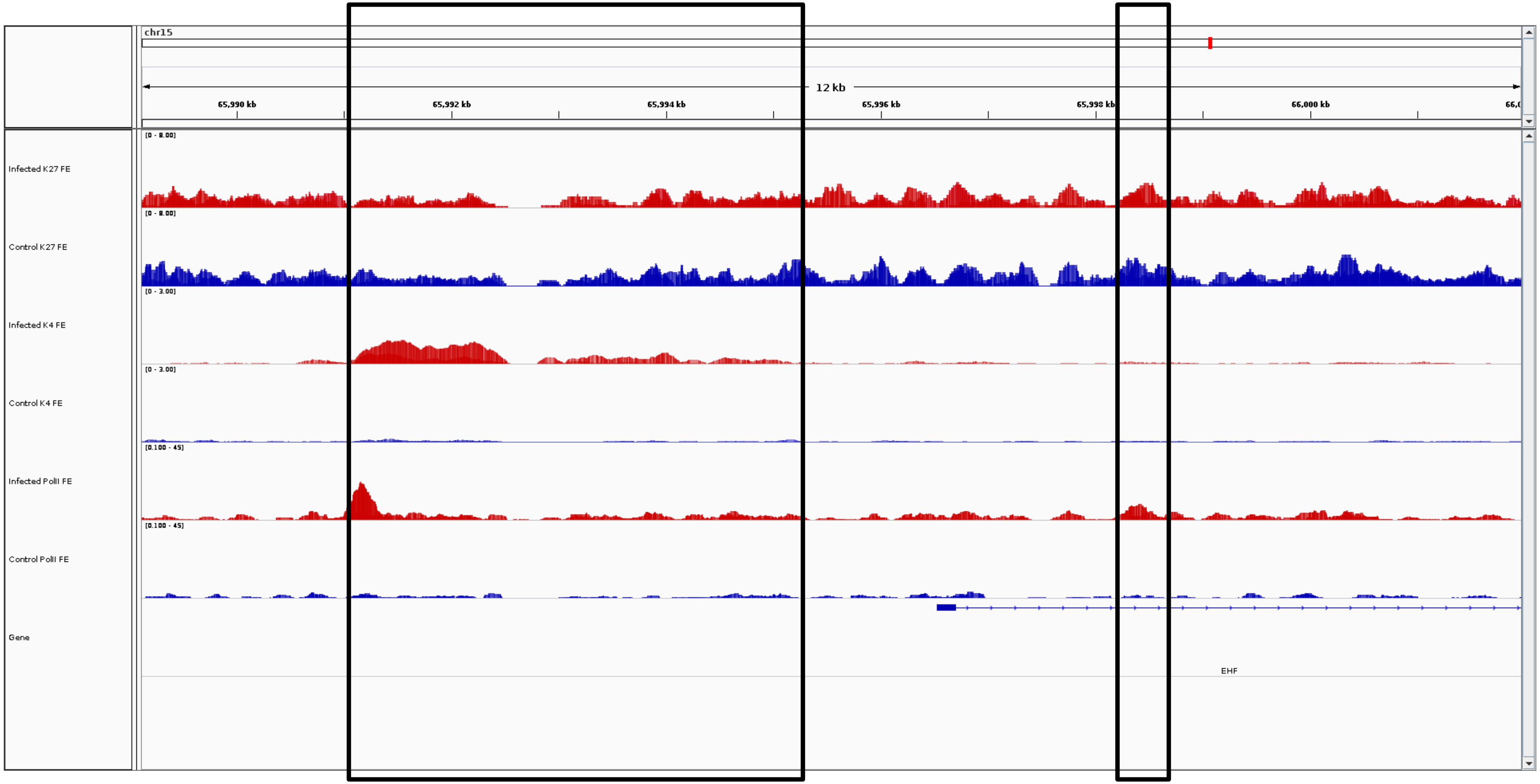

chr15
12 kb
65,990 kb
65,992 kb
65,994 kb
65,996 kb
65,998 kb
66,000 kb
Infected K27 FE
Control K27 FE
Infected K4 FE
Control K4 FE
Infected PolII FE
Control PolII FE
Gene
EHF


***EHF***

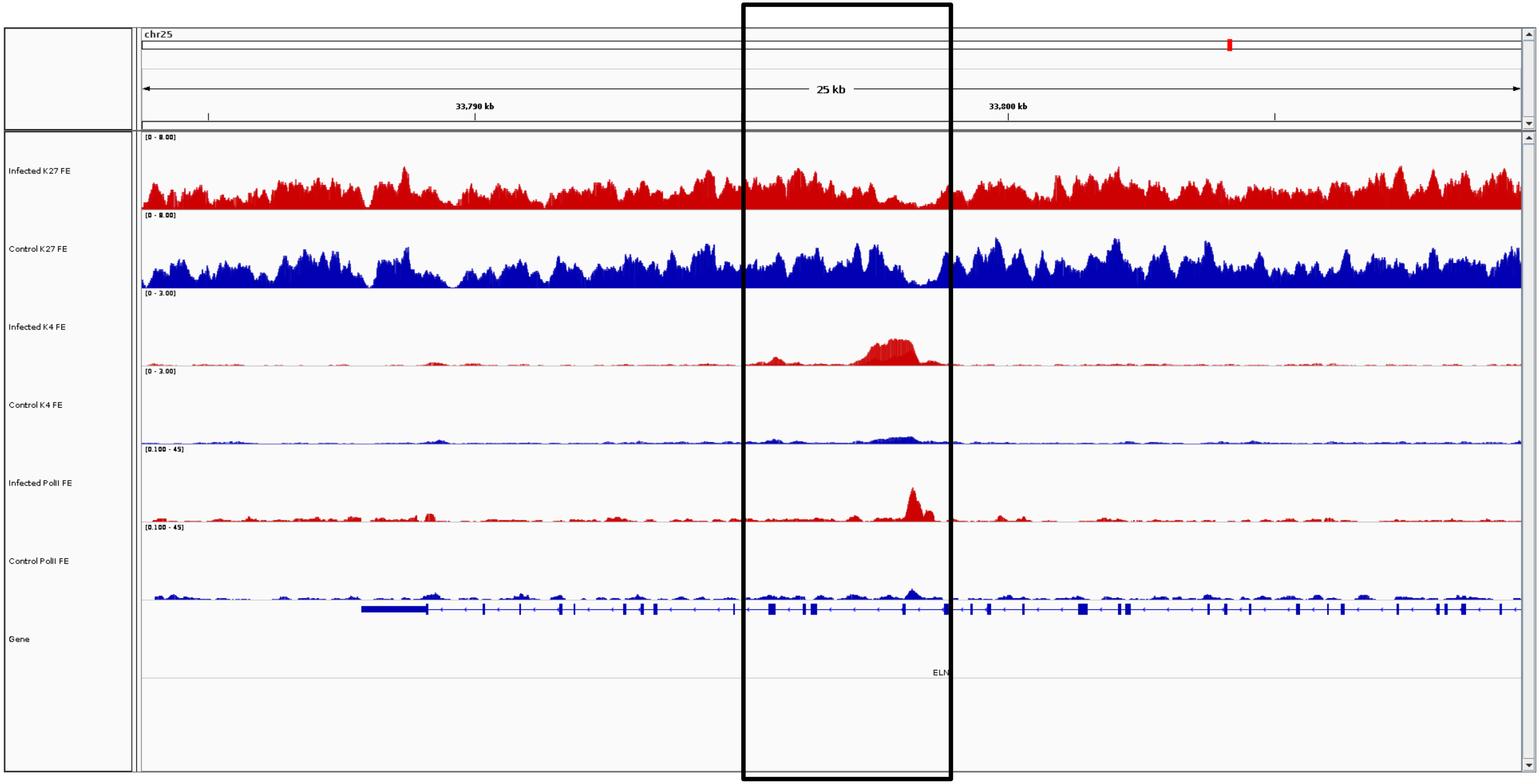
chr25
25 kb
33,790 kb
33,800 kb
[0 - 8.00]
Infected K27 FE
[0 - 8.00]
Control K27 FE
[0 - 3.00]
Infected K4 FE
[0 - 3.00]
Control K4 FE
[0.100 - 45]
Infected PolII FE
[0.100 - 45]
Control PolII FE
Gene
ELN

***ELN***

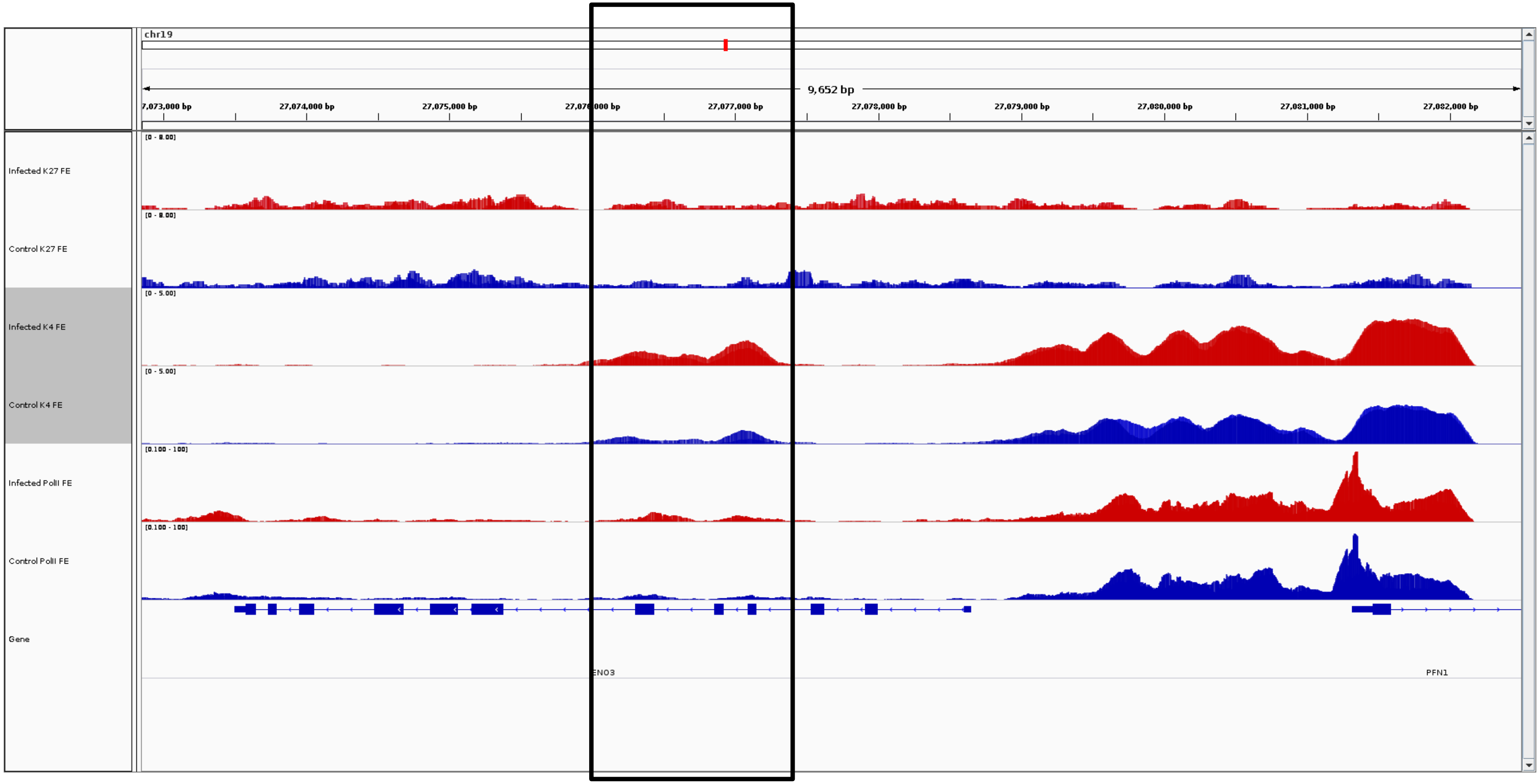
chr19
9,652 bp
7,073,000 bp
27,074,000 bp
27,075,000 bp
27,077,000 bp
27,078,000 bp
27,079,000 bp
27,080,000 bp
27,081,000 bp
27,082,000 bp
Infected K27 FE
Control K27 FE
Infected K4 FE
Control K4 FE
Infected PolII FE
Control PolII FE
Gene
ENO3
PFN1

***ENO3***

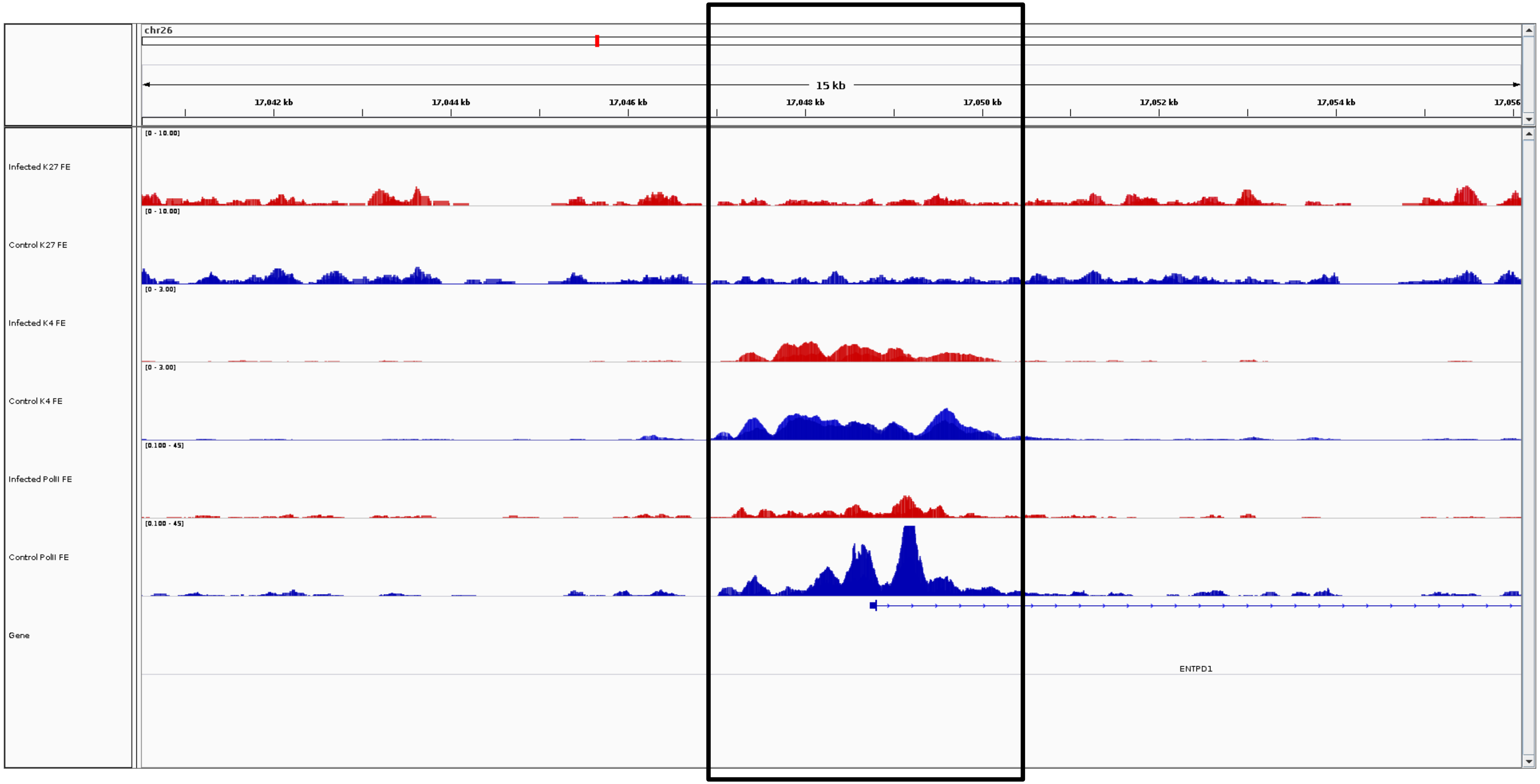
chr26
15 kb
17,042 kb
17,044 kb
17,046 kb
17,048 kb
17,050 kb
17,052 kb
17,054 kb
17,056
Infected K27 FE
Control K27 FE
Infected K4 FE
Control K4 FE
Infected PolII FE
Control PolII FE
Gene
ENTPD1

***ENTPD1***

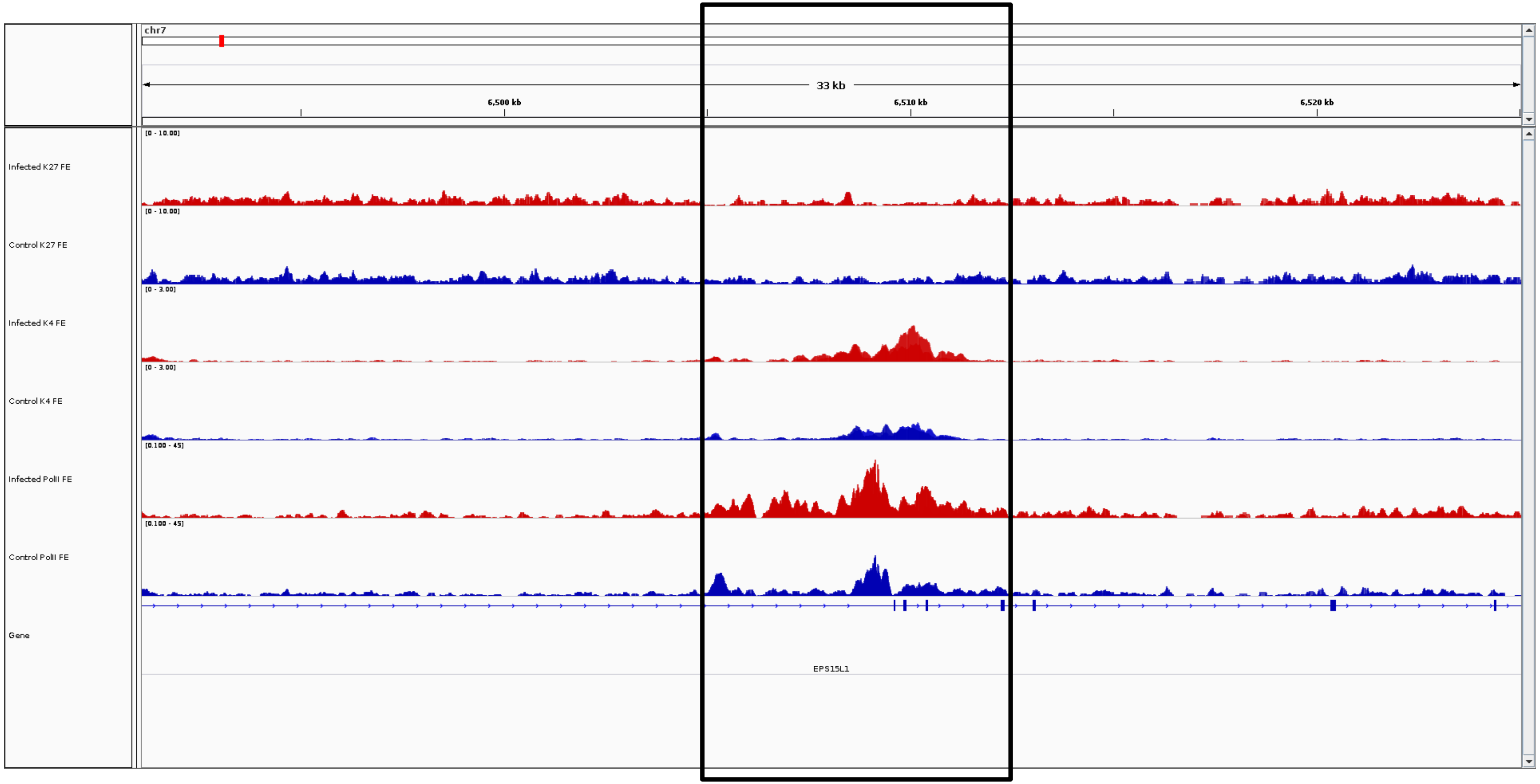
chr7
33 kb
6,500 kb
6,510 kb
6,520 kb
Infected K27 FE
[0 - 10.00]
Control K27 FE
[0 - 10.00]
Infected K4 FE
[0 - 3.00]
Control K4 FE
[0 - 3.00]
Infected PolII FE
[0.100 - 45]
Control PolII FE
[0.100 - 45]
Gene
EPS15L1

***EPS15L1***

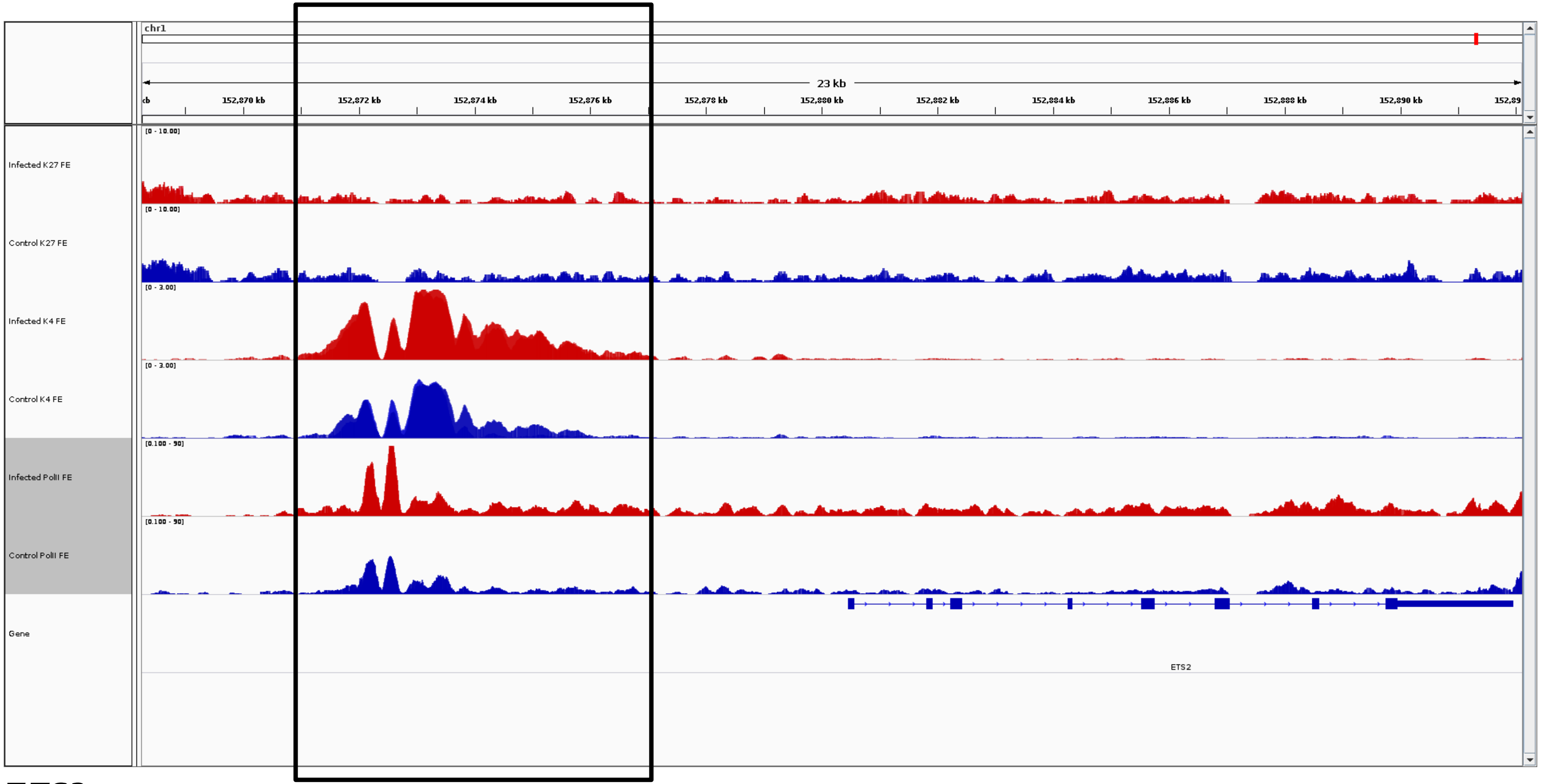
chr1
23 kb
152,870 kb
152,872 kb
152,874 kb
152,876 kb
152,878 kb
152,880 kb
152,882 kb
152,884 kb
152,886 kb
152,888 kb
152,890 kb
Infected K27 FE
Control K27 FE
Infected K4 FE
Control K4 FE
Infected PolII FE
Control PolII FE
Gene
ETS2

***ETS2***

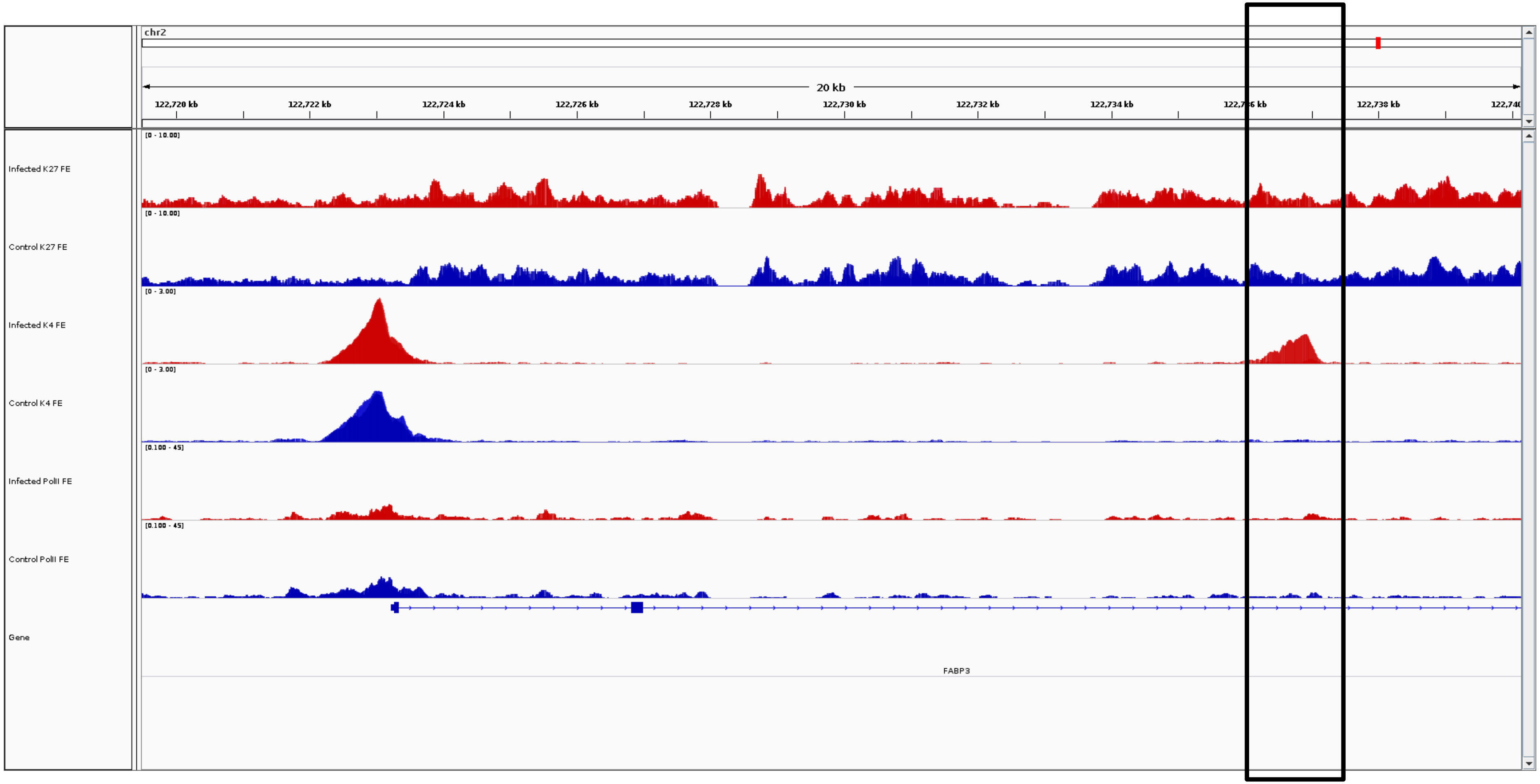
chr2
20 kb
122,720 kb
122,722 kb
122,724 kb
122,726 kb
122,728 kb
122,730 kb
122,732 kb
122,734 kb
122,738 kb
[0 - 10.00]
Infected K27 FE
Control K27 FE
[0 - 3.00]
Infected K4 FE
Control K4 FE
[0.100 - 45]
Infected PolII FE
Control PolII FE
Gene
FABP3

***FABP3***

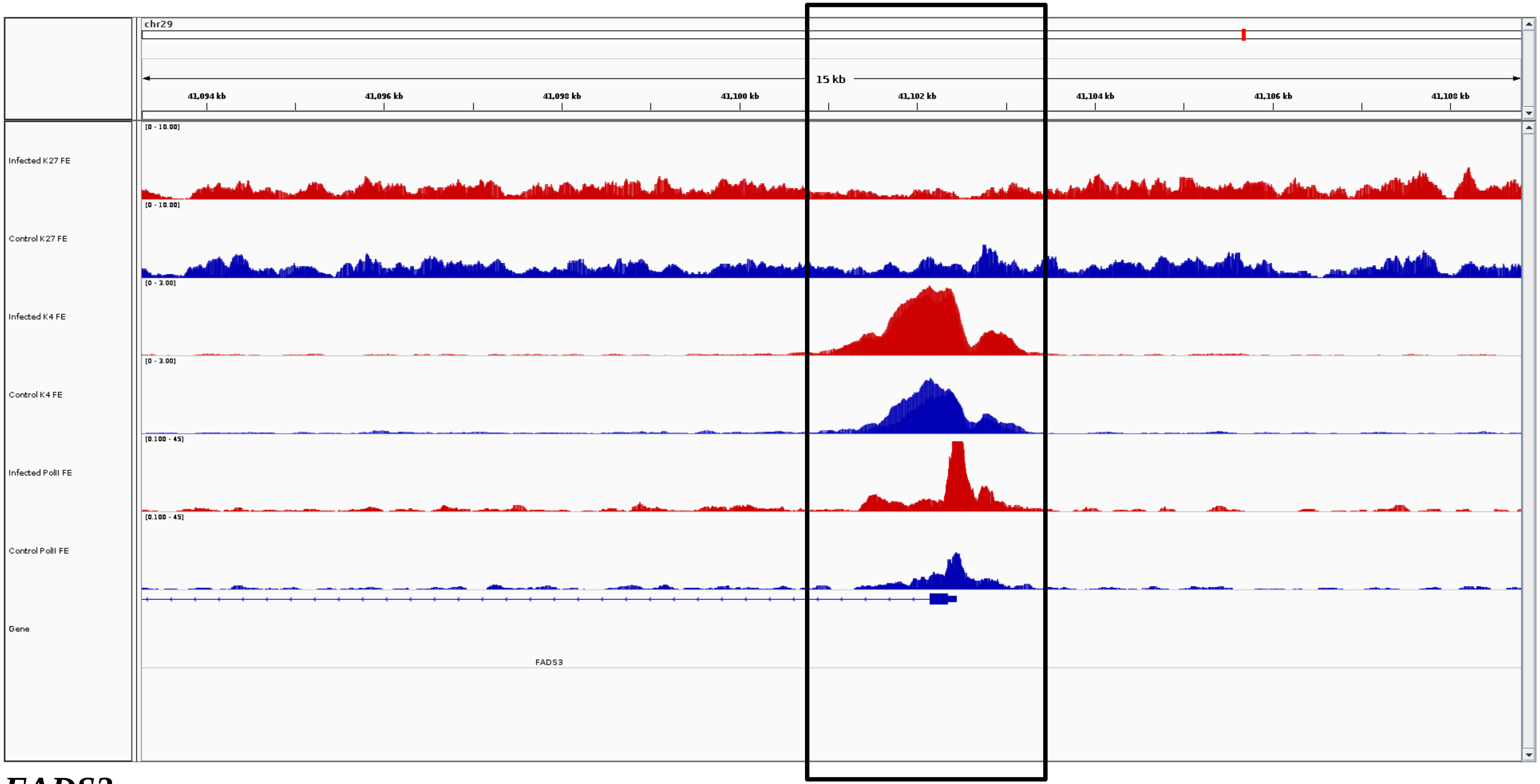
chr29
15 kb
41,094 kb
41,096 kb
41,098 kb
41,100 kb
41,102 kb
41,104 kb
41,106 kb
41,108 kb
Infected K27 FE
Control K27 FE
Infected K4 FE
Control K4 FE
Infected PolII FE
Control PolII FE
Gene
FADS3

***FADS3***

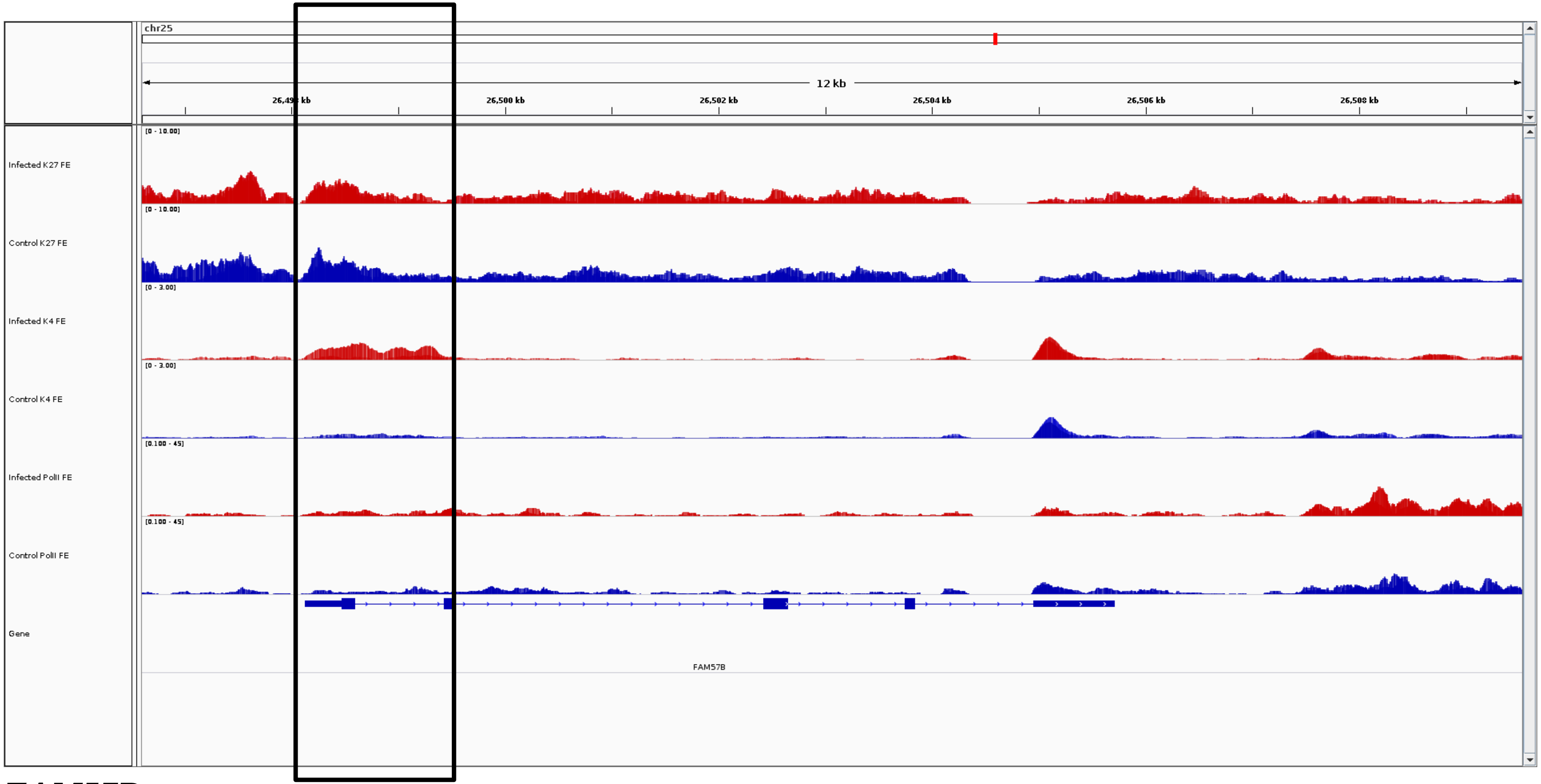
chr25
12 kb
26,500 kb
26,502 kb
26,504 kb
26,506 kb
26,508 kb
[0 - 10.00]
Infected K27 FE
[0 - 10.00]
Control K27 FE
[0 - 3.00]
Infected K4 FE
[0 - 3.00]
Control K4 FE
[0.100 - 45]
Infected PolII FE
[0.100 - 45]
Control PolII FE
Gene
FAM57B

***FAM57B***

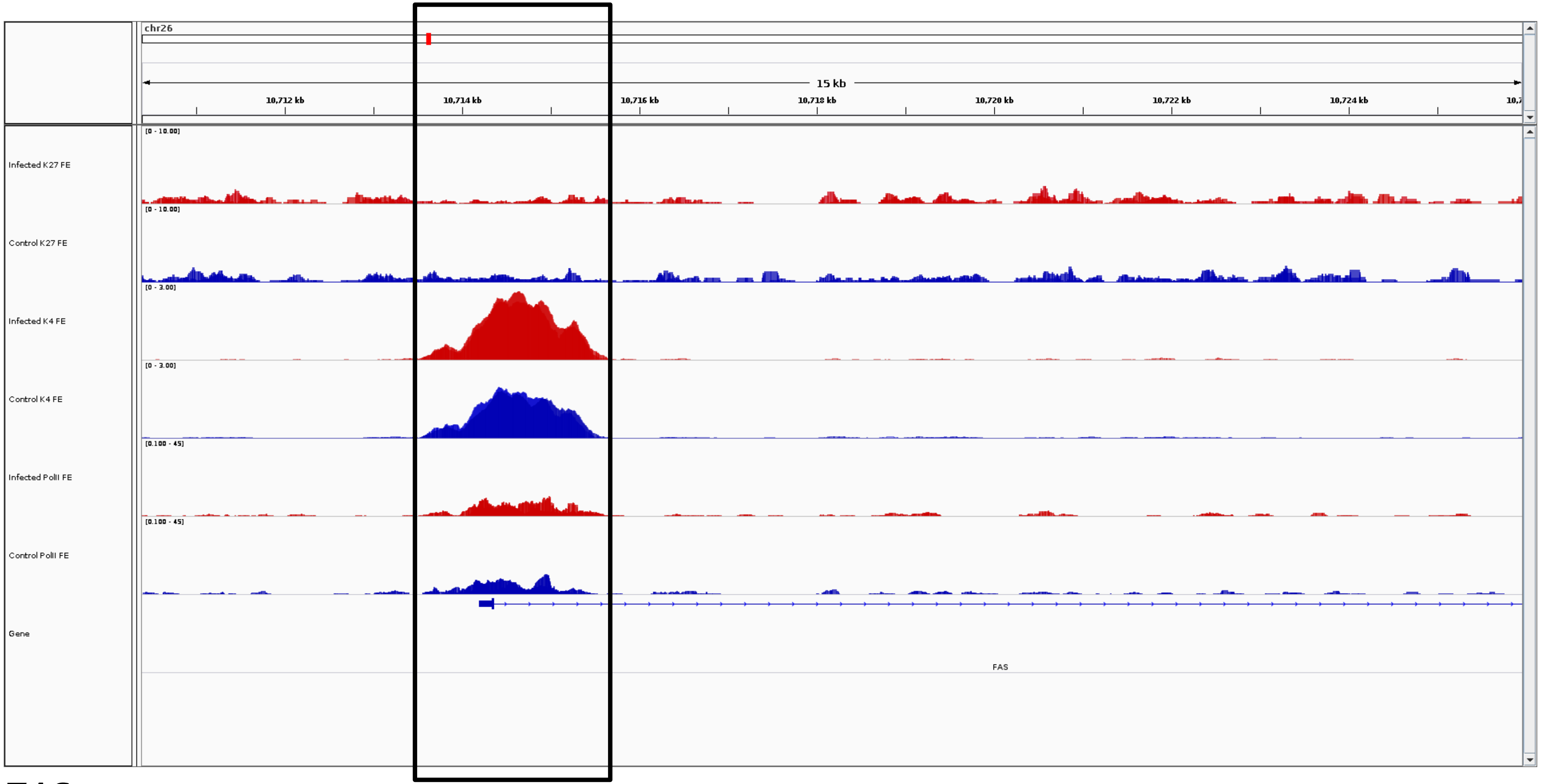
chr26
15 kb
10,712 kb
10,714 kb
10,716 kb
10,718 kb
10,720 kb
10,722 kb
10,724 kb
Infected K27 FE
[0 - 10.00]
Control K27 FE
[0 - 10.00]
Infected K4 FE
[0 - 3.00]
Control K4 FE
[0 - 3.00]
Infected PolII FE
[0.100 - 45]
Control PolII FE
[0.100 - 45]
Gene
FAS

***FAS***

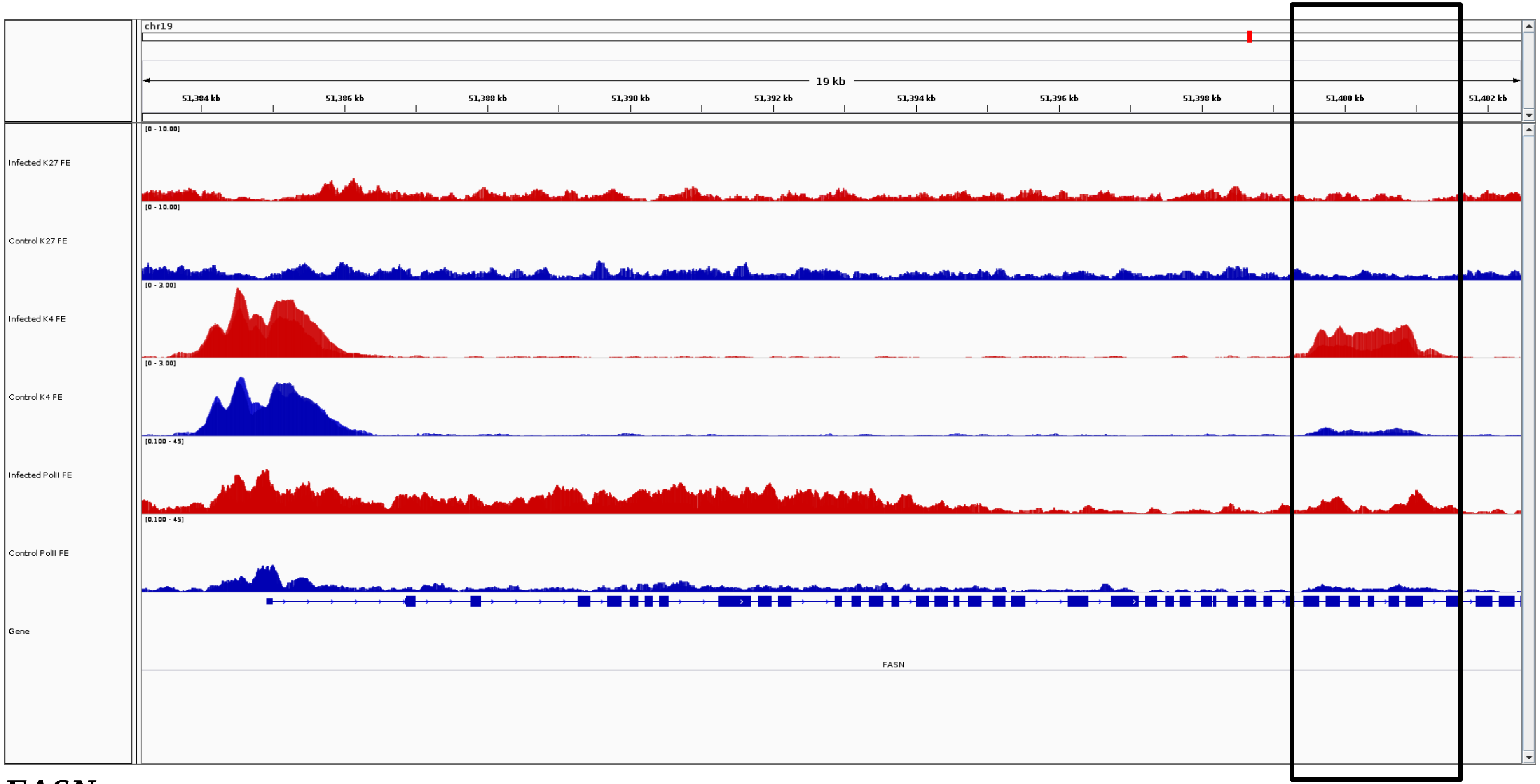
chr19
19 kb
51,384 kb
51,386 kb
51,388 kb
51,390 kb
51,392 kb
51,394 kb
51,396 kb
51,398 kb
51,400 kb
51,402 kb
Infected K27 FE
[0 - 10.00]
Control K27 FE
[0 - 10.00]
Infected K4 FE
[0 - 3.00]
Control K4 FE
[0 - 3.00]
Infected PolII FE
[0.100 - 45]
Control PolII FE
[0.100 - 45]
Gene
FASN

***FASN***

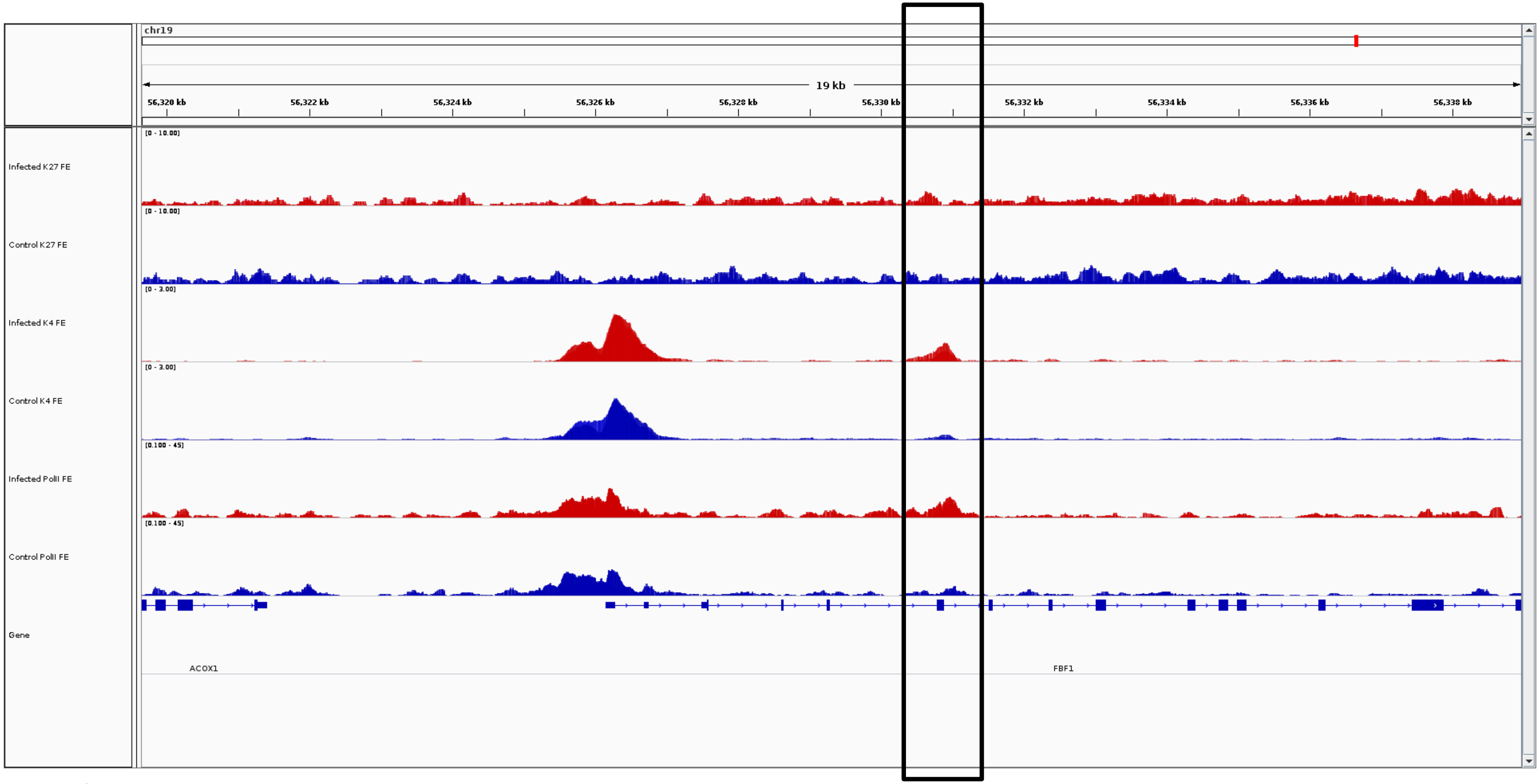
chr19
19 kb
56,320 kb
56,322 kb
56,324 kb
56,326 kb
56,328 kb
56,330 kb
56,332 kb
56,334 kb
56,336 kb
56,338 kb
Infected K27 FE
[0 - 10.00]
Control K27 FE
[0 - 10.00]
Infected K4 FE
[0 - 3.00]
Control K4 FE
[0 - 3.00]
Infected PolII FE
[0.100 - 45]
Control PolII FE
[0.100 - 45]
Gene
ACOX1
FBF1

***FBF1***

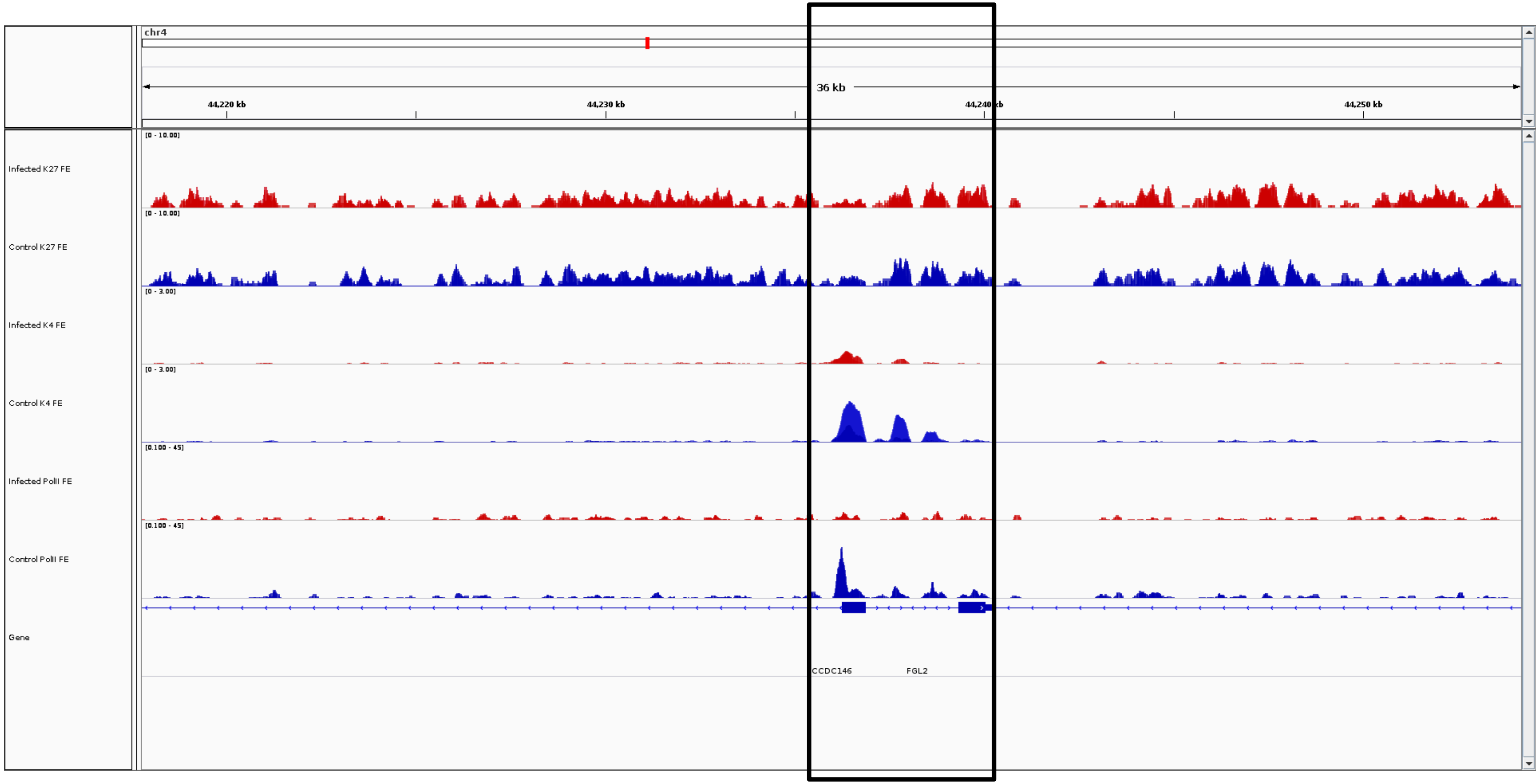
chr4
36 kb
44,220 kb
44,230 kb
44,240 kb
44,250 kb
Infected K27 FE
Control K27 FE
Infected K4 FE
Control K4 FE
Infected PolII FE
Control PolII FE
Gene
CCDC146
FGL2

***FGL2***

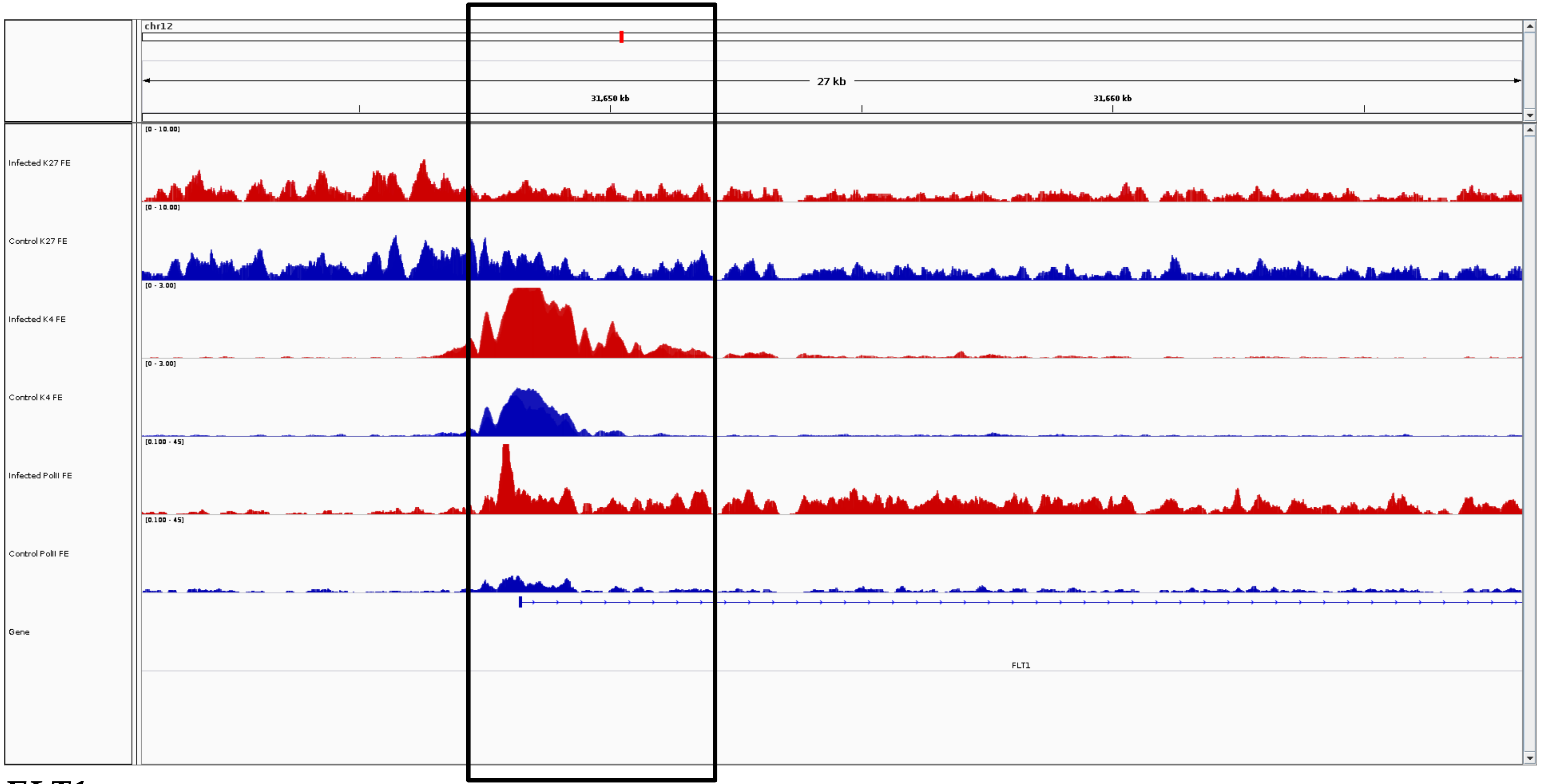
chr12
27 kb
31,650 kb
31,660 kb
[0 - 10.00]
Infected K27 FE
[0 - 10.00]
Control K27 FE
[0 - 3.00]
Infected K4 FE
[0 - 3.00]
Control K4 FE
[0.100 - 45]
Infected PolII FE
[0.100 - 45]
Control PolII FE
Gene
FLT1

***FLT1***

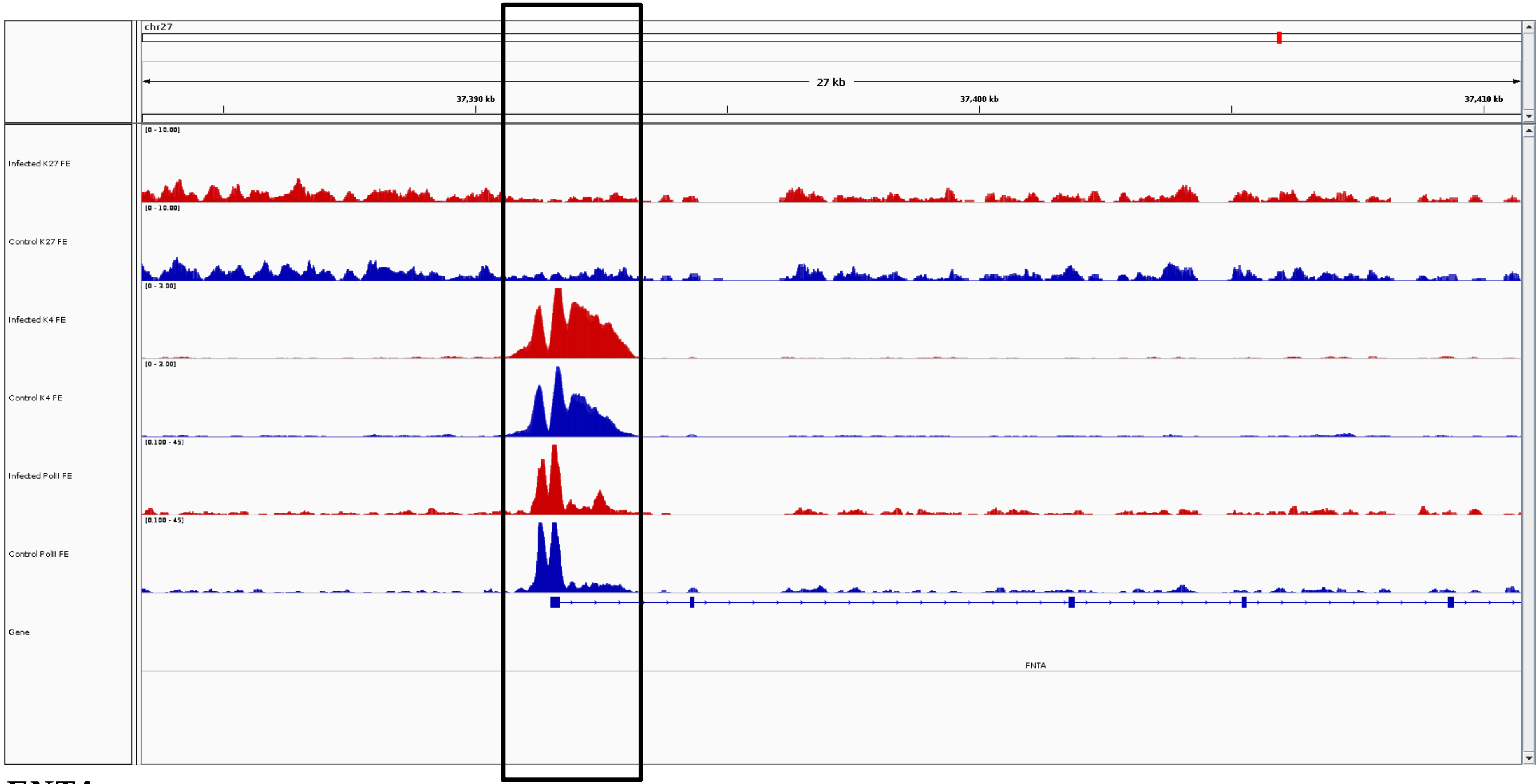
chr27
27 kb
37,390 kb
37,400 kb
37,410 kb
Infected K27 FE
Control K27 FE
Infected K4 FE
Control K4 FE
Infected PolII FE
Control PolII FE
Gene
FNTA

***FNTA***

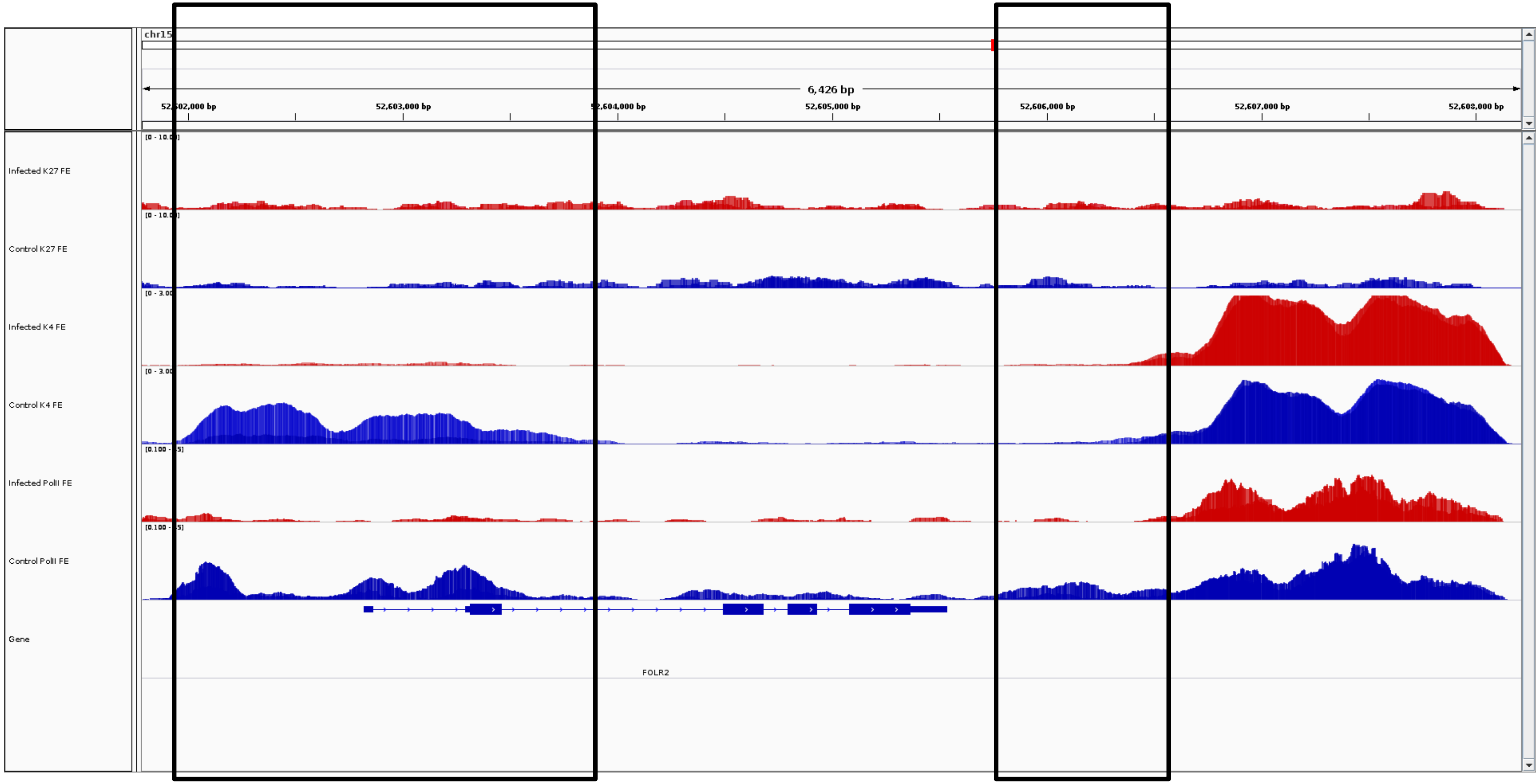

chr15
6,426 bp
52,602,000 bp
52,603,000 bp
52,604,000 bp
52,605,000 bp
52,606,000 bp
52,607,000 bp
52,608,000 bp
Infected K27 FE
Control K27 FE
Infected K4 FE
Control K4 FE
Infected PolII FE
Control PolII FE
Gene
FOLR2


***FOLR2***

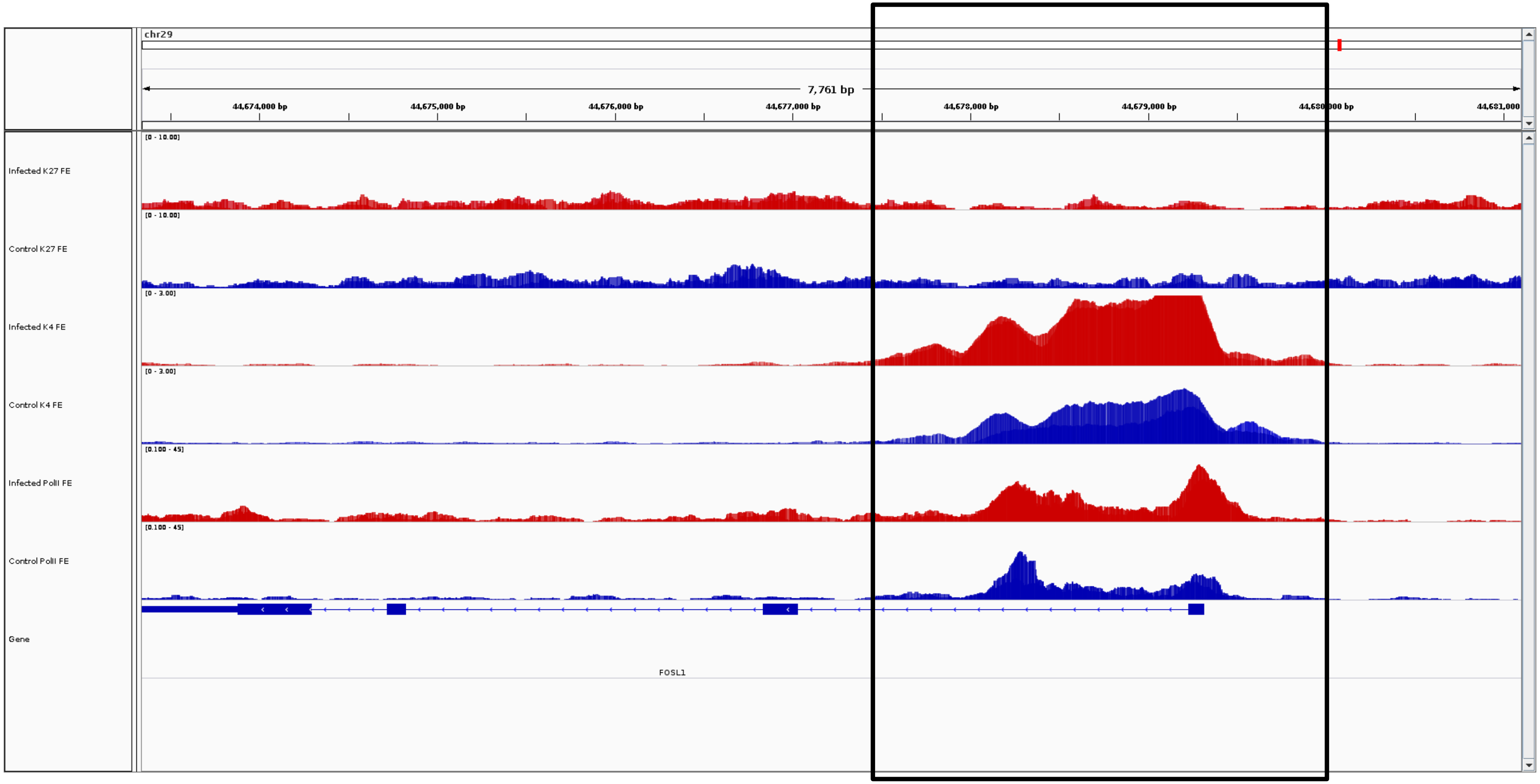
chr29
7,761 bp
44,674,000 bp
44,675,000 bp
44,676,000 bp
44,677,000 bp
44,678,000 bp
44,679,000 bp
44,681,000
Infected K27 FE
Control K27 FE
Infected K4 FE
Control K4 FE
Infected PolII FE
Control PolII FE
Gene
FOSL1

***FOSL1***

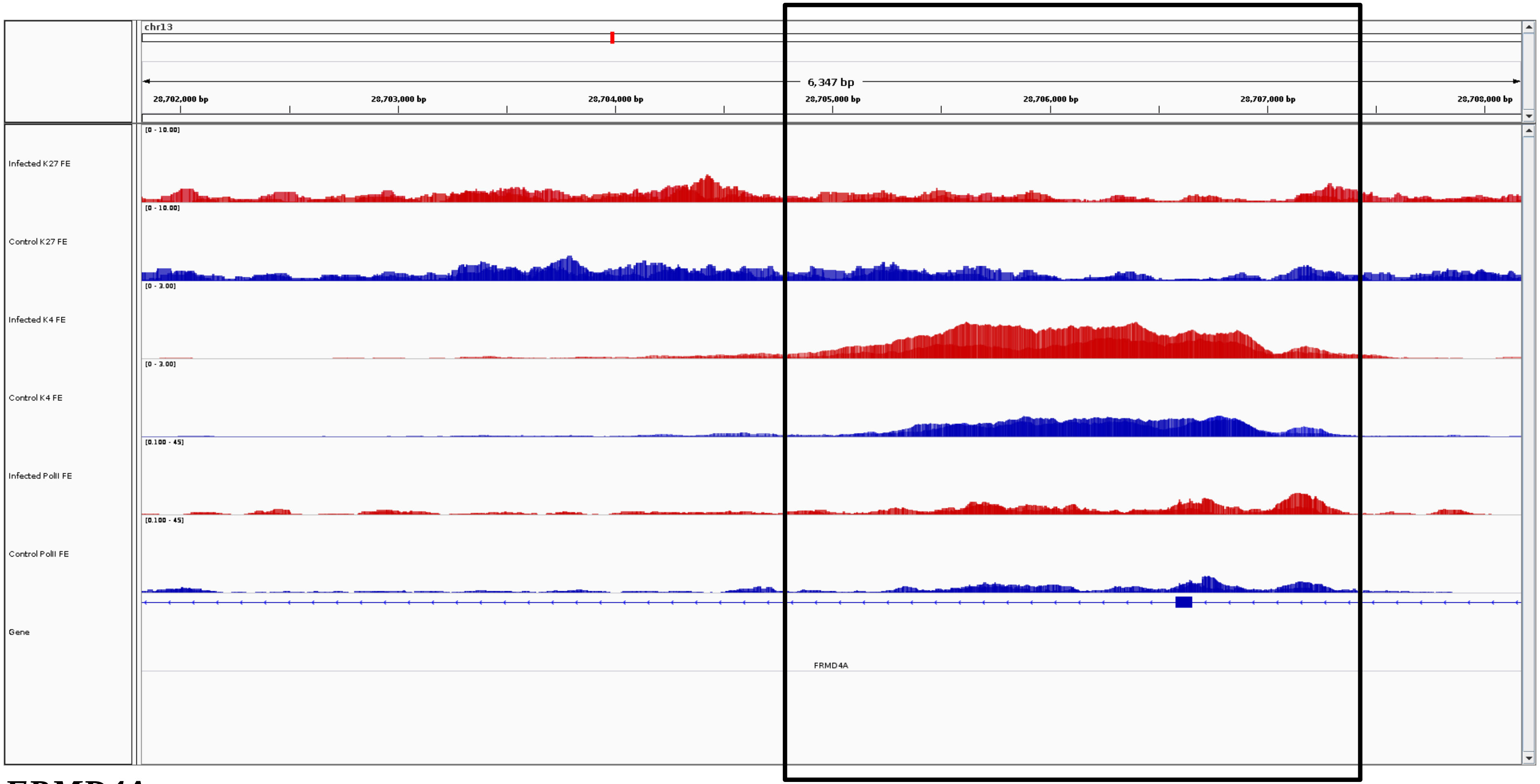
chr13
6,347 bp
28,702,000 bp
28,703,000 bp
28,704,000 bp
28,705,000 bp
28,706,000 bp
28,707,000 bp
28,708,000 bp
Infected K27 FE
[0 - 10.00]
Control K27 FE
[0 - 10.00]
Infected K4 FE
[0 - 3.00]
Control K4 FE
[0 - 3.00]
Infected PolII FE
[0.100 - 45]
Control PolII FE
[0.100 - 45]
Gene
FRMD4A

***FRMD4A***

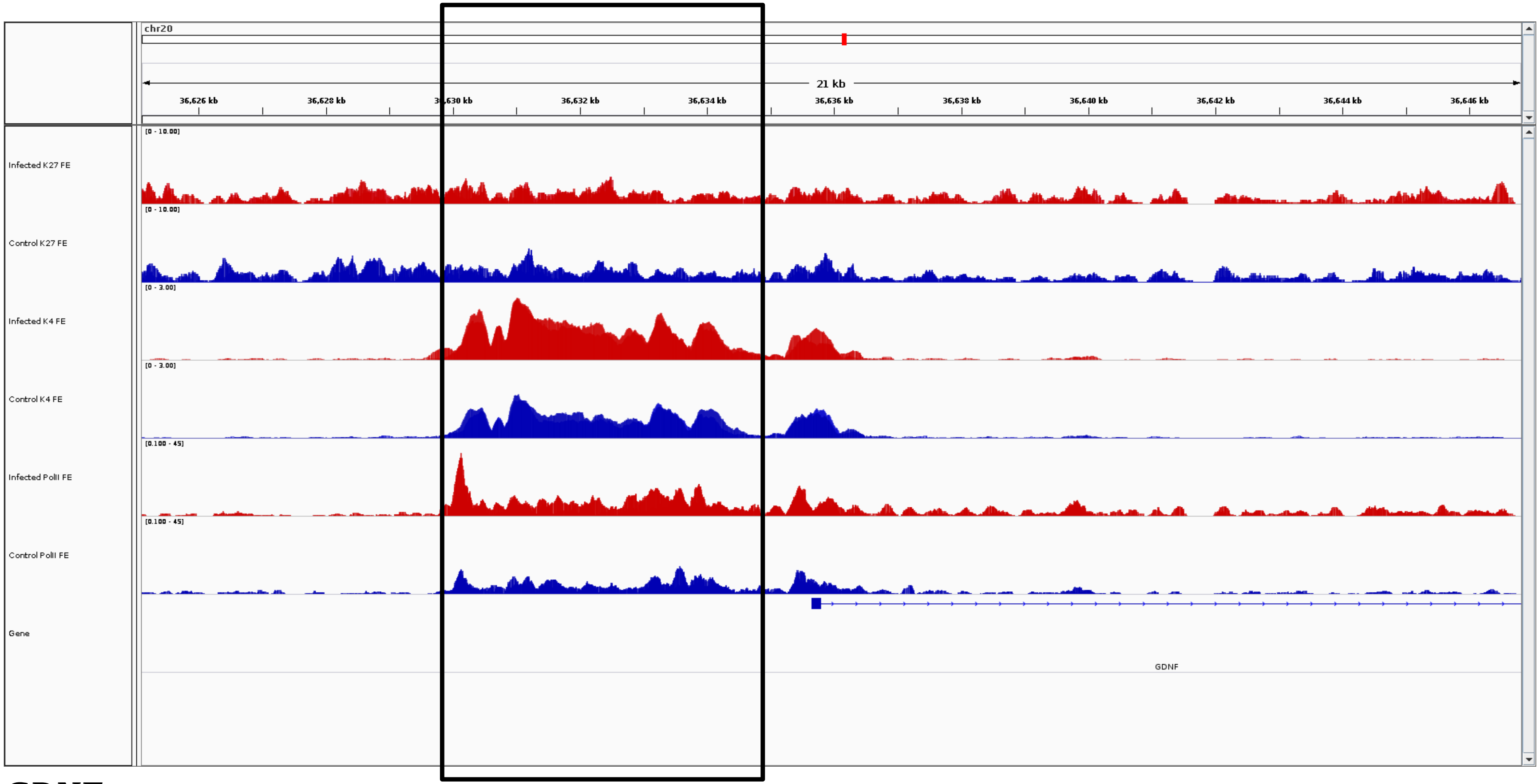
chr20
21 kb
36,626 kb
36,628 kb
36,630 kb
36,632 kb
36,634 kb
36,636 kb
36,638 kb
36,640 kb
36,642 kb
36,644 kb
36,646 kb
Infected K27 FE
Control K27 FE
Infected K4 FE
Control K4 FE
Infected PolII FE
Control PolII FE
Gene
GDNF

***GDNF***

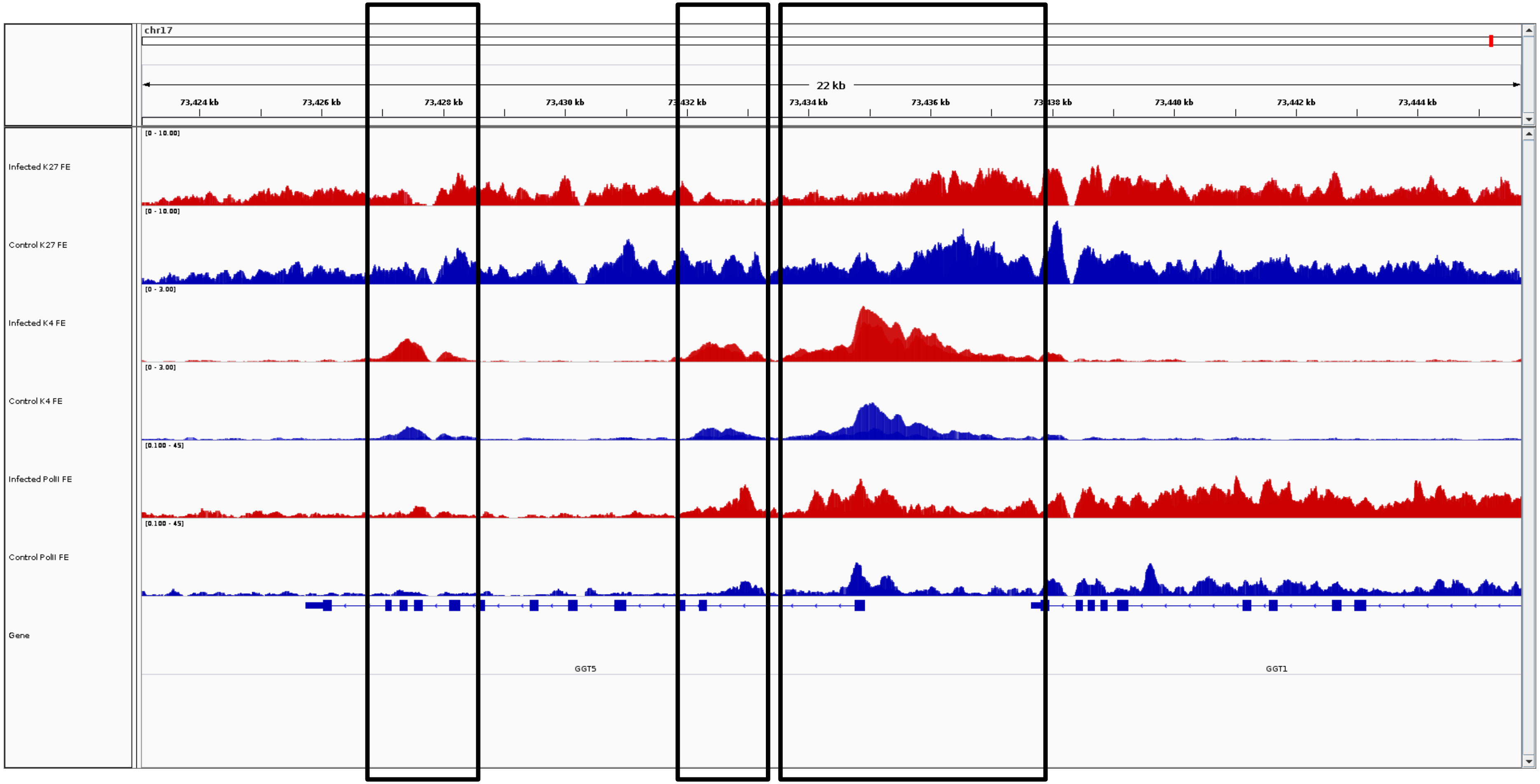
chr17
22 kb
73,424 kb
73,426 kb
73,428 kb
73,430 kb
73,434 kb
73,436 kb
73,440 kb
73,442 kb
73,444 kb
Infected K27 FE
[0 - 10.00]
Control K27 FE
[0 - 10.00]
Infected K4 FE
[0 - 3.00]
Control K4 FE
[0 - 3.00]
Infected PolII FE
[0.100 - 45]
Control PolII FE
[0.100 - 45]
Gene
GGT5
GGT1

***GGT5, GGT1***

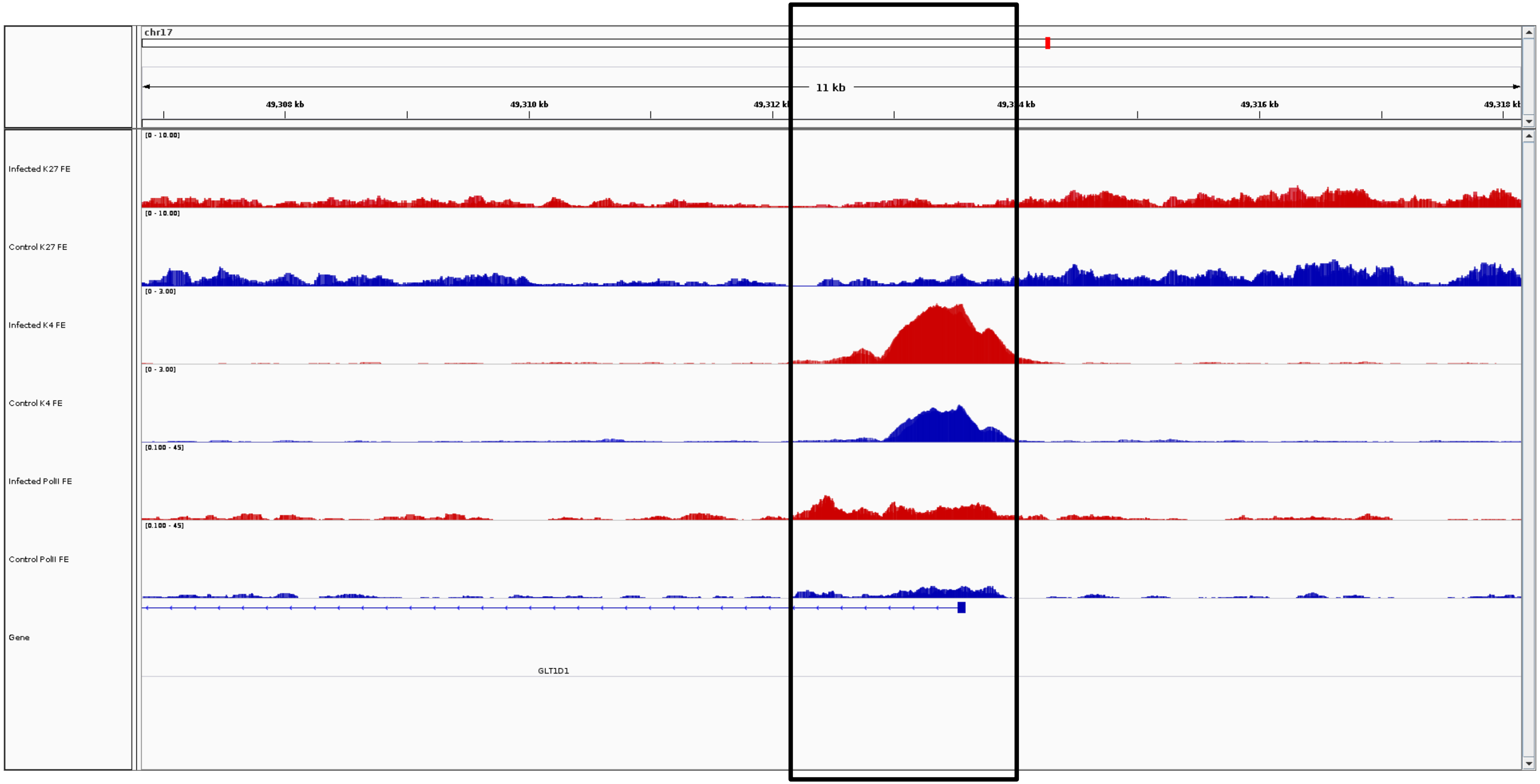
chr17
11 kb
49,308 kb
49,310 kb
49,316 kb
Infected K27 FE
Control K27 FE
Infected K4 FE
Control K4 FE
Infected PolII FE
Control PolII FE
Gene
GLT1D1

***GLT1D1***

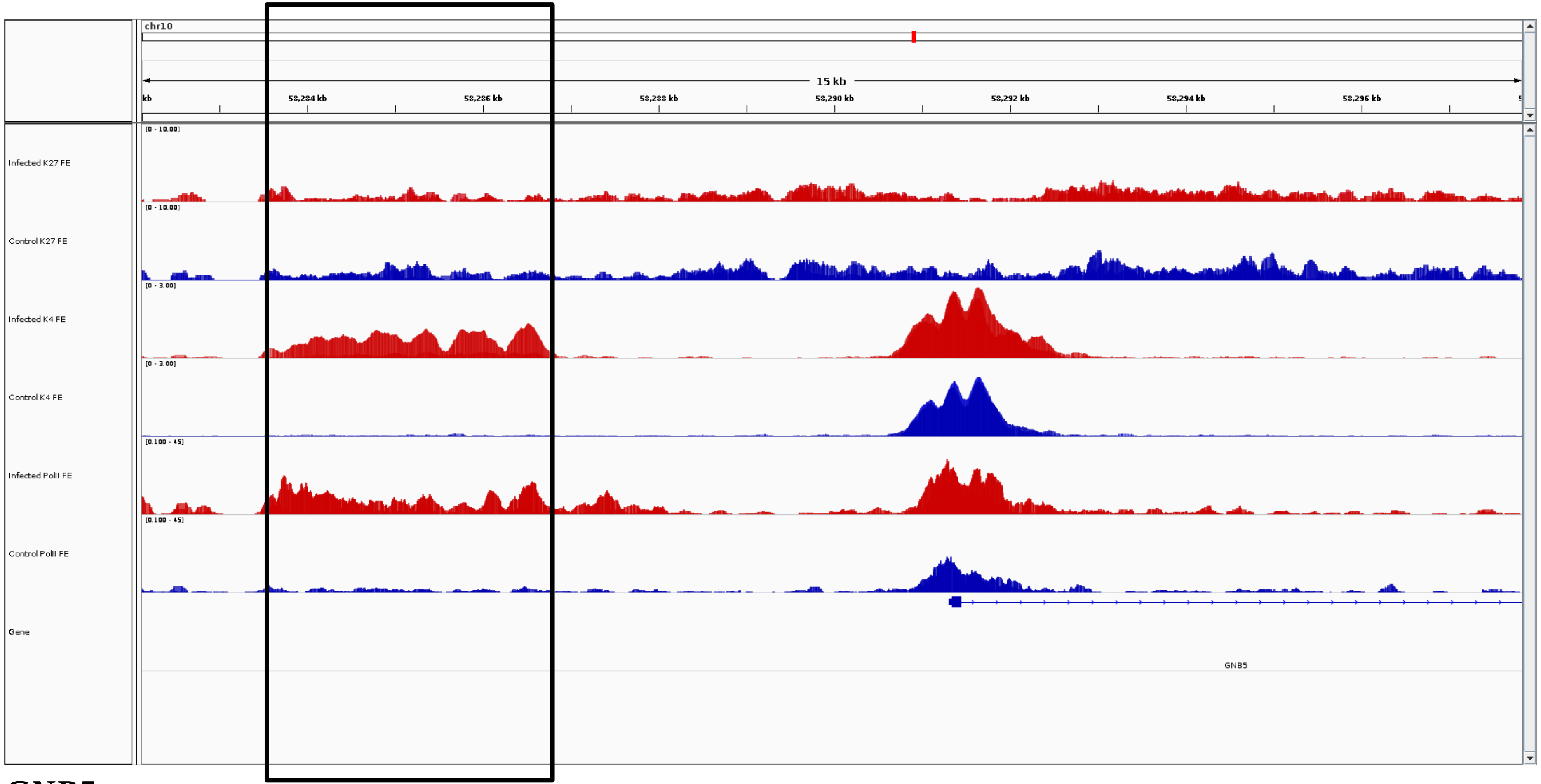
chr10
15 kb
58,284 kb
58,286 kb
58,288 kb
58,290 kb
58,292 kb
58,294 kb
58,296 kb
Infected K27 FE
Control K27 FE
Infected K4 FE
Control K4 FE
Infected PolII FE
Control PolII FE
Gene
GNB5

***GNB5***

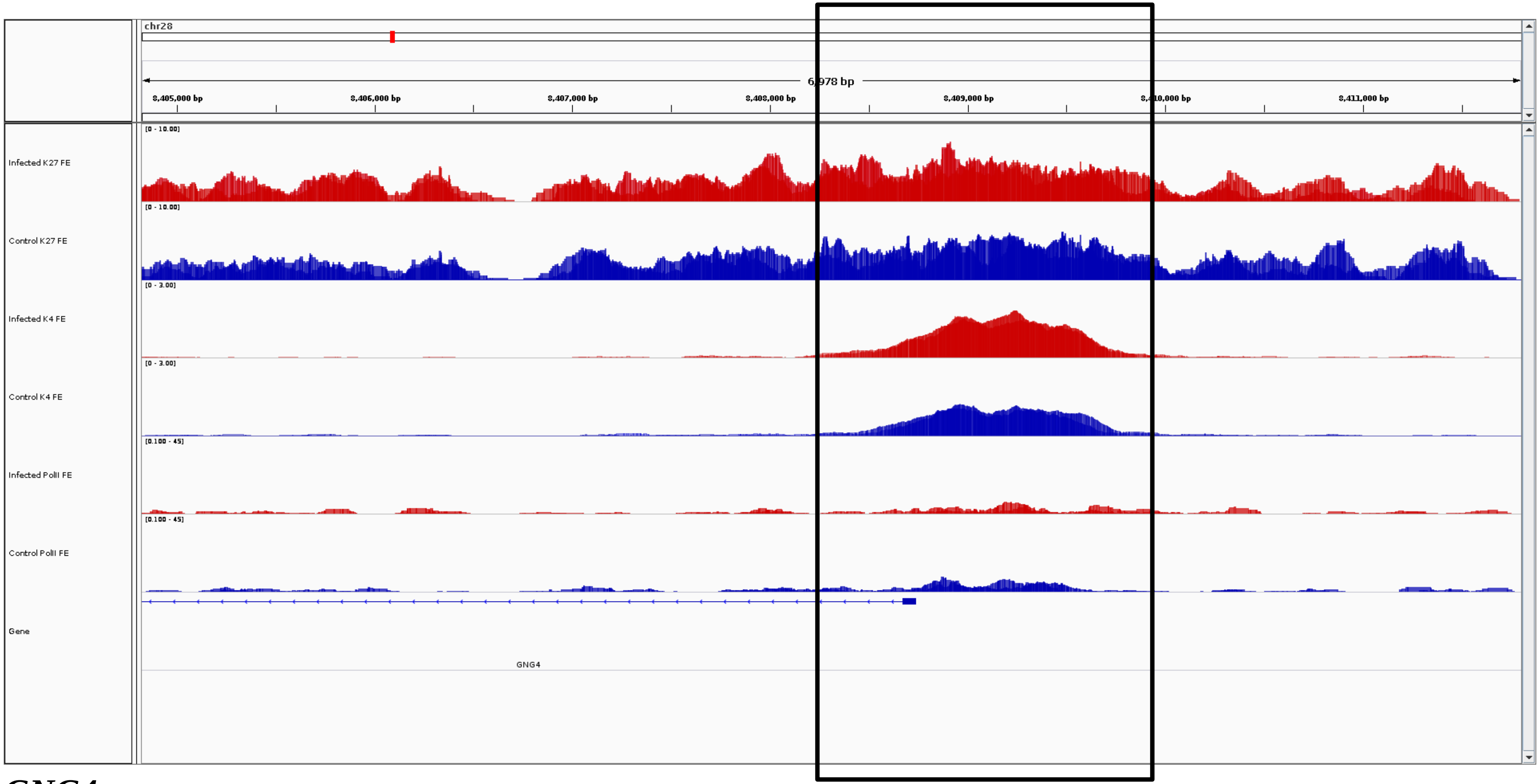
chr28
6,978 bp
8,405,000 bp
8,406,000 bp
8,407,000 bp
8,408,000 bp
8,409,000 bp
8,410,000 bp
8,411,000 bp
Infected K27 FE
Control K27 FE
Infected K4 FE
Control K4 FE
Infected PolII FE
Control PolII FE
Gene
GNG4

***GNG4***

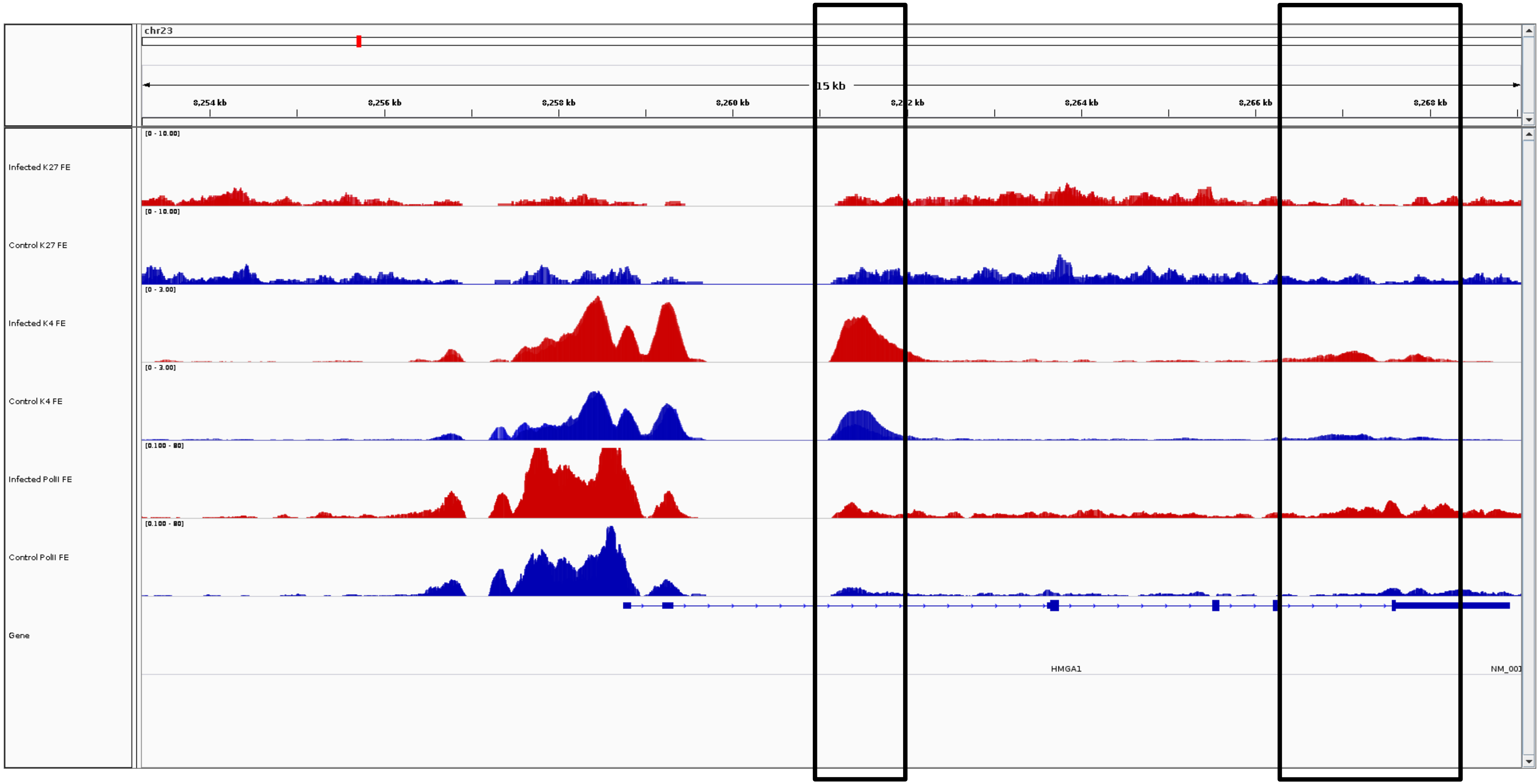
chr23
15 kb
8,254 kb
8,256 kb
8,258 kb
8,260 kb
8,264 kb
8,266 kb
8,268 kb
Infected K27 FE
Control K27 FE
Infected K4 FE
Control K4 FE
Infected PolII FE
Control PolII FE
Gene
HMGA1

***HMGA1***

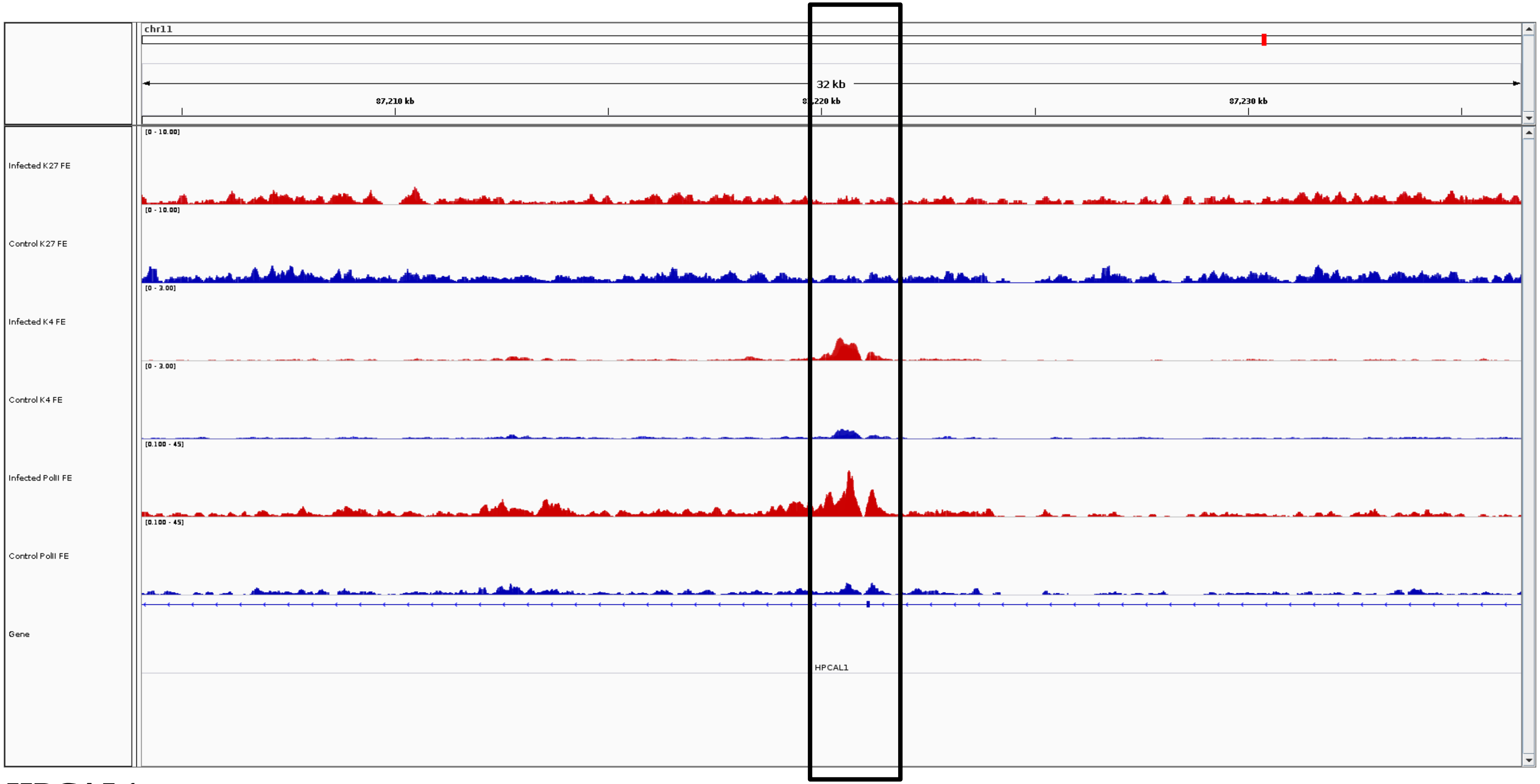
chr11
32 kb
87,210 kb
87,220 kb
87,230 kb
Infected K27 FE
[0 - 10.00]
Control K27 FE
[0 - 10.00]
Infected K4 FE
[0 - 3.00]
Control K4 FE
[0 - 3.00]
Infected PolII FE
[0.100 - 45]
Control PolII FE
[0.100 - 45]
Gene
HPCAL1

***HPCAL1***

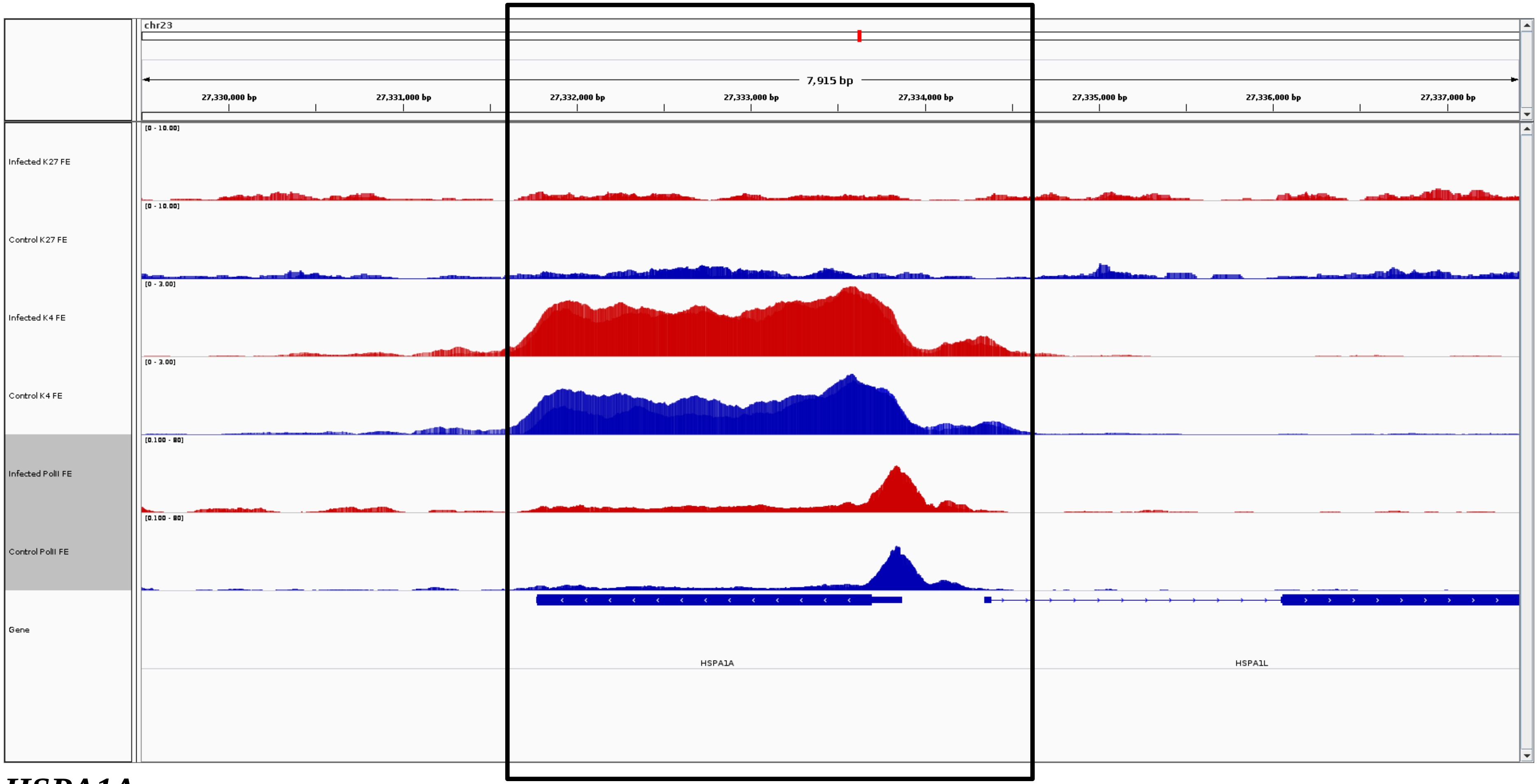
chr23
7,915 bp
27,330,000 bp
27,331,000 bp
27,332,000 bp
27,333,000 bp
27,334,000 bp
27,335,000 bp
27,336,000 bp
27,337,000 bp
Infected K27 FE
[0 - 10.00]
Control K27 FE
[0 - 10.00]
Infected K4 FE
[0 - 3.00]
Control K4 FE
[0 - 3.00]
Infected PolII FE
[0.100 - 80]
Control PolII FE
[0.100 - 80]
Gene
HSPA1A
HSPA1L

***HSPA1A***

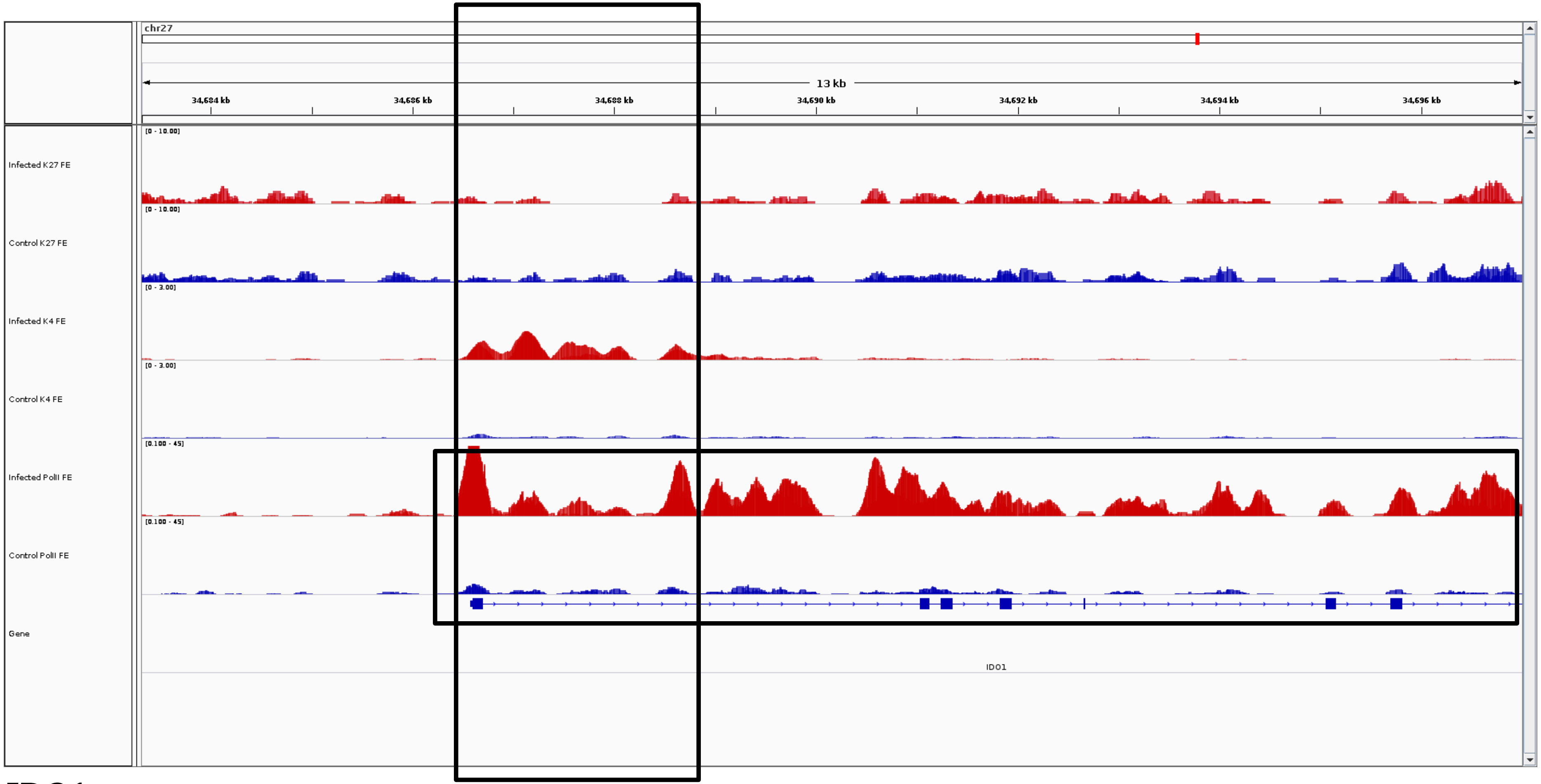
chr27
13 kb
34,684 kb
34,686 kb
34,688 kb
34,690 kb
34,692 kb
34,694 kb
34,696 kb
Infected K27 FE
[0 - 10.00]
Control K27 FE
[0 - 10.00]
Infected K4 FE
[0 - 3.00]
Control K4 FE
[0 - 3.00]
Infected PolII FE
[0.100 - 45]
Control PolII FE
[0.100 - 45]
Gene
IDO1

***IDO1***

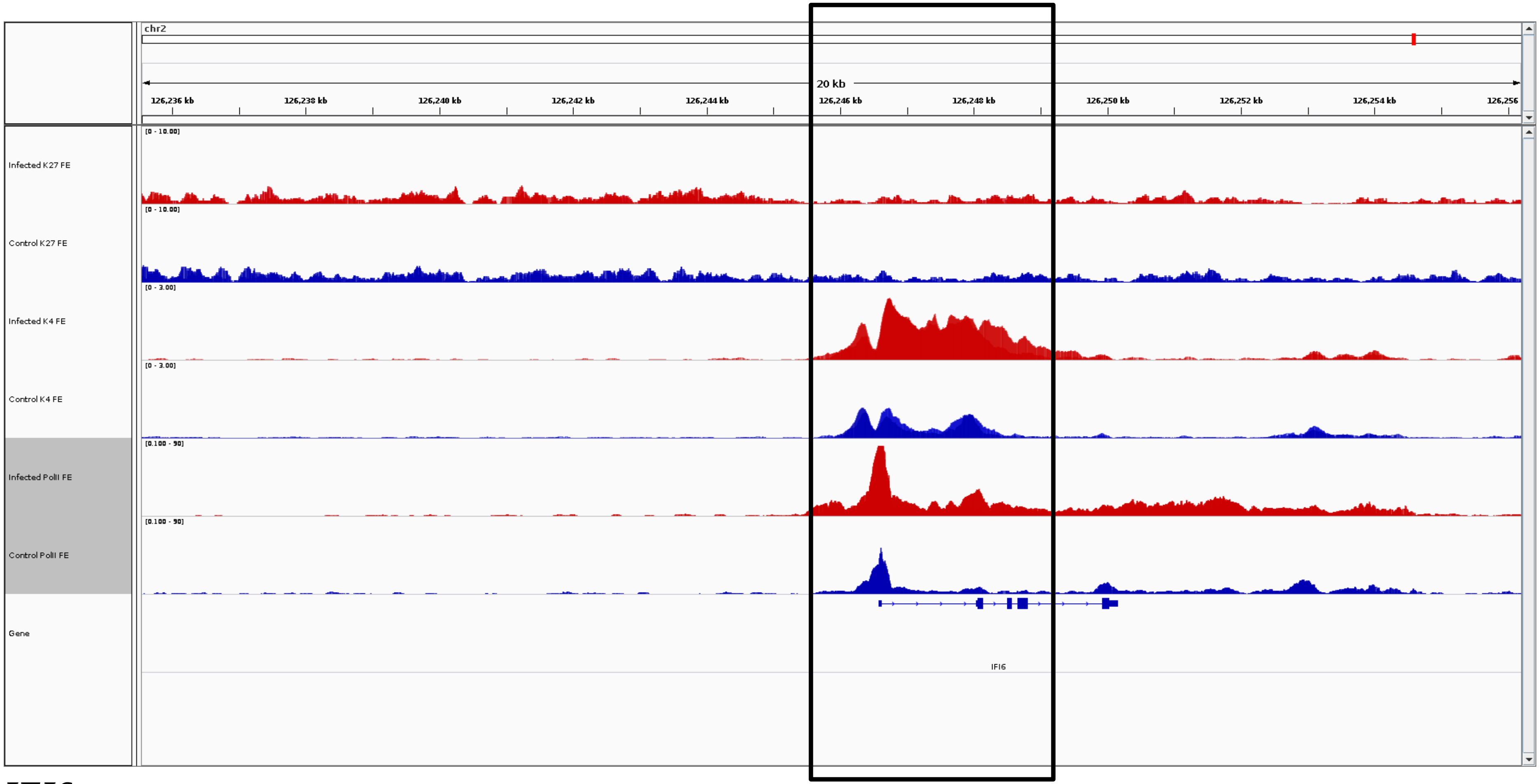
chr2
20 kb
126,236 kb
126,238 kb
126,240 kb
126,242 kb
126,244 kb
126,246 kb
126,248 kb
126,250 kb
126,252 kb
126,254 kb
126,256
Infected K27 FE
Control K27 FE
Infected K4 FE
Control K4 FE
Infected PolII FE
Control PolII FE
Gene
IFI6

***IFI6***

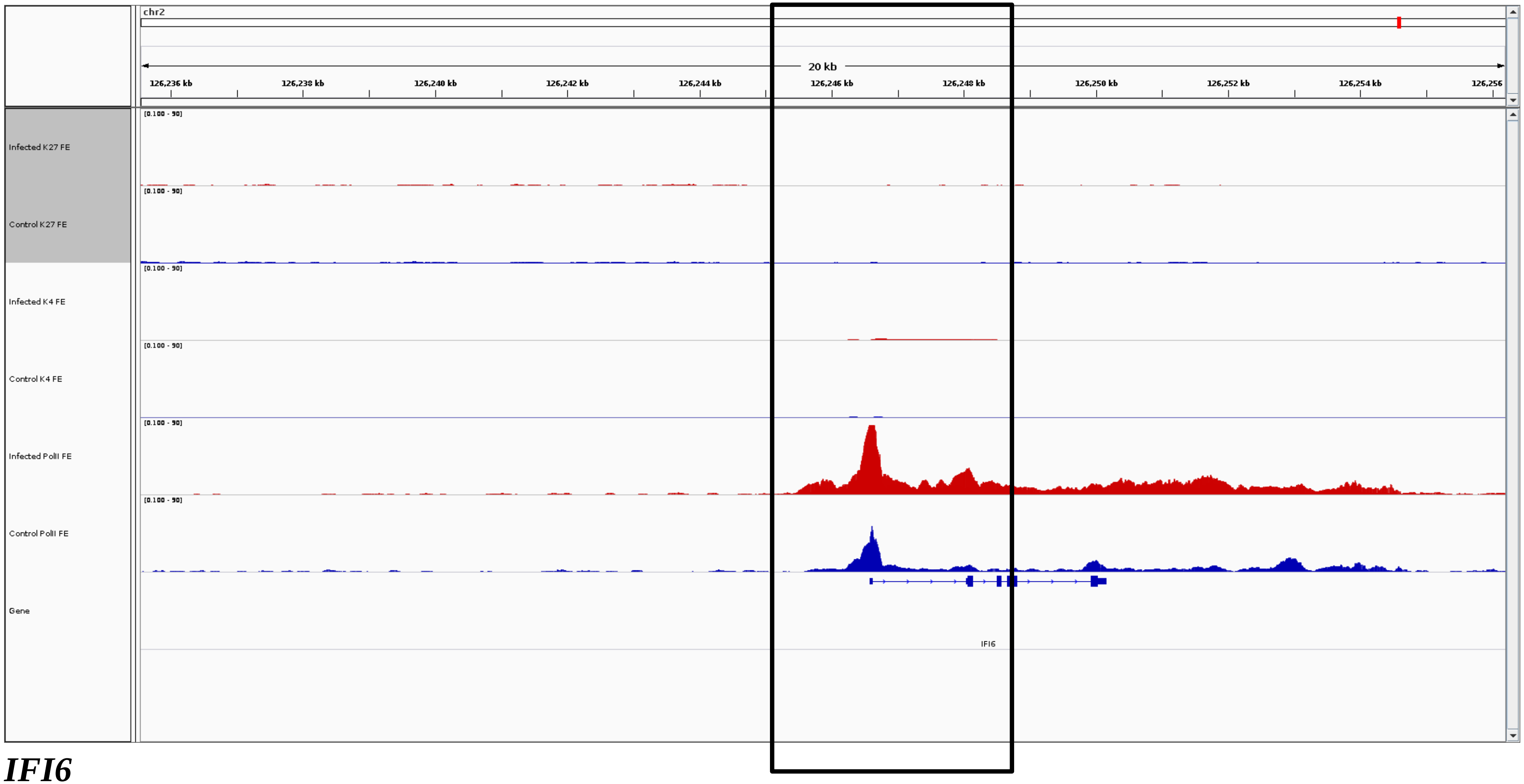
chr2
20 kb
126,236 kb
126,238 kb
126,240 kb
126,242 kb
126,244 kb
126,246 kb
126,248 kb
126,250 kb
126,252 kb
126,254 kb
126,256
Infected K27 FE
Control K27 FE
Infected K4 FE
Control K4 FE
Infected PolII FE
Control PolII FE
Gene
[0.100 - 90]
IFI6

***IFI6***

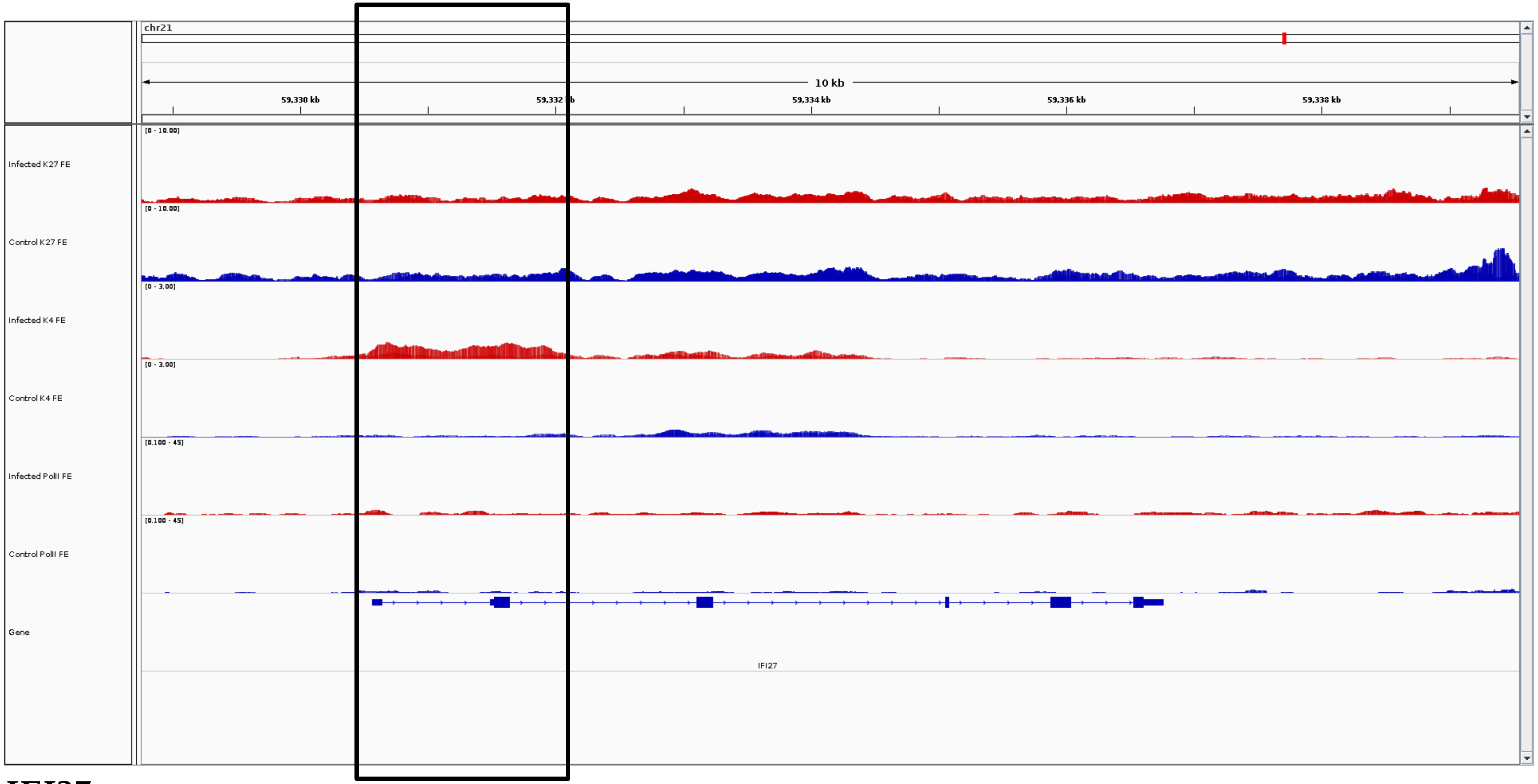
chr21
10 kb
59,330 kb
59,332 kb
59,334 kb
59,336 kb
59,338 kb
[0 - 10.00]
Infected K27 FE
[0 - 10.00]
Control K27 FE
[0 - 3.00]
Infected K4 FE
[0 - 3.00]
Control K4 FE
[0.100 - 45]
Infected PolII FE
[0.100 - 45]
Control PolII FE
Gene
IFI27

***IFI27***

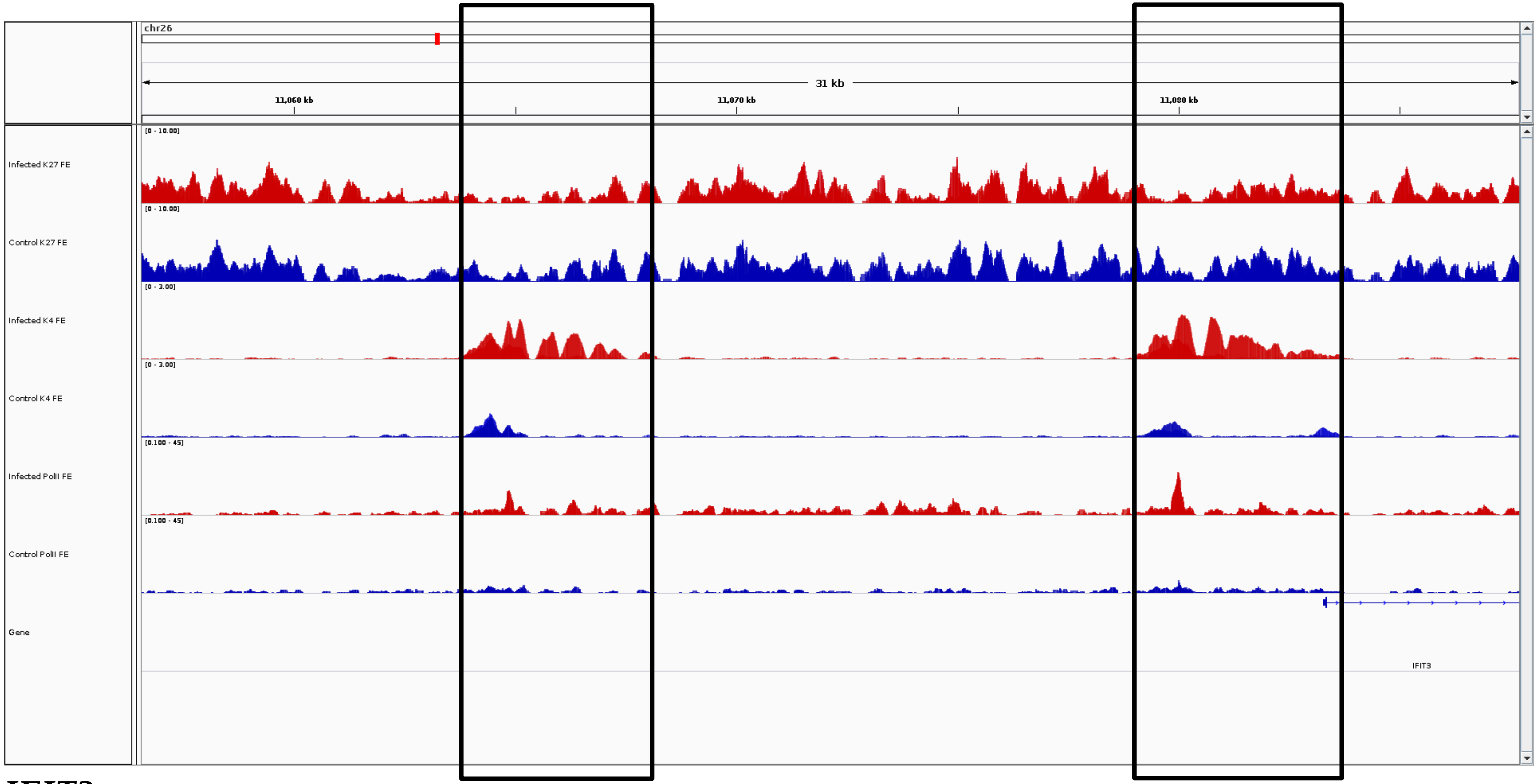
chr26
31 kb
11,060 kb
11,070 kb
11,080 kb
Infected K27 FE
[0 - 10.00]
Control K27 FE
[0 - 10.00]
Infected K4 FE
[0 - 3.00]
Control K4 FE
[0 - 3.00]
Infected PolII FE
[0.100 - 45]
Control PolII FE
[0.100 - 45]
Gene
IFIT3

***IFIT3***

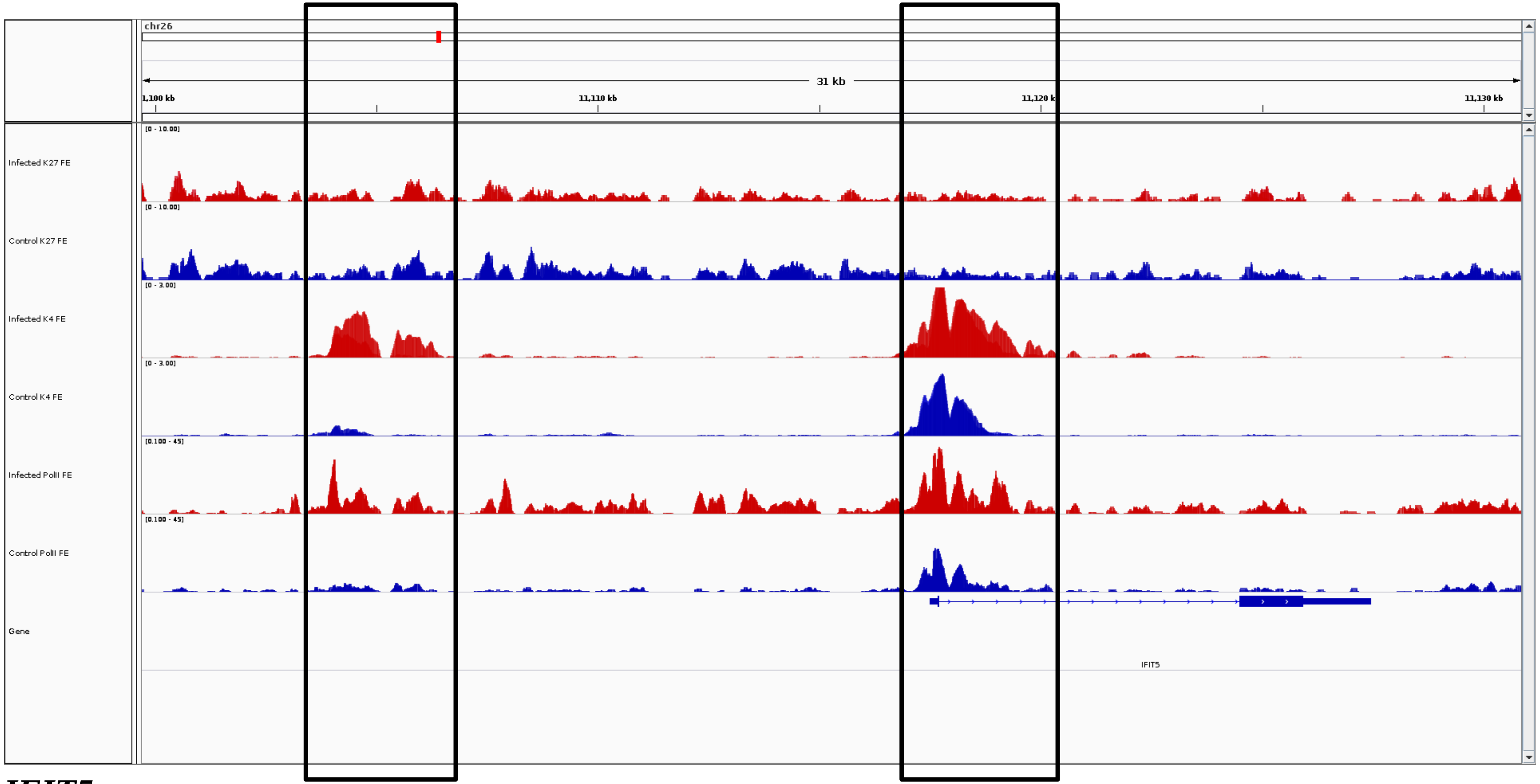
chr26
31 kb
1,100 kb
11,110 kb
11,120 kb
11,130 kb
Infected K27 FE
[0 - 10.00]
Control K27 FE
[0 - 10.00]
Infected K4 FE
[0 - 3.00]
Control K4 FE
[0 - 3.00]
Infected PolII FE
[0.100 - 45]
Control PolII FE
[0.100 - 45]
Gene
IFIT5

***IFIT5***

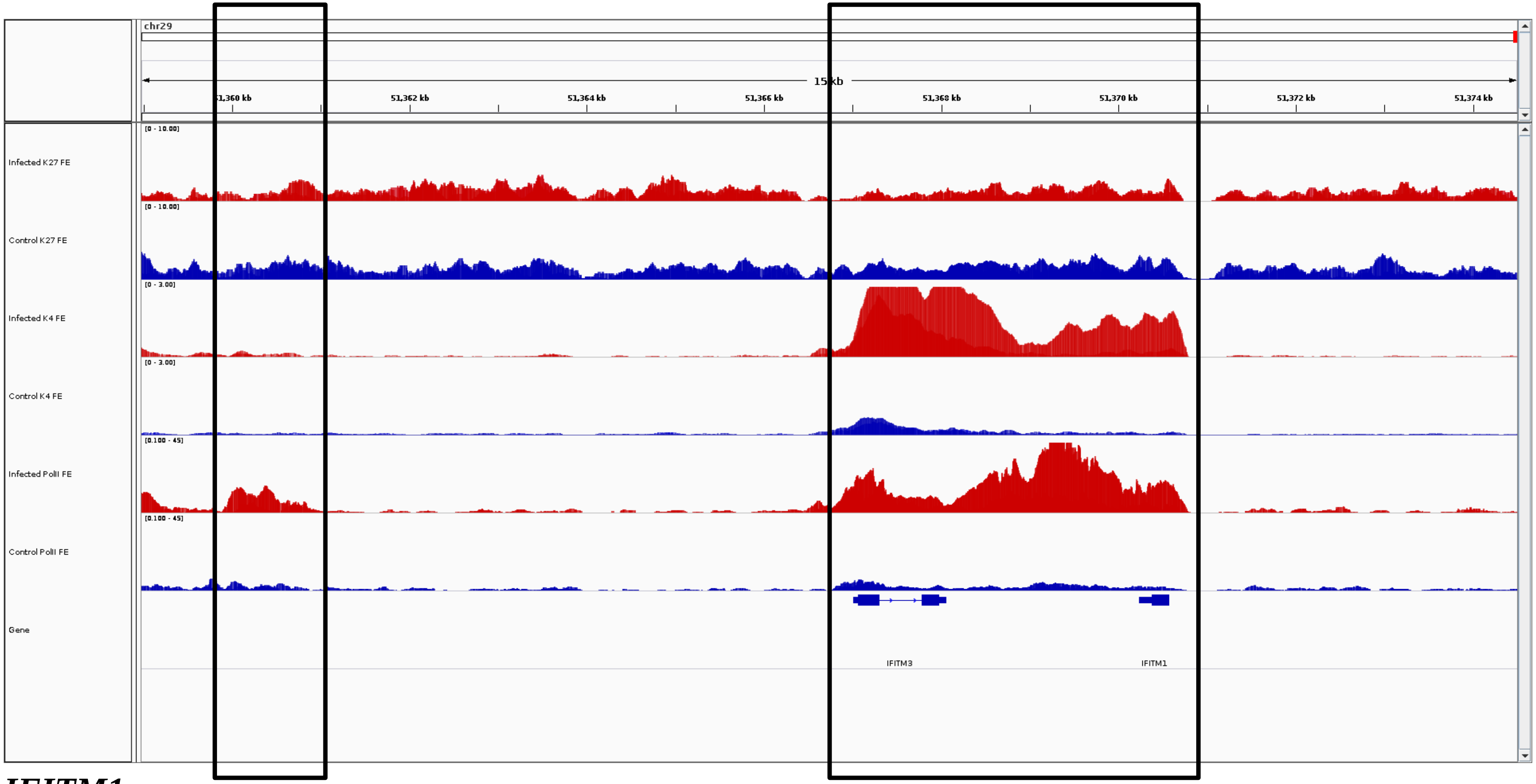
chr29
15 kb
51,360 kb
51,362 kb
51,364 kb
51,366 kb
51,368 kb
51,370 kb
51,372 kb
51,374 kb
Infected K27 FE
Control K27 FE
Infected K4 FE
Control K4 FE
Infected PolII FE
Control PolII FE
Gene
IFITM3
IFITM1

***IFITM1***

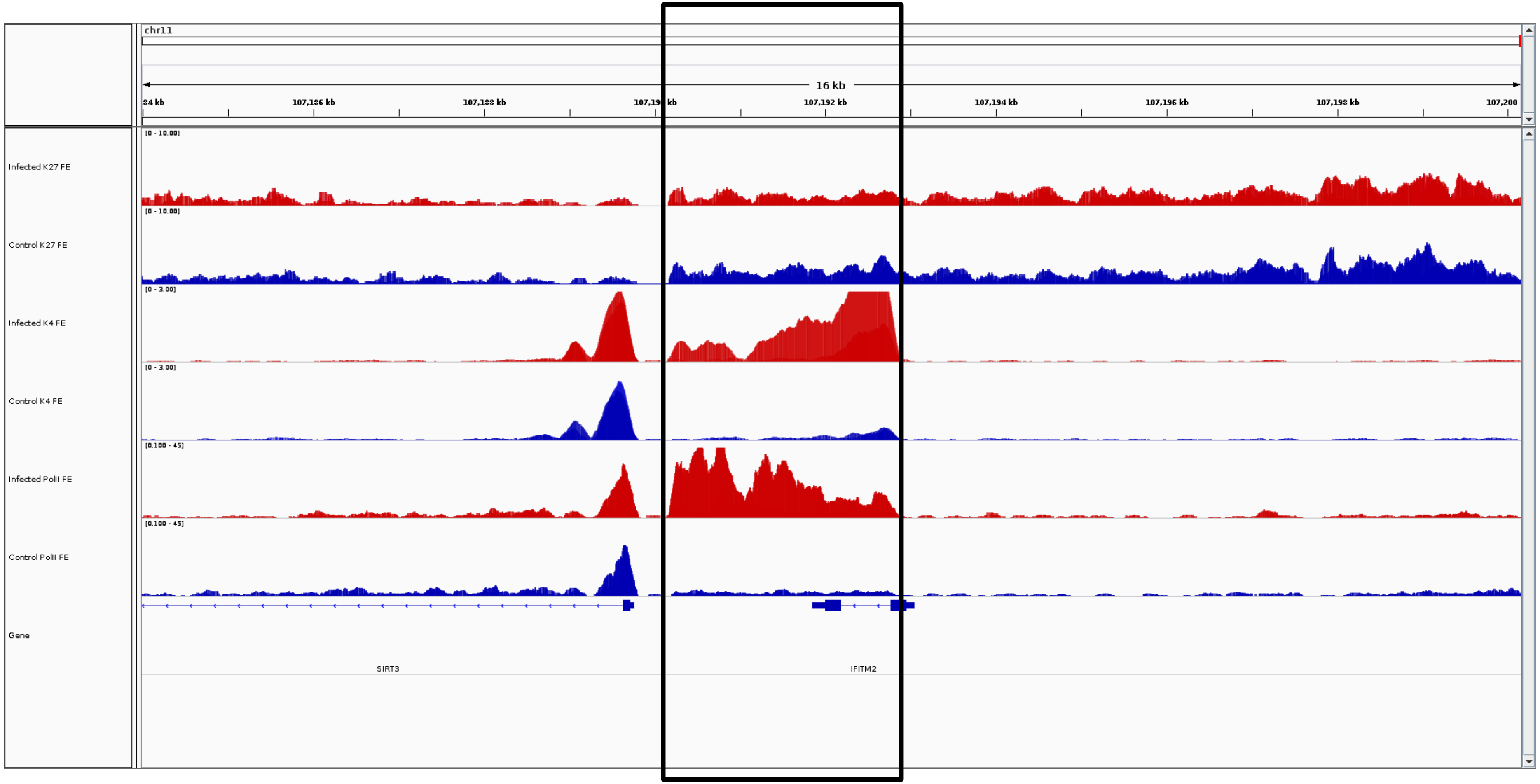
chr11
16 kb
107,186 kb
107,188 kb
107,192 kb
107,194 kb
107,196 kb
107,198 kb
107,200
Infected K27 FE
Control K27 FE
Infected K4 FE
Control K4 FE
Infected PolII FE
Control PolII FE
Gene
SIRT3
IFITM2

***IFITM2***

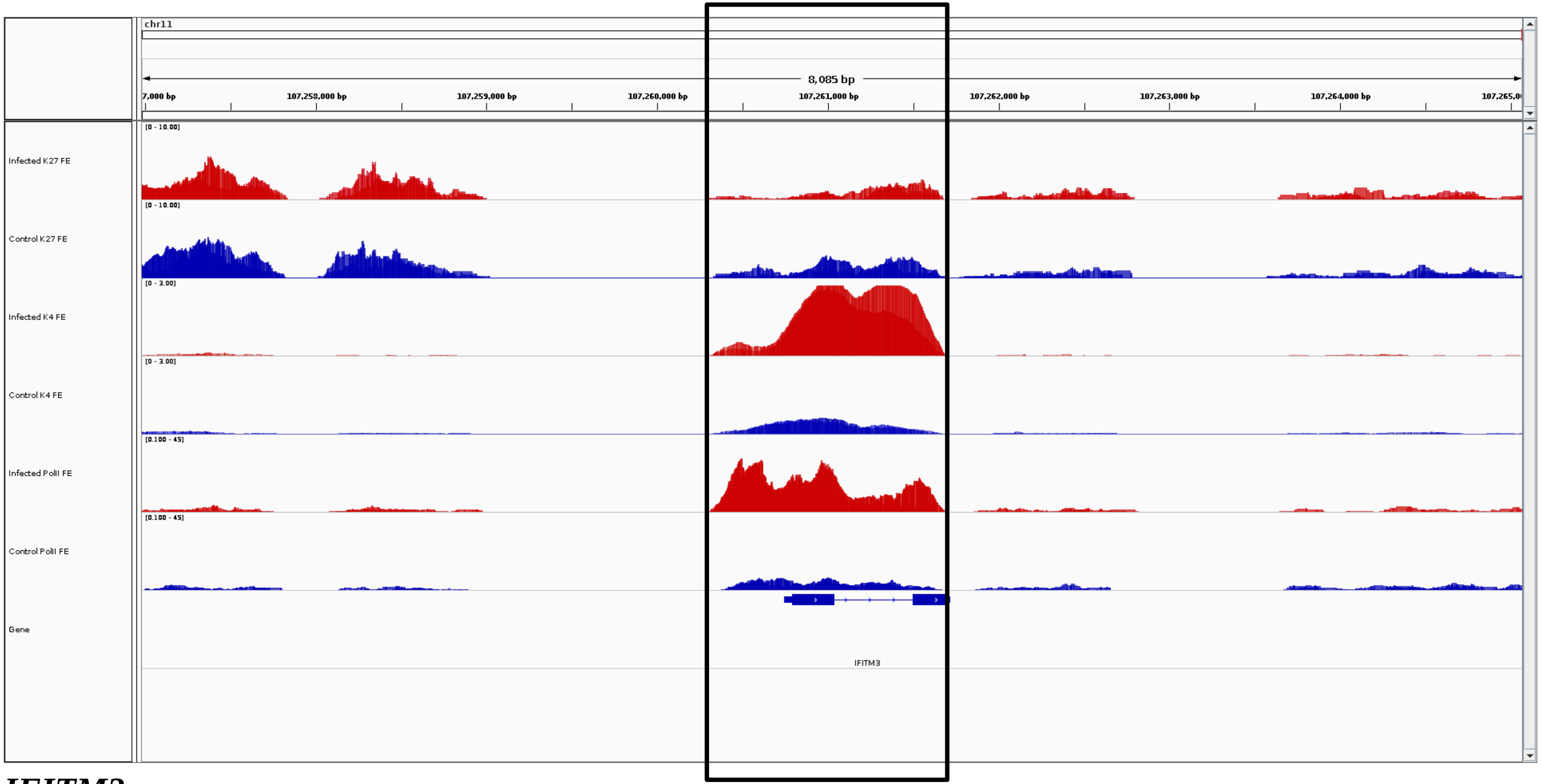
chr11
8,085 bp
7,000 bp
107,258,000 bp
107,259,000 bp
107,260,000 bp
107,261,000 bp
107,262,000 bp
107,263,000 bp
107,264,000 bp
Infected K27 FE
[0 - 10.00]
Control K27 FE
[0 - 10.00]
Infected K4 FE
[0 - 3.00]
Control K4 FE
[0 - 3.00]
Infected PolII FE
[0.100 - 45]
Control PolII FE
[0.100 - 45]
Gene
IFITM3

***IFITM3***

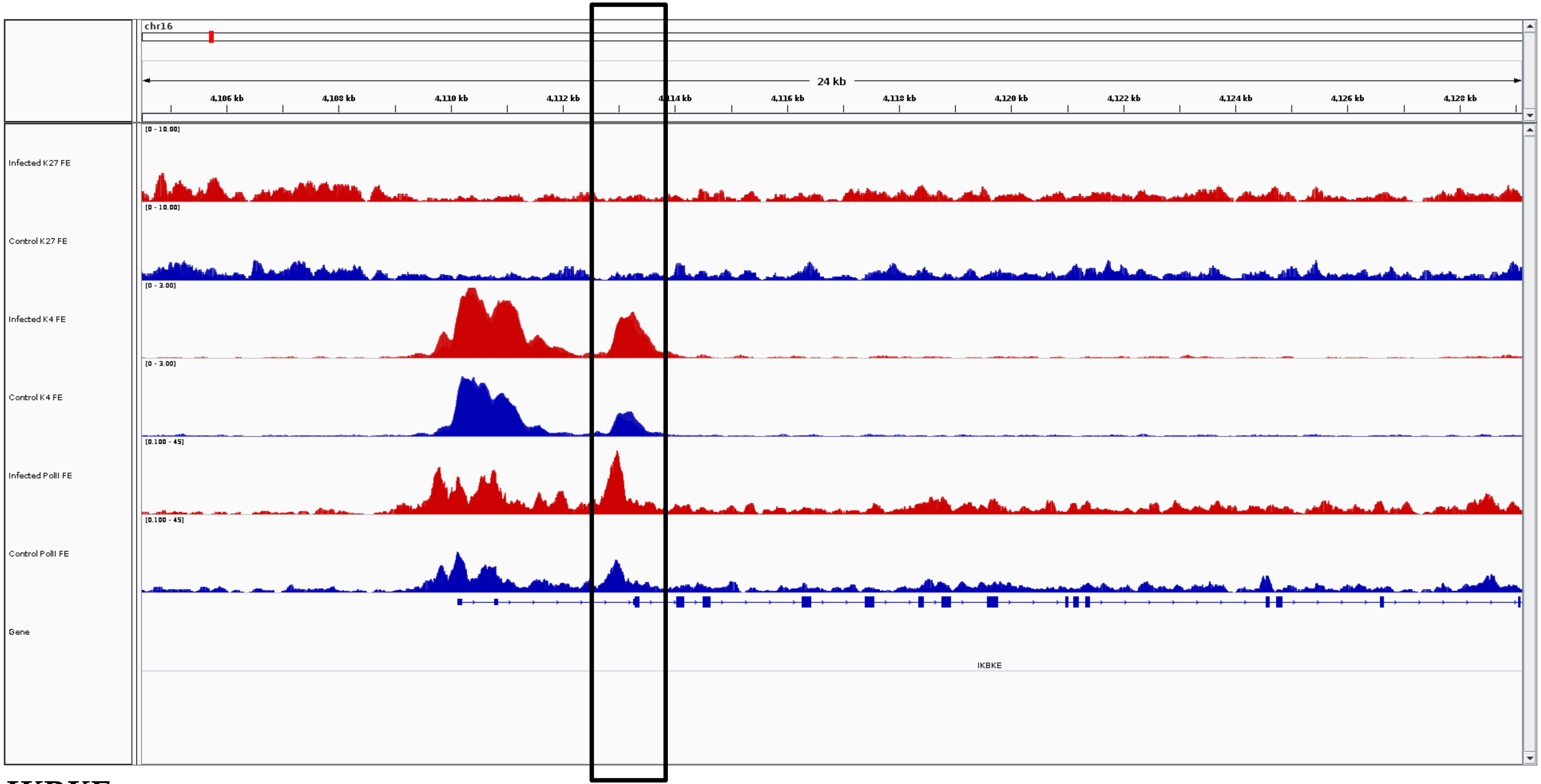
chr16
24 kb
4,106 kb
4,108 kb
4,110 kb
4,112 kb
4,116 kb
4,118 kb
4,120 kb
4,122 kb
4,124 kb
4,126 kb
4,128 kb
Infected K27 FE
Control K27 FE
Infected K4 FE
Control K4 FE
Infected PolII FE
Control PolII FE
Gene
IKBKE

***IKBKE***

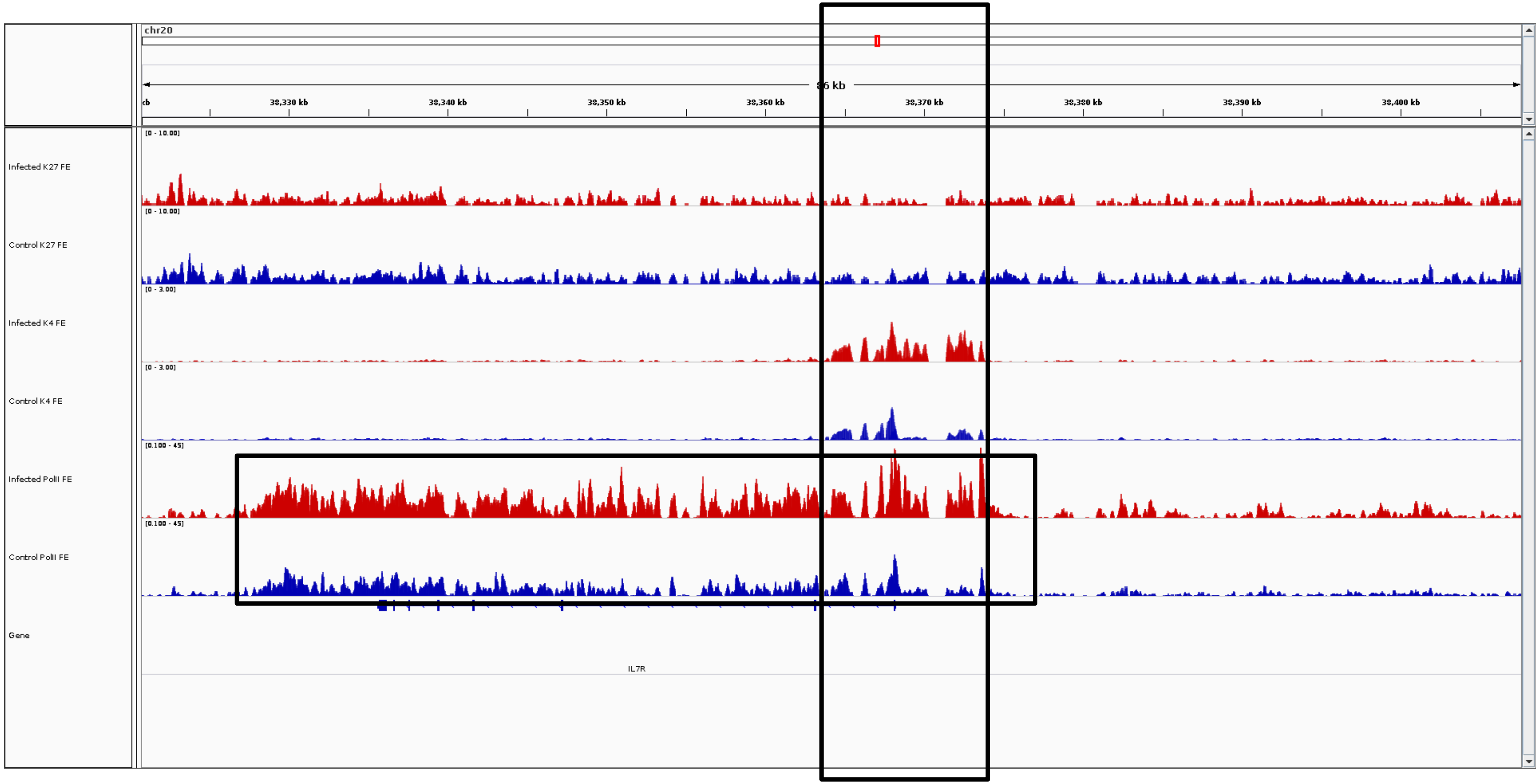
chr20
86 kb
38,330 kb
38,340 kb
38,350 kb
38,360 kb
38,370 kb
38,380 kb
38,390 kb
38,400 kb
Infected K27 FE
Control K27 FE
Infected K4 FE
Control K4 FE
Infected PolII FE
Control PolII FE
Gene
IL7R

***IL7R***

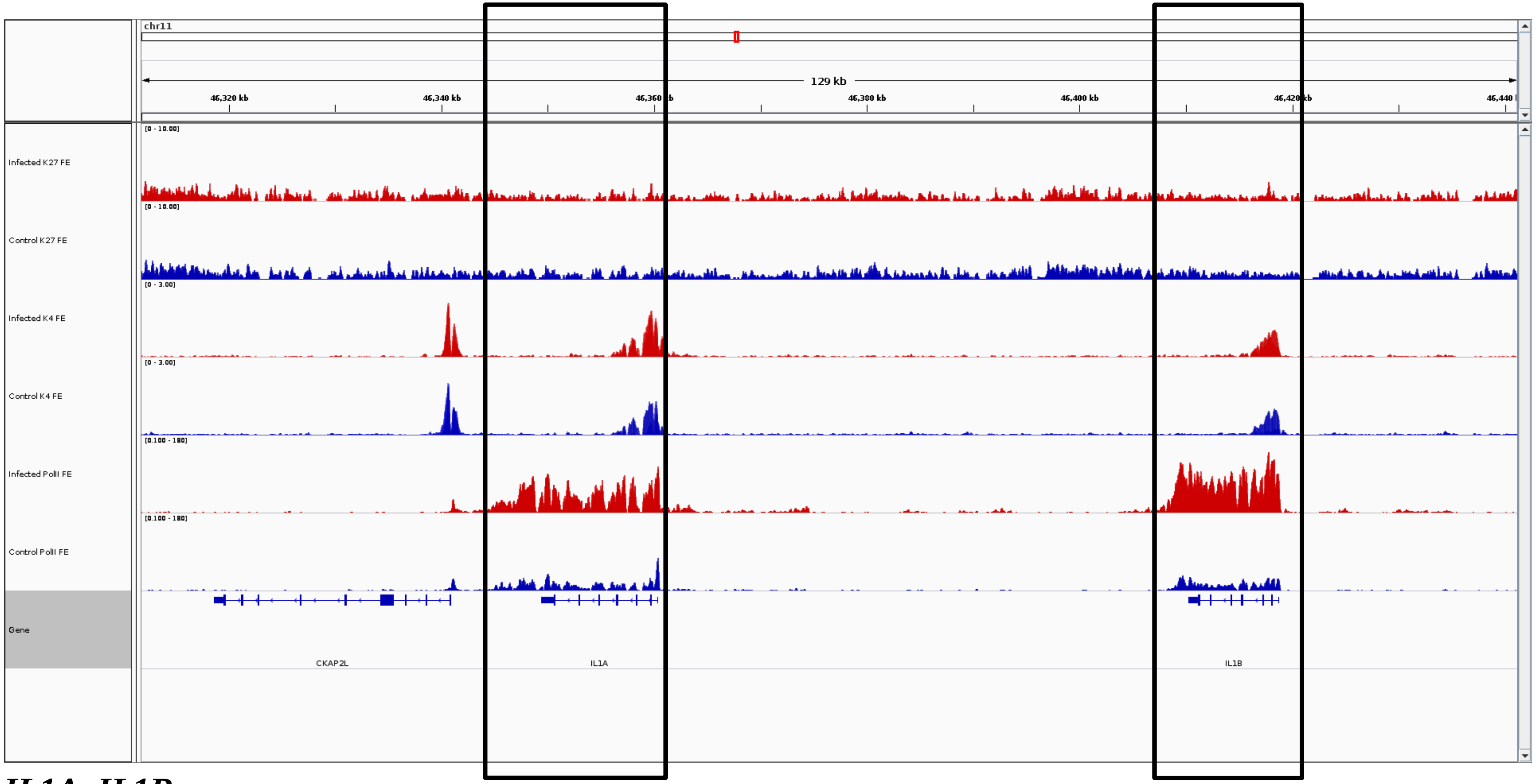
chr11
129 kb
46,320 kb
46,340 kb
46,360 kb
46,380 kb
46,400 kb
46,420 kb
46,440 kb
Infected K27 FE
Control K27 FE
Infected K4 FE
Control K4 FE
Infected PolII FE
Control PolII FE
Gene
CKAP2L
IL1A
IL1B

***IL1A, IL1B***

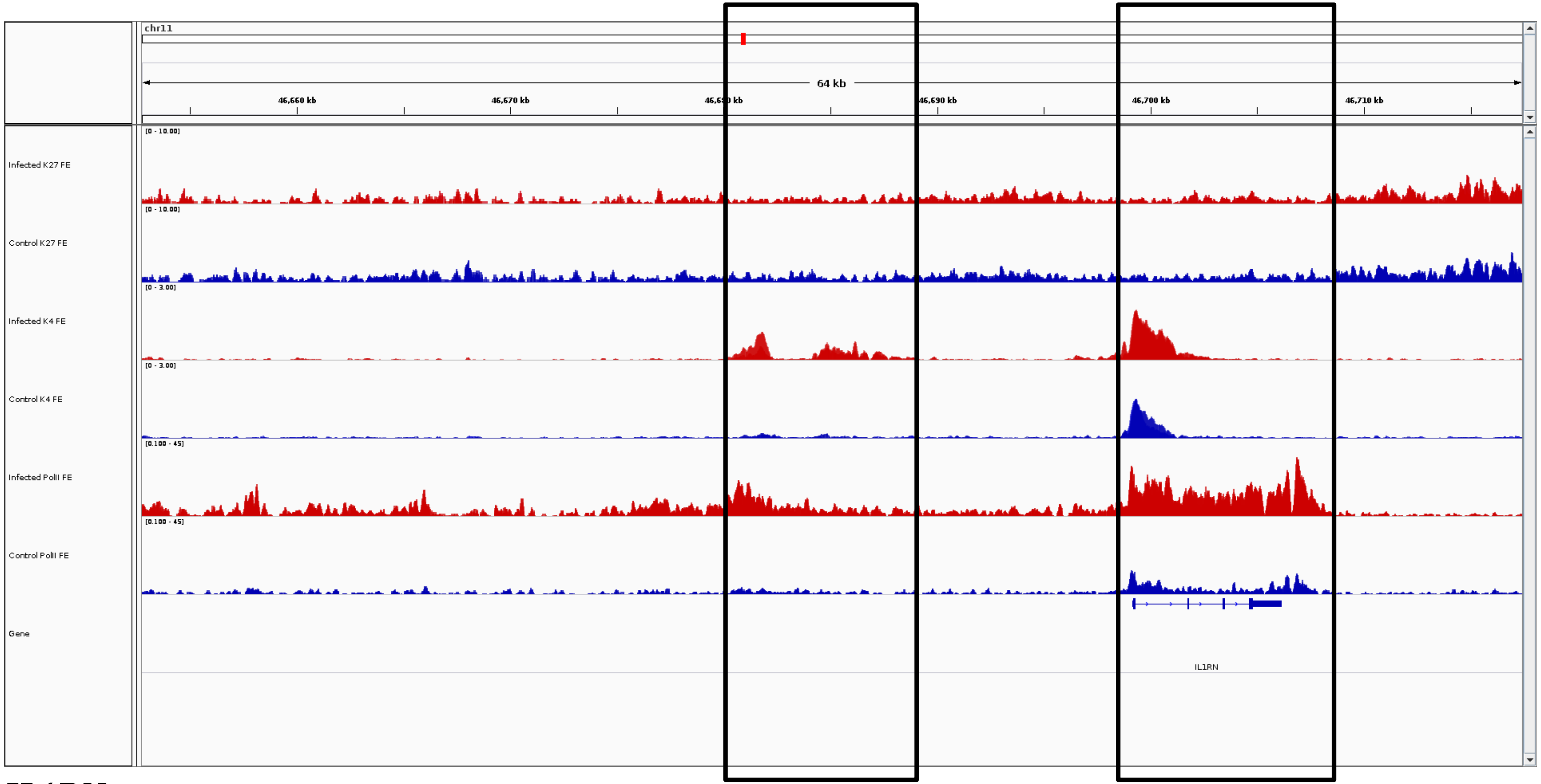
chr11
64 kb
46,660 kb
46,670 kb
46,690 kb
46,700 kb
46,710 kb
Infected K27 FE
Control K27 FE
Infected K4 FE
Control K4 FE
Infected PolII FE
Control PolII FE
Gene
IL1RN

***IL1RN***

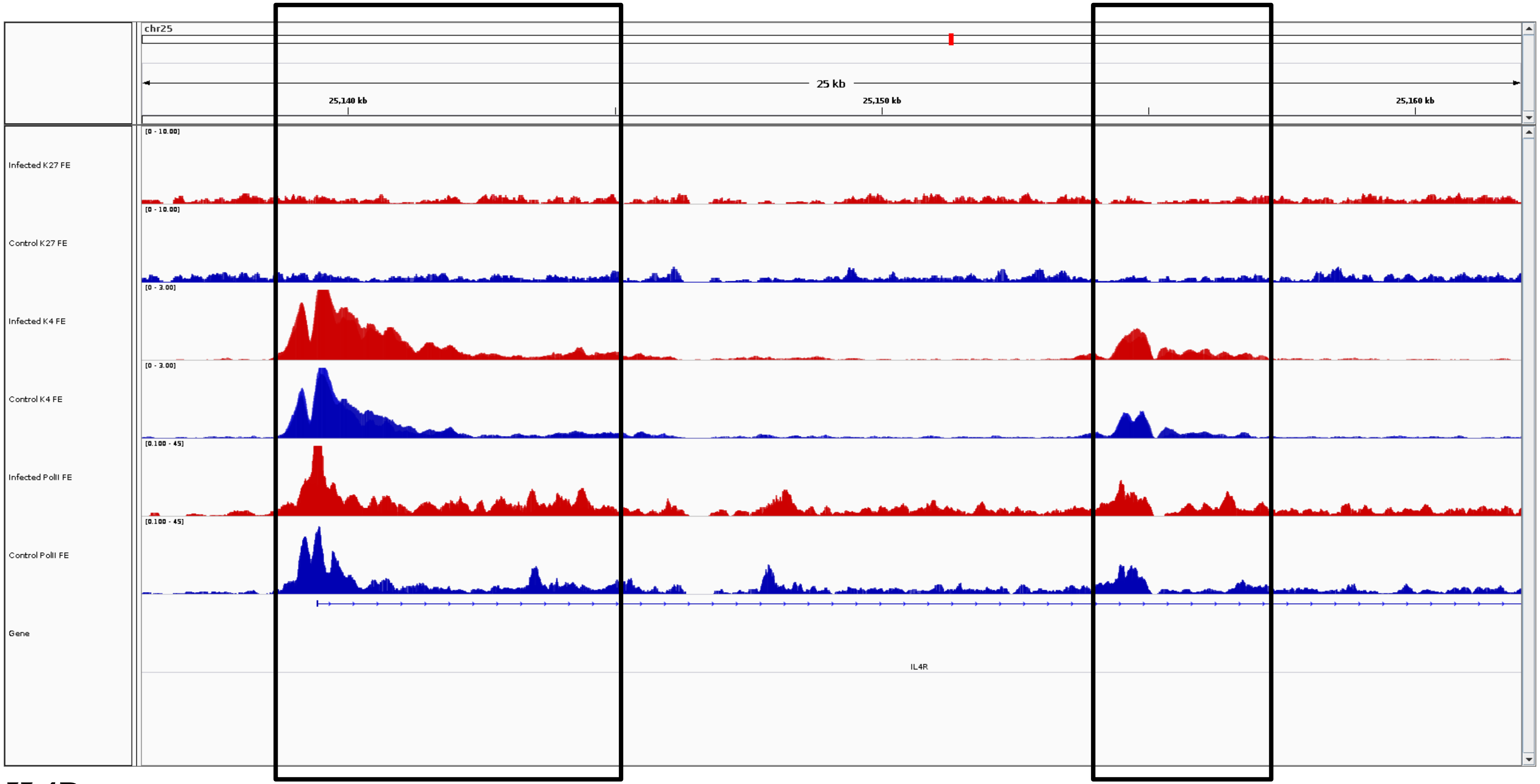
chr25
25 kb
25,140 kb
25,150 kb
25,160 kb
[0 - 10.00]
Infected K27 FE
[0 - 10.00]
Control K27 FE
[0 - 3.00]
Infected K4 FE
[0 - 3.00]
Control K4 FE
[0.100 - 45]
Infected PolII FE
[0.100 - 45]
Control PolII FE
Gene
IL4R

***IL4R***

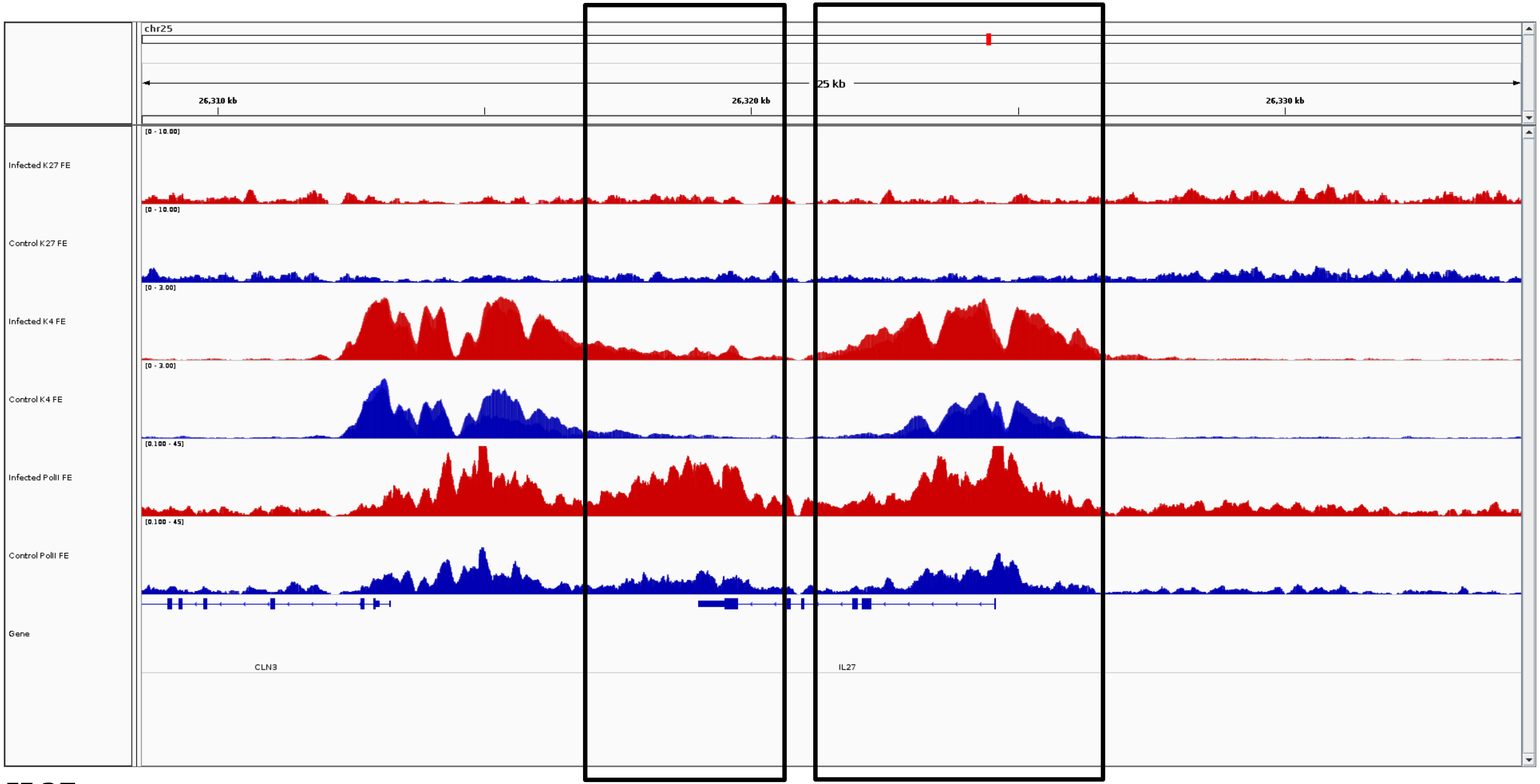
chr25
25 kb
26,310 kb
26,320 kb
26,330 kb
Infected K27 FE
[0 - 10.00]
Control K27 FE
[0 - 10.00]
Infected K4 FE
[0 - 3.00]
Control K4 FE
[0 - 3.00]
Infected PolII FE
[0.100 - 45]
Control PolII FE
[0.100 - 45]
Gene
CLN3
IL27

***IL27***

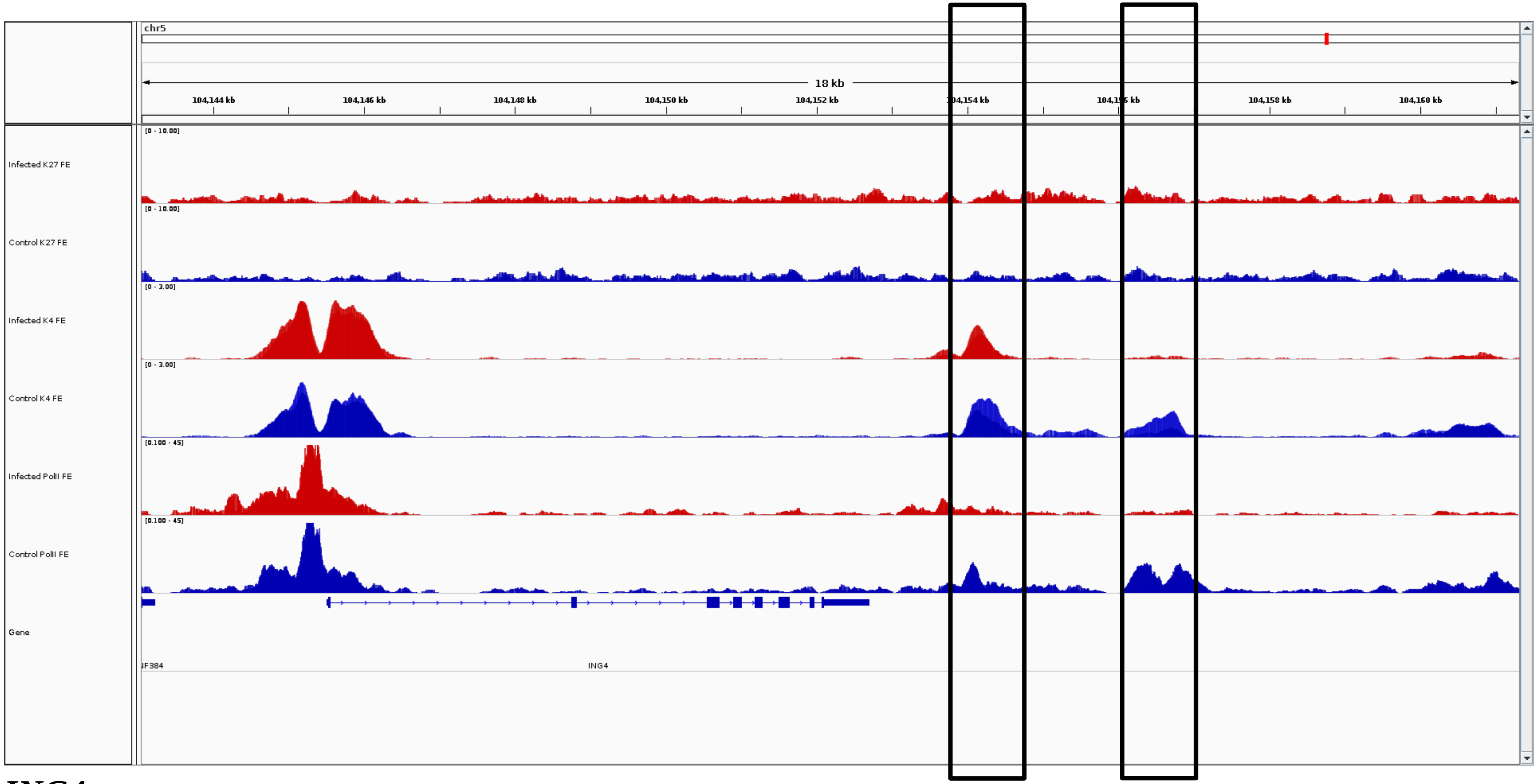
chr5
18 kb
104,144 kb
104,146 kb
104,148 kb
104,150 kb
104,152 kb
104,154 kb
104,158 kb
104,160 kb
Infected K27 FE
[0 - 10.00]
Control K27 FE
[0 - 10.00]
Infected K4 FE
[0 - 3.00]
Control K4 FE
[0 - 3.00]
Infected PolII FE
[0.100 - 45]
Control PolII FE
[0.100 - 45]
Gene
ING4

***ING4***

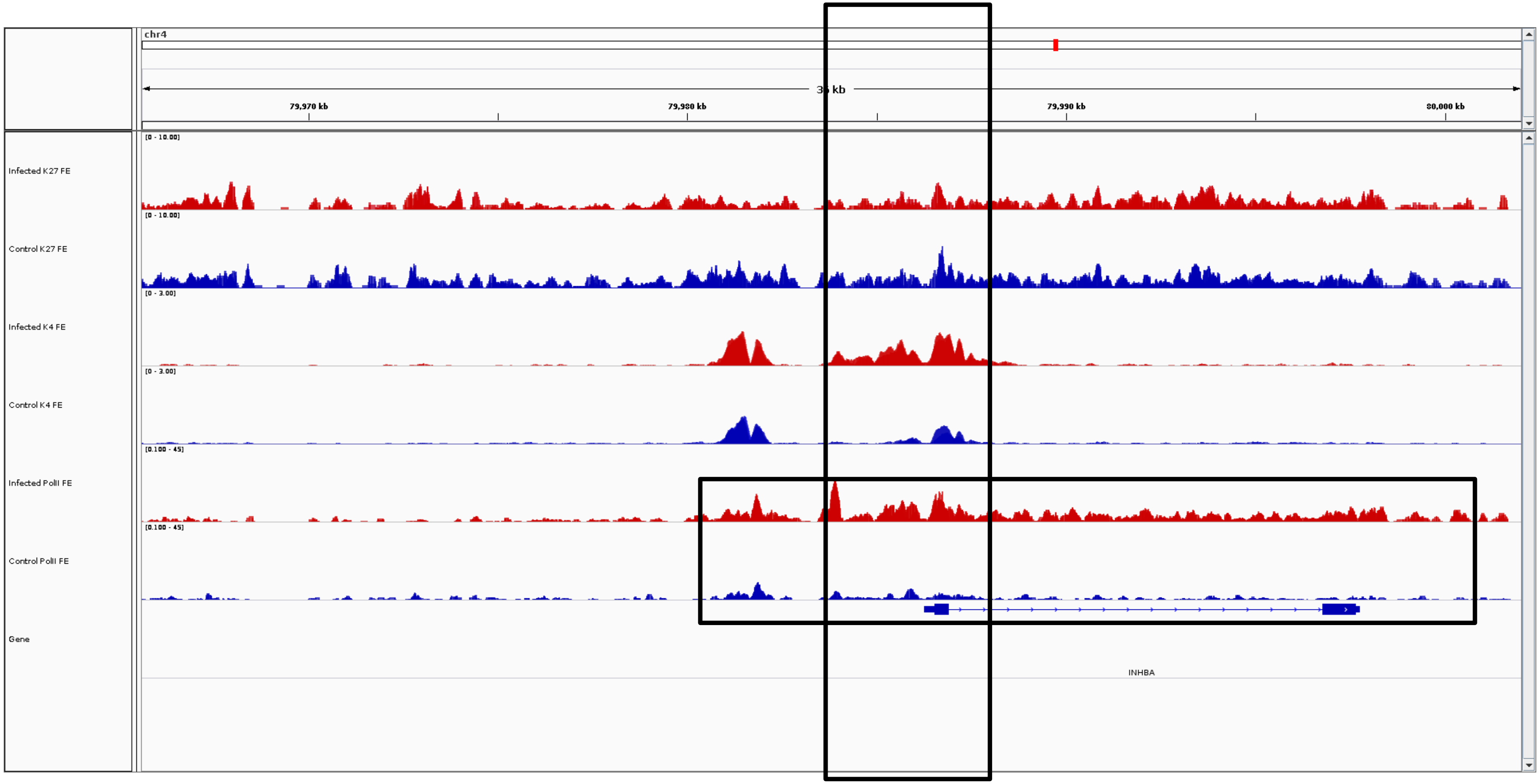
chr4
35 kb
79,970 kb
79,980 kb
79,990 kb
80,000 kb
Infected K27 FE
[0 - 10.00]
Control K27 FE
[0 - 10.00]
Infected K4 FE
[0 - 3.00]
Control K4 FE
[0 - 3.00]
Infected PolII FE
[0.100 - 45]
Control PolII FE
[0.100 - 45]
Gene
INHBA

***INHBA***

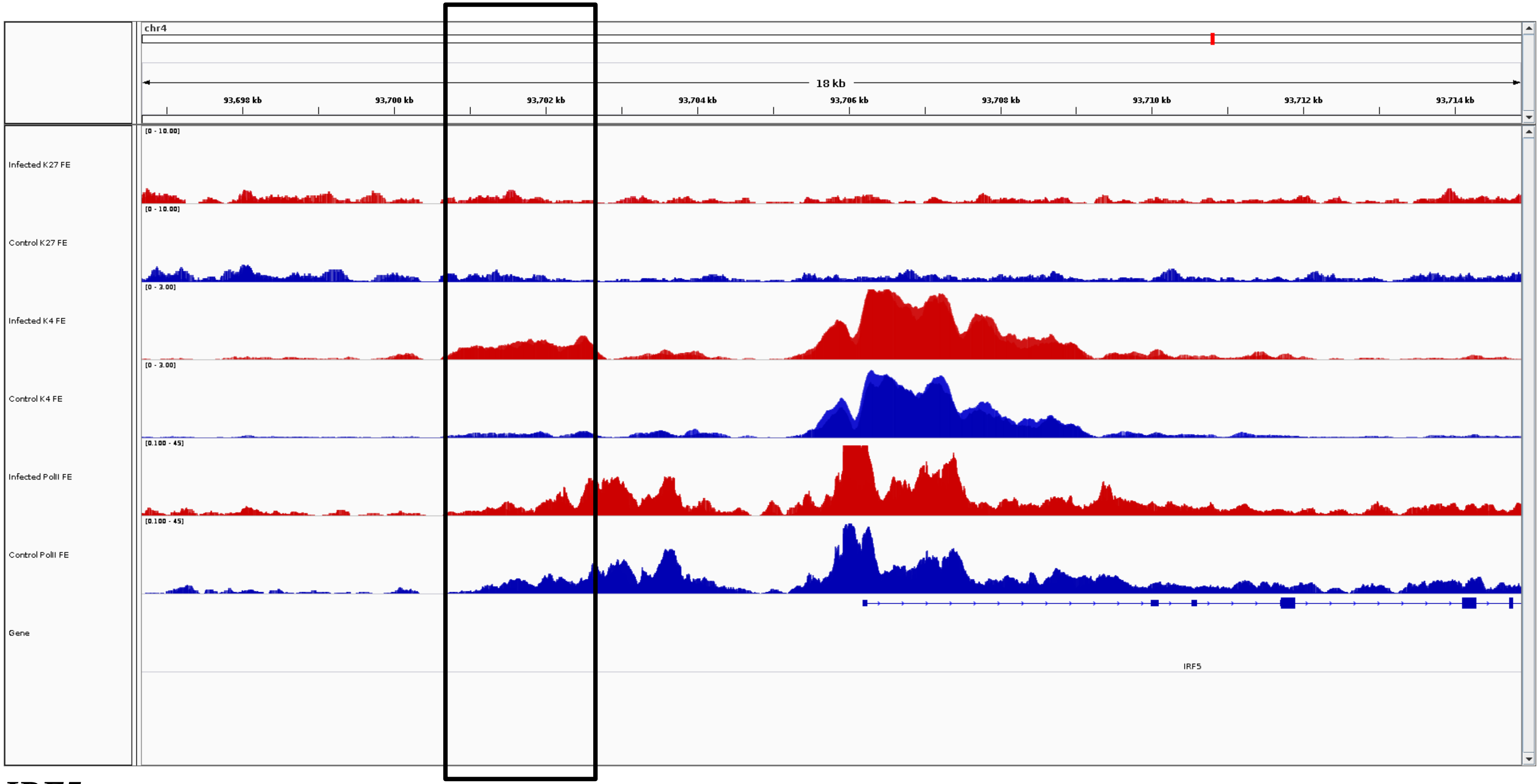
chr4
18 kb
93,698 kb
93,700 kb
93,702 kb
93,704 kb
93,706 kb
93,708 kb
93,710 kb
93,712 kb
93,714 kb
Infected K27 FE
Control K27 FE
Infected K4 FE
Control K4 FE
Infected PolII FE
Control PolII FE
Gene
IRF5

***IRF5***

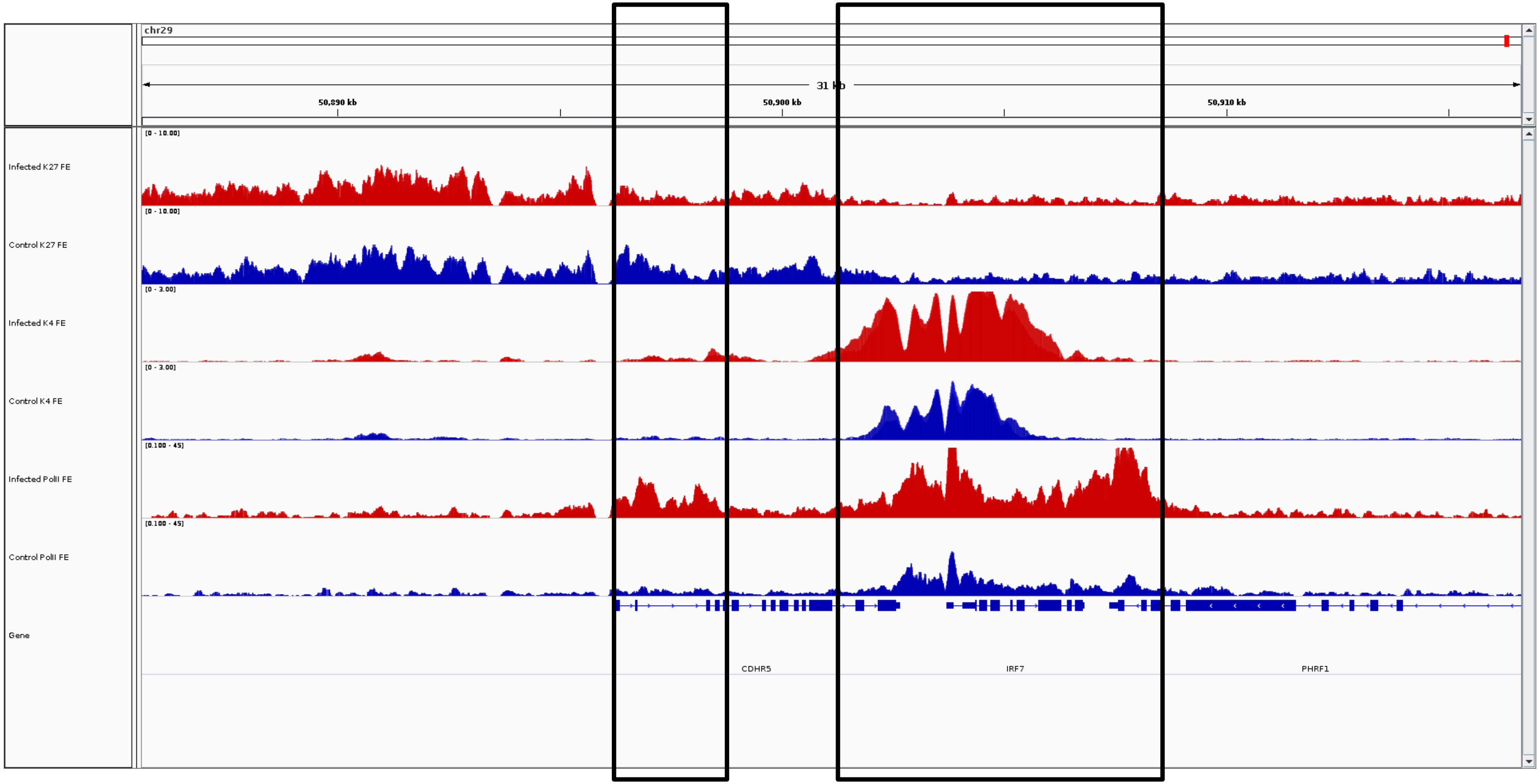
chr29
31 kb
50,890 kb
50,900 kb
50,910 kb
Infected K27 FE
Control K27 FE
Infected K4 FE
Control K4 FE
Infected PolII FE
Control PolII FE
Gene
[0 - 10.00]
[0 - 10.00]
[0 - 3.00]
[0 - 3.00]
[0.100 - 45]
[0.100 - 45]
CDHR5
IRF7
PHRF1

***IRF7***

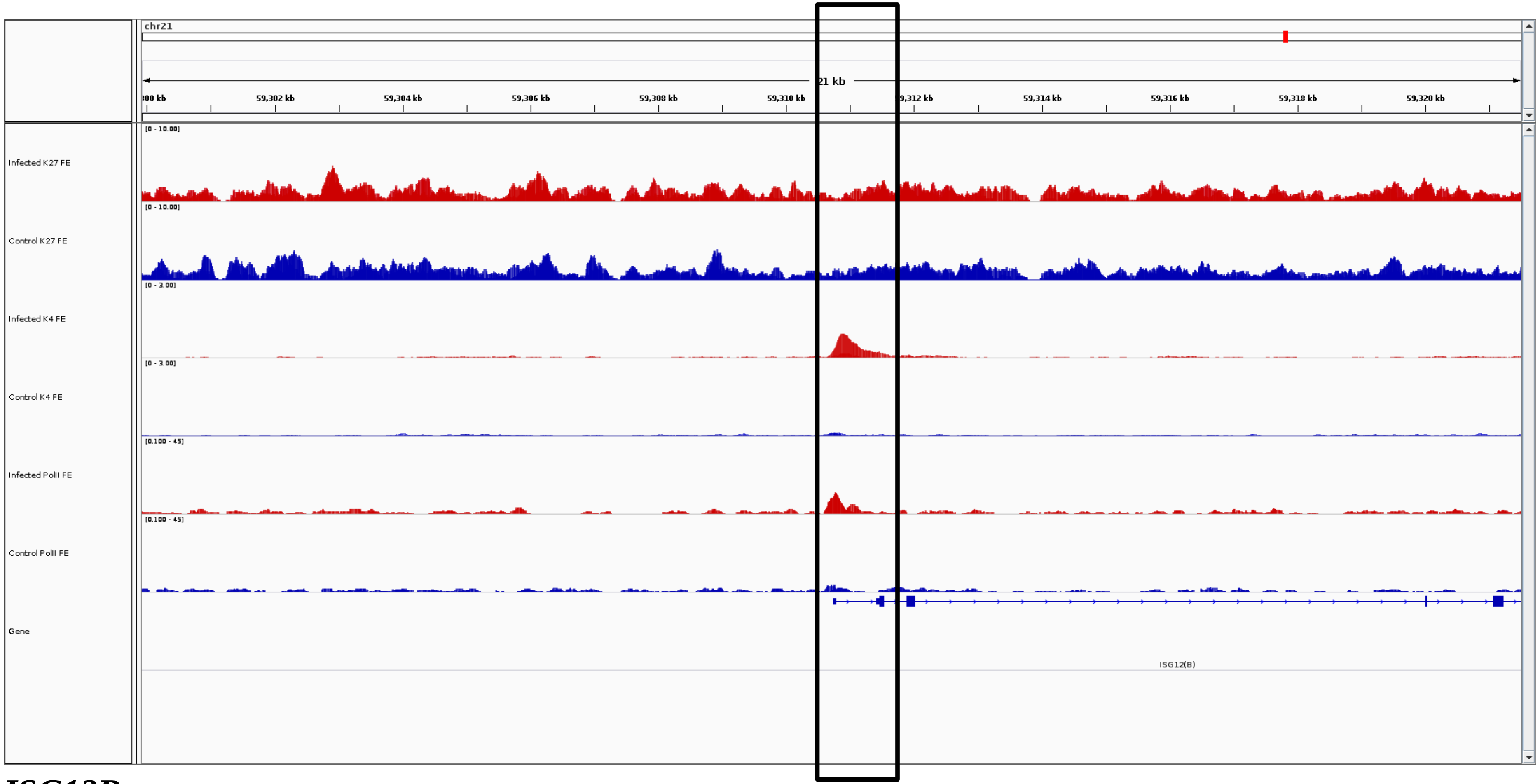
chr21
21 kb
59,302 kb
59,304 kb
59,306 kb
59,308 kb
59,310 kb
59,314 kb
59,316 kb
59,318 kb
59,320 kb
Infected K27 FE
[0 - 10.00]
Control K27 FE
[0 - 10.00]
Infected K4 FE
[0 - 3.00]
Control K4 FE
[0 - 3.00]
Infected PolII FE
[0.100 - 45]
Control PolII FE
[0.100 - 45]
Gene
ISG12(B)

***ISG12B***

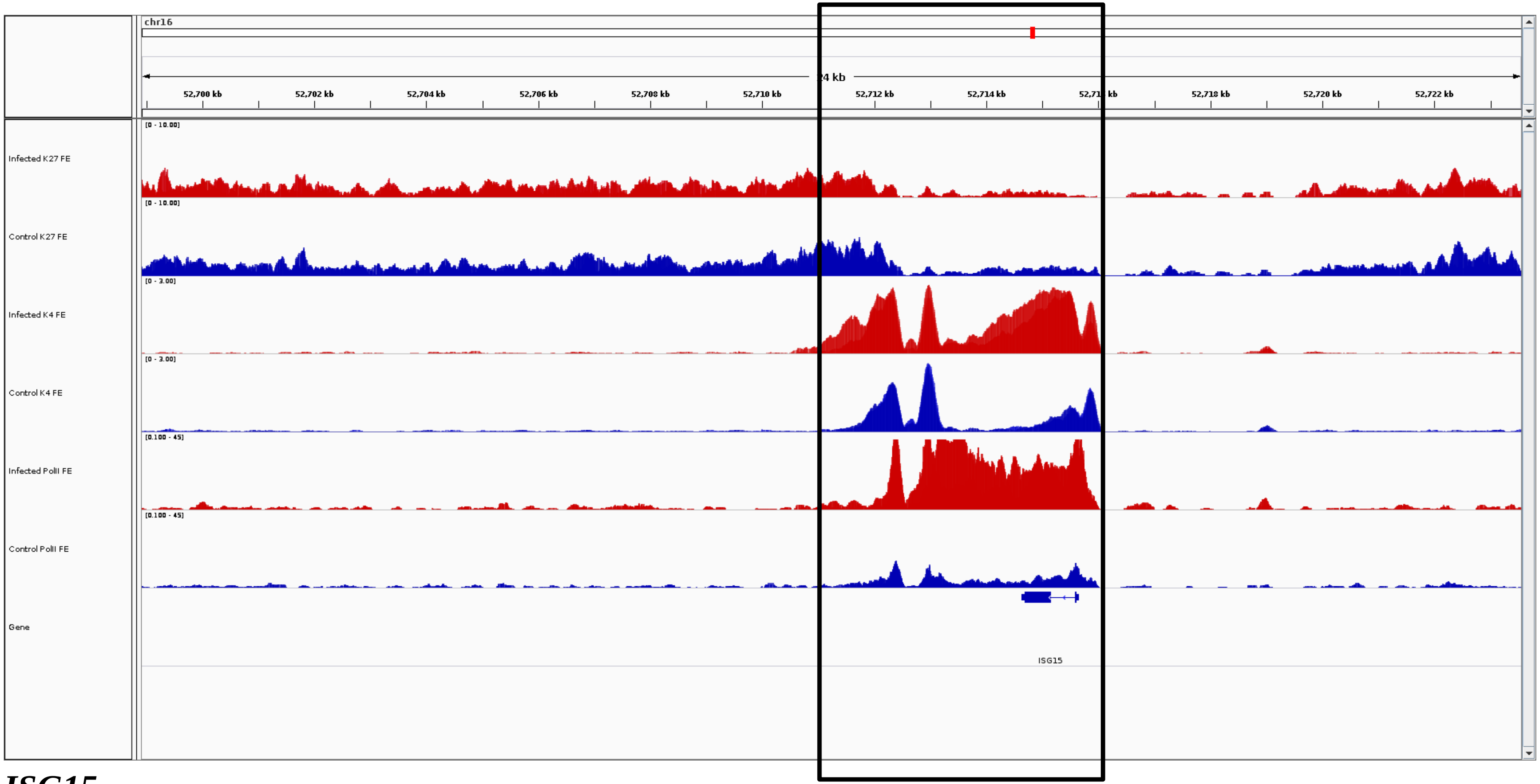
chr16
24 kb
52,700 kb
52,702 kb
52,704 kb
52,706 kb
52,708 kb
52,710 kb
52,712 kb
52,714 kb
52,718 kb
52,720 kb
52,722 kb
Infected K27 FE
[0 - 10.00]
Control K27 FE
[0 - 10.00]
Infected K4 FE
[0 - 3.00]
Control K4 FE
[0 - 3.00]
Infected PolII FE
[0.100 - 45]
Control PolII FE
[0.100 - 45]
Gene
ISG15

***ISG15***

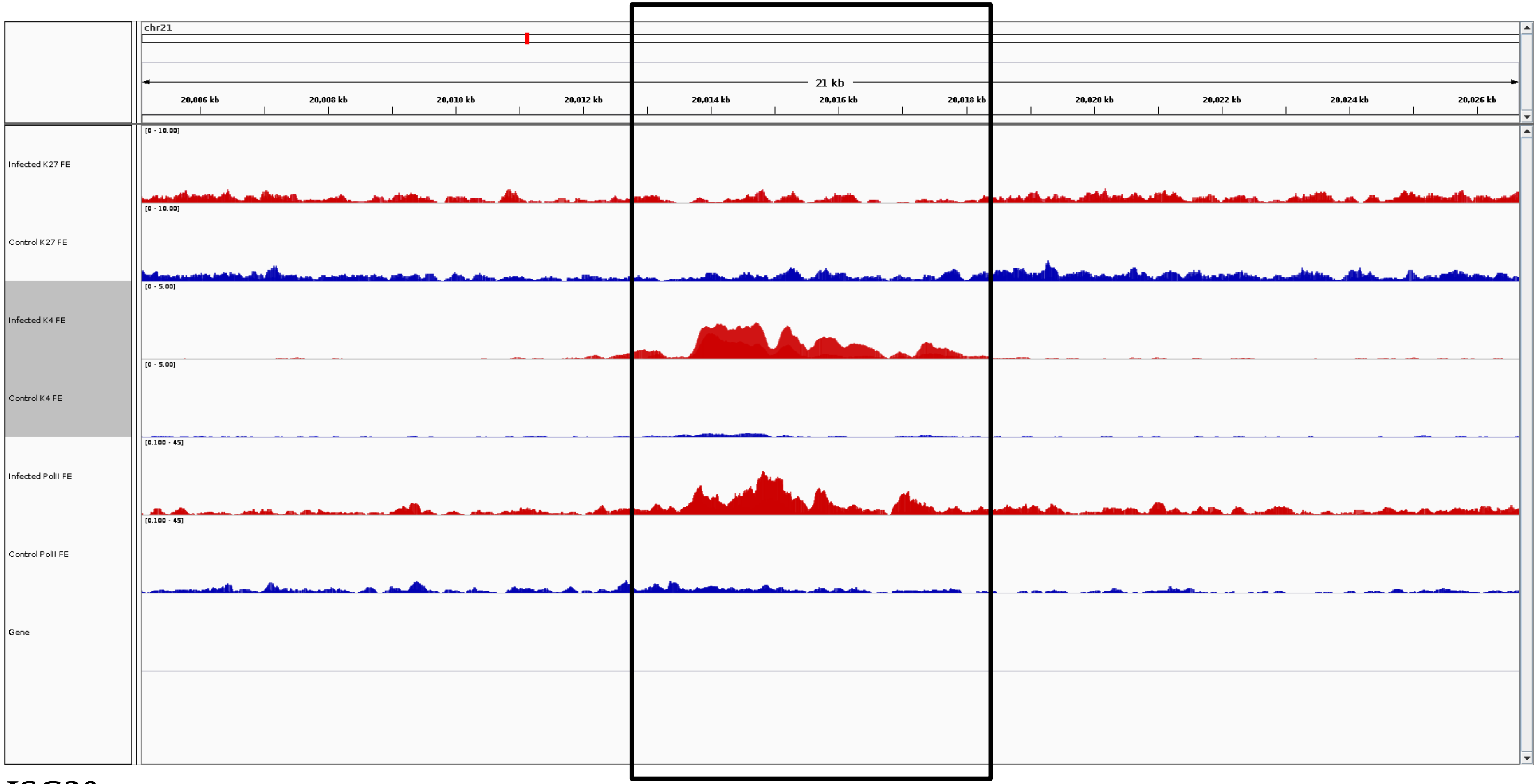

chr21
21 kb
20,006 kb
20,008 kb
20,010 kb
20,012 kb
20,014 kb
20,016 kb
20,018 kb
20,020 kb
20,022 kb
20,024 kb
20,026 kb
Infected K27 FE
Control K27 FE
Infected K4 FE
Control K4 FE
Infected PolII FE
Control PolII FE
Gene


***ISG20***

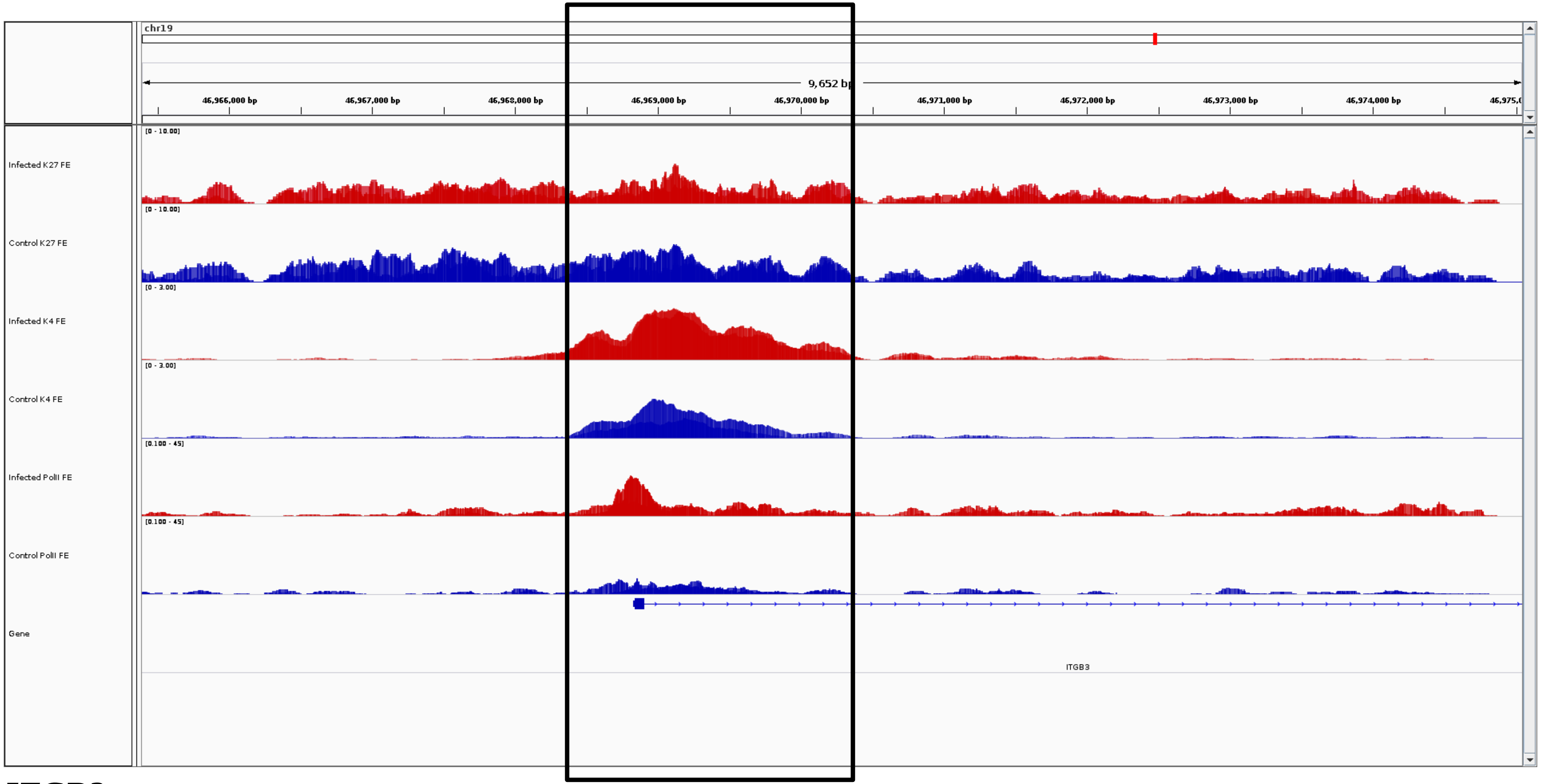
chr19
9,652 bp
46,966,000 bp
46,967,000 bp
46,968,000 bp
46,969,000 bp
46,970,000 bp
46,971,000 bp
46,972,000 bp
46,973,000 bp
46,974,000 bp
Infected K27 FE
Control K27 FE
Infected K4 FE
Control K4 FE
Infected PolII FE
Control PolII FE
Gene
ITGB3

***ITGB3***

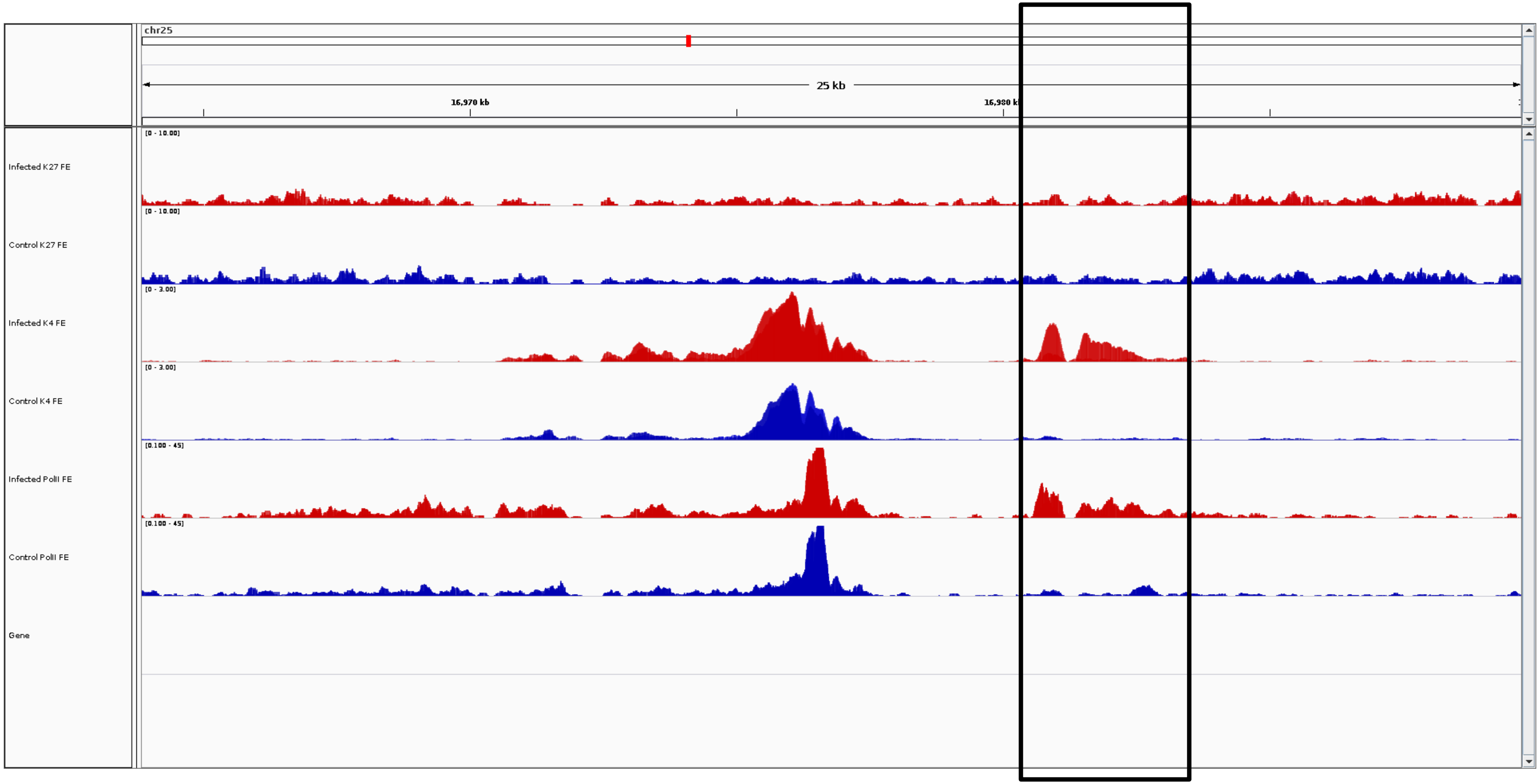
chr25
25 kb
16,970 kb
16,980 kb
Infected K27 FE
[0 - 10.00]
Control K27 FE
[0 - 10.00]
Infected K4 FE
[0 - 3.00]
Control K4 FE
[0 - 3.00]
Infected PolII FE
[0.100 - 45]
Control PolII FE
[0.100 - 45]
Gene

***ITPRIPL2***

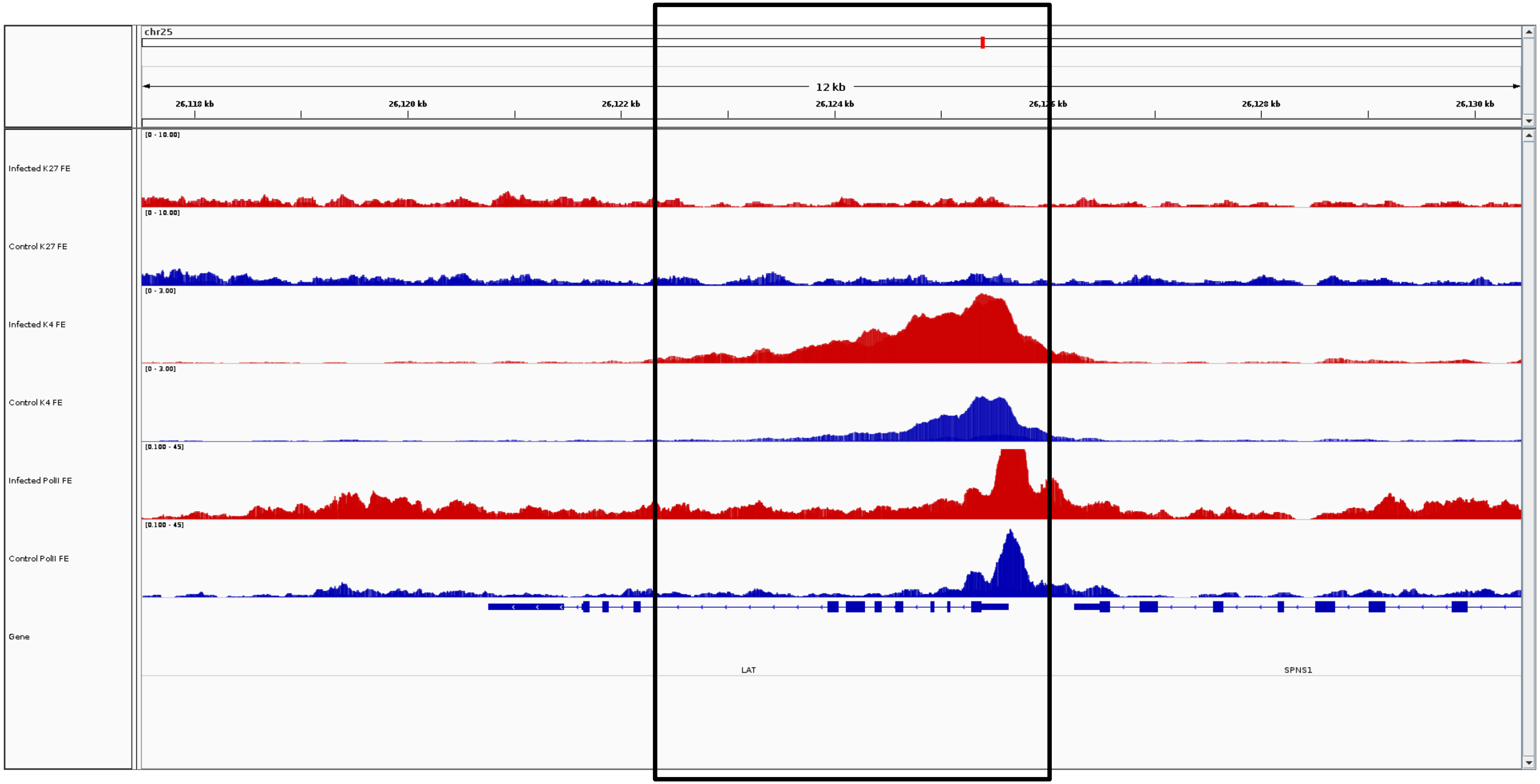
chr25
12 kb
26,118 kb
26,120 kb
26,122 kb
26,124 kb
26,128 kb
26,130 kb
Infected K27 FE
Control K27 FE
Infected K4 FE
Control K4 FE
Infected PolII FE
Control PolII FE
Gene
LAT
SPNS1

***LAT***

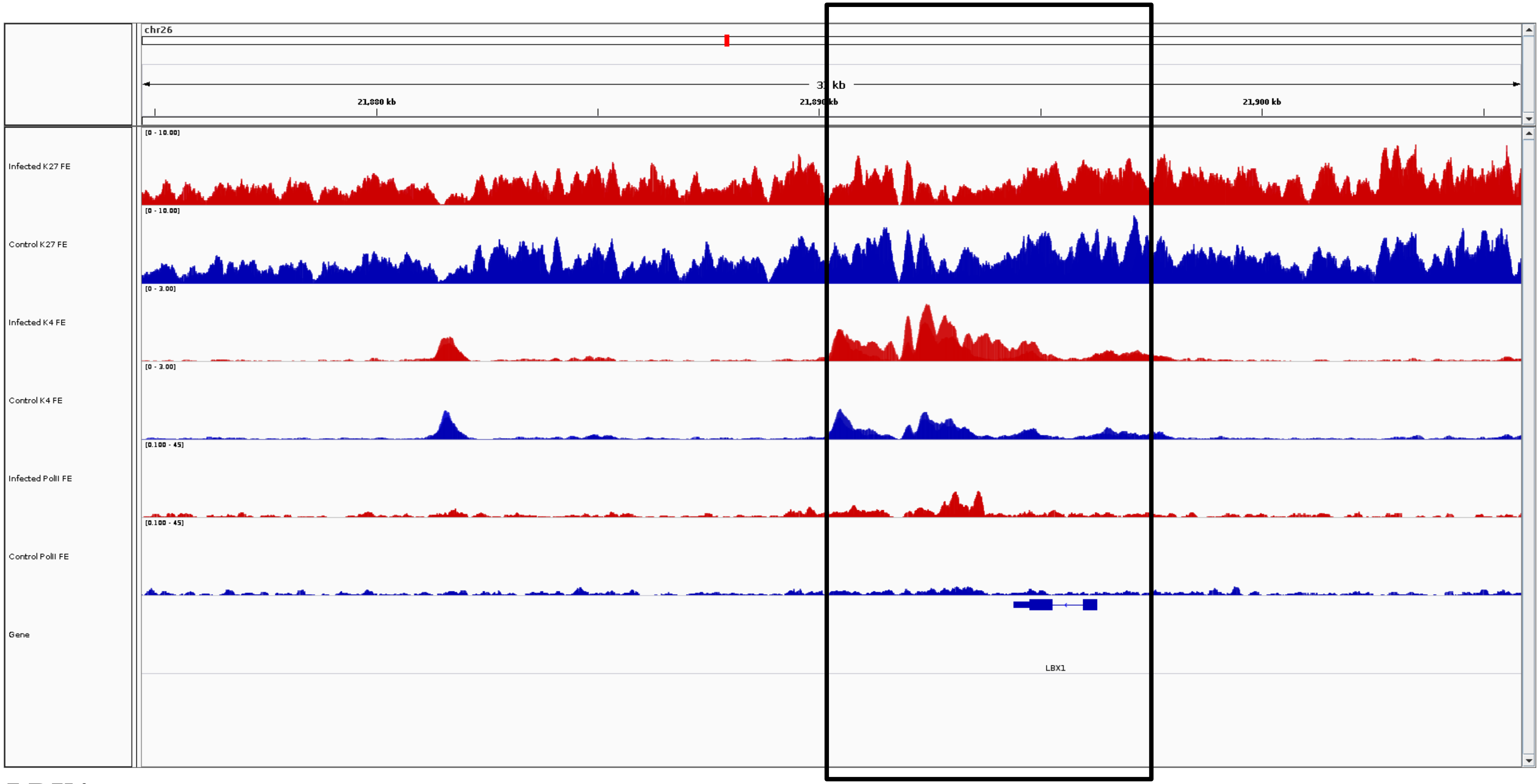
chr26
31 kb
21,880 kb
21,890 kb
21,900 kb
[0 - 10.00]
Infected K27 FE
[0 - 10.00]
Control K27 FE
[0 - 3.00]
Infected K4 FE
[0 - 3.00]
Control K4 FE
[0.100 - 45]
Infected PolII FE
[0.100 - 45]
Control PolII FE
Gene
LBX1

***LBX1***

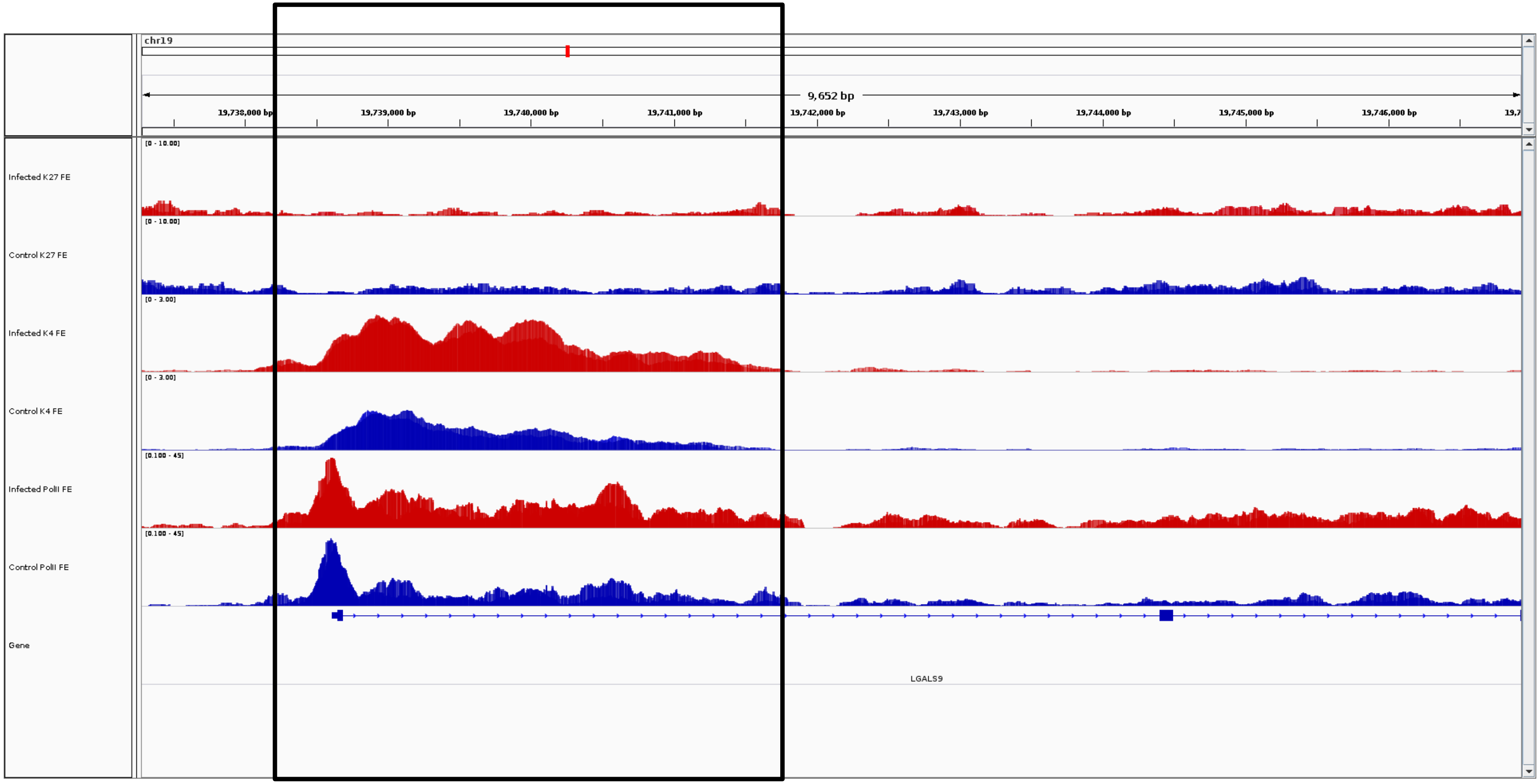
chr19
9,652 bp
19,738,000 bp
19,739,000 bp
19,740,000 bp
19,741,000 bp
19,742,000 bp
19,743,000 bp
19,744,000 bp
19,745,000 bp
19,746,000 bp
Infected K27 FE
[0 - 10.00]
Control K27 FE
[0 - 10.00]
Infected K4 FE
[0 - 3.00]
Control K4 FE
[0 - 3.00]
Infected PolII FE
[0.100 - 45]
Control PolII FE
[0.100 - 45]
Gene
LGALS9

***LGALS9***

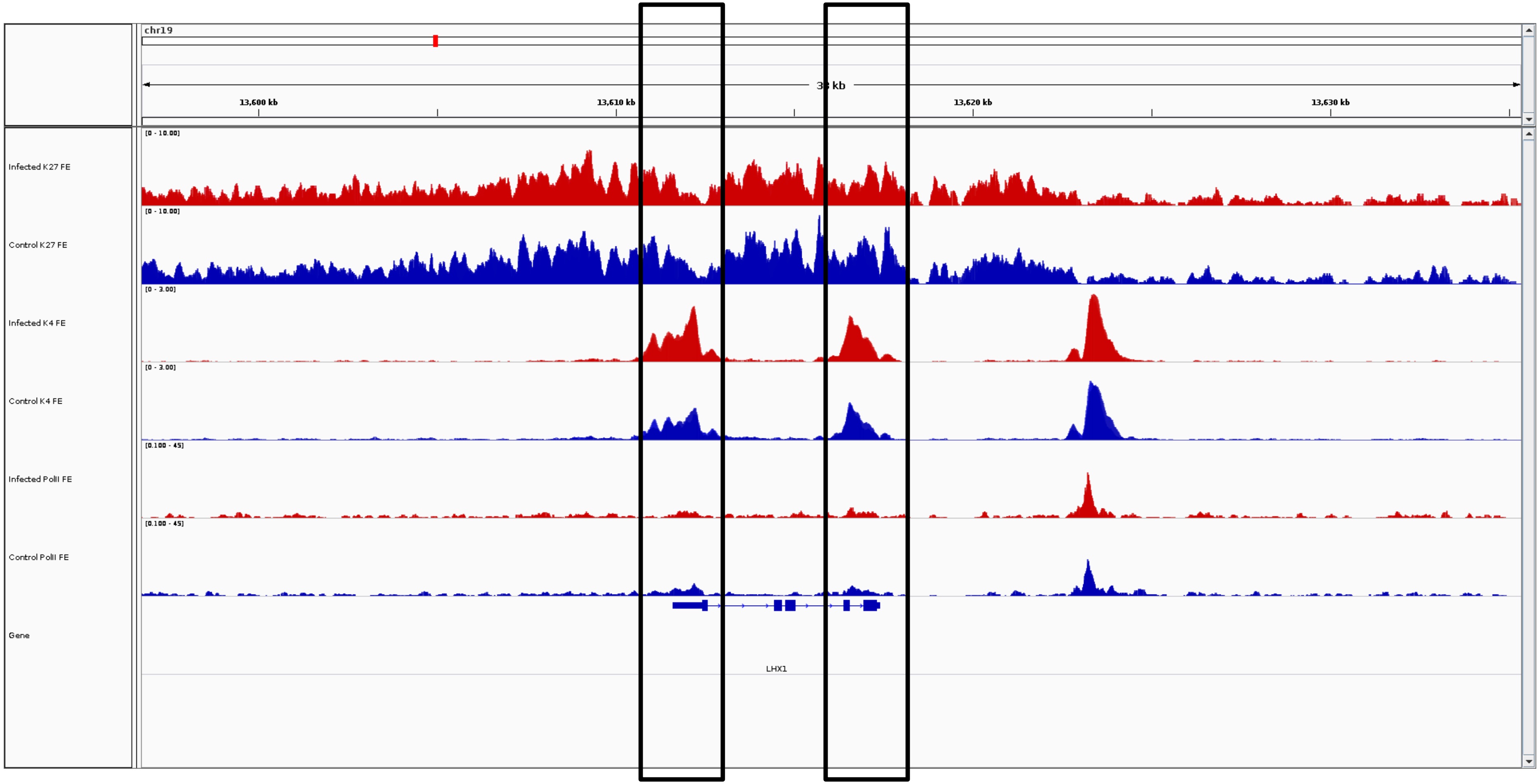
chr19
3[illegible] kb
13,600 kb
13,610 kb
13,620 kb
13,630 kb
Infected K27 FE
Control K27 FE
Infected K4 FE
Control K4 FE
Infected PolII FE
Control PolII FE
Gene
LHX1

***LHX1***

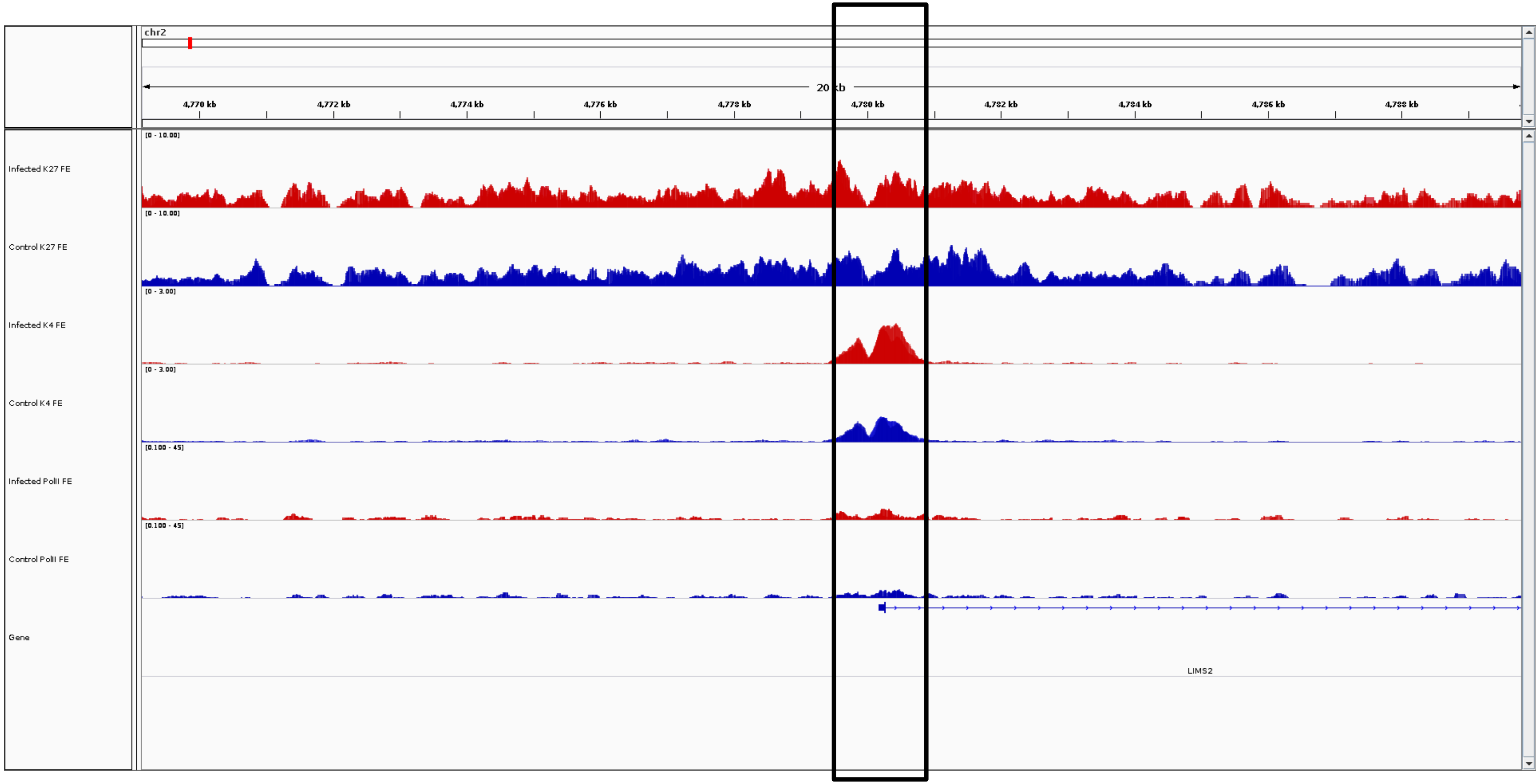
chr2
20 kb
4,770 kb
4,772 kb
4,774 kb
4,776 kb
4,778 kb
4,780 kb
4,782 kb
4,784 kb
4,786 kb
4,788 kb
Infected K27 FE
[0 - 10.00]
Control K27 FE
[0 - 10.00]
Infected K4 FE
[0 - 3.00]
Control K4 FE
[0 - 3.00]
Infected PolII FE
[0.100 - 45]
Control PolII FE
[0.100 - 45]
Gene
LIMS2

***LIMS2***

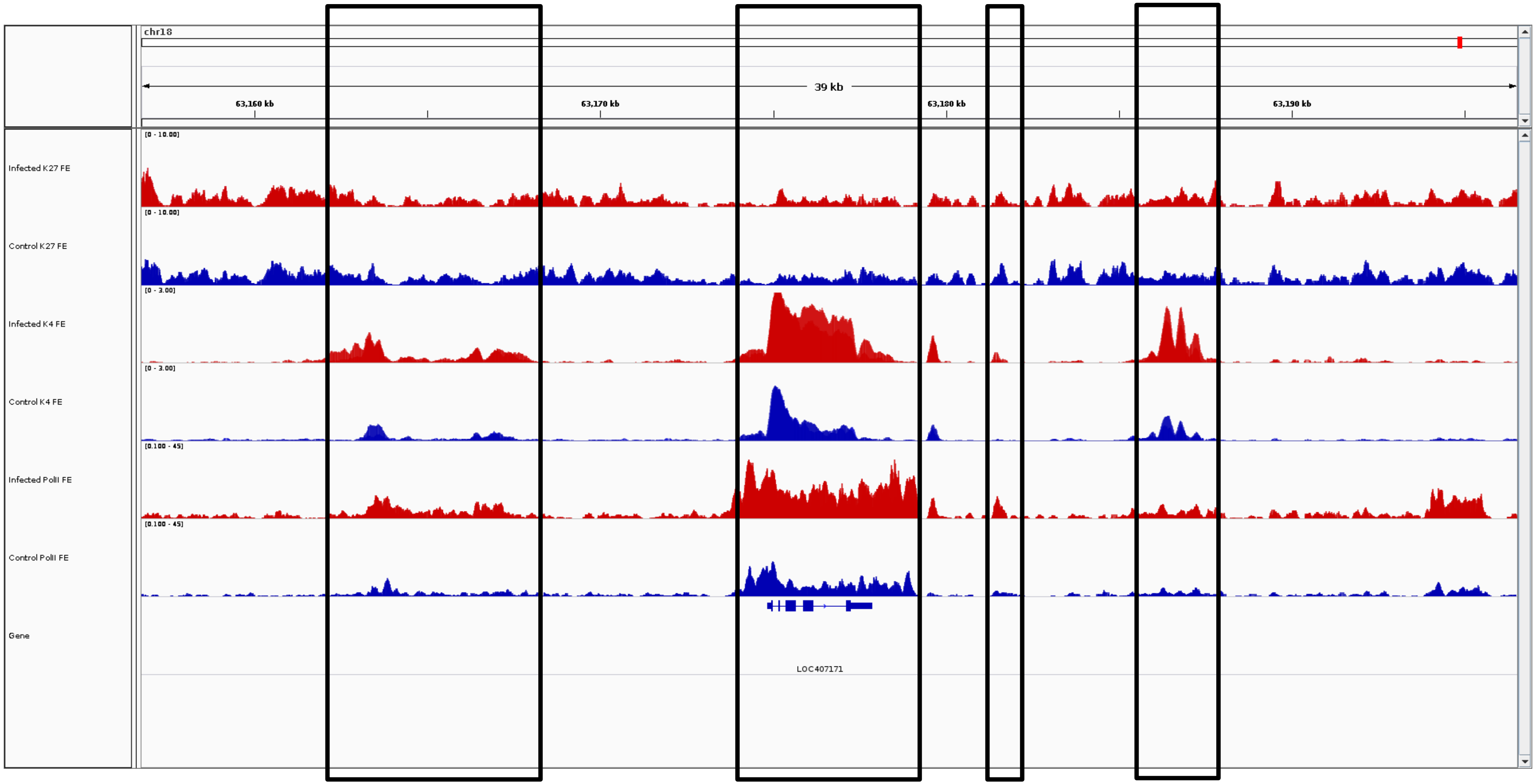
chr18
39 kb
63,160 kb
63,170 kb
63,180 kb
63,190 kb
Infected K27 FE
[0 - 10.00]
Control K27 FE
[0 - 10.00]
Infected K4 FE
[0 - 3.00]
Control K4 FE
[0 - 3.00]
Infected PolII FE
[0.100 - 45]
Control PolII FE
[0.100 - 45]
Gene
LOC407171

***LOC407171***

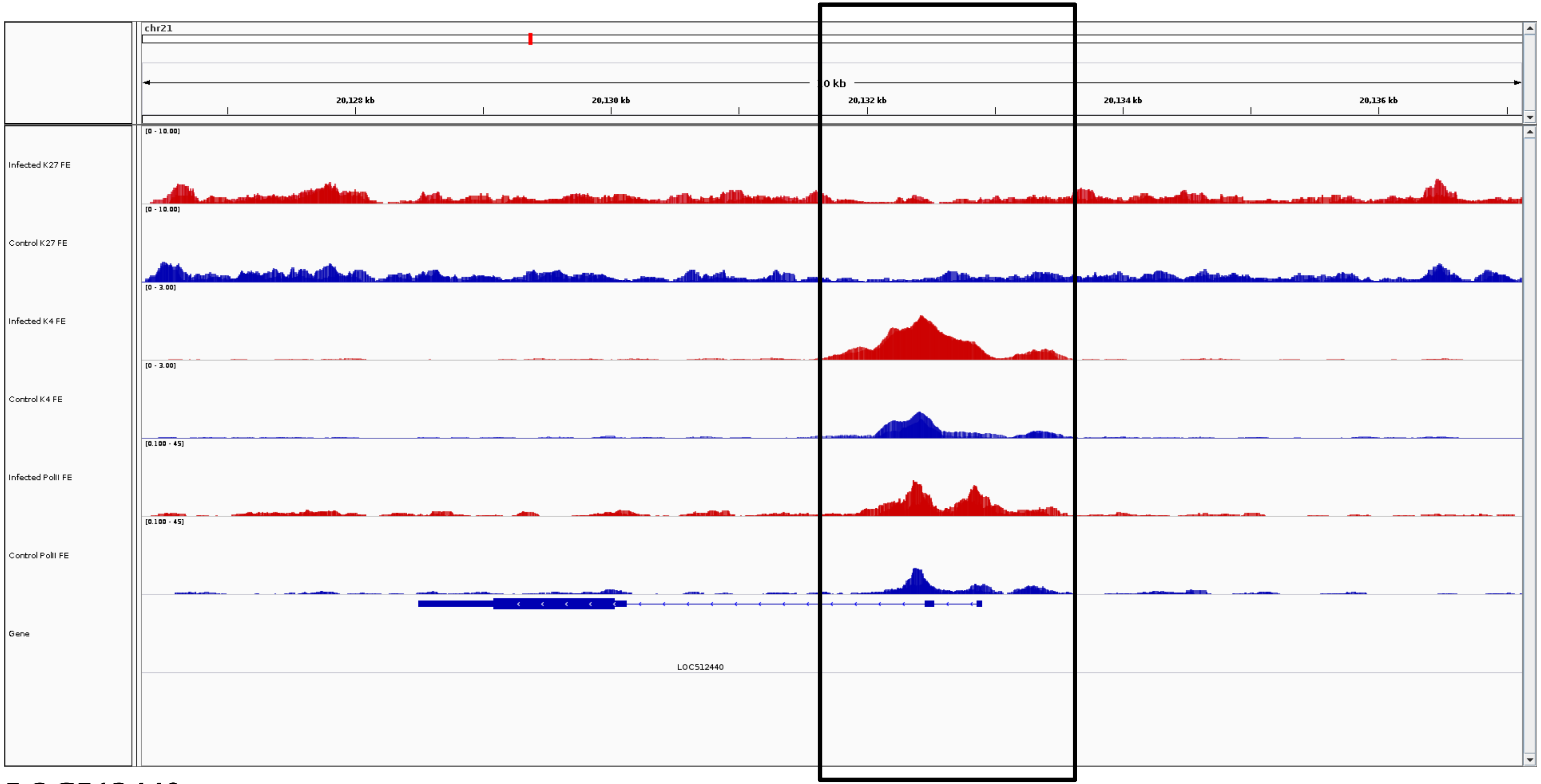
chr21
0 kb
20,128 kb
20,130 kb
20,132 kb
20,134 kb
20,136 kb
Infected K27 FE
Control K27 FE
Infected K4 FE
Control K4 FE
Infected PolII FE
Control PolII FE
Gene
LOC512440

***LOC512440***

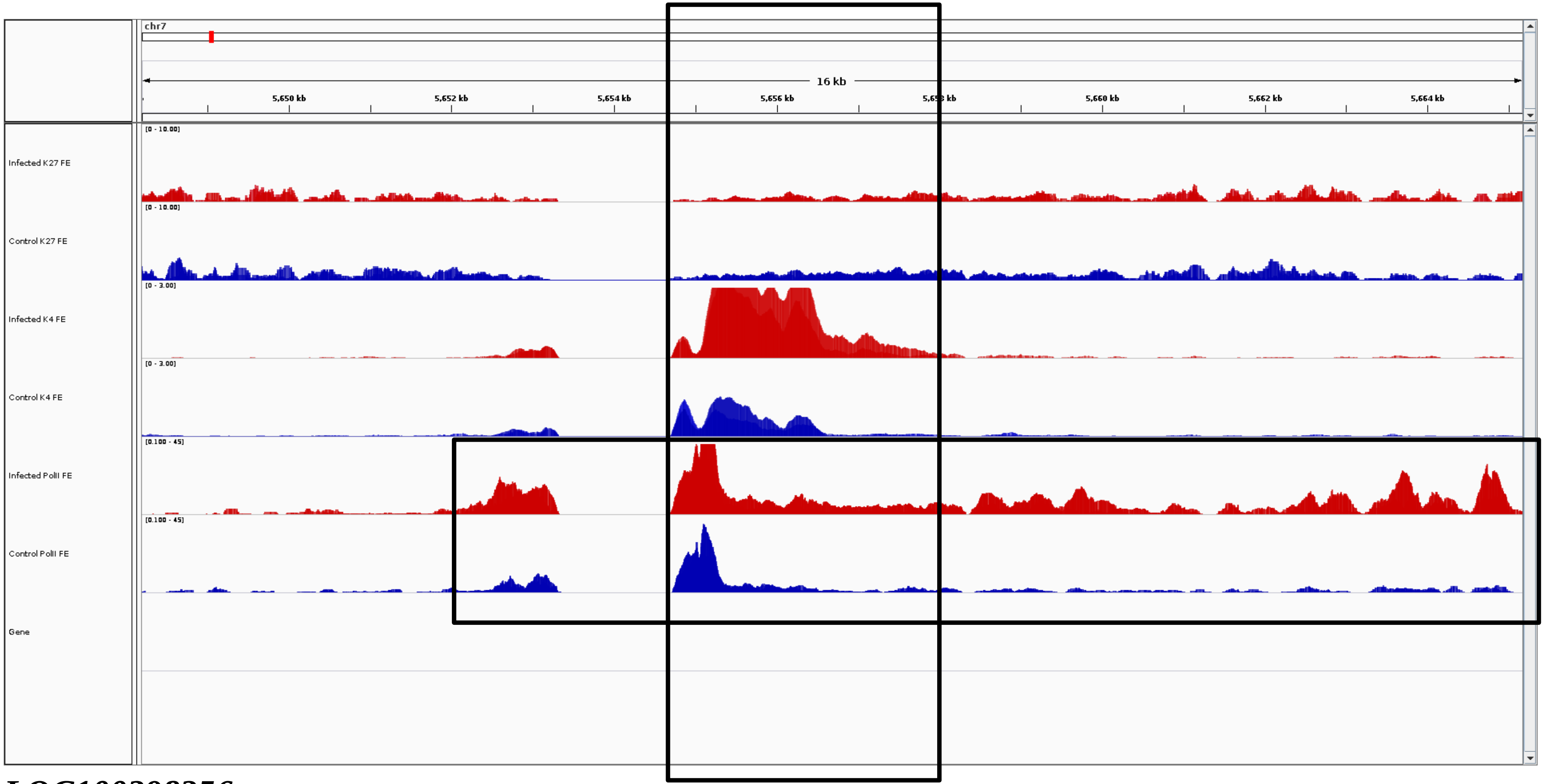


***LOC100298356***

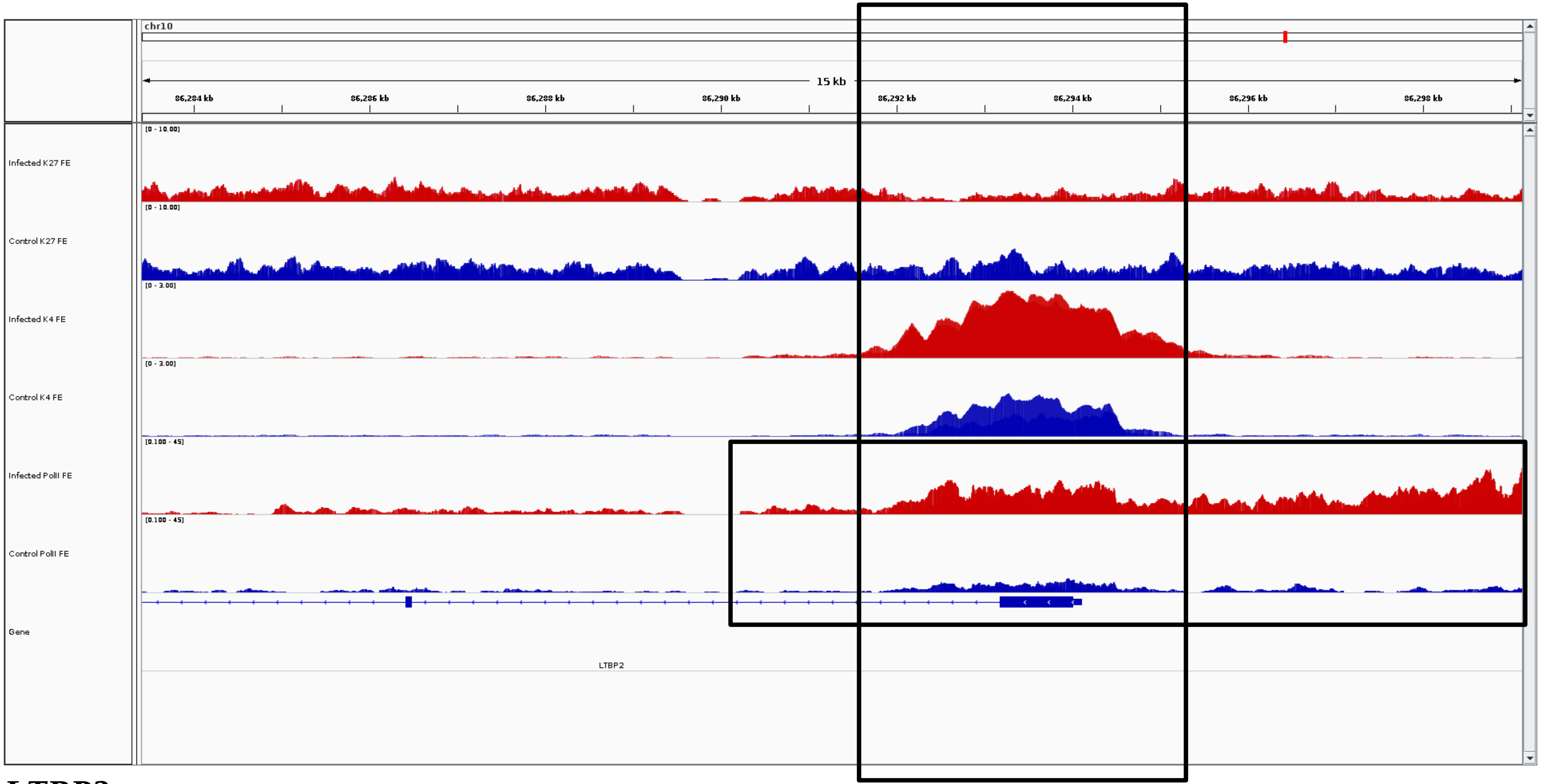
chr10
15 kb
86,284 kb
86,286 kb
86,288 kb
86,290 kb
86,292 kb
86,294 kb
86,296 kb
86,298 kb
Infected K27 FE
[0 - 10.00]
Control K27 FE
[0 - 10.00]
Infected K4 FE
[0 - 3.00]
Control K4 FE
[0 - 3.00]
Infected PolII FE
[0.100 - 45]
Control PolII FE
[0.100 - 45]
Gene
LTBP2

***LTBP2***

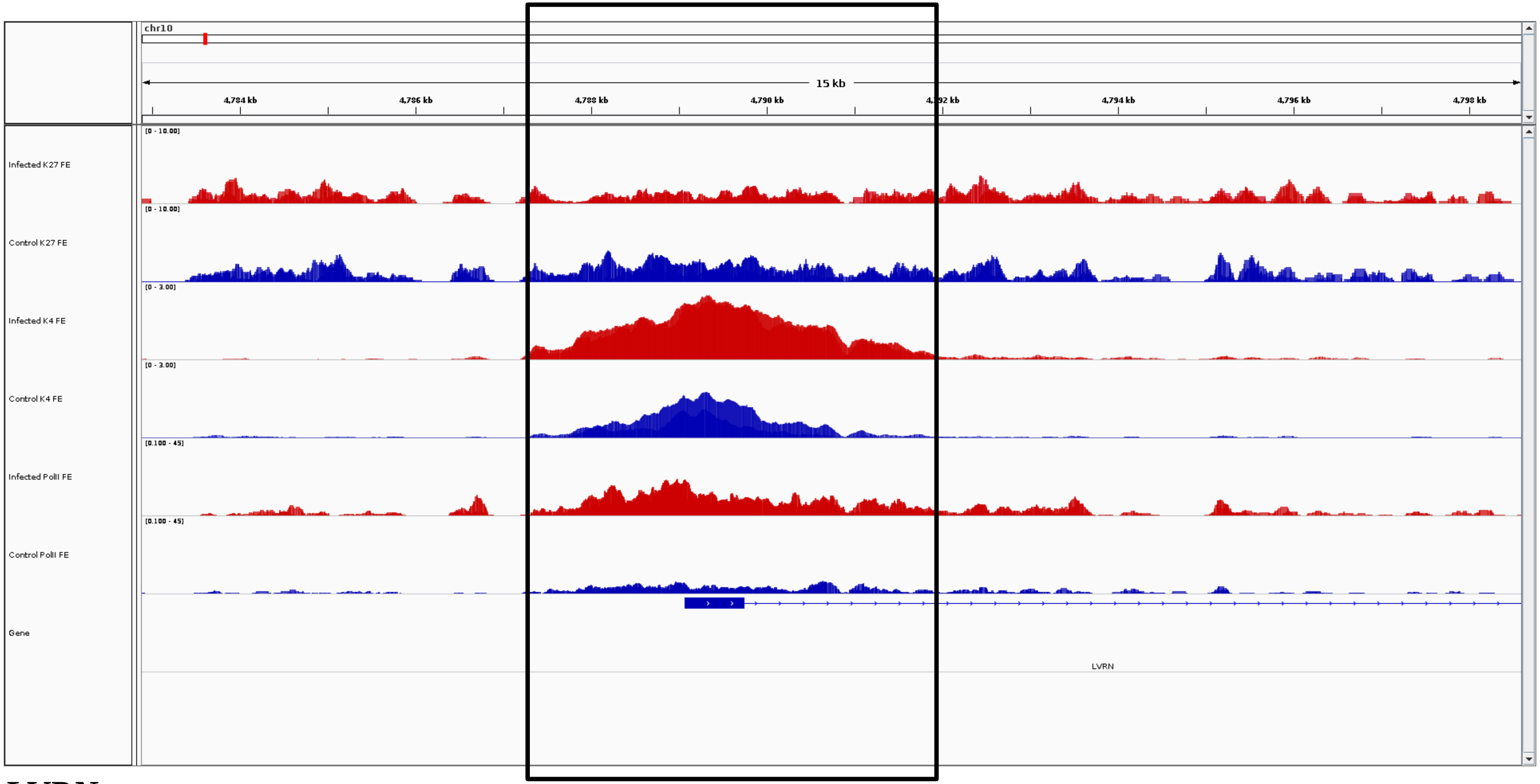
chr10
15 kb
4,784 kb
4,786 kb
4,788 kb
4,790 kb
4,794 kb
4,796 kb
4,798 kb
Infected K27 FE
Control K27 FE
Infected K4 FE
Control K4 FE
Infected PolII FE
Control PolII FE
Gene
LVRN

***LVRN***

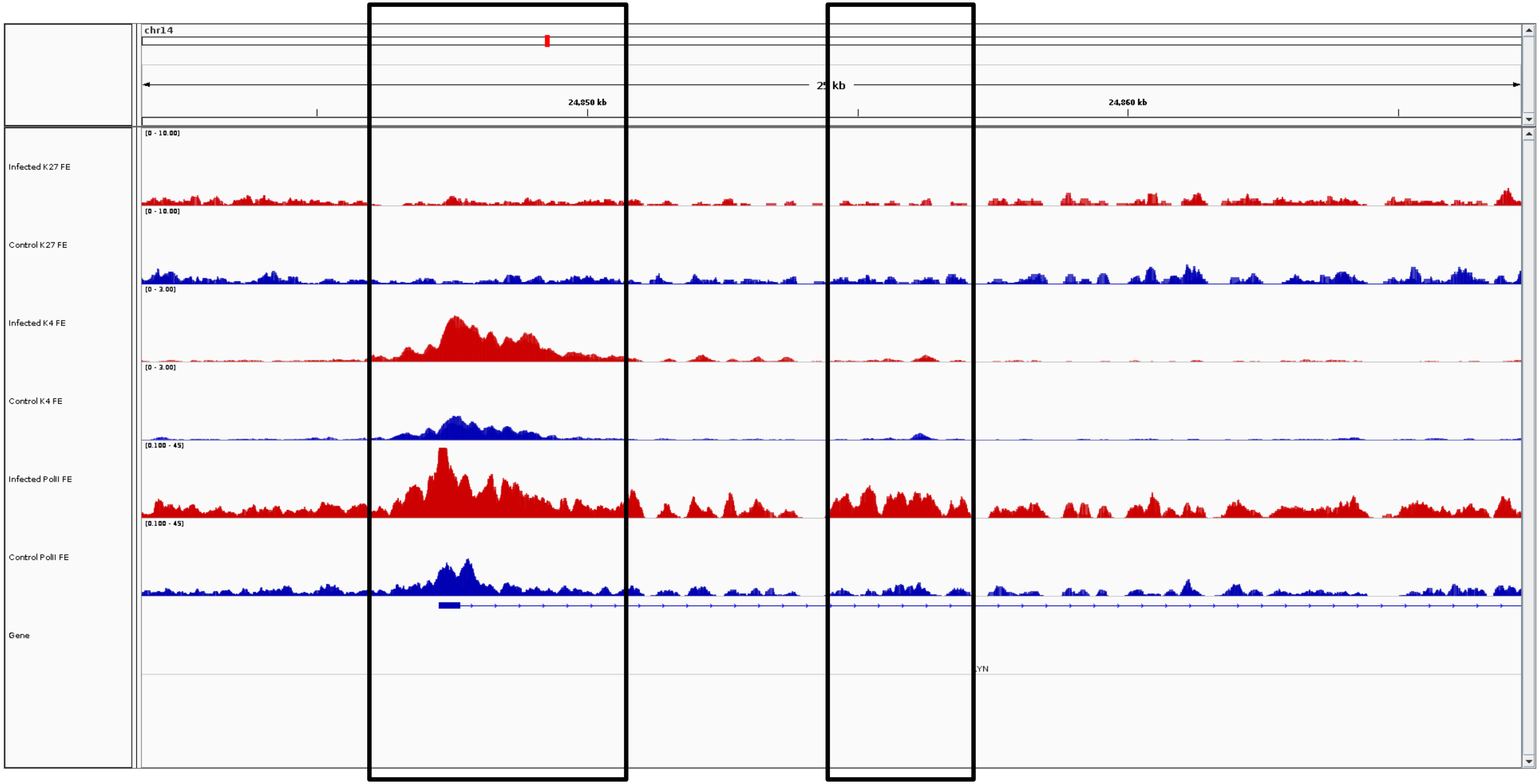
chr14
25 kb
24,850 kb
24,860 kb
Infected K27 FE
Control K27 FE
Infected K4 FE
Control K4 FE
Infected PolII FE
Control PolII FE
Gene
LYN

***LYN***

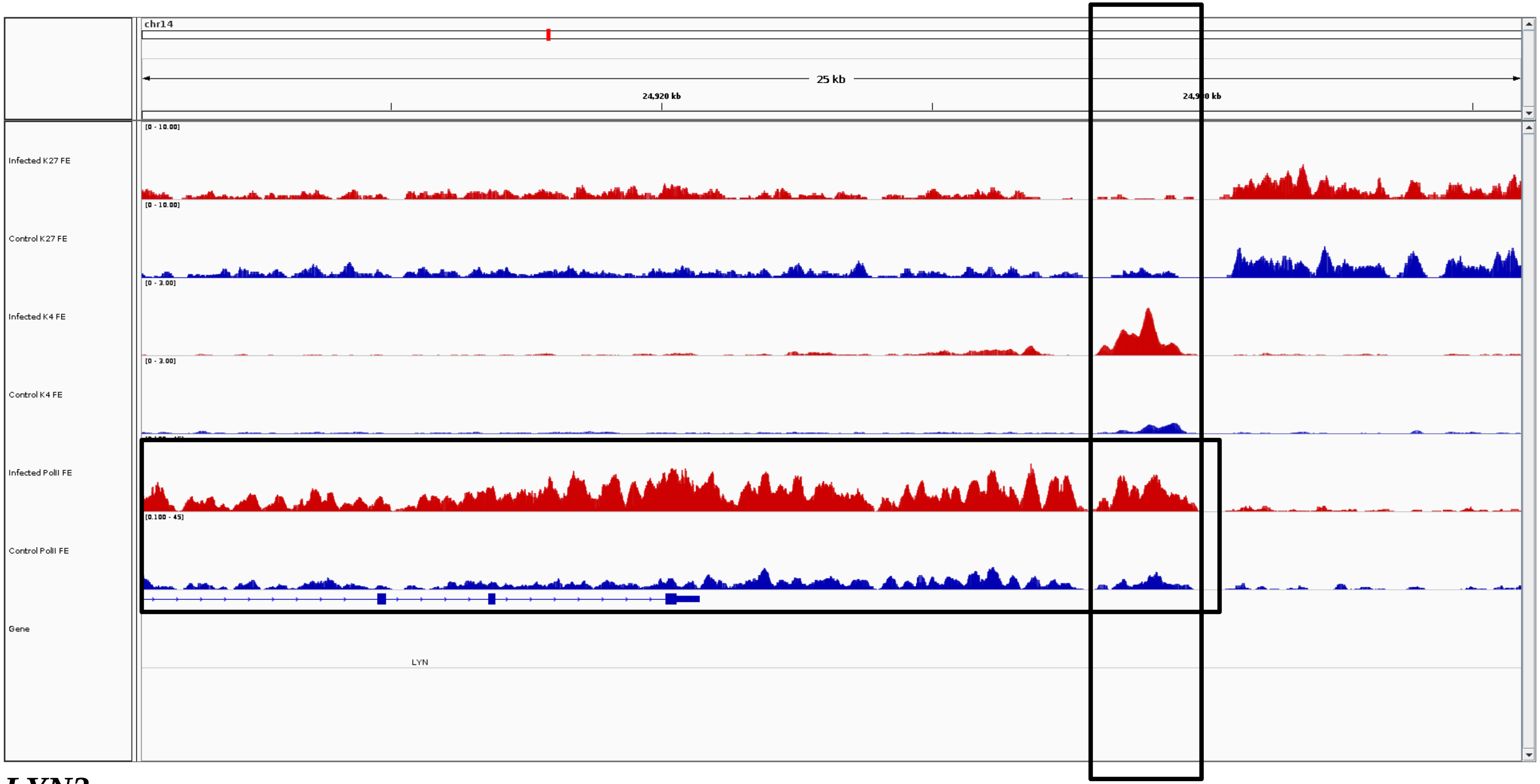
chr14
25 kb
24,920 kb
Infected K27 FE
Control K27 FE
Infected K4 FE
Control K4 FE
Infected PolII FE
Control PolII FE
Gene
LYN

***LYN3***

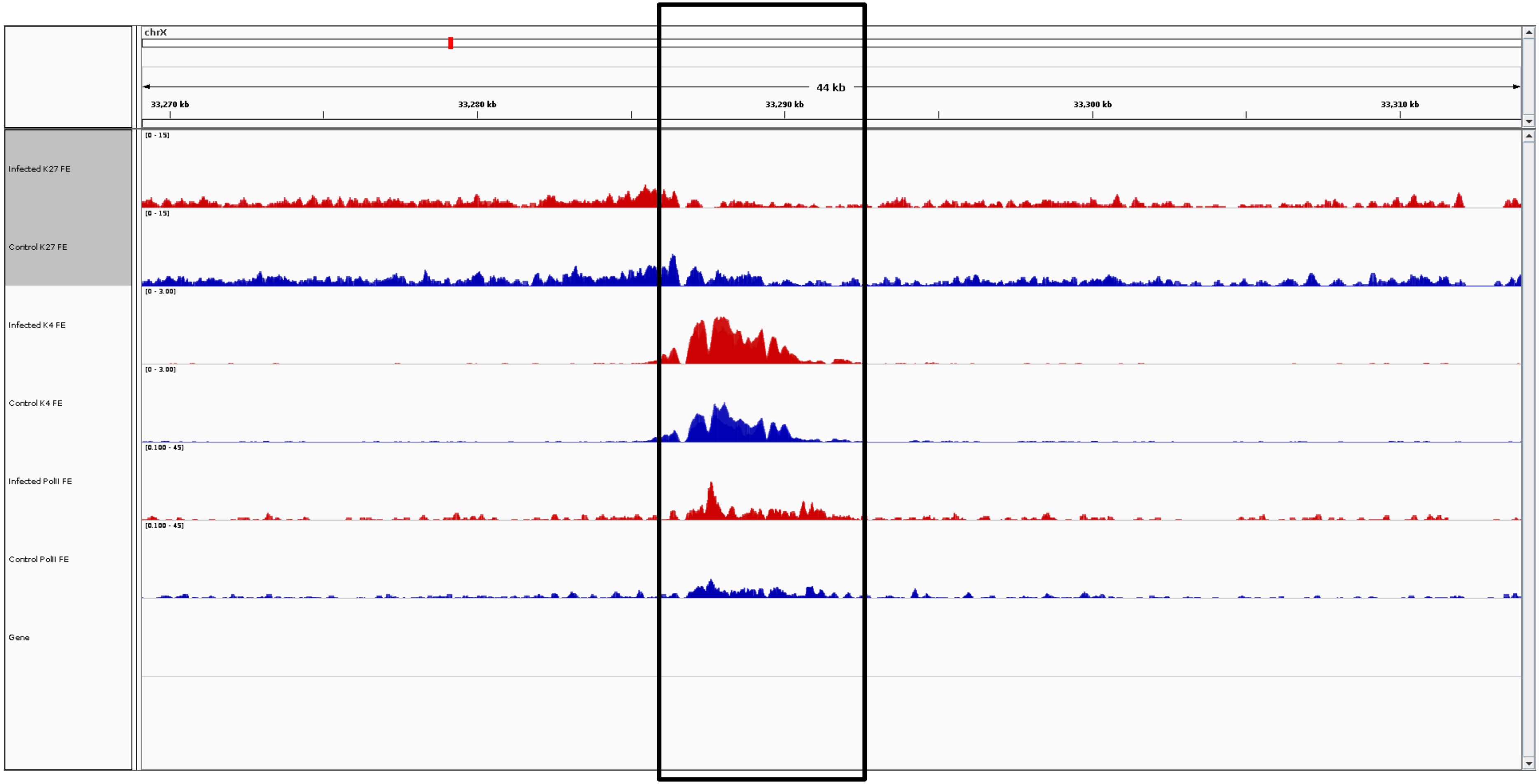
chrX
44 kb
33,270 kb
33,280 kb
33,290 kb
33,300 kb
33,310 kb
Infected K27 FE
[0 - 15]
Control K27 FE
[0 - 15]
Infected K4 FE
[0 - 3.00]
Control K4 FE
[0 - 3.00]
Infected PolII FE
[0.100 - 45]
Control PolII FE
[0.100 - 45]
Gene

***MAMLD1***

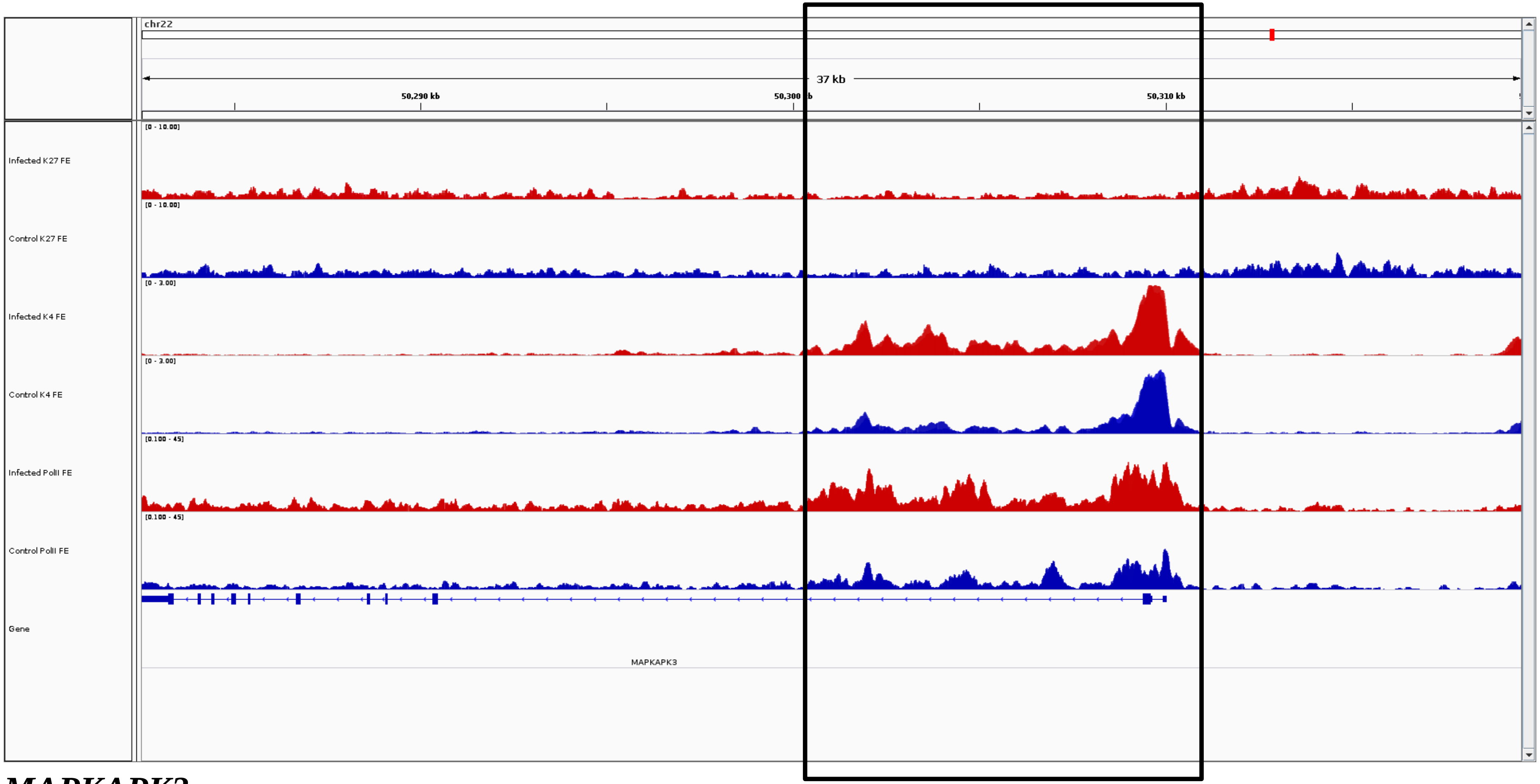

chr22
37 kb
50,290 kb
50,300 kb
50,310 kb
Infected K27 FE
[0 - 10.00]
Control K27 FE
[0 - 10.00]
Infected K4 FE
[0 - 3.00]
Control K4 FE
[0 - 3.00]
Infected PolII FE
[0.100 - 45]
Control PolII FE
[0.100 - 45]
Gene
MAPKAPK3


***MAPKAPK3***

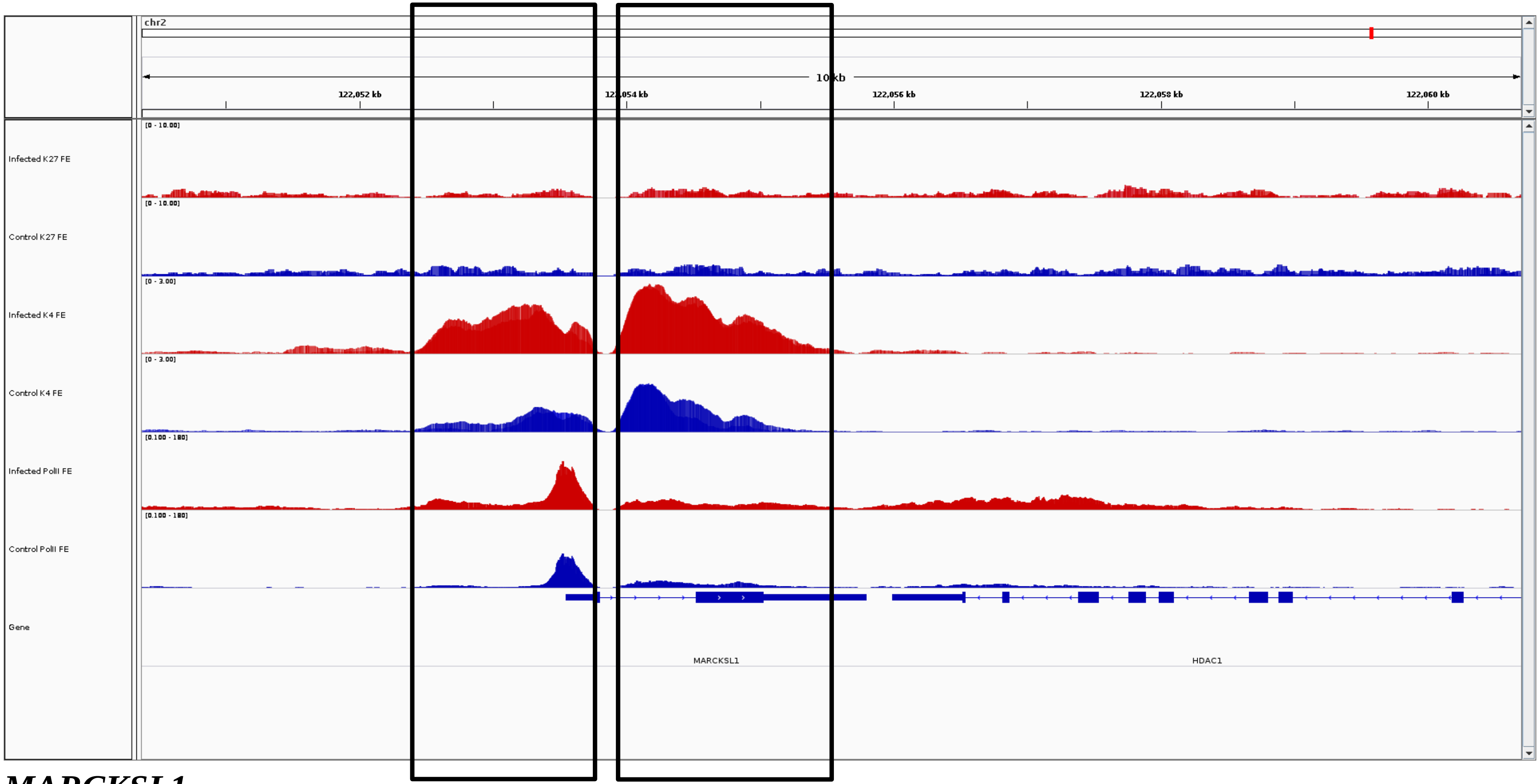
chr2
10 kb
122,052 kb
122,054 kb
122,056 kb
122,058 kb
122,060 kb
Infected K27 FE
Control K27 FE
Infected K4 FE
Control K4 FE
Infected PolII FE
Control PolII FE
Gene
MARCKSL1
HDAC1

***MARCKSL1***

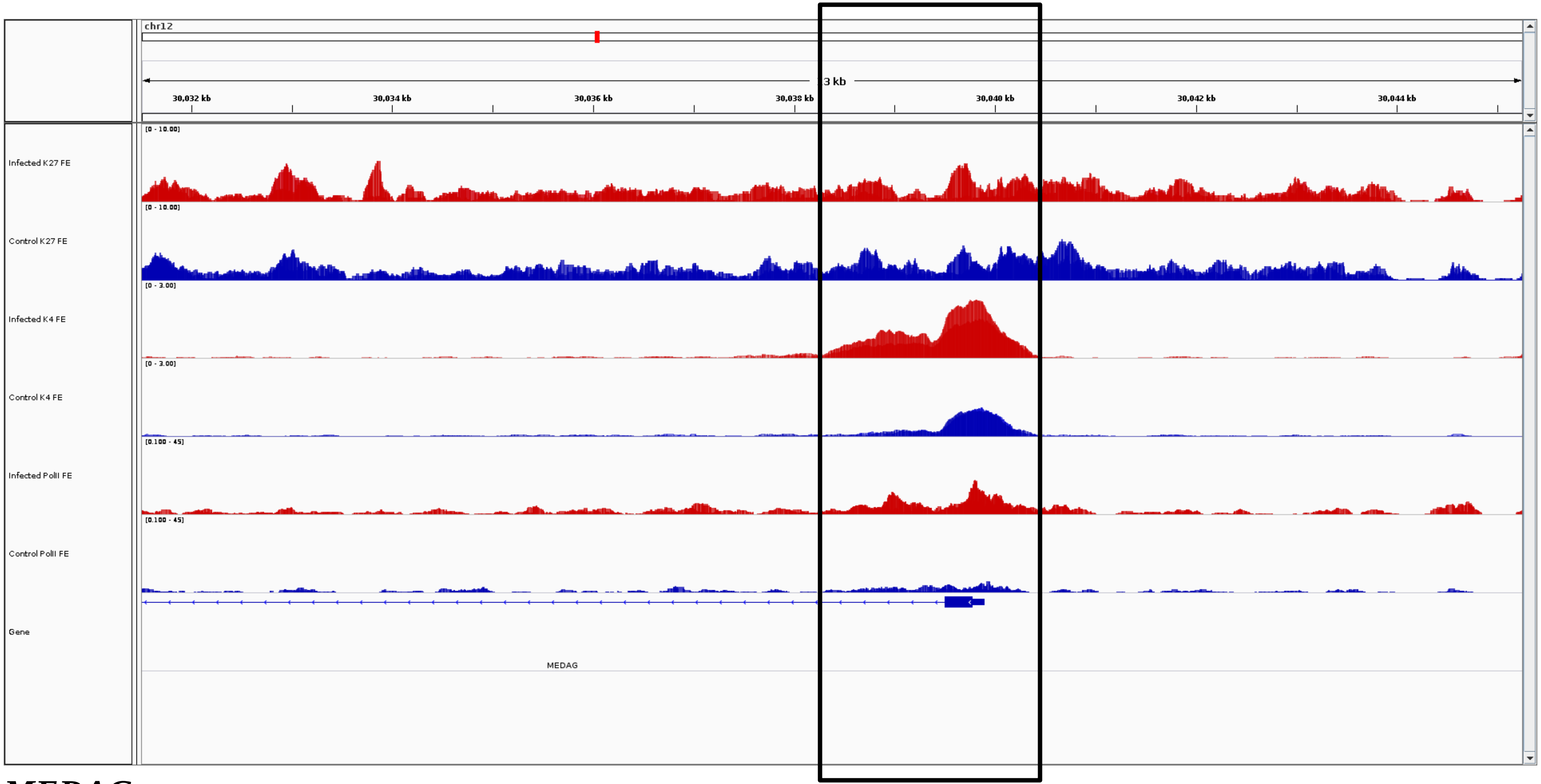
chr12
3 kb
30,032 kb
30,034 kb
30,036 kb
30,038 kb
30,040 kb
30,042 kb
30,044 kb
Infected K27 FE
[0 - 10.00]
Control K27 FE
[0 - 10.00]
Infected K4 FE
[0 - 3.00]
Control K4 FE
[0 - 3.00]
Infected PolII FE
[0.100 - 45]
Control PolII FE
[0.100 - 45]
Gene
MEDAG

***MEDAG***

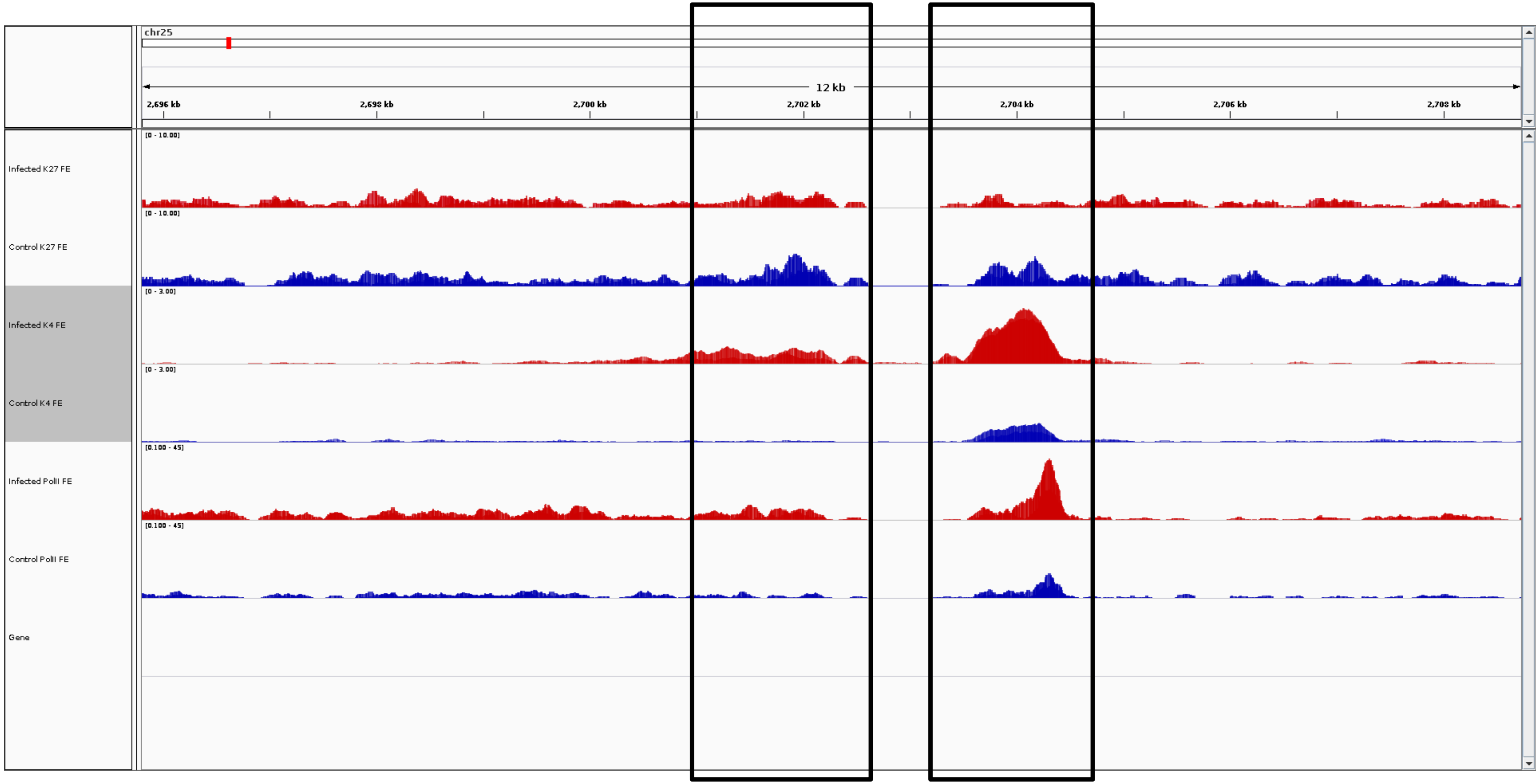
chr25
12 kb
2,696 kb
2,698 kb
2,700 kb
2,702 kb
2,704 kb
2,706 kb
2,708 kb
Infected K27 FE
[0 - 10.00]
Control K27 FE
[0 - 10.00]
Infected K4 FE
[0 - 3.00]
Control K4 FE
[0 - 3.00]
Infected PolII FE
[0.100 - 45]
Control PolII FE
[0.100 - 45]
Gene

***MEFV***

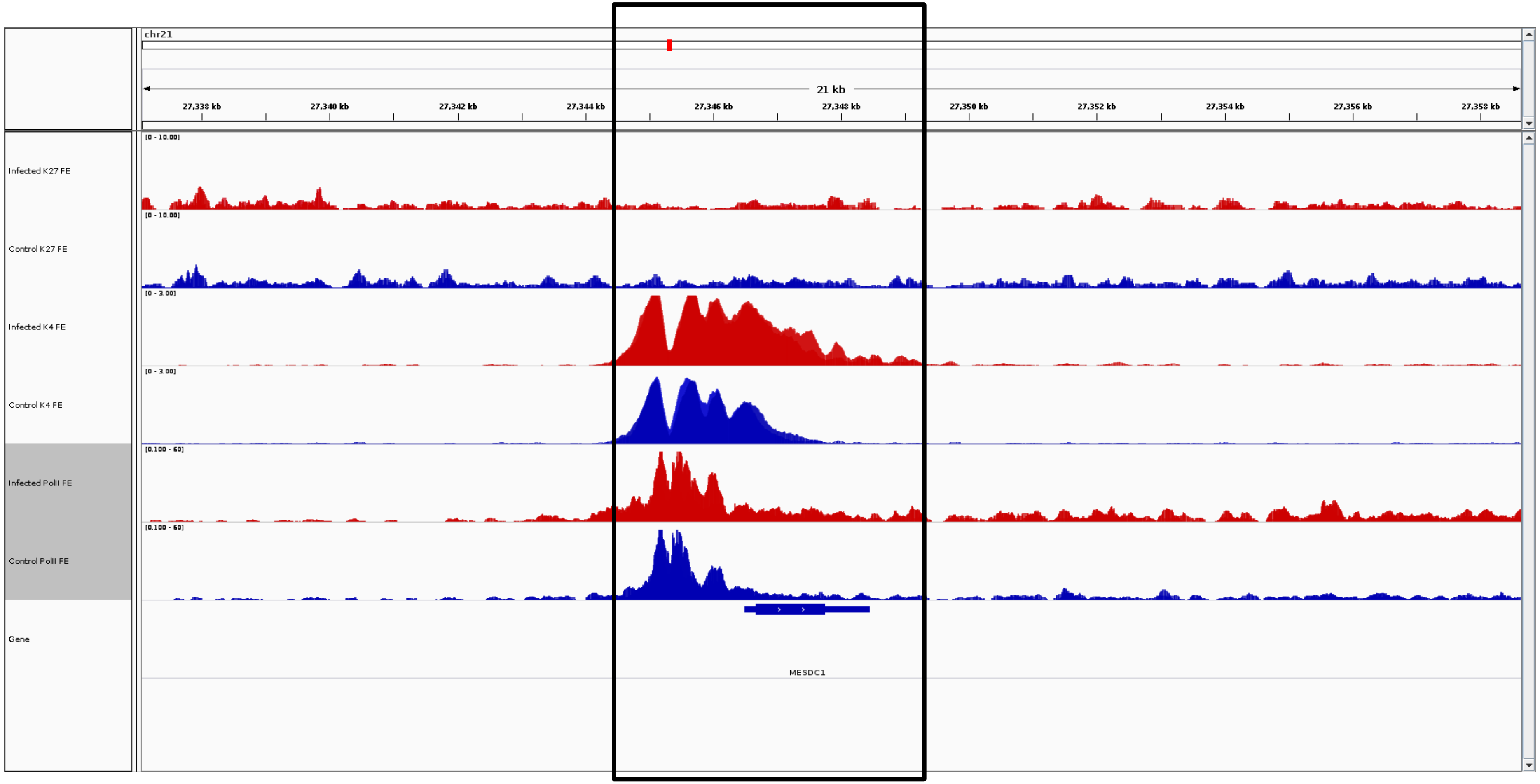
chr21
21 kb
27,338 kb
27,340 kb
27,342 kb
27,344 kb
27,346 kb
27,348 kb
27,350 kb
27,352 kb
27,354 kb
27,356 kb
27,358 kb
Infected K27 FE
[0 - 10.00]
Control K27 FE
[0 - 10.00]
Infected K4 FE
[0 - 3.00]
Control K4 FE
[0 - 3.00]
Infected PolII FE
[0.100 - 60]
Control PolII FE
[0.100 - 60]
Gene
MESDC1

***MESDC1***

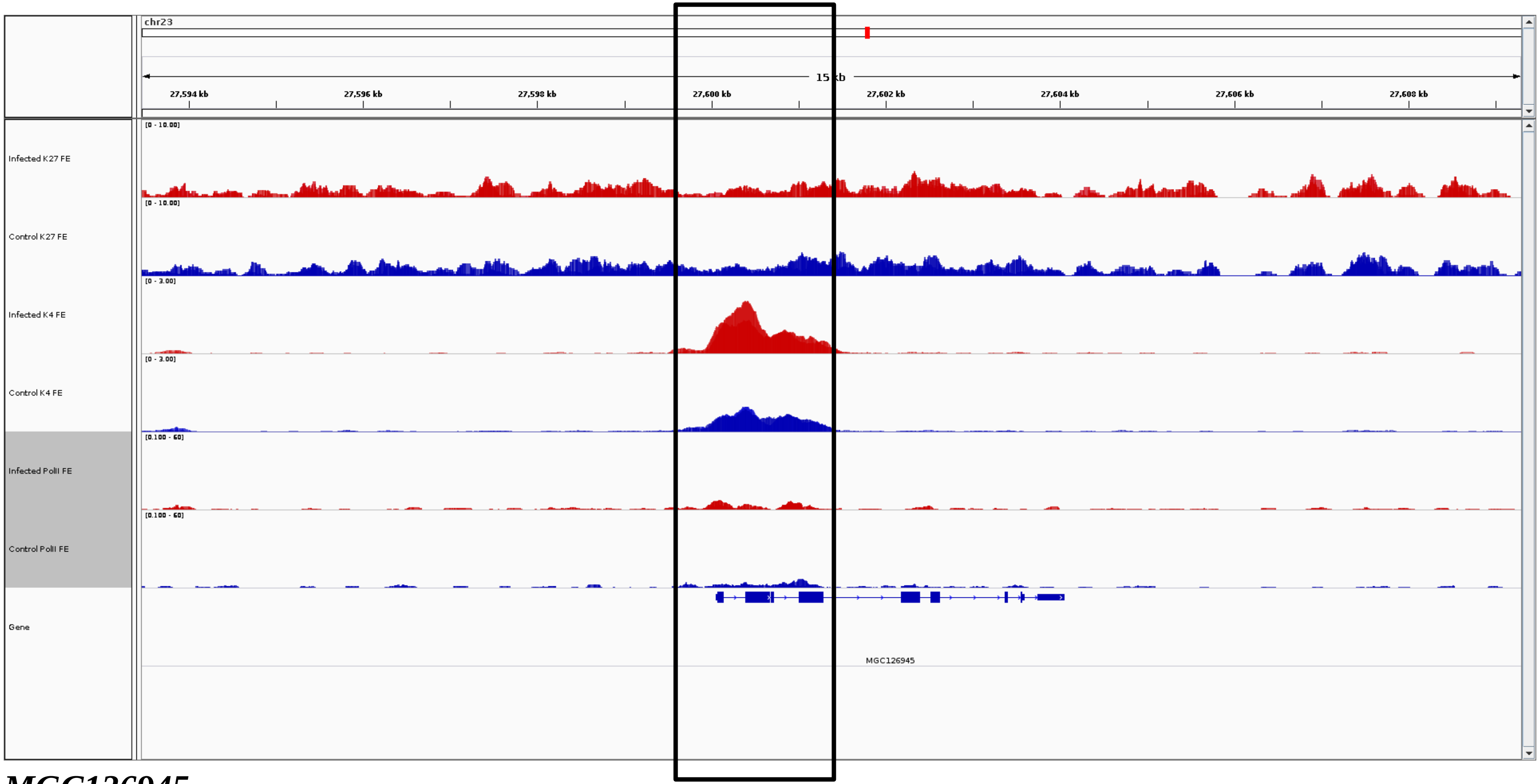
chr23
15 kb
27,594 kb
27,596 kb
27,598 kb
27,600 kb
27,602 kb
27,604 kb
27,606 kb
27,608 kb
Infected K27 FE
[0 - 10.00]
Control K27 FE
[0 - 10.00]
Infected K4 FE
[0 - 3.00]
Control K4 FE
[0 - 3.00]
Infected PolII FE
[0.100 - 60]
Control PolII FE
[0.100 - 60]
Gene
MGC126945

***MGC126945***

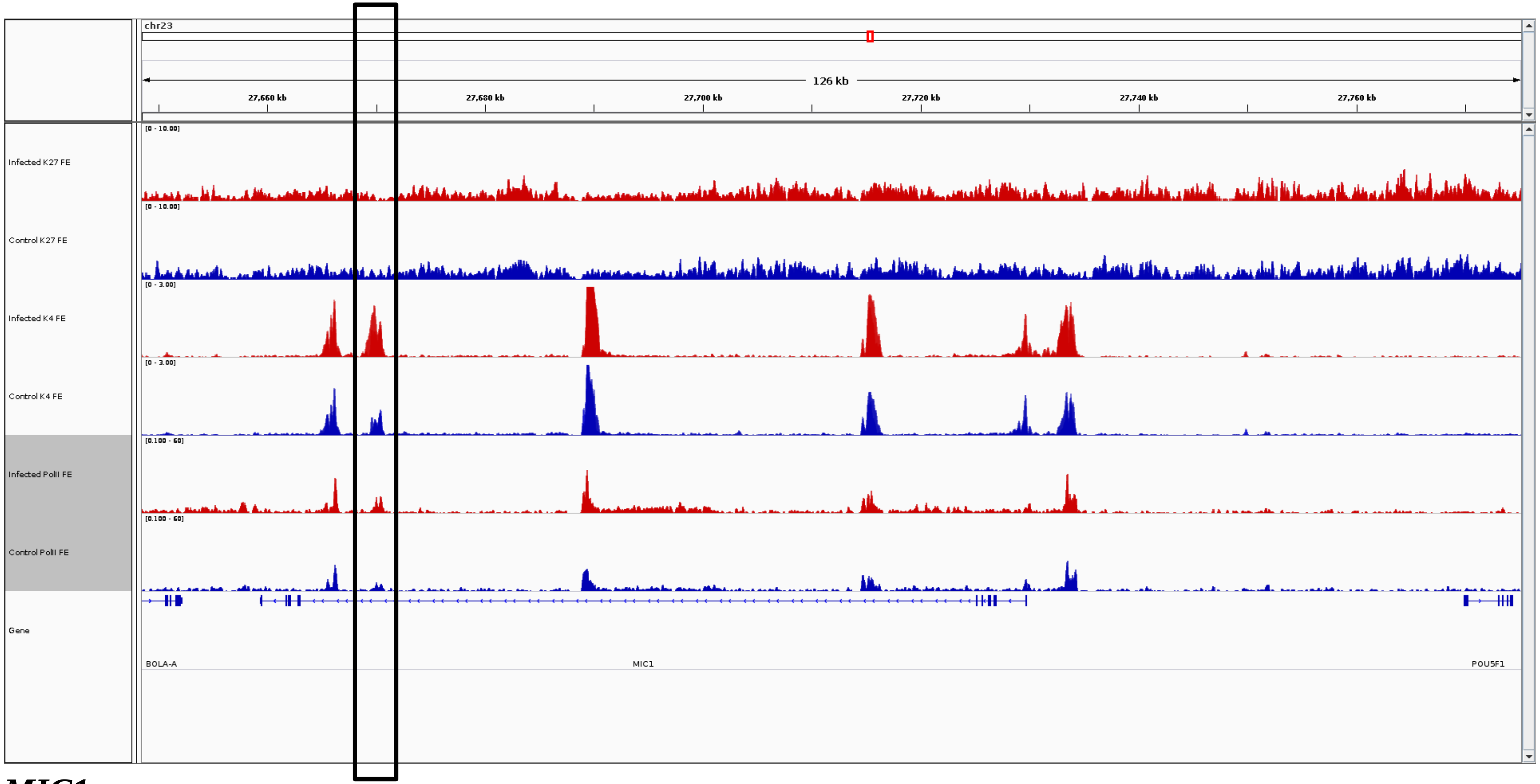
chr23
126 kb
27,660 kb
27,680 kb
27,700 kb
27,720 kb
27,740 kb
27,760 kb
Infected K27 FE
Control K27 FE
Infected K4 FE
Control K4 FE
Infected PolII FE
Control PolII FE
Gene
BOLA-A
MIC1
POU5F1

***MIC1***

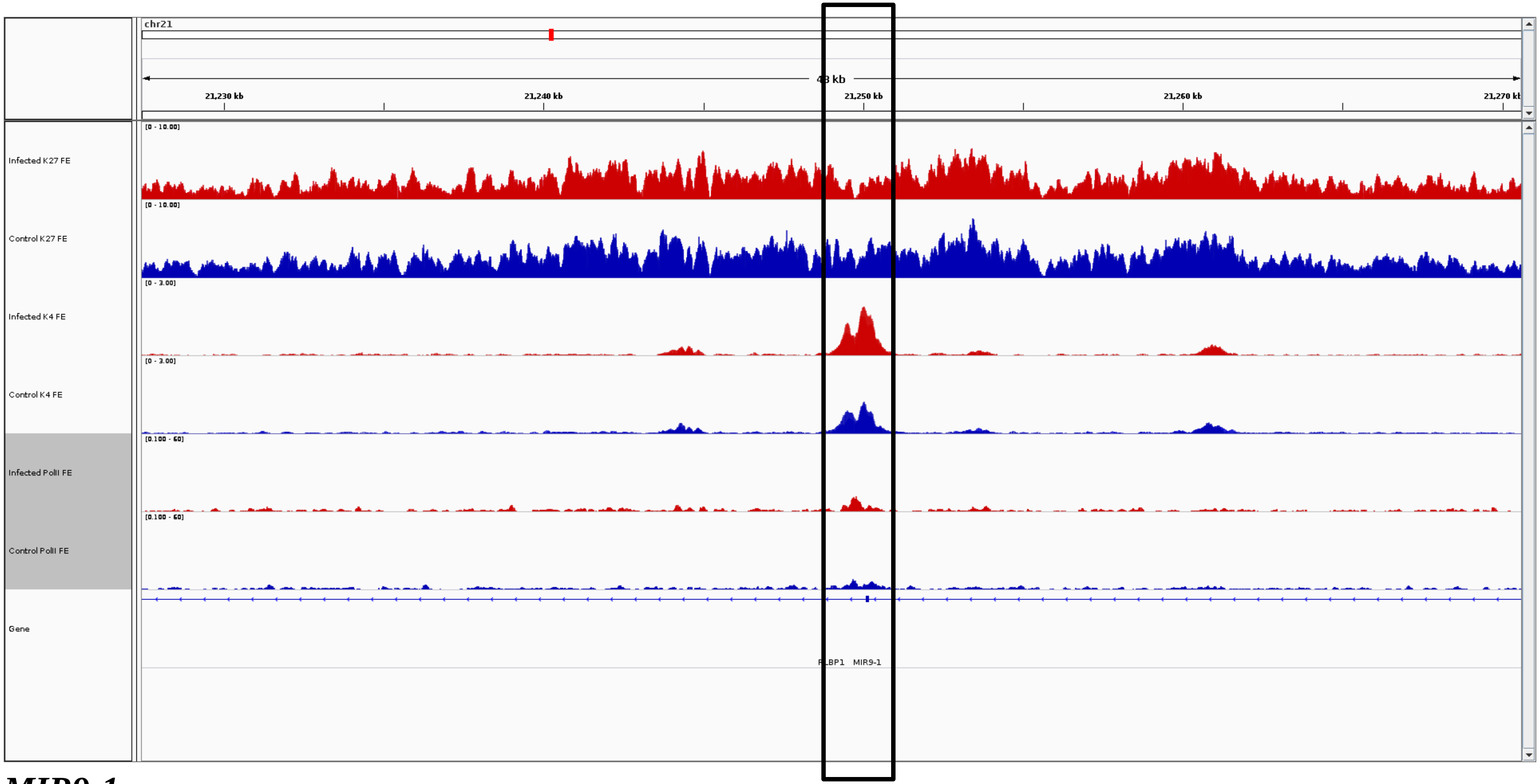
chr21
48 kb
21,230 kb
21,240 kb
21,250 kb
21,260 kb
21,270 kb
Infected K27 FE
[0 - 10.00]
Control K27 FE
[0 - 10.00]
Infected K4 FE
[0 - 3.00]
Control K4 FE
[0 - 3.00]
Infected PolII FE
[0.100 - 60]
Control PolII FE
[0.100 - 60]
Gene
MIR9-1

***MIR9-1***

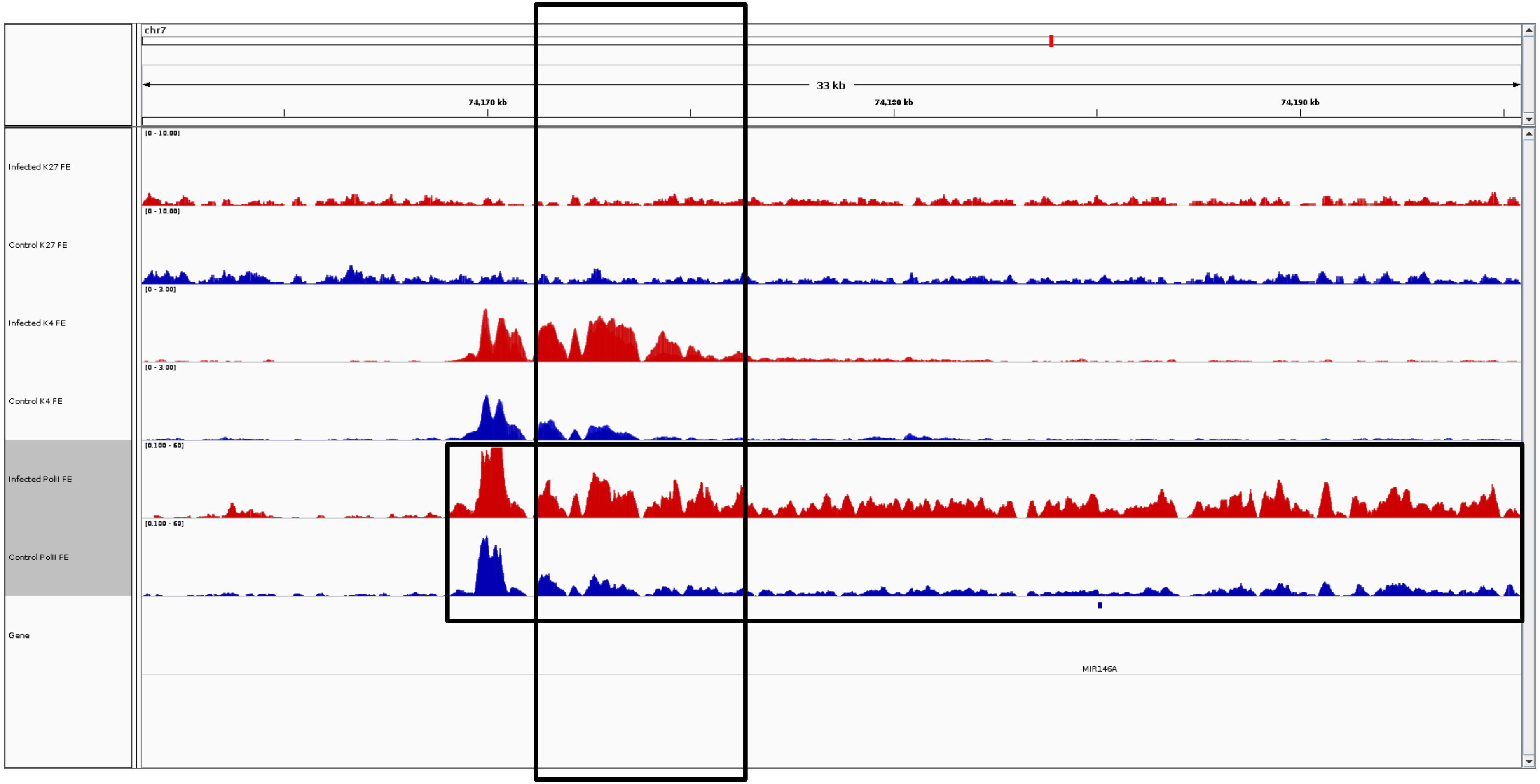
chr7
33 kb
74,170 kb
74,180 kb
74,190 kb
Infected K27 FE
Control K27 FE
Infected K4 FE
Control K4 FE
Infected PolII FE
Control PolII FE
Gene
MIR146A

***MIR146A***

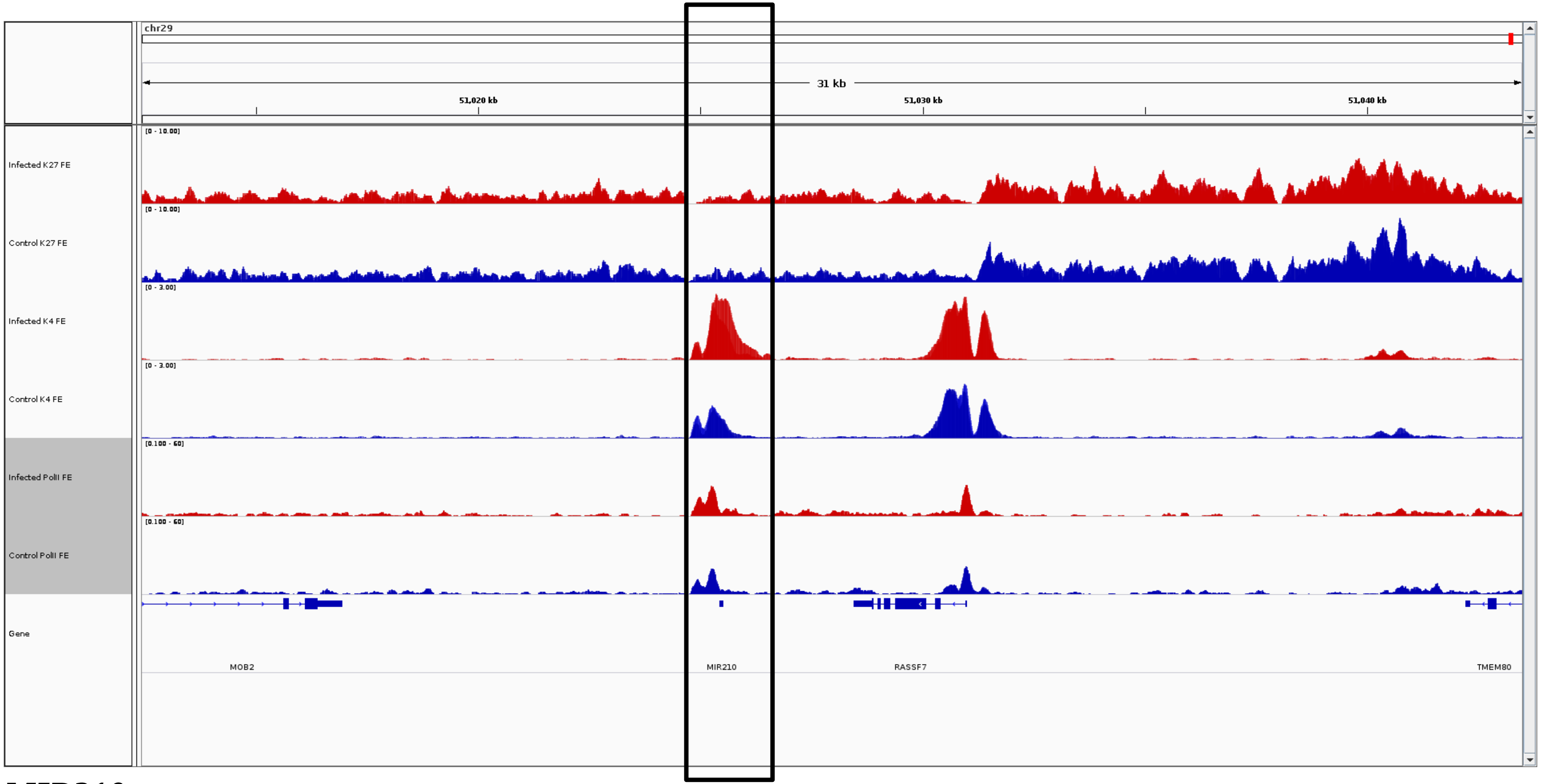
chr29
31 kb
51,020 kb
51,030 kb
51,040 kb
Infected K27 FE
[0 - 10.00]
Control K27 FE
[0 - 10.00]
Infected K4 FE
[0 - 3.00]
Control K4 FE
[0 - 3.00]
Infected PolII FE
[0.100 - 60]
Control PolII FE
[0.100 - 60]
Gene
MOB2
MIR210
RASSF7
TMEM80

***MIR210***

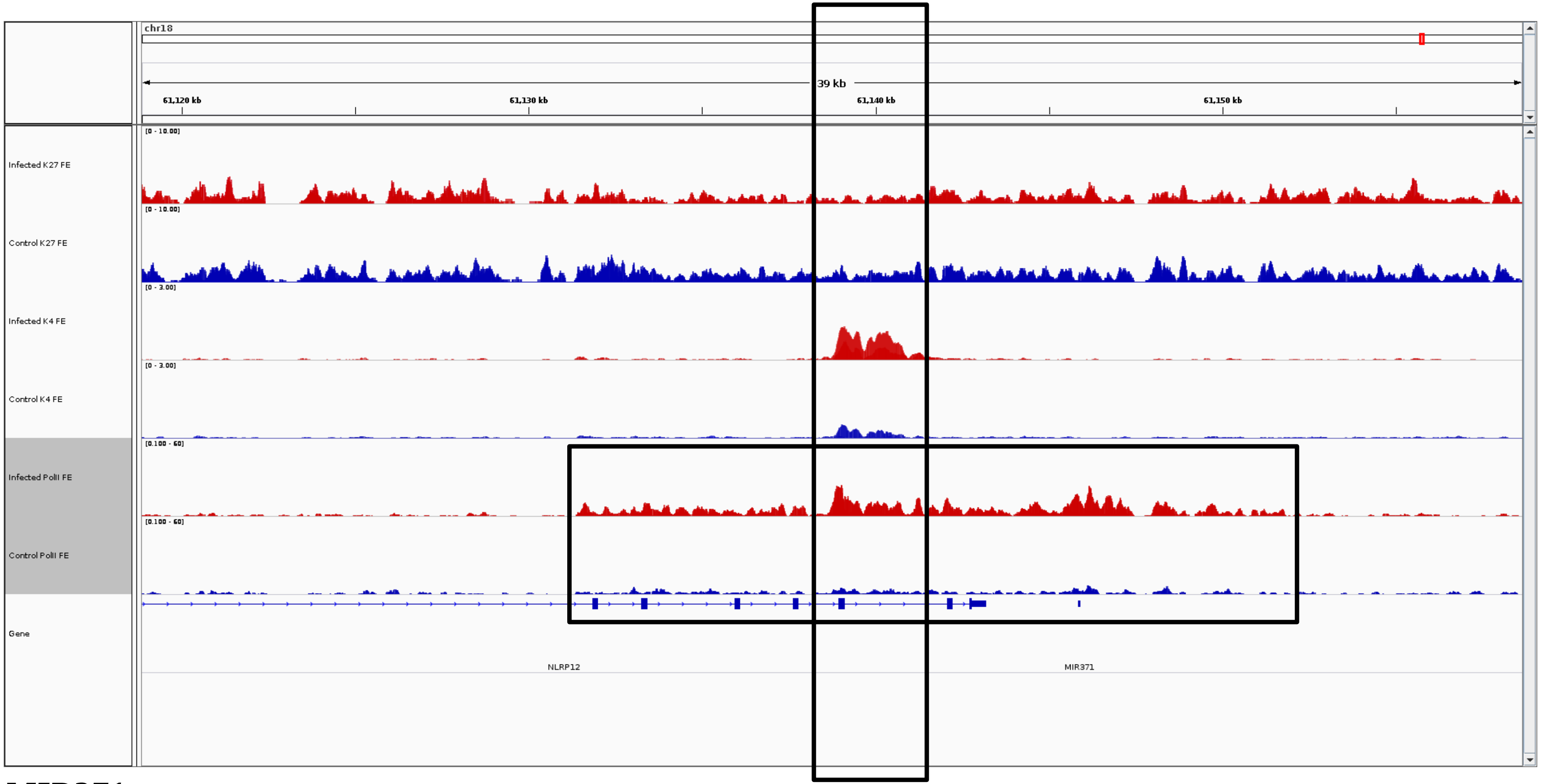
chr18
39 kb
61,120 kb
61,130 kb
61,140 kb
61,150 kb
Infected K27 FE
[0 - 10.00]
Control K27 FE
[0 - 10.00]
Infected K4 FE
[0 - 3.00]
Control K4 FE
[0 - 3.00]
Infected PolII FE
[0.100 - 60]
Control PolII FE
[0.100 - 60]
Gene
NLRP12
MIR371

***MIR371***

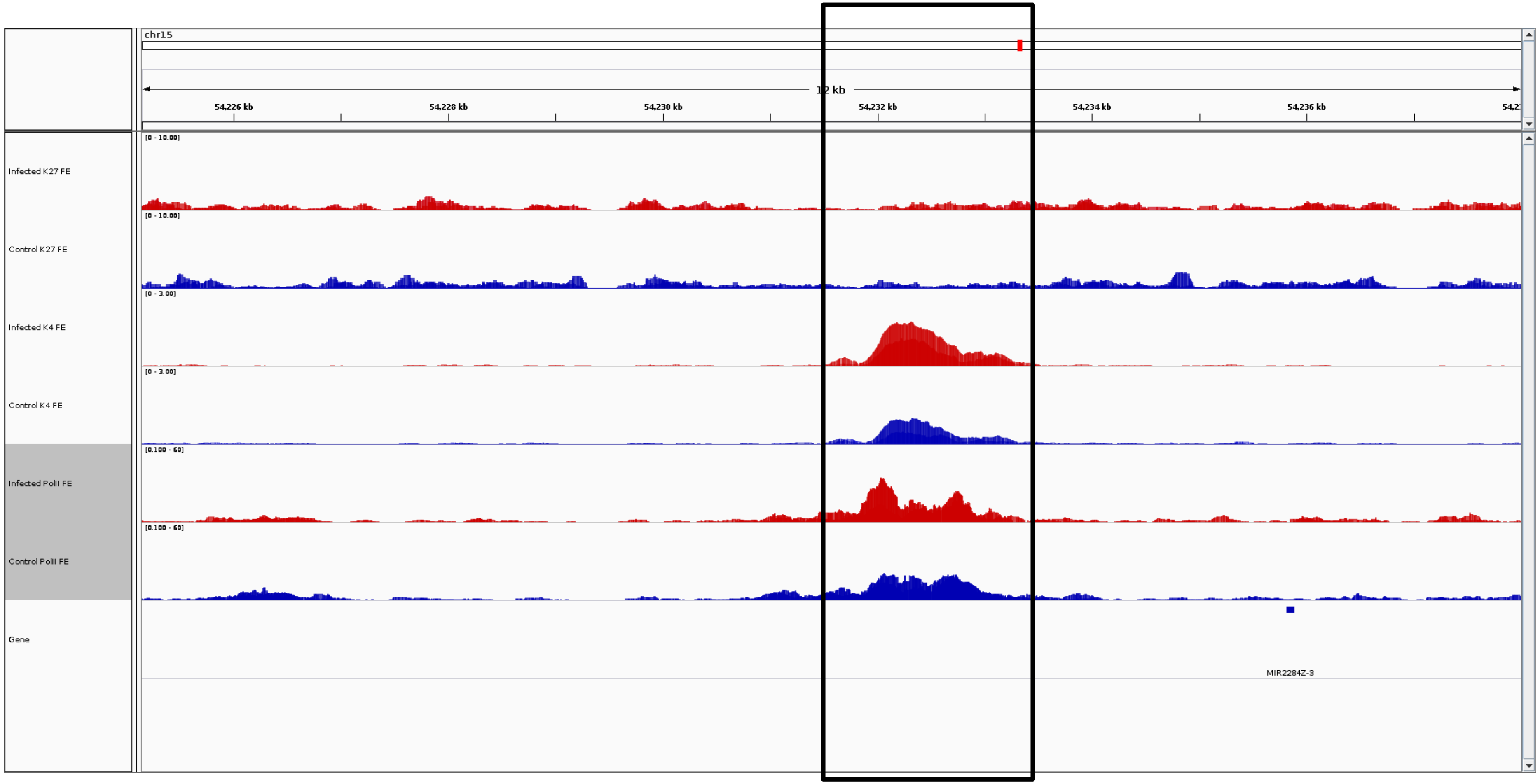
chr15
12 kb
54,226 kb
54,228 kb
54,230 kb
54,232 kb
54,234 kb
54,236 kb
Infected K27 FE
[0 - 10.00]
Control K27 FE
[0 - 10.00]
Infected K4 FE
[0 - 3.00]
Control K4 FE
[0 - 3.00]
Infected PolII FE
[0.100 - 60]
Control PolII FE
[0.100 - 60]
Gene
MIR2284Z-3

***MIR2284Z-3***

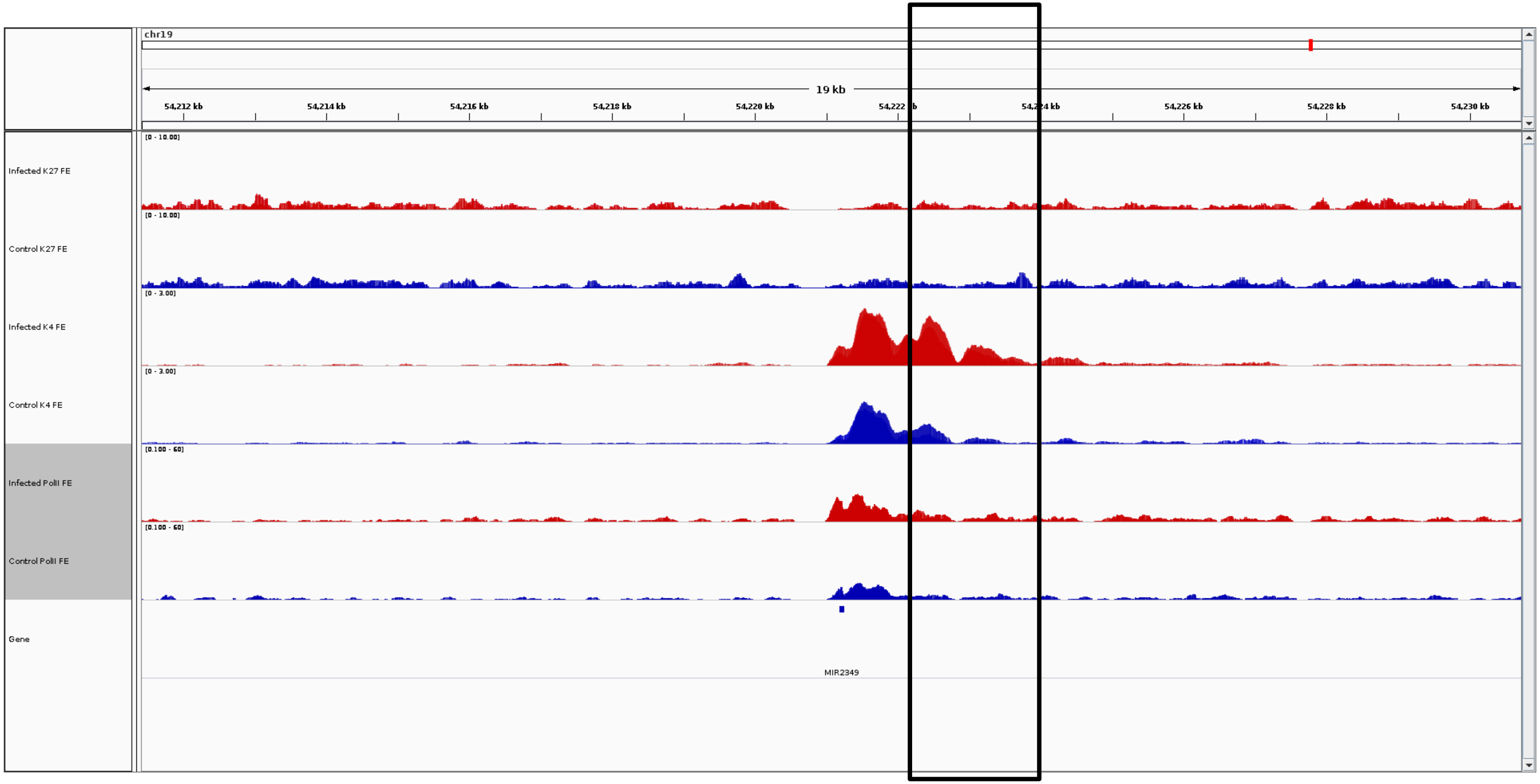
chr19
19 kb
54,212 kb
54,214 kb
54,216 kb
54,218 kb
54,220 kb
54,226 kb
54,228 kb
54,230 kb
Infected K27 FE
Control K27 FE
Infected K4 FE
Control K4 FE
Infected PolII FE
Control PolII FE
Gene
MIR2349

***MIR2349***

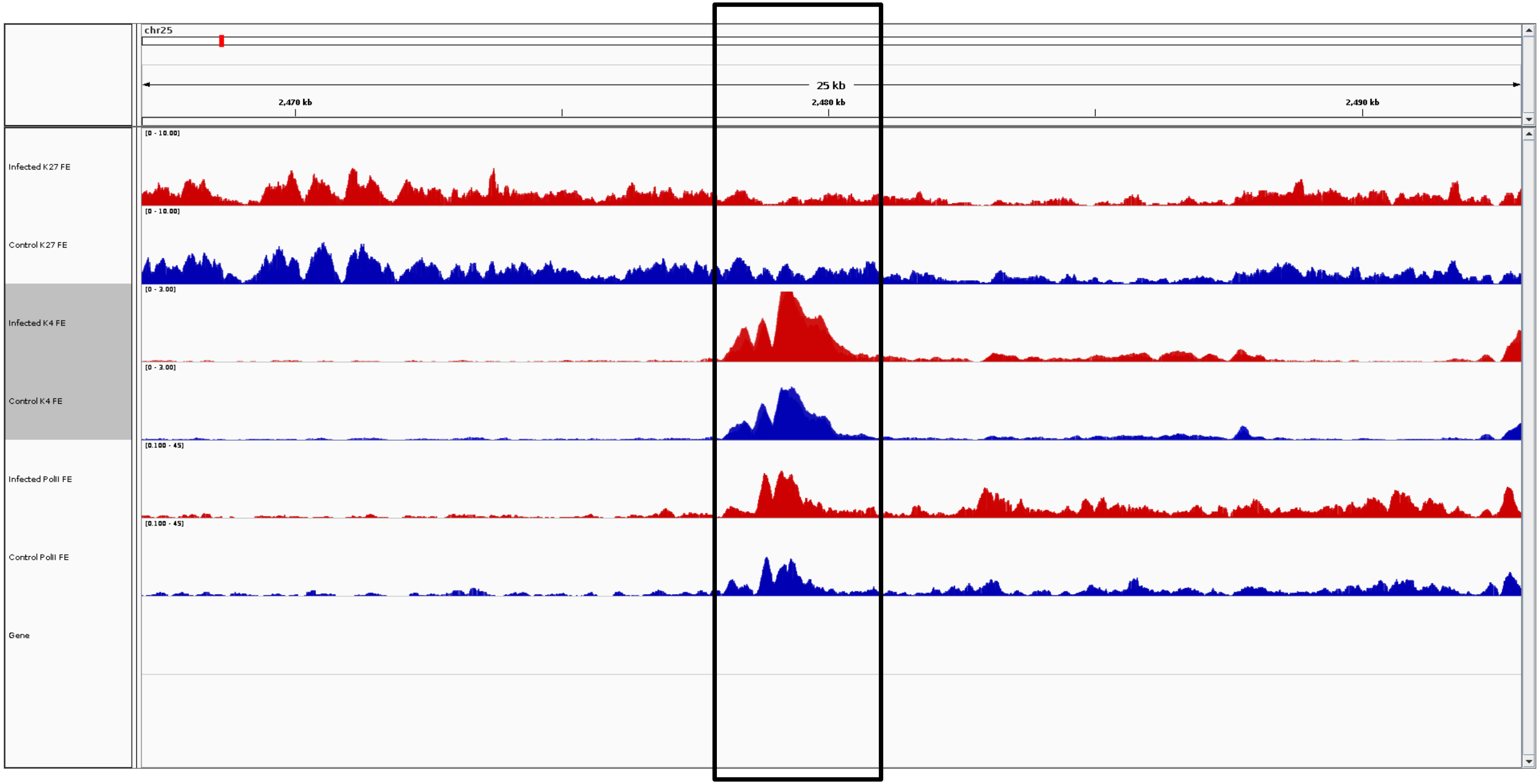
chr25
25 kb
2,470 kb
2,480 kb
2,490 kb
[0 - 10.00]
Infected K27 FE
[0 - 10.00]
Control K27 FE
[0 - 3.00]
Infected K4 FE
[0 - 3.00]
Control K4 FE
[0.100 - 45]
Infected PolII FE
[0.100 - 45]
Control PolII FE
Gene

***MMP25***

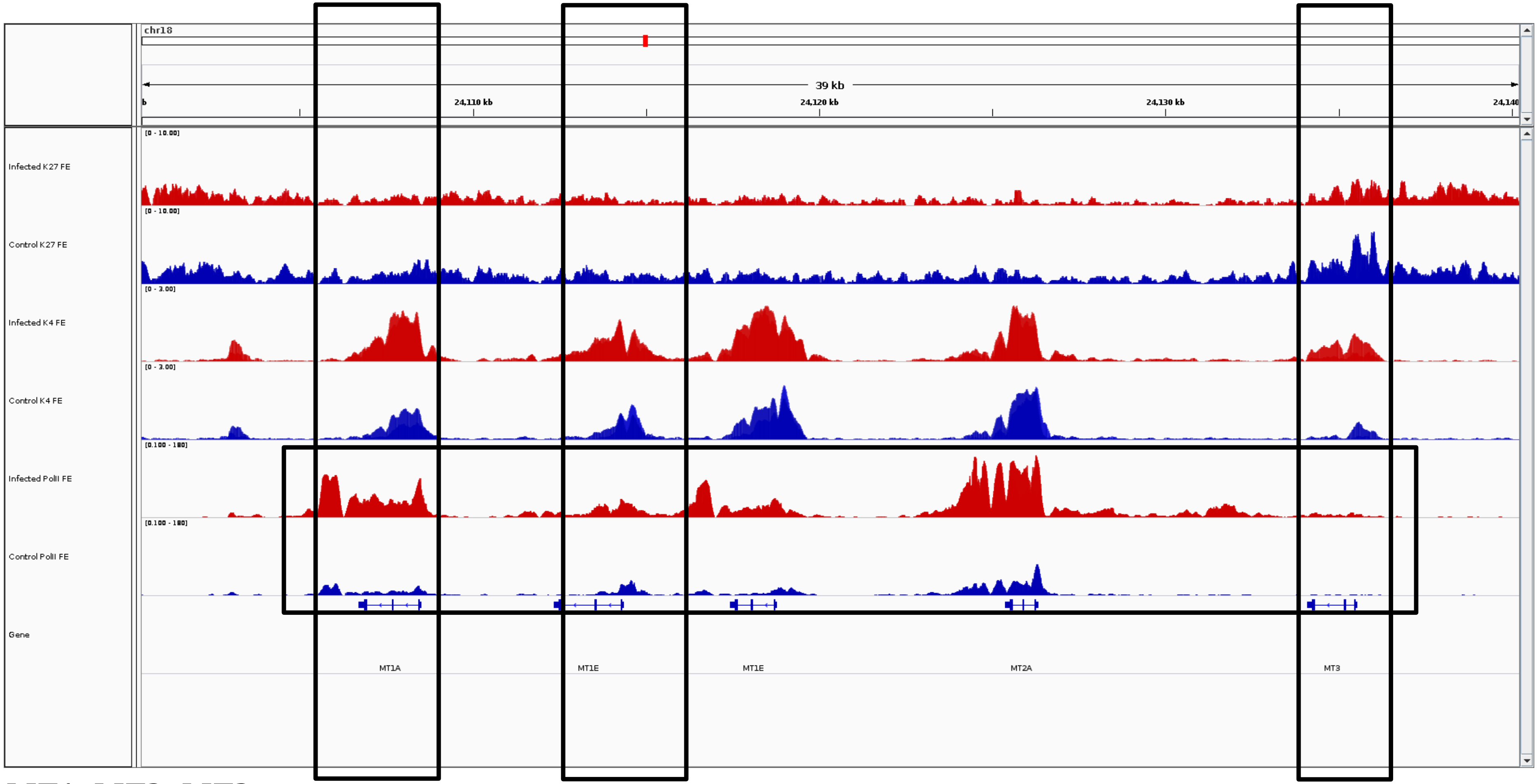
chr18
39 kb
24,110 kb
24,120 kb
24,130 kb
Infected K27 FE
[0 - 10.00]
Control K27 FE
[0 - 10.00]
Infected K4 FE
[0 - 3.00]
Control K4 FE
[0 - 3.00]
Infected PolII FE
[0.100 - 180]
Control PolII FE
[0.100 - 180]
Gene
MT1A
MT1E
MT1E
MT2A
MT3

***MT1, MT2, MT3***

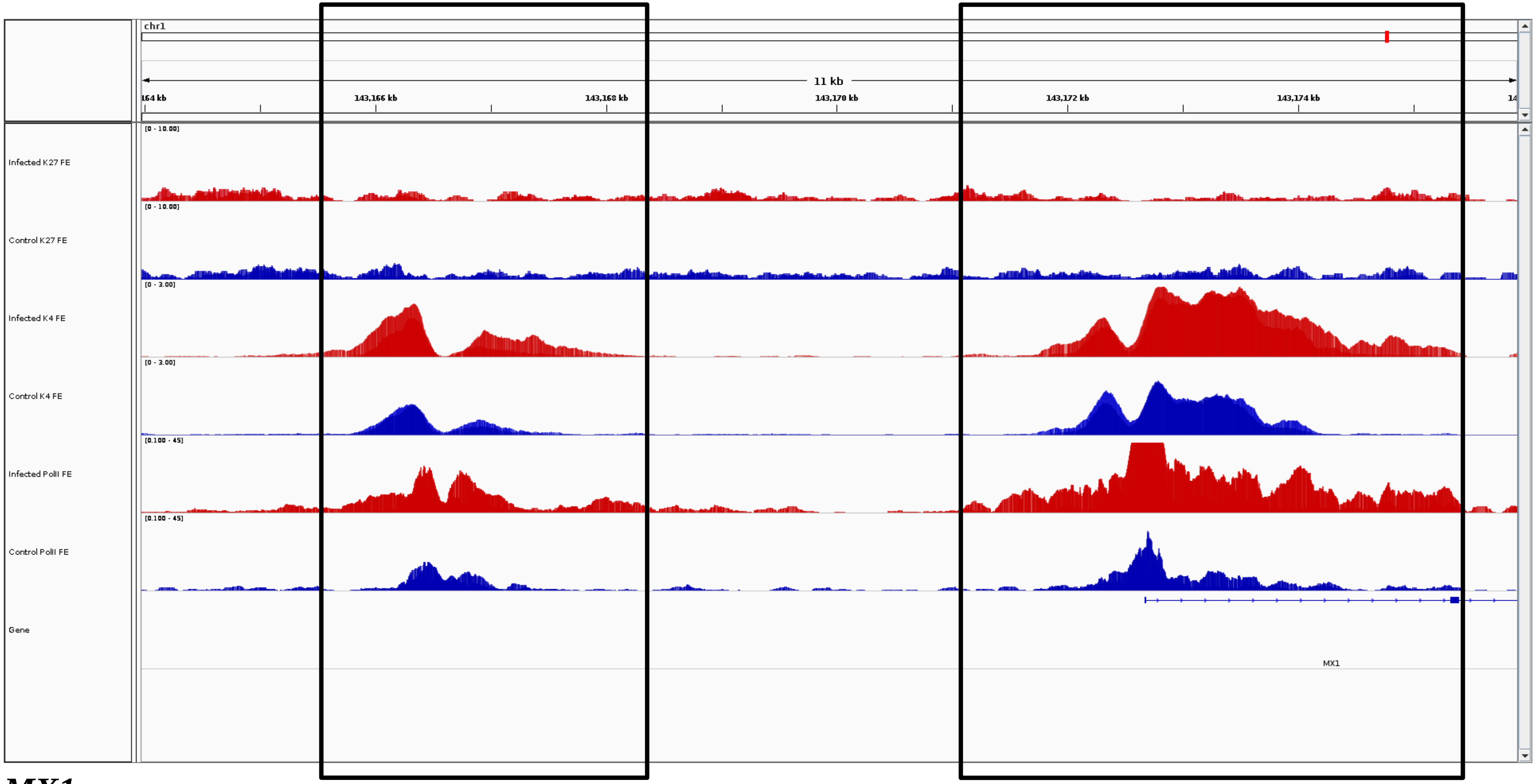
chr1
11 kb
164 kb
143,166 kb
143,168 kb
143,170 kb
143,172 kb
143,174 kb
Infected K27 FE
[0 - 10.00]
Control K27 FE
[0 - 10.00]
Infected K4 FE
[0 - 3.00]
Control K4 FE
[0 - 3.00]
Infected PolII FE
[0.100 - 45]
Control PolII FE
[0.100 - 45]
Gene
MX1

***MX1***

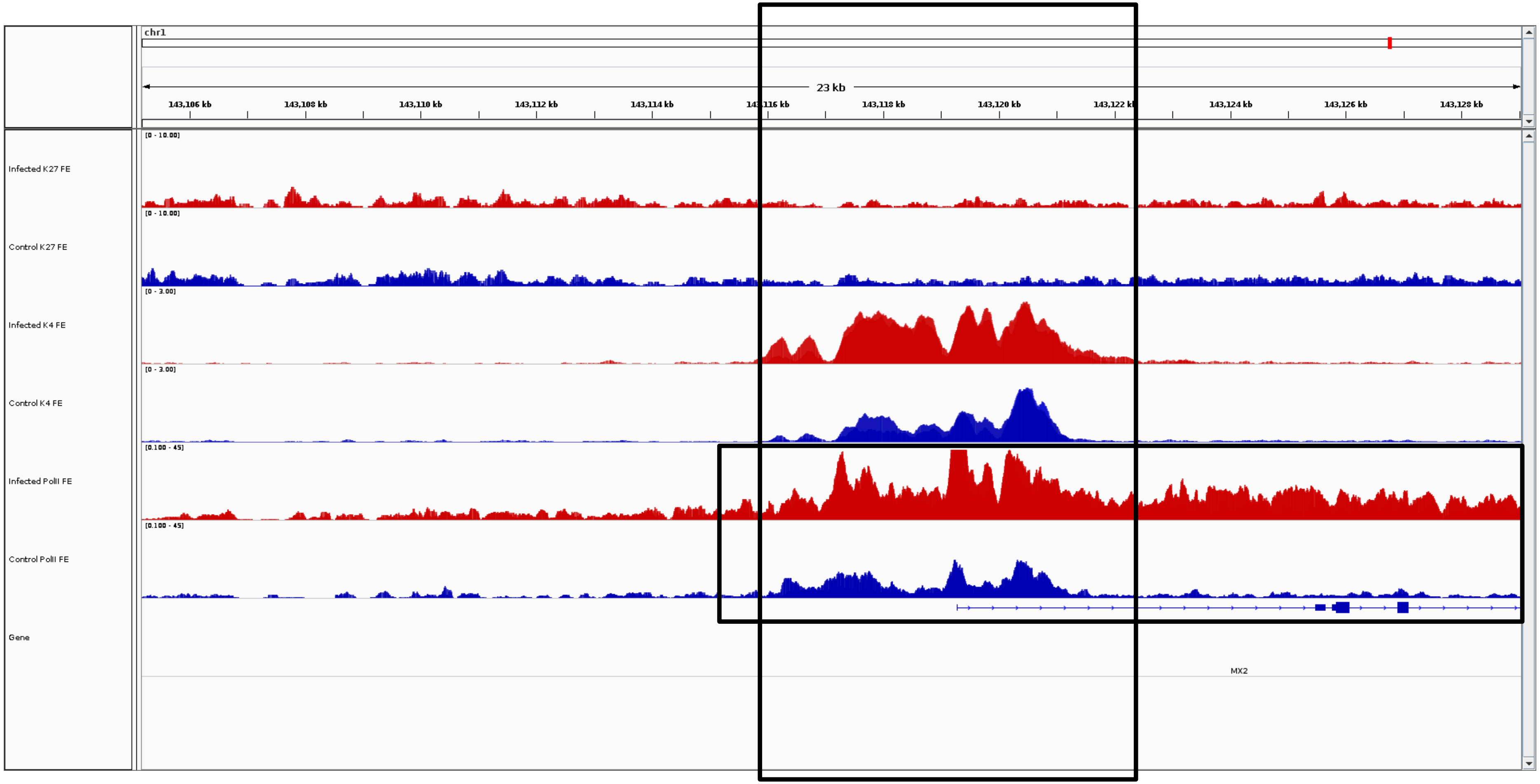
chr1
23 kb
143,106 kb
143,108 kb
143,110 kb
143,112 kb
143,114 kb
143,118 kb
143,120 kb
143,124 kb
143,126 kb
143,128 kb
Infected K27 FE
[0 - 10.00]
Control K27 FE
[0 - 10.00]
Infected K4 FE
[0 - 3.00]
Control K4 FE
[0 - 3.00]
Infected PolII FE
[0.100 - 45]
Control PolII FE
[0.100 - 45]
Gene
MX2

***MX2***

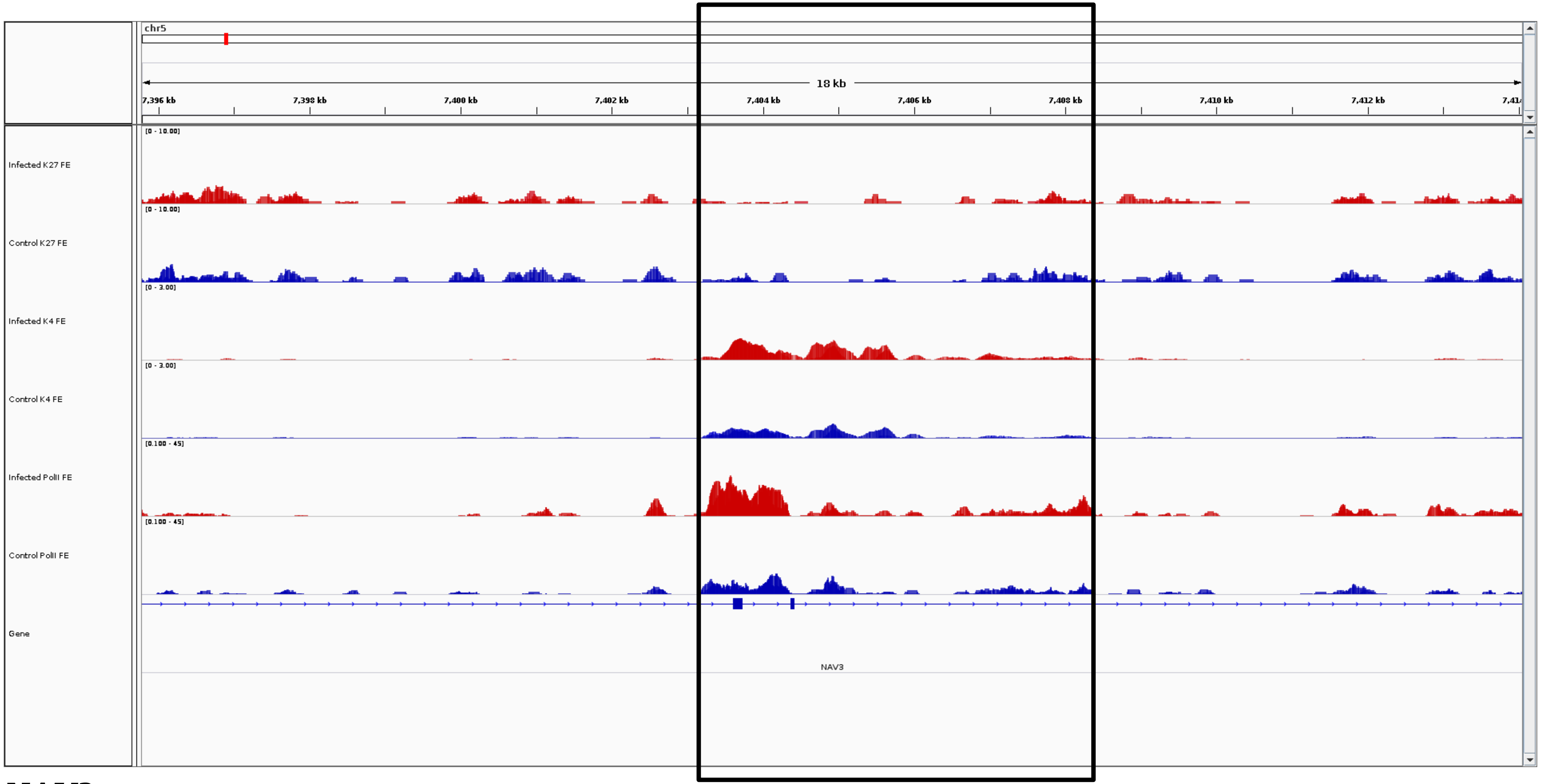
chr5
18 kb
7,396 kb
7,398 kb
7,400 kb
7,402 kb
7,404 kb
7,406 kb
7,408 kb
7,410 kb
7,412 kb
Infected K27 FE
[0 - 10.00]
Control K27 FE
[0 - 10.00]
Infected K4 FE
[0 - 3.00]
Control K4 FE
[0 - 3.00]
Infected PolII FE
[0.100 - 45]
Control PolII FE
[0.100 - 45]
Gene
NAV3

***NAV3***

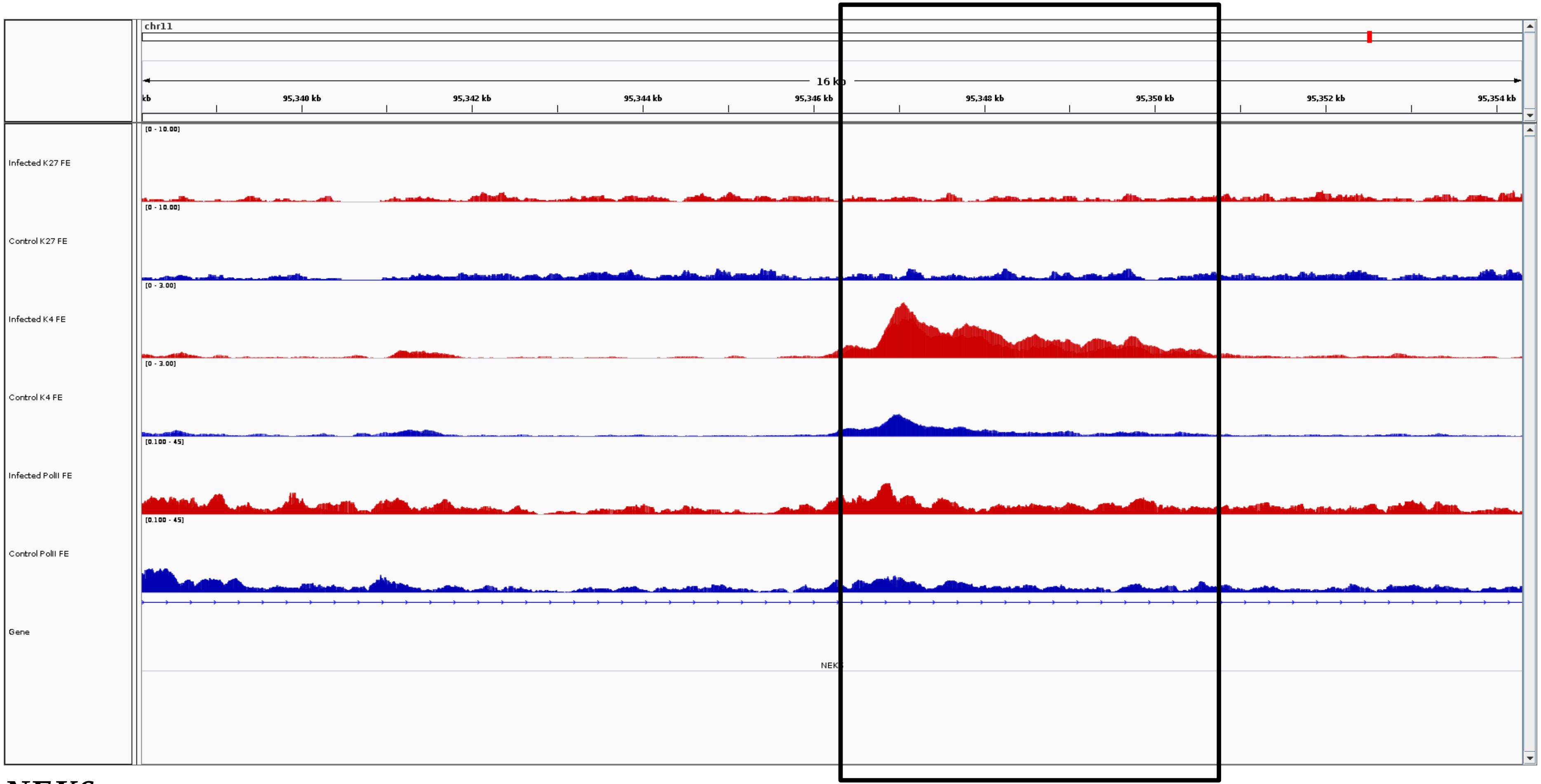
chr11
16 kb
95,340 kb
95,342 kb
95,344 kb
95,346 kb
95,348 kb
95,350 kb
95,352 kb
95,354 kb
Infected K27 FE
[0 - 10.00]
Control K27 FE
[0 - 10.00]
Infected K4 FE
[0 - 3.00]
Control K4 FE
[0 - 3.00]
Infected PolII FE
[0.100 - 45]
Control PolII FE
[0.100 - 45]
Gene

***NEK6***

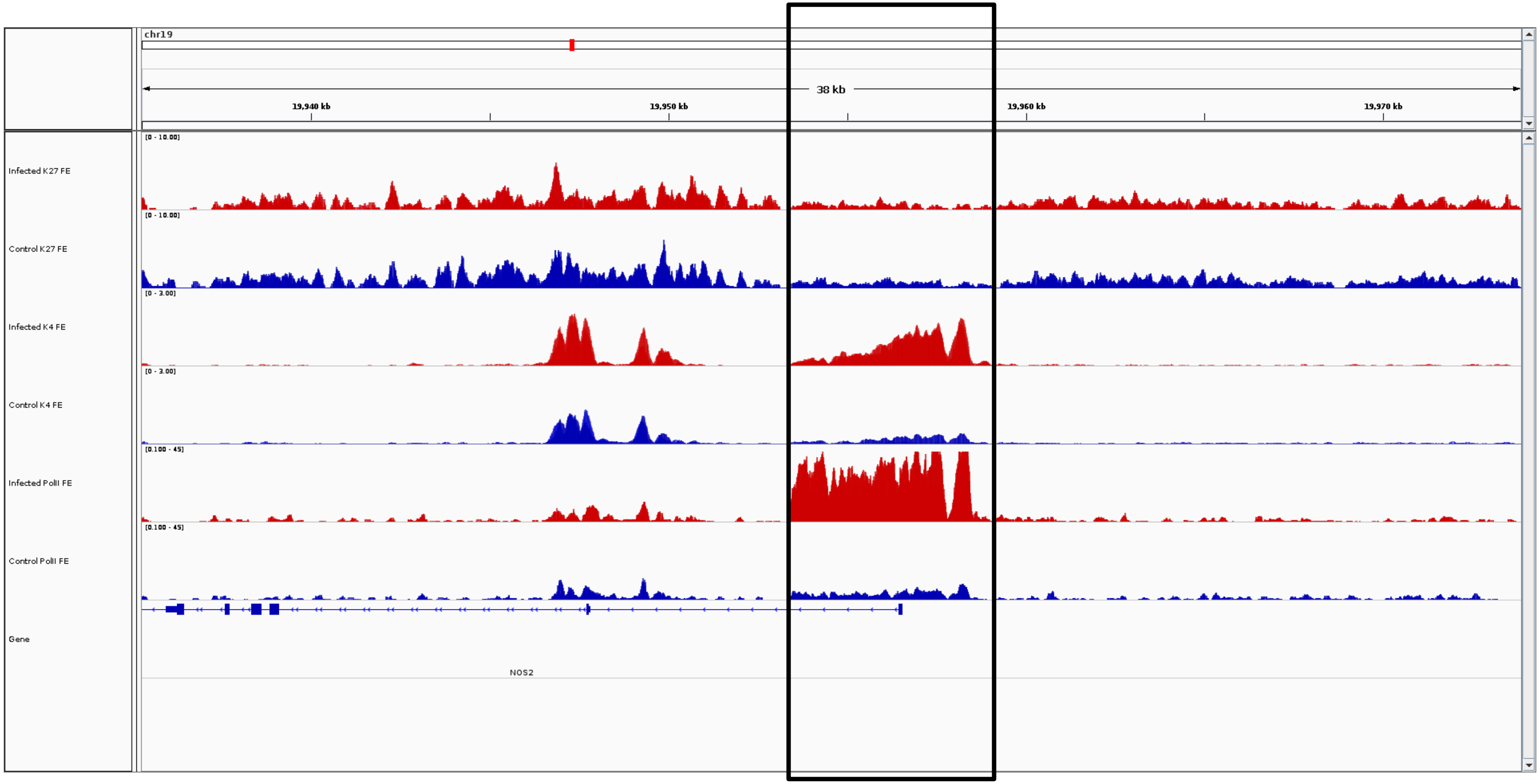
chr19
38 kb
19,940 kb
19,950 kb
19,960 kb
19,970 kb
Infected K27 FE
Control K27 FE
Infected K4 FE
Control K4 FE
Infected PolII FE
Control PolII FE
Gene
NOS2

***NOS2***

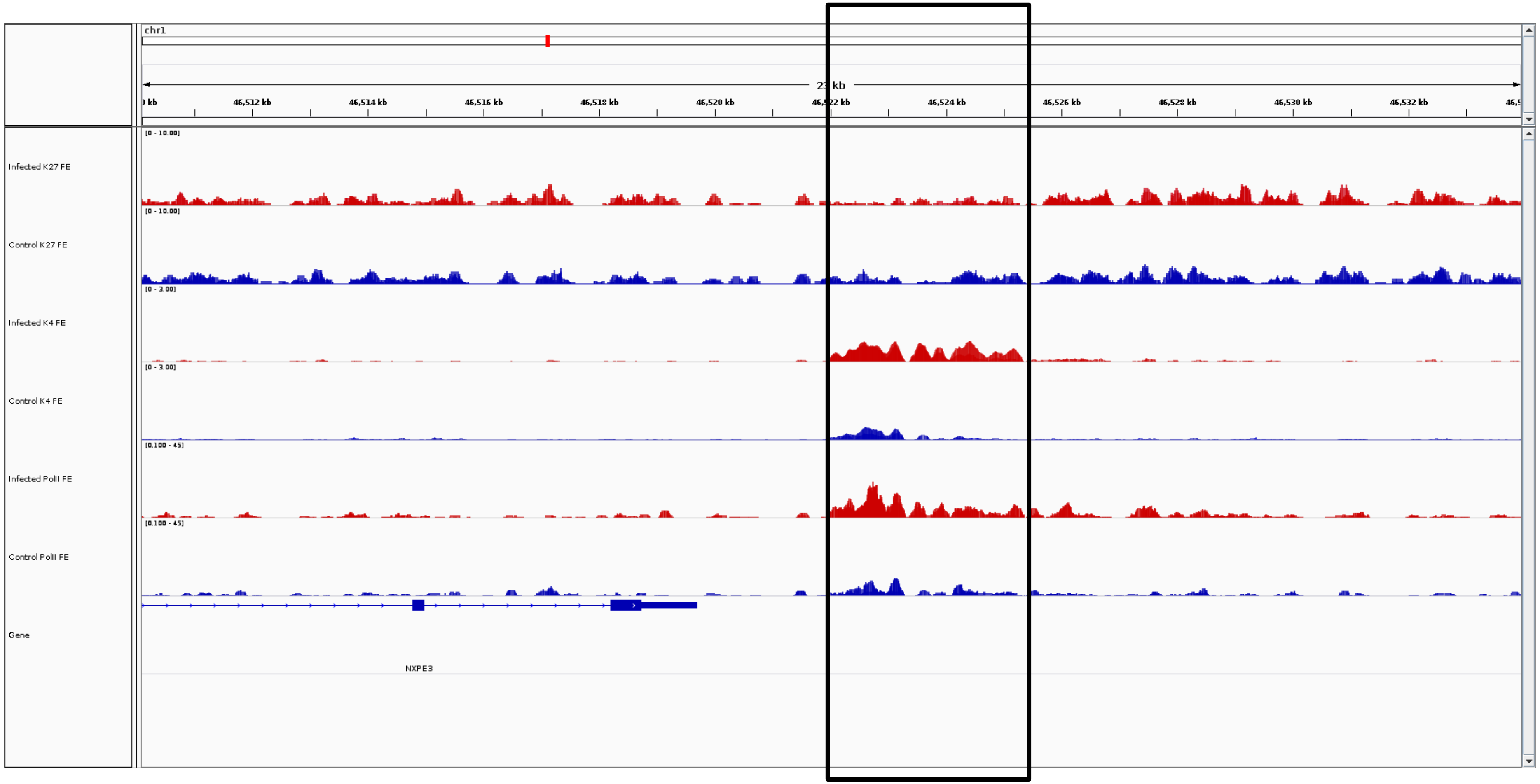
chr1
23 kb
46,512 kb
46,514 kb
46,516 kb
46,518 kb
46,520 kb
46,524 kb
46,526 kb
46,528 kb
46,530 kb
46,532 kb
Infected K27 FE
[0 - 10.00]
Control K27 FE
[0 - 10.00]
Infected K4 FE
[0 - 3.00]
Control K4 FE
[0 - 3.00]
Infected PolII FE
[0.100 - 45]
Control PolII FE
[0.100 - 45]
Gene
NXPE3

***NXPE3***

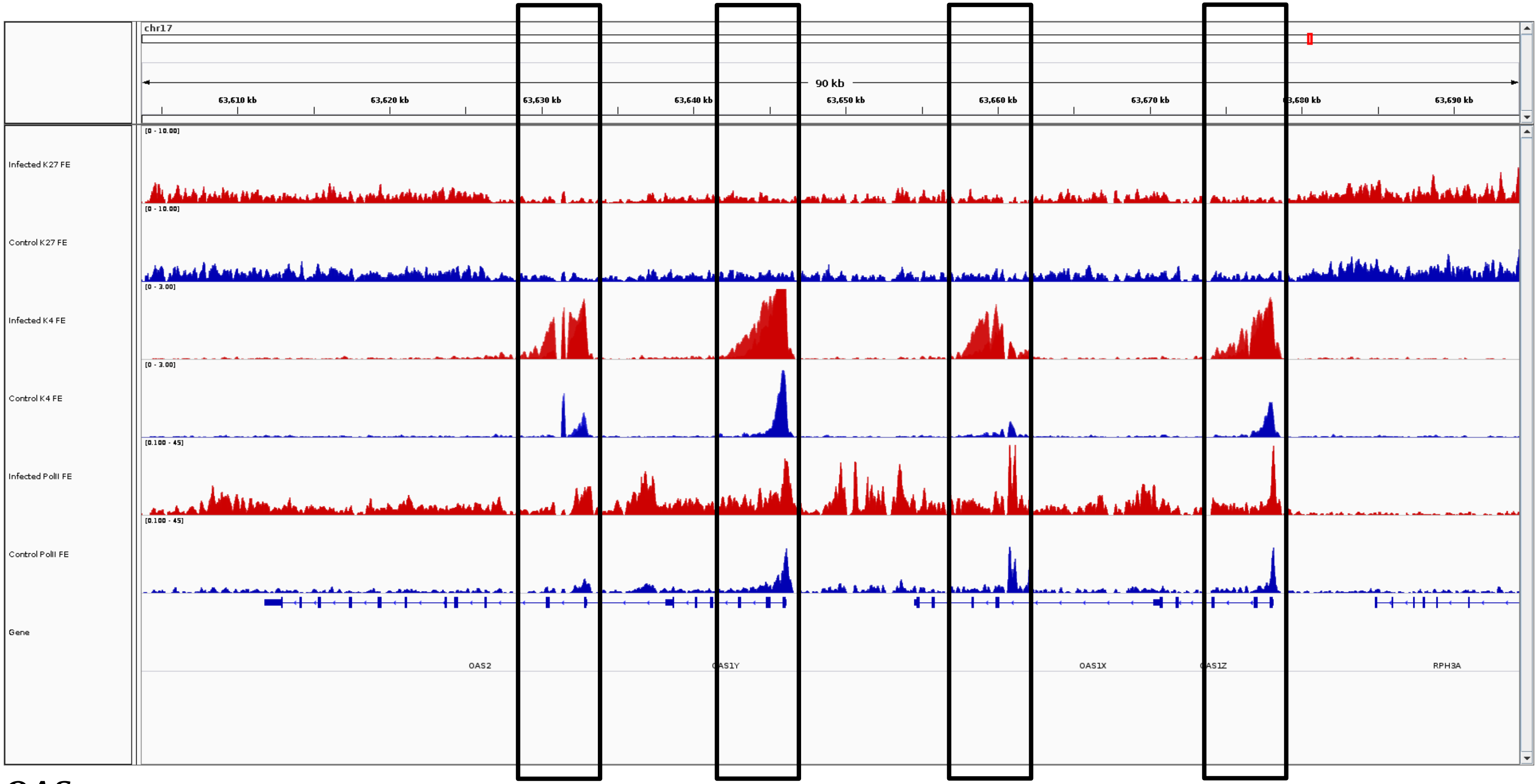
chr17
90 kb
63,610 kb
63,620 kb
63,630 kb
63,640 kb
63,650 kb
63,660 kb
63,670 kb
63,690 kb
Infected K27 FE
Control K27 FE
Infected K4 FE
Control K4 FE
Infected PolII FE
Control PolII FE
Gene
OAS2
OAS1X
RPH3A

### *OAS*

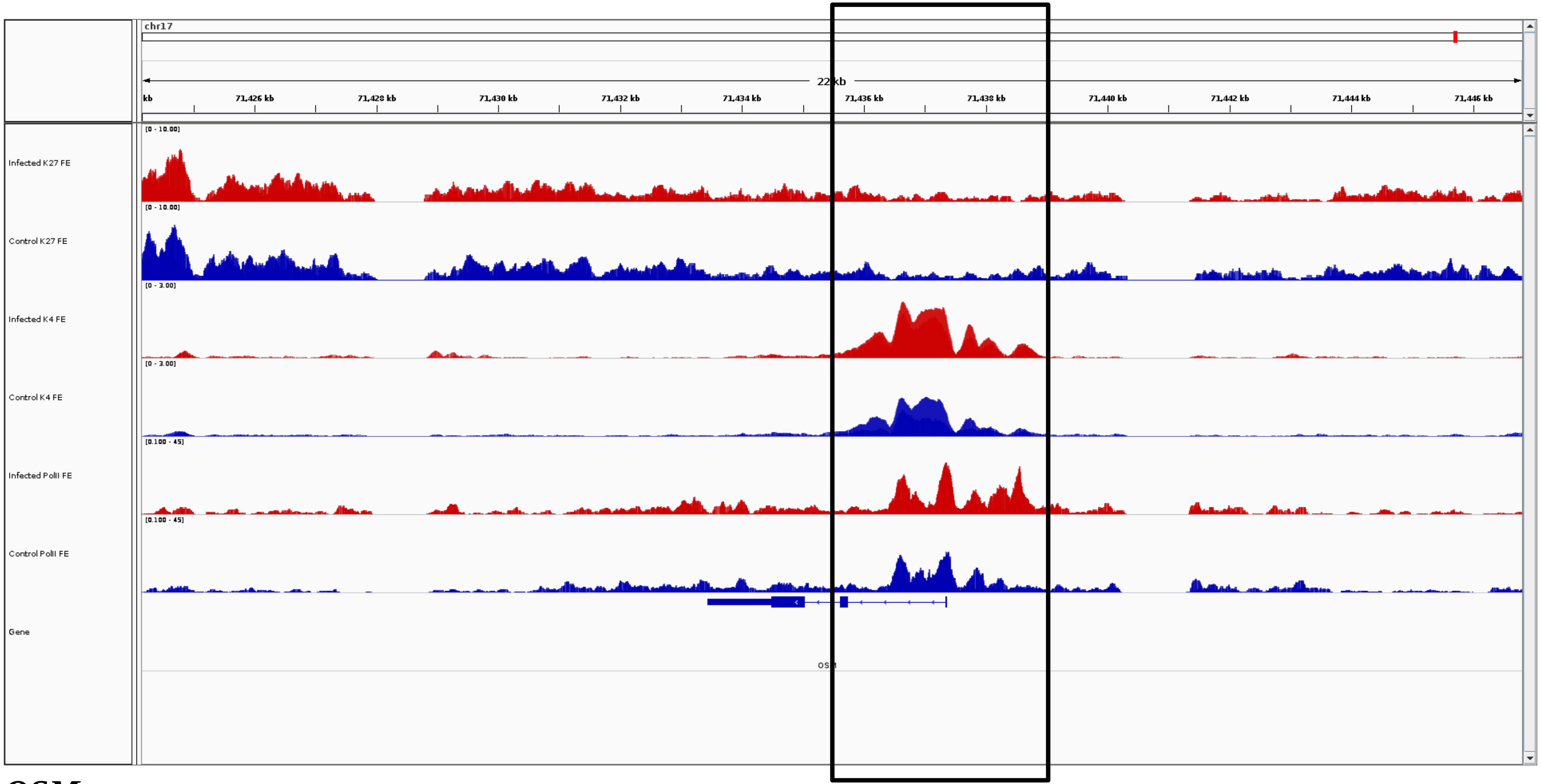
chr17
22 kb
71,426 kb
71,428 kb
71,430 kb
71,432 kb
71,434 kb
71,436 kb
71,438 kb
71,440 kb
71,442 kb
71,444 kb
71,446 kb
Infected K27 FE
[0 - 10.00]
Control K27 FE
[0 - 10.00]
Infected K4 FE
[0 - 3.00]
Control K4 FE
[0 - 3.00]
Infected PolII FE
[0.100 - 45]
Control PolII FE
[0.100 - 45]
Gene

***OSM***

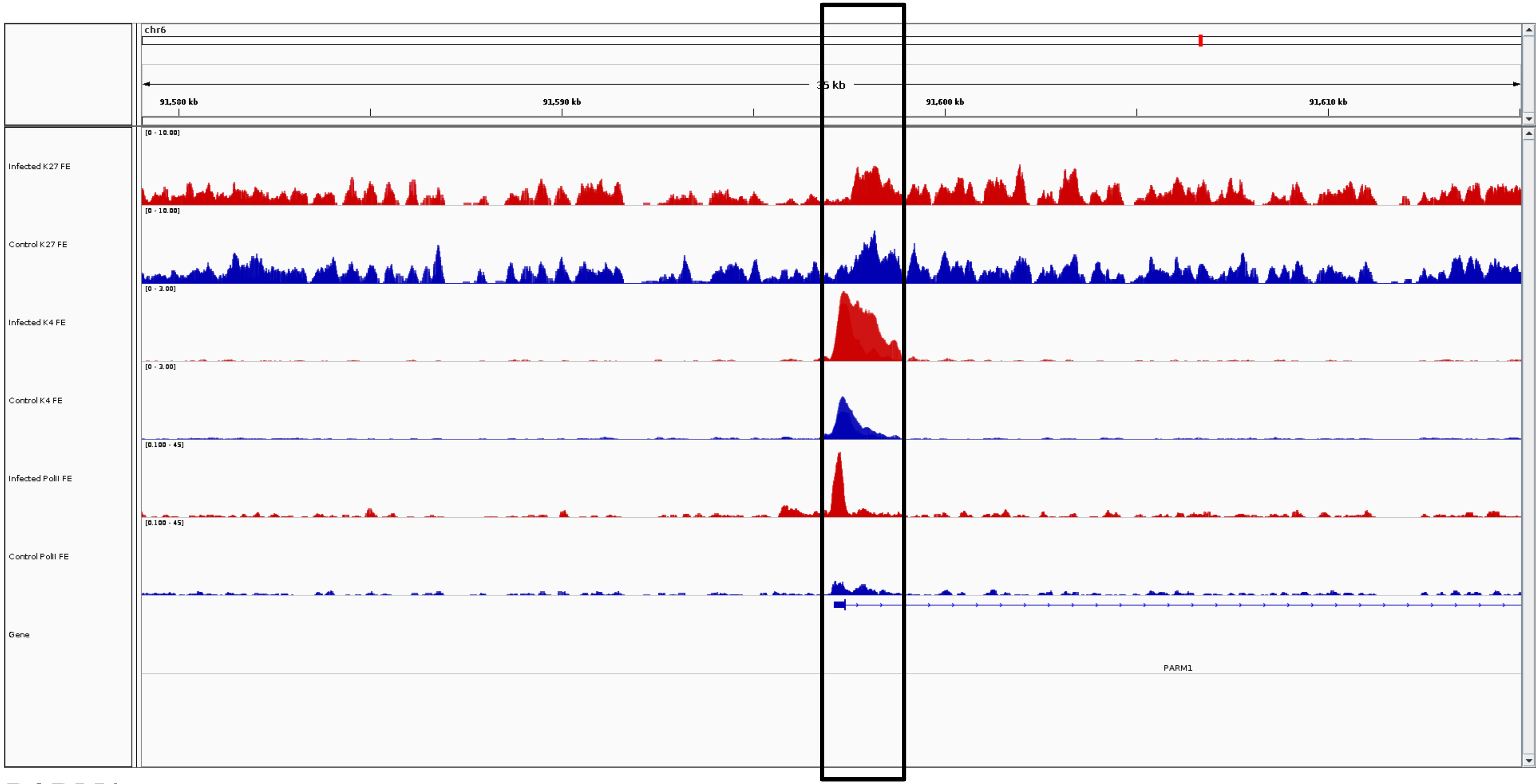
chr6
35 kb
91,580 kb
91,590 kb
91,600 kb
91,610 kb
Infected K27 FE
[0 - 10.00]
Control K27 FE
[0 - 10.00]
Infected K4 FE
[0 - 3.00]
Control K4 FE
[0 - 3.00]
Infected PolII FE
[0.100 - 45]
Control PolII FE
[0.100 - 45]
Gene
PARM1

***PARM1***

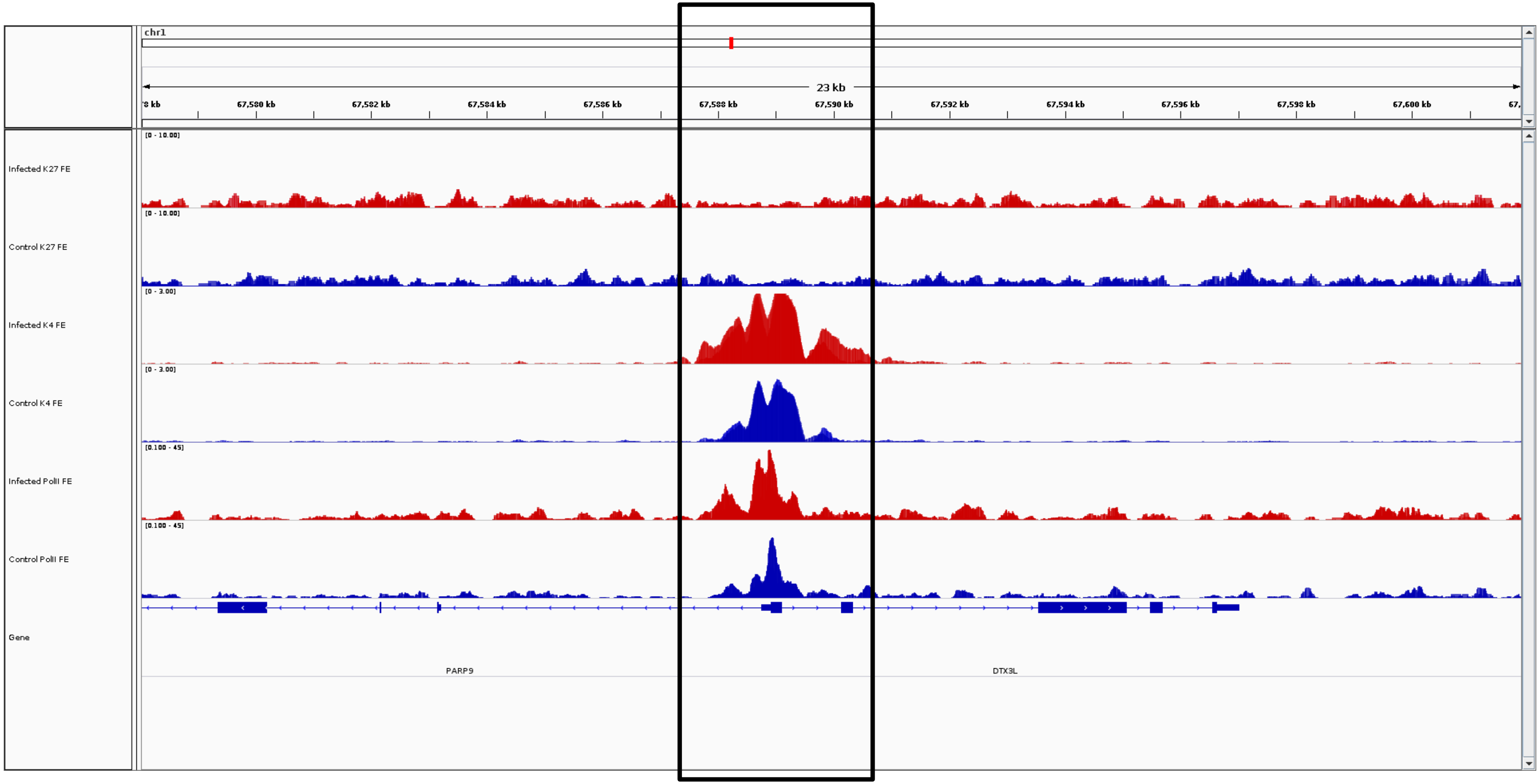
chr1
23 kb
67,580 kb
67,582 kb
67,584 kb
67,586 kb
67,588 kb
67,590 kb
67,592 kb
67,594 kb
67,596 kb
67,598 kb
67,600 kb
Infected K27 FE
[0 - 10.00]
Control K27 FE
[0 - 10.00]
Infected K4 FE
[0 - 3.00]
Control K4 FE
[0 - 3.00]
Infected PolII FE
[0.100 - 45]
Control PolII FE
[0.100 - 45]
Gene
PARP9
DTX3L

***PARP9***

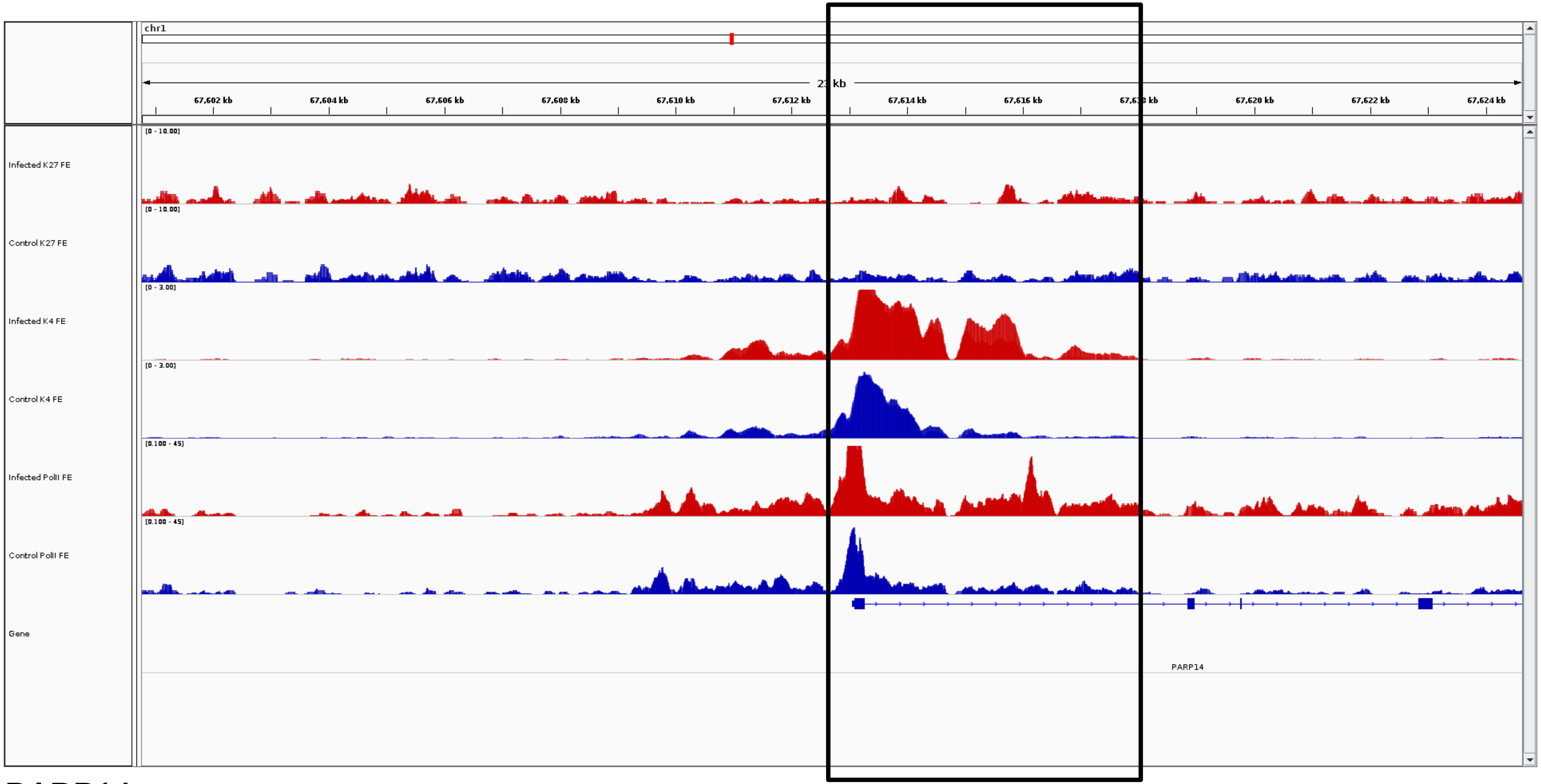
chr1
21 kb
67,602 kb
67,604 kb
67,606 kb
67,608 kb
67,610 kb
67,612 kb
67,614 kb
67,616 kb
67,620 kb
67,622 kb
67,624 kb
Infected K27 FE
[0 - 10.00]
Control K27 FE
[0 - 10.00]
Infected K4 FE
[0 - 3.00]
Control K4 FE
[0 - 3.00]
Infected PolII FE
[0.100 - 45]
Control PolII FE
[0.100 - 45]
Gene
PARP14

***PARP14***

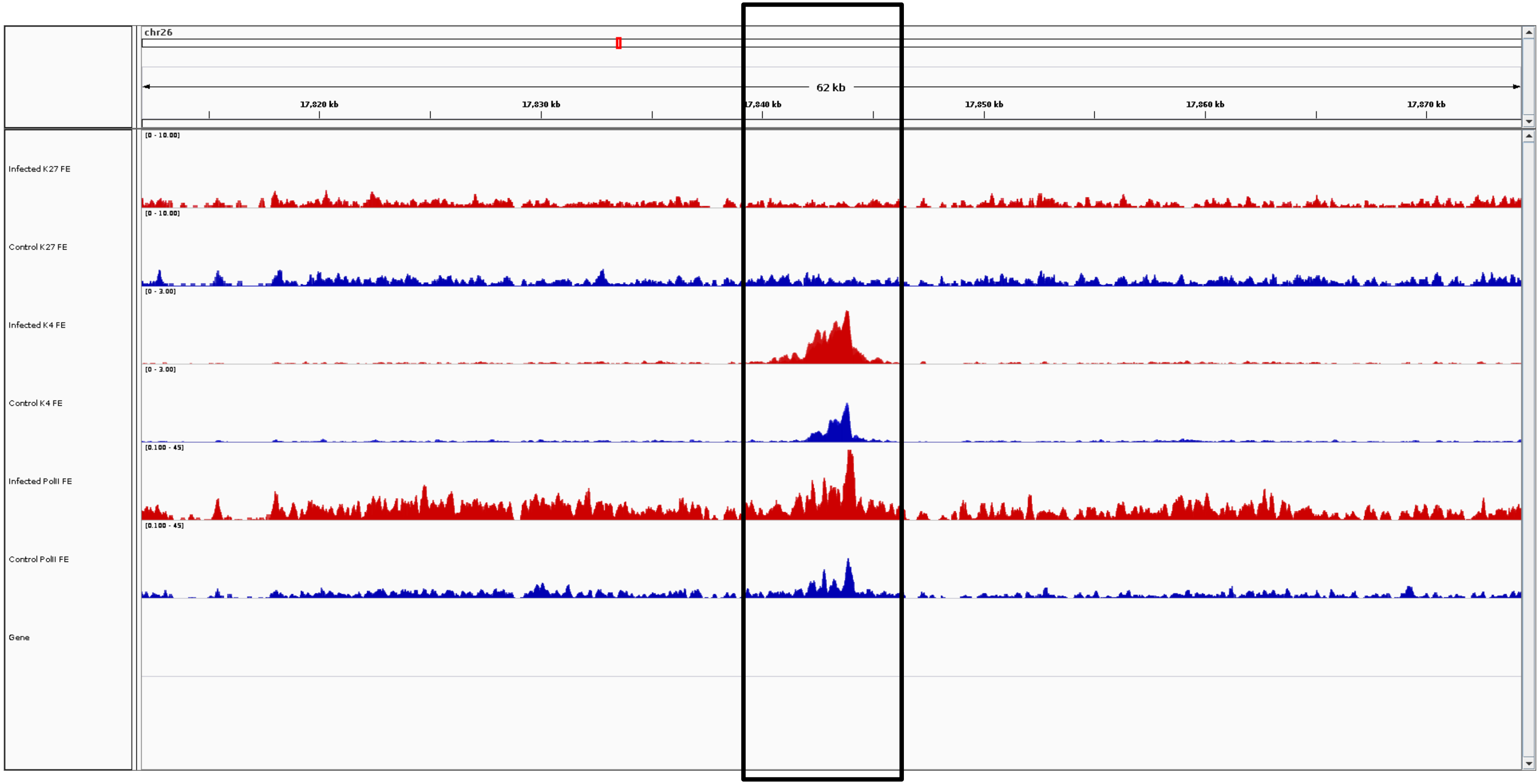
chr26
62 kb
17,820 kb
17,830 kb
17,840 kb
17,850 kb
17,860 kb
17,870 kb
Infected K27 FE
[0 - 10.00]
Control K27 FE
[0 - 10.00]
Infected K4 FE
[0 - 3.00]
Control K4 FE
[0 - 3.00]
Infected PolII FE
[0.100 - 45]
Control PolII FE
[0.100 - 45]
Gene

***PIK3AP1***

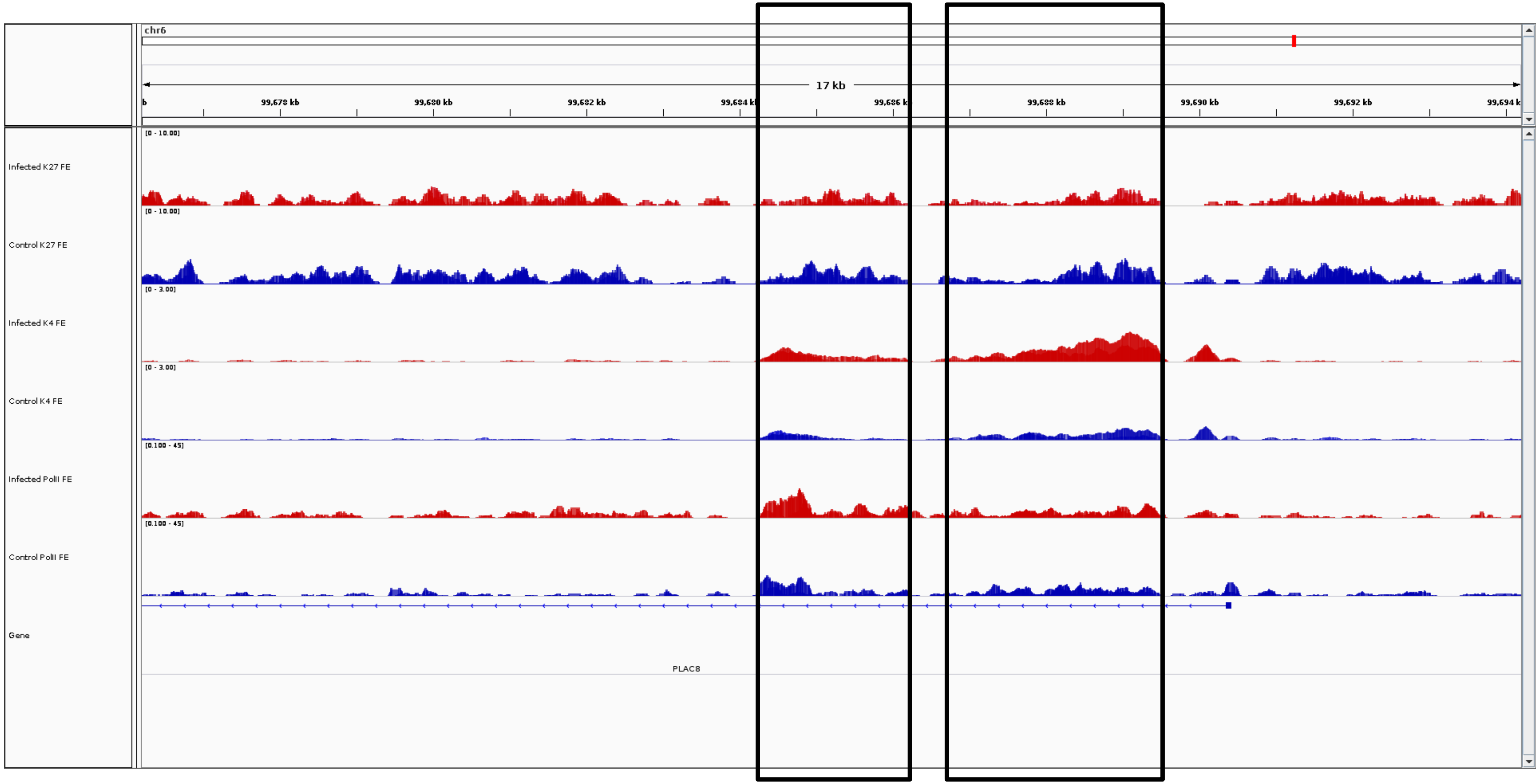
chr6
17 kb
99,678 kb
99,680 kb
99,682 kb
99,684 kb
99,686 kb
99,688 kb
99,690 kb
99,692 kb
99,694 kb
Infected K27 FE
[0 - 10.00]
Control K27 FE
[0 - 10.00]
Infected K4 FE
[0 - 3.00]
Control K4 FE
[0 - 3.00]
Infected PolII FE
[0.100 - 45]
Control PolII FE
[0.100 - 45]
Gene
PLAC8

***PLAC8***

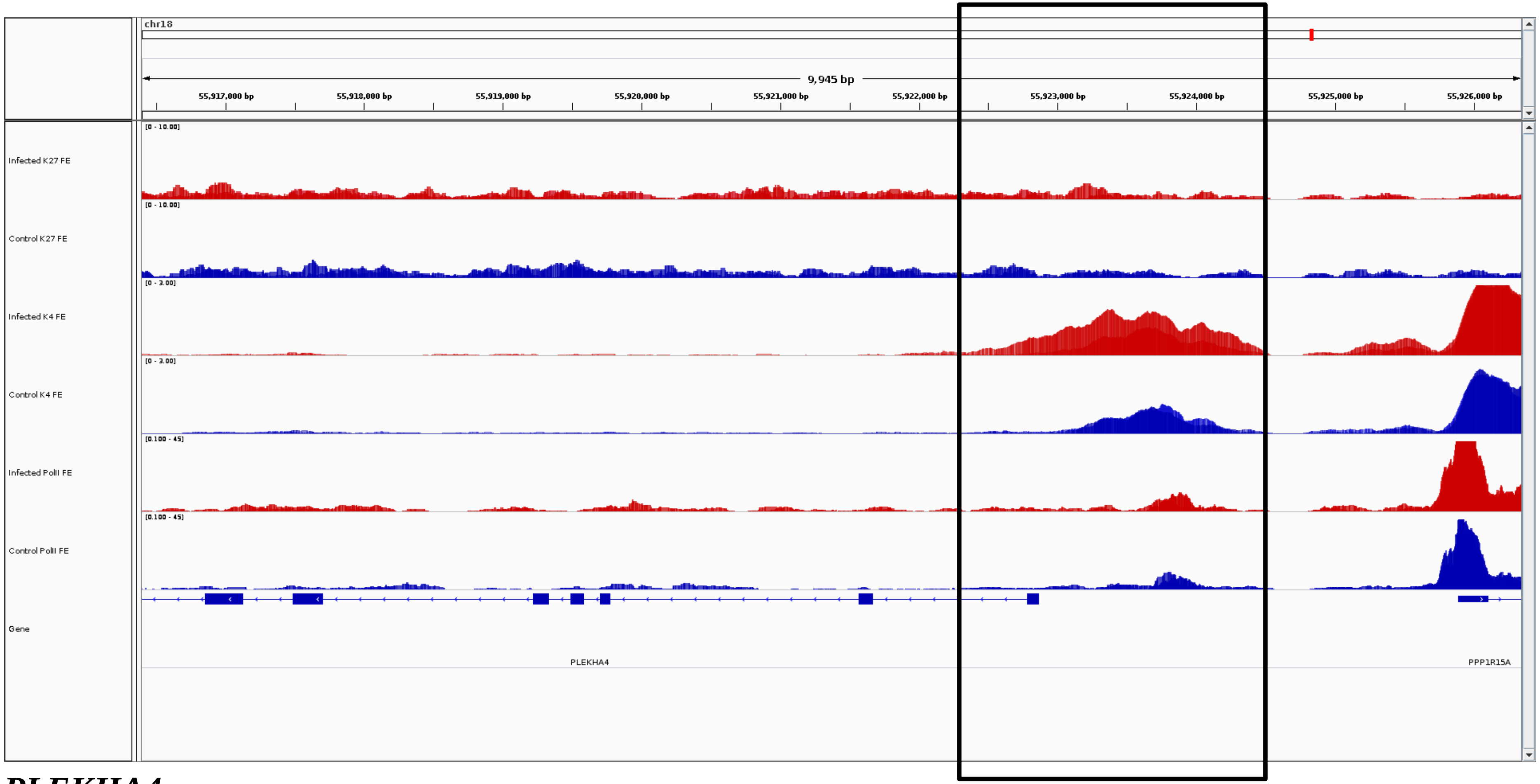
chr18
9,945 bp
55,917,000 bp
55,918,000 bp
55,919,000 bp
55,920,000 bp
55,921,000 bp
55,922,000 bp
55,923,000 bp
55,924,000 bp
55,925,000 bp
55,926,000 bp
Infected K27 FE
Control K27 FE
Infected K4 FE
Control K4 FE
Infected PolII FE
Control PolII FE
Gene
PLEKHA4
PPP1R15A

***PLEKHA4***

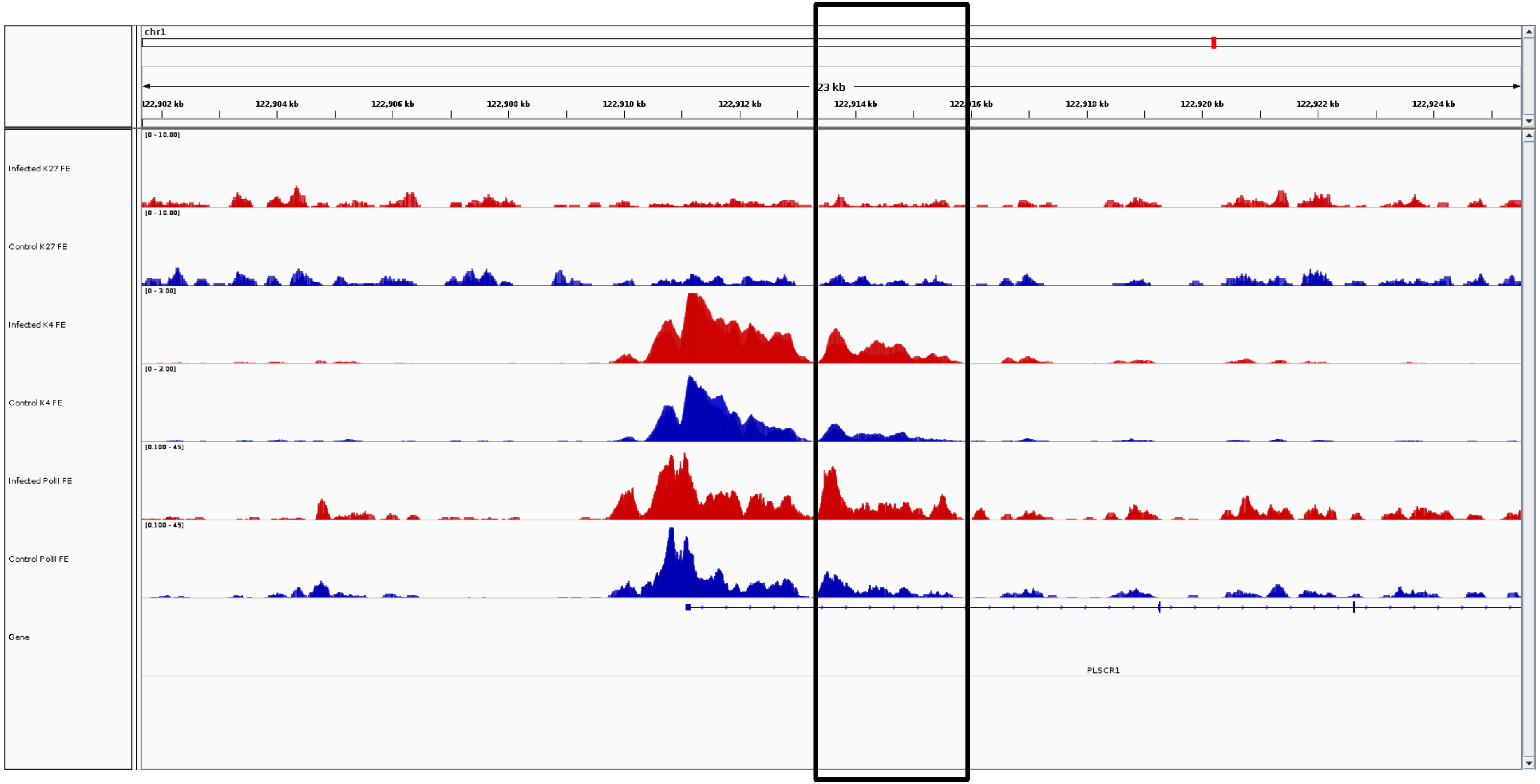
chr1
23 kb
122,902 kb
122,904 kb
122,906 kb
122,908 kb
122,910 kb
122,912 kb
122,914 kb
122,916 kb
122,918 kb
122,920 kb
122,922 kb
122,924 kb
Infected K27 FE
[0 - 10.00]
Control K27 FE
[0 - 10.00]
Infected K4 FE
[0 - 3.00]
Control K4 FE
[0 - 3.00]
Infected PolII FE
[0.100 - 45]
Control PolII FE
[0.100 - 45]
Gene
PLSCR1

***PLSCR1***

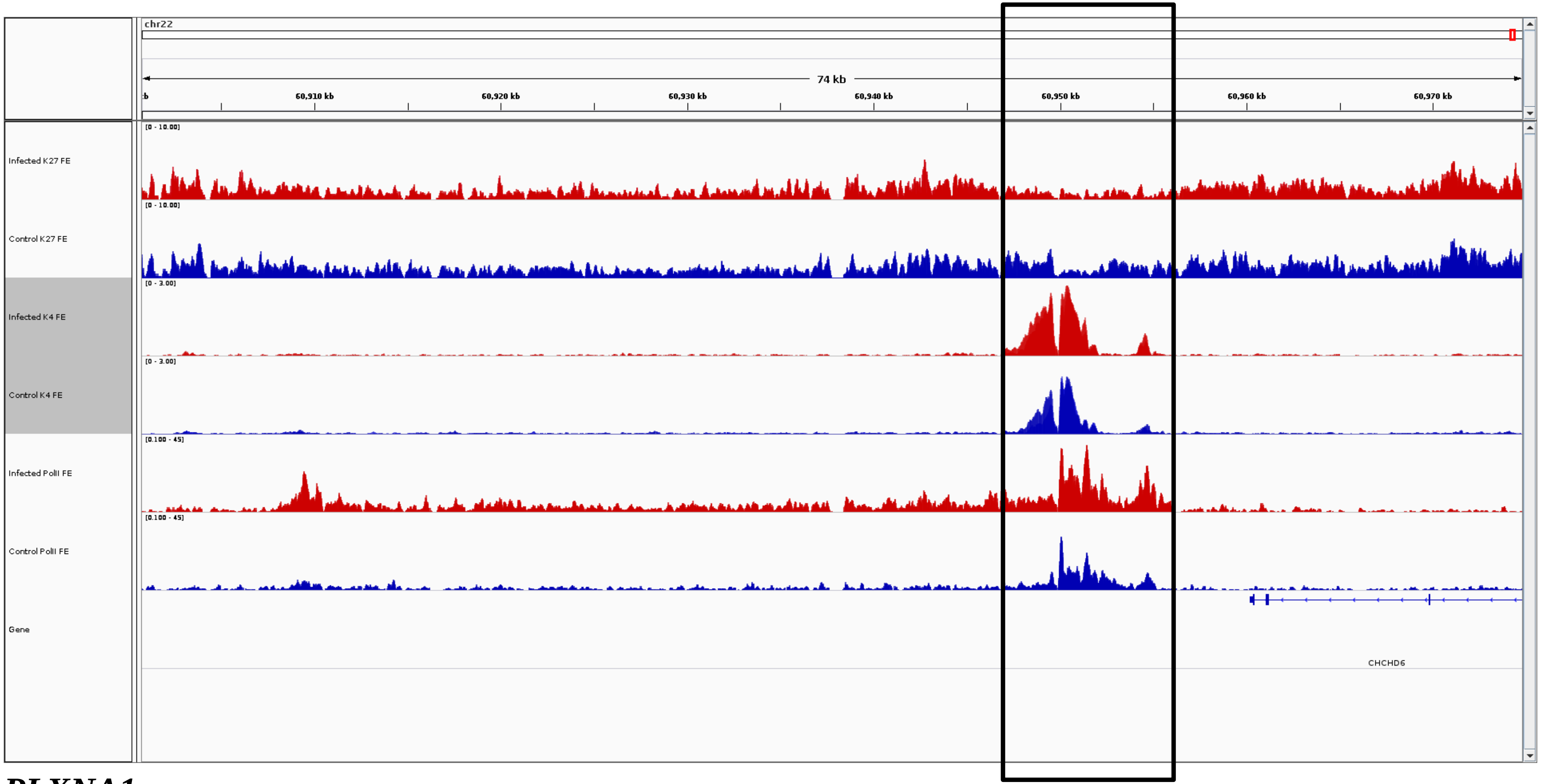
chr22
74 kb
60,910 kb
60,920 kb
60,930 kb
60,940 kb
60,950 kb
60,960 kb
60,970 kb
Infected K27 FE
Control K27 FE
Infected K4 FE
Control K4 FE
Infected PolII FE
Control PolII FE
Gene
CHCHD6

***PLXNA1***

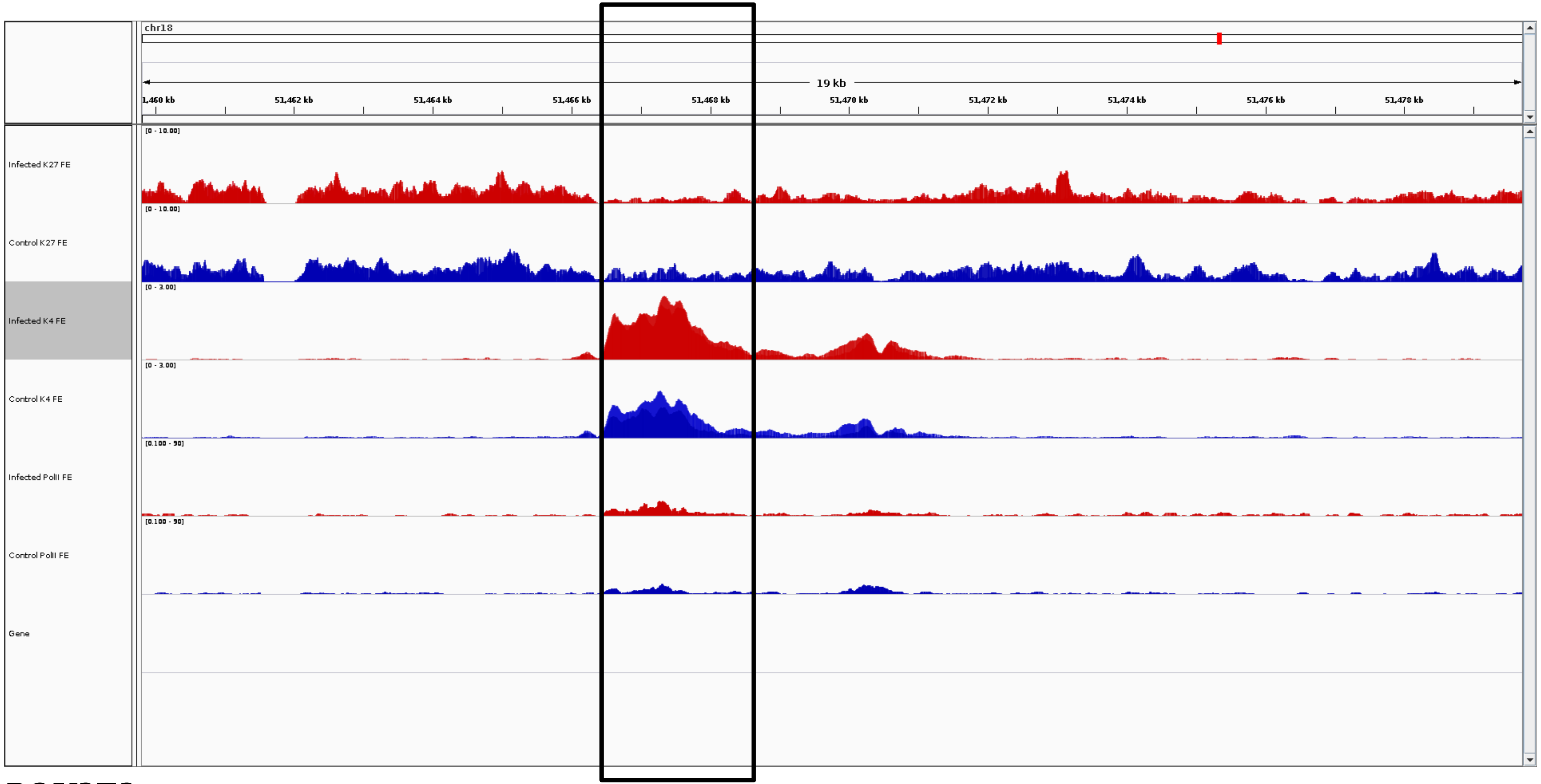
chr18
19 kb
1,460 kb
51,462 kb
51,464 kb
51,466 kb
51,468 kb
51,470 kb
51,472 kb
51,474 kb
51,476 kb
51,478 kb
Infected K27 FE
[0 - 10.00]
Control K27 FE
[0 - 10.00]
Infected K4 FE
[0 - 3.00]
Control K4 FE
[0 - 3.00]
Infected PolII FE
[0.100 - 50]
Control PolII FE
[0.100 - 50]
Gene

***POU2F2***

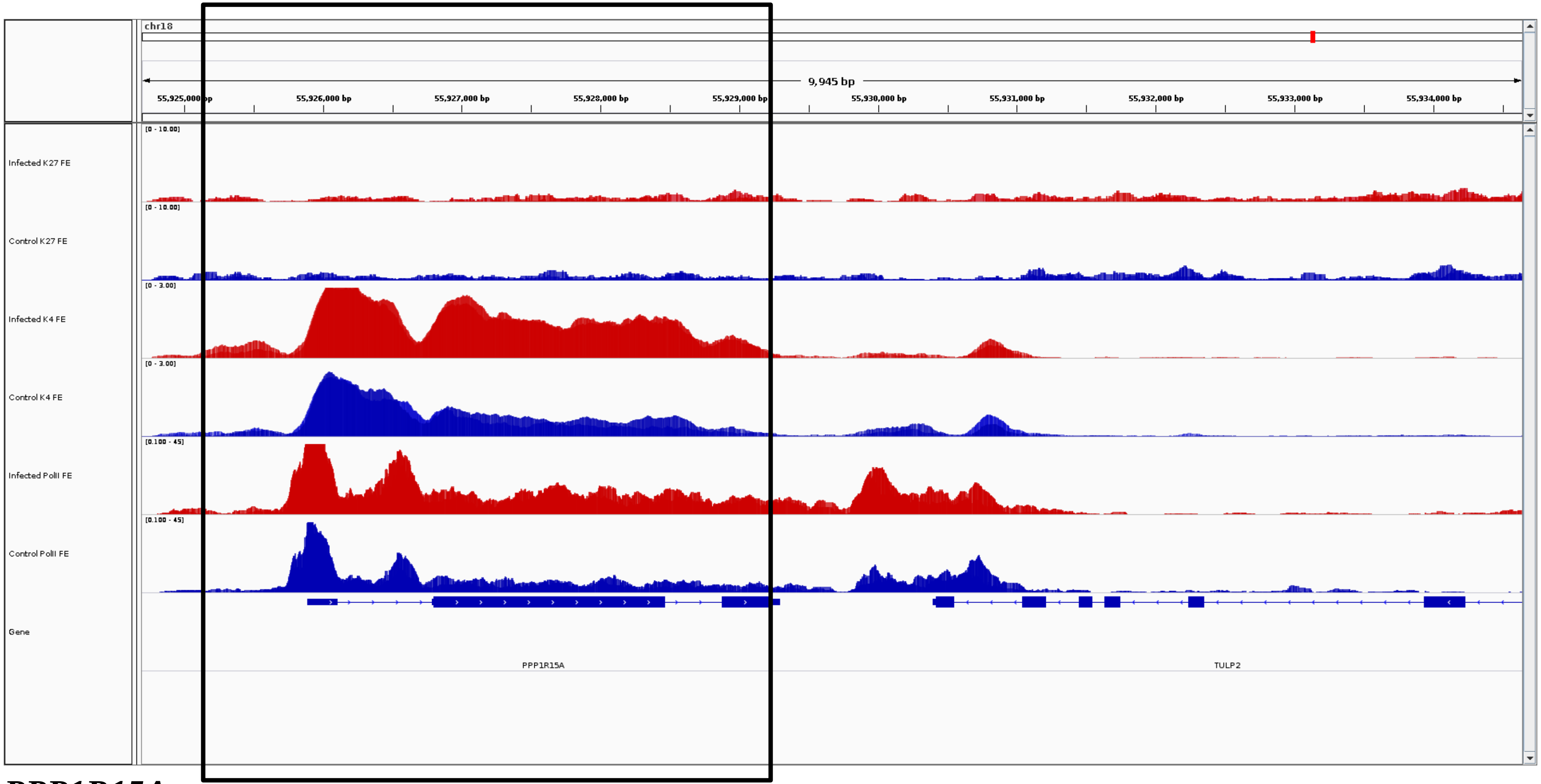
chr18
9,945 bp
55,925,000 bp
55,926,000 bp
55,927,000 bp
55,928,000 bp
55,929,000 bp
55,930,000 bp
55,931,000 bp
55,932,000 bp
55,933,000 bp
55,934,000 bp
Infected K27 FE
Control K27 FE
Infected K4 FE
Control K4 FE
Infected PolII FE
Control PolII FE
Gene
PPP1R15A
TULP2

***PPP1R15A***

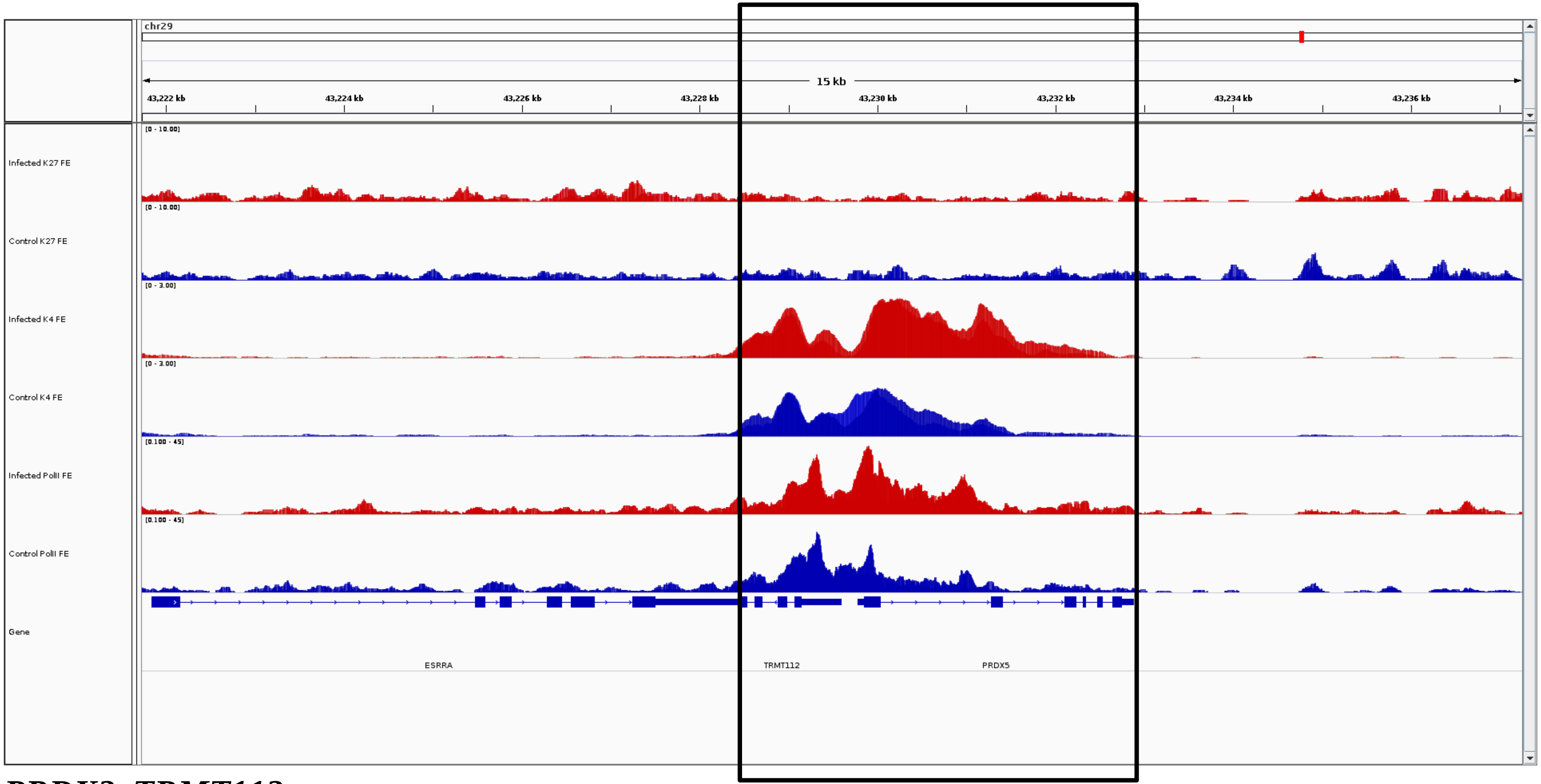


***PRDX2, TRMT112***

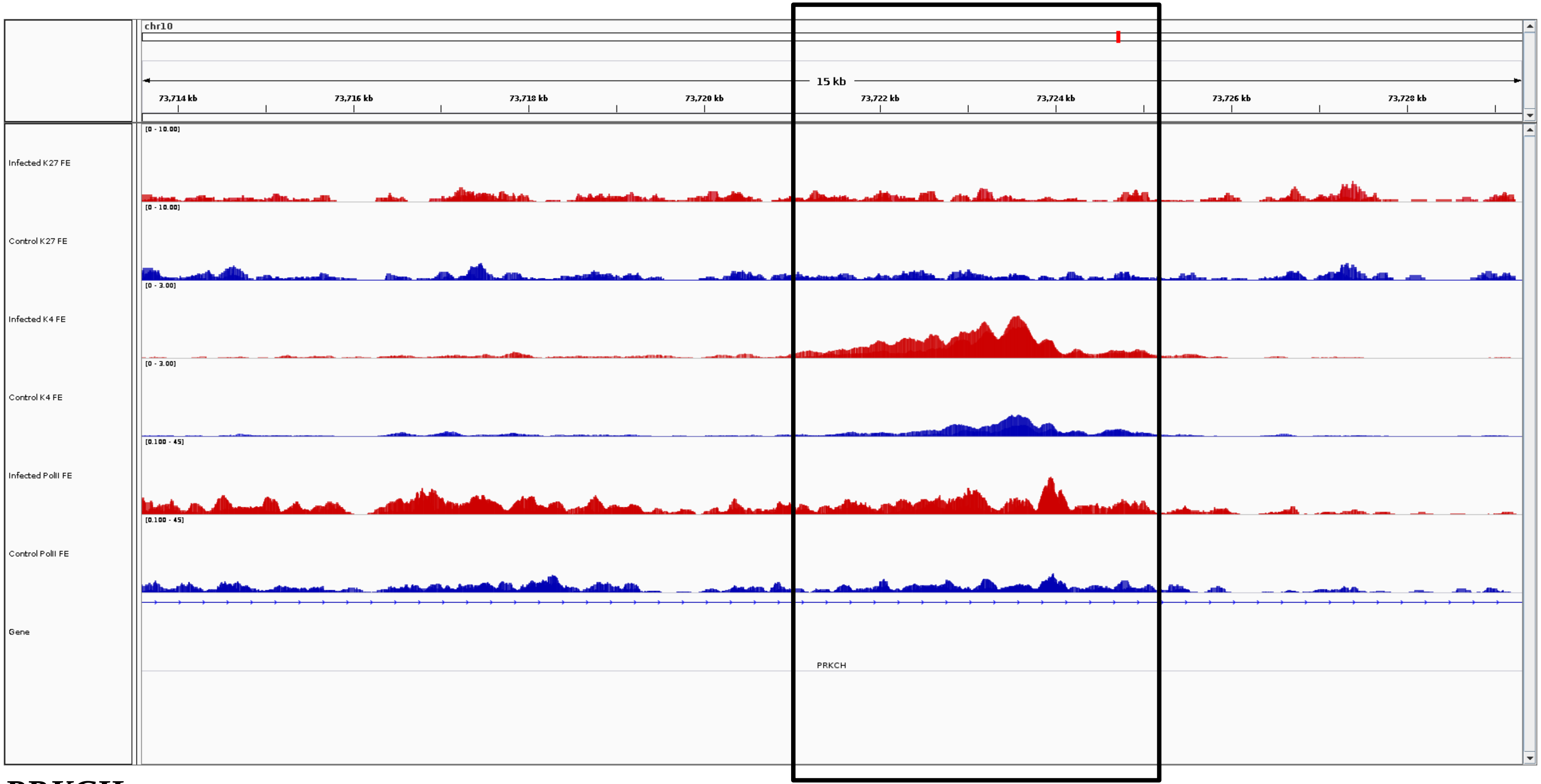
chr10
15 kb
73,714 kb
73,716 kb
73,718 kb
73,720 kb
73,722 kb
73,724 kb
73,726 kb
73,728 kb
Infected K27 FE
[0 - 10.00]
Control K27 FE
[0 - 10.00]
Infected K4 FE
[0 - 3.00]
Control K4 FE
[0 - 3.00]
Infected PolII FE
[0.100 - 45]
Control PolII FE
[0.100 - 45]
Gene
PRKCH

***PRKCH***

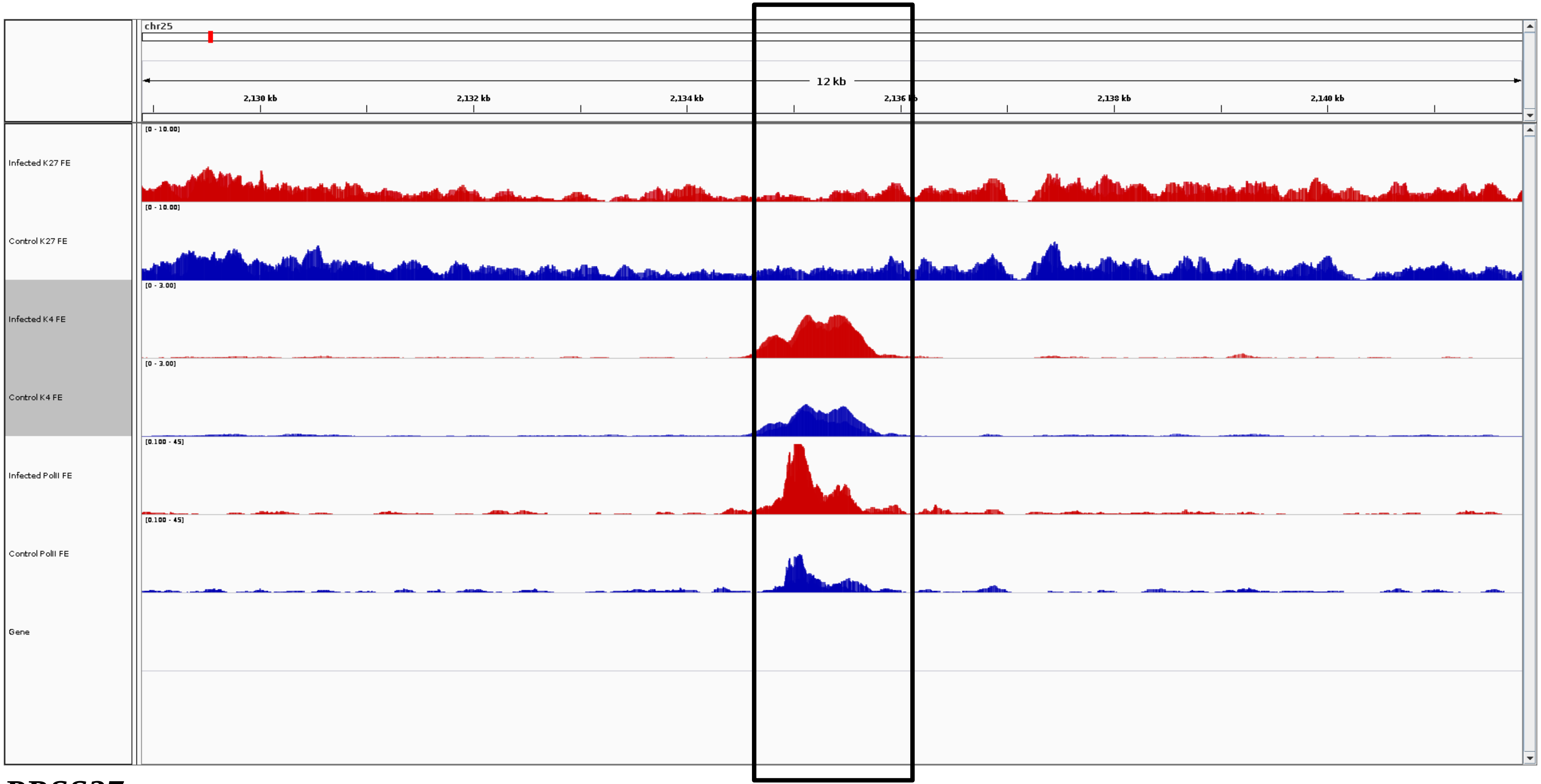
chr25
12 kb
2,130 kb
2,132 kb
2,134 kb
2,136 kb
2,138 kb
2,140 kb
[0 - 10.00]
Infected K27 FE
[0 - 10.00]
Control K27 FE
[0 - 3.00]
Infected K4 FE
[0 - 3.00]
Control K4 FE
[0.100 - 45]
Infected PolII FE
[0.100 - 45]
Control PolII FE
Gene

***PRSS27***

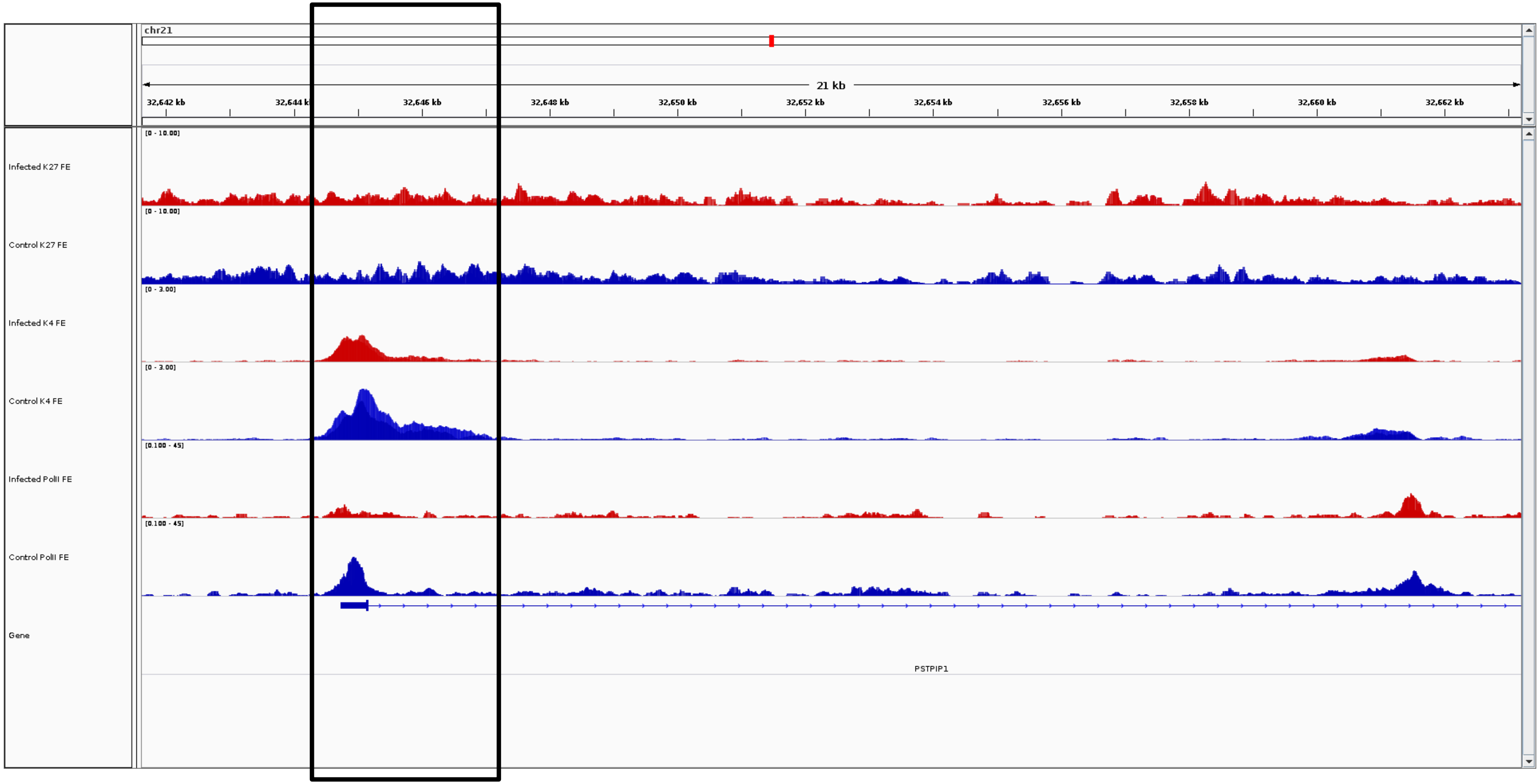
chr21
21 kb
32,642 kb
32,644 kb
32,646 kb
32,648 kb
32,650 kb
32,652 kb
32,654 kb
32,656 kb
32,658 kb
32,660 kb
32,662 kb
Infected K27 FE
Control K27 FE
Infected K4 FE
Control K4 FE
Infected PolII FE
Control PolII FE
Gene
PSTPIP1

***PSTPIP1***

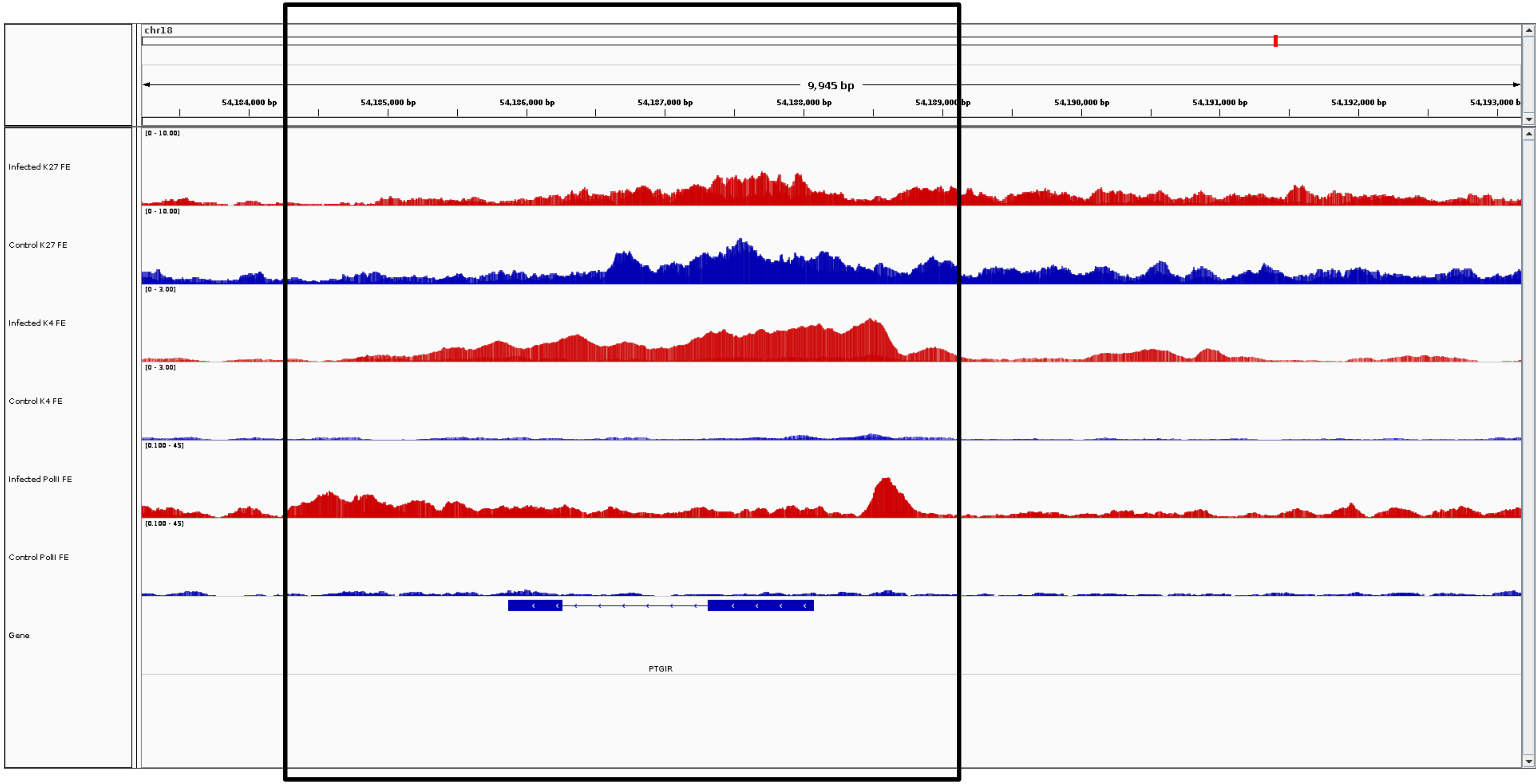
chr18
9,945 bp
54,184,000 bp
54,185,000 bp
54,186,000 bp
54,187,000 bp
54,188,000 bp
54,189,000 bp
54,190,000 bp
54,191,000 bp
54,192,000 bp
Infected K27 FE
Control K27 FE
Infected K4 FE
Control K4 FE
Infected PolII FE
Control PolII FE
Gene
PTGIR

***PTGIR***

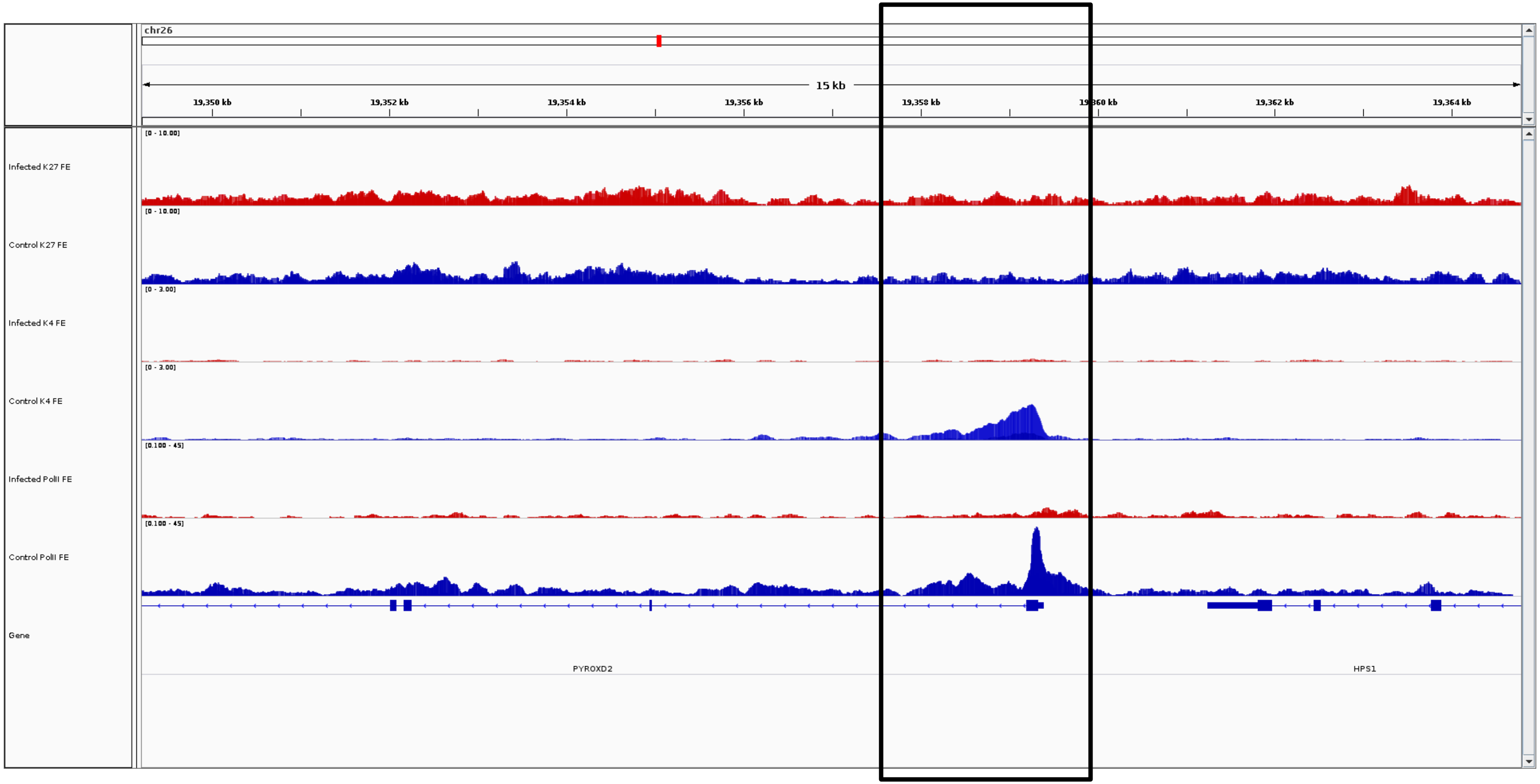
chr26
15 kb
19,350 kb
19,352 kb
19,354 kb
19,356 kb
19,358 kb
19,360 kb
19,362 kb
19,364 kb
Infected K27 FE
Control K27 FE
Infected K4 FE
Control K4 FE
Infected PolII FE
Control PolII FE
Gene
PYROXD2
HPS1

***PYROXD2***

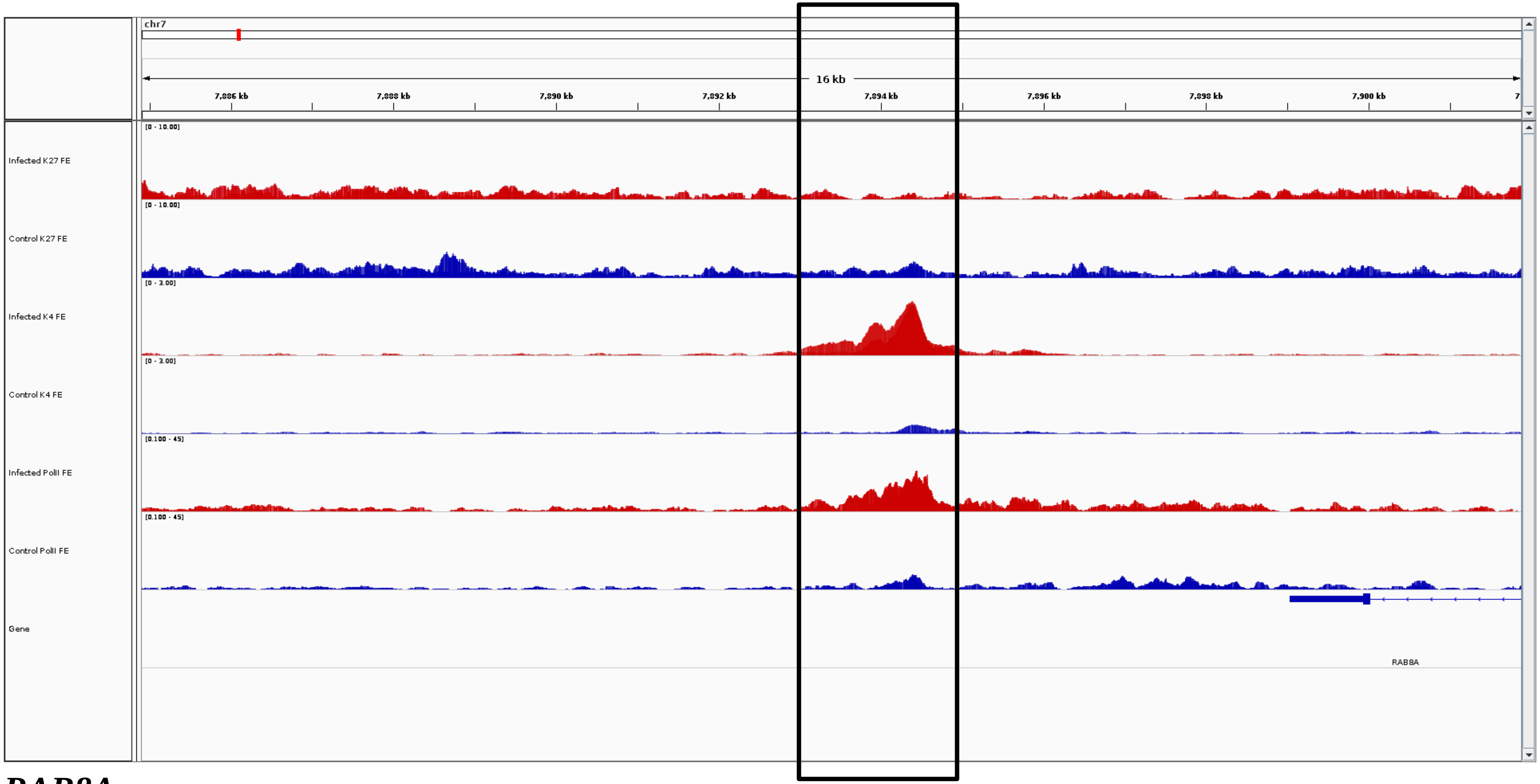
chr7
16 kb
7,886 kb
7,888 kb
7,890 kb
7,892 kb
7,894 kb
7,896 kb
7,898 kb
7,900 kb
Infected K27 FE
[0 - 10.00]
Control K27 FE
[0 - 10.00]
Infected K4 FE
[0 - 3.00]
Control K4 FE
[0 - 3.00]
Infected PolII FE
[0.100 - 45]
Control PolII FE
[0.100 - 45]
Gene
RAB8A

***RAB8A***

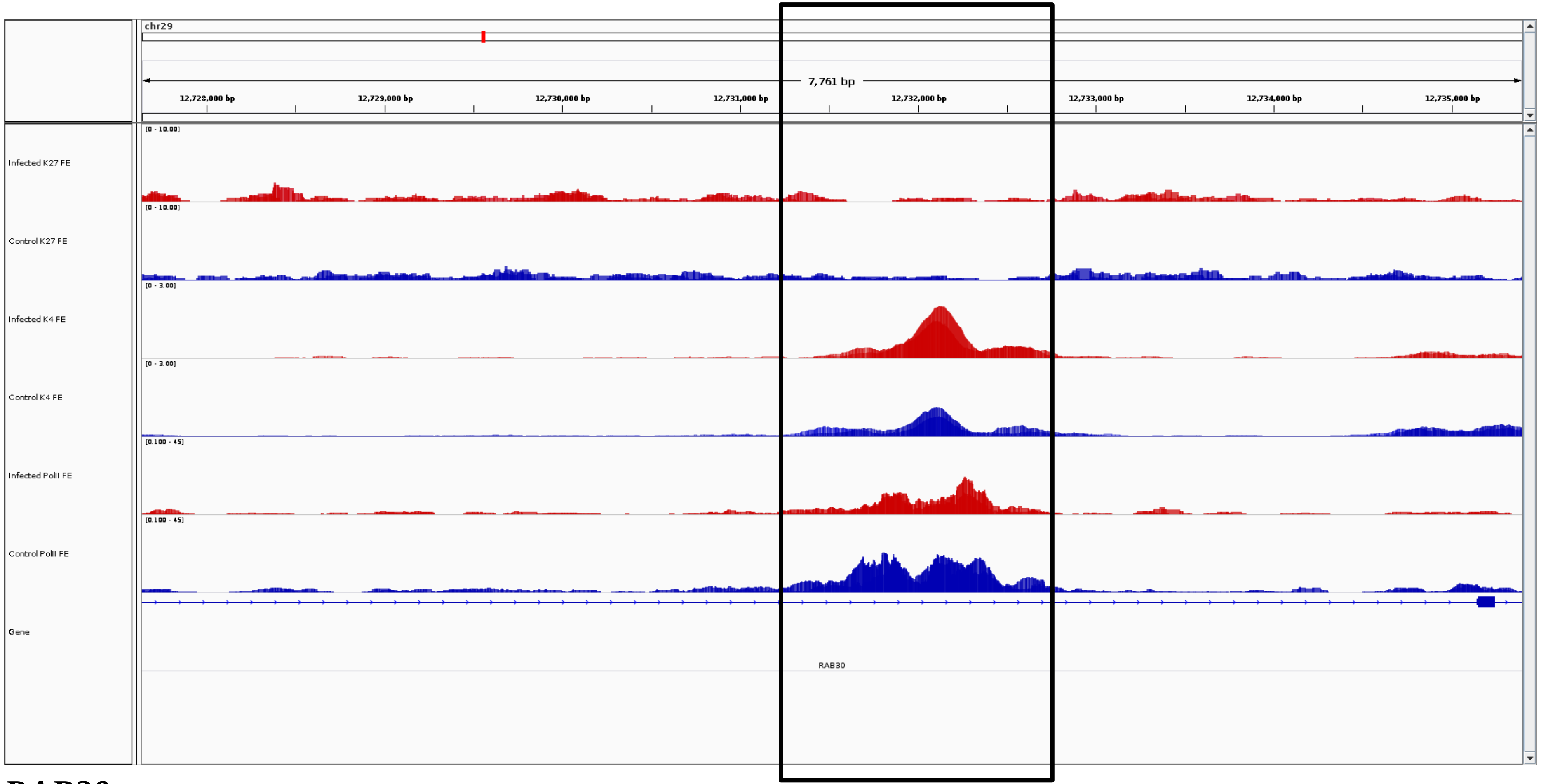
chr29
7,761 bp
12,728,000 bp
12,729,000 bp
12,730,000 bp
12,731,000 bp
12,732,000 bp
12,733,000 bp
12,734,000 bp
12,735,000 bp
Infected K27 FE
[0 - 10.00]
Control K27 FE
[0 - 10.00]
Infected K4 FE
[0 - 3.00]
Control K4 FE
[0 - 3.00]
Infected PolII FE
[0.100 - 45]
Control PolII FE
[0.100 - 45]
Gene
RAB30

***RAB30***

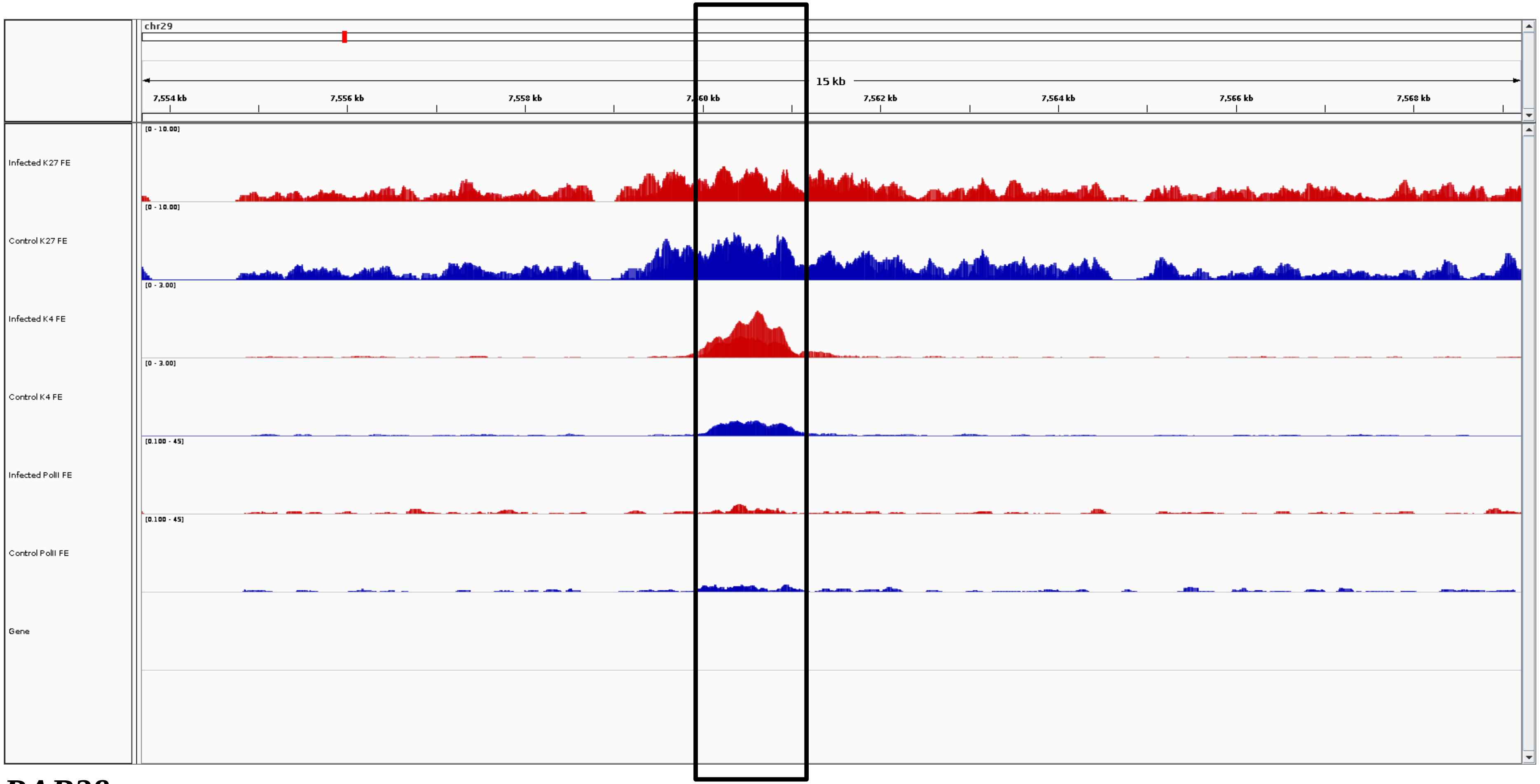
chr29
15 kb
7,554 kb
7,556 kb
7,558 kb
7,560 kb
7,562 kb
7,564 kb
7,566 kb
7,568 kb
Infected K27 FE
[0 - 10.00]
Control K27 FE
[0 - 10.00]
Infected K4 FE
[0 - 3.00]
Control K4 FE
[0 - 3.00]
Infected PolII FE
[0.100 - 45]
Control PolII FE
[0.100 - 45]
Gene

***RAB38***

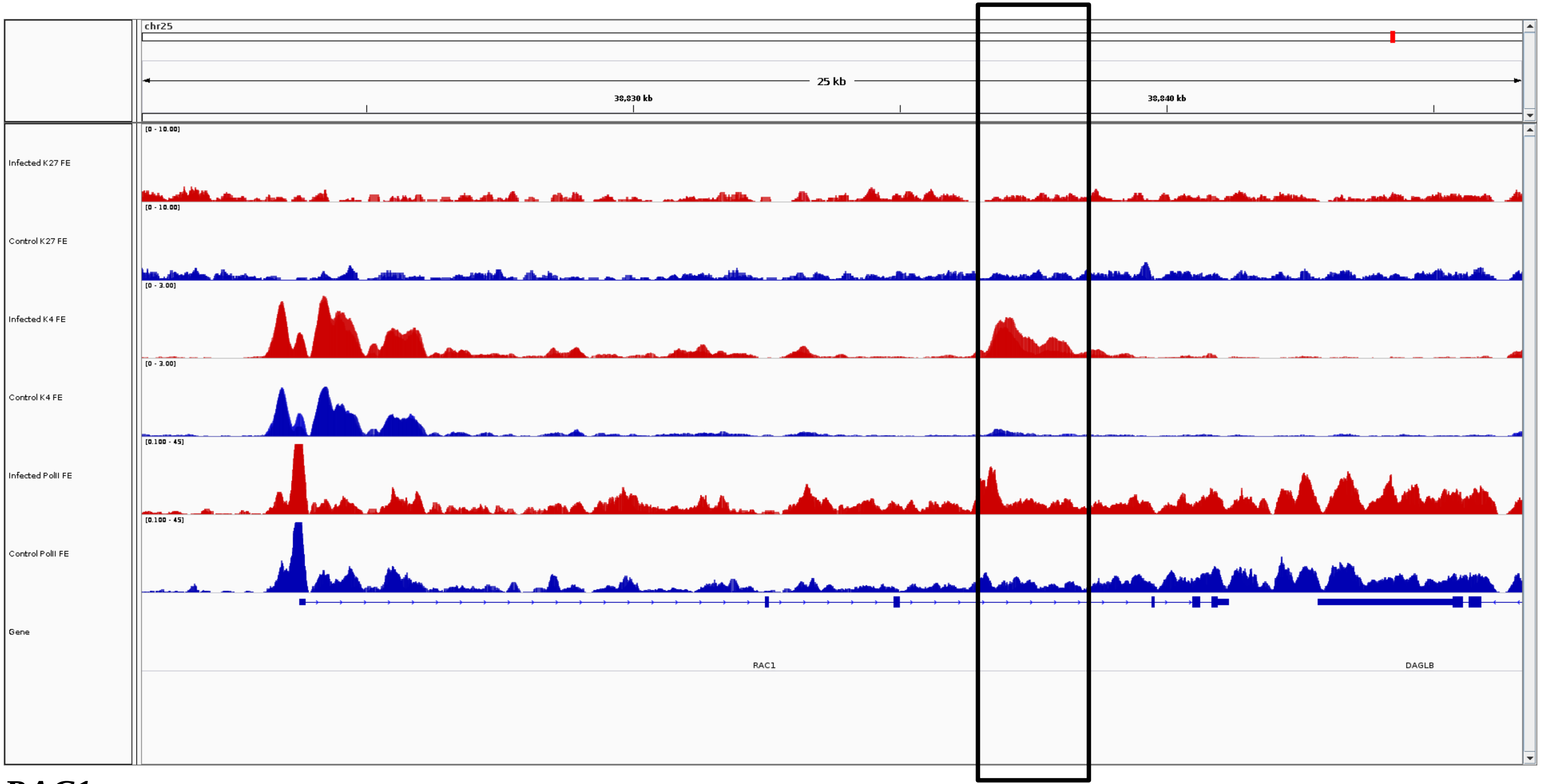
chr25
25 kb
38,830 kb
38,840 kb
Infected K27 FE
Control K27 FE
Infected K4 FE
Control K4 FE
Infected PolII FE
Control PolII FE
Gene
RAC1
DAGLB

***RAC1***

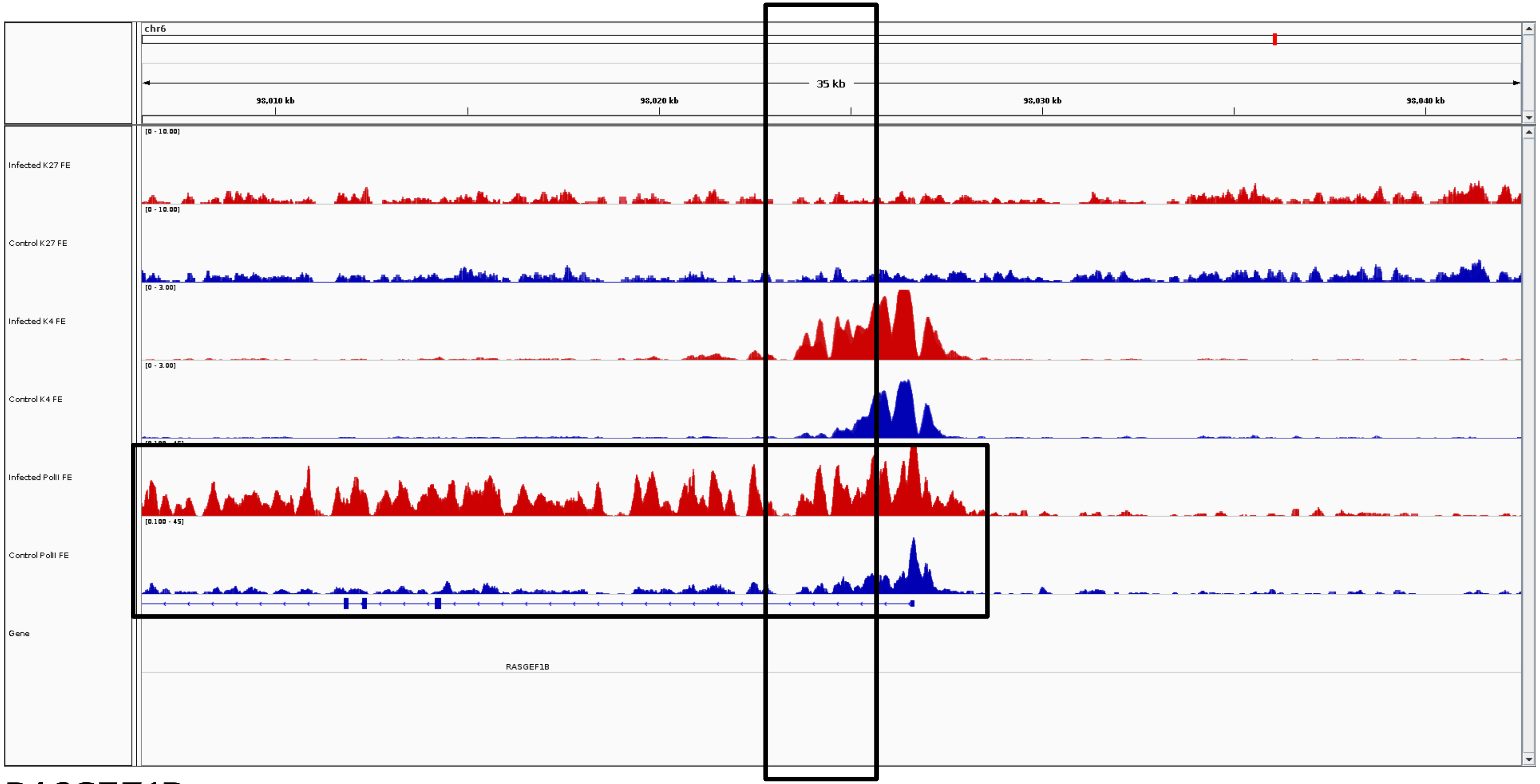
chr6
35 kb
98,010 kb
98,020 kb
98,030 kb
98,040 kb
Infected K27 FE
Control K27 FE
Infected K4 FE
Control K4 FE
Infected PolII FE
Control PolII FE
Gene
RASGEF1B

***RASGEF1B***

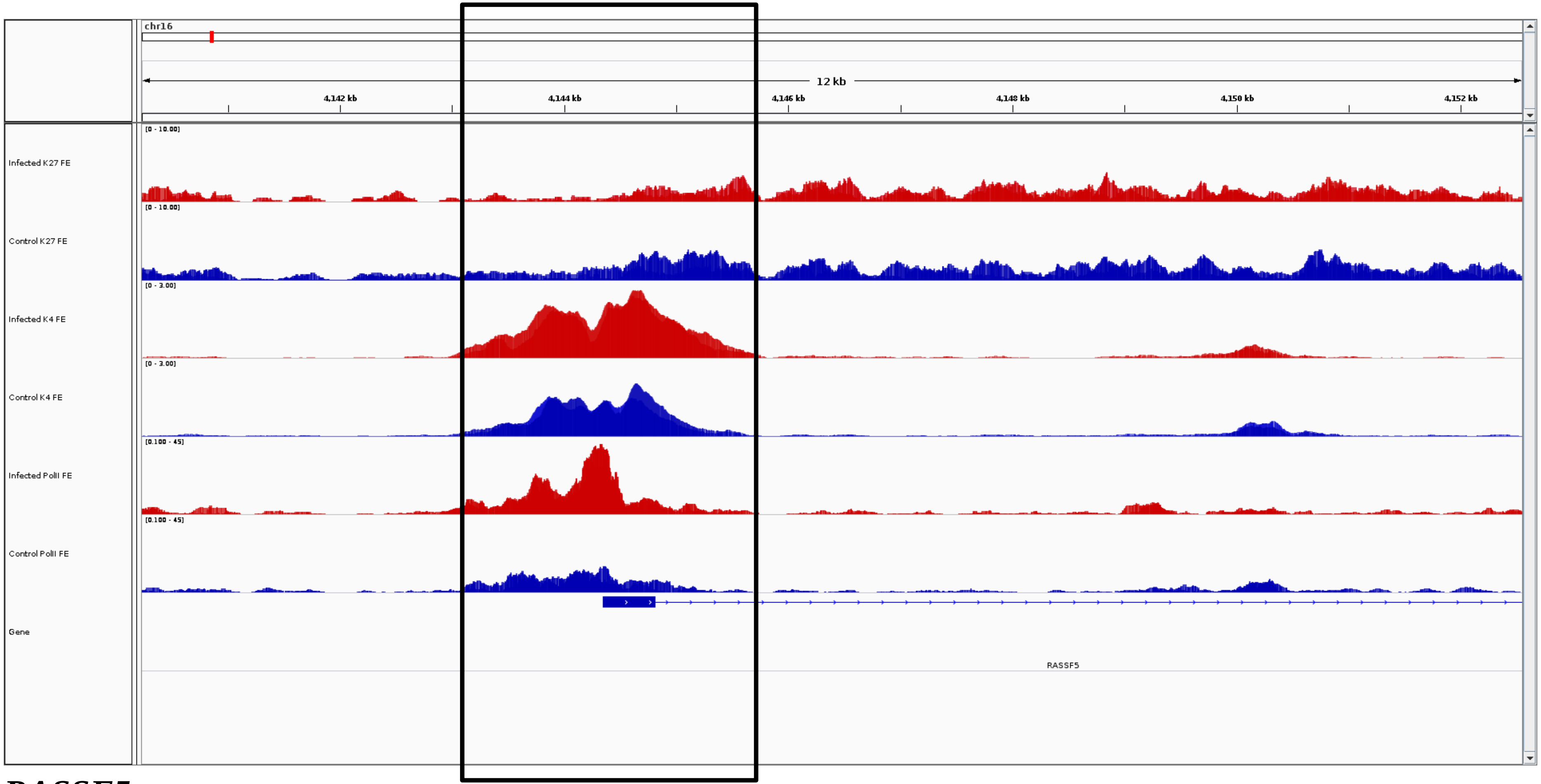
chr16
12 kb
4,142 kb
4,144 kb
4,146 kb
4,148 kb
4,150 kb
4,152 kb
Infected K27 FE
[0 - 10.00]
Control K27 FE
[0 - 10.00]
Infected K4 FE
[0 - 3.00]
Control K4 FE
[0 - 3.00]
Infected PolII FE
[0.100 - 45]
Control PolII FE
[0.100 - 45]
Gene
RASSF5

***RASSF5***

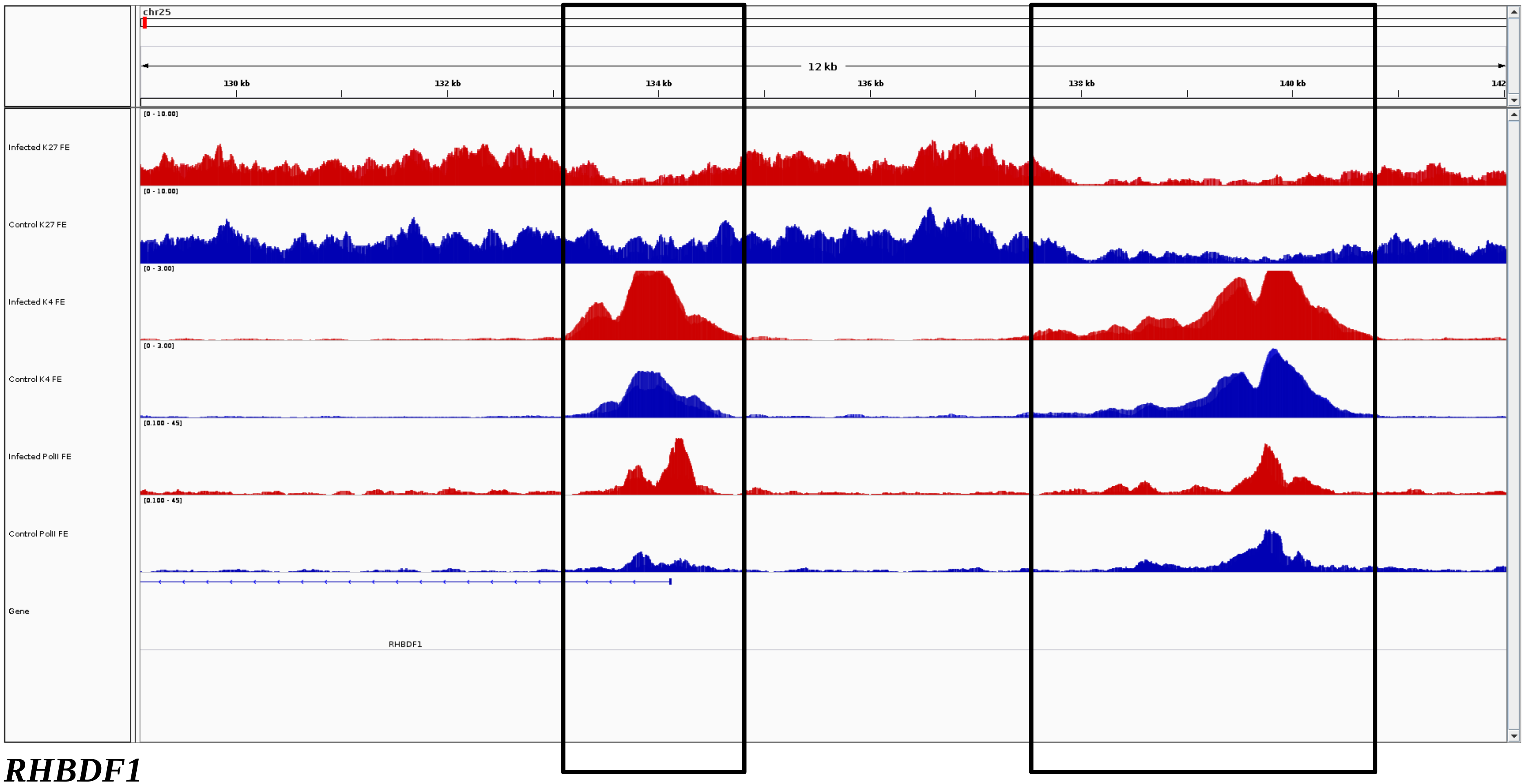
chr25
12 kb
130 kb
132 kb
134 kb
136 kb
138 kb
140 kb
142
Infected K27 FE
Control K27 FE
Infected K4 FE
Control K4 FE
Infected PolII FE
Control PolII FE
Gene
RHBDF1

***RHBDF1***

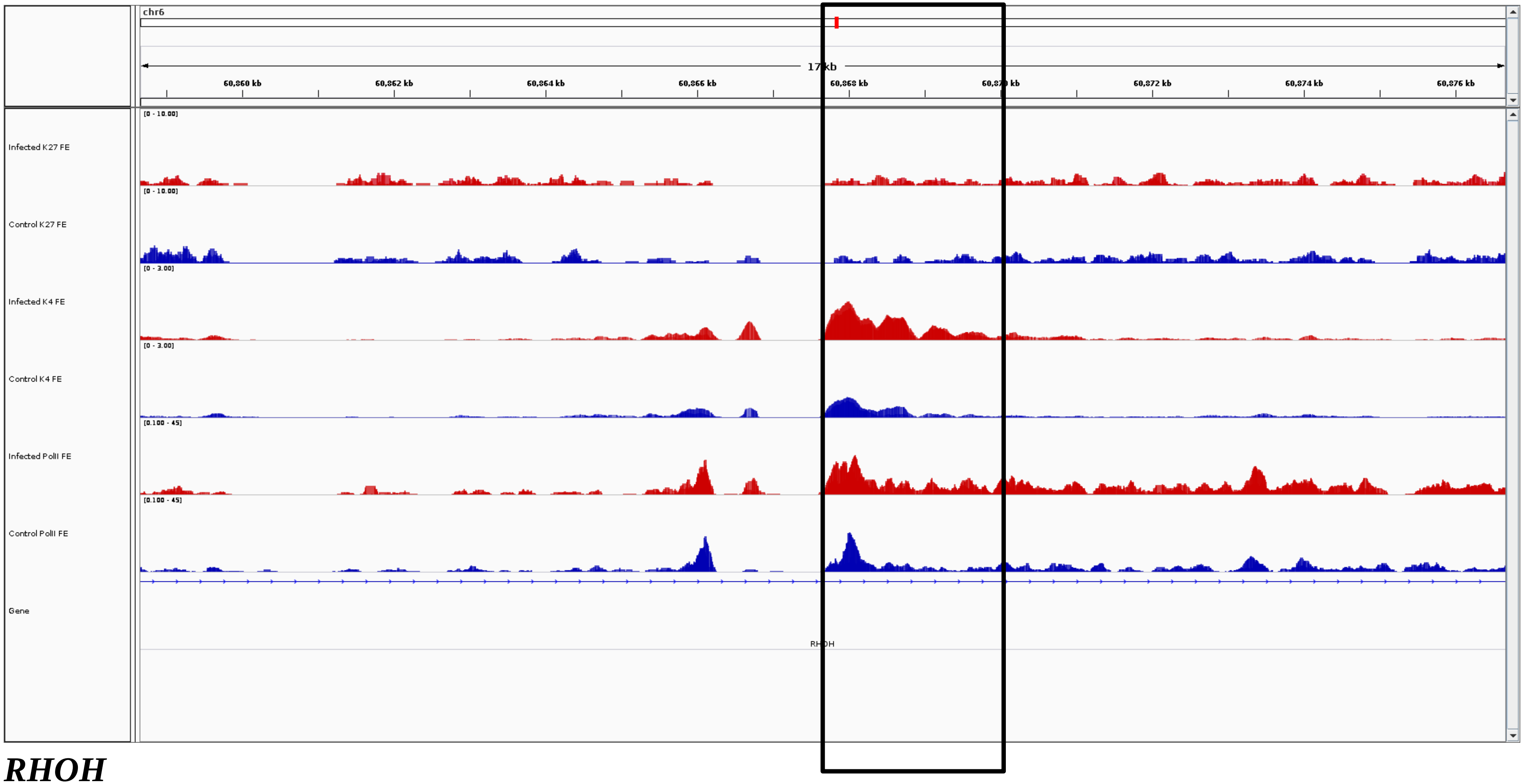
chr6
17 kb
60,860 kb
60,862 kb
60,864 kb
60,866 kb
60,868 kb
60,872 kb
60,874 kb
60,876 kb
Infected K27 FE
[0 - 10.00]
Control K27 FE
[0 - 10.00]
Infected K4 FE
[0 - 3.00]
Control K4 FE
[0 - 3.00]
Infected PolII FE
[0.100 - 45]
Control PolII FE
[0.100 - 45]
Gene
RHOH

***RHOH***

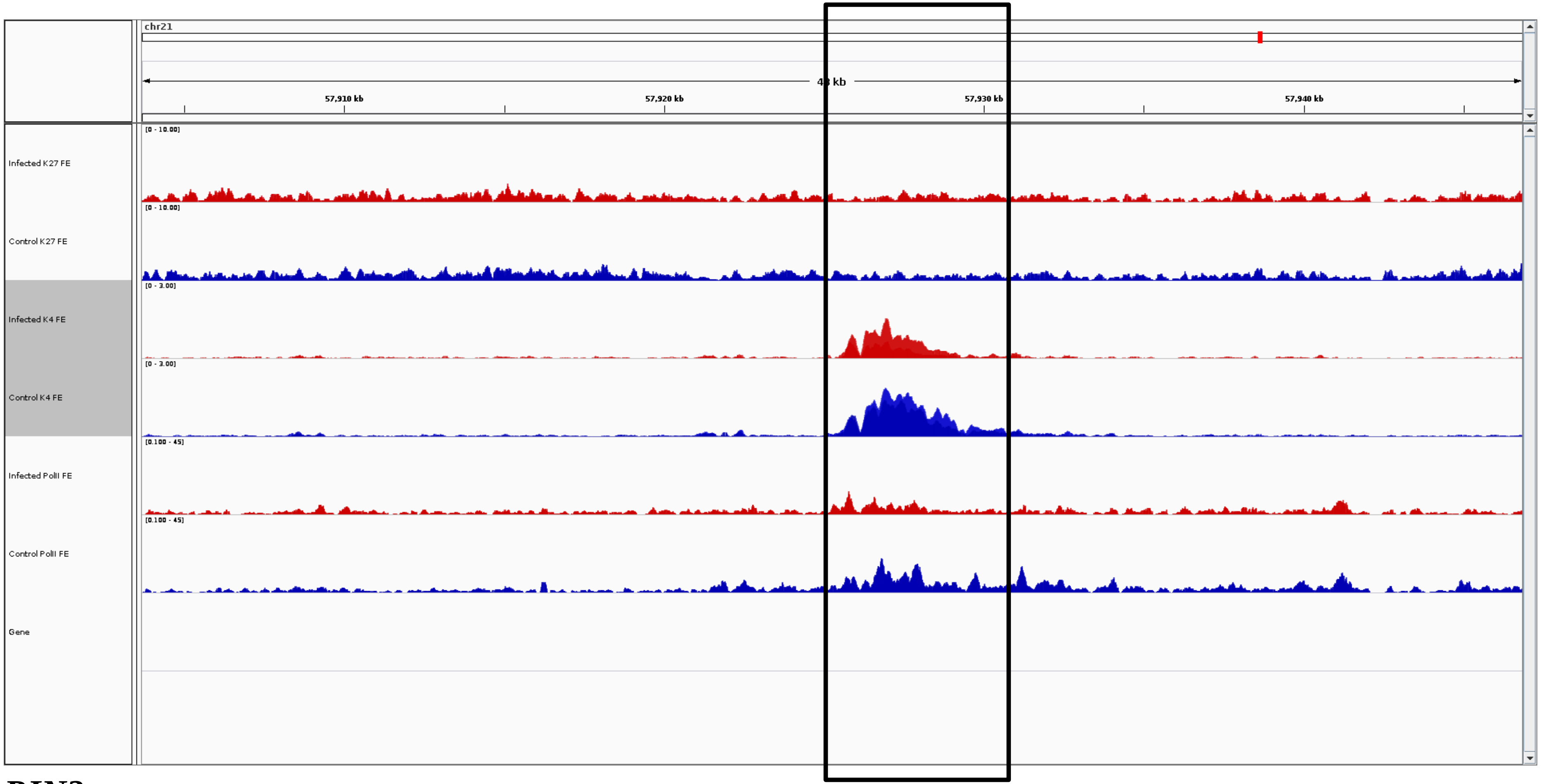
chr21
kb
57,910 kb
57,920 kb
57,930 kb
57,940 kb
[0 - 10.00]
Infected K27 FE
[0 - 10.00]
Control K27 FE
[0 - 3.00]
Infected K4 FE
[0 - 3.00]
Control K4 FE
[0.100 - 45]
Infected PolII FE
[0.100 - 45]
Control PolII FE
Gene

***RIN3***

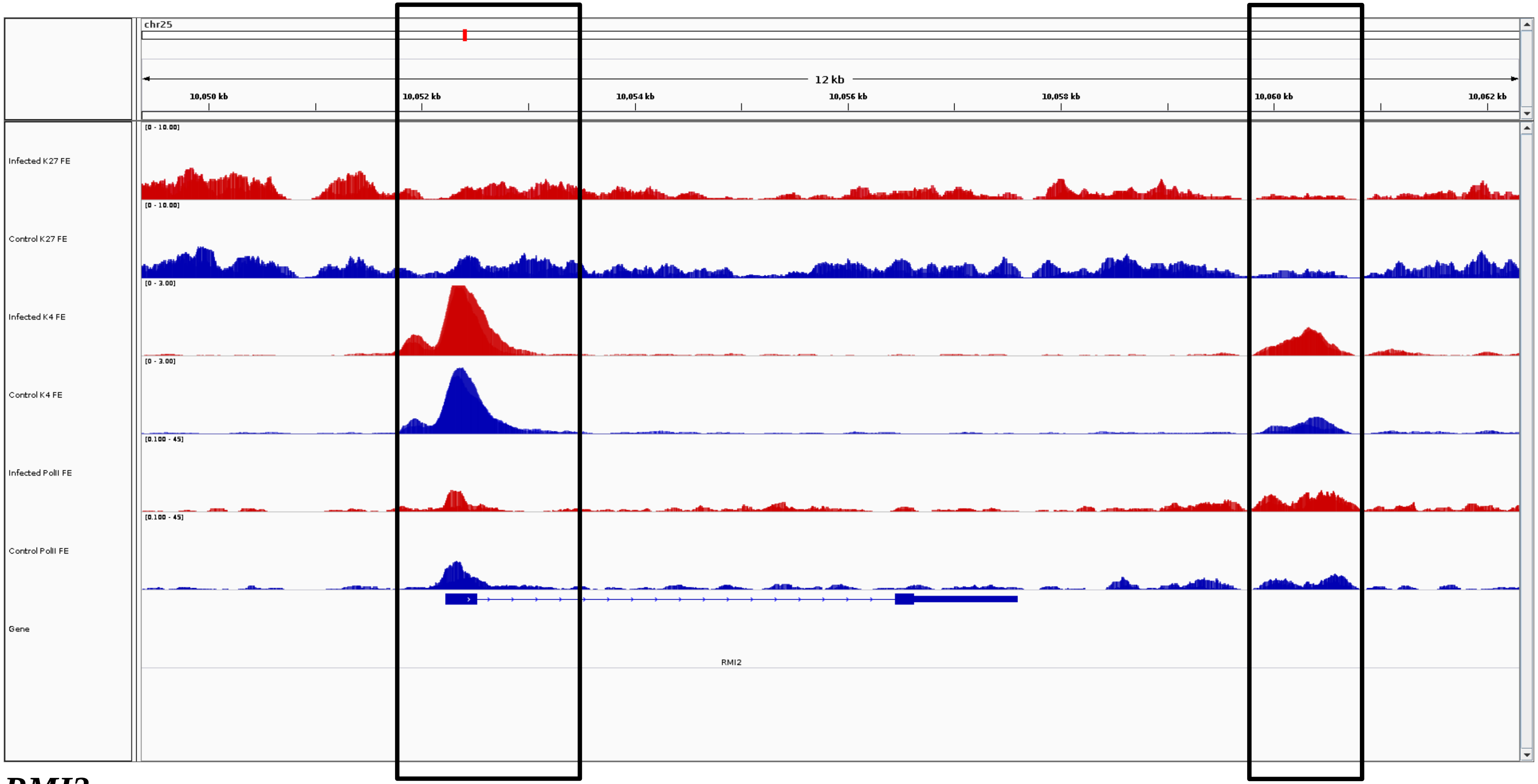
chr25
12 kb
10,050 kb
10,052 kb
10,054 kb
10,056 kb
10,058 kb
10,060 kb
10,062 kb
Infected K27 FE
Control K27 FE
Infected K4 FE
Control K4 FE
Infected PolII FE
Control PolII FE
Gene
RMI2

***RMI2***

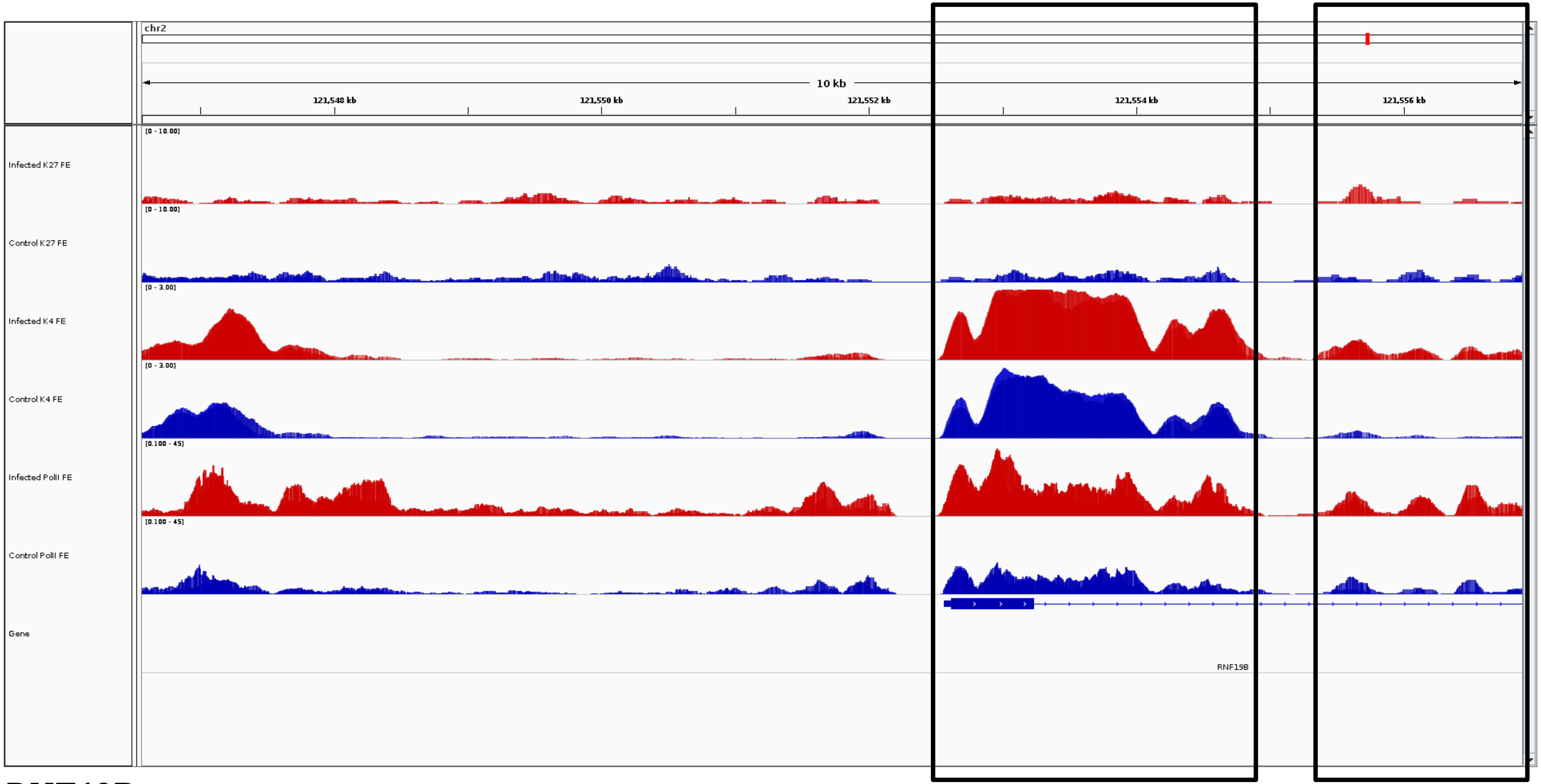
chr2
10 kb
121,548 kb
121,550 kb
121,552 kb
121,554 kb
121,556 kb
Infected K27 FE
Control K27 FE
Infected K4 FE
Control K4 FE
Infected PolII FE
Control PolII FE
Gene
RNF19B

***RNF19B***

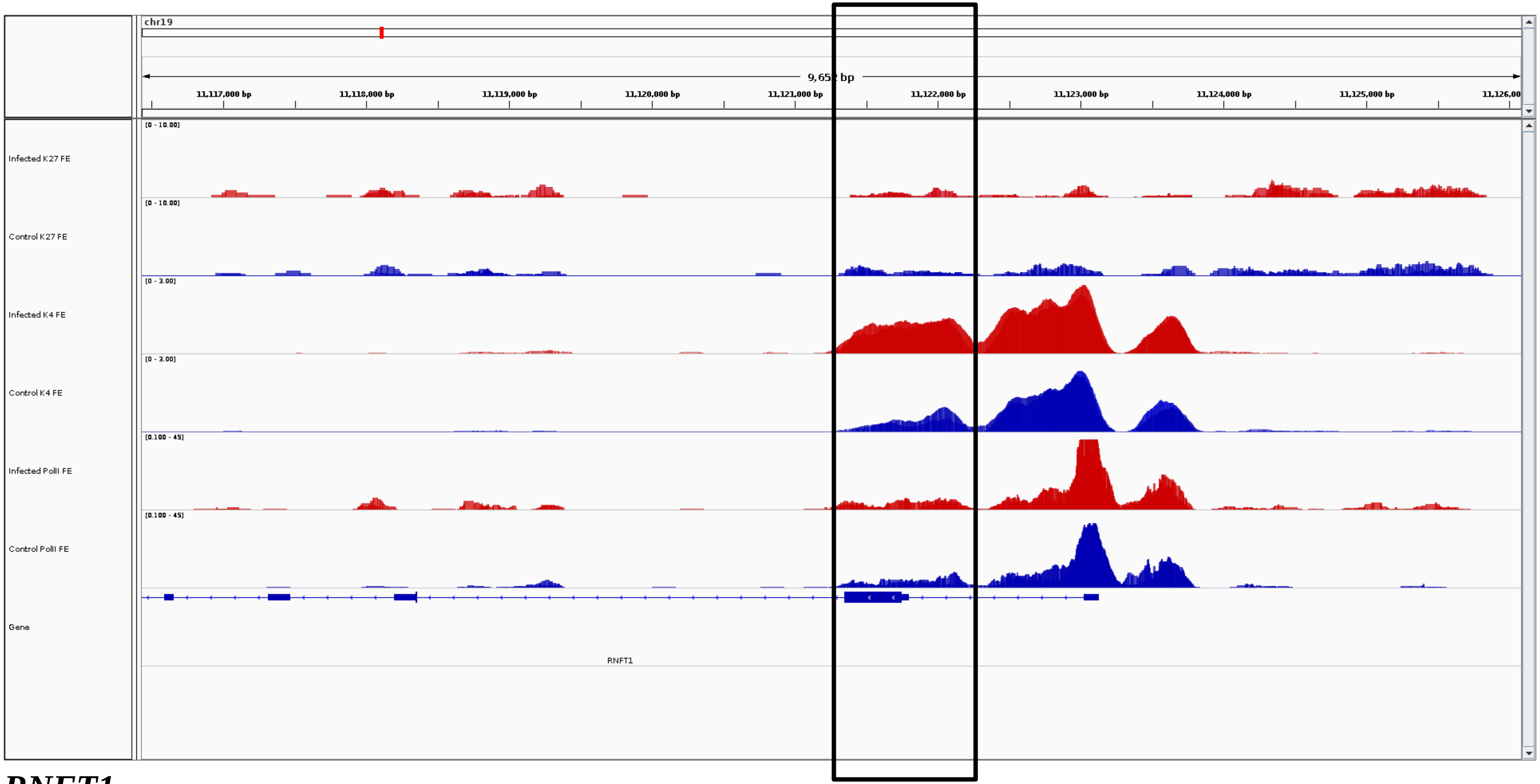
chr19
9,652 bp
11,117,000 bp
11,118,000 bp
11,119,000 bp
11,120,000 bp
11,121,000 bp
11,122,000 bp
11,123,000 bp
11,124,000 bp
11,125,000 bp
Infected K27 FE
Control K27 FE
Infected K4 FE
Control K4 FE
Infected PolII FE
Control PolII FE
Gene
RNFT1

***RNFT1***

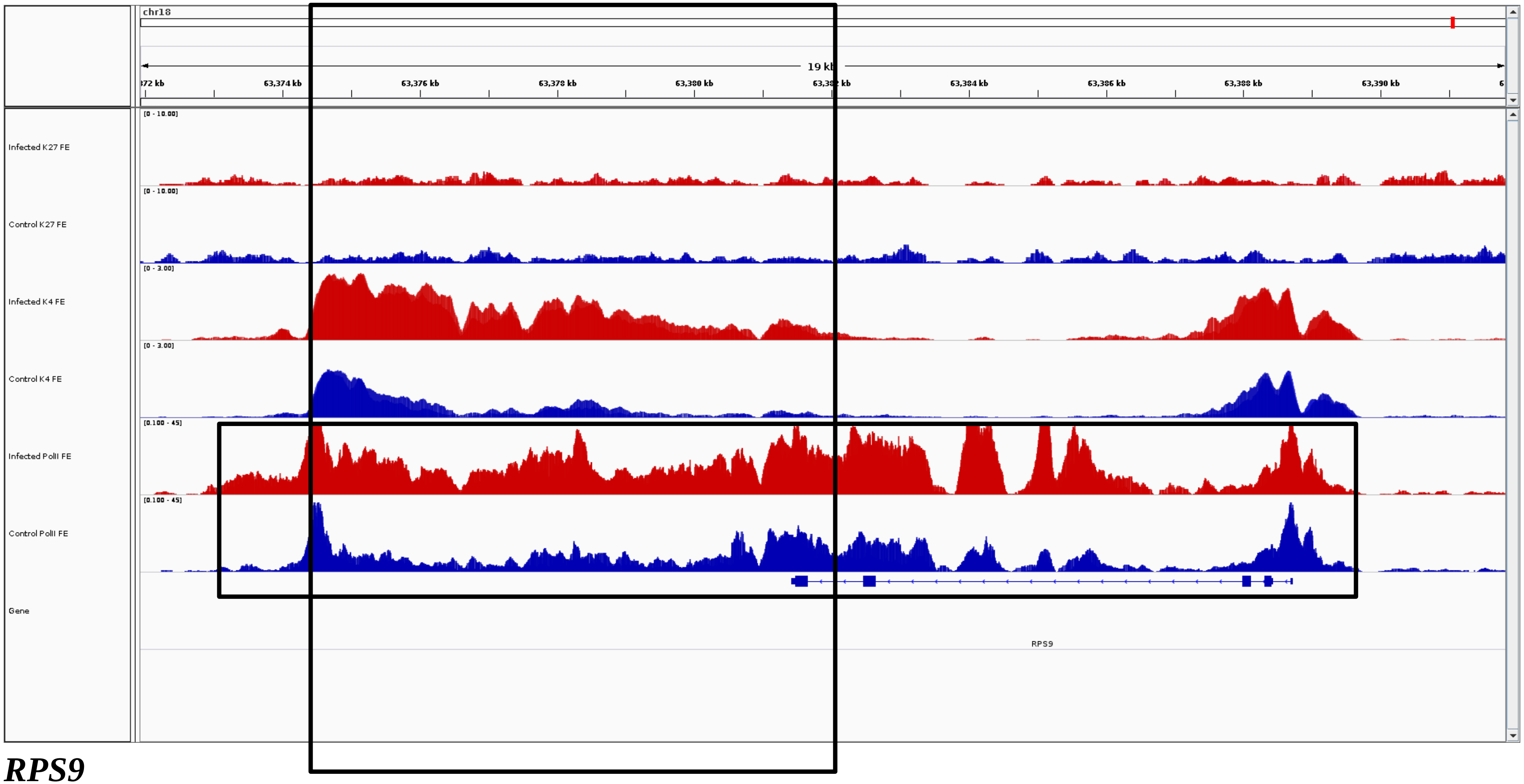
chr18
19 kb
63,374 kb
63,376 kb
63,378 kb
63,380 kb
63,384 kb
63,386 kb
63,388 kb
63,390 kb
Infected K27 FE
Control K27 FE
Infected K4 FE
Control K4 FE
Infected PolII FE
Control PolII FE
Gene
[0 - 10.00]
[0 - 3.00]
[0.100 - 45]
RPS9

***RPS9***

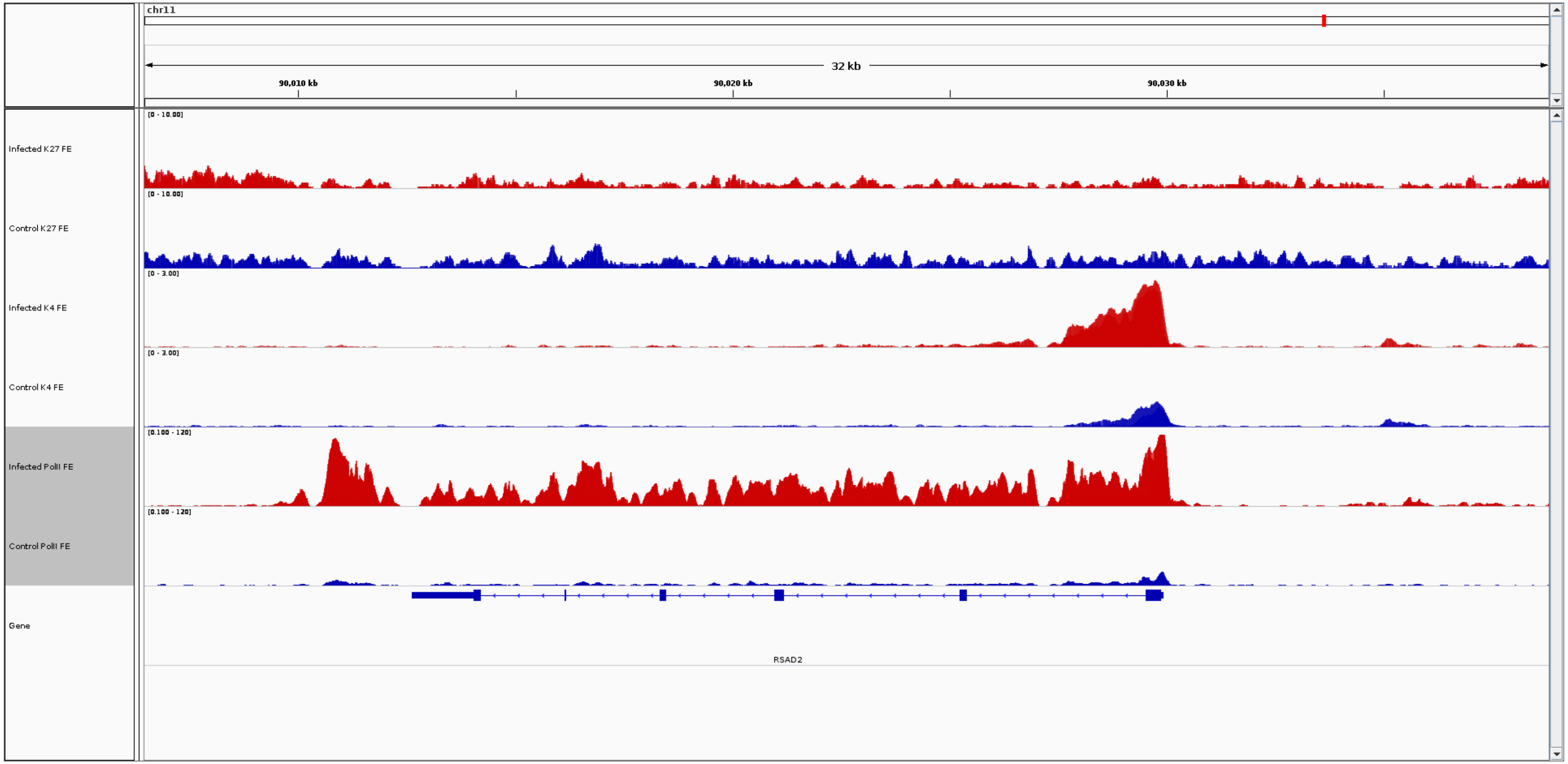
chr11
32 kb
90,010 kb
90,020 kb
90,030 kb
Infected K27 FE
[0 - 10.00]
Control K27 FE
[0 - 10.00]
Infected K4 FE
[0 - 3.00]
Control K4 FE
[0 - 3.00]
Infected PolII FE
[0.100 - 120]
Control PolII FE
[0.100 - 120]
Gene
RSAD2

***RSAD2***

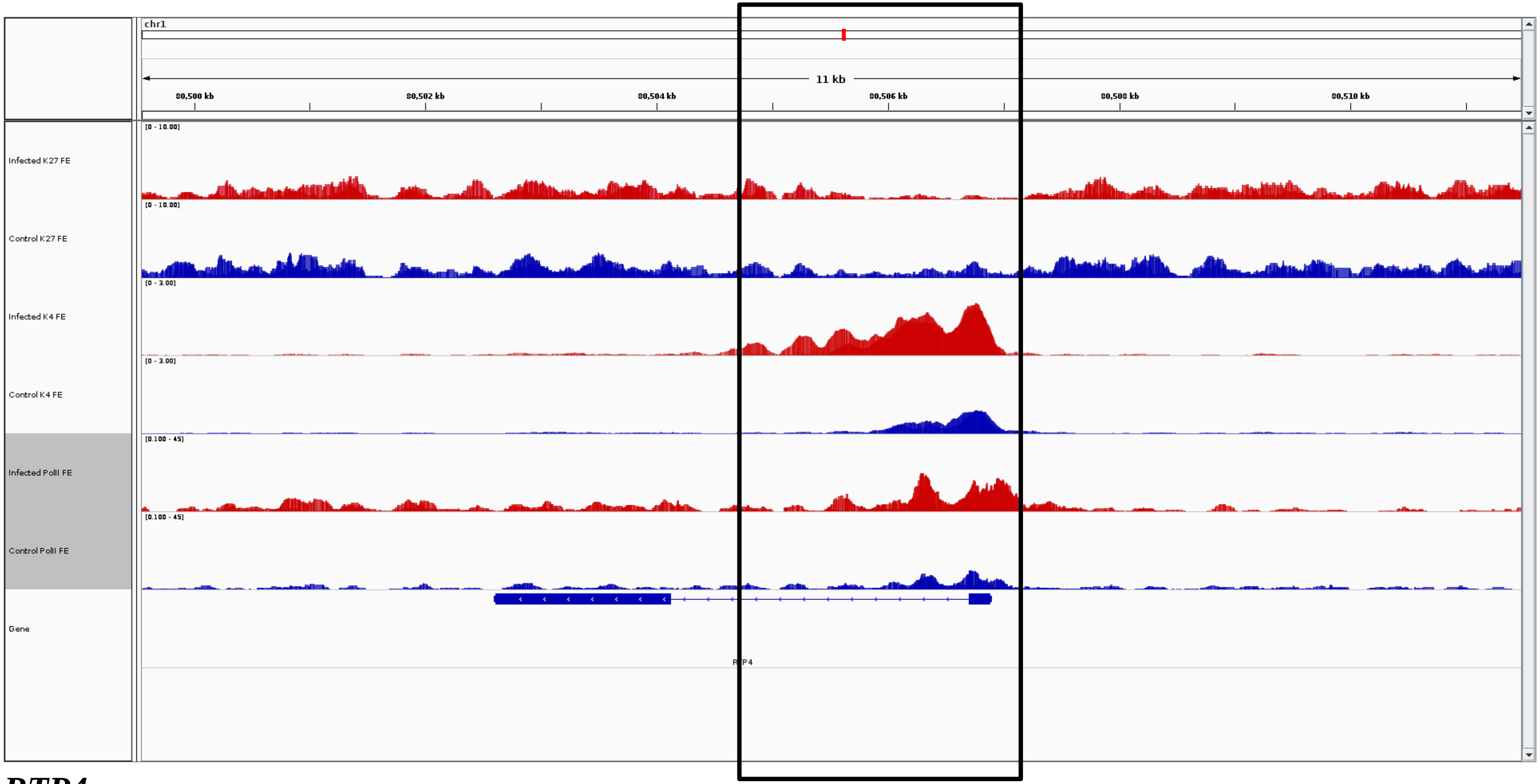
chr1
11 kb
80,500 kb
80,502 kb
80,504 kb
80,506 kb
80,508 kb
80,510 kb
Infected K27 FE
Control K27 FE
Infected K4 FE
Control K4 FE
Infected PolII FE
Control PolII FE
Gene

***RTP4***

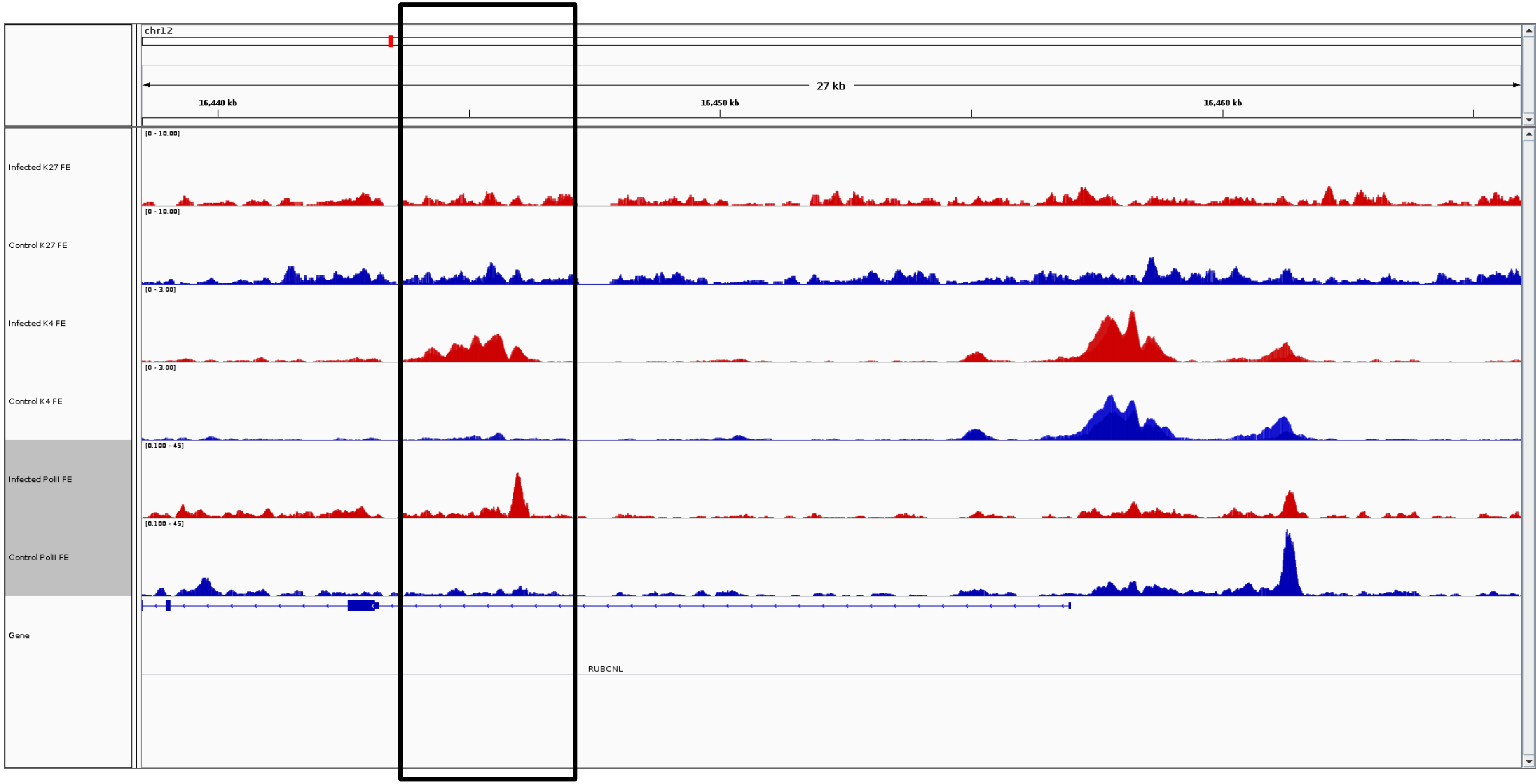
chr12
27 kb
16,440 kb
16,450 kb
16,460 kb
Infected K27 FE
[0 - 10.00]
Control K27 FE
[0 - 10.00]
Infected K4 FE
[0 - 3.00]
Control K4 FE
[0 - 3.00]
Infected PolII FE
[0.100 - 45]
Control PolII FE
[0.100 - 45]
Gene
RUBCNL

***RUBCNL***

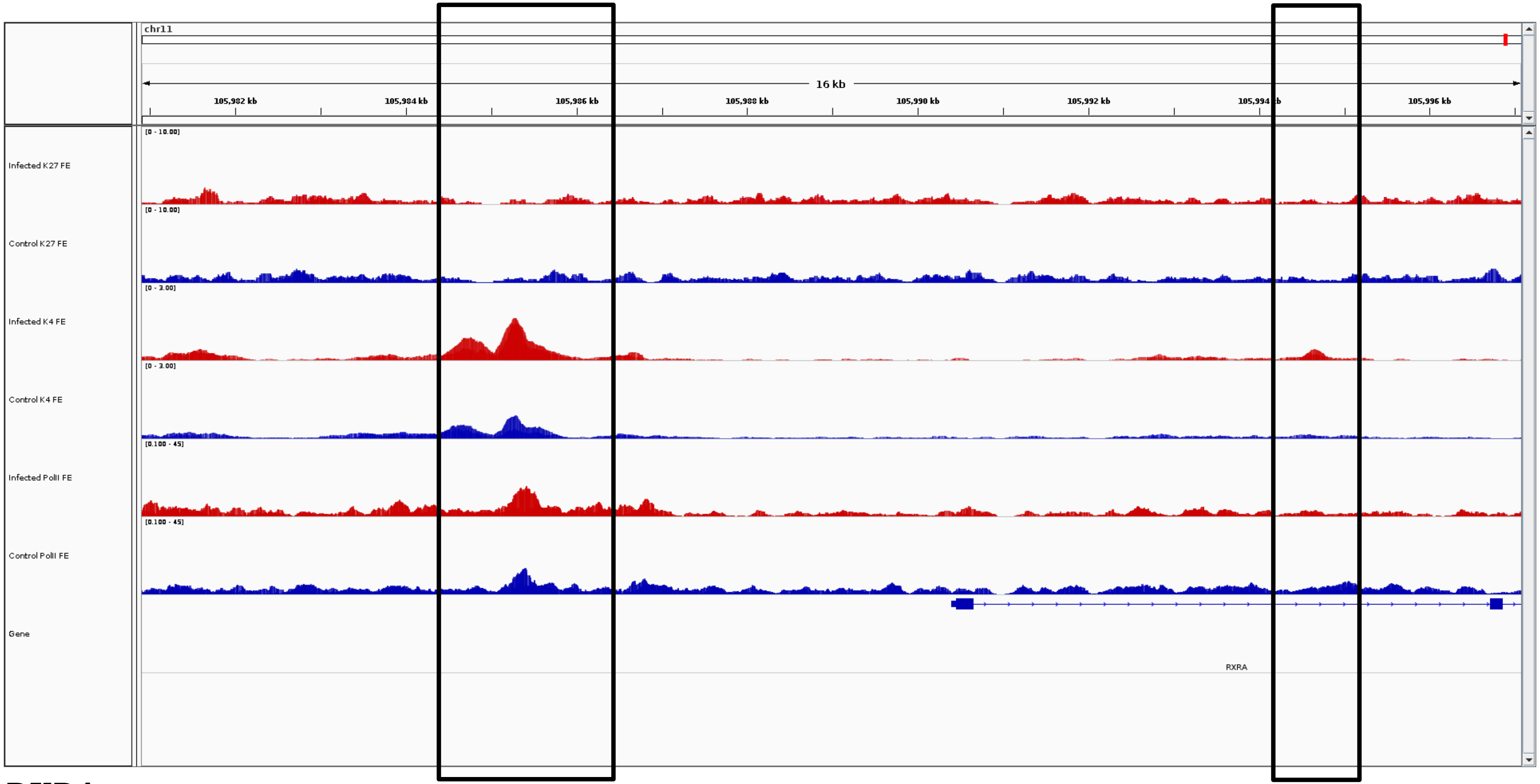
chr11
16 kb
105,982 kb
105,984 kb
105,986 kb
105,988 kb
105,990 kb
105,992 kb
105,994 kb
105,996 kb
Infected K27 FE
Control K27 FE
Infected K4 FE
Control K4 FE
Infected PolII FE
Control PolII FE
Gene
RXRA

***RXRA***

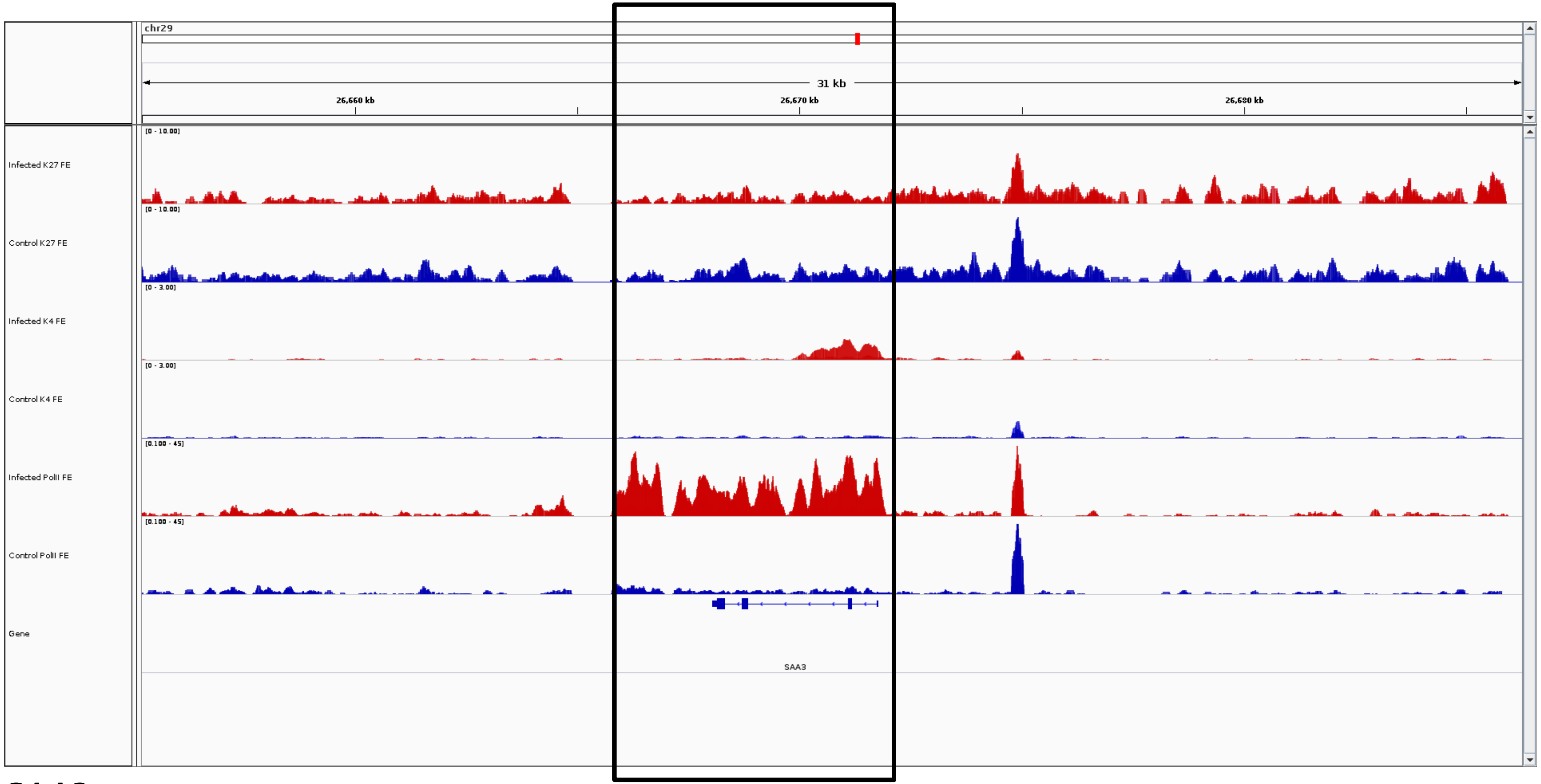

chr29
31 kb
26,660 kb
26,670 kb
26,680 kb
Infected K27 FE
Control K27 FE
Infected K4 FE
Control K4 FE
Infected PolII FE
Control PolII FE
Gene
SAA3


***SAA3***

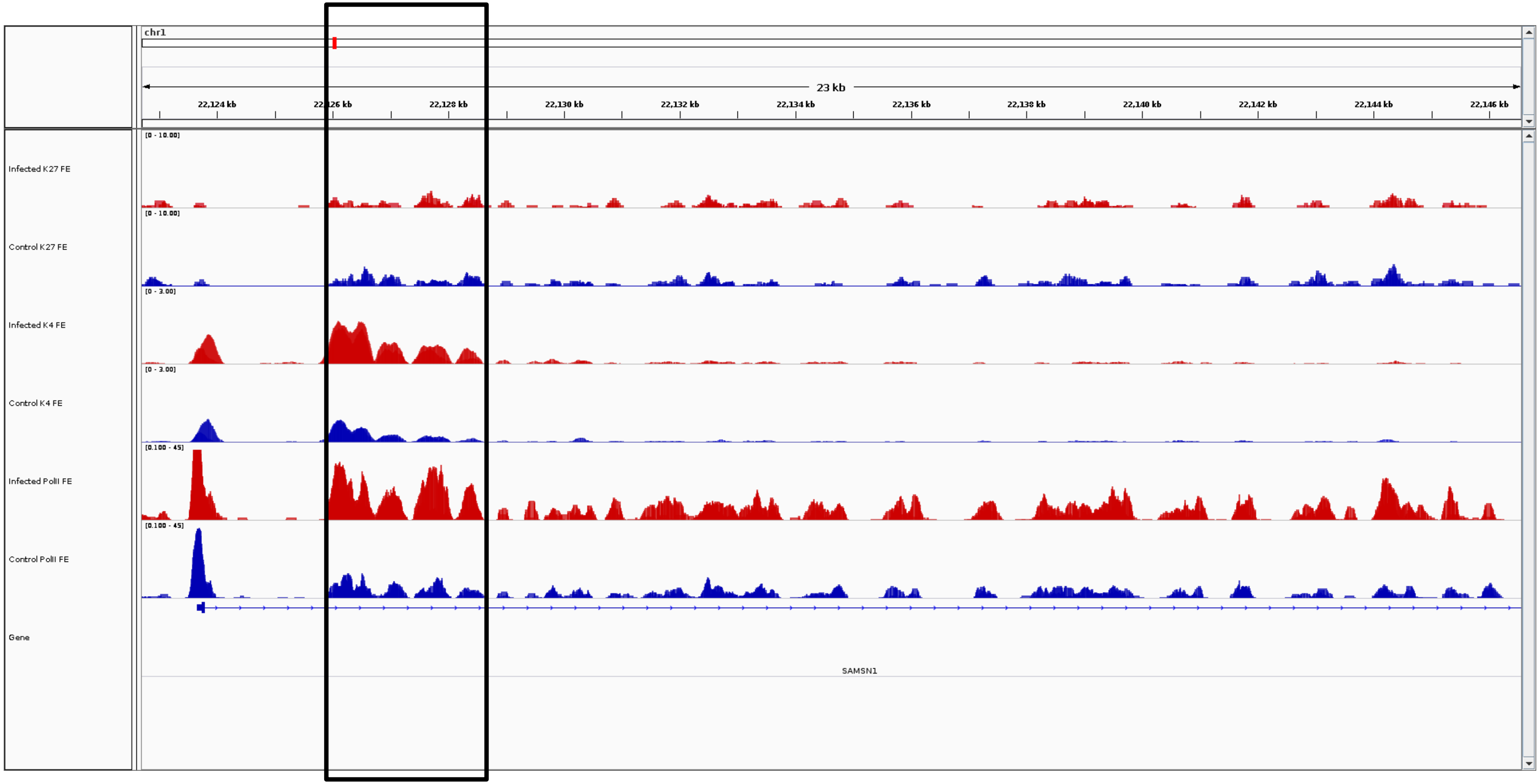
chr1
23 kb
22,124 kb
22,126 kb
22,128 kb
22,130 kb
22,132 kb
22,134 kb
22,136 kb
22,138 kb
22,140 kb
22,142 kb
22,144 kb
22,146 kb
Infected K27 FE
[0 - 10.00]
Control K27 FE
[0 - 10.00]
Infected K4 FE
[0 - 3.00]
Control K4 FE
[0 - 3.00]
Infected PolII FE
[0.100 - 45]
Control PolII FE
[0.100 - 45]
Gene
SAMSN1

***SAMSN1***

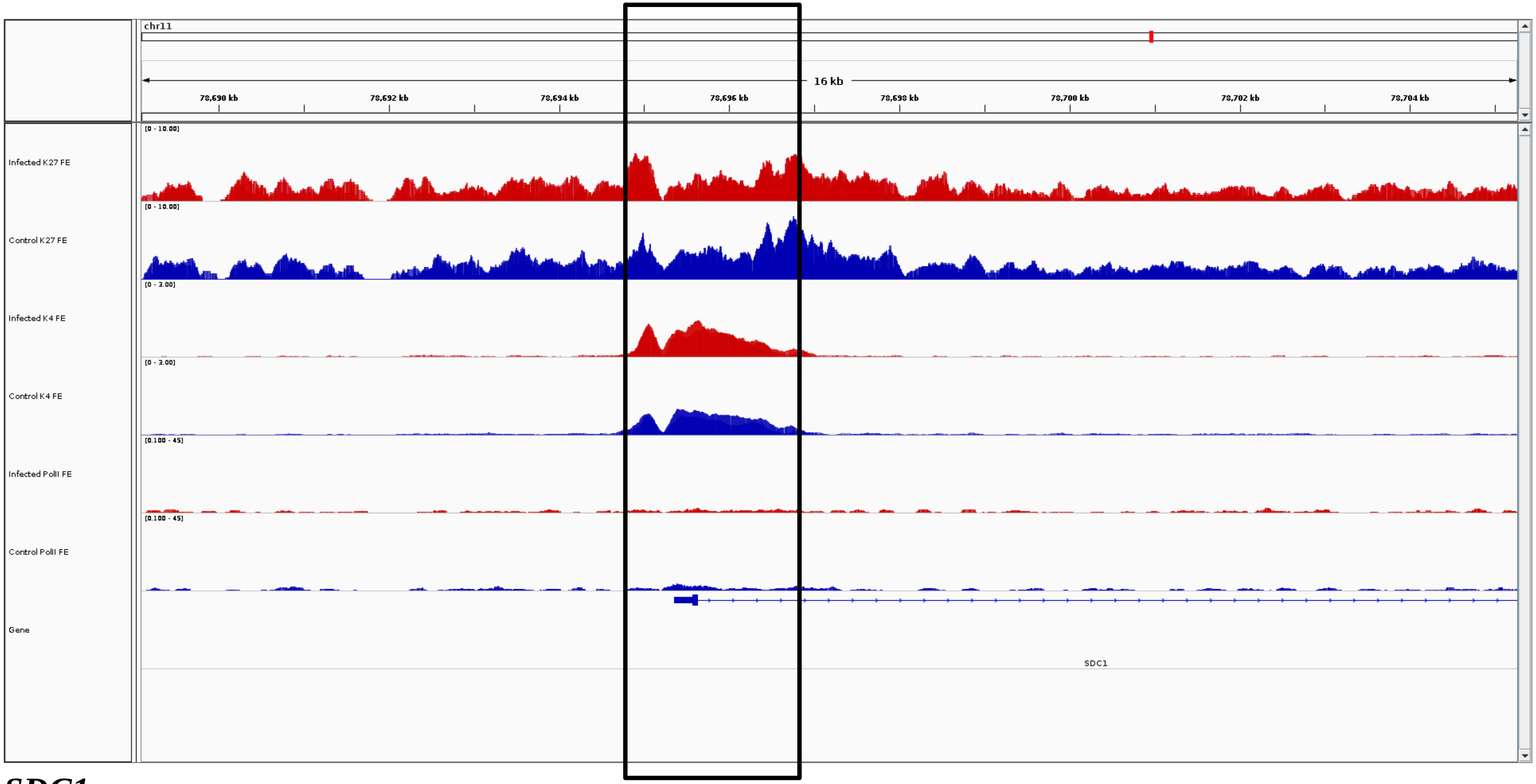
chr11
16 kb
78,690 kb
78,692 kb
78,694 kb
78,696 kb
78,698 kb
78,700 kb
78,702 kb
78,704 kb
Infected K27 FE
Control K27 FE
Infected K4 FE
Control K4 FE
Infected PolII FE
Control PolII FE
Gene
SDC1

***SDC1***

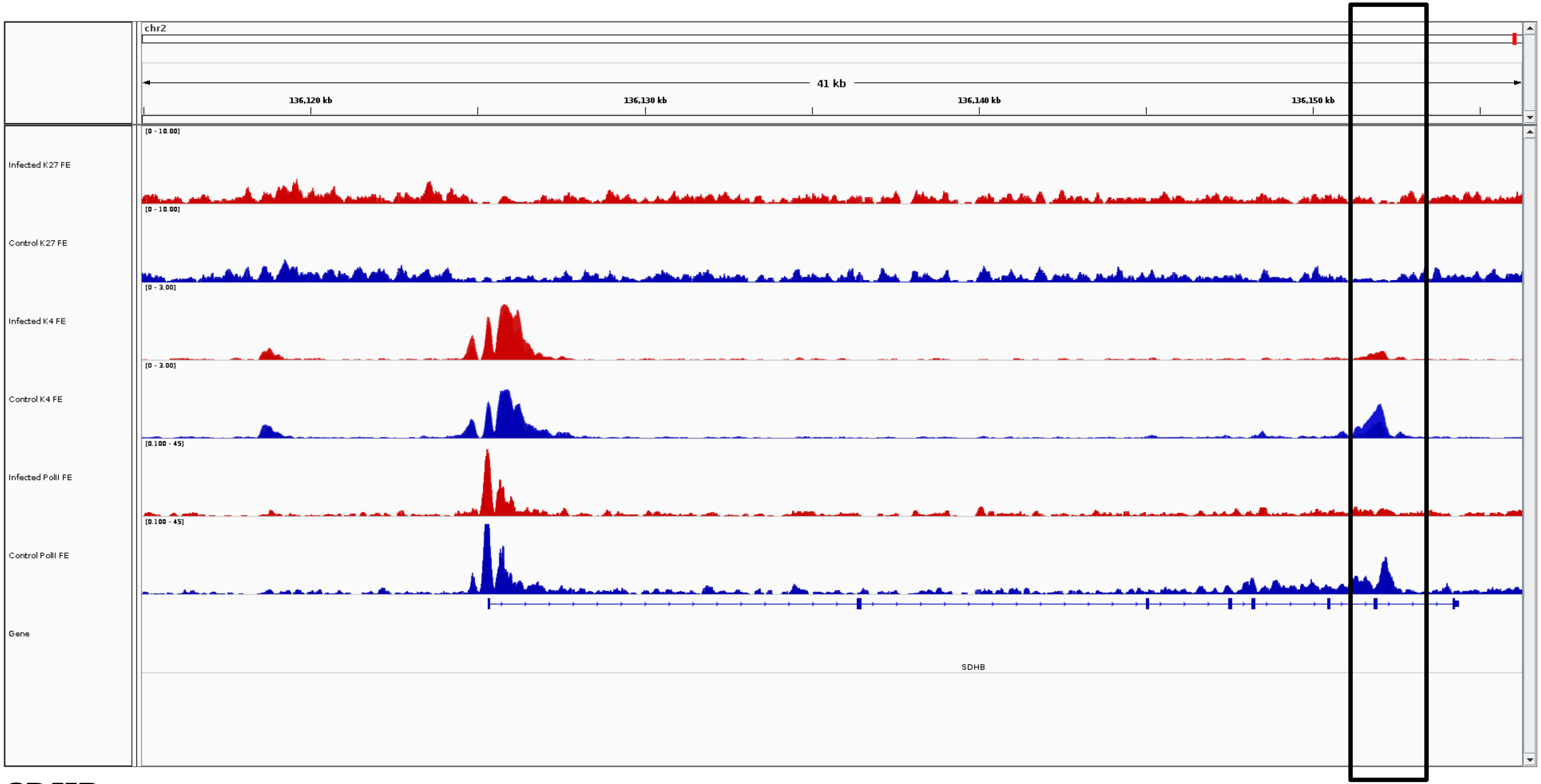

chr2
41 kb
136,120 kb
136,130 kb
136,140 kb
136,150 kb
Infected K27 FE
[0 - 10.00]
Control K27 FE
[0 - 10.00]
Infected K4 FE
[0 - 3.00]
Control K4 FE
[0 - 3.00]
Infected PolII FE
[0.100 - 45]
Control PolII FE
[0.100 - 45]
Gene
SDHB


***SDHB***

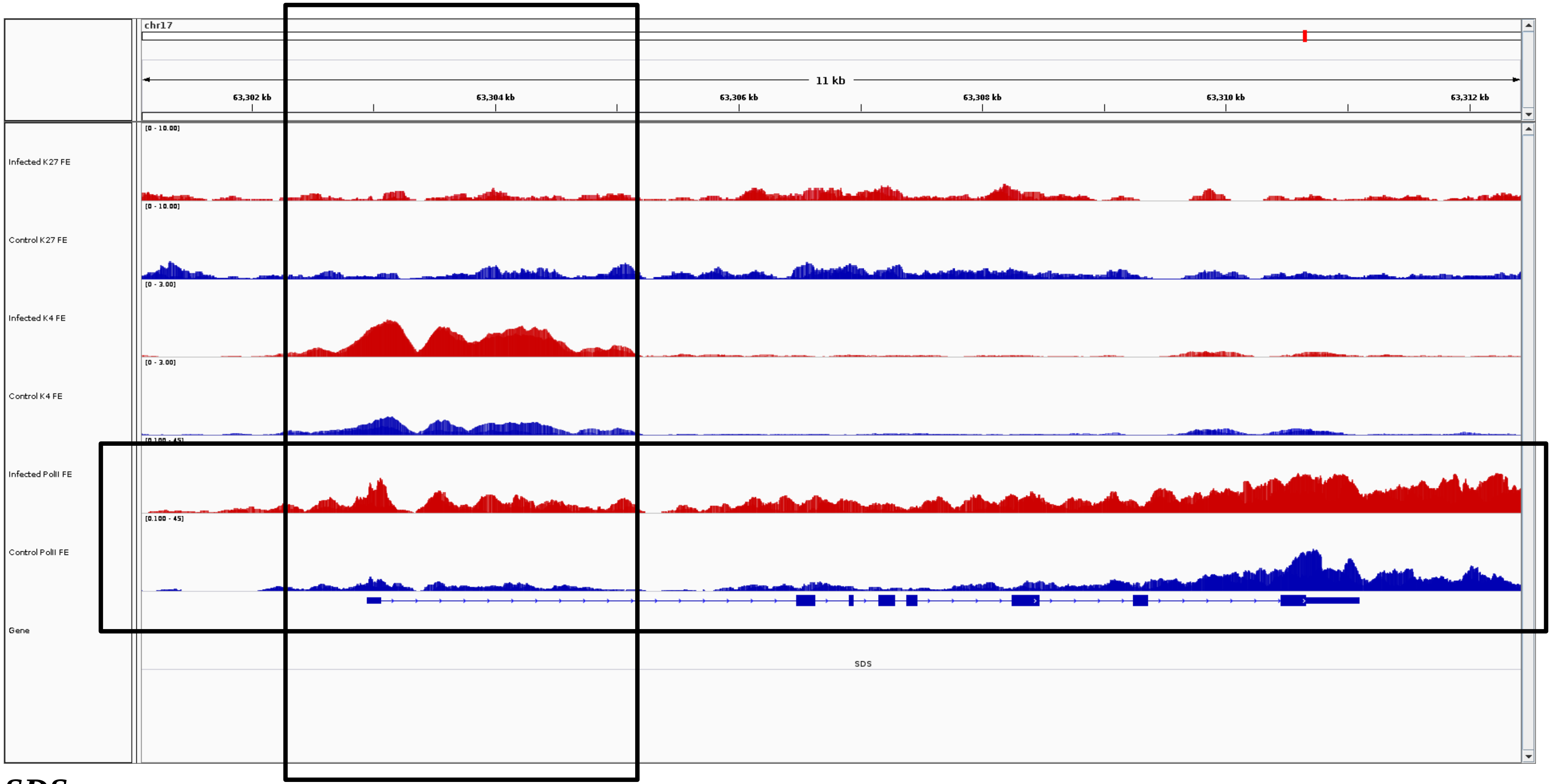
chr17
11 kb
63,302 kb
63,304 kb
63,306 kb
63,308 kb
63,310 kb
63,312 kb
Infected K27 FE
Control K27 FE
Infected K4 FE
Control K4 FE
Infected PolII FE
Control PolII FE
Gene
SDS

***SDS***

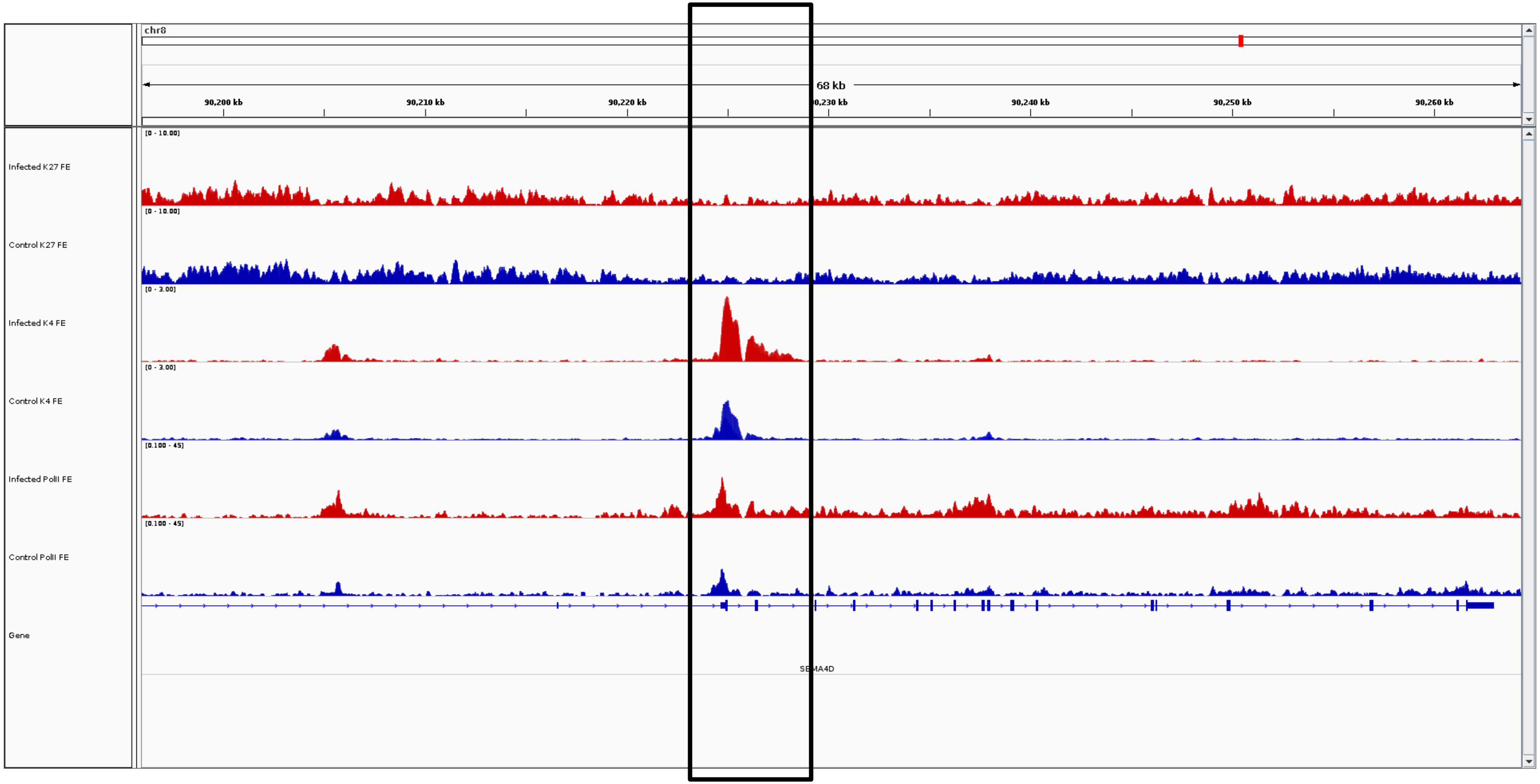
chr8
68 kb
90,200 kb
90,210 kb
90,220 kb
0,230 kb
90,240 kb
90,250 kb
90,260 kb
[0 - 10.00]
Infected K27 FE
[0 - 10.00]
Control K27 FE
[0 - 3.00]
Infected K4 FE
[0 - 3.00]
Control K4 FE
[0.100 - 45]
Infected PolII FE
[0.100 - 45]
Control PolII FE
Gene
SEMA4D

***SEMA4D***

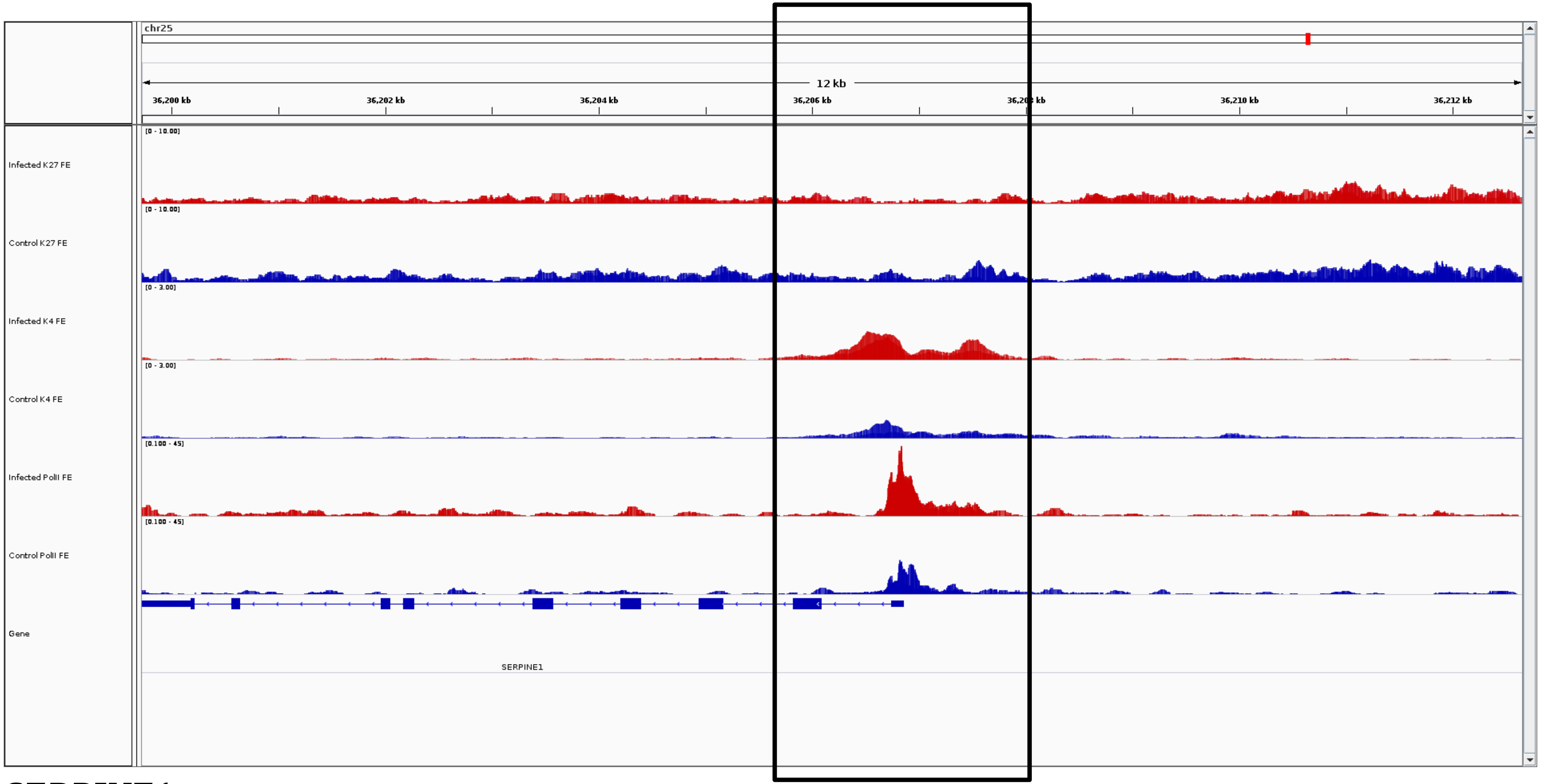
chr25
12 kb
36,200 kb
36,202 kb
36,204 kb
36,206 kb
36,210 kb
36,212 kb
[0 - 10.00]
Infected K27 FE
Control K27 FE
[0 - 3.00]
Infected K4 FE
Control K4 FE
[0.100 - 45]
Infected PolII FE
Control PolII FE
Gene
SERPINE1

***SERPINE1***

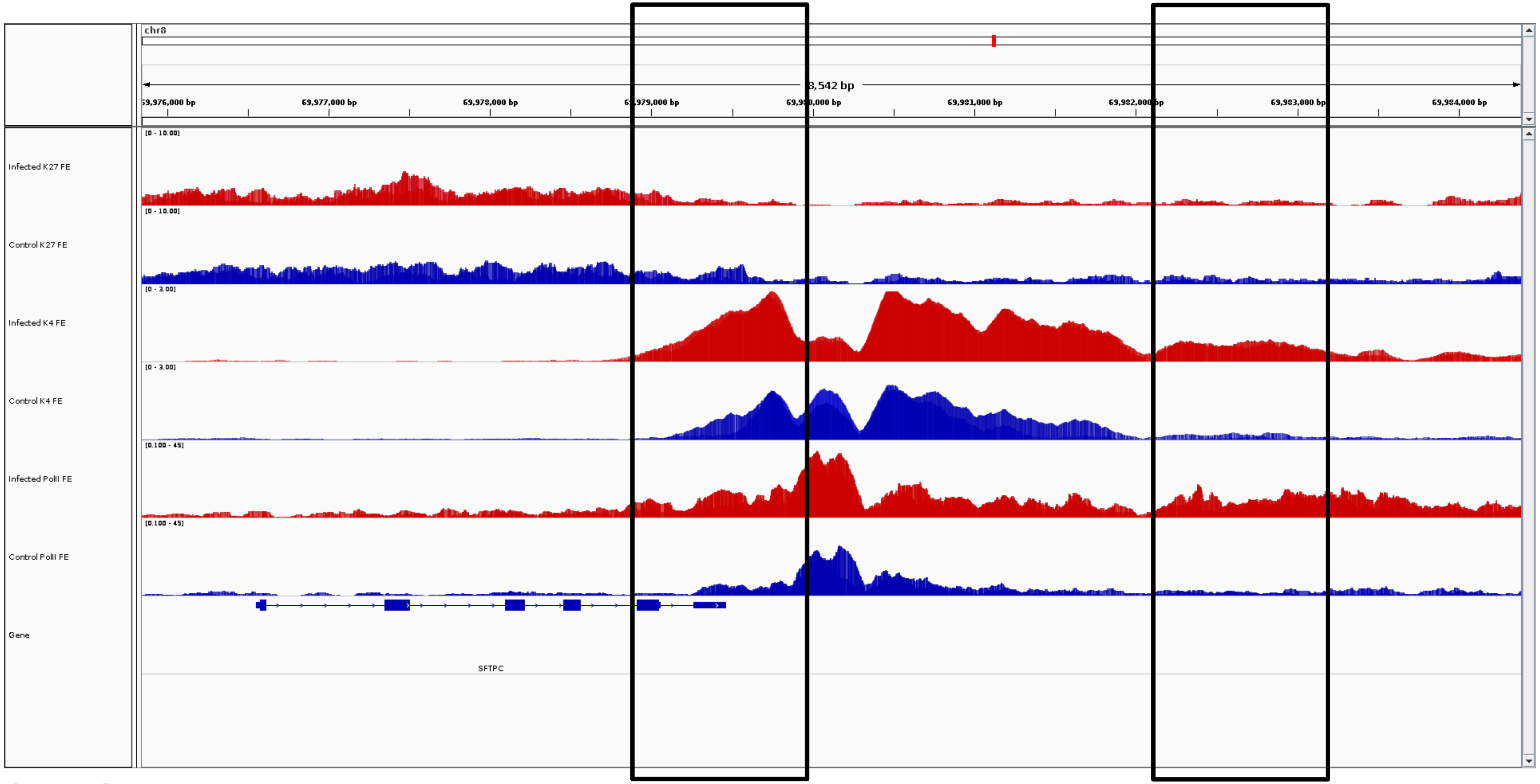

chr8
8,542 bp
69,976,000 bp
69,977,000 bp
69,978,000 bp
69,979,000 bp
69,980,000 bp
69,981,000 bp
69,982,000 bp
69,983,000 bp
69,984,000 bp
Infected K27 FE
[0 - 10.00]
Control K27 FE
[0 - 10.00]
Infected K4 FE
[0 - 3.00]
Control K4 FE
[0 - 3.00]
Infected PolII FE
[0.100 - 45]
Control PolII FE
[0.100 - 45]
Gene
SFTPC


***SFTPC***

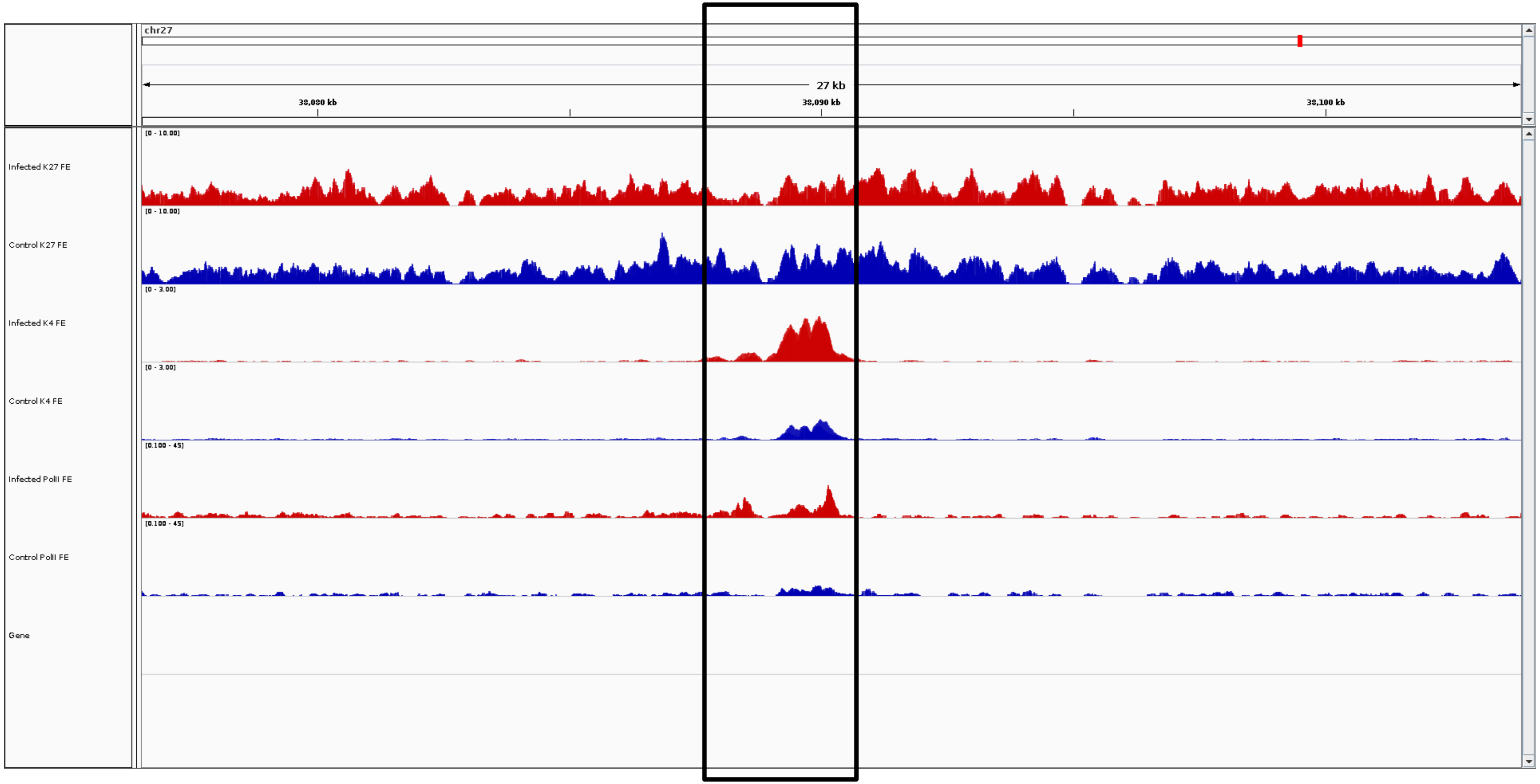
chr27
27 kb
38,080 kb
38,090 kb
38,100 kb
Infected K27 FE
[0 - 10.00]
Control K27 FE
[0 - 10.00]
Infected K4 FE
[0 - 3.00]
Control K4 FE
[0 - 3.00]
Infected PolII FE
[0.100 - 45]
Control PolII FE
[0.100 - 45]
Gene

***SH2D4A***

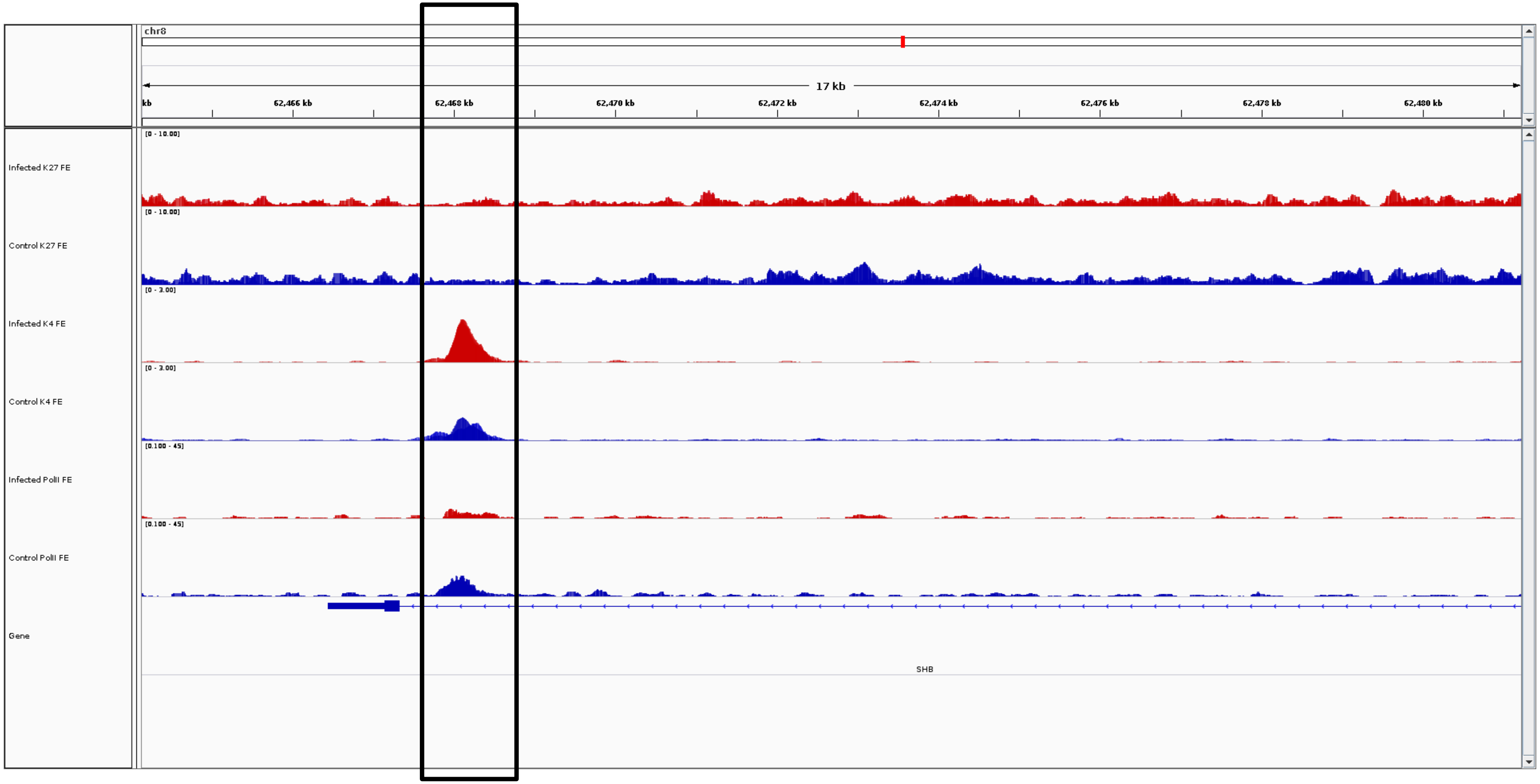
chr8
17 kb
kb
62,466 kb
62,468 kb
62,470 kb
62,472 kb
62,474 kb
62,476 kb
62,478 kb
62,480 kb
Infected K27 FE
[0 - 10.00]
Control K27 FE
[0 - 10.00]
Infected K4 FE
[0 - 3.00]
Control K4 FE
[0 - 3.00]
Infected PolII FE
[0.100 - 45]
Control PolII FE
[0.100 - 45]
Gene
SHB

***SHB***

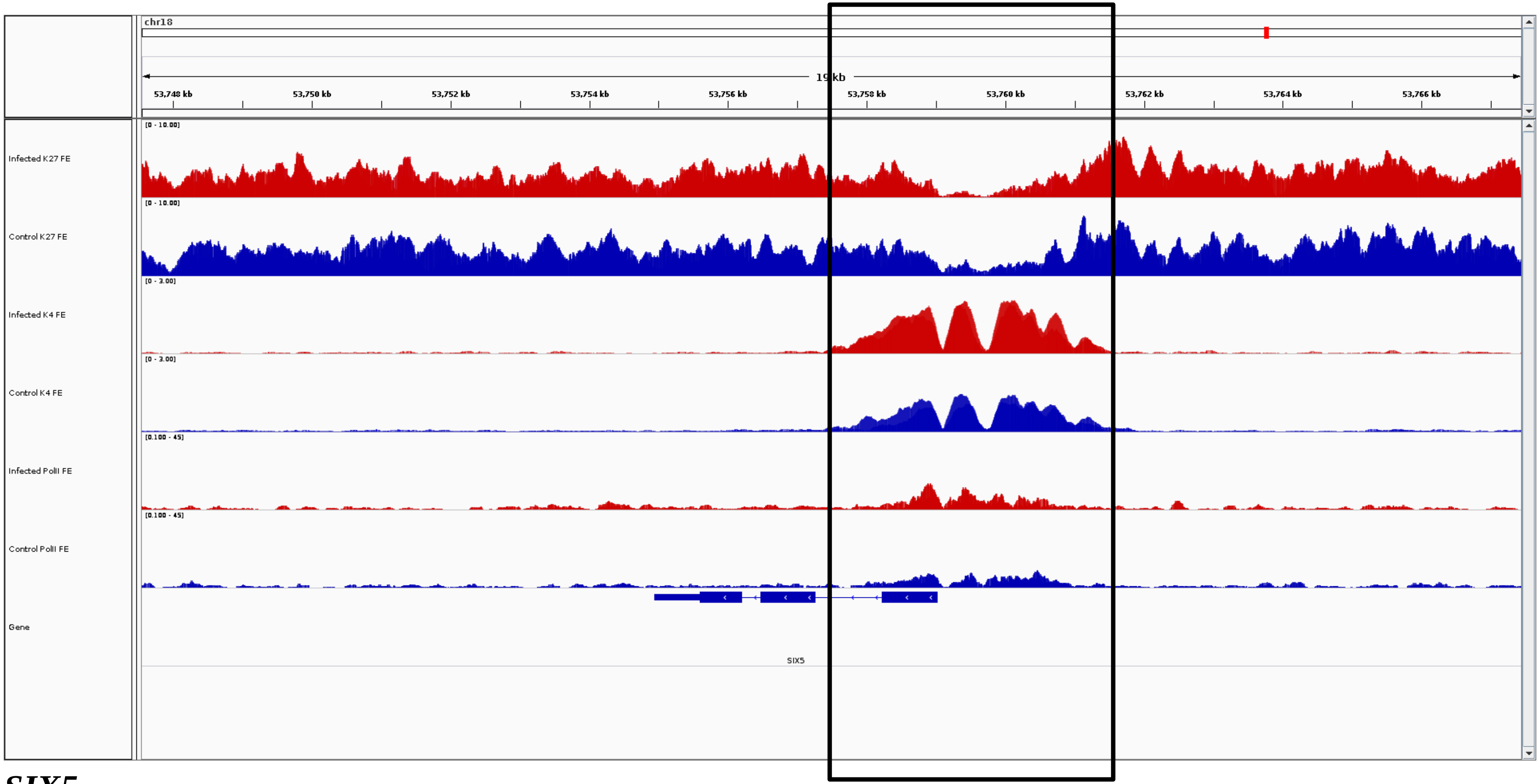
chr18
19 kb
53,748 kb
53,750 kb
53,752 kb
53,754 kb
53,756 kb
53,758 kb
53,760 kb
53,762 kb
53,764 kb
53,766 kb
Infected K27 FE
[0 - 10.00]
Control K27 FE
[0 - 10.00]
Infected K4 FE
[0 - 3.00]
Control K4 FE
[0 - 3.00]
Infected PolII FE
[0.100 - 45]
Control PolII FE
[0.100 - 45]
Gene
SIX5

***SIX5***

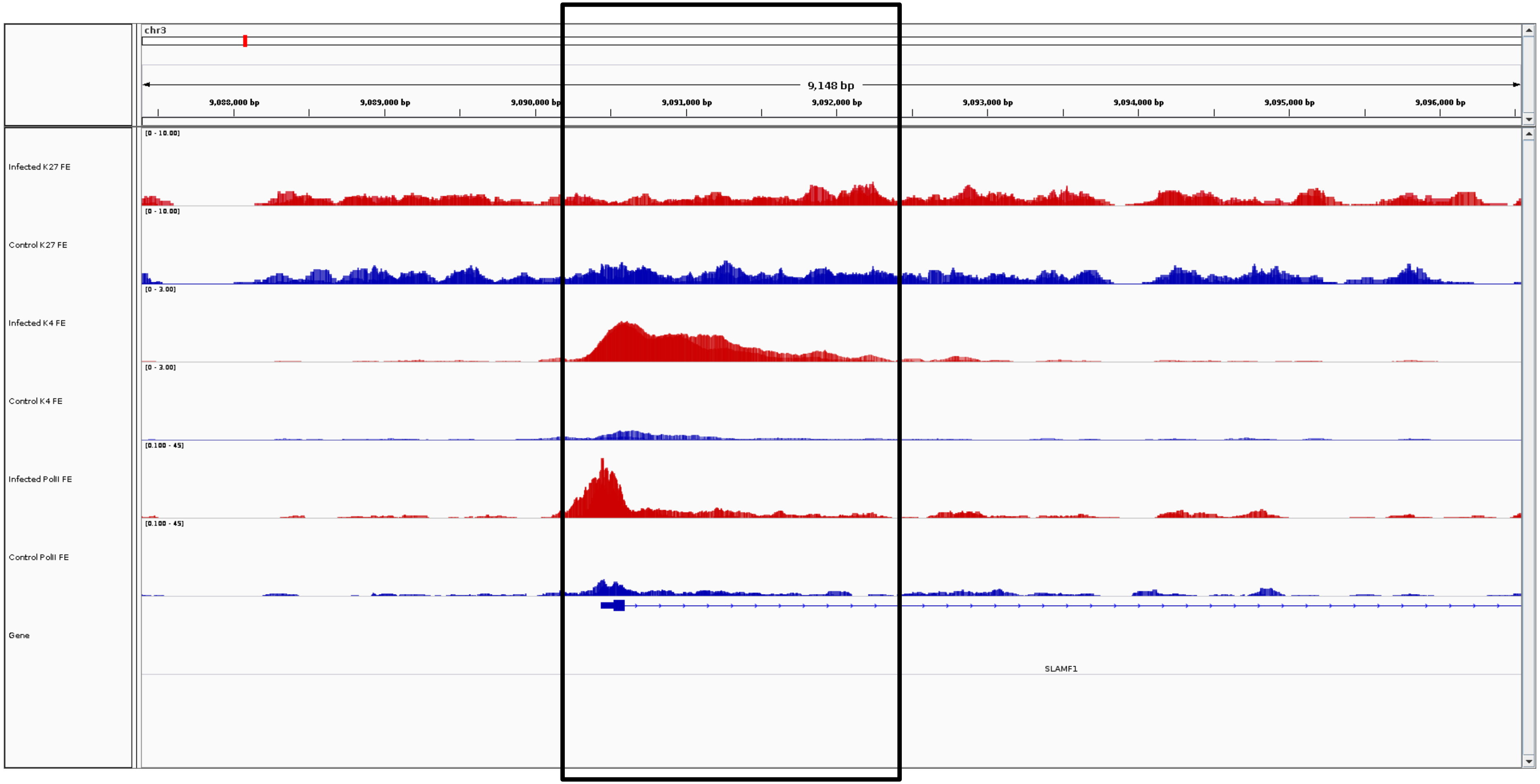
chr3
9,148 bp
9,088,000 bp
9,089,000 bp
9,090,000 bp
9,091,000 bp
9,092,000 bp
9,093,000 bp
9,094,000 bp
9,095,000 bp
9,096,000 bp
Infected K27 FE
[0 - 10.00]
Control K27 FE
[0 - 10.00]
Infected K4 FE
[0 - 3.00]
Control K4 FE
[0 - 3.00]
Infected PolII FE
[0.100 - 45]
Control PolII FE
[0.100 - 45]
Gene
SLAMF1

***SLAMF1***

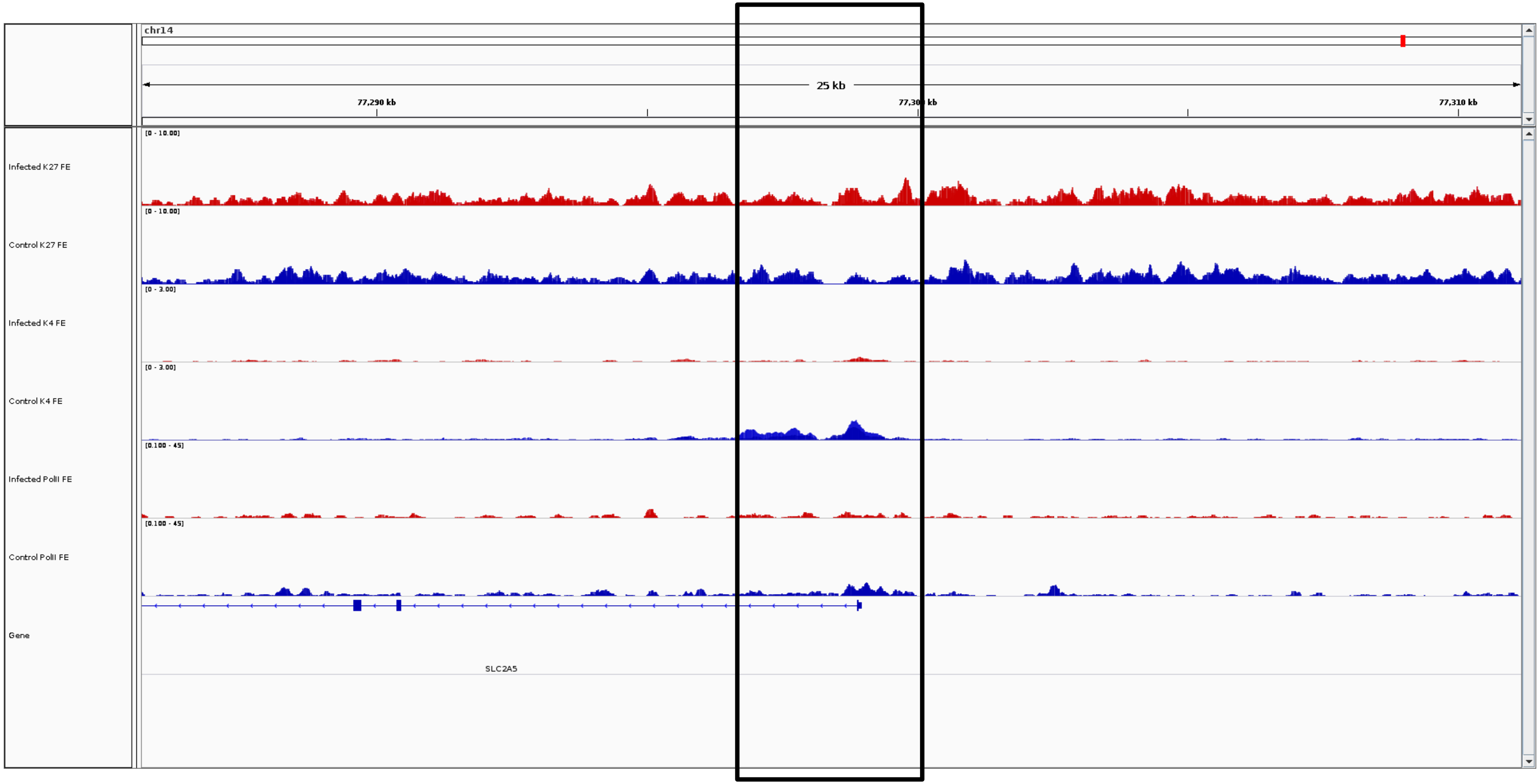
chr14
25 kb
77,290 kb
77,310 kb
Infected K27 FE
[0 - 10.00]
Control K27 FE
[0 - 10.00]
Infected K4 FE
[0 - 3.00]
Control K4 FE
[0 - 3.00]
Infected PolII FE
[0.100 - 45]
Control PolII FE
[0.100 - 45]
Gene
SLC2A5

***SLC2A5***

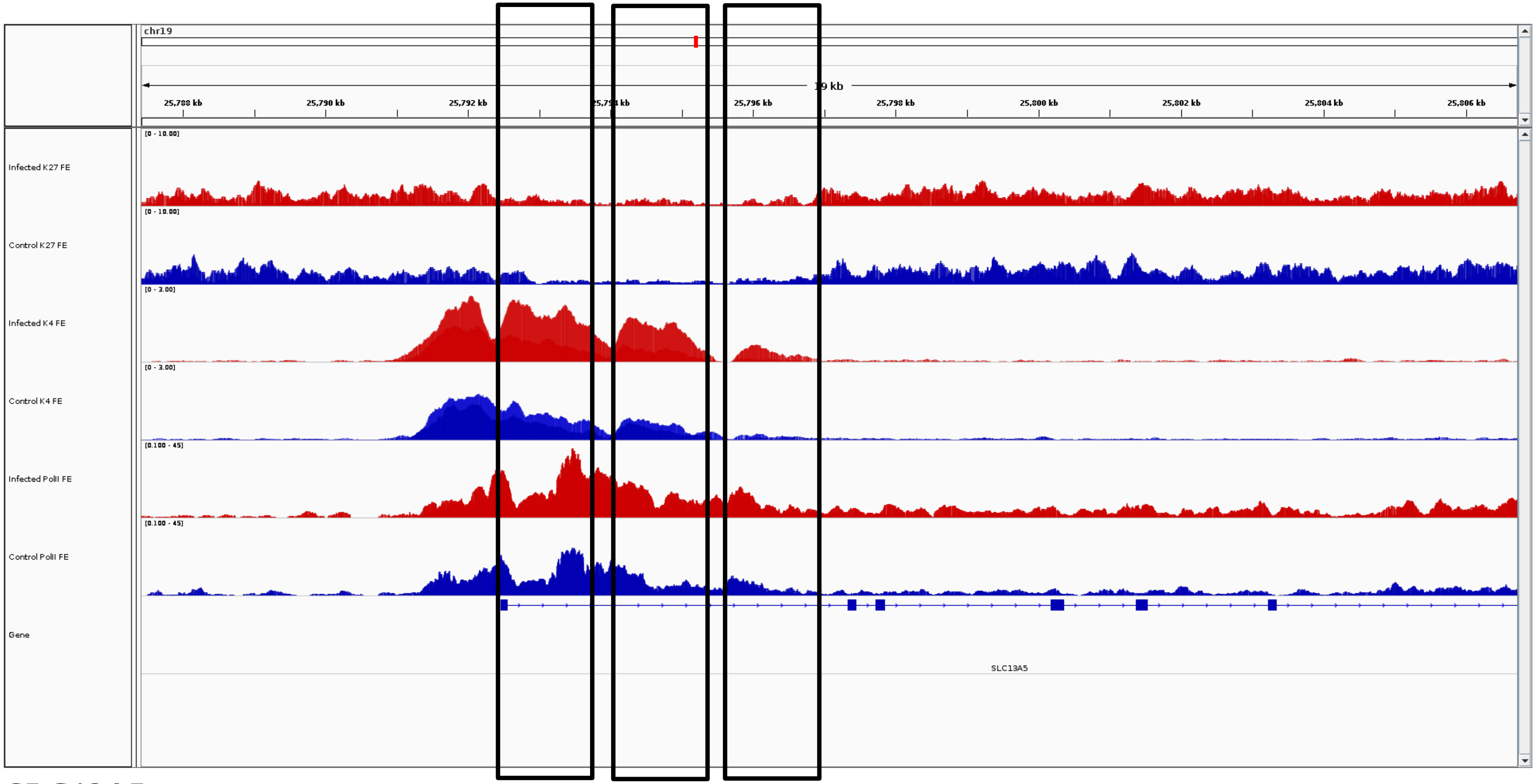
chr19
19 kb
25,788 kb
25,790 kb
25,792 kb
25,796 kb
25,798 kb
25,800 kb
25,802 kb
25,804 kb
25,806 kb
Infected K27 FE
Control K27 FE
Infected K4 FE
Control K4 FE
Infected PolII FE
Control PolII FE
Gene
SLC13A5

***SLC13A5***

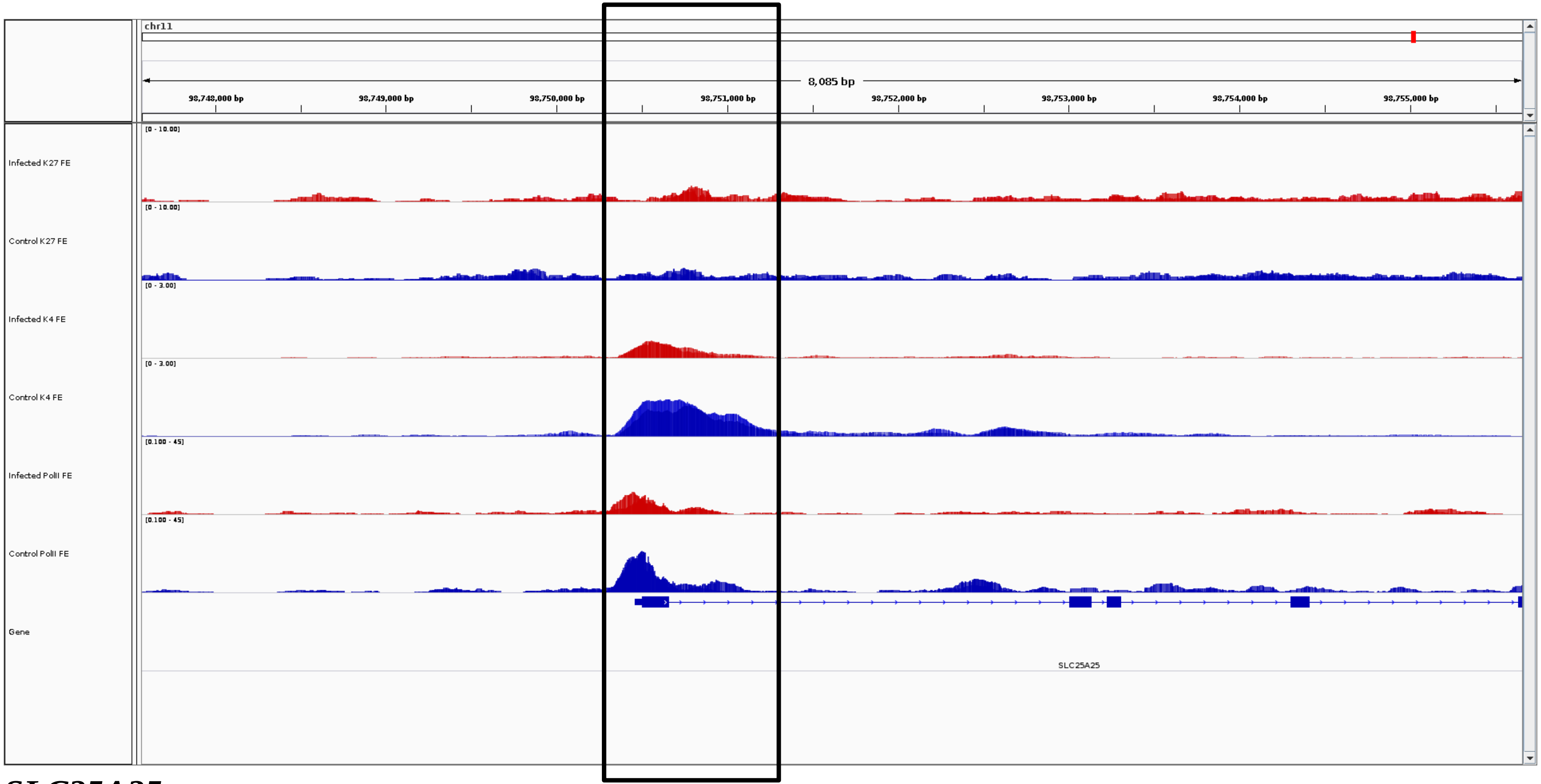
chr11
8,085 bp
98,748,000 bp
98,749,000 bp
98,750,000 bp
98,751,000 bp
98,752,000 bp
98,753,000 bp
98,754,000 bp
98,755,000 bp
Infected K27 FE
[0 - 10.00]
Control K27 FE
[0 - 10.00]
Infected K4 FE
[0 - 3.00]
Control K4 FE
[0 - 3.00]
Infected PolII FE
[0.100 - 45]
Control PolII FE
[0.100 - 45]
Gene
SLC25A25

***SLC25A25***

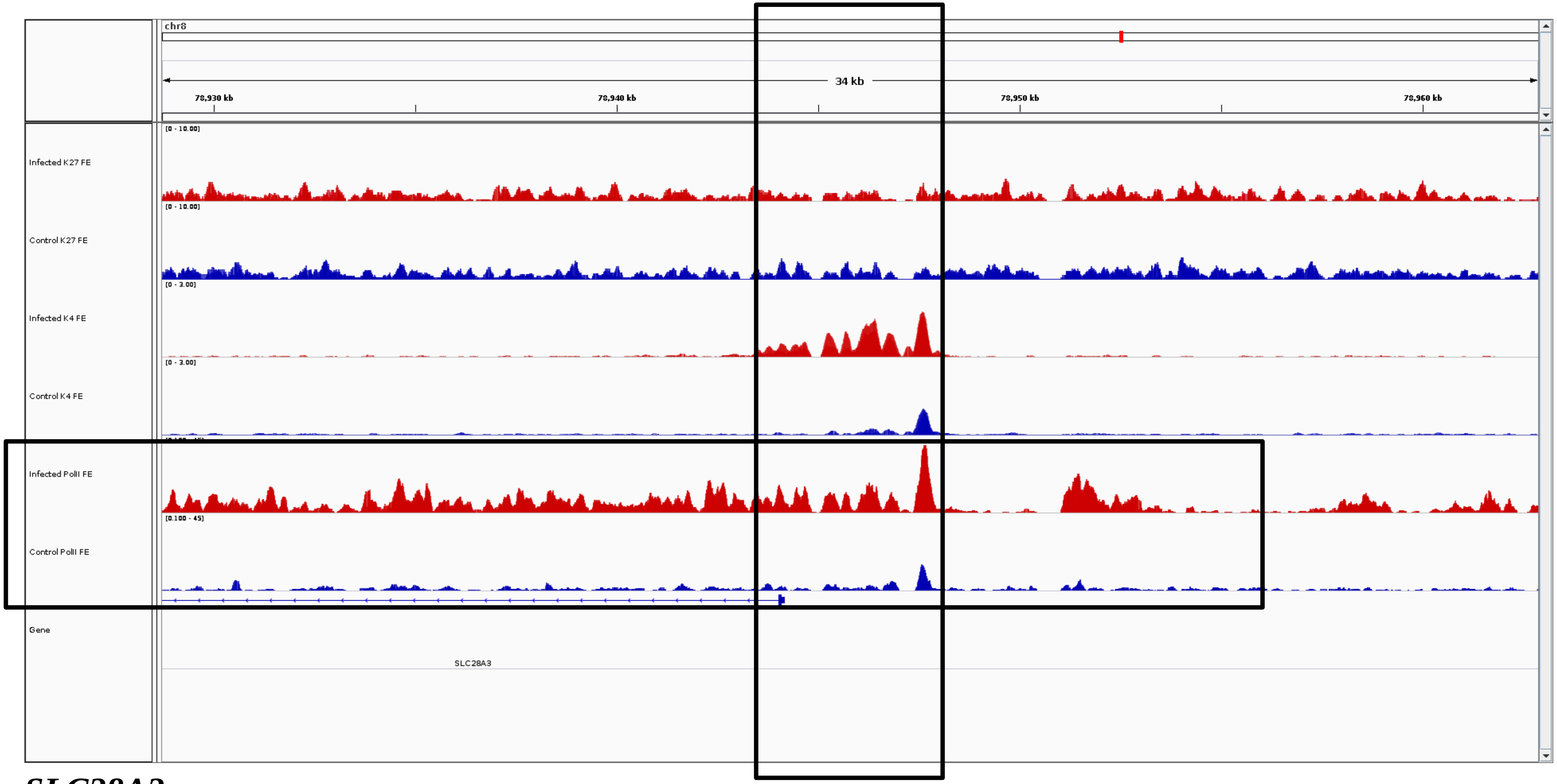
chr8
34 kb
78,930 kb
78,940 kb
78,950 kb
78,960 kb
Infected K27 FE
Control K27 FE
Infected K4 FE
Control K4 FE
Infected PolII FE
Control PolII FE
Gene
SLC28A3

***SLC28A3***

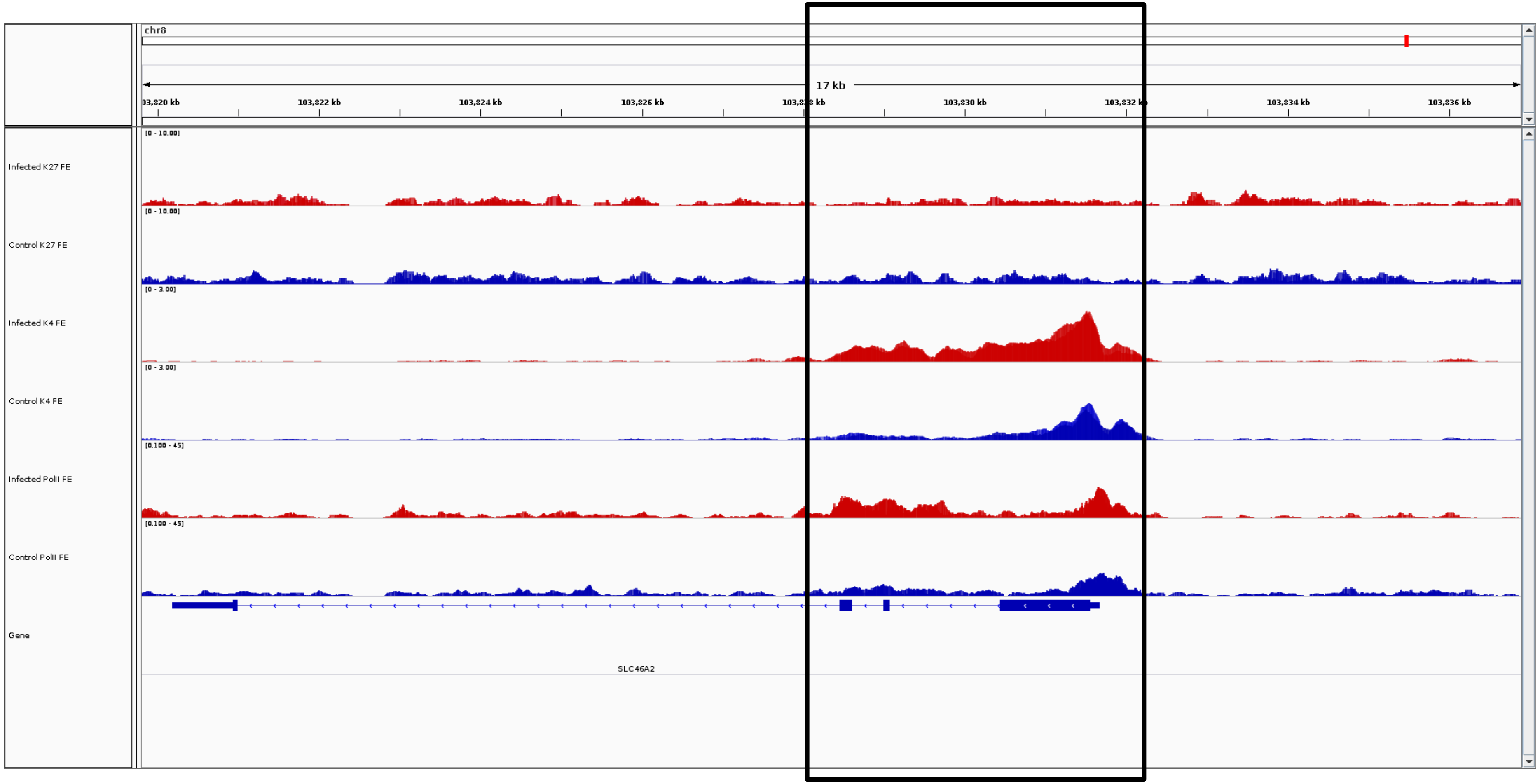
chr8
17 kb
03,820 kb
103,822 kb
103,824 kb
103,826 kb
103,830 kb
103,834 kb
103,836 kb
Infected K27 FE
Control K27 FE
Infected K4 FE
Control K4 FE
Infected PolII FE
Control PolII FE
Gene
SLC46A2

***SLC46A2***

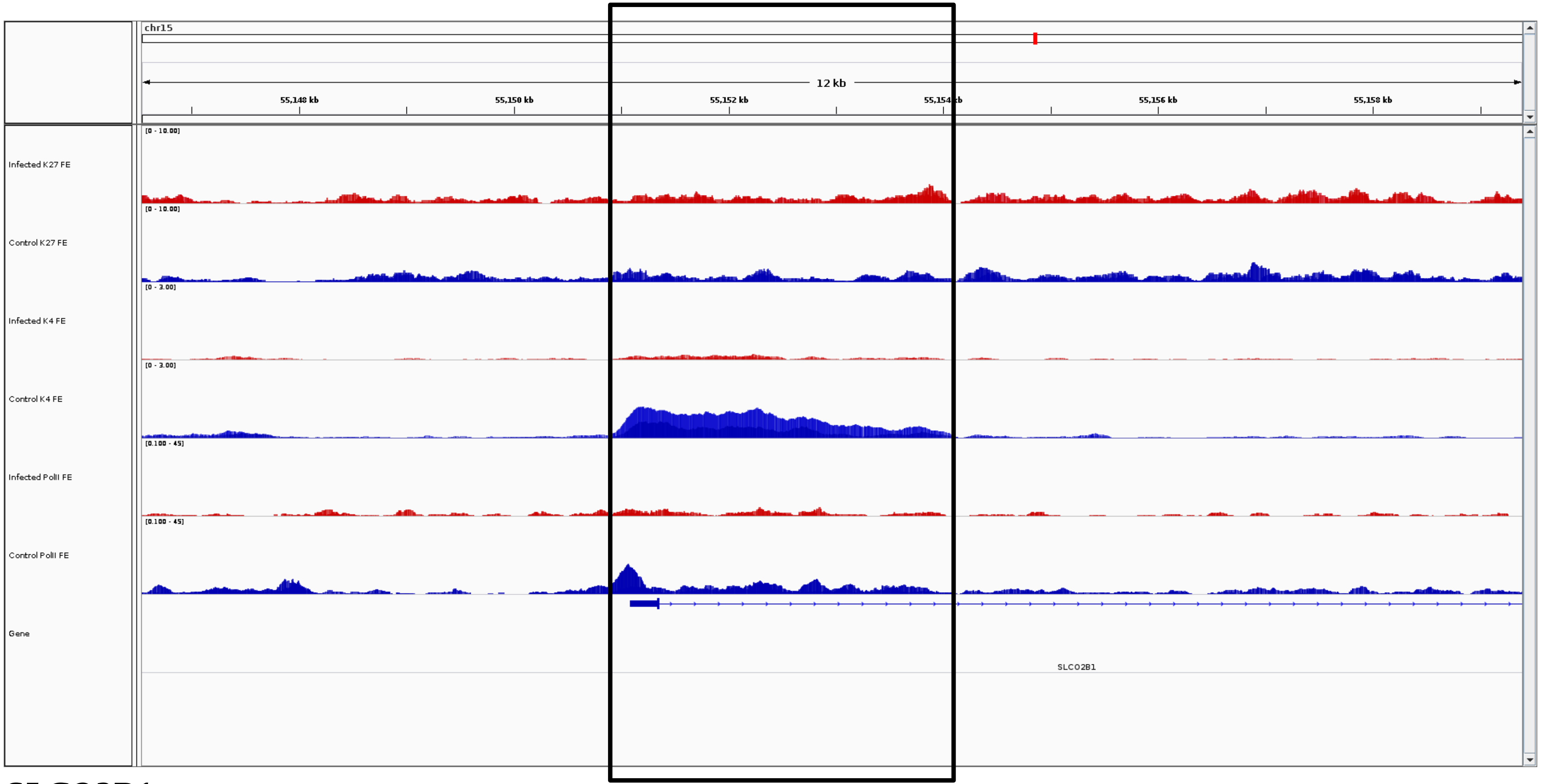
chr15
12 kb
55,148 kb
55,150 kb
55,152 kb
55,154 kb
55,156 kb
55,158 kb
Infected K27 FE
[0 - 10.00]
Control K27 FE
[0 - 10.00]
Infected K4 FE
[0 - 3.00]
Control K4 FE
[0 - 3.00]
Infected PolII FE
[0.100 - 45]
Control PolII FE
[0.100 - 45]
Gene
SLCO2B1

***SLCO2B1***

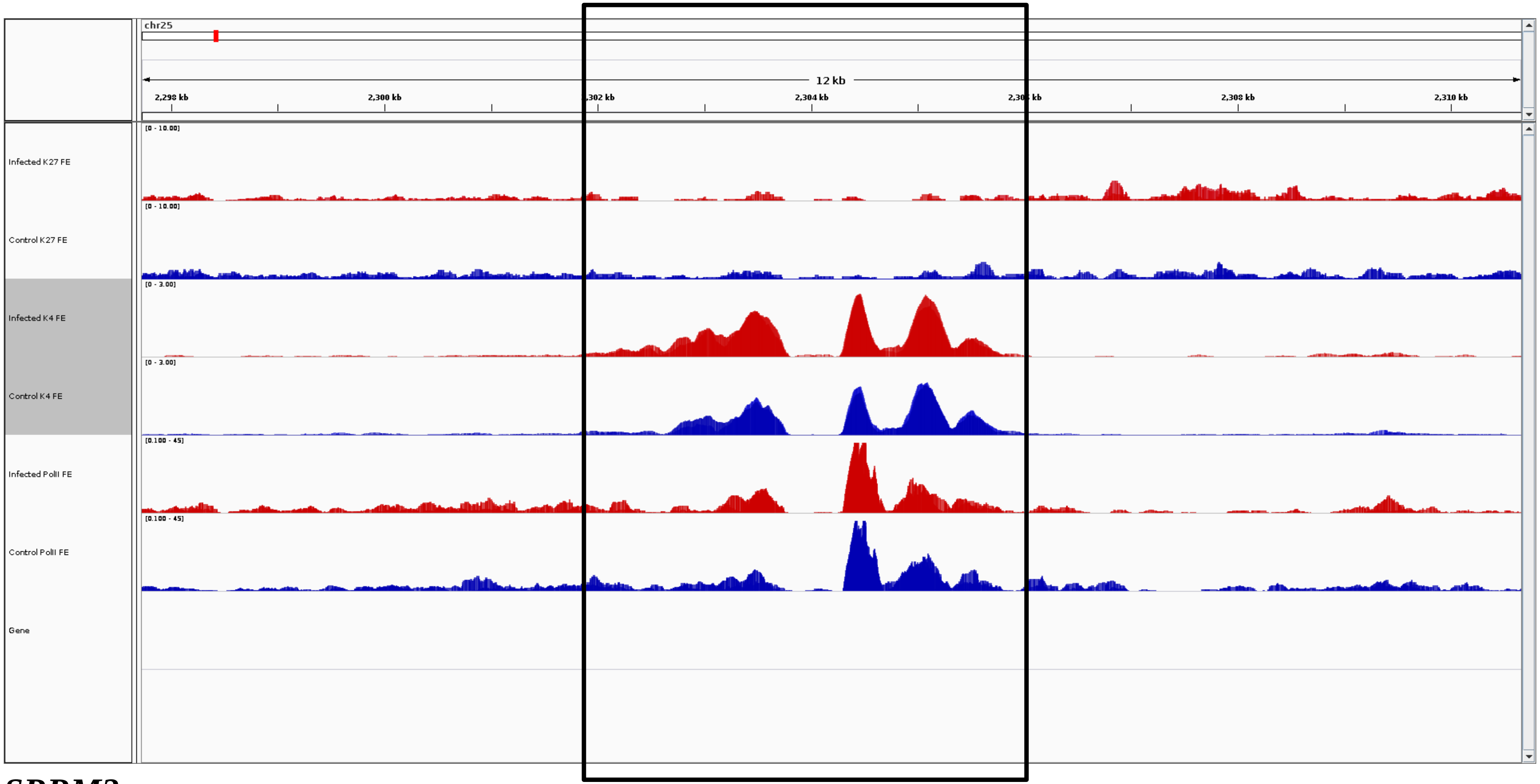
chr25
12 kb
2,298 kb
2,300 kb
2,302 kb
2,304 kb
2,306 kb
2,308 kb
2,310 kb
Infected K27 FE
[0 - 10.00]
Control K27 FE
[0 - 10.00]
Infected K4 FE
[0 - 3.00]
Control K4 FE
[0 - 3.00]
Infected PolII FE
[0.100 - 45]
Control PolII FE
[0.100 - 45]
Gene

***SRRM2***

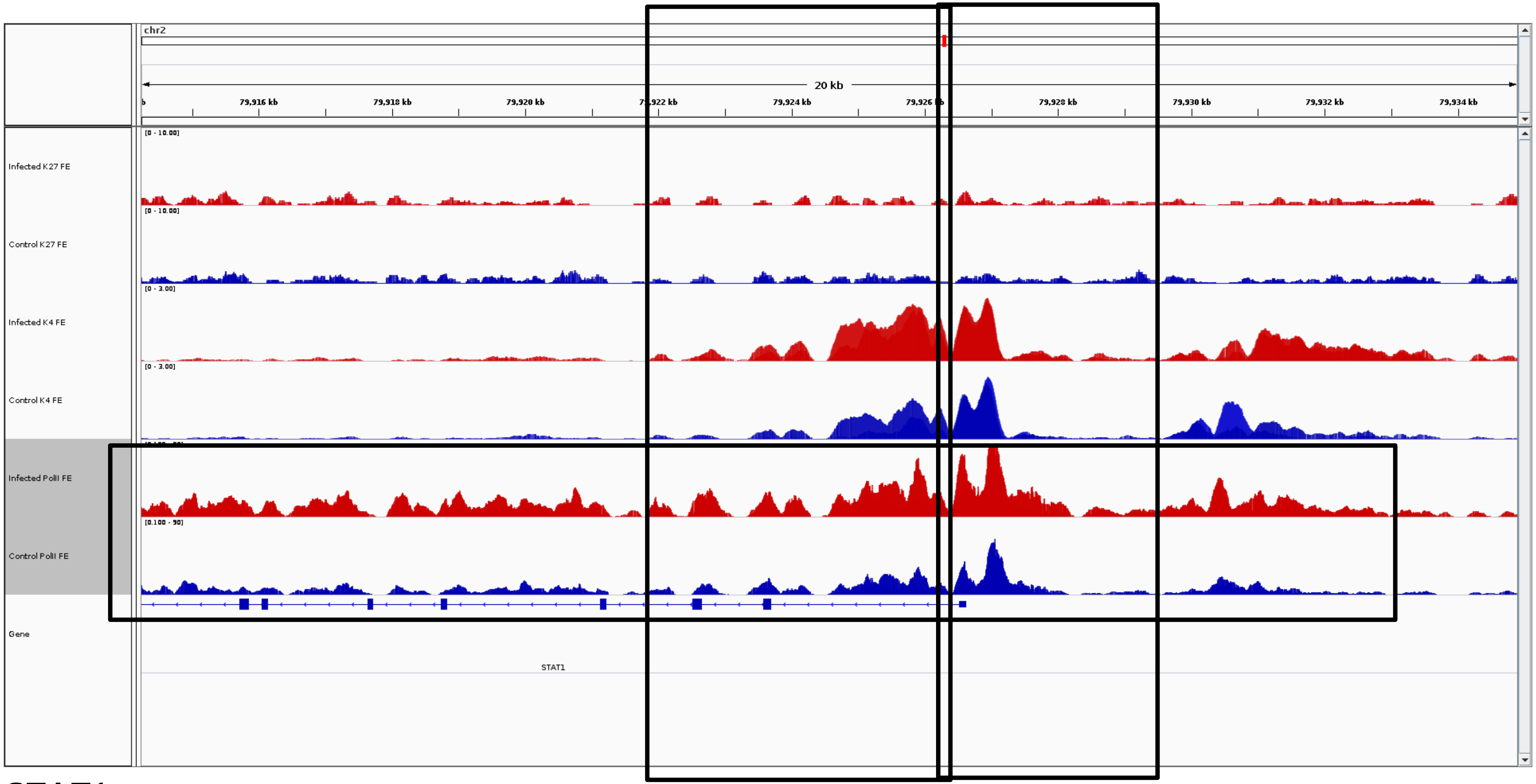
chr2
20 kb
79,916 kb
79,918 kb
79,920 kb
79,924 kb
79,928 kb
79,930 kb
79,932 kb
79,934 kb
Infected K27 FE
Control K27 FE
Infected K4 FE
Control K4 FE
Infected PolII FE
Control PolII FE
Gene
STAT1

***STAT1***

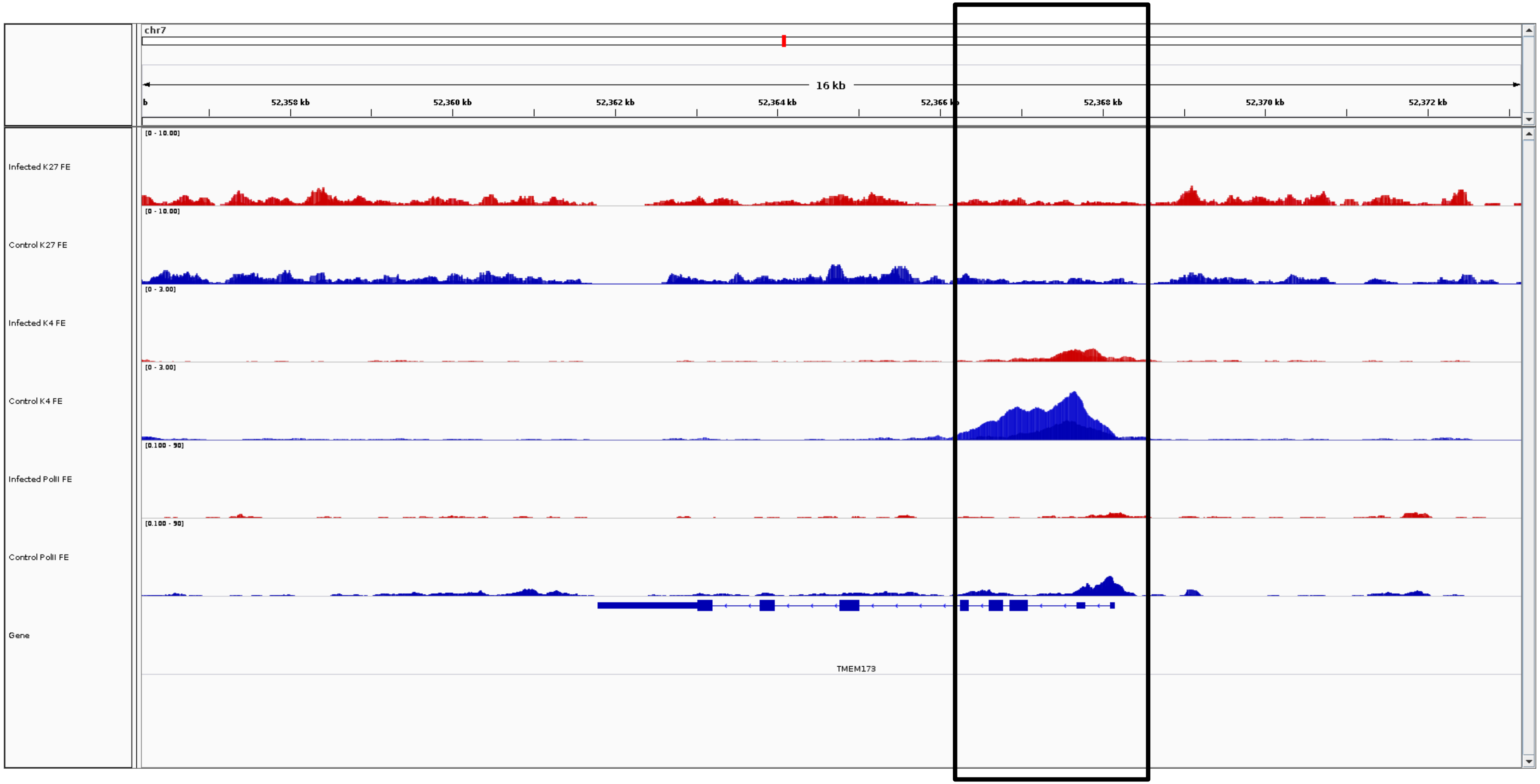
chr7
16 kb
52,358 kb
52,360 kb
52,362 kb
52,364 kb
52,366 kb
52,368 kb
52,370 kb
52,372 kb
Infected K27 FE
[0 - 10.00]
Control K27 FE
[0 - 10.00]
Infected K4 FE
[0 - 3.00]
Control K4 FE
[0 - 3.00]
Infected PolII FE
[0.100 - 90]
Control PolII FE
[0.100 - 90]
Gene
TMEM173

***STING***

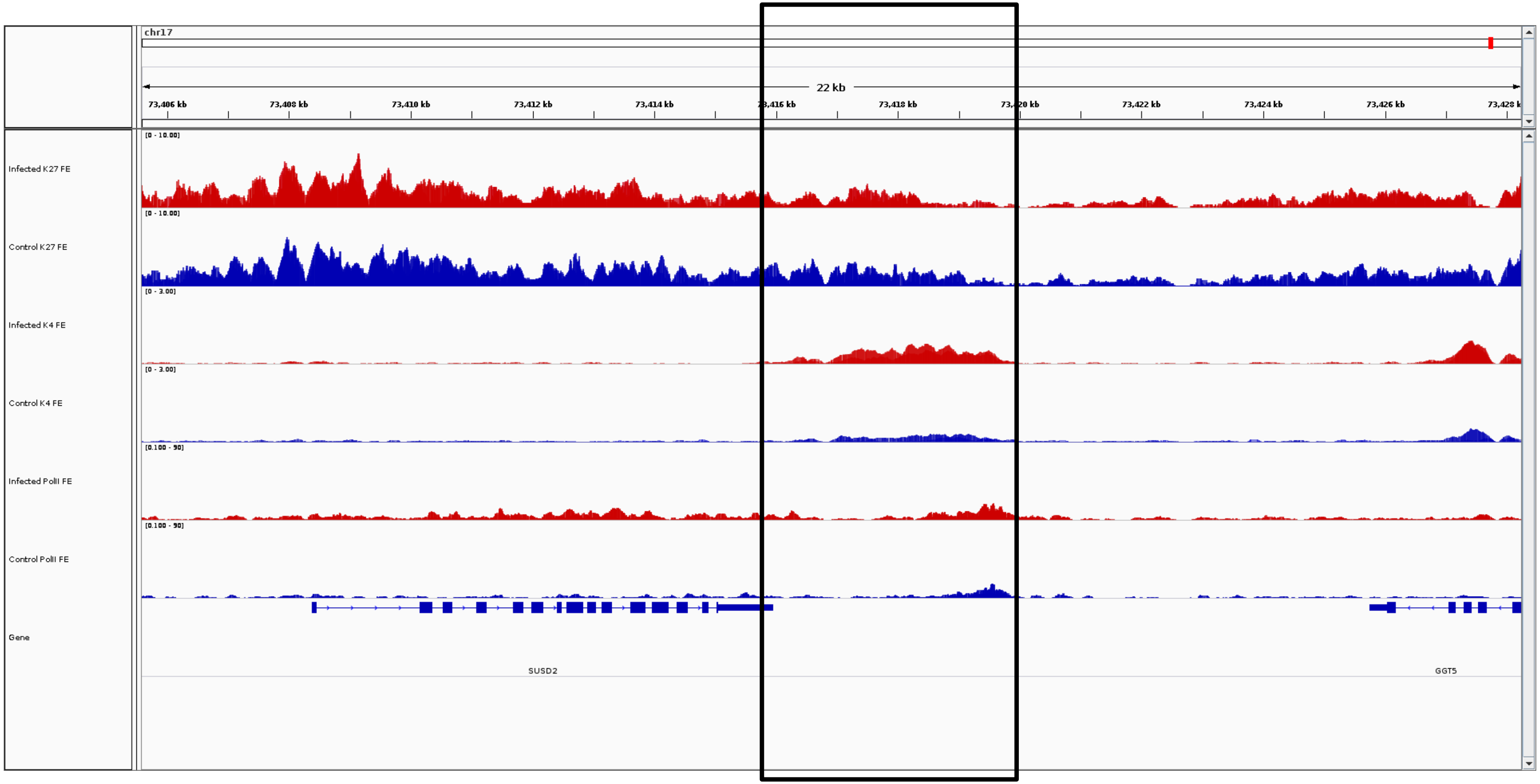
chr17
22 kb
73,406 kb
73,408 kb
73,410 kb
73,412 kb
73,414 kb
73,418 kb
73,422 kb
73,424 kb
73,426 kb
Infected K27 FE
Control K27 FE
Infected K4 FE
Control K4 FE
Infected PolII FE
Control PolII FE
Gene
SUSD2
GGT5

***SUSD2***

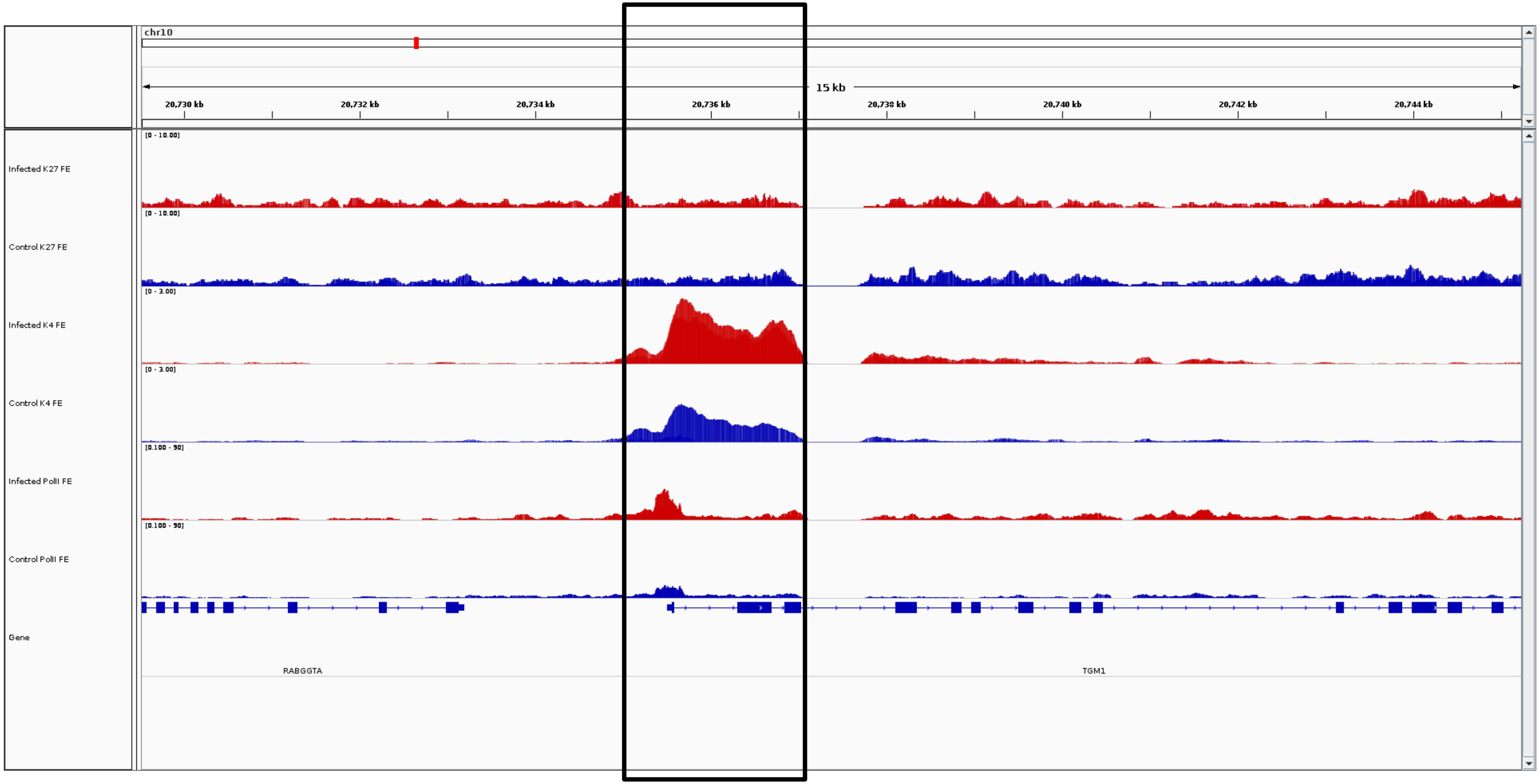
chr10
15 kb
20,730 kb
20,732 kb
20,734 kb
20,736 kb
20,738 kb
20,740 kb
20,742 kb
20,744 kb
Infected K27 FE
[0 - 10.00]
Control K27 FE
[0 - 10.00]
Infected K4 FE
[0 - 3.00]
Control K4 FE
[0 - 3.00]
Infected PolII FE
[0.100 - 90]
Control PolII FE
[0.100 - 90]
Gene
RABGGTA
TGM1

***TGM1***

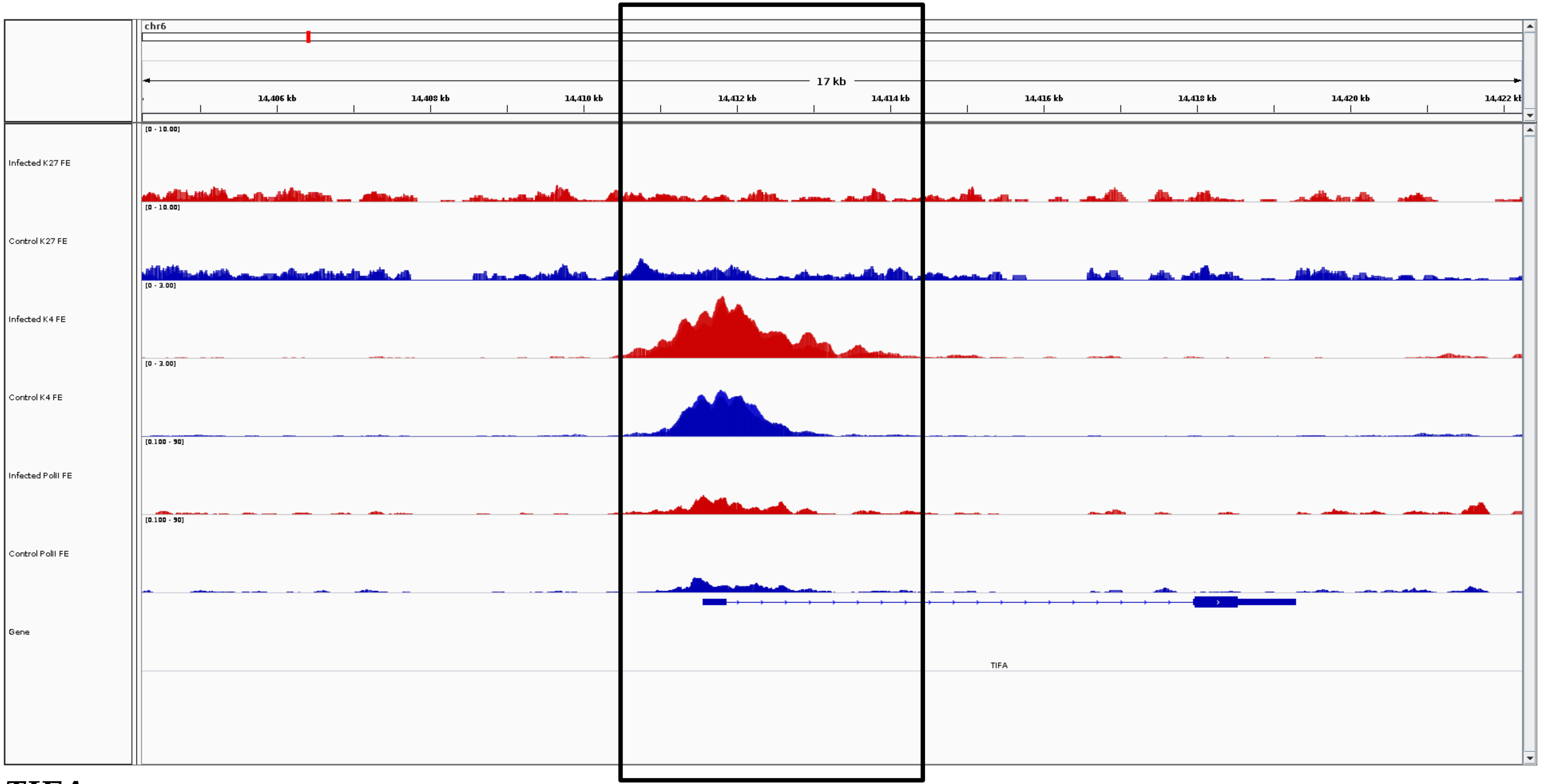
chr6
17 kb
14,406 kb
14,408 kb
14,410 kb
14,412 kb
14,414 kb
14,416 kb
14,418 kb
14,420 kb
Infected K27 FE
Control K27 FE
Infected K4 FE
Control K4 FE
Infected PolII FE
Control PolII FE
Gene
TIFA

***TIFA***

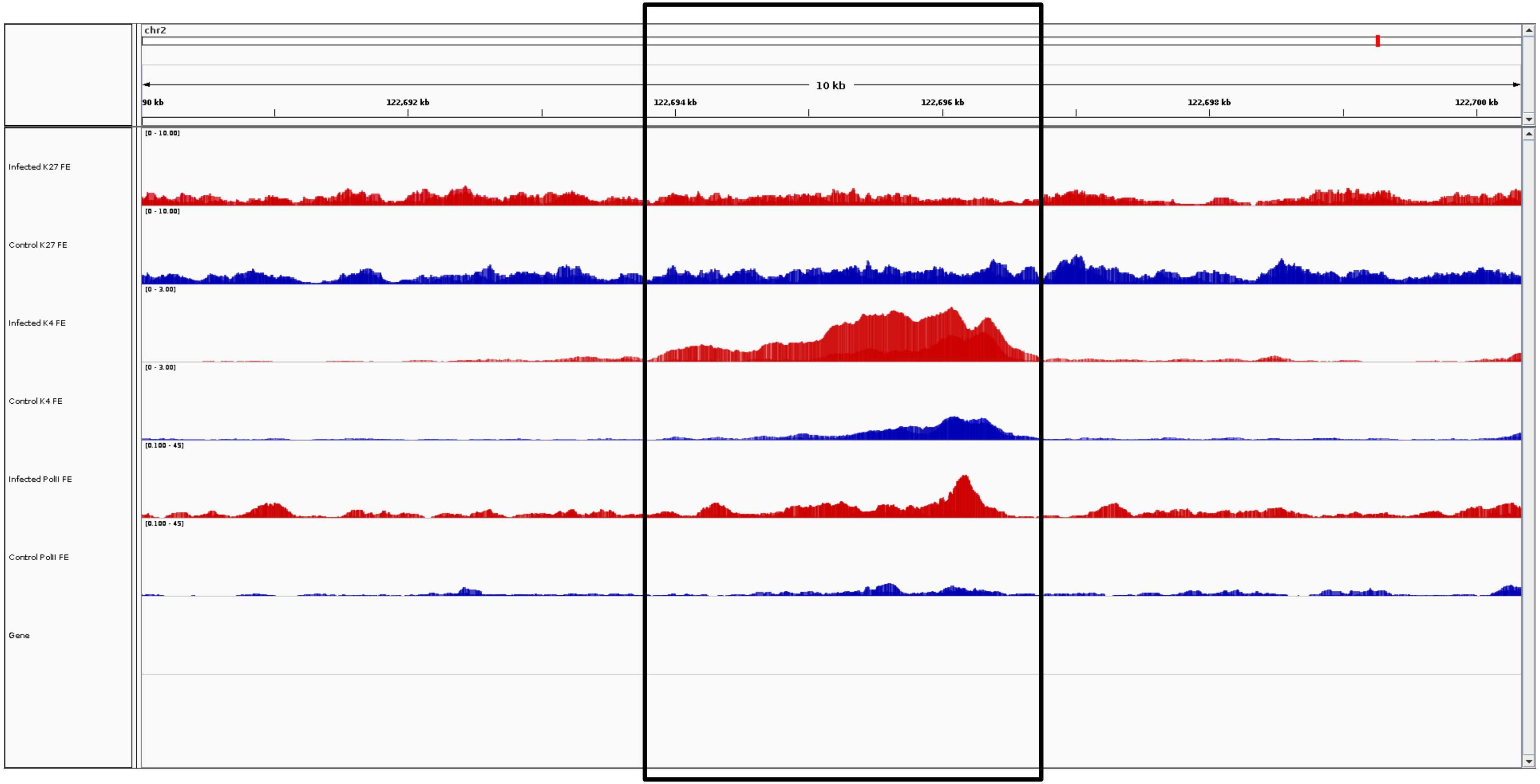
chr2
10 kb
90 kb
122,692 kb
122,694 kb
122,696 kb
122,698 kb
122,700 kb
Infected K27 FE
[0 - 10.00]
Control K27 FE
[0 - 10.00]
Infected K4 FE
[0 - 3.00]
Control K4 FE
[0 - 3.00]
Infected PolII FE
[0.100 - 45]
Control PolII FE
[0.100 - 45]
Gene

***TINAGL1***

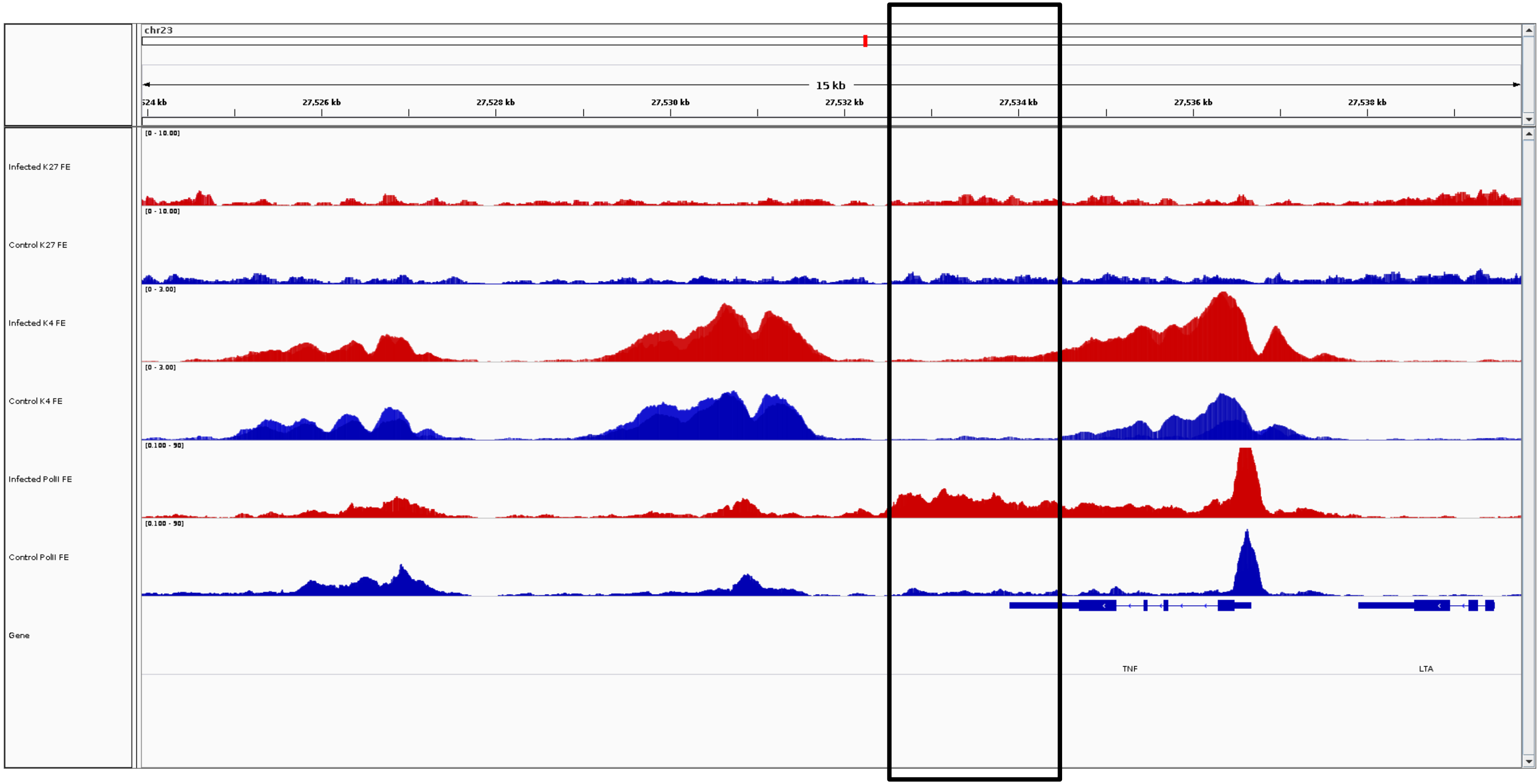
chr23
15 kb
27,524 kb
27,526 kb
27,528 kb
27,530 kb
27,532 kb
27,534 kb
27,536 kb
27,538 kb
Infected K27 FE
[0 - 10.00]
Control K27 FE
[0 - 10.00]
Infected K4 FE
[0 - 3.00]
Control K4 FE
[0 - 3.00]
Infected PolII FE
[0.100 - 50]
Control PolII FE
[0.100 - 50]
Gene
TNF
LTA

***TNF***

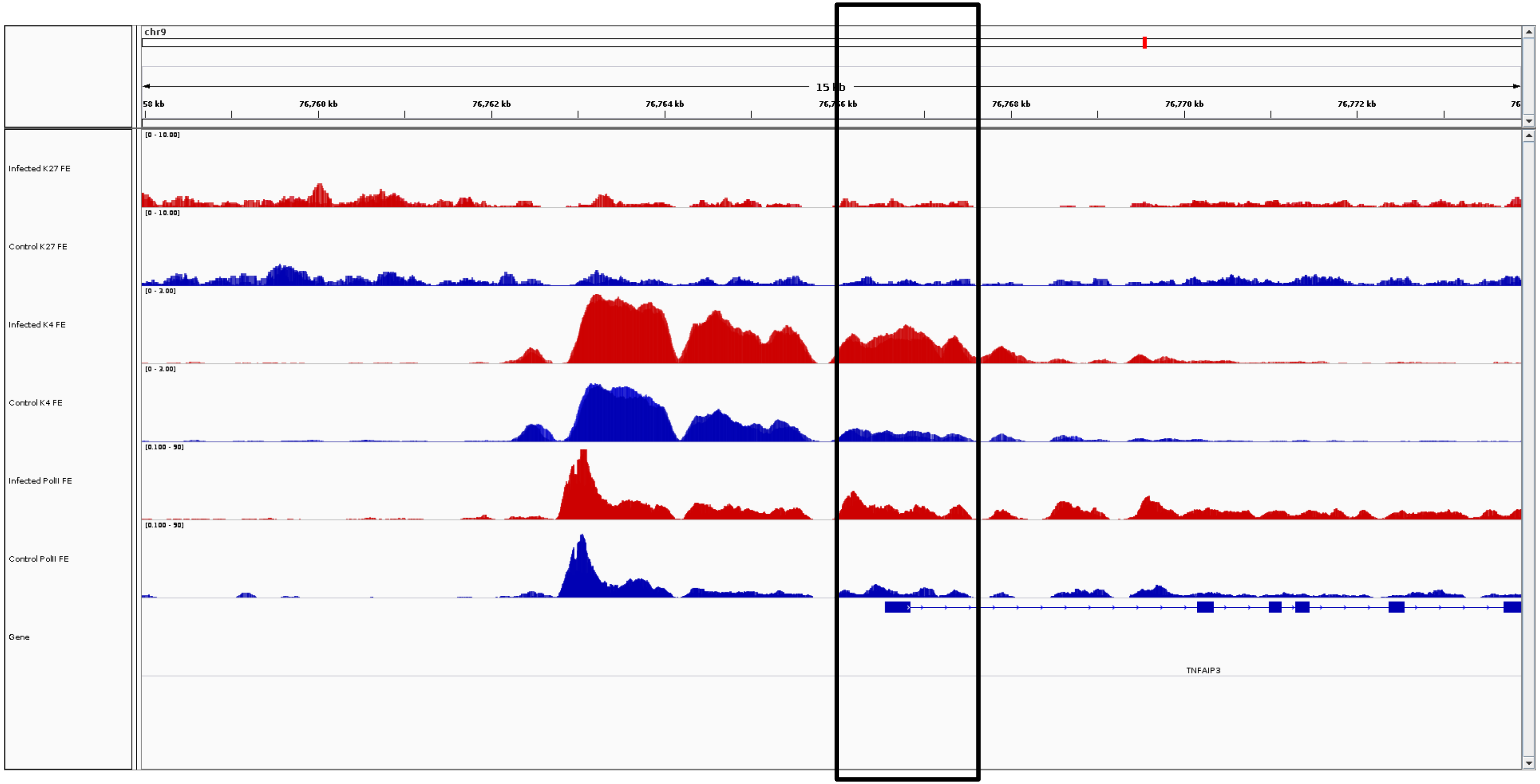
chr9
15 kb
76,760 kb
76,762 kb
76,764 kb
76,768 kb
76,770 kb
76,772 kb
Infected K27 FE
Control K27 FE
Infected K4 FE
Control K4 FE
Infected PolII FE
Control PolII FE
Gene
TNFAIP3

***TNFAIP3***

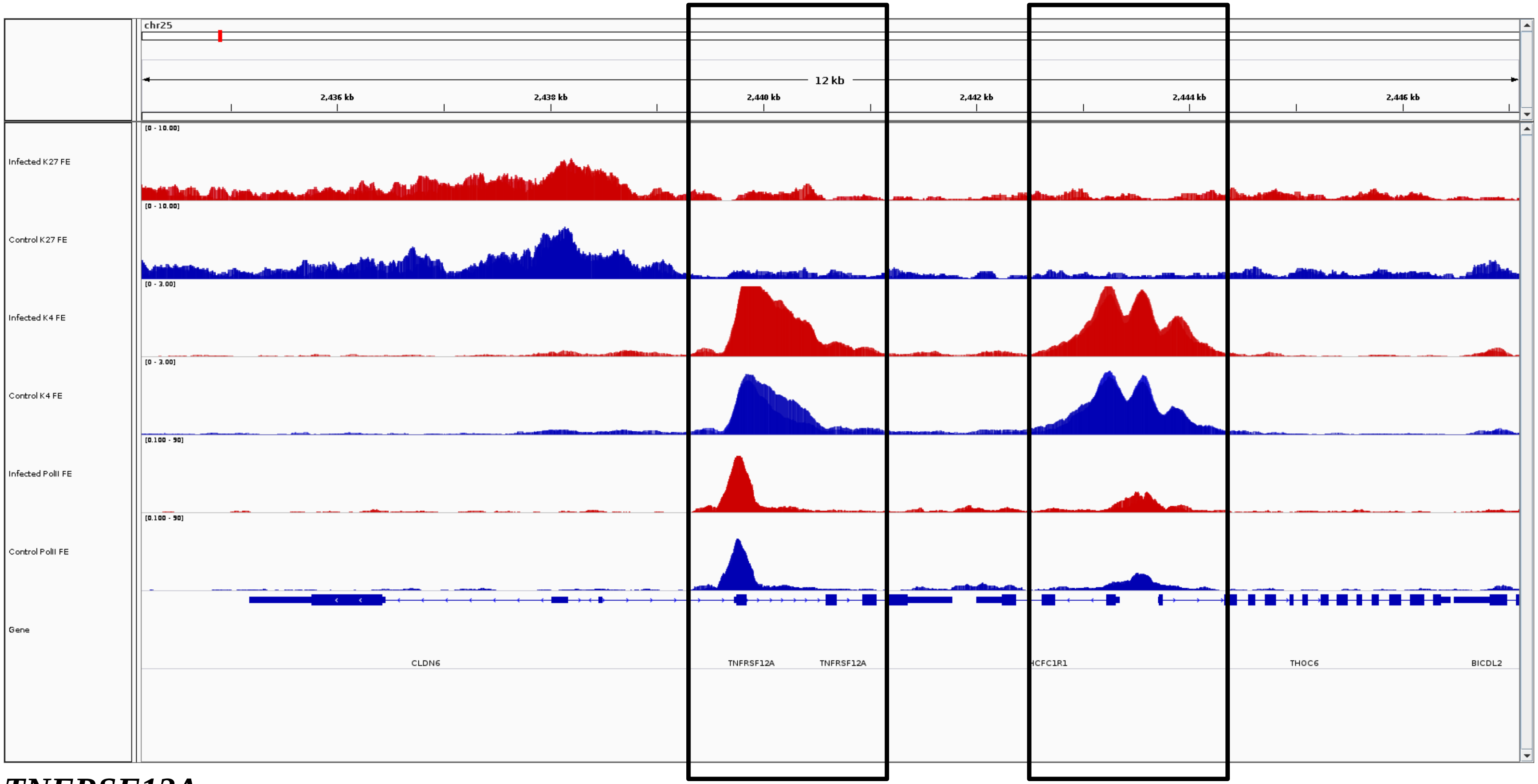
chr25
12 kb
2,436 kb
2,438 kb
2,440 kb
2,442 kb
2,444 kb
2,446 kb
Infected K27 FE
[0 - 10.00]
Control K27 FE
[0 - 10.00]
Infected K4 FE
[0 - 3.00]
Control K4 FE
[0 - 3.00]
Infected PolII FE
[0.100 - 90]
Control PolII FE
[0.100 - 90]
Gene
CLDN6
TNFRSF12A
TNFRSF12A
HCFC1R1
THOC6
BICDL2

***TNFRSF12A***

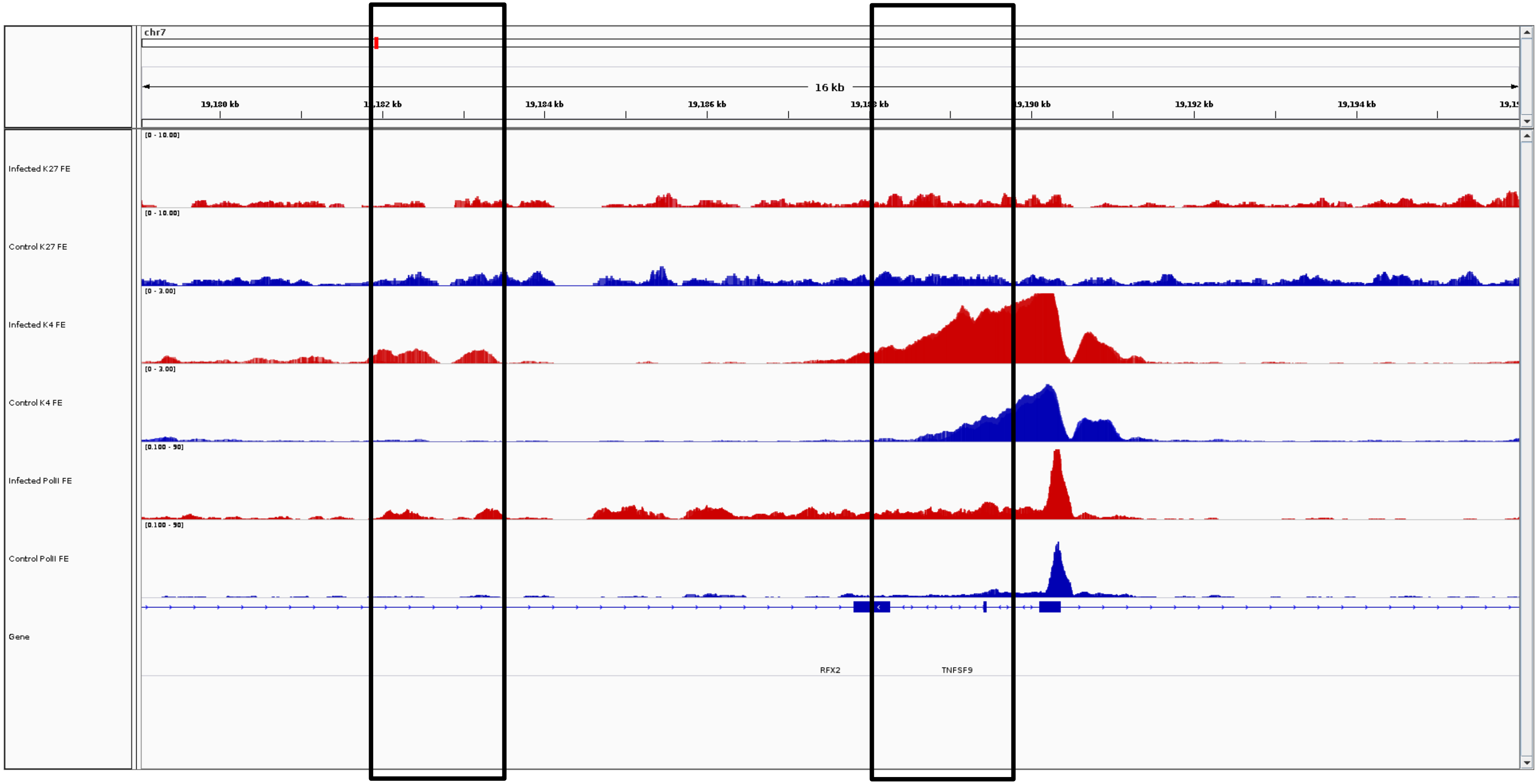
chr7
16 kb
19,180 kb
19,184 kb
19,186 kb
19,190 kb
19,192 kb
19,194 kb
[0 - 10.00]
Infected K27 FE
Control K27 FE
[0 - 3.00]
Infected K4 FE
Control K4 FE
[0.100 - 90]
Infected PolII FE
Control PolII FE
Gene
RFX2
TNFSF9

***TNFSF9***

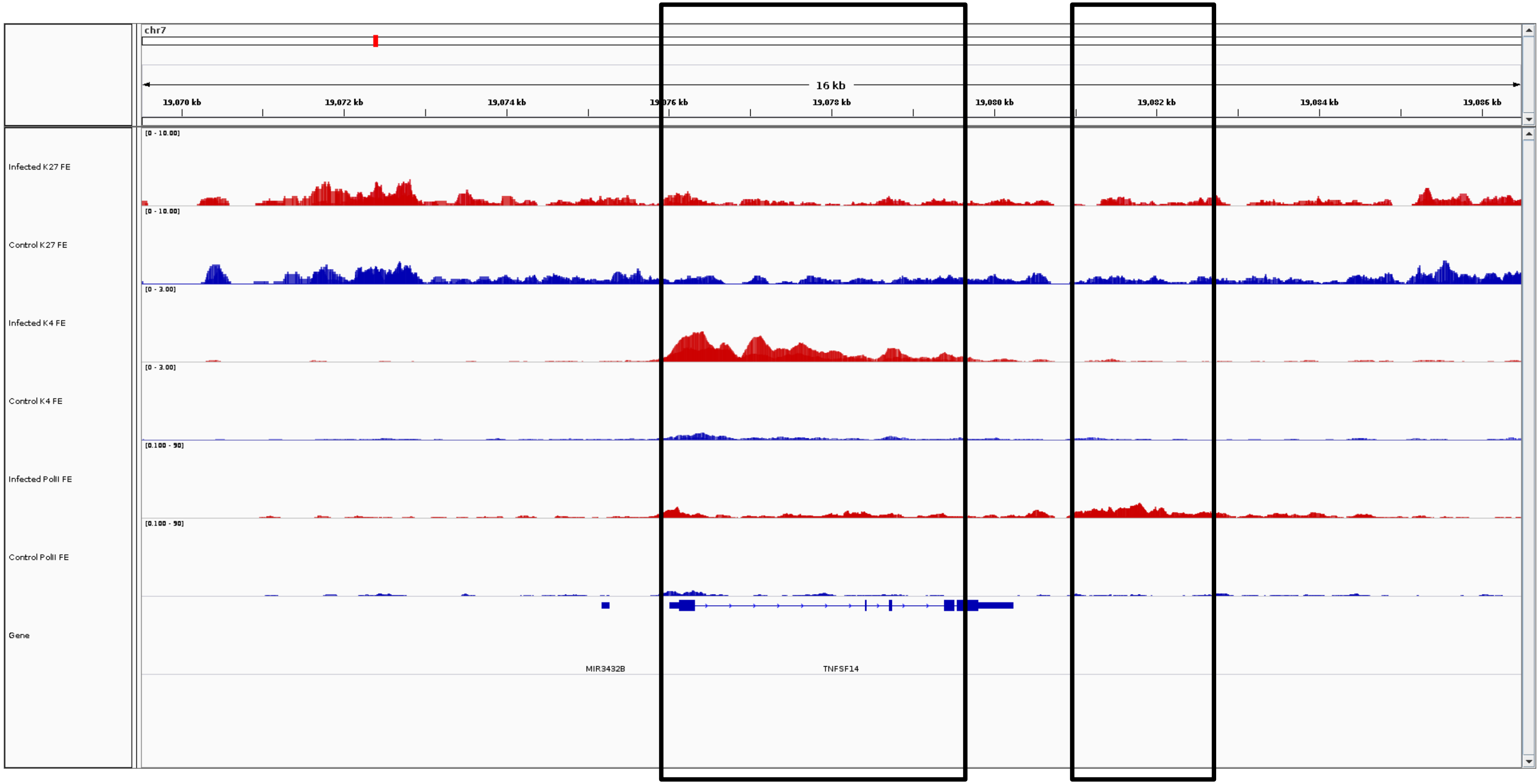
chr7
16 kb
19,070 kb
19,072 kb
19,074 kb
19,076 kb
19,078 kb
19,080 kb
19,082 kb
19,084 kb
19,086 kb
Infected K27 FE
[0 - 10.00]
Control K27 FE
[0 - 10.00]
Infected K4 FE
[0 - 3.00]
Control K4 FE
[0 - 3.00]
Infected PolII FE
[0.100 - 50]
Control PolII FE
[0.100 - 50]
Gene
MIR3432B
TNFSF14

***TNFSF14***

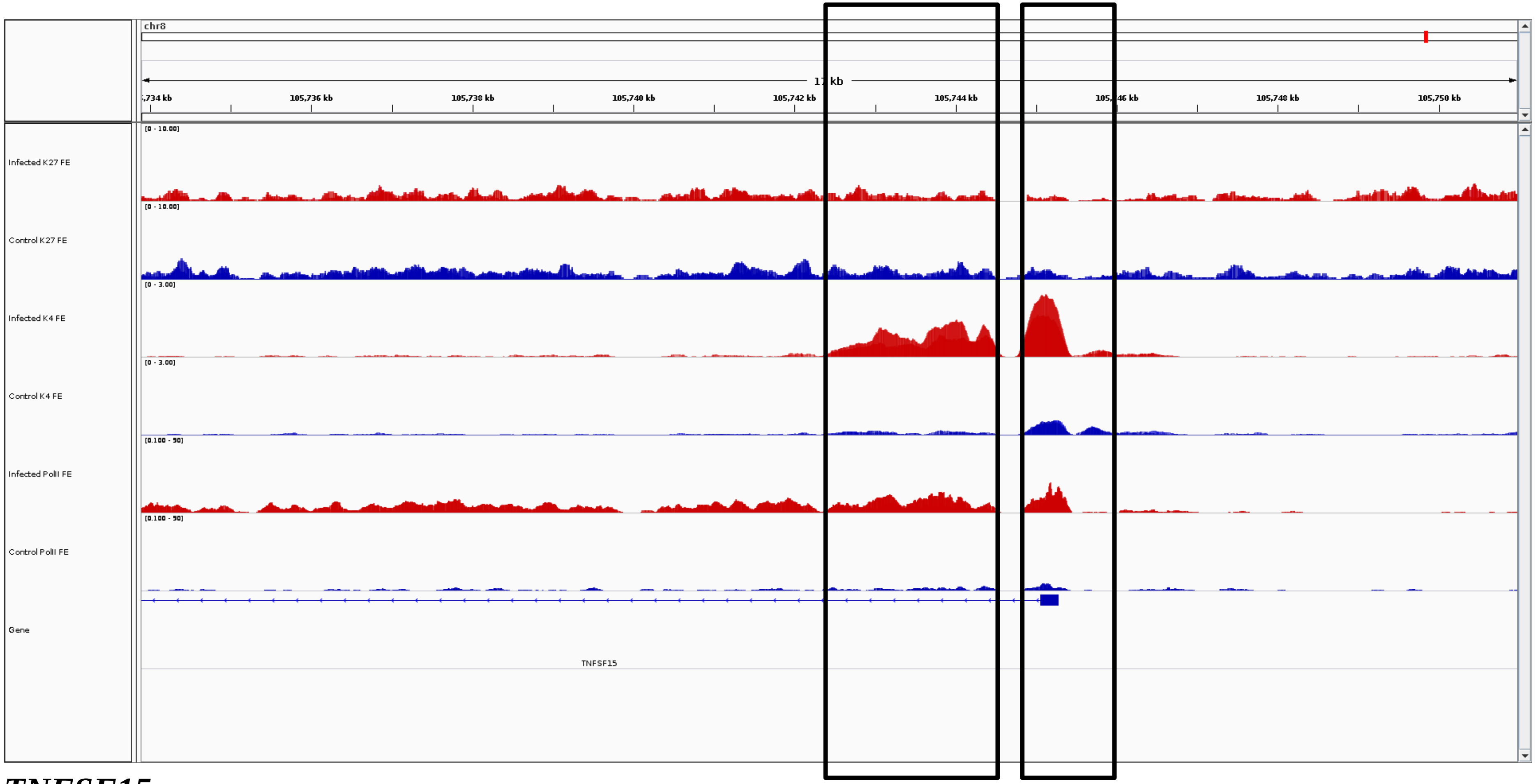
chr8
17 kb
,734 kb
105,736 kb
105,738 kb
105,740 kb
105,742 kb
105,744 kb
105,746 kb
105,748 kb
105,750 kb
[0 - 10.00]
Infected K27 FE
[0 - 10.00]
Control K27 FE
[0 - 3.00]
Infected K4 FE
[0 - 3.00]
Control K4 FE
[0.100 - 50]
Infected PolII FE
[0.100 - 50]
Control PolII FE
Gene
TNFSF15

***TNFSF15***

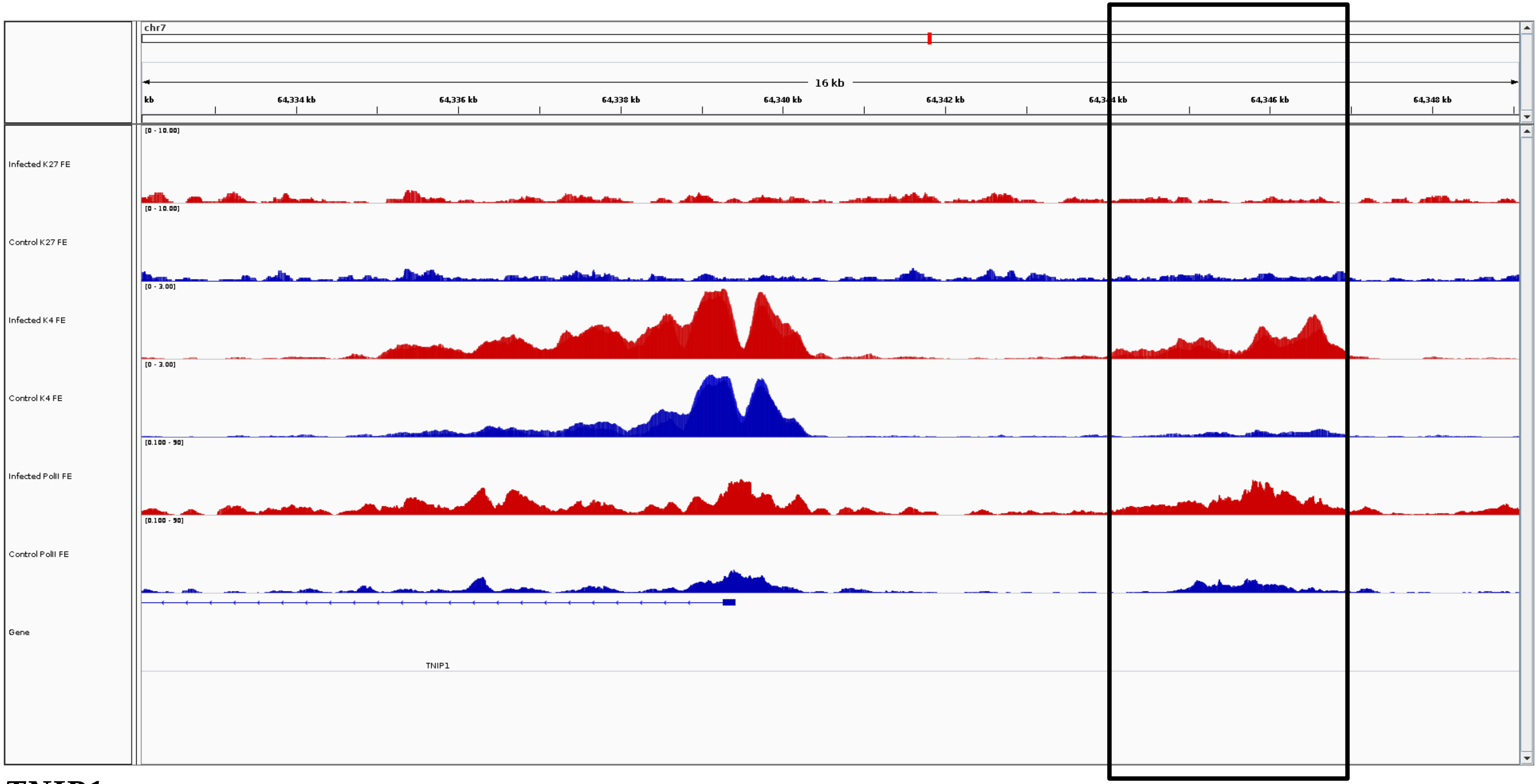
chr7
16 kb
kb
64,334 kb
64,336 kb
64,338 kb
64,340 kb
64,342 kb
64,346 kb
64,348 kb
Infected K27 FE
Control K27 FE
Infected K4 FE
Control K4 FE
Infected PolII FE
Control PolII FE
Gene
TNIP1

***TNIP1***

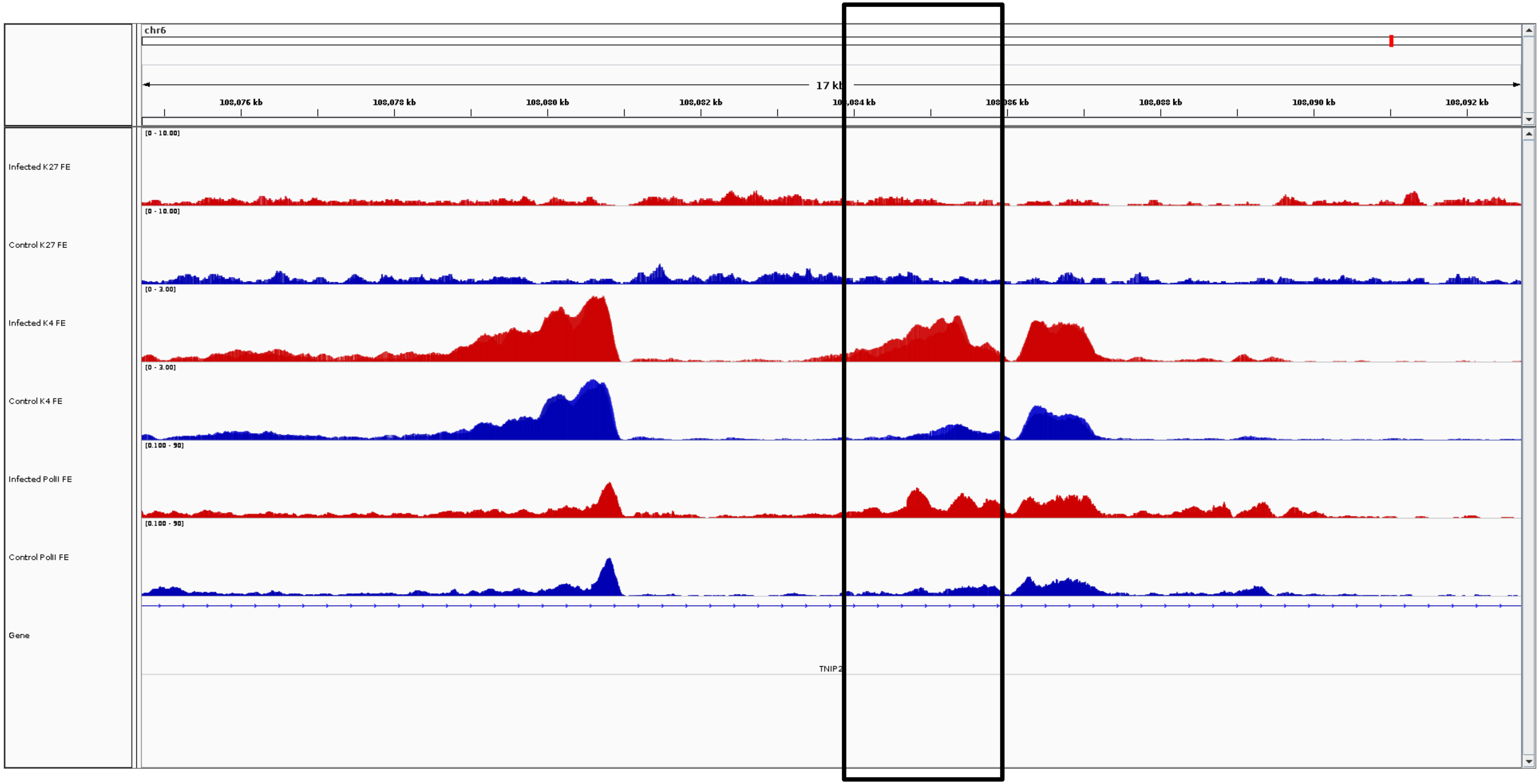
chr6
17 kb
108,076 kb
108,078 kb
108,080 kb
108,082 kb
108,086 kb
108,088 kb
108,090 kb
108,092 kb
Infected K27 FE
[0 - 10.00]
Control K27 FE
[0 - 10.00]
Infected K4 FE
[0 - 3.00]
Control K4 FE
[0 - 3.00]
Infected PolII FE
[0.100 - 50]
Control PolII FE
[0.100 - 50]
Gene
TNIP2

***TNIP2***

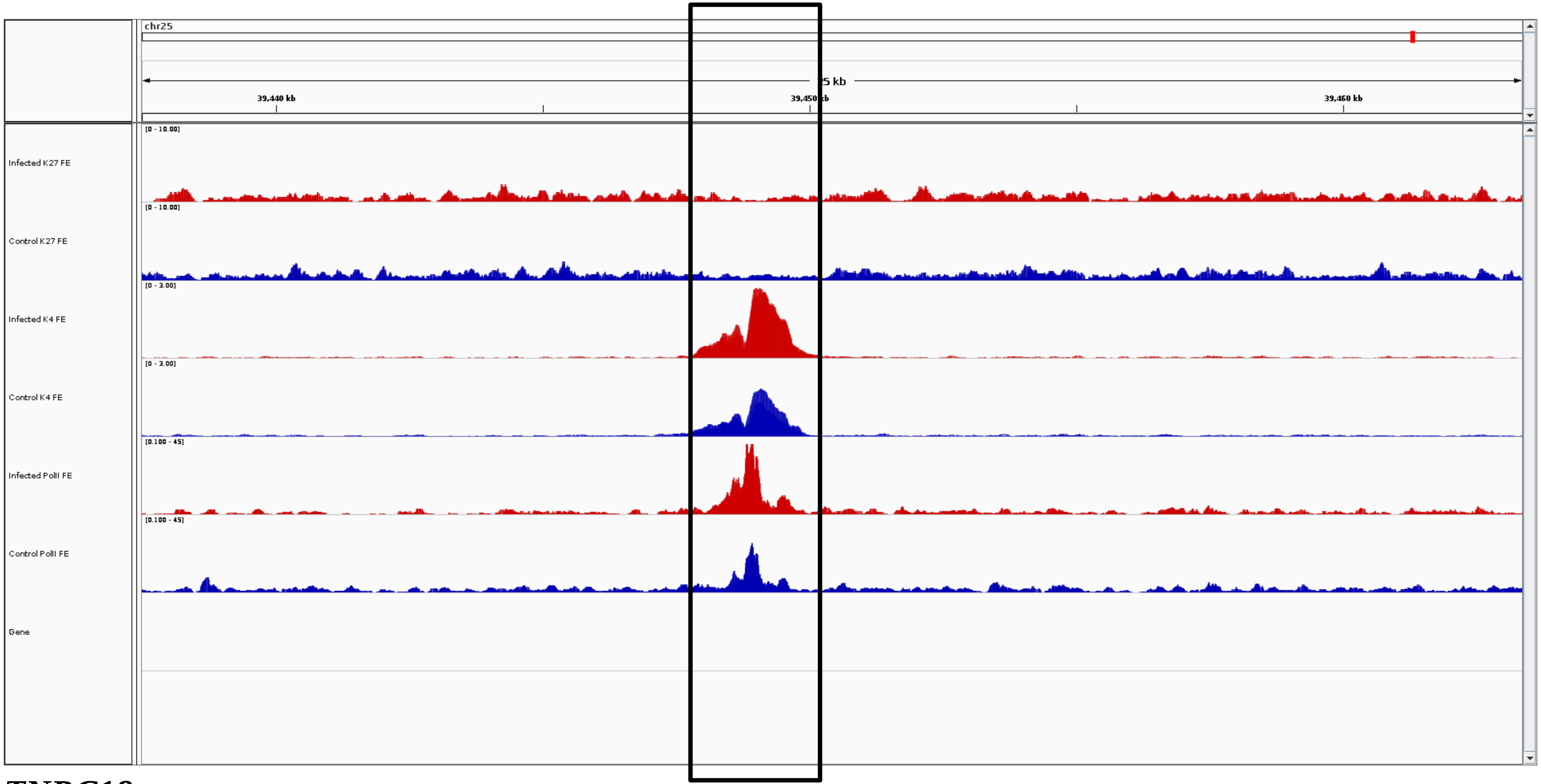
chr25
25 kb
39,440 kb
39,450 kb
39,460 kb
Infected K27 FE
[0 - 10.00]
Control K27 FE
[0 - 10.00]
Infected K4 FE
[0 - 3.00]
Control K4 FE
[0 - 3.00]
Infected PolII FE
[0.100 - 45]
Control PolII FE
[0.100 - 45]
Gene

***TNRC18***

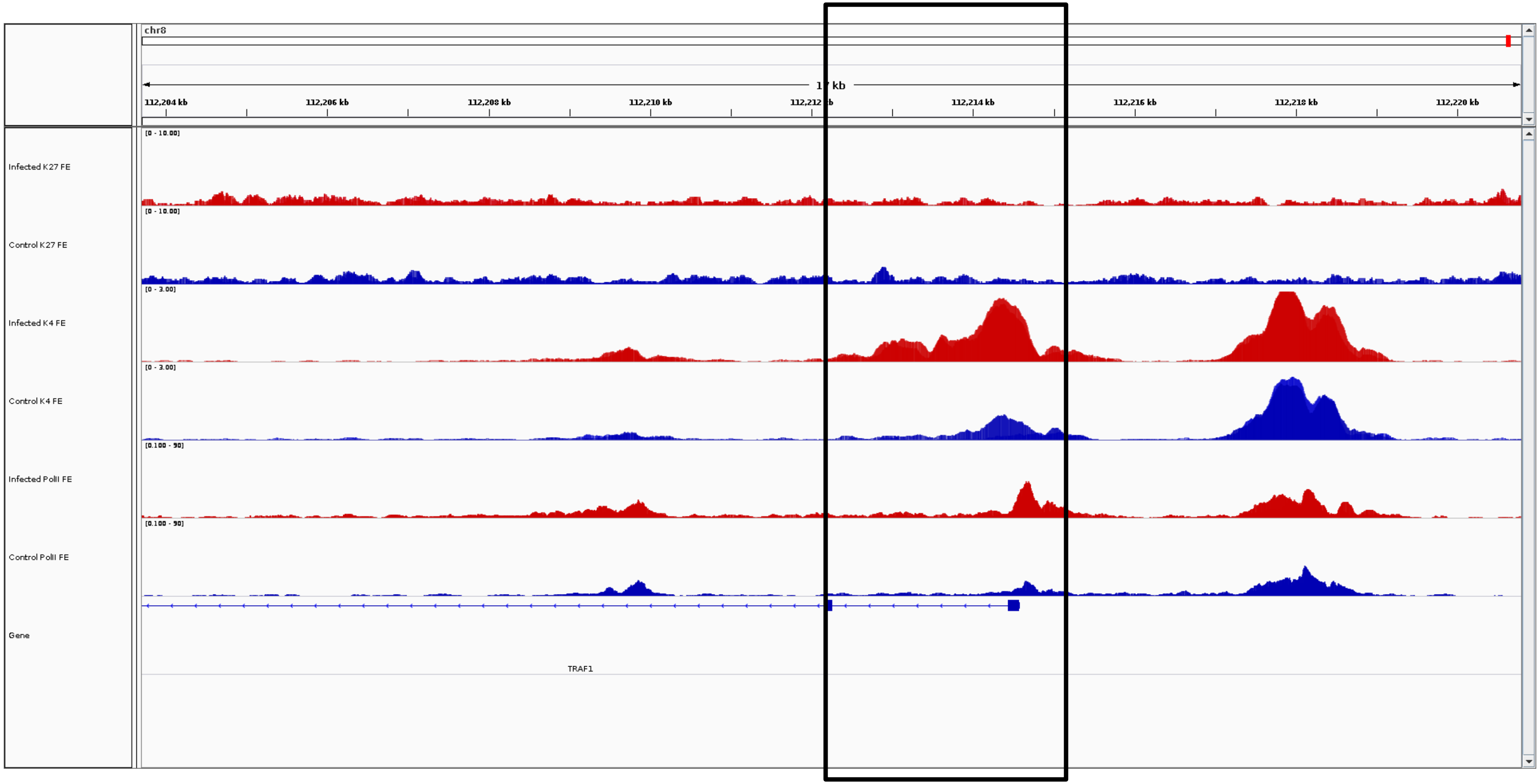
chr8
17 kb
112,204 kb
112,206 kb
112,208 kb
112,210 kb
112,212 kb
112,214 kb
112,216 kb
112,218 kb
112,220 kb
Infected K27 FE
[0 - 10.00]
Control K27 FE
[0 - 10.00]
Infected K4 FE
[0 - 3.00]
Control K4 FE
[0 - 3.00]
Infected PolII FE
[0.100 - 90]
Control PolII FE
[0.100 - 90]
Gene
TRAF1

***TRAF1***

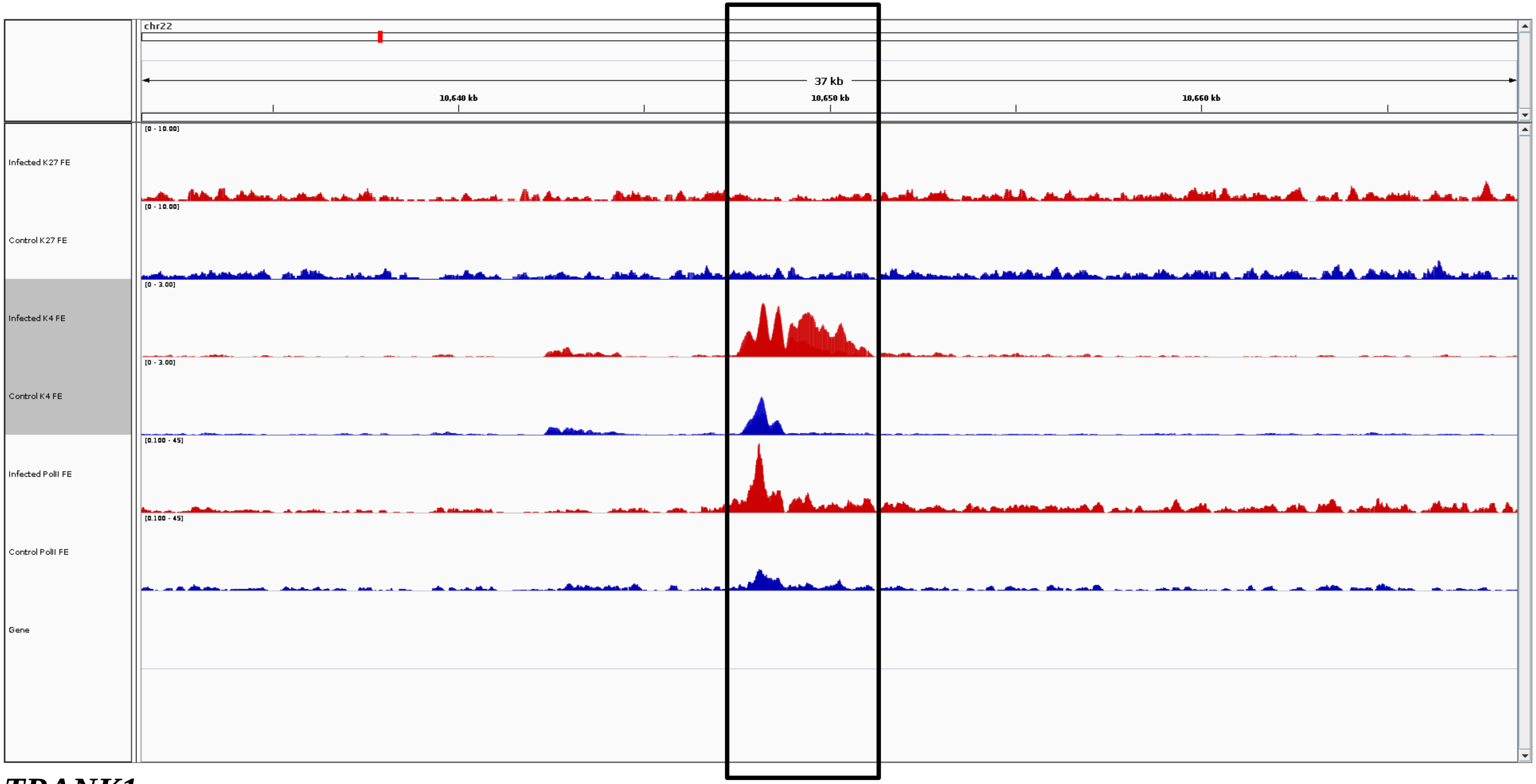
chr22
37 kb
10,640 kb
10,650 kb
10,660 kb
[0 - 10.00]
Infected K27 FE
[0 - 10.00]
Control K27 FE
[0 - 3.00]
Infected K4 FE
[0 - 3.00]
Control K4 FE
[0.100 - 45]
Infected PolII FE
[0.100 - 45]
Control PolII FE
Gene

***TRANK1***

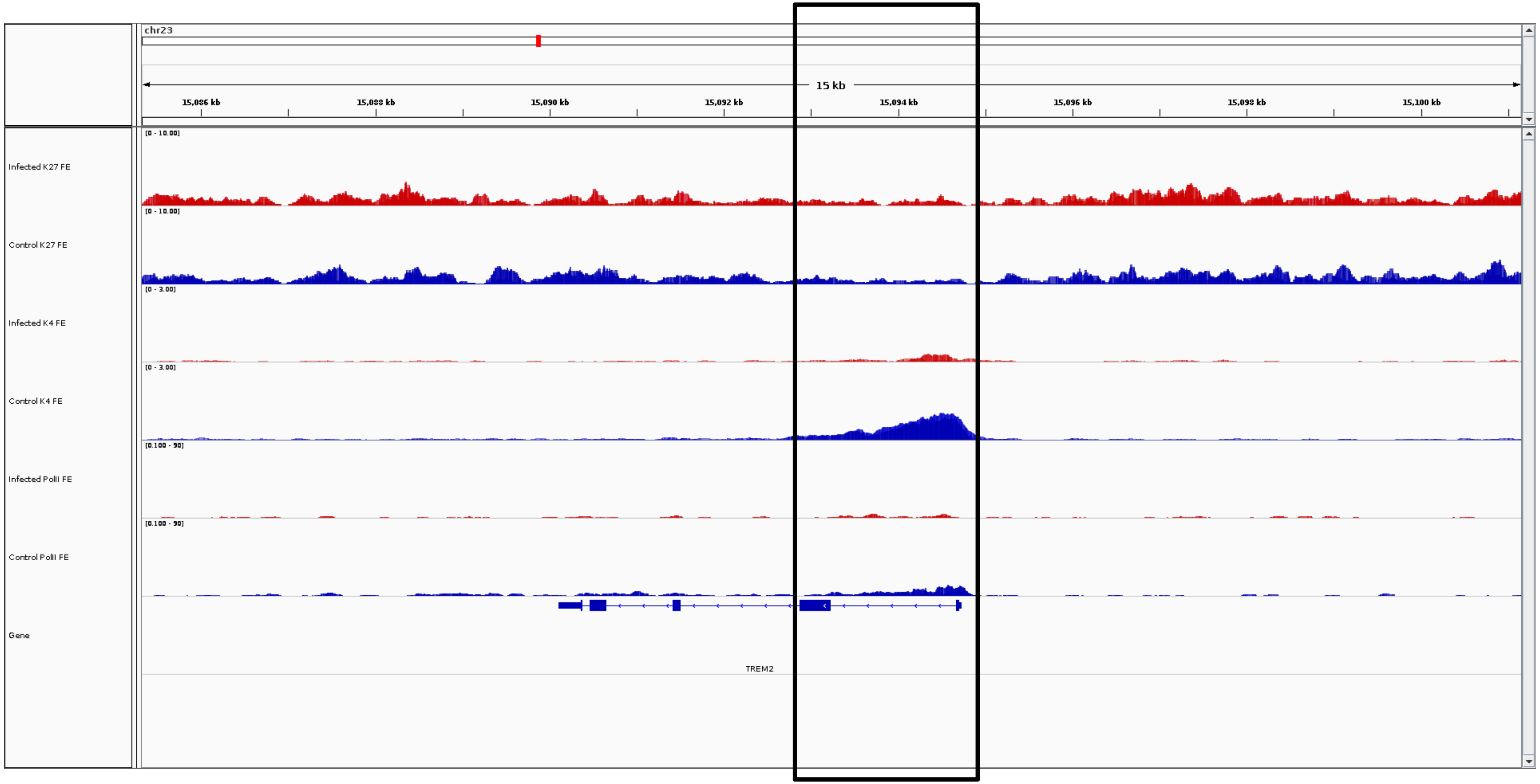
chr23
15 kb
15,086 kb
15,088 kb
15,090 kb
15,092 kb
15,094 kb
15,096 kb
15,098 kb
15,100 kb
Infected K27 FE
[0 - 10.00]
Control K27 FE
[0 - 10.00]
Infected K4 FE
[0 - 3.00]
Control K4 FE
[0 - 3.00]
Infected PolII FE
[0.100 - 50]
Control PolII FE
[0.100 - 50]
Gene
TREM2

***TREM2***

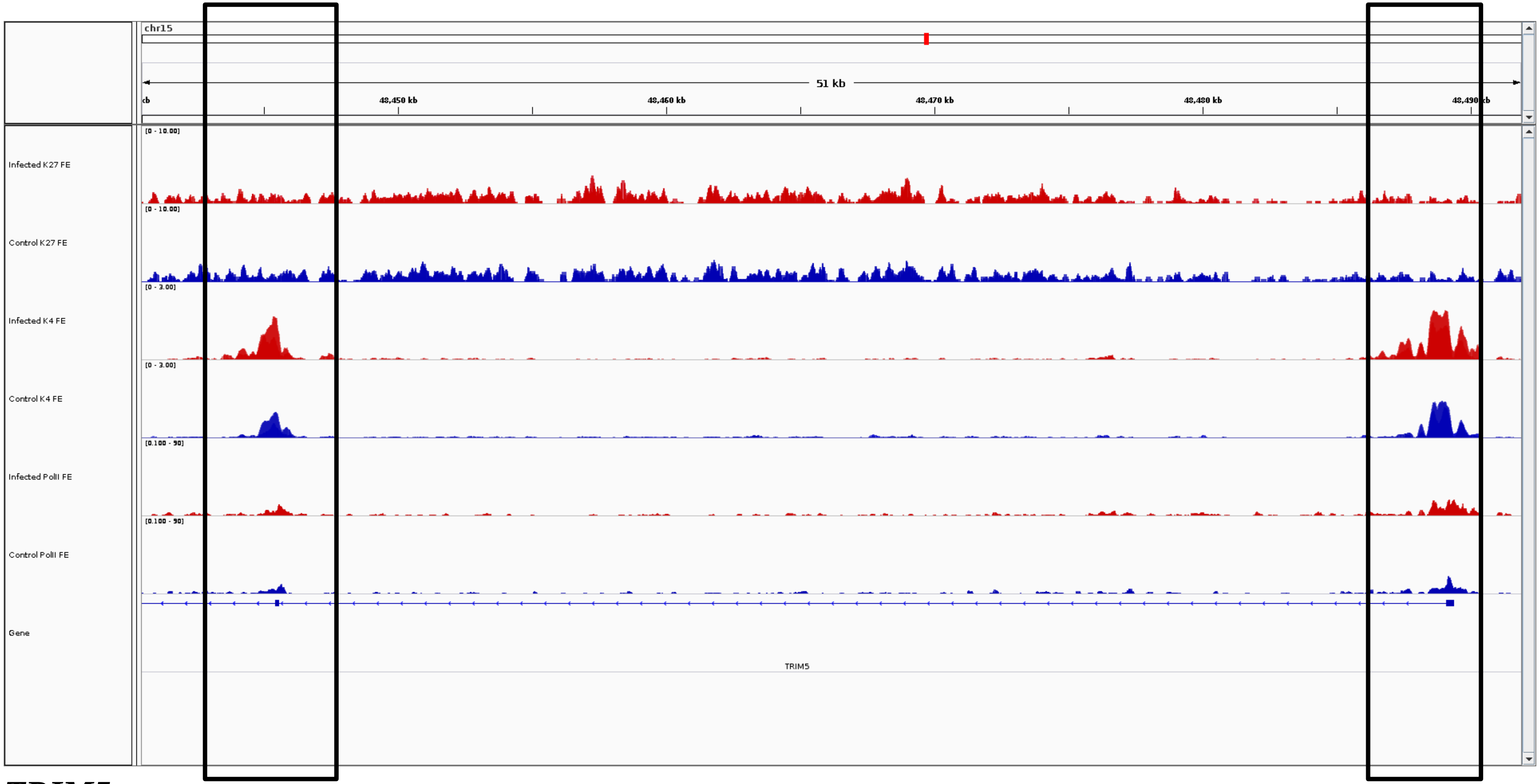
chr15
51 kb
48,450 kb
48,460 kb
48,470 kb
48,480 kb
48,490 kb
Infected K27 FE
Control K27 FE
Infected K4 FE
Control K4 FE
Infected PolII FE
Control PolII FE
Gene
TRIM5

***TRIM5***

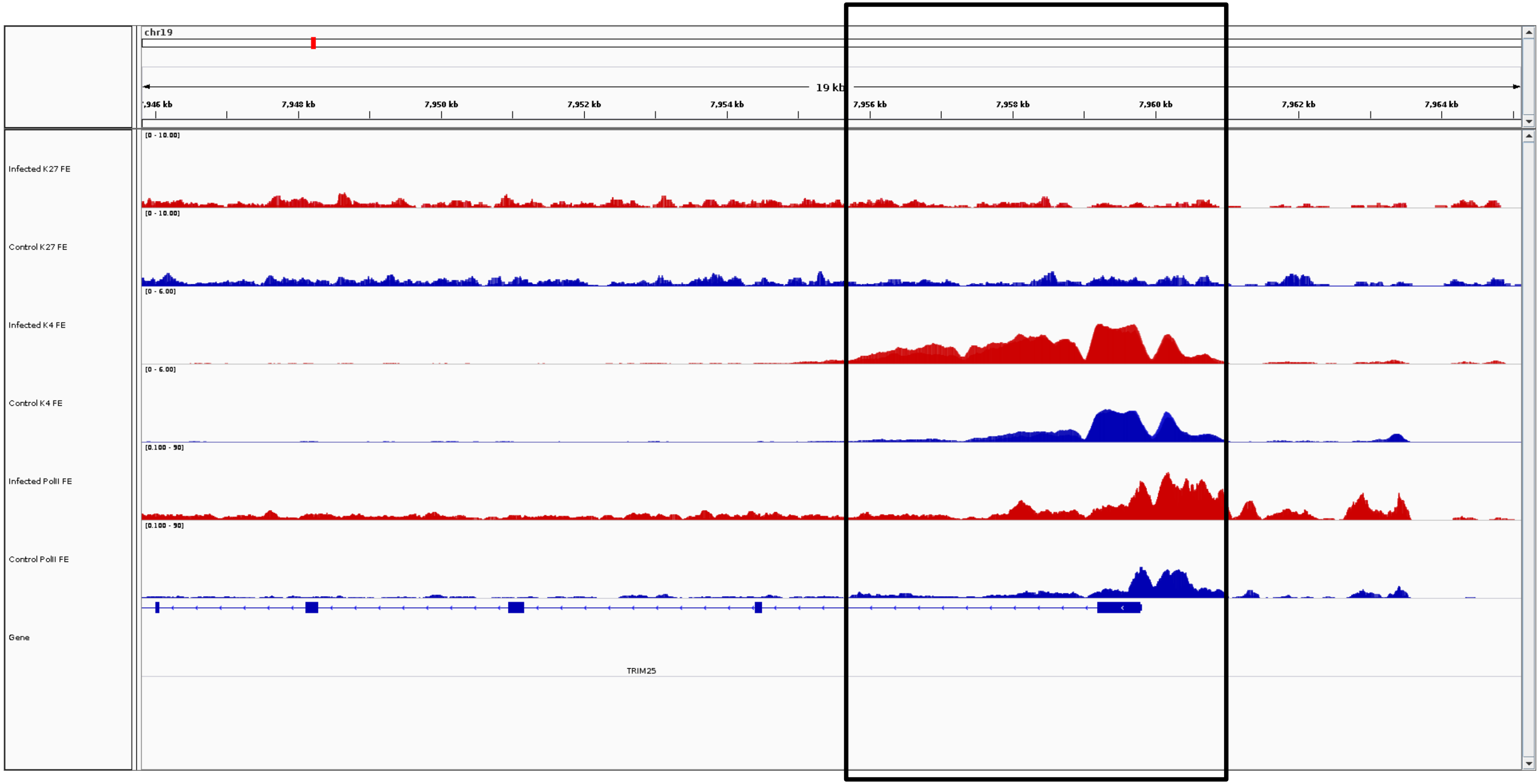
chr19
19 kb
7,946 kb
7,948 kb
7,950 kb
7,952 kb
7,954 kb
7,956 kb
7,958 kb
7,960 kb
7,962 kb
7,964 kb
Infected K27 FE
[0 - 10.00]
Control K27 FE
[0 - 10.00]
Infected K4 FE
[0 - 6.00]
Control K4 FE
[0 - 6.00]
Infected PolII FE
[0.100 - 90]
Control PolII FE
[0.100 - 90]
Gene
TRIM25

***TRIM25***

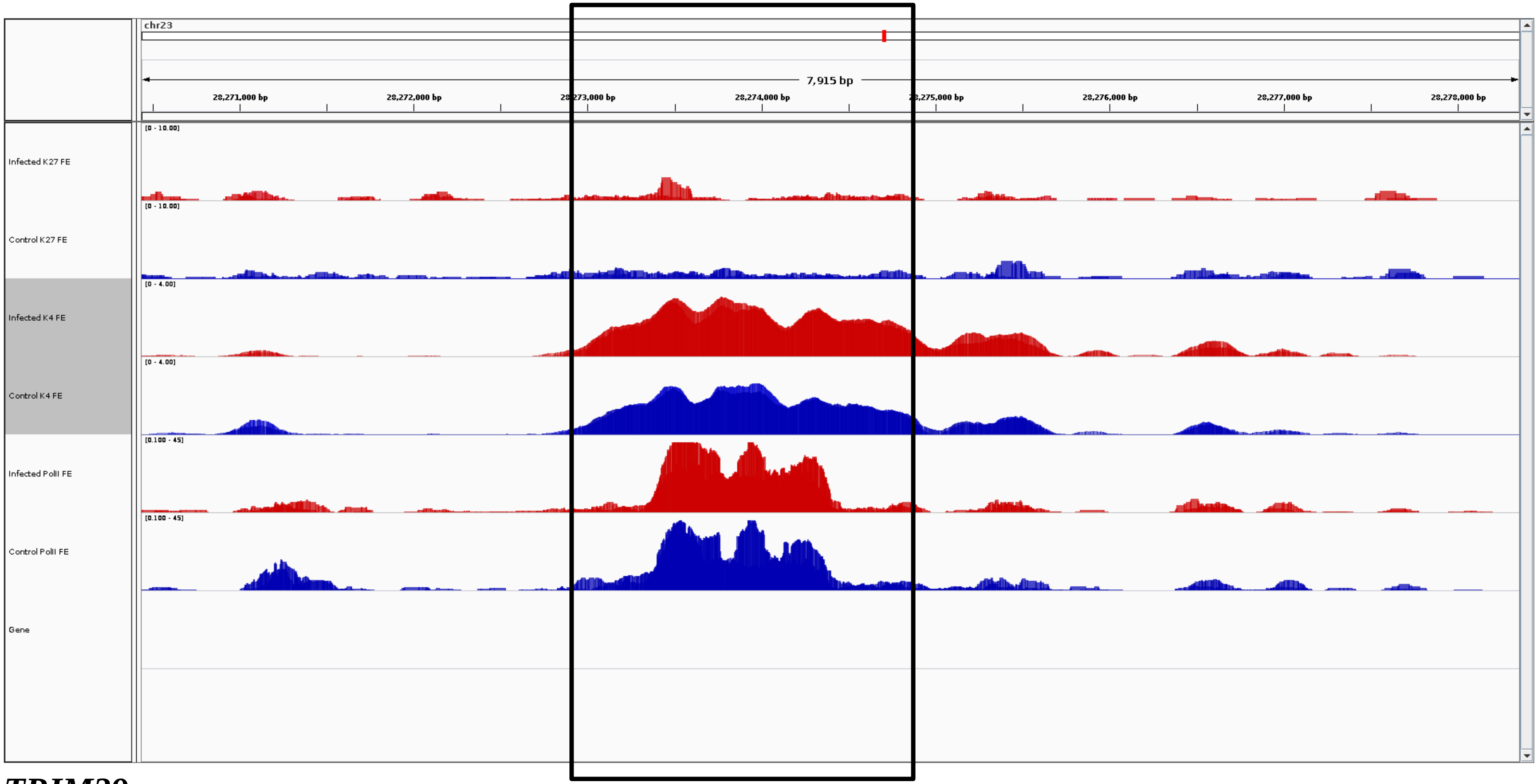
chr23
7,915 bp
28,271,000 bp
28,272,000 bp
28,273,000 bp
28,274,000 bp
28,275,000 bp
28,276,000 bp
28,277,000 bp
28,278,000 bp
Infected K27 FE
[0 - 10.00]
Control K27 FE
[0 - 10.00]
Infected K4 FE
[0 - 4.00]
Control K4 FE
[0 - 4.00]
Infected PolII FE
[0.100 - 45]
Control PolII FE
[0.100 - 45]
Gene

***TRIM39***

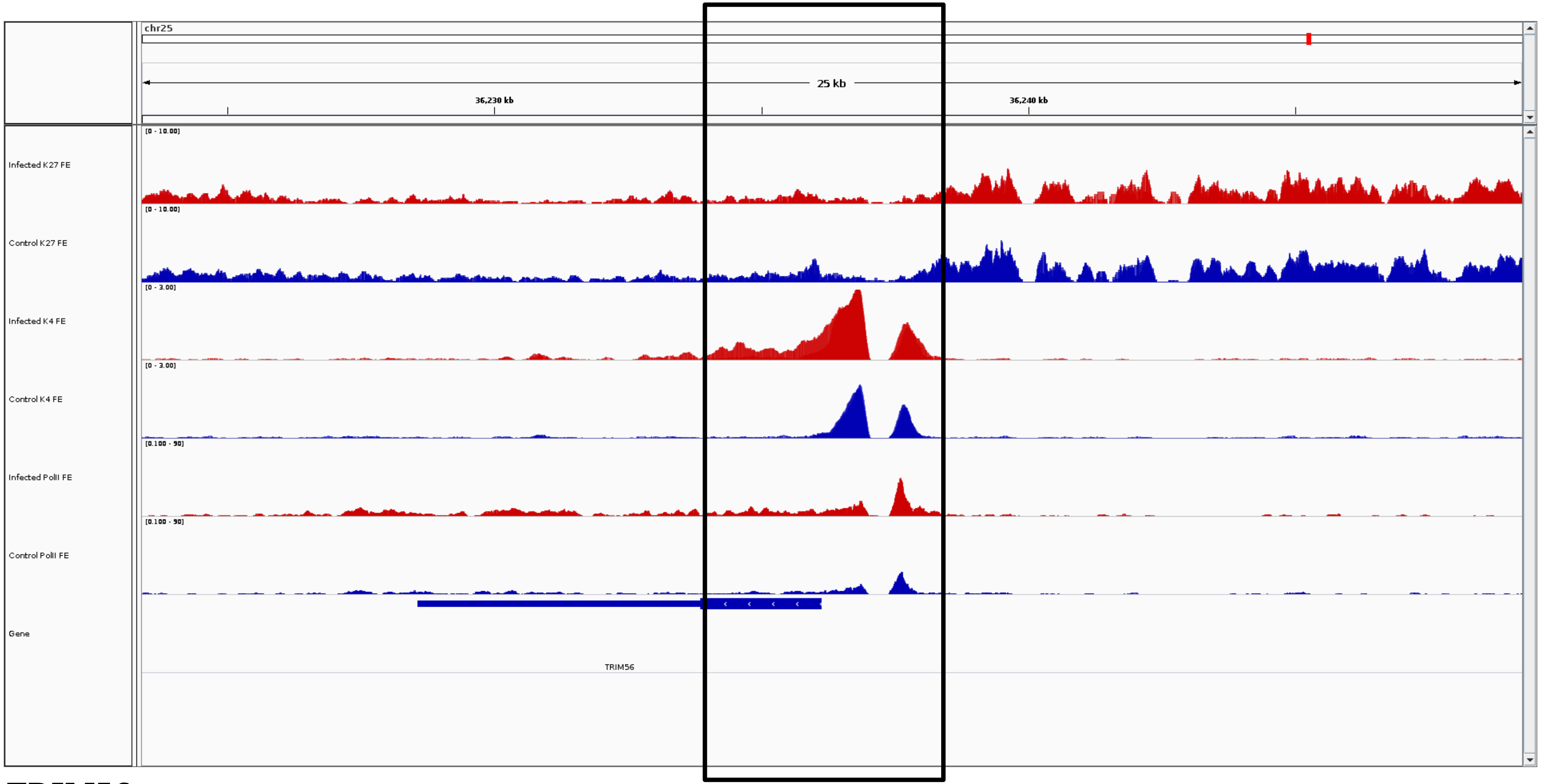
chr25
25 kb
36,230 kb
36,240 kb
Infected K27 FE
[0 - 10.00]
Control K27 FE
[0 - 10.00]
Infected K4 FE
[0 - 3.00]
Control K4 FE
[0 - 3.00]
Infected PolII FE
[0.100 - 90]
Control PolII FE
[0.100 - 90]
Gene
TRIM56

***TRIM56***

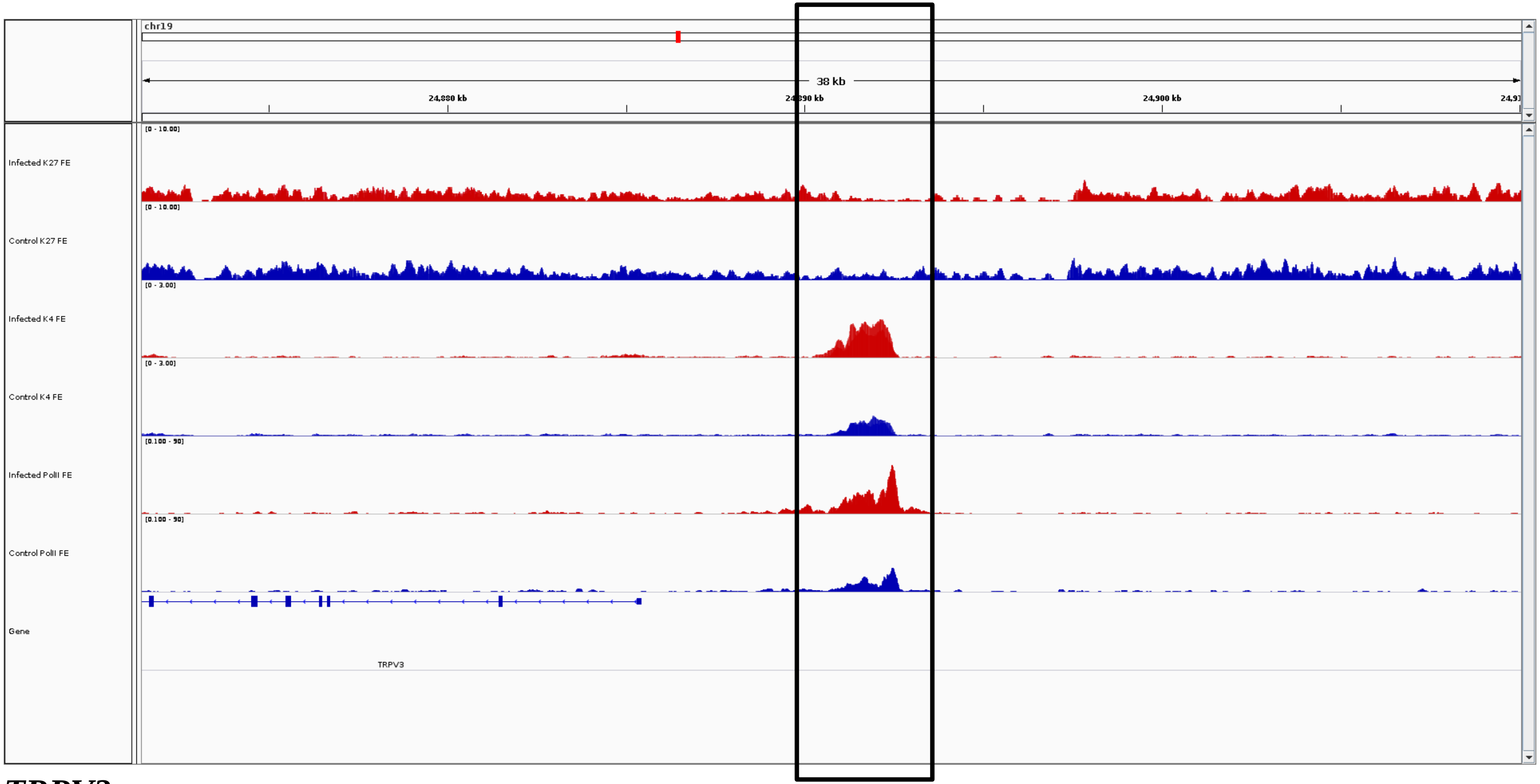
chr19
38 kb
24,880 kb
24,890 kb
24,900 kb
Infected K27 FE
[0 - 10.00]
Control K27 FE
[0 - 10.00]
Infected K4 FE
[0 - 3.00]
Control K4 FE
[0 - 3.00]
Infected PolII FE
[0.100 - 90]
Control PolII FE
[0.100 - 90]
Gene
TRPV3

***TRPV3***

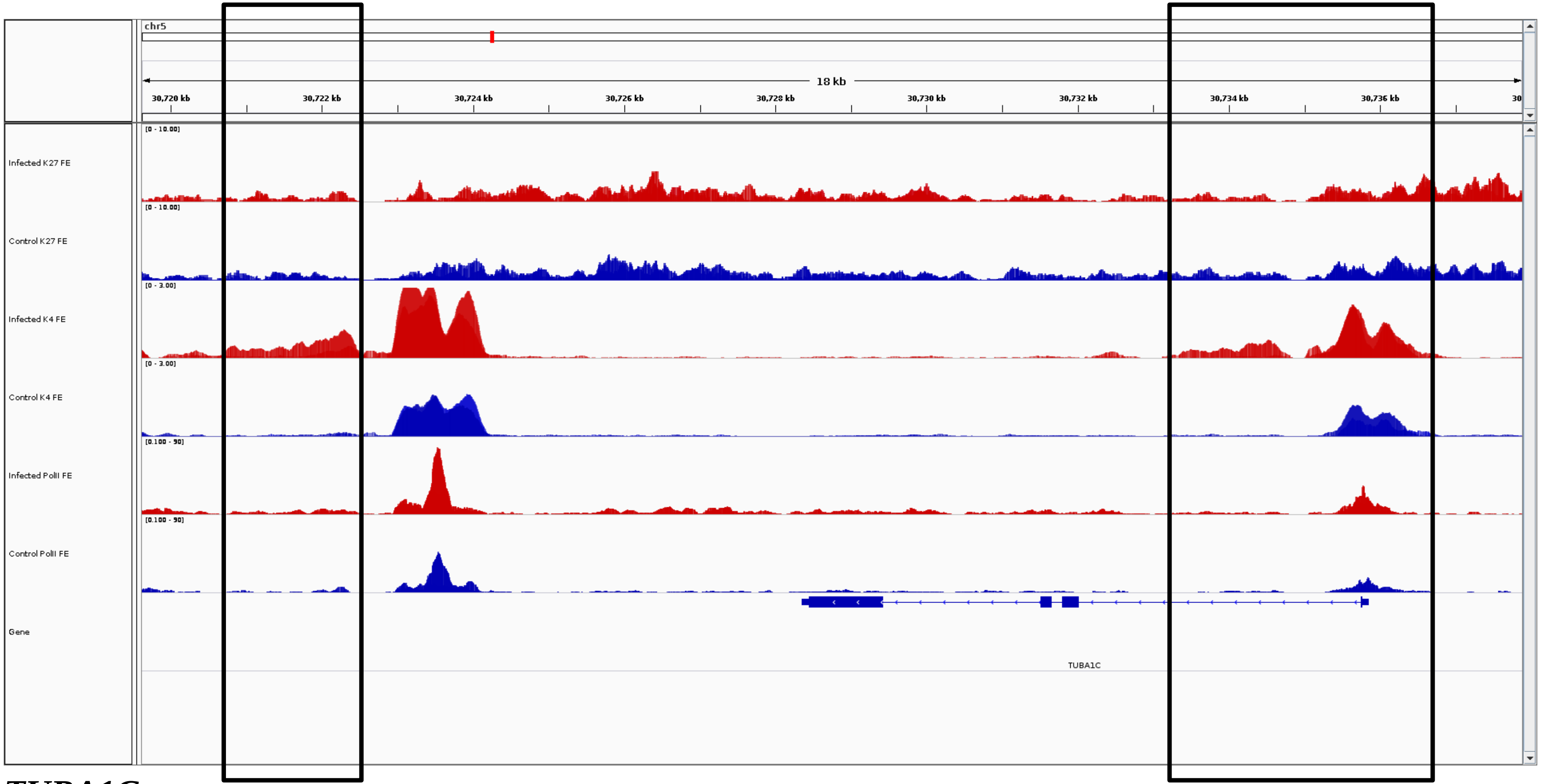
chr5
18 kb
30,720 kb
30,722 kb
30,724 kb
30,726 kb
30,728 kb
30,730 kb
30,732 kb
30,734 kb
30,736 kb
Infected K27 FE
Control K27 FE
Infected K4 FE
Control K4 FE
Infected PolII FE
Control PolII FE
Gene
TUBA1C

***TUBA1C***

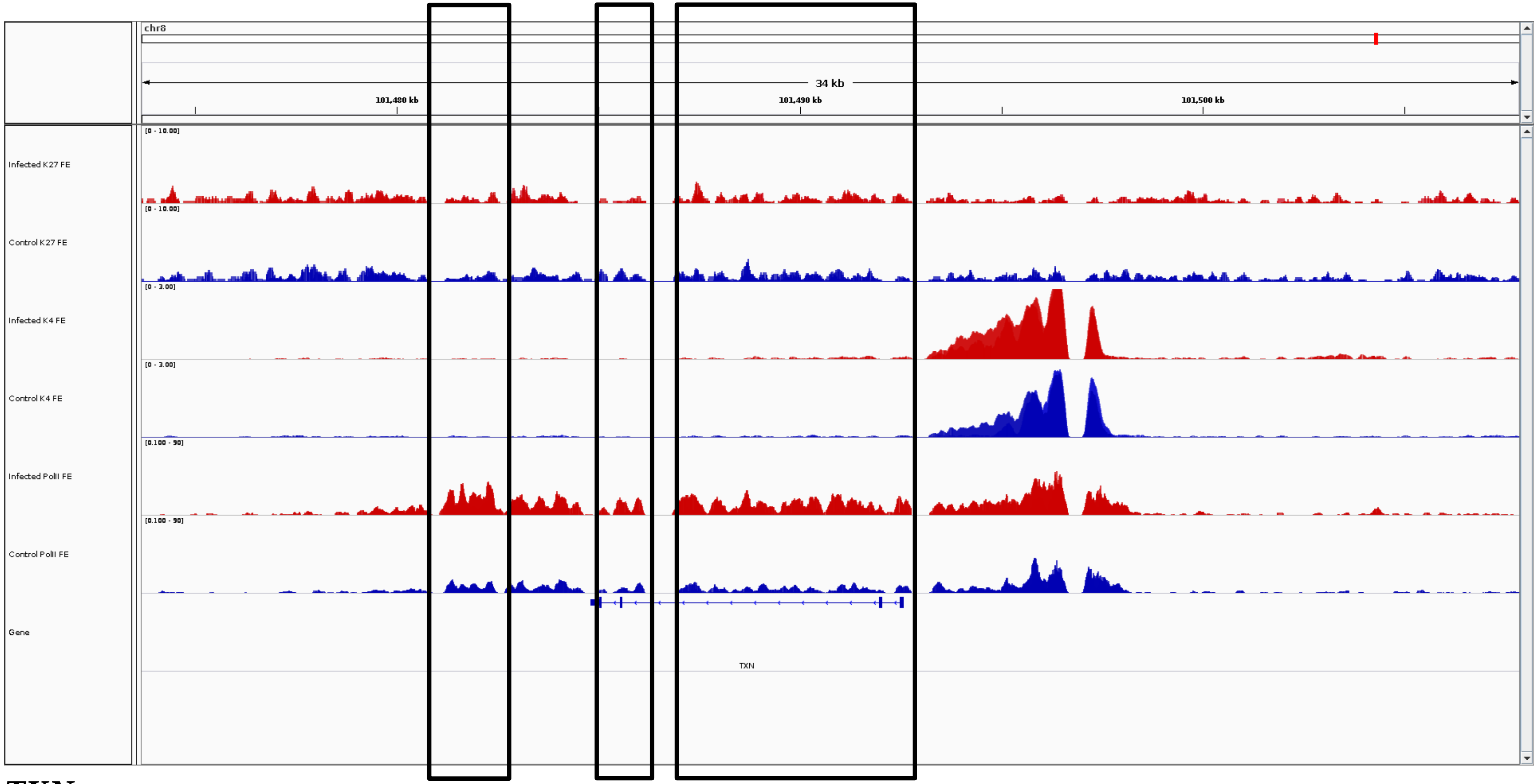
chr8
34 kb
101,480 kb
101,490 kb
101,500 kb
[0 - 10.00]
Infected K27 FE
[0 - 10.00]
Control K27 FE
[0 - 3.00]
Infected K4 FE
[0 - 3.00]
Control K4 FE
[0.100 - 50]
Infected PolII FE
[0.100 - 50]
Control PolII FE
Gene
TXN

***TXN***

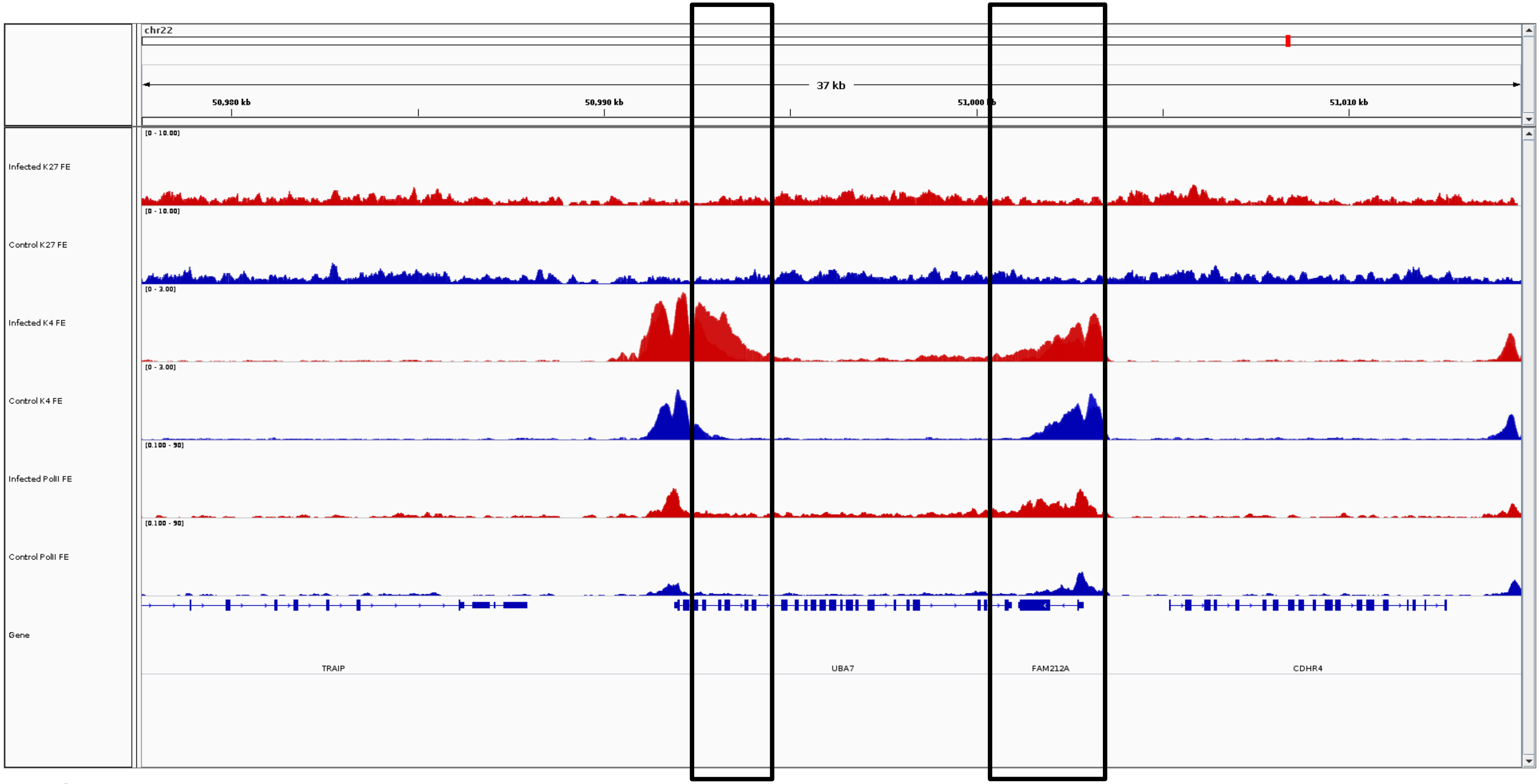
chr22
37 kb
50,980 kb
50,990 kb
51,000 kb
51,010 kb
[0 - 10.00]
Infected K27 FE
Control K27 FE
[0 - 3.00]
Infected K4 FE
Control K4 FE
[0.100 - 50]
Infected PolII FE
Control PolII FE
Gene
TRAIP
UBA7
FAM212A
CDHR4

***UBA7***

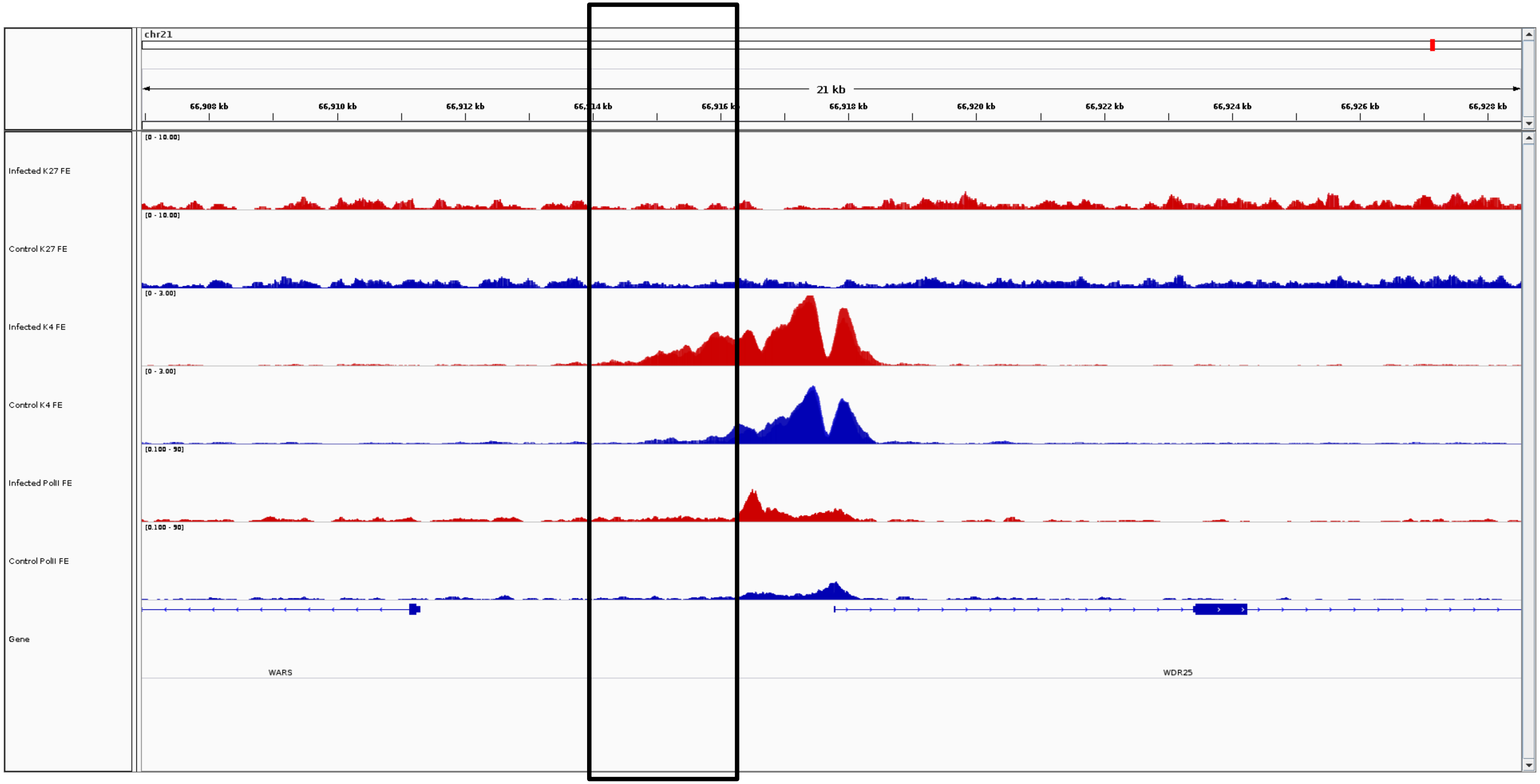
chr21
21 kb
66,908 kb
66,910 kb
66,912 kb
66,914 kb
66,916 kb
66,918 kb
66,920 kb
66,922 kb
66,924 kb
66,926 kb
66,928 kb
Infected K27 FE
[0 - 10.00]
Control K27 FE
[0 - 10.00]
Infected K4 FE
[0 - 3.00]
Control K4 FE
[0 - 3.00]
Infected PolII FE
[0.100 - 50]
Control PolII FE
[0.100 - 50]
Gene
WARS
WDR25

***WDR25***

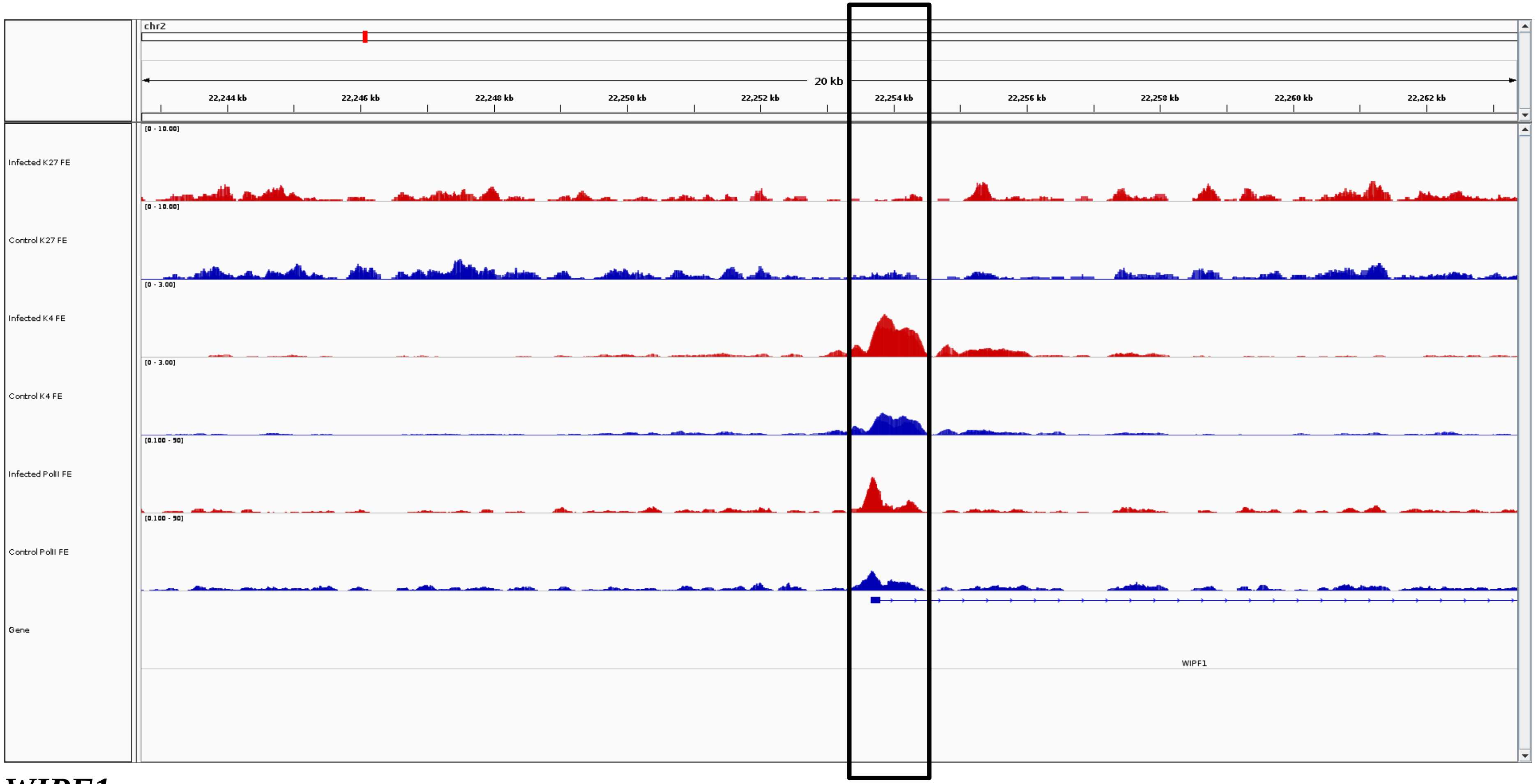

chr2
20 kb
22,244 kb
22,246 kb
22,248 kb
22,250 kb
22,252 kb
22,254 kb
22,256 kb
22,258 kb
22,260 kb
22,262 kb
Infected K27 FE
Control K27 FE
Infected K4 FE
Control K4 FE
Infected PolII FE
Control PolII FE
Gene
WIPF1


***WIPF1***

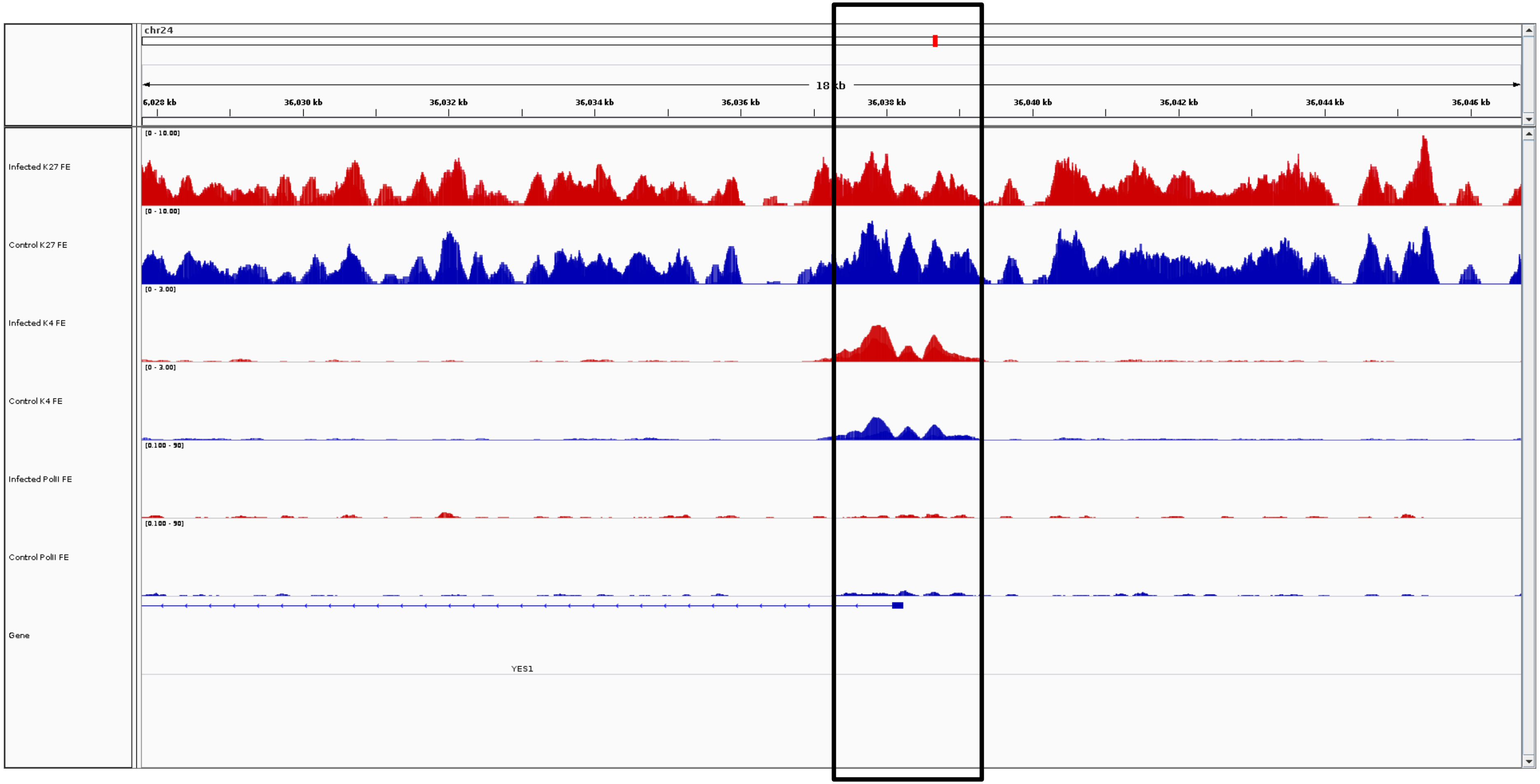
chr24
18 kb
36,028 kb
36,030 kb
36,032 kb
36,034 kb
36,036 kb
36,038 kb
36,040 kb
36,042 kb
36,044 kb
36,046 kb
Infected K27 FE
Control K27 FE
Infected K4 FE
Control K4 FE
Infected PolII FE
Control PolII FE
Gene
YES1

***YES1***

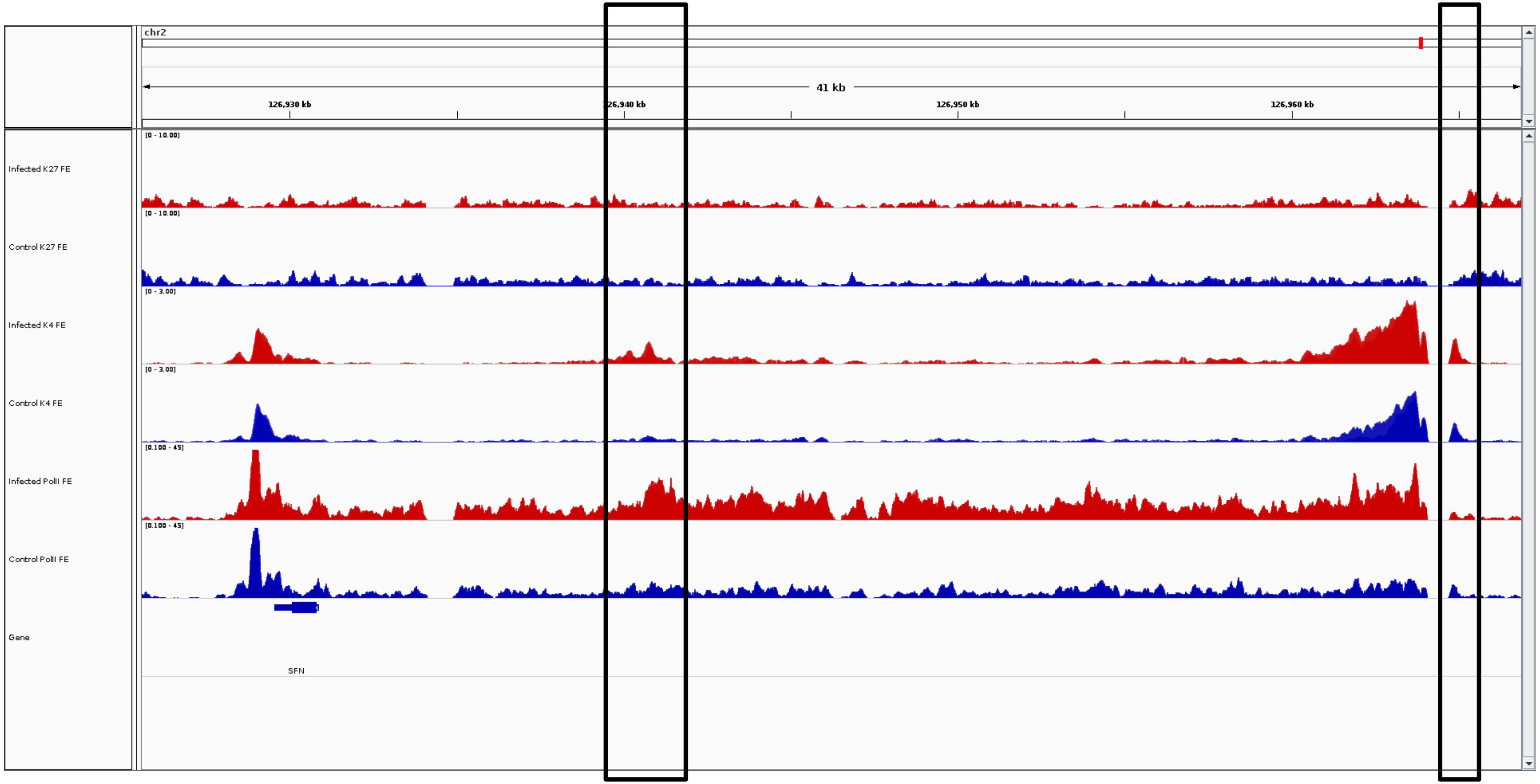
chr2
41 kb
126,930 kb
126,940 kb
126,950 kb
126,960 kb
Infected K27 FE
[0 - 10.00]
Control K27 FE
[0 - 10.00]
Infected K4 FE
[0 - 3.00]
Control K4 FE
[0 - 3.00]
Infected PolII FE
[0.100 - 45]
Control PolII FE
[0.100 - 45]
Gene
SFN

***ZDHHC18-201***

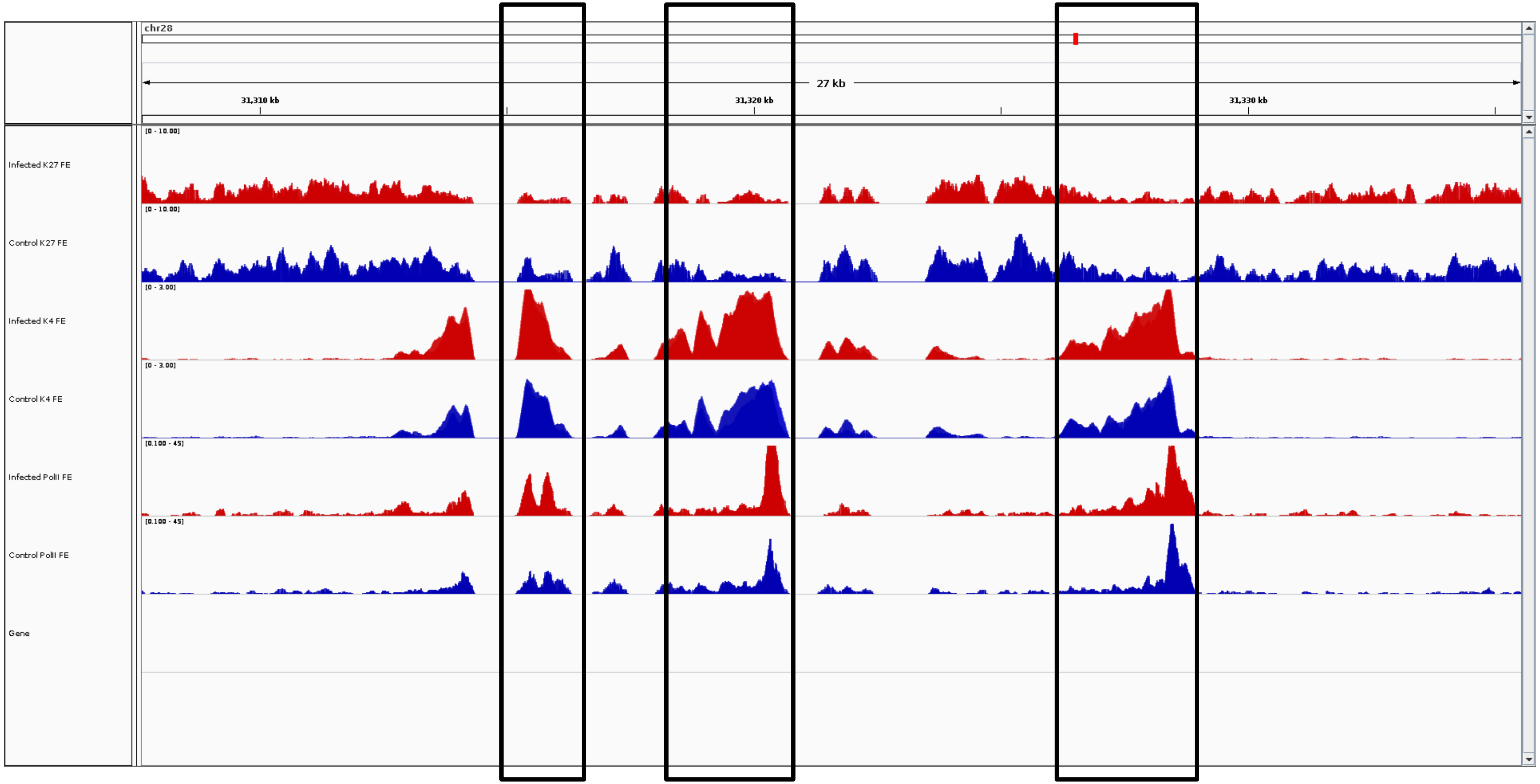
chr28
27 kb
31,310 kb
31,320 kb
31,330 kb
Infected K27 FE
Control K27 FE
Infected K4 FE
Control K4 FE
Infected PolII FE
Control PolII FE
Gene

***ZNF503***
